# The swan genome and transcriptome: its not all black and white

**DOI:** 10.1101/2022.05.02.490350

**Authors:** Anjana C. Karawita, Yuanyuan Cheng, Keng Yih Chew, Arjun Challgula, Robert Kraus, Ralf C. Mueller, Marcus Z. W. Tong, Katina D. Hulme, Helle Beielefeldt-Ohmann, Lauren E. Steele, Melanie Wu, Julian Sng, Ellesandra Noye, Timothy J. Bruxner, Gough G. Au, Suzanne Lowther, Julie Blommaert, Alexander Suh, Alexander J. McCauley, Parwinder Kaur, Olga Dudchenko, Erez Aiden, Olivier Fedrigo, Giulio Formenti, Jacquelyn Mountcastle, William Chow, Fergal J. Martin, Denye N. Ogeh, Françoise Thiaud-Nissen, Kerstin Howe, Joanna Collins, Alan Tracey, Jacqueline Smith, Richard I. Kuo, Marilyn B. Renfree, Takashi Kimura, Yoshihiro Sakoda, Mathew McDougall, Hamish G. Spencer, Michael Pyne, Conny Tolf, Jonas Waldenström, Erich D. Jarvis, Michelle L. Baker, David W. Burt, Kirsty R. Short

## Abstract

The Australian black swan (*Cygnus atratus*) is an iconic species with contrasting plumage to that of the closely related Northern Hemisphere white swans. The relative geographic isolation of the black swan may have resulted in a limited immune repertoire and increased susceptibility to infectious disease, notably infectious diseases from which Australia has been largely shielded. Indeed, unlike Mallard ducks and the mute swan (*Cygnus olor*), the black swan is extremely sensitive to severe highly pathogenic avian influenza (HPAI). Understanding this susceptibility has been impaired by the absence of any available swan genome and transcriptome information. Here, we generate the first chromosome-length annotated black and mute swan genomes annotated with transcriptome data, all using long-read based pipelines generated for vertebrate species. We used these genomes and transcriptomes, to show that unlike other wild waterfowl, black swans lack an expanded immune gene repertoire, lack a key viral pattern-recognition receptor in endothelial cells and mount a poorly controlled inflammatory response to HPAI. We also implicate genetic differences in *SLC45A2* in the iconic plumage of the Australian black swan. Together, these data suggest that the immune system of the black swan is such that should any avian viral infection become established in its native habitat the survival of the black swan would be in significant peril.

## INTRODUCTION

The distinctive black plumage of the native Australian black swan (*Cygnus atratus*) is in stark contrast to the white swans that are native to Europe and North America. This unique feature has resulted in the black swan playing an important role in Australian heraldry and culture. The limited native geographic range (Australia) and relative isolation of the black swan has direct consequences for its immune repertoire and susceptibility to infectious disease common to other parts of the world. Specifically, geographic isolation can result in founder effects and reduced immune diversity as a result of limited pathogen challenge^1^.

The native Australian black swan has a remarkably distinct response to infection by highly pathogenic avian influenza (HPAI) virus compared to the closely related white swans (e.g. the mute swan; *Cygnus olor*) and other waterfowl^2,3^. Unlike Mallard ducks and mute swans, the black swan is extremely sensitive to HPAI, succumbing to the disease within 2 to 3 days post-infection. This disease pathogenesis mirrors that of infected chickens, viewed as the most susceptible species to HPAI^3^. One of the striking features common to both black swans and chickens is that HPAI viruses preferentially infect endothelial cells, which may contribute to the disease severity in these two species^3^. These experimental studies are consistent with reports of natural infections, which suggest that captive black swans quickly succumb to HPAI whilst co-housed mute swans survive the infection^4^.

Comparative genomics has played an important role in understanding species-dependent differences in HPAI pathogenesis^5^ whilst also revealing the unique immune systems of many native Australian fauna^6^. However, comparative genomics is contingent upon the availability of high-quality species-specific genomes and transcriptomes.

Here, we generate the first black and mute swan reference genomes and transcriptomes, including the transcriptional response of primary black swan endothelial cells to HPAI. These data show that the black swan has numerous unique characteristics including (i) lack of an expanded immune gene repertoire (ii) undetectable *Toll-like Receptor (TLR) 7* gene expression in infected endothelial cells and (iii) a dysregulated pro-inflammatory response to viral infection that is likely to leave the species highly susceptible to viral infections such as HPAI. It is also likely that genetic differences in melanin production contribute to the distinctive black plumage of the black swan.

## RESULTS

### GENOME LANDSCAPE

The chromosome-length reference genomes for black and mute swan were constructed according to a Pacbio continuous long-read (CLR) DNA Zoo pipeline and Vertebrate Genomes Pipeline 1.5^7^, respectively (Supplementary Figures 1 and 2). This included scaffolding contigs with Hi-C and Bionano maps. Curations of the assemblies identified scaffolds representing 34 autosomes plus the Z sex chromosome, and were named according to the descending order of the physical size. We assigned final scaffolds to 34 chromosomes (including the male sex chromosome) based on the physical size. The W chromosome was not assigned a scaffold as the DNA was from a male. The expected diploid number of chromosomes in the mute swan is 80^8^. The black swan genome was sequenced using 90x PacBio CLR coverage, generating a 1.12 Gb reference assembly. The mute swan genome was sequenced using 60x PacBio CLR coverage, generating a 1.13 Gb reference assembly. The details of both genomes, including a comparison to the latest VGP chicken (*Gallus gallus*) and Mallard duck (*Anas platyrhynchos*) genomes, are shown in Table 1. The total classified repeat content of the genome was 10.56% for the black swan, with 1.71% unclassified repeats, and 10.76% for the mute swan genome, with 1.55% unclassified. This is lower than the repeat content recorded in the chicken (15%) and the Mallard duck (17%) ^9,10^.

**Table 1:** Comparison of genomes between black and mute swans (generated herein) and published high-quality avian genomes generated by the VGP.

| Species | Assembly size (Gb) | G + C content (%)<br>Macro-chromosome | G + C content (%)<br>Micro-chromosome | No. scaffolds | Scaffold N50 (Mb) | Scaffold L50 | ISO-seq<br>ISO-seq coverage* | Coding genes | Non-coding genes | Pseudogenes | Ref |
| --- | --- | --- | --- | --- | --- | --- | --- | --- | --- | --- | --- |
| Black swan | 1.12 | 47% | 52% | 568 | 84 | 4 | 68.87% of the genome | 15478 | 1792 | 104 | This study |
| Mute swan | 1.13 | 47% | 53% | 181 | 40.61 | 6 | 54.36% of the genome | 14791 | 2720 | 126 | This study |
| Chicken | 1.09 | 40% | 48% | 214 | 18 | 18 | NA | 16878 | 7166 | 312 | bGalGal1.mat.broiler.GRCg7b (GCF_016699485.2) |
| Duck | 1.18 | 40.72 | 46.51 | 756 | 76.3 | 5 | NA | 16618 | 9285 | 49 | ZJU1.0 (GCF_015476345.1) |

\* of the largest 34 scaffolds \*micro-chromosomes < 10 MBps and macro-choromosomes > 10 Mbps

The black and mute swan genomes were annotated with RNASeq and IsoSeq transcriptome data, homology based alignments with other species and with bioinformatically inferred gene predictions, according to the methods listed in Supplementary Figure 5.

The completeness of the black and mute swan genomes was assessed using the Core Eukaryotic Genes Mapping (CEGMA) and Benchmark Universal Single Copy Orthologues (BUSCO) analyses and compared to the chicken and Mallard duck genome (Supplementary Table 1 & 2). Notably, the black swan genome had the highest complete BUSCOs (8093), followed by the chicken (8054) and the mute swan (8010) genomes. Whilst the chicken genome had the highest complete core-eukaryotic gene content (224), this was only marginally higher than that of the mute (221) and black swan (219) genomes.

One-to-one alignment between the black and mute swan genomes showed 98.35% average nucleotide identity between the 34 autosomes of the black and mute swans (Supplementary Figure 3). The Z chromosome of the black and mute swan had several large (>10kb) structural variants (Supplementary Figure 4), but were otherwise largely consistent. These structural differences in the Z chromosome may be associated with the speciation of the black and mute swan from their last common ancestor (approximately 6.1 million years ago) ^11^.

Structural genomic differences have been associated with the differential susceptibility of chickens and ducks to avian influenza^12^. We found no substantive inversions between the black and mute swan. However, given that both ducks and swans are *Anseriformes* we compared structural genomic differences between the susceptible black swan and mute swan and relatively resistant duck, using the ostrich as an outgroup. We investigated genes in the inverted genomic regions present in the duck but absent in the swans on chromosome 1. Strikingly, we found that 53 inverted genes (out of 1758 total genes) in chromosome one of the duck genome were mapped to immune system processes (Supplementary Table 3). However, given the absence of substantive structural variants between the black and mute swan, it is likely that any immunological consequences of these structural variants would be present in multiple swan species.

The black and mute swan genomes were then annotated according to the methods listed in Supplementary Figure 5. Sixteen thousand two hundred four (16,204) gene models were obtained through Evidence Modeler as the final gene models in the black swan and 15,789 in the mute swan. Protein alignment against the UniProt/Swiss-Prot database was used to infer 15,478 gene model names for the black swan and 14,791 gene model names for the mute swan.

### MUTATIONS IN *SLC45A2* MAY ACCOUNT FOR DIFFERENCES IN SWAN PLUMAGE

One of the most remarkable features of the black swan is its distinct plumage pattern. To determine if the highly annotated genomes presented herein could offer insight into the iconic plumage we examined four genes (*SLC45A2, SLMO2, ATP5e* and *EDN3*) known to be involved in avian plumage colour^13,14^.Three of the four genes, *SLMO2 (PERLI3B), ATP5e* and *EDN3* shared 100% amino acid identity between the black and mute swan (data not shown). The exception, *SLC45A2* in the mute swan had a nucleotide deletion in the first open reading frame of the mute swan, instigating a frame-shift mutation and an in-frame early stop codon (Figure 2). Multiple non-homologous nucleotides were detected in the chicken and the duck *SLC45A2* relative to that of the black swan (Figure 2). *SLC45A2* encodes a membrane-associated transport protein which regulates the tyrosinase activity and the melanin content of melanosomes^15^. The knockdown of this gene causes low melanin content and reduced tyrosinase activity in human melanoma cell lines^16^. These results suggest that this deletion in *SLC45A2* is a candidate genetic change that could be responsible for the white plumage in white swans in the genus *Cygnus*.

**Figure 2:**
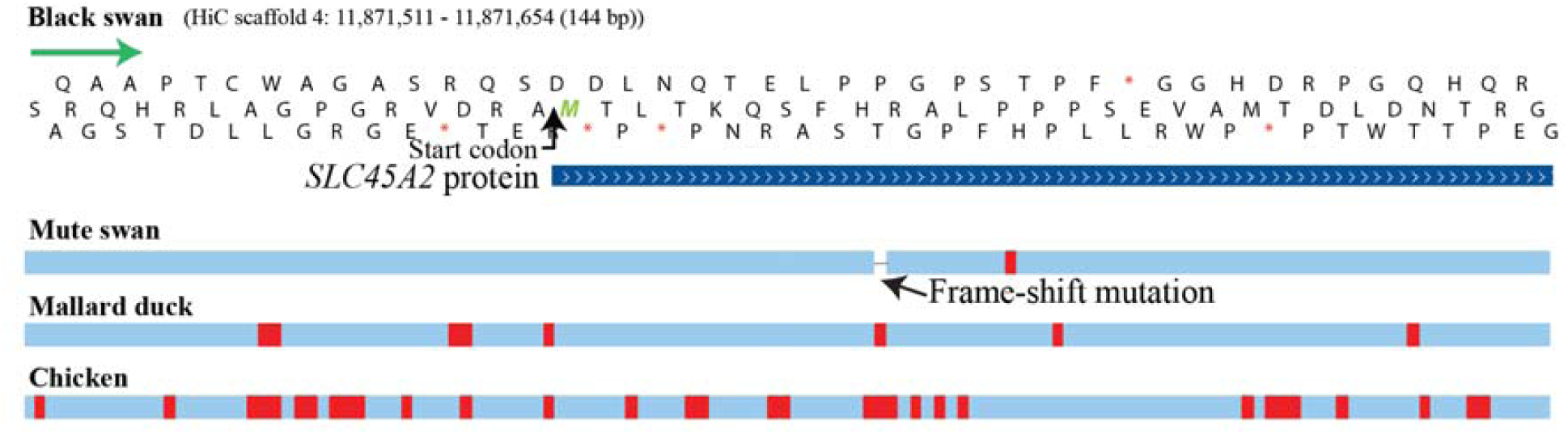
Position of the frame-shift mutation in the mute swan *SLC45A2* compared to the black swan genome. Red vertical lines represent non-homologous nucleotides compared to the black swan. The figure was created with whole-genome HAL alignment produced from CACTUS. The alignment was visualised with the University of California, Santa Cruz (UCSC) Genome Assembly in a Box (GBiB). The predicted mute swan gene *SLC45A2* was aligned using the BLAT alignment algorithm.

### IMMUNE GENE FAMILIES ARE EXPANDED IN THE MALLARD DUCK AND MUTE SWAN GENOMES

To understand whether the relative isolation of the black swan has resulted in an altered immune gene repertoire, evolutionary gene gain and loss were determined (Figure 3). (*p-value* < 0.05). The black swan genome was estimated to be contractive, indicating that the total gene gain was less than the gene loss from the last common ancestor. The biological function of expanded genes in the black swan, chicken, mute swan, and duck was then investigated.

**Figure 3:**
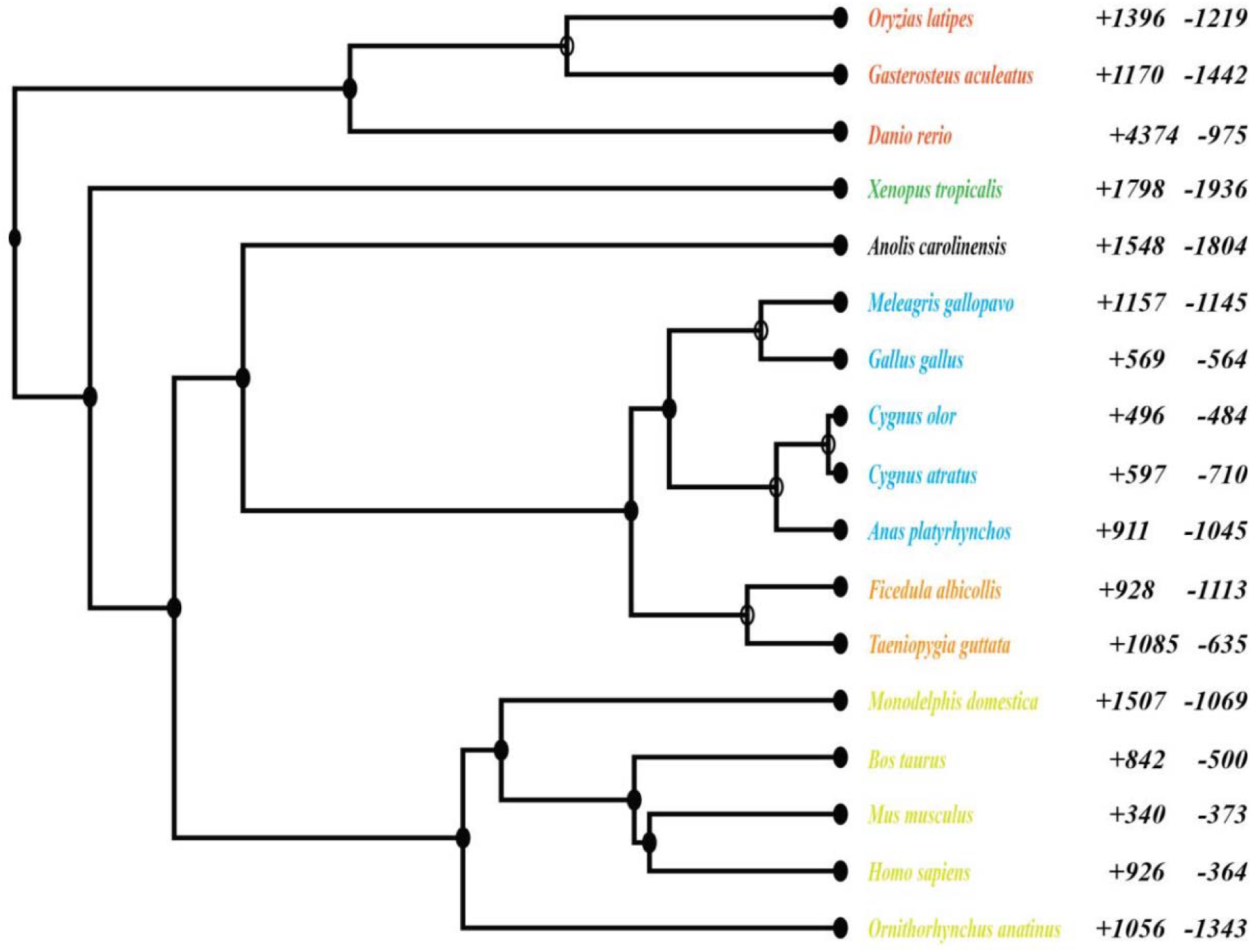
Gene losses and gains across vertebrate species tree. Data are shown for 17 vertebrate species, three teleost (red), an amphibian (green), six reptilians (including birds) (black, blue, and orange) and five mammals (yellow). The numbers of gene family gains (+) and losses (-) are given to the right of the taxa (*p-value* < 0.05) compared to the taxon’s last common ancestor. The rate of gene birth and death for clades derived from the most recent common ancestor (MRCA) for *Gallianseriformes* is 0.0016 (per gene per million years).

Strikingly, immune system processes (e.g. GO0002376) were only expanded in the Mallard duck and the mute swan genomes (Figure 4). In contrast, in chickens, expanded gene families were associated with regulation of GTPase activity, extracellular matrix and structure organization, whilst the over-represented functional terms for expanded black swan gene families included cell-matrix adhesion, cell-substrate adhesion, extracellular matrix organization and extracellular structure organization. To specifically compare the immune gene repertoire of black and mute swans we used human and mouse immune genes to identify immune gene families in *Cygnus* species. Thirty-nine immune-related gene families of the black swan were contractive compared to the mute swan (Supplementary Table 4). The PANTHER pathways related to these genes included apoptosis signalling, cadherin signalling, general transcription by RNA polymerase, gonadotropin-releasing hormone receptor, inflammation mediated by chemokine and cytokine signalling, interleukin signalling, TGF-beta signalling and Wnt signalling pathways.

**Figure 4:**
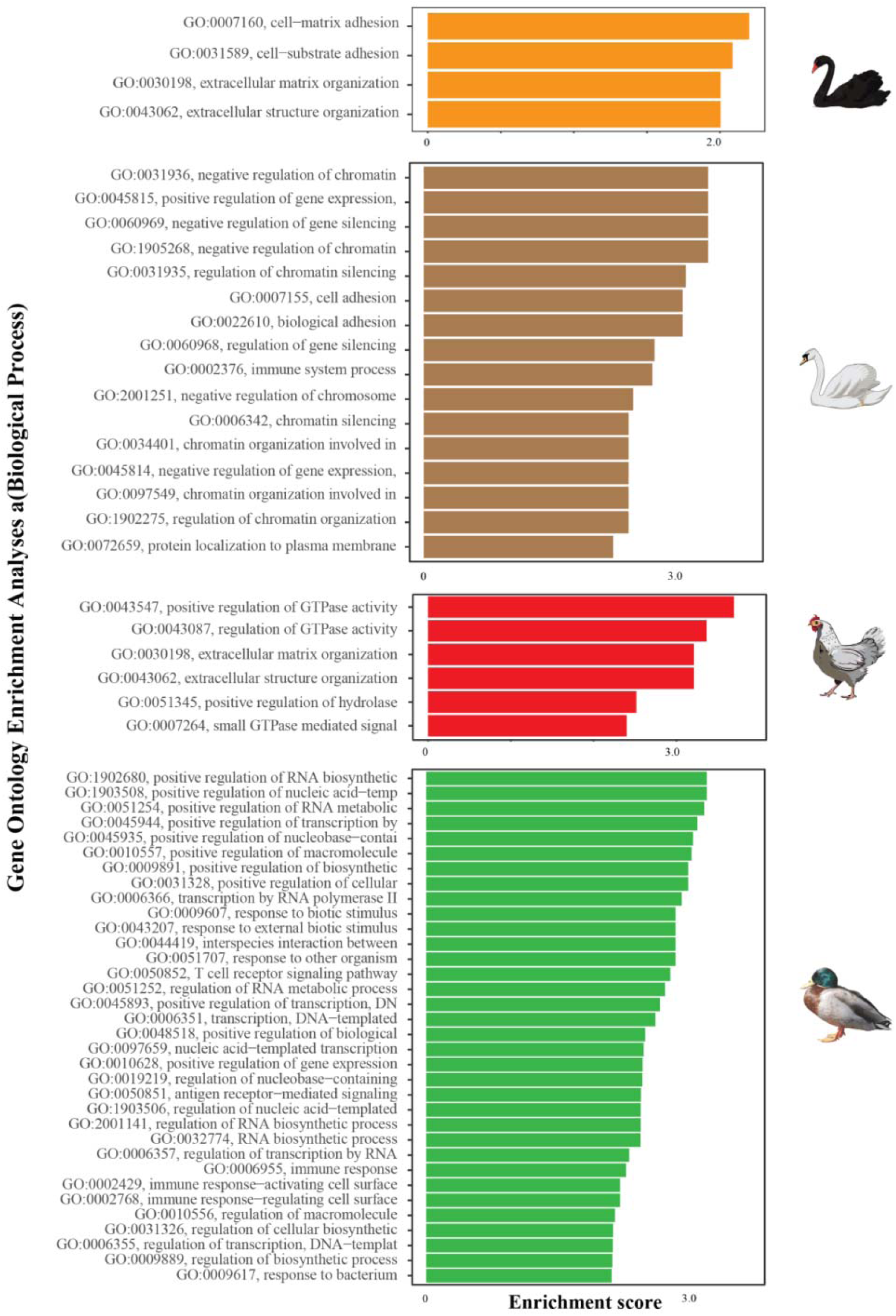
GO enrichment analysis for over-represented GO terms associated with significantly expanded gene families in black swan, mute swan, chicken, and mallard duck.

### BLACK AND MUTE SWAN CLASSICAL MHC CLASS LOCI ARE HOMOLOGOUS

MHC diversity is altered in some avian species ^17^, which may affect susceptibility to disease^18^. We therefore compared MHC loci between black and mute swans. Two MHC class I and MHC class II loci were identified in the black and mute swan (Supplementary Figure 6). These were located on chromosome 33 in the swan genomes. A similar number of MHC complex associated genes were identified in each species. None of these genes as appeared to be pseudogenes. Compared to mammals, both mute and the black swans have a compact, relatively simple MHC B locus (Supplementary Figure 7), with two class IIb (so-called *BLB*) genes followed by a pair of class I (so-called *BF*) genes that flank the *TAP* genes. The *TAPBP* gene in both birds, unlike chickens, does not flank the two-class-IIb genes ^19^. Overall, the MHC region of both the black and mute swan share a similar genome landscape and represent a so-called minimal essential MHC similar to chicken and Mallard duck^20^. It is therefore unlikely that differences in the MHC complex contribute to species-specific differences in the response to HPAI virus infection.

### TOLL LIKE 7 RECEPTOR (TLR7) EXPRESSION CANNOT BE DETECTED IN BLACK SWAN ENDOTHELIAL CELLS

TLR7 signalling has been implicated in influenza A virus recognition in mammals and birds where it functions as a pathogen recognition receptor (PRR) that recognizes single-stranded viral-RNA^21^. *TLR7* has been duplicated independently in several avian species^22^ and differences in *TLR7* tropism and function have been associated with the increased resistance of ducks to HPAI^23^ There was no notable structural difference in the *TLR7* gene between the black and mute swan genomes. However, strikingly *TLR7* expression signals were detected in ISO-Seq analysis of mute swans but not in the ISO-seq analysis of black swan (Supplementary Figure 8). To independently confirm these data, we investigated the expression of *TLR7* using qRT-PCR in black swan tissues collected post-mortem. *TLR7* mRNA could not be detected in any of the collected black swan tissues (Supplementary Table 5). As *TLR7* expression can be induced by interferon we reasoned that gene expression in the black swan may only be detected in the presence of virus infection. Accordingly, we sought to establish an *in vitro* model of HPAI infection in black swans. In black swans experimentally infected with HPAI virus endothelial cells are the primary target of infection^3^. We observed similar infection of endothelial cells in swans naturally infected with HPAI (Supplementary Figure 9). We therefore cultured primary black swan endothelial cells according to our previously described protocol for avian species^24^ and endothelial cell identity was confirmed by tube formation, uptake of acetylated low density lipoprotein, von Willebrand factor expression and the absence of *CD45* expression (Supplementary Figure 10). Chicken, duck, and black swan endothelial cells were infected with A/Chicken/Vietnam/008/2004/H5N1 (VN04) and six hours later gene expression was examined by RNASeq. PCA plots showed that mock and virus-infected samples clustered separately for all three species (Figure 5). Viral RNA was detected in the infected endothelial cells of all three species (data not shown). Importantly, whilst *TLR7* transcription was upregulated (although not statistically significant) in infected duck and chicken endothelial cells, *TLR7* transcription could not be detected in infected or naïve black swan endothelial cells (Table 2). Moreover, whilst *MyD88*, the downstream adaptor of *TLR7* was upregulated in infected duck and chicken endothelial cells it was downregulated in infected black swan endothelial cells (Table 2). These data are consistent with the absence of *TLR7* expression in black swan endothelial cells, despite an apparently intact *TLR7* gene in the genome.

**Table 2:** Quantified *TLR7* and *MyD88* expression following VN04 infection of avian endothelial cells

| Species | Gene | Fold-change | Adjusted p value | Ensemble gene ID |
| --- | --- | --- | --- | --- |
| Duck | *TLR7* | 3.13 | >0.1(0.57) | ENSAPLG00000026818 |
| Duck | *MyD88* | 1.37 | <0.1(9.87e-8) | ENSAPLG00000026818 |
| Chicken | *TLR7* | 2.37 | >0.1(0.23) | ENSGALG00000016590 |
| Chicken | *MyD88* | 1.31 | <0.1(1.9e-7) | ENSGALG00000005947 |
| Black swan | *TLR7* | NE | NA | *ENSACYG00000003732 |
| Black swan | *MyD88* | -1.22 | >0.1(0.15857) | *ENSACYG00000016507 |

\*corresponding gene ID Ensembl annotation for black swan, NA – Not applicable, NE – Not expressed/detected.

### BLACK SWAN ENDOTHELIAL CELLS DISPLAY A DYSREGULATED PRO-INFLAMMATORY RESPONSE TO HPAI VIRUS INFECTION

The transcriptomic data generated herein offer a unique insight into the transcriptomic response of black swans to HPAI virus infection. To explore this matter further, we performed GO enrichment analysis using significantly differentially expressed genes in infected chicken, duck, and black swan endothelial cells (Supplementary Table 6-8). The most significantly upregulated gene in black swans was *IFIT5, IL8* in chickens, and *BCOR* in ducks. *IL6* was significantly upregulated in the black swan (log _2_fold change = 1.89) and chicken (log _2_fold change = 1.21), indicating a strong pro-inflammatory response while not differentially expressed in ducks. Black swan, chicken and duck endothelial cells differentially expressed other cytokines/chemokines and their receptors in response to HPAI VN04 infection (Supplementary Table 9). Typically, black swan and chicken endothelial cells upregulated more cytokines and cytokine receptors than duck endothelial cells in response to HPAI VN04 infection. Indeed, the highest number of cytokines and cytokine receptors were upregulated by infected chicken endothelial cells (Supplementary Table 9). In infected black swan endothelial cells 113 GO terms were significantly enriched (Supplementary Table 10; Figure 6A). Many of these GO terms were associated with innate immunity, the cytokine signalling response and chemokine signalling. Several innate immunity pathways were increased in response to viral infection (z – score > 0) whilst GO terms such as negative regulation of MAP kinase activity and negative regulation of JNK cascade were decreased (z-score < 0). Similarly, 123 enriched GO terms in infected chicken endothelial cells included positive regulation of viral response and regulation of leukocyte chemotaxis (Supplementary Table 11; Figure 6B). Terms such as leukocyte mediated cytotoxicity were increased after infection (z-score > 0) whilst negative regulation of apoptotic signalling and the positive regulation of innate immune responses were decreased. Strikingly, GO biological process terms enriched in infected duck endothelial cells were not primarily associated with the innate immune response (Supplementary Table 12; Figure 6C). Rather, most genes were linked to cellular biological signalling and activity. This finding is consistent with our previous study of HPAI viruses in duck endothelial cells^25^. Interestingly, in direct contrast to black swans the inactivation of MAPK activity was significantly increased in ducks (z-score < 0). Due to the wide-ranging role of the MAPK cascade, including pro-inflammatory responses, we further investigated the expression profiles of the genes and identified ten genes involved in the “inactivation of MAPK pathway,” five of which were significantly downregulated genes (i.e., *DUSP1, DUSP4, DUSP7, DUSP10*, and *RGS3*) in black swans (Supplementary Figure 13). Dual-specificity phosphatases (DUSPs) are negative regulators of MAPKs and their associated pro-inflammatory effects^26^. Accordingly, we specifically examined the differential expression of DUSPs across the three avian species. In the black swan, all DUSPs were either not differentially expressed or down regulated in response to infection. In contrast, in the duck all DUSPS (except for *DUSP15*) were upregulated. Similar, in the chicken, *DUSP1, DUSP5, DUSP7, DUSP10, DUSP15 DUSP16* were significantly downregulated in response to infection (Supplementary Table 13). In contrast, in the duck, all DUSPS (except for DUSP15) were upregulated. This transcriptional profile is consistent with poor regulation of a pro-inflammatory response to HPAI virus in black swans.

**Figure 5:**
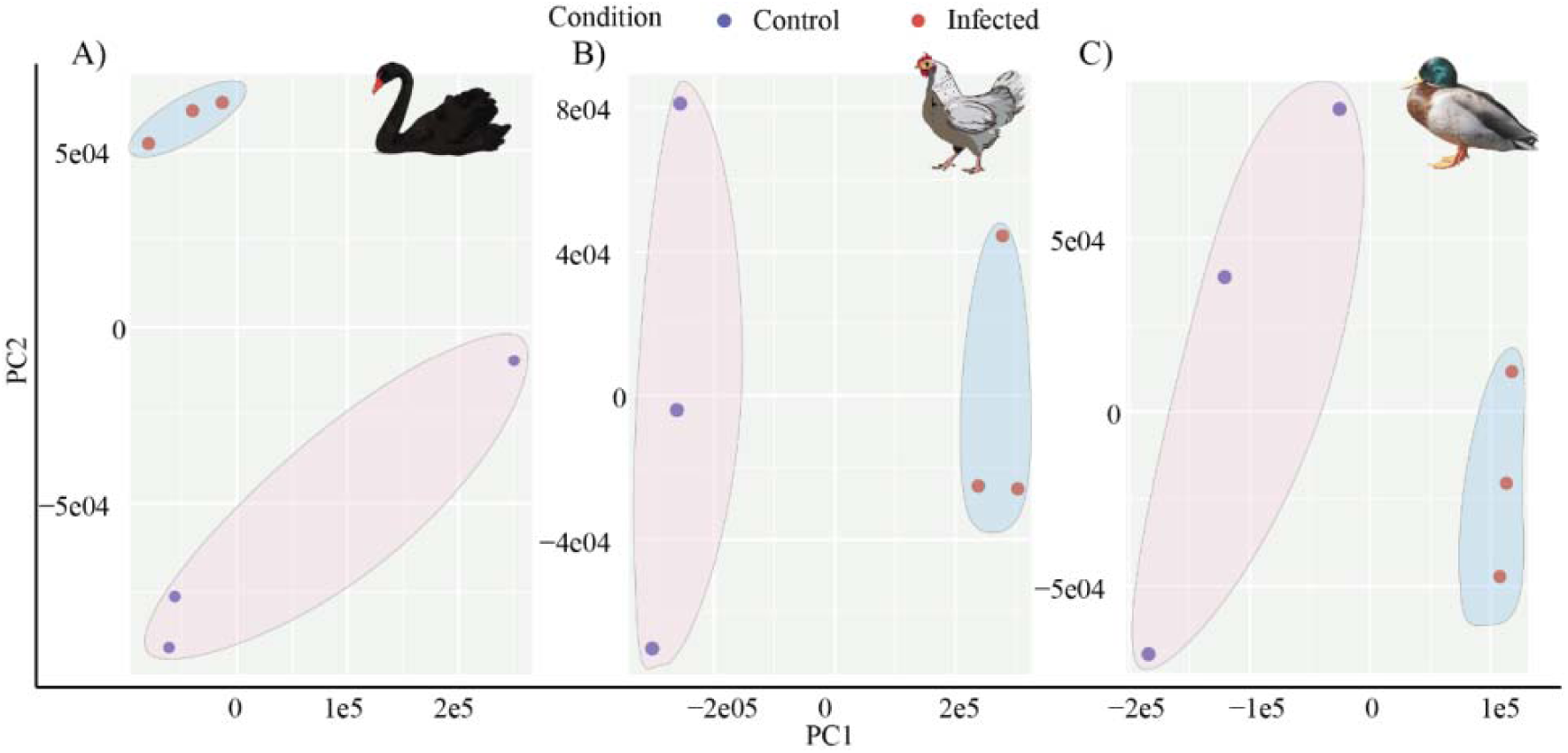
Principal component analysis (PCA) of infected and uninfected (A) black swan (B) chicken (C) duck endothelial cells. The control and the infected groups showed intergroup clustering, indicating differences in whole transcriptome profiles between the control and the infected group in each species.

**Figure 6:**
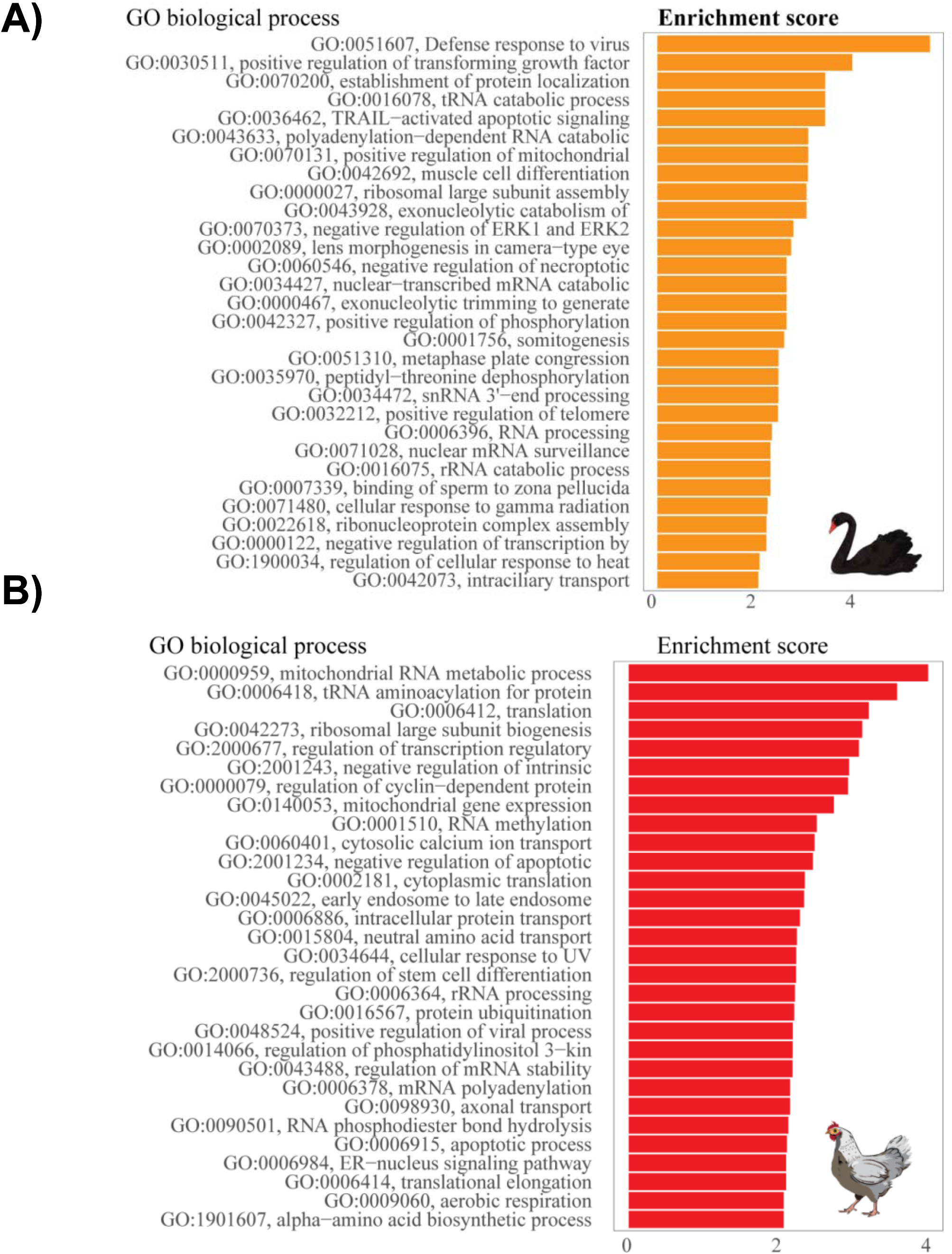
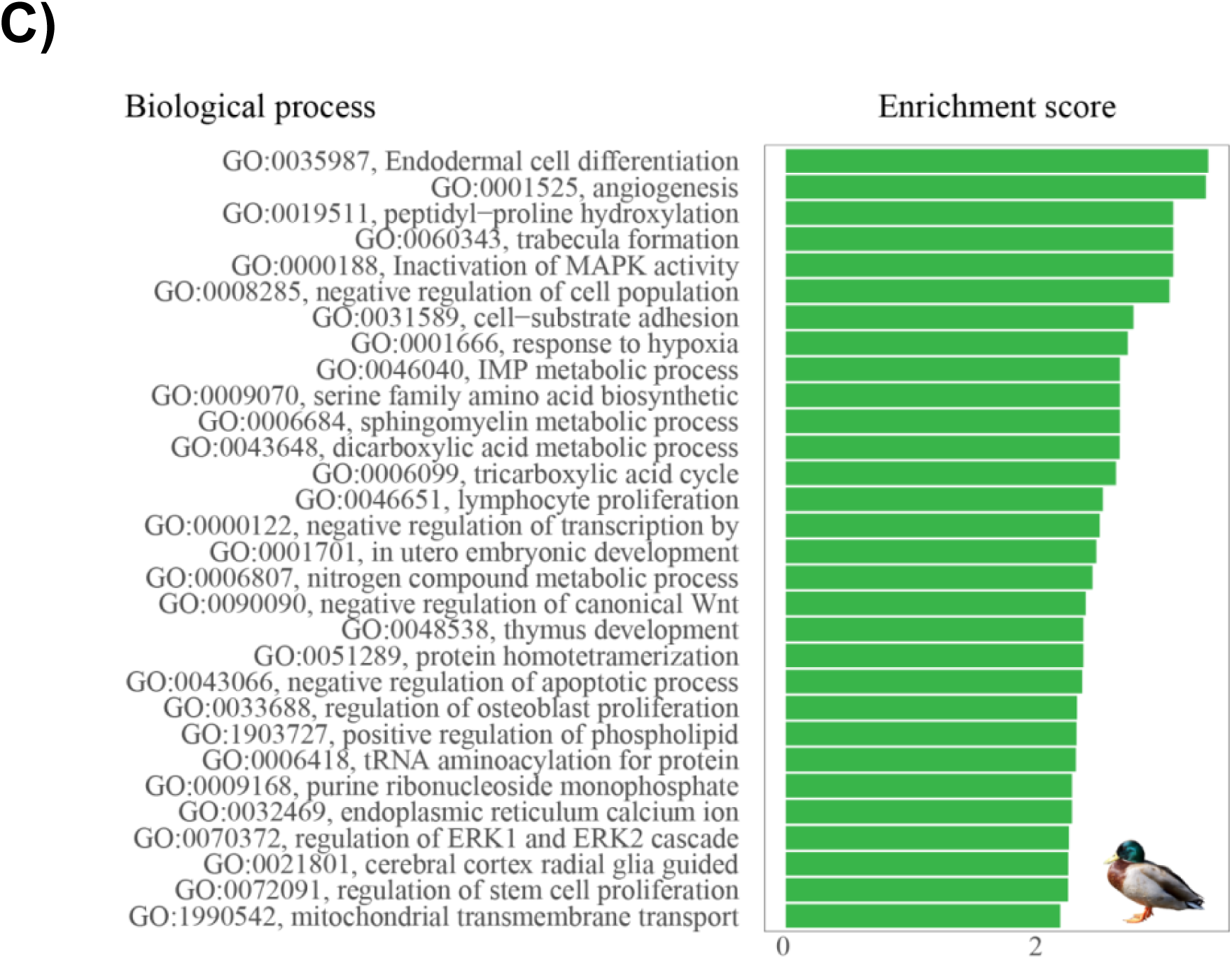
Top 30 GO biological process terms that were significantly enriched in infected A) black swan B) chicken and C) duck endothelial cells. The bars represent the enrichment score for the corresponding GO biological term with a p-value <0.05.

## DISCUSSION

The black and mute swan reference genomes provided herein represent the first publicly available swan genomes. The analyses of these genomes, together with the first black swan transcriptome in response to HPAI virus infection, has provided a unique insight into the plumage and immune system of the black swan.

The genomic insights provided by the present study were only possible due to the growing availability and accessibility of third generation sequencing. Specifically, older technologies that generate short read sequences can result in incorrect assembly, annotation errors and a large amount of manual effort to correct individual genes ^7^. In contrast, and consistent with the broader goals of the Vertebrate Genomes Project ^7^, the use of longer read sequences herein allowed us to generate black swan and mute swan genomes that were scaffolded to near chromosomal length and that were of comparable quality to the well-annotated chicken reference genome.

Genomic analysis of four genes known to be associated with plumage colour in other birds ^27^ identified a potential frame-shift mutation in the first exon of *SLC45A2* in the mute swan, which may have led to pseudogenization of this gene. *SLC45A2* encodes a transporter protein involved in melanin synthesis and is considered one of the most important proteins affecting human pigmentation ^28^. Mutations in the *SLC45A2* gene have been reported in albinism in humans ^29^. Furthermore, mutations in the gene have been associated with plumage colour variation in Japanese quails ^30^, indicating the importance of the *SLA45A2* in avian plumage. Interestingly, should a mutation of *SLC45A2* have resulted in the differential plumage of the black and mute swan it would suggest that the last common ancestor of these birds was, in fact, black. This is direct contrast to the metaphor of ‘black swan events’ that are so defined because of their unprecedented and unexpected nature. Instead, it would appear that at one point in history black plumage for the swan was the norm rather than the exception.

Compared to the last common ancestor, mute swan and mallard duck gene families involved in immune system processes were expansive. In contrast, no expansion in immune gene families was noted in the chicken or the black swan. This differential immune gene expansion, and its implications for susceptibility to HPAIV, are likely compounded by the observed impaired expression of TLR7 in the endothelial cells of black swans. Interestingly, other genes that have been observed to be differentially expressed between chickens and ducks, and implicated in susceptibility to HPAIV, were not differentially expressed between infected black swan and duck endothelial cells (e.g. RIG-I and IFITM3) ^31-34^. It is interesting to speculate as to whether mute swan endothelial cells would express *TLR7*. However, the presence or absence of *TLR7* in the endothelial cells of mute swans is perhaps irrelevant to the pathogenesis of HPAIV, as the virus is not heavily endothelial tropic in this species ^3^. In the black swan the observed differences in *TLR7* expression in endothelial cells speaks to the value of combining genomics with both primary cell culture and transcriptomics, as has recently been suggested as the new standard for comparative genomics by Stephan and colleagues ^35^. Either as a result of, or in addition to, these observed immune differences, black swan endothelial cells also mounted a markedly pro-inflammatory response to HPAIV infection. We have previously reported a similar pro-inflammatory in infected chicken endothelial cells (compared to those of ducks) and speculated that this inflammatory response leads to immunopathology in chickens *in vivo*^36^. Whether disease severity in black swans is driven by immunopathology remains to be determined, although it is consistent with the observed pathology in infected birds^37,38^. In sum, it is likely that this combination of species-specific differences in the immune response contribute to the marked susceptibility of both the black swan and chicken to HPAIVs.

The observed species dependent differences in the immune responses of swans raises the intriguing question as to why the black swan continues to thrive in its native Australia as well as in New Zealand (where it was introduced in 19^th^ century). This may be due to the fact that HPAI is not endemic in Australia and New Zealand. Indeed, captive populations of black swans located in parts of the world frequently exposed to HPAI are highly susceptible to severe disease^4^. The data presented in this study would therefore suggest that should HPAI become more prevalent in the Oceania region the ongoing survival of the black swan would be at significant risk. Moreover, many of the immune limitations described herein are not specific to avian influenza viruses. For example, TLR7 is essential in the immune recognition of a wide number of viral pathogens including avian coronaviruses^39^. These data suggest that should any avian endothelial specific viral infection become established in the native habitat of the black swan the survival of this iconic species would be in significant peril.

## ACKNOWLEDGEMENTS

We would like to acknowledge colleagues from the Friedrich-Loeffler-Institut (Jens Peter Teifke, Robert Klopfleisch and Angele Breithaupt) for providing FFPE material. Special thanks to Christoph Schulze from the state diagnostic laboratory was involved in the autopsy of most of the samples use for FFPE material collection and Ashling Charles from DNA Zoo Australia team for routine data processing support. Hi-C data for the Black Swan was created by the DNA Zoo Consortium (www.dnazoo.org). DNA Zoo sequencing effort is supported by Illumina, Inc. E.L.A. was supported by the Welch Foundation (Q-1866-20210327), an NIH Encyclopedia of DNA Elements Mapping Center Award (UM1HG009375), a US-Israel Binational Science Foundation Award (2019276), the Behavioral Plasticity Research Institute (NSF DBI-2021795), NSF Physics Frontiers Center Award (NSF PHY-2019745), and an NIH CEGS (RM1HG011016-01A1).

This study was funded by the Australian Department of Agriculture and Water Resources (CSIRO Science and Innovation Award; KRS), the Australian Department of Agriculture and Water Resources (Australian Eggs Innovation Award; KRS) and an Australian Research Council DECRA (DE180100512; KRS). This research was funded in part by the Wellcome Trust [Grant numbers 108749/Z/15/Z, 222155/Z/20/Z]. For the purpose of open access, the author has applied a CC BY public copyright licence to any Author Accepted Manuscript version arising from this submission. ACK was supported by the Australian Government’s research training program (RTP) scholarship. P.K. is supported by the University of Western Australia with additional computational resources and support from the Pawsey Supercomputing Centre with funding from the Australian Government and Government of Western Australia.

