## Supplementary material for "The swan genome and transcriptome: its not all black and white": Methods

A full description of the methods can be found in the Supplementary Note. No statistical methods were used to predetermine sample size.

#### Ethics statement

Animal ethics approval was obtained from the animal ethics unit, the University of Queensland's research ethics office (ANFRA/311/18; SCMB/002/18). A wildlife research permit was obtained from the Queensland Department of Environment, Land, and Water Planning under the Wildlife Act 1975 (permit number: 10008904). For bone marrow tissue collection in New Zealand, we obtained permission from the Department of Conservation under Wildlife Act Authority on non-public conservation land (70705-FAU & 70709-DOA). Ethical approval to sample mute swans was obtained by the Linköping Animal Ethics board (permit 01125-2020).

### Black swan genome

#### Sequencing

For genomic DNA and RNA extraction, liver, spleen, kidney, skeletal muscles, lung, heart, stomach, intestines, and gonads were obtained from an adult male black swan that had to be euthanised at Currumbin Wildlife Sanctuary hospital (https://currumbinsanctuary.com.au/wildlife-hospital) due to injury. Genomic DNA from the stomach tissue was extracted using a commercially available kit (MagAttract HMW DNA kit – Qiagen®). A DNA library was prepared using Single Molecular Real-Time (SMRT) bell template Prep Kit 1.0 (Pacific Biosciences) with unsheared gDNA with a size cut-off of 20kb using a Blue Pippin instrument (Sage Science ©) according to the manufacturer's protocol. SMRT sequencing was performed using PacBio Sequel instrument at the Institute of Molecular Biosciences, the University of Queensland using 1-6 pica Molar (pM) loading concentration, version 3.0 sequencing chemistry, diffusion loading, and 10-hours movies over 10 SMRT cells (1M V.3).

#### Genome Assembly

Raw SMRT sequencing reads were used for *de novo* black swan genome assembly using open-source FLACON(v2018. 31-08-03.0)/FALCON-UNZIP (6.0.0.47841) diploid aware genome assembly algorithms to produce primary and alternative haplotigs as the primary assembly. For FALCON assembly, the estimated genome size (haplotype) used for the black swan was 1.4181 x 10^9 base pairs. Additional parameters used for FALCON assembly are listed in Supplementary Table 14. Raw FASTA sequences were extracted from the subread BAM files produced by SMRT sequencing using PacBio® BAM2fasta tool. Raw FASTA sequences from 10 SMRT cells were given as the input for the FALCON assembler, and the FALCON assembly parameters were provided using a configuration file. Once the FALCON run was completed, FALCON-UNZIP was executed in the same directory as the initial FALCON run. For the FALCON-UNZIP run, the subread BAM data was also used. Once the FALCON UNZIP run was complete, two genome polishing steps were undertaken through PacBio® SMRT Link version 7.0.0 resequencing pipeline using PacBio CLR reads with default settings (-x 30) in addition to FALCON_UNZIP integrated arrow polishing to produce the primary PacBio assembly. This PacBio assembly was screened for the presence of mitochondrial/non-chromosomal DNA suing NCBI Blast+(v2.9.0+) against the black swan mitochondrial genome[1]. The mitochondrial DNA sequences were removed from the primary PacBio assembly.

PURGE_DUPS (v1.2.5) was next employed to remove false haplotypic duplications from the concatenated primary PacBio assembly and alternate haplotigs. The PacBio assembly completeness was assessed via BUSCO V.5.1.2 against avian orthoDB10. The presence of highly conserved core vertebrate genes was evaluated using the CEGMA pipeline, available online in gVolante (https://gvolante.riken.jp/).

The FALCON draft was scaffolded to chromosome-length by the DNA Zoo Consortium following the methodology described here: www.dnazoo.org/methods. Briefly, the Hi-C data was processed using Juicer [Durand, Shamim et al., 2016], and used as input into the 3D-DNA pipeline [Dudchenko et al., 2017] to produce a candidate chromosome-length genome assembly. We performed additional finishing on the scaffolds using Juicebox Assembly Tools [Durand, Robinson et al., 2016; Dudchenko et al., 2018]. The contact matrices generated by aligning the Hi-C data to the genome assembly before and after the Hi-C scaffolding are available for browsing at multiple resolutions on https://www.dnazoo.org/assemblies/Cygnus_atratus visualized using Juicebox.js, a cloud-based visualization system for Hi-C data [Robinson et al., 2018]. Finally, the Hi-C assembly was subjected to gap closing using the TGS-GapCloser(v1.0.1) algorithm[2] with default settings. PacBio continuous long reads (CLRs) were used as input for the software tool for gap closing.

The *Anatidae* family's established terminology for the chromosome-length scaffolds was based on the homology between black swan scaffolds/chromosomes and the closely related mute swan. MUMMER(v3) was used to establish a 1:1 homology between the black swan and the mute swan before assigning the macro and microchromosomes terminology to black swan chromosomes.

### Mute swan genome

#### Sequencing

The mute swan genome was sequenced by the vertebrate genome project (VGP) according to the "vertebrate genome project assembly phase I" pipeline. The genomic DNA was extracted from the stomach tissue of a male adult mute swan. The sequencing technology and data used to build the mute swan haploid chromosome length hybrid assembly included PacBio Sequel I CLR, Illumina Novaseq®, Arima® Hi-C linkage data, and BioNano optical maps

#### Assembly

FALCON (v2018. 31-08-03.0)/FALCON-UNZIP (6.0.0.47841) was used to generate the phased contigs. To remove the false duplications in the phased haplotigs, Purge_Haplotigs(v1.0.3+) was used. After that, two rounds of scaffolding were performed using 10x Genomics link data with scaff10x (v4.1.0) (https://github.com/wtsi-hpag/Scaff10X) for the first step of scaffolding with 10xGenomics® link data to build the S1 scaffold set. Next, the second step of scaffolding was performed on S1 using BioNano data.

BioNano solve (v3.2.1_04_122018) software was used to produce the S2 scaffold. The last scaffolding was performed to scaffold the S2 set using Arima® Hi-C linkage data with SALSA (v2.2)[3,4]. Primary and alternate haplotigs were concatenated, and bases were polished with Arrow (SMRTanalysis v6.0.0.47841) using PacBio CLR reads. Two more rounds of polishing were performed with linked reads by aligning with Longranger (v2.2.2) and variant calling with FreeBayes (v1.3.1) followed by consensus calling with bcftools consensus. Finally, manual curation of the assembly was performed using gEVAL (v2019-09-26).

The chromosome assignment was performed using evidence from Hi-C. A scaffold was considered a chromosome with Hi-C evidence when there is a clear unbroken diagonal in the Hi-C maps in the Juicebox. Every Hi-C box from the largest to the smallest for evidence-validated scaffolds was considered as a chromosome. Subsequently, established terminology for the chromosomes was given for the *Anatidae* family.

**Genome annotation**

The final assembly's structural and functional annotation of genes was conducted using *de novo* homology-based and evidence-based methods (for details see online supplementary method). Firstly, the primary PacBio assembly was annotated using NCBI eukaryotic and Ensembl gene annotation pipelines. A set of transcriptomic data (of both short-reads and long reads) derived from the liver, spleen, peripheral blood mononuclear cells and primary bone marrow derived endothelial cells was used for black swan genome annotation, and the ISOseq data for mute swan annotation was derived from the peripheral blood.

**Structural variant assessment of the genes associated with plumage colour in swans**
Structural variants in solute carrier family 45 member A2 (*SLC45A2*), PRELI domain containing 3B (*SLMO2*), ATP synthase, $H_+$ transporting, mitochondrial F1 complex, epsilon subunit (*ATP5e*), and Endothelin3 (*EDN3)* were examined using the University of California, Santa Cruz (UCSC) genome browser with multiple whole-genome alignments. We first aligned four Gallianseriformes' whole genomes (black swan, mute swan, chicken and mallard duck) with CACTUS to produce a HAL alignment file[5]. The HAL file alignment was then visualised in the Genome assembly In a Box (GBiB) using the HAL tools package (V2.1). We then used predicted protein and mRNA sequences encoded by the *SLC45A2, SLMO2, ATP5e,* and *EDN3* genes (from all four species) as queries to perform protein coding region alignment with the BLAT built in the GBiB and compare with each species in the HAL alignment. We validated putative open reading frames (ORF) for each gene from each species by visual inspection. Validity was assessed based on the presence or absence of putative donor-recipient splice sites, start codon and in-frame stop codons.

**Gene family evolution**

The longest peptide sequence for a given protein-coding gene for 17 species was retrieved from Ensembl (release 104) and NCBI Annotation release 103, including Zebrafish (*Danio rerio*) (GCA_000002035.4), Japanese rice fish (*Oryzias latipes*) (GCF_002234675.1), three-pinned stickleback fish (*Gasterosteus aculeatus*) (BROAD S1), Tropical clawed fish (*Xenopus tropicalis*) (GCA_000004195.3), Platypus (*Ornithorhynchus anatinus*) (GCA_004115215.2), Gray short-tailed opossum (*Monodelphis domestic*) (GCA_000002295.1), Mouse (*Mus musculus*) (GCA_000001635.9), Human (*Homo sapiens*) (GCA_000001405.28), Cattle (*Bos taurus*) (GCA_002263795.2), Anole lizard (*Anolis carolinensis*) (GCA_000090745.2), Mallard duck (*Anas platyrhynchos*) (GCA_002743455.1), Turkey (*Meleagris gallopavo*) (GCA_000146605.4), Chicken (*Gallus gallus*) (GCA_000002315.5), Zebra finch (*Taeniopygia guttata*) (GCA_003957565.2), and collared flycatcher (*Ficedula albicollis*) (GCA_000247815.2). The longest amino acid sequences from coding genes in the final reference annotations were used for the black and mute swan. The phylogenetic tree for the above 17 species was constructed using the Orthofinder pipeline with – M msa -S blast -I 1.3 settings. The final rooted species tree was inferred using STAG from the Orthofinder, which was then made ultra-metric with the root-age of 435 million years (obtained from TimeTree.org database). Peptide sequences in 17 species were scored using PANTHER (v15.0) database and clustered into PANTHER families and subfamilies and then identified by a family-specific PANTHER identifier. The maximum likelihood of gene family evolution for a given taxon was estimated by running CAFÉ(v5) with the PANTHER assigned protein families of the 17 species and the Orthofinder-based ultrametric species tree (significant at *p-value* < 0.05).

**Functional profiling of expanded and contracted gene families in swans**

Each PANTHER sub-family identification with a significant gene family evolution was assigned to the gene ontology category (GOslim identification number) to examine the biological role of significantly evolving (either by expansion or contraction) (Vertibri *p-value* < 0.05) gene families through the r-package PANTHER.db. The significant gene families that appeared to have expanded or contracted were separated. Subsequently, GO enrichment analyses were conducted using topGO[6] to test for over-represented functional terms associated with expanded and contracted gene families(a given gene family is considered expanded if the number of genes are increased compared to last common ancestor). The significant gene sub-families annotated with GOslim identifications were used as the foreground, and the GOslim annotated gene subfamilies of the last common ancestor with at least one gene present in the family were used as the background. Fisher's exact test was used for statistical testing, and the terms with corrected *p-value* < 0.01 were chosen as overrepresented GO terms.

**Comparison of the immune gene repertoire**

Selected immune databases were used to obtain the list of known immune genes from Ensembl release and were compared to that of the black swan and mute swan genome (see online methods for details)

**Major histocompatibility complex (MHC) class annotation in black and mute swan genomes**

The major histocompatibility complex (MHC) loci (MHC class I and MHC class IIb) for both mute and the black swan genomes were identified as previously described [7]. Briefly, the avian order-level consensus sequences for MHC I and MHC II loci were built using several different bird species covering at least three exons (Exon 2, exon 3, and exon 4). The blast algorithm aligned the order-level consensus sequences to the black swan and mute swan genomes. MHC loci were estimated as the number of blast hits that contained all three exons within 2kb of each other. Each exon was examined for in-frame stop codons, and it was eliminated as a locus if present. Genes present in regions 100kb upstream and downstream of each locus were manually annotated and the gene location plotted through genes to identify the additional genes of each predicted locus. Each predicted MHC locus was manually inspected for premature stop codons, and if present, were eliminated. Identified MHC molecules and MHC B loci associated genes were visualized for black and the mute swan for comparison using Circlize (R-package).

***TLR7* expression in black swan tissues**

Total RNA was extracted from tissues of an adult male black swan (stomach, kidney, and liver) using an RNA plus mini kit (Qiagen, Hilden, Germany) according to the manufacturer's instructions. Genomic DNA was extracted from the stomach muscles using the MagAttract HMW DNA kit (Qiagen®, Hilden, Germany) for the positive control, following the manufacturer's recommendation. The black swan-specific *TLR7 (FW: TTGCACTTCCACACTCCAAG; RVTCAGTCCAATTGCACCTCTG; Probe:CTCCGAAACAATCGCATTCAACGG)* and *18S (FW: CCTGCGGCTTAATTTGACTC; RV: AGACAAATCGCTCCACCAAC; Probe:TTGAGAGCTCTTTCTCGATTCCGTGG)* primers and probes were designed using the primer3plus web tool

**Immunohistochemistry**

Immunohistochemistry for influenza A virus nucleoprotein (NP) was performed according to the online supplement.

**Isolation of primary black swan, chicken and duck endothelial cells**

We used previously isolated and characterized chicken and duck primary aortic endothelial cells as the controls for black swan primary endothelial cell characterization[8,9]. Briefly, seventeen-day-old chicken and twenty-one-day-old Pekin duck embryonated eggs were purchased from Darling Downs Hatchery (Queensland, Australia). Primary aortic endothelial cells were cultured from the aortic arches of chicken and duck embryos as described previously using EGM-2MV medium (Lonza, Basel, Switzerland) with 10% FBS (Gibco, Waltham, MA, USA) [305, 307]. All avian cells were cultured at 40 °C 5% $CO_2$ unless otherwise stated.

The black swans used for cell isolation were sourced from Currumbin Wildlife Animal Hospital (-28.14, 153.48), Queensland, Australia, and Tauranga Harbor, Bay of Plenty, New Zealand (-37.57, 175.97). The animals were either euthanized due to a terminal illness or as part of a government approved cull. Tibiotarsi and femurs were obtained from all birds. The bones were then partially opened in the middle to expose the bone marrow and then submerged in pre-chilled (4◦C) Dulbecco's modified Eagle medium (DMEM) in T175 flasks. Flasks were then transferred to a personal containment level 2 (PC2) laboratory for further processing within four to six hours *post mortem*.

Bone marrow was subsequently removed and resuspended in pre-chilled DMEM. A 15ml suspension of DMEM containing bone marrow was then layered after filtering through a 40µm sterile filter over 15ml of Lymphoprep® in a 50 ml SepMate® tube. The layer of cells at the DMEM-Lymphoprep interface was separated according to the SepMate manufacturer's instructions. The separated cell layer was then resuspended in 1ml EGM-2MV media and enumerated. The cell concentration was adjusted to approximately 1 x $10^6$ cells per ml before transferring to a Petri dish with 10ml EGM-2MV media. The Petri dish was incubated at 40◦C with 5% $CO_2$. The media was replaced every other day until cells became confluent. Bone marrow-derived primary cells (EPC) of passage 4 were cultured to confluency and incubated with 1.75µg/ml Alexa Fluor™ 488 Acetylated low-density lipoprotein (ac-LDL) (Invitrogen, USA) for 4 hours. The cells were detached by washing twice with PBS and incubating with 0.05% Trypsin-EDTA (Thermo Fisher®, USA). The cells were mixed in 2% (v/v) FBS in PBS and subsequently sorted with the Moflo Astrios ® High-Speed cell sorter (Beckman Coulter®, CA, USA) under sterile conditions. Ac-LDL-stained cells were sorted based on prior knowledge of high and low ac-LDL intake [303] using BD FACSDiva™ (version 8.0.1 – BD©). The sorted cells were counted and transferred to gelatine-coated plates and grown in EGM-2MV (Endothelial Basal Media with EGM – 2MV bullet kit (Lonza©). Cells at post-sorted passages were known as black swan primary endothelial cells (bsEPCs). The purity of bsEPCs was confirmed as described by the online supplementary methods.

Modelling HPAI in avian endothelial cells

The experiment was designed in to include two groups viz. challenge and control, from each of the species, i.e., Black swan, chicken and duck. Each group consisted of three biological replicates of endothelial cells.

**Viral infection**

A/Chicken/Vietnam/0008/2004(H5N1) (VN04) virus was amplified in MDCK cells. All experiments using HPAIVs were performed under physical containment level 3 (PC3) settings at the Australian Centre for Disease Preparedness (Geelong, Australia). Primary endothelial cells were grown to at least 60% confluency and were inoculated with 5 x $10^5$ PFU/mL of VN04 for 60 min at 40◦C under physical containment 3 (PC3) settings. The inoculum was removed, and the cells were washed once with PBS, and EGM-2MV media without FBS was added. The cells were incubated for an additional five hours.

**RNA extraction**

According to the manufacturer's instructions, cellular RNA was extracted from cultured cells using the RNeasy Plus Kit (Qiagen, Hilden, Germany). Subsequently, 3M of sodium acetate (NaOAc, pH 5.5) and 100% ethanol were added for RNA precipitation.

**RNA sequencing**

Paired-end, 150bp long, RNAseq was performed by Macrogen (Macrogen, Seoul, South Korea) approximately 40 million reads per sample. Library preparation was performed using the SMARTer® ultra-low kit (Takara Bio® USA, Inc.). All the libraries were barcoded and subsequently sequenced using the Novaseq® 6000 (Illumina, San Diego, CA, USA) with Novaseq® 6000 S4 Reagent Kit (Illumina, San Diego, CA, USA) as per the manufacturer's instructions using within a single lane in a single cell to avoid potential technical variations. RNAseq reads were separated into original samples based on the corresponding bar code.

**Differential gene expression analysis**

RNAseq-based differential gene expression analysis was performed after assessing the quality of the raw reads using the FastQC tool before and after adaptor and quality trimming with TrimGalore version 0.6.2 (RRID: SCR_011847). RNAseq reads were then quantified at the transcript level using Salmon v1.5.2 [319]. First, the indexes were created for the black swan reference transcriptome and the Ensembl version 104 chicken (GRCg6a) and duck (CAU_Duck1.0) transcriptomes with following parameters (salmon quant -i (params.ind) -l (params.libtype) -g (params.genemap) -r input_r1 -f input_r2 --validateMappings -o (output.dir)). The transcript abundance data of each species were imported through the R package 'tximport' by transforming data from transcripts to gene-level quantification. The expression data from each species (black swan, chicken, and duck) were independently analysed for differential gene expression. Once RNAseq read libraries were normalized for sequencing depth and RNA composition, differential gene expression analysis was performed using DESeq2 version 1.30.1[11]. A Bonferroni adjusted *p-value* (*adj.p.val*) cut-off of 0.05 and $\log_2$fold enrichment of 0.58 (fold-change 1.5) were used as the significance threshold (differential gene expression: mock vs. infected).

**Gene Ontology enrichment analysis**

Gene Ontology (GO) terms were assigned to all quantified genes in chicken and duck using the BioConductor package BioMart[12], and the GO terms were assigned to all expressed and quantified genes using InterProScan5[13]. For black swan genes, we used InterProScan 5[13] for GO annotation. Using the GO annotation, we then performed GO enrichment analysis with topGO[6]. For the statistical enrichment, Fisher's exact test was used with the 'weight01' algorithm. GO enrichment analyses were performed for all three GO categories, including 'biological process,' 'cellular component,' and 'molecular function'. Results of the GO enrichment were visualized with 'ggplot2' and 'GOplot'[14] R packages. We calculated a score (named as "z-score" in the GOplot package) to evaluate the trend (increasing or decreasing biological process) using GOplot in-built mathematical formula (Formula 1) for enriched terms of the GO biological process category in each species.

**Code availability**

The software tools for black swan genome assembly are available as a bioconda repository - Pb-falcon v0.3.4 (https://anaconda.org/bioconda/pb-falcon). Further details are in Table 1.

**Table 1: The script usage and availability**

| Usage | Script repository link |
|---|---|
| Genome assembly | https://github.com/akaraw/Black_swan_genome_assembly |
| MHC class annotation | https://github.com/akaraw/MHC_complex_annotation_of_swans |
| Genome annotation | https://github.com/akaraw/swan_genome_annotation |
| RNAseq data analysis | https://github.com/akaraw/RNA-Seq_Endothelial_cells |
| Gene family evolution | https://github.com/akaraw/Gene_family_evolution_in_swans |
| Comparison of immune gene repertoire | https://github.com/akaraw/Gene_family_evolution_in_swans |

**Data availability**

| | | | | | |
|---|---|---|---|---|---|
| Swan assembly - primary | Black swan | GCA_013377495.1 | PRJNA640810 | PRJNA640810 | NCBI Refseq |
| swan assembly – Final | Black swan | N/A | N/A | https://www.dnazoo.org/assemblies/Cygnus_atratus | DNAzoo |
| swan assembly – Final | Mute swan | GCA_009769625.2 | GCA_009769625.2 | GCA_009769625.2 | NCBI Refseq |
| Swan ISOseq | Black swan | N/A | PRJEB38872 | https://www.ncbi.nlm.nih.gov/bioproject/PRJEB38872 | NCBI Refseq |
| Swan ISOseq | Mute swan | N/A | PRJEB38872 | https://www.ncbi.nlm.nih.gov/bioproject/PRJEB38872 | NCBI Refseq |
| PacBio raw reads | Black swan | | PRJEB44719 | https://www.ncbi.nlm.nih.gov/bioproject/?term=PRJEB44719 | NCBI Refseq |
| PacBio raw reads | Mute swan | N/A | GCA_009769625.2 | https://vgp.github.io/genomeark/Cygnus_olor/ | Genome Ark |
| RNAseq | Black swan | RNAseq - Endothelial cells | PRJEB45365 | https://www.ncbi.nlm.nih.gov/bioproject/?term=PRJEB45365 | NCBI Refseq |
| Scaffolding | Black swan | Hi-C Illumina data | PRJNA512907 | https://www.ncbi.nlm.nih.gov/sra?LinkName=biosample_sra&from_uid=21582233 | NCBI SRA |

Raw Hi-C data is available on SRA (PRJNA512907, SRX12373281).

1 Jiang, F. *et al.* The complete mitochondrial genomes of the whistling duck (Dendrocygna javanica) and black swan (Cygnus atratus): dating evolutionary divergence in Galloanserae. *Molecular biology reports* **37**, 3001-3015 (2010).

2 Xu, M. *et al.* TGS-GapCloser: a fast and accurate gap closer for large genomes with low coverage of error-prone long reads. *Gigascience* **9**, giaa094 (2020).
3 Ghurye, J., Pop, M., Koren, S., Bickhart, D. & Chin, C.-S. Scaffolding of long read assemblies using long range contact information. *BMC genomics* **18**, 1-11 (2017).
4 Ghurye, J. *et al.* Integrating Hi-C links with assembly graphs for chromosome-scale assembly. *PLoS computational biology* **15**, e1007273 (2019).
5 Hickey, G., Paten, B., Earl, D., Zerbino, D. & Haussler, D. HAL: a hierarchical format for storing and analyzing multiple genome alignments. *Bioinformatics* **29**, 1341-1342 (2013).
6 Alexa, A. & Rahnenfuhrer, J. topGO: enrichment analysis for gene ontology. *R package version* **2**, 2010 (2010).
7 He, K., Minias, P. & Dunn, P. O. Long-read genome assemblies reveal extraordinary variation in the number and structure of MHC loci in birds. *Genome biology and evolution* **13**, evaa270 (2021).
8 Davis, R. L. *et al.* The culture of primary duck endothelial cells for the study of avian influenza. *BMC microbiology* **18**, 1-9 (2018).
9 Tong, Z. W. M. *et al.* Primary Chicken and Duck Endothelial Cells Display a Differential Response to Infection with Highly Pathogenic Avian Influenza Virus. *Genes* **12**, 901 (2021).
10 Short, K. R. *et al.* Using bioluminescent imaging to investigate synergism between Streptococcus pneumoniae and influenza A virus in infant mice. *JoVE (Journal of Visualized Experiments)*, e2357 (2011).
11 Love, M. I., Huber, W. & Anders, S. Moderated estimation of fold change and dispersion for RNA-seq data with DESeq2. *Genome biology* **15**, 1-21 (2014).
12 Durinck, S., Spellman, P. T., Birney, E. & Huber, W. Mapping identifiers for the integration of genomic datasets with the R/Bioconductor package biomaRt. *Nature protocols* **4**, 1184-1191 (2009).
13 Jones, P. *et al.* InterProScan 5: genome-scale protein function classification. *Bioinformatics* **30**, 1236-1240 (2014).
14 Walter, W., Sánchez-Cabo, F. & Ricote, M. GOplot: an R package for visually combining expression data with functional analysis. *Bioinformatics* **31**, 2912-2914 (2015).
