## Supplementary material for "The swan genome and transcriptome: its not all black and white": Online Methods

### ISO-Seq full-length transcript data for evidence-based annotation

Total RNA from four different cell/tissue types (the kidney, the liver, peripheral blood mononuclear cells, and primary endothelial cells) from black swan and blood cells from mute swan were extracted using RNA plus mini kit (Qiagen) and Invitrogen™ RiboPure™ RNA Purification Kit (Thermo Fisher Scientific), respectively, following the manufacturer's protocol. ISO-Seq libraries were prepared using the NEBNext™ Single Cell/Low Input cDNA Synthesis and Amplification Module (NEB, E6421S) and the SMRTbell™ Express Template Prep Kit 2.0 (PacBio, 100-938-900) according to the standard protocol for ISO-Seq Express Template Preparation for Sequel and Sequel II Systems (PacBio, Part # 101-763-800 v1). Total RNA (300 ng per sample) was reverse transcribed into cDNA and amplified to generate at least 160 ng of cDNA for library preparation. During amplification, different unique barcodes were incorporated into the cDNA fragments for each sample. The amplified cDNA was purified with ProNex™ beads (Promega, NG2001) using the standard workflow indicated in the ISO-Seq protocol. The refined, size-selected cDNA for each sample was pooled equimolar, and a single library was prepared for the pooled cDNA. 160-500 ng of pooled/purified, size-selected cDNA was used in a DNA damage repair reaction, followed by an end-repair/A-tailing reaction. Overhang adapters were ligated to the poly A-tailed library fragments, followed by purification with ProNex™ beads. The final purified libraries were quantified on the Qubit fluorometer using the Qubit double-stranded DNA (dsDNA) HS assay kit (Invitrogen, Q32854) to assess the concentration and the Agilent BioAnalyzer 2100® using the High Sensitivity DNA kit (Agilent, 5067-4626) to assess the fragment size. Sequencing was performed using the PacBio Sequel II (software/chemistry v8.0.0). The library was prepared for sequencing using the standard protocol for diffusion loading with ProNex™ bead purification, Sequencing Primer v4, Sequel II Binding Kit v2.0 and the Sequel II DNA Internal Control v1. The polymerase-bound library was sequenced on 1 SMRT Cell, each with a 24 h movie time plus a two-hour pre-extension using the Sequel II Sequencing 2.0 Kit (PacBio, 101-820-200) and an SMRT Cell 8M (PacBio, 101-389-001). Library preparation and sequencing were performed at the Institute for Molecular Bioscience Sequencing Facility (The University of Queensland) (black swan) and the National Genomic Infrastructure of Sweden (mute swan).

The generated PacBio single-molecule sequencing data were processed using ISo-Seq version 3 incorporated in SMRT Link v8.0. The FLNC BAM files were used to extract FLNC fasta ("SAMTOOLS fasta" command: SAMTOOLS package) files for downstream analysis with Transcriptome Annotation by Modular Algorithms (TAMA) package available on GitHub.

The FLNC fasta files (from different tissue/cell libraries) were aligned to the final black swan genome assembly using minimap2 with the following options: "-ax splice -uf --secondary=no --splice-flank=no -C5 -O6,24 -B4". The sorted BAM files were created for each FLNC alignment using the SAMTOOLS package [221]. These BAM files were then processed using TAMA collapse in two phases. The first TAMA collapse was executed with options: "-p 1006_first -a 100 -z 100 -x no_cap" and the second phase with the options: "-a 100 -z 100 -x no_cap -sjt". Once both runs were over, the results were merged to obtain the final transcript models using TAMA merge. Finally, the "bedtools getfasta" tool was used to obtain the full-length transcripts in fasta format for downstream analysis with MAKER3 genome annotations software.

### Short-read RNA-seq for evidence-based annotation

Short read RNA-seq data was generated using six independent black swan endothelial cell cultures. RNA-seq reads after quality control from these six different libraries were aligned to the final black swan genome assembly using STAR software version 2.7.0a to produce BAM alignment files. These BAM files were used for downstream genome annotations with MAKER3.

**Homology-based genome annotation**

Non-redundant transcript evidence from the NCBI RefSeq database for Chicken and Zebra finch was downloaded. These transcripts were then used as evidence for black swan genome annotation using MAKER3. Avian proteomes, including chicken, mallard duck, and zebra finch, downloaded from the Ensembl or NCBI RefSeq databases, were used as protein homology evidence for the annotation with GeMoMa. Results of both NCBI and Ensembl annotation pipelines for the publicly available black swan primary assembly were also obtained as evidence for homology-based annotation of the final black swan genome.

**Ensembl annotation of the primary black swan genome**

Annotation of the primary assembly was created in collaboration with the Ensembl gene annotation system. A set of potential transcripts was generated primarily through alignment of transcriptomic data (both Illumina short reads and PacBio ISo-Seq long reads) and secondarily through gap filling with homology-based evidence. For constructing the transcriptome-based models, the short-read RNA-seq data were sourced from endothelial cells generated as part of this project (PRJEB13246) and aligned to the genome as described in detail in Aken et al. (2016)[1]. The long-read data were sourced from blood, endothelial cells, and the kidney generated as part of this project (PRJEB38871) and aligned using minimap2. Homology-based models were generated in two ways. Firstly, UniProt vertebrate proteins with experimental evidence at the protein or transcript level (termed protein existence levels 1 and 2 in UniProt) were downloaded and aligned in a splice-aware manner to the primary black swan genome using GenBlast. The second method involved mapping the coding exons for each canonical transcript in the human GENCODE reference gene set to the black swan assembly using the lastZ pairwise whole-genome alignment between the GRCh38 human assembly and the black swan assembly. Low-quality transcript models were removed at each locus, and the data were collapsed and consolidated into a final gene model. When collapsing the data, priority was given to models derived from transcriptomic data. The most extended open reading coverage for each putative transcript was assessed with known vertebrate proteins to help differentiate between true isoforms and fragments. In loci where the transcriptomic data were fragmented or missing, homology data took precedence, with preference given to longer transcripts with strong intron support from the short-read data. Gene models from the above process were classified into three main types: protein-coding, pseudogene, and long non-coding RNA. Models with BLAST [206] hits to known vertebrate proteins and few structural abnormalities (i.e., they had canonical splice sites, introns passing a minimum size threshold, and low repeat coverage) were classified as protein-coding.

**The National Centre for Biotechnology Information (NCBI/USA) annotation of the primary black swan genome and the mute swan genome**

The NCBI annotation pipeline was used for both swan genomes as described previously[2].

**MAKER3 annotation**

The MAKER3 annotation pipeline was used for both the black and mute swans based on homology and *ab initio*-based methods. MAKER3 annotation was performed in three iterative steps. The first round was performed on the repeat-masked genomes with homology-based evidence. The second round included *de novo* gene prediction using SNAP and Augustus software with pre-trained parameters. The Hidden Markov Models (HMMs) for annotated genes were generated using the first MAKER3 annotation results. The custom Augustus parameters were trained from the evidence supported by the ISO-Seq and RNAseq data and the genome to protein alignments. The second round of MAKER3 incorporated homology and *de novo* gene prediction evidence from the first round. For round three, the Augustus and SNAP were again retrained for gene prediction to be used in MARKER3. Both new *de novo* gene predictions and homology evidence were incorporated into the final MAKER3 annotation round.

**GeMoMa annotation**

Gene Model Mapper (GeMoMa) was used to identify the protein homology with closely related species. ISO-Seq reads aligned to the black and mute swan genomes were also used in the GeMoMa pipeline as evidence in addition to the homology evidence provided by other species (chicken, Mallard duck, and Zebra finch). GeMoMa was executed with the options "tblastn=false r=MAPPED ERE.c=false AnnotationFinalizer.r=NO GeMoMa.e=0.00001 d=DENOISE DenoiseIntrons.m=50000".

**PASA (Program to Assemble Spliced Alignments) transcript annotation**

PASA was used to annotate the non-redundant transcripts built from the TAMA pipeline to create transcript annotation in Gene Finding Format (GFF), which was then directly used with EvidenceModeller.

**Combining different annotations with evidence modeller**

EvidenceModeller was used to construct the final reference annotation by combining different annotations/evidence from different sources, including GeMoMa, Ensembl, NCBI RefSeq, PASA transcript and MAKER3 annotations. A "weight" file was created, which informed the weights or priorities of different annotation sources. *Ab initio* gene predictions were assigned with the lowest score, while the transcript-based annotation received the highest.

**Functional genome annotation**

Functional annotation of the final genome annotation reference set was conducted using BLAST and InterProScan tools. Final protein models of black and mute swan were aligned against UniProt/Swiss-Prot database (release 2021_3: accessed on June 2021) using the blastp in BLAST+ version 2.3.0. Protein models were scanned against several resources, including PANTHER-15.0, Pfam-33.1, ProSiteProfiles-2021_01, SUPERFAMILY-1.75, and TIGRFAM-15.0 (using InterPro version 86.0 data) with the pre-calculated match lookup service using InterProScan 5.52-86.0.

**AGAT (Another gene annotation tool)**

AGAT tools were used to curate the final reference gene sets of mute swan and black swan and parse the functional annotation arising from various resources, including the BLAST results against the UniProt/Swiss-Prot protein database and the results from the InterProScan tool. We used agat_sp_manage_functional_annotation.pl function in the AGAT tools to parse the functional annotation from protein level to gene level.

**Repeat annotation**

Custom repeat libraries were created for the mute and black swan genome assemblies. RepeatModeler2 [3] (v2.0.1, with the TE Tools container v1.2 of Dfam [4]; https://github.com/Dfam-consortium/TETools/tree/1.2) was run on each genome with default options plus the -LTRStruct option to detect long terminal repeat (LTR) retrotransposons. This produced 346 and 351 consensus sequences for the mute swan and black swan respectively. Of these, 242 were classified as unknown and 83 were LTRs in mute swan, and 244 were classified as unknown and 60 were LTRs in black swan.

Following this, each repeat library was compared to the curated bird repeat library from [5] and consensus sequences were removed from the swan libraries only if the classification was "unknown" and had a hit to the bird repeat library that was >95% identical across 90% of the swan repeat, and the bird repeat library was classified not as "unknown". Following this, consensus sequences were compared to swissprot [6] proteins via BLAST [7] to identify potential high-copy host genes which may have ended up in the swan repeat libraries. For sequences with BLASTX hits with an e-value below 0.1, the best hit per sequence was selected (based on highest score) to define gene identity. If the gene was not one with a domain known from major groups of transposable elements (e.g. reverse transcriptase, transposase) *sensu* [8], the sequence was considered a potential host gene and removed from the repeat library. Finally, each of the filtered swan repeat libraries were individually combined with the curated bird repeat library from [5] and these combined, species-specific libraries were used to mask each swan genome using RepeatMasker [9] with default settings and within the TE Tools container. Subsequently, each repeat library was compared to the curated bird repeat library from[10]. The consensus sequences were removed from the swan libraries only if, (i) the classification was "unknown" and had a hit to the bird repeat library that was >95% identical across 90% of the swan repeat and (ii) the bird repeat library was classified not as "unknown." Next, consensus sequences were compared to Swiss-Prot proteins via BLAST to identify potential high-copy host genes in the swan repeat libraries. For sequences with BLASTX hits with an e-value below 0.1, the best hit per sequence was selected (based on the highest score) to define gene identity. If the gene was not one with a domain known from major groups of transposable elements (e.g., reverse transcriptase, transposase), the sequence was considered a potential host gene and removed from the repeat library. Finally, each of the filtered swan repeat libraries was individually combined with the curated bird repeat library from[10]. These combined, species-specific libraries were used to mask each swan genome using RepeatMasker with default settings within the TE Tools container.

**Comparison of immune gene repertoire**

The immune databases included Immunogenetic Related information source (IRIS)[11] InnateDB (www.innatedb.com), Septic Shock group, MAPK/NFKB network, Import, and Immunome database (https://www.innatedb.com/redirect.do?go=resourcesGeneLists). The longest peptide sequences of these genes were obtained from Ensembl (release 104) and scored using InterProscan5 to identify and assign the gene to a gene sub-family. The number of gene members in each species (of the 17 species above) for the specified immune gene family was then calculated using a custom R-script. Proteins sequences derived from the InnateDB were assigned a gene sub-family separately to identify the genes belonging to innate immune gene families in each species.

Moreover, we evaluated the contractive immune gene repertoire by manual inspection to retain only the legitimate gene families where black swans have undergone contraction. Each immune gene family member was thoroughly examined individually and compared to the four proteomes, of the black swan, chicken, mute swan, and Mallard duck using BLAST. If a BLAST hit did not have over 80% identity with over 80% query coverage with proteins in black swans, it was suspected as an actual gene loss. Furthermore, those suspected genes were searched against Ensembl and NCBI black swan annotation through BLAST to validate the observation. Next, we generated phylogenetic trees (neighbour-joining trees for the given gene in a given family through the BLAST hits). If a given gene was present in at least one of the two annotations, the protein sub-family was removed from the black swan's contractive immune gene family list. If a protein was not present or flagged as a pseudogene, we examined both RNAseq and ISO-Seq alignments to validate if it lacked transcription, where if the pseudogene is not transcribed, then it was considered highly likely to be absent in the species. If the annotation for a given gene in black swan was doubtful (based on the phylogenetic tree) within a contracted immune gene family, we removed the family without further consideration.

**Immunohistochemistry for influenza virus antigen**

Immunohistochemical (IHC) labelling was performed on histological sections from a naturally infected black swan (A/Black swan/Akita/1/2016 (H5N6)). The sections from the black swan brain, spleen, and liver were kindly provided by Hiono *et al.*[12] Unstained paraffin sections were de-waxed and rehydrated by a series of xylene and ethanol washes. Slides were then incubated in 0.1% protease from *Streptomyces griseus* (Sigma-Aldrich, Missouri, USA) for 10 min at 37ºC and washed with deionized water. Endogenous peroxidase was blocked by incubating the slides in 3% $H_2O_2$ for 10 minutes at room temperature, followed by a wash with Tris-buffered saline (TBS) with 0.2% Tween 20 (TBST) (Sigma-Aldrich). Aldehydes were blocked by incubating the slides with 0.15M glycine in phosphate-buffered saline (PBS) for 15 minutes at room temperature, followed by a brief wash with TBST. Slides were then blocked using Dako antibody diluent with background reducing components (Agilent, California, USA) for 30 minutes at room temperature. Slides were incubated with a 1:400 dilution of monoclonal antibody (clone HB65 IgG2a) directed against the nucleoprotein (NP) of influenza A virus (American Type Culture Collection, Manassas, VA, USA) overnight at 4ºC followed by a series of TBST washes over 15 minutes. Slides were then incubated for 30 minutes at room temperature with HRP-conjugated anti-mouse immunoglobulin (EnVision kit, Agilent). Slides were subject to another series of TBST washes over 15 minutes. AEC chromogen substrate (Agilent) was added to the slides, and signal development was monitored on a microscope. Following the development, AEC was discarded, and slides were counterstained with Meyer's hematoxylin for 2 minutes. Following this, slides were washed briefly using TBST and finally with deionized water. The slides were mounted with VectaMount permanent mounting medium (Vector Laboratories, Burlingame, CA, USA). Brightfield IHC influenza and nuclei immunolabeling were visualized at 200x magnification using an Olympus microscope (Olympus©, Japan) and associated camera. Image processing was performed using ZEN blue software (version 3.3) (Zeiss).

### Confirmation of black swan endothelial cell cultures

Black swan endothelial cell (bsEPCs) identity was confirmed by micro tubule formation on bovine basement membrane extract, uptake of acetylated low-density lipoprotein, von Willibrand factor (vWBF) expression and the absence of CD45 expression.

bsEPCs, duck and chicken aortic endothelial cells of post-sorted passages 2-6 or MDCKs were grown to confluency and subsequently treated with 0.05% trypsin (Gibco®). Approximately 7.5 × $10_3$ cells per cell type, diluted in EGM-2MV, were aliquoted into a well of a 96-plate coated with 50 µL of solidified Cultrex® Basement Membrane Extract Type 2 (Trevigen, Gaithersburg, MD, USA). The plate was left to incubate at 37 °C 5% CO2 for 4h. The media were removed, washed gently with 1X PBS, and the cells visualized under BX51 Upright Light Microscope (Olympus, Tokyo, Japan).

Ac-LDL uptake was visualized under BX51 Upright Light Microscope (Olympus, Tokyo, Japan)

Primers for the *von Willebrand factor* (*vWBF*) a key endothelial marker, were designed for the black swan, chicken, and duck;

| Species | Gene | Forward | Reverse |
|---|---|---|---|
| **Black swan** | *vWBF* | GGGAAGTTCAACTCGGCATA | GCATGGTTCTGGTCACACAC |
| | *CD45* | ACTCGCTGTGAGGAAGGAAA | GCACTGCAATGAACCAC |
| **Chicken** | *vWBF* | TGTGCCCACAAAGCTGT | GGCCAAGTCCATCATCTTG |
| | *CD45* | ACTCGCTGTGAGGAAGGAAA | GCACTGCAATGAACCAC |
| **Duck** | *vWBF* | TGTGCCCACAAAGCTGT | GGCCAAGTCCATCATCTTG |
| | *CD45* | ACTCGCTGTGAGGAAGGAAA | GCACTGCAATGAACCAC |

In passage four, mRNA extracted from black swan primary cells, complementary DNA (cDNA) was synthesized with the High-Capacity cDNA Reverse Transcription Kit (Applied Biosystems, Waltham, CA, USA) and random primers according to the manufacturer's guidelines. RT-PCR was then conducted using the Phusion high-fidelity PCR kit (Biolabs, San Diego, CA, USA)

In these experiments. Madin-Darby Canine Kidney (MDCK) cells were used for control purposes. MDCK cells were obtained from the American Type Culture Collection (ATCC), and cultured in DMEM (Gibco, Waltham, MA, USA) supplemented with 10% fetal bovine serum (FBS) (Gibco, Waltham, MA, USA) and 1% penicillin/streptomycin (Gibco, Waltham, MA, USA). All cell lines of mammalian origin were incubated at 37 °C with 5% $CO_2$, whilst all avian cells were cultured at 40 °C with 5% $CO_2$ unless otherwise stated.


1 Aken, B. L. *et al.* The Ensembl gene annotation system. *Database (Oxford)* **2016**, doi:10.1093/database/baw093 (2016).

2 Thibaud-Nissen, F. *et al.* P8008 The NCBI Eukaryotic Genome Annotation Pipeline. *Journal of Animal Science* **94**, 184-184, doi:10.2527/jas2016.94supplement4184x (2016).

3 Flynn, J. M. *et al.* RepeatModeler2: automated genomic discovery of transposable element families. *bioRxiv*, 856591, doi:10.1101/856591 (2019).

4 Hubley, R. *et al.* The Dfam database of repetitive DNA families. *Nucleic Acids Res.* **44**, D81-D89, doi:10.1093/nar/gkv1272 (2016).

5 Peona, V. *et al.* The avian W chromosome is a refugium for endogenous retroviruses with likely effects on female-biased mutational load and genetic incompatibilities. *bioRxiv*, 2020.2007.2031.230854, doi:10.1101/2020.07.31.230854 (2020).

6 Boeckmann, B. *et al.* The SWISS-PROT protein knowledgebase and its supplement TrEMBL in 2003. *Nucleic Acids Res.* **31**, 365-370, doi:10.1093/nar/gkg095 (2003).

7 Altschul, S. F., Gish, W., Miller, W., Myers, E. W. & Lipman, D. J. Basic local alignment search tool. *J. Mol. Biol.* **215**, 403–410, doi:10.1016/S0022-2836(05)80360-2 (1990).

8 Wicker, T. *et al.* A unified classification system for eukaryotic transposable elements. *Nat. Rev. Genet.* **8**, 973-982 (2007).

9 Smit, A., Hubley, R. & Green, P. RepeatMasker Open-3.3.0. http://www.repeatmasker.org. (1996-2010).

10 Peona, V. *et al.* Identifying the causes and consequences of assembly gaps using a multiplatform genome assembly of a bird‐of‐paradise. *Molecular ecology resources* **21**, 263-286 (2021).

11 Kelley, J., de Bono, B. & Trowsdale, J. IRIS: a database surveying known human immune system genes. *Genomics* **85**, 503-511, doi:10.1016/j.ygeno.2005.01.009 (2005).

12 Hiono, T. *et al.* Characterization of H5N6 highly pathogenic avian influenza viruses isolated from wild and captive birds in the winter season of 2016–2017 in Northern Japan. *Microbiology and immunology* **61**, 387-397 (2017).
