## Supplementary Files for "The swan genome and transcriptome: its not all black and white"

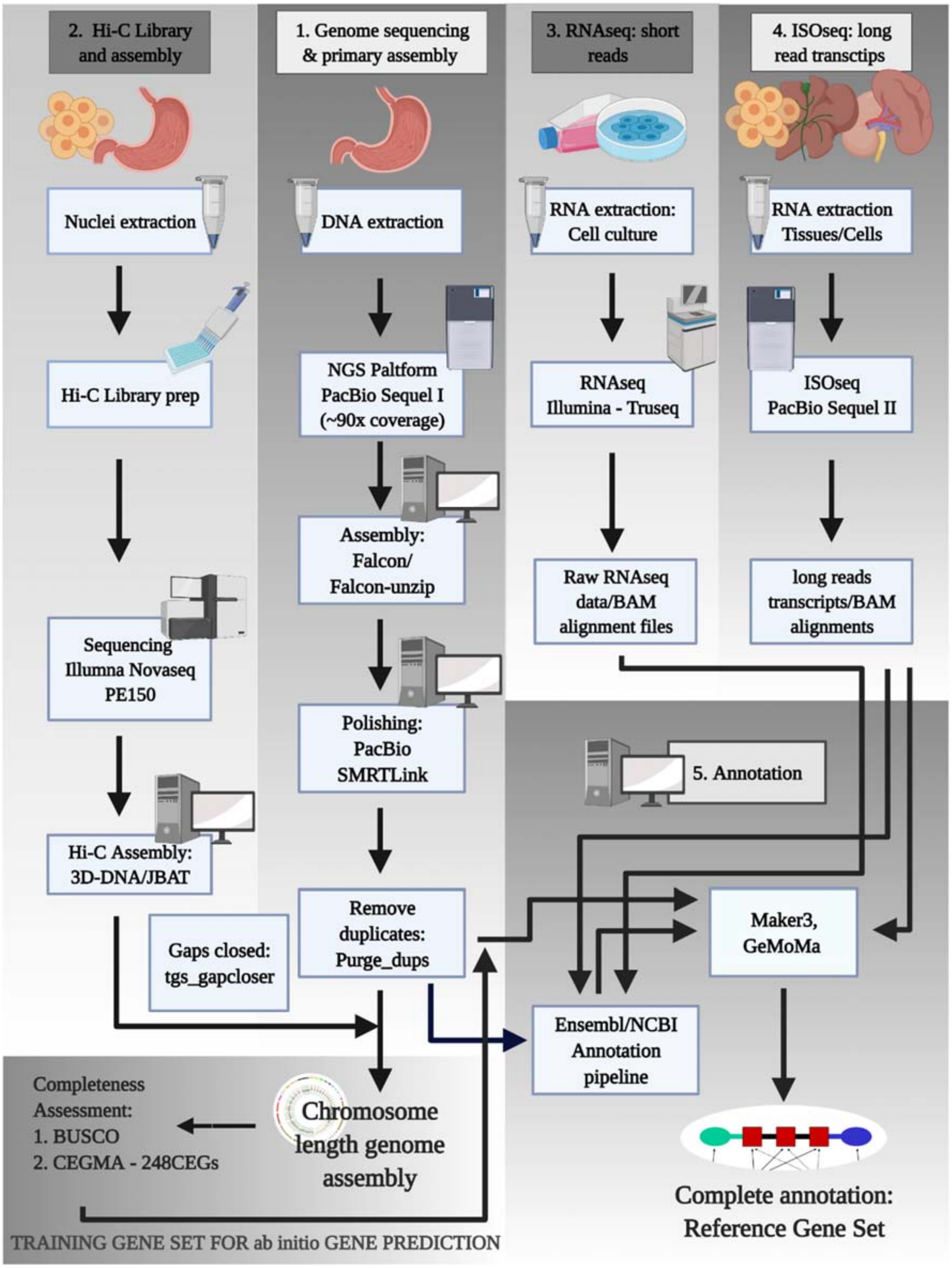


**Supplementary Figure 1:** Schematic overview of the process used to construct the black swan genome.

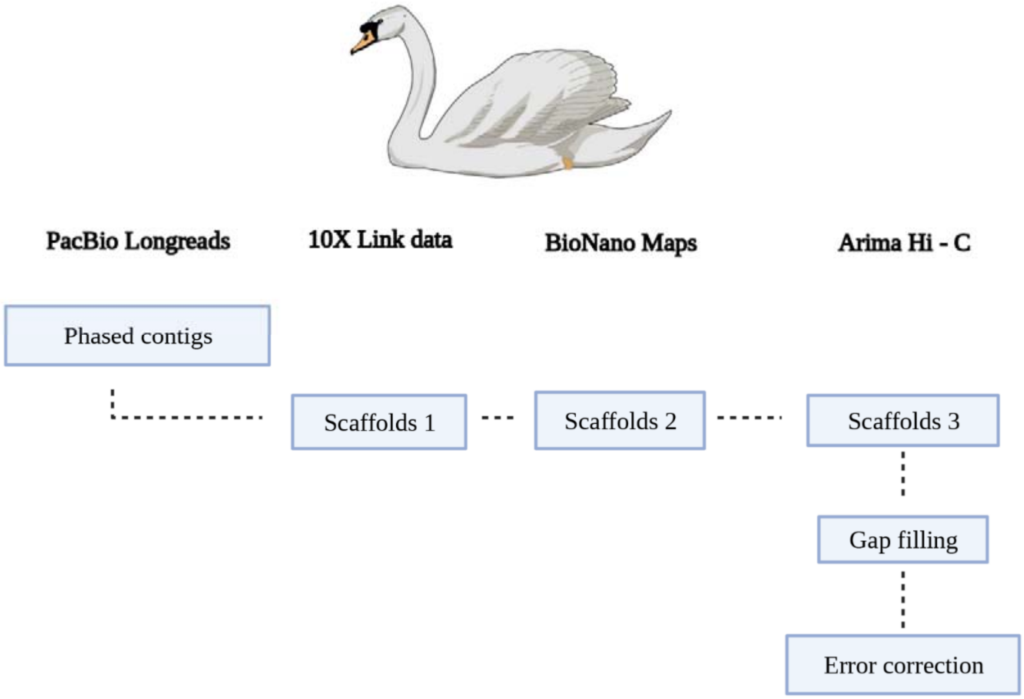


**Supplementary Figure 2:** Schematic overview of the assembly pipeline version 1.5 of the vertebrate genome project used to construct the mute swan genome.

**Supplementary Table 1:** BUSCO analysis of the final genomes

| BUSCOs categories | Black swan | Mute swan | Chicken (bGalGal1) |
|---|---|---|---|
| **Complete** | 8093 | 8010 | 8054 |
| **Complete single copy** | 8061 | 7967 | 8021 |
| **Complete duplicated** | 32 | 43 | 33 |
| **Fragmented** | 56 | 66 | 67 |
| **Missing** | 189 | 262 | 216 |

**Supplementary Table 2:** CEGMA analysis of the final genomes

| CEGMA (core eukaryotic genes) | Black swan | Mute swan | Chicken (bGalGal1) |
|---|---|---|---|
| **Complete** | 219 | 221 | 224 |
| **Complete and partial** | 226 | 228 | 229 |
| **Missing** | 22 | 20 | 19 |
| **Average ortho per gene** | 1.27 | 1.24 | 1.21 |
| **% of detected core genes that have more than one ortholog** | 20.55 | 19.46 | 16.52 |

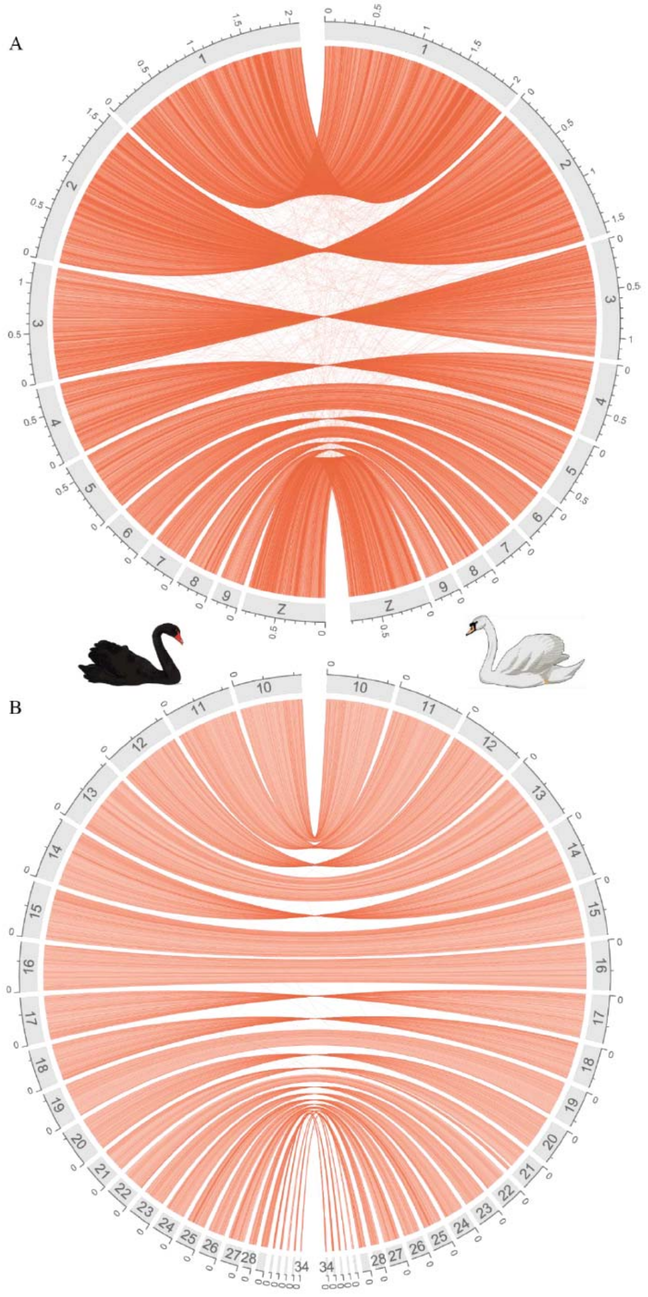

**Supplementary Figure 3: The largest 34 chromosomes aligned between the black swan (left) and the mute swan (right). A)** Macrochromosomes **B)** Microchromosomes. There was a 1-to-1 alignment identity of 98.35% between the two species. Figures were created using MUMMER and Circlize.

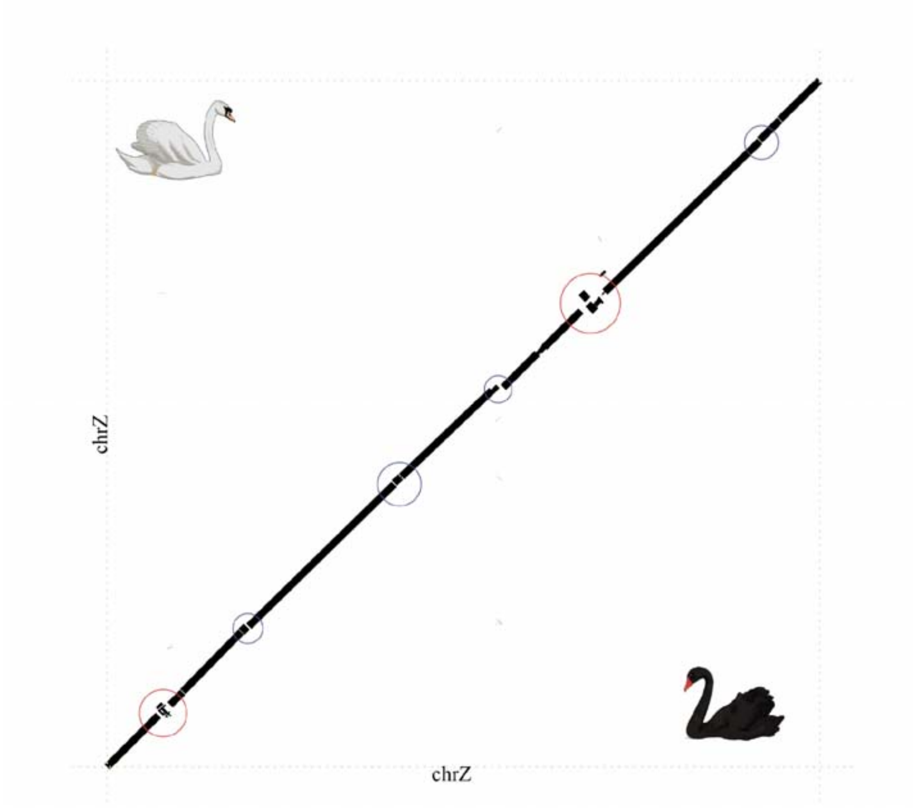


**Supplementary Figure 4: Collinearity analysis of the Z chromosome between the mute swan (left) and black swan (right).** Genomic inversions are circled in red and indels are circled in blue.

**Supplementary Table 3: GO biological process annotation of inverted genes in the Mallard duck (relative to the black swan)**

| GO biological process | Gene count |
|---|---|
| Cellular process (GO:0009987) | 319 |
| Reproductive process (GO:0022414) | 1 |
| Localisation (GO0051179) | 58 |
| Interspecies interaction between organisms (GO:0044419) | 4 |
| Reproduction (GO:0000003) | 1 |
| Biological regulation (GO:0065007) | 235 |
| Response to stimulus (GO:0050896) | 75 |
| Behaviour (GO:0007610) | 6 |
| Signalling (GO:0023052) | 69 |
| Developmental process (GO: 0032502) | 35 |
| Multicellular organismal process (GO:0032501) | 89 |
| Locomotion (GO:0040011) | 7 |
| Biological adhesion (GO0022610) | 6 |
| Metabolic process (GO:0008152) | 235 |
| Immune system process (GO:0002376) | 53 |

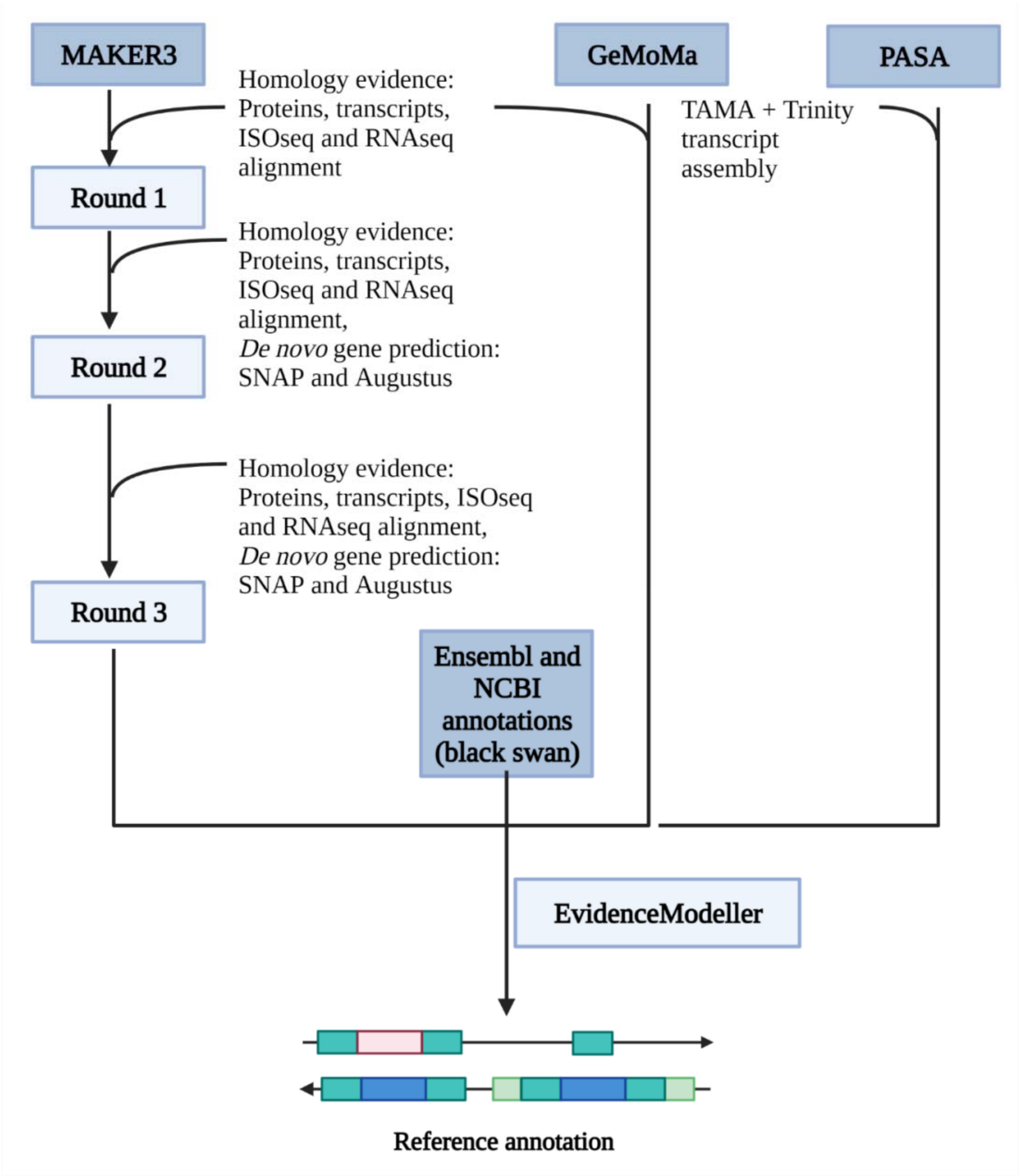


**Supplementary Figure 5: Schematic overview of the gene annotation pipeline.**

**Supplementary Table 4: Immune gene families are contractive in black swans compared to mute swans.** The number of genes in each immune gene sub-family identified is given in each column for the corresponding species

| PANTHER protein sub-family | Family name | *Anas platyrhynchos* | *Gallus gallus* | *Cygnus atratus* | *Cygnus olor* |
|---|---|---|---|---|---|
| PTHR10621:SF29 | UV EXCISION REPAIR PROTEIN RAD23 HOMOLOG A | 1 | 1 | 0 | 1 |
| PTHR11073:SF16 | CALRETICULIN | 1 | 1 | 0 | 1 |
| PTHR11829:SF68 | FORKHEAD BOX PROTEIN C1 | 1 | 0 | 0 | 1 |
| PTHR11955:SF56 | RETINOL BINDING PROTEIN 1 | 1 | 1 | 0 | 1 |
| PTHR12306:SF10 | CELL DEATH ACTIVATOR CIDE-B | 1 | 0 | 0 | 1 |
| PTHR12610:SF23 | SINGLE-STRANDED DNA BINDING PROTEIN 2 | 1 | 1 | 0 | 2 |
| PTHR13843:SF11 | MIRCOTUBULE ASSOCIATED PROTEIN 1S | 1 | 1 | 0 | 1 |
| PTHR15492:SF1 | CYCLIN-D1-BINDING PROTEIN 1 | 1 | 1 | 1 | 2 |
| PTHR20875:SF2 | EF-HAND CALCIUM-BINDING DOMAIN-CONTAINING PROTEIN 6 | 1 | 2 | 0 | 4 |
| PTHR21540:SF3 | E3 UBIQUITIN-PROTEIN LIGASE ZSWIM2 | 1 | 0 | 0 | 2 |
| PTHR22923:SF69 | COMPLEMENT C1Q-LIKE PROTEIN 2 | 2 | 1 | 0 | 1 |

| | | | | | |
|---|---|---|---|---|---|
| PTHR22940:SF4 | PROTEIN TIMELESS HOMOLOG | 1 | 1 | 0 | 1 |
| PTHR23238:SF5 | RNA-BINDING PROTEIN FUS | 0 | 1 | 0 | 1 |
| PTHR23347:SF5 | USHER SYNDROME TYPE-1C PROTEIN-BINDING PROTEIN 1 | 1 | 2 | 0 | 1 |
| PTHR24026:SF36 | CADHERIN EGF LAG SEVEN-PASS G-TYPE RECEPTOR 1 | 1 | 0 | 0 | 1 |
| PTHR24346:SF56 | SERINE/THREONINE-PROTEIN KINASE MARK2 | 2 | 1 | 0 | 1 |
| PTHR31770:SF2 | CHEMOKINE-LIKE PROTEIN TAFA-1 | 1 | 1 | 0 | 1 |
| PTHR44329:SF6 | RECEPTOR-INTERACTING SERINE/THREONINE-PROTEIN KINASE 1 | 1 | 1 | 0 | 1 |
| PTHR45636:SF26 | PAIRED BOX PROTEIN PAX-7 | 1 | 1 | 0 | 1 |
| PTHR45664:SF11 | HOMEOBOX PROTEIN HOX-B3 | 1 | 1 | 0 | 1 |
| PTHR45807:SF6 | NON-RECEPTOR TYROSINE-PROTEIN KINASE TYK2 | 1 | 1 | 0 | 1 |
| PTHR45894:SF1 | RNA-BINDING PROTEIN 8A | 1 | 1 | 0 | 1 |
| PTHR46318:SF4 | NUCLEOLAR TRANSCRIPTION FACTOR 1 | 1 | 1 | 0 | 1 |
| PTHR10155:SF6 | SRC-LIKE-ADAPTER 2 | 2 | 1 | 1 | 2 |

| | | | | | |
|---|---|---|---|---|---|
| PTHR10460:SF2 | ABL INTERACTOR 1 | 2 | 1 | 1 | 2 |
| PTHR11412:SF86 | COMPLEMENT C4-A-RELATED | 2 | 2 | 1 | 2 |
| PTHR11753:SF45 | AP-1 COMPLEX SUBUNIT SIGMA-2 | 2 | 1 | 1 | 2 |
| PTHR13703:SF42 | MOTHERS AGAINST DECAPENTAPLEGIC HOMOLOG 2 | 3 | 4 | 1 | 4 |
| PTHR14879:SF2 | E3 UBIQUITIN-PROTEIN LIGASE RIFIFYLIN | 2 | 1 | 1 | 2 |
| PTHR16830:SF13 | FYN-BINDING PROTEIN 1 | 3 | 2 | 1 | 3 |
| PTHR22741:SF11 | SICKLE TAIL PROTEIN HOMOLOG | 1 | 1 | 1 | 2 |
| PTHR32546:SF11 | G-PROTEIN COUPLED RECEPTOR 158-RELATED | 2 | 1 | 1 | 2 |
| PTHR43607:SF1 | V-TYPE PROTON ATPASE CATALYTIC SUBUNIT A | 2 | 2 | 1 | 2 |
| PTHR48021:SF59 | SOLUTE CARRIER FAMILY 2, FACILITATED GLUCOSE TRANSPORTER MEMBER 6 | 2 | 1 | 1 | 2 |
| PTHR22826:SF104 | TRIPLE FUNCTIONAL DOMAIN PROTEIN | 4 | 4 | 2 | 3 |
| PTHR23036:SF29 | INTERLEUKIN-11 RECEPTOR SUBUNIT ALPHA | 1 | 1 | 1 | 2 |

| | | | | | |
|---|---|---|---|---|---|
| PTHR11267:SF104 | T-BOX TRANSCRIPTION FACTOR TBX1 | 4 | 2 | 3 | 4 |
| PTHR24046:SF4 | SIGNAL PEPTIDE, CUB AND EGF-LIKE DOMAIN-CONTAINING PROTEIN 1 | 4 | 3 | 3 | 4 |
| PTHR11709:SF226 | CERULOPLASMIN | 8 | 8 | 4 | 5 |

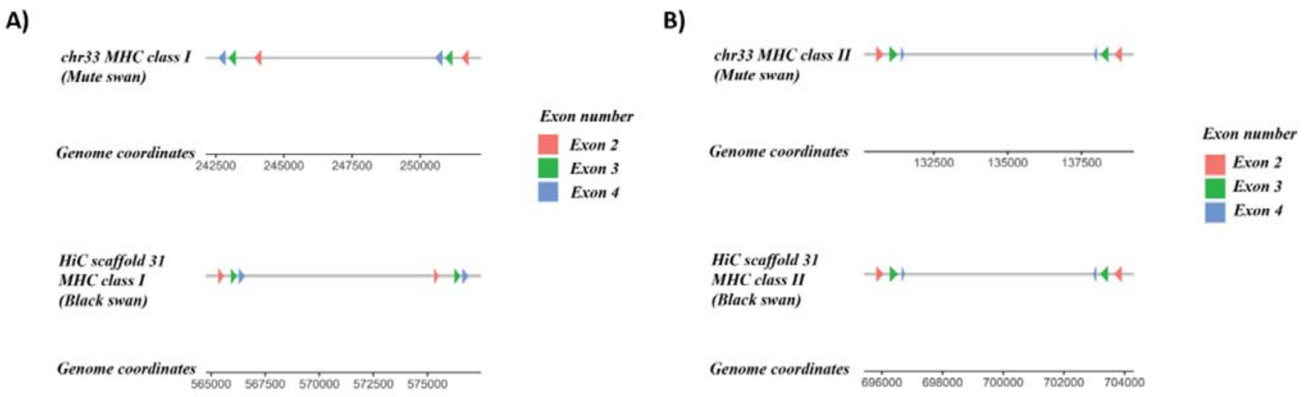


**Supplementary Figure 6: MHC Class I (A) and MHC Class II (B) aligned regions of order-level consensus exons of the mute and black swan.**

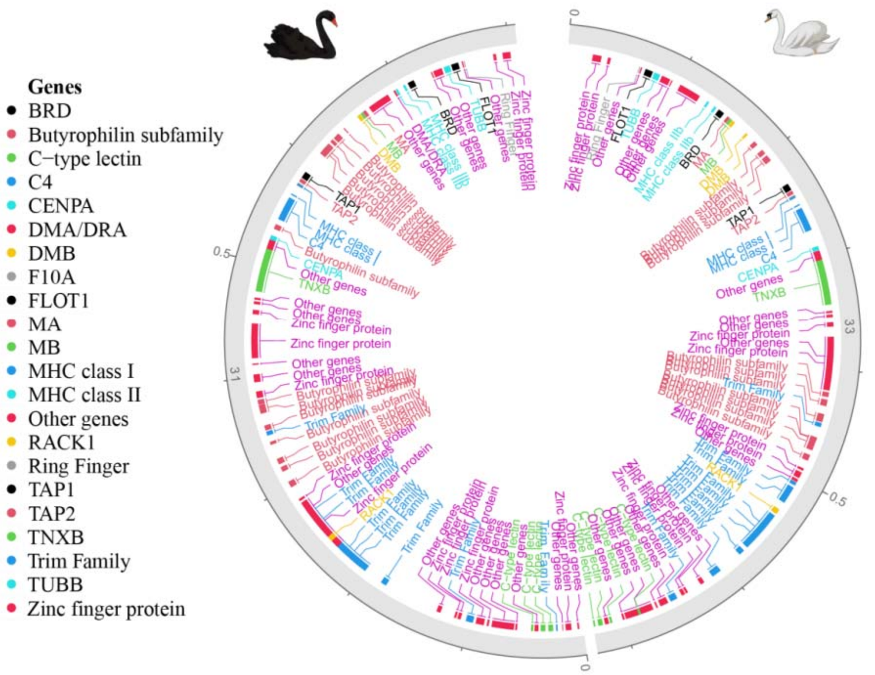


**Supplementary Figure 7: Relative locations of MHC complex and associated genes in black and mute swan chromosome 33.** *BRD* – Bromodomain 2 containing gene, *C4* – complement factor 4 – like gene, *CENPA* – Centromere protein A, *DMA* – Major histocompatibility complex II DM alpha, *DRA* - Major histocompatibility complex II DR alpha, FLT1- Flotillin, *MA* - Major histocompatibility complex II M alpha, *MB* - Major histocompatibility complex II M beta, *RACK1* – Receptor for activated C kinase 1, *TAP1* - Transporter 1, ATP Binding Cassette Subfamily B Member, *TAP2* - Transporter 2, ATP Binding Cassette Subfamily B Member, *TNXB* – Tenascin XB, *TUBB* – Tubulin beta class I, Trim family – Tripartite motif-containing family of proteins, Ring finger – Ringer finger domain-containing proteins, Zinc finger protein – Zinc finger domain-containing proteins

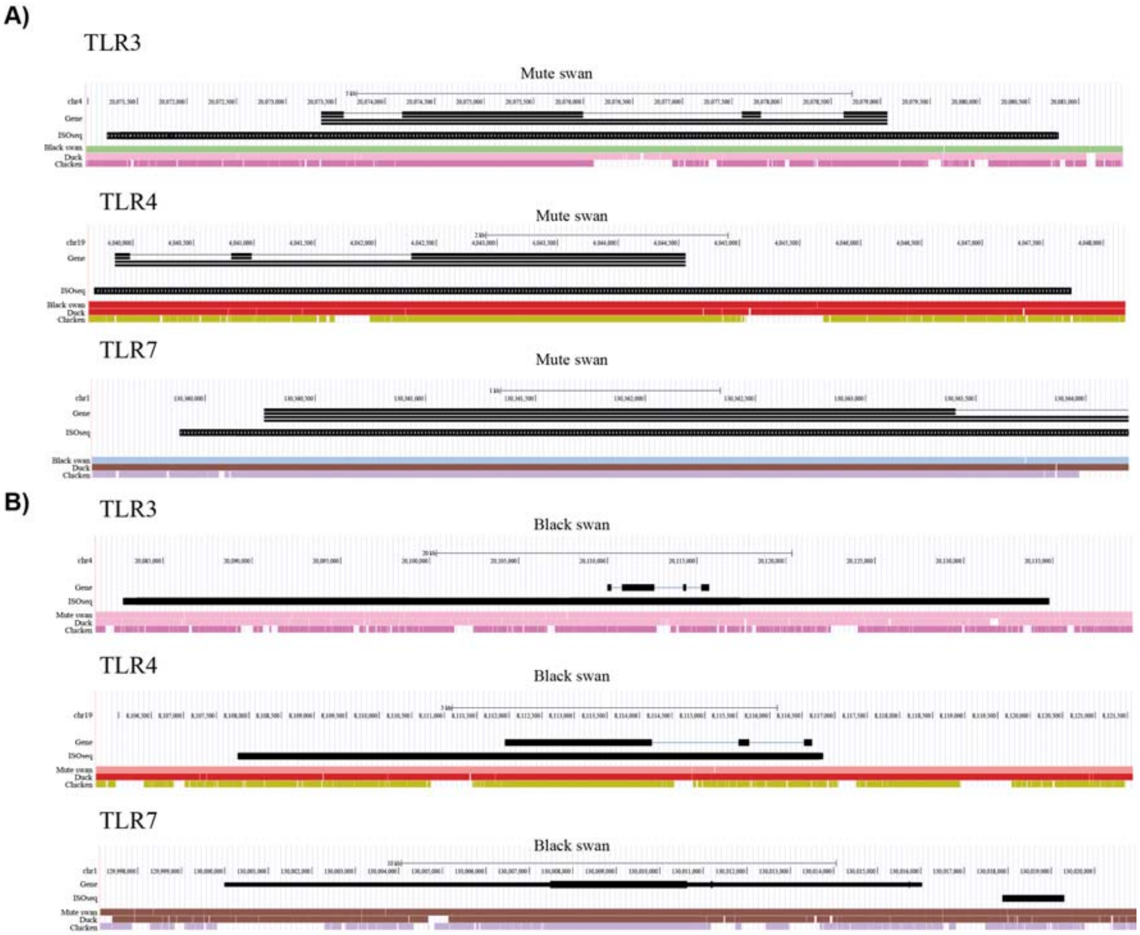


**Supplementary Figure 8: TLR7 expression can be detected in ISO Seq analysis of the mute swan (A) but not the black swan (B).** Chr: chromosome; Gene: the annotated gene model; ISOseq: ISOseq read alignment.

**Supplementary Table 5: TLR7 expression could not be detected in black swan spleen, liver and kidney by qRT-PCR. *ND: Not detected**

| Sample | 18S Ct | TLR7 Ct |
|---|---|---|
| Black swan 1 Kidney | 16.90791 | ND* |
| Black swan 1 Spleen | 29.62261 | ND |
| Black swan 1 Liver | 14.65389 | ND |
| Black swan 2 Kidney | 15.75131 | ND |
| Black swan 2 Liver | 20.61559 | ND |
| Black swan genomic DNA | 25.61549 | 29.36461 |
| Black swan RNA | ND | ND |

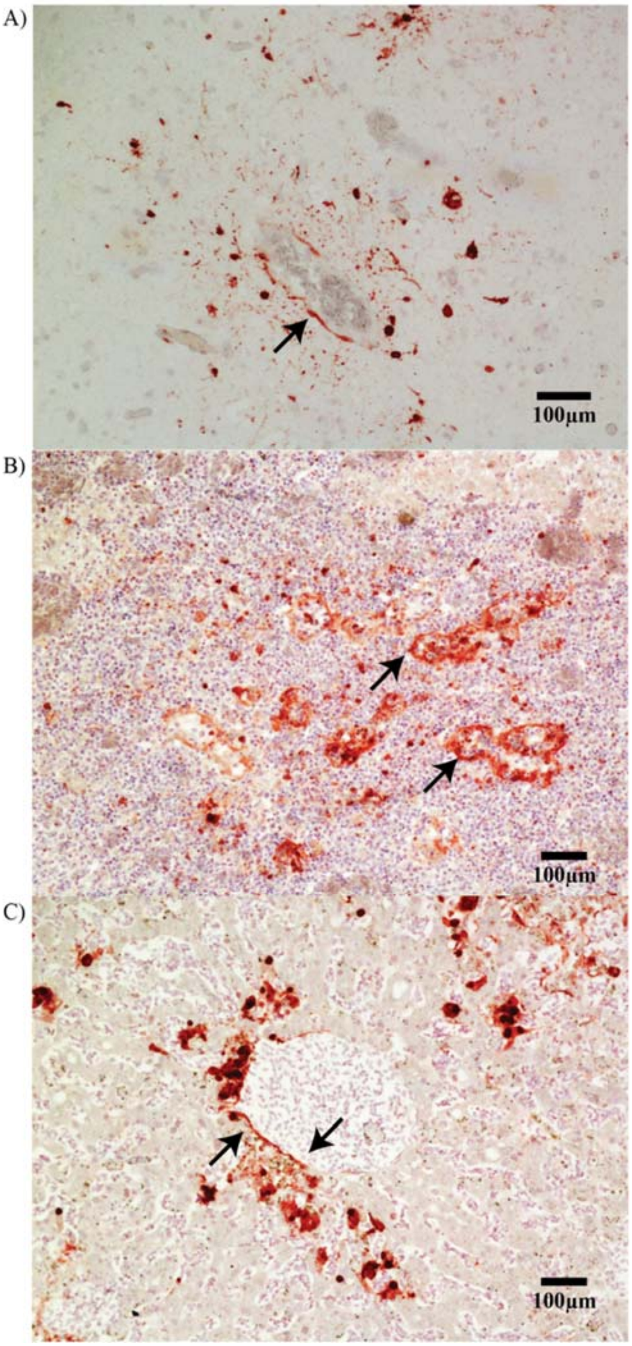


**Supplementary Figure 9: IAV NP antigen distribution in tissues from a black swan naturally infected with A/Black swan/Akita/2/2016 (H5N6) in 2016 in Akita prefecture, Japan.** Immunohistochemical detection of H5N6 nuclear protein in paraffin-embedded sections of black swan **A)** brain, **B)** spleen and **C)** liver (bar = 100μm). Arrows point to the IAV-positive endothelial cell lining. The presence of NP viral antigen is indicated by red/brown stain. No NP viral antigen was detected in the control stained serial sections.

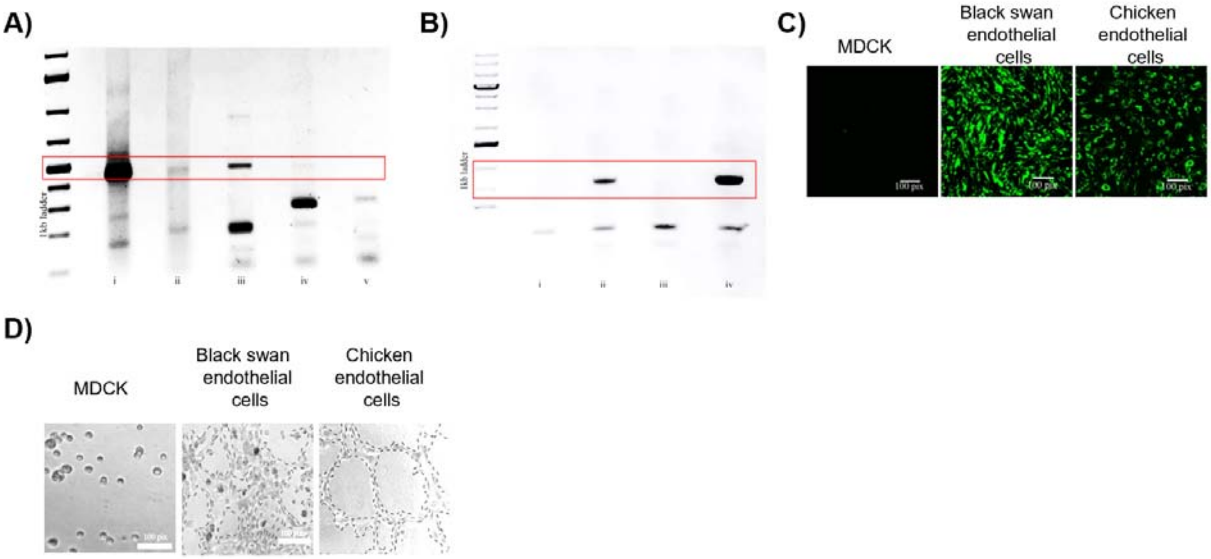


**Supplementary Figure 10: The successful culture of primary black swan endothelial cells confirmed by qRT-PCR, immunofluorescence and tube formation. A)** von Willebrand factor (vWF) expression in endothelial cells. i) duck aortic endothelial cells ii) duck bone marrow cells in culture for one day iii) black swan endothelial cells iv) black swan bone marrow cells in culture for one day v) black swan peripheral blood mononuclear cells. PCR amplicon size = ~450bp. **B)** CD45 expression in endothelial cells i) duck aortic endothelial cells ii) duck bone marrow cells in culture for one day iii) black swan endothelial cells iv) black swan bone marrow cells in culture for one day iv) black swan peripheral blood mononuclear cells. PCR amplicon size = ~310bp. **C)** Immunofluorescence of cells following a four hour incubation with FITC-labelled acetylated low-density lipoprotein. **D)** Cellular tube formation on solidified Cultrex® Basement Membrane Extract Type 2.

**Supplementary Table 6: Differentially expressed genes in infected black swan endothelial cells**

| Gene Name | log2FoldChange | p-value | p-adj |
|---|---|---|---|
| LOC118246182 | 4.580175048 | 2.79E-91 | 3.68E-87 |
| ENSACYG00000016114 | 6.195069474 | 7.14E-76 | 4.71E-72 |
| RSAD2 | 3.695484293 | 2.36E-74 | 1.04E-70 |
| LOC118254221 | 3.58506455 | 3.62E-60 | 1.19E-56 |
| USP18 | 3.502175442 | 2.68E-59 | 7.07E-56 |
| HELZ2 | 3.645057686 | 2.48E-46 | 5.46E-43 |
| RGS2 | -2.383023552 | 3.86E-44 | 7.28E-41 |
| ENSACYG00000016981 | 3.41066558 | 8.38E-42 | 1.38E-38 |
| LOC118247687 | 3.652176405 | 1.30E-34 | 1.91E-31 |
| SPRY1 | -1.782290425 | 8.67E-25 | 1.14E-21 |
| DHX58 | 2.148332185 | 2.75E-24 | 3.30E-21 |
| CCL20 | 3.574765402 | 3.36E-23 | 3.43E-20 |
| TNFSF8 | 4.466255163 | 3.38E-23 | 3.43E-20 |
| LOC118247453 | 2.445280893 | 2.39E-22 | 2.25E-19 |
| CMPK2 | 1.816769385 | 6.59E-21 | 5.61E-18 |
| CCN2 | -1.484283988 | 6.80E-21 | 5.61E-18 |
| EIF2AK2 | 1.642487532 | 1.29E-20 | 1.00E-17 |
| LOC118253574 | 5.72343541 | 7.52E-20 | 5.51E-17 |
| LOC118257698 | 7.095131916 | 2.51E-19 | 1.74E-16 |
| EDN1 | -1.588052126 | 8.88E-19 | 5.86E-16 |
| JUN | -1.379333866 | 4.92E-18 | 3.09E-15 |
| LOC118243513 | 3.561929001 | 1.83E-17 | 1.10E-14 |
| ENSACYG00000007562 | 4.962435236 | 1.14E-16 | 6.55E-14 |
| ENSACYG00000015363 | 2.617776861 | 1.63E-16 | 8.98E-14 |
| GRN | -1.185595554 | 8.03E-16 | 4.24E-13 |
| DDIT4 | -1.795782961 | 1.27E-15 | 6.44E-13 |
| ENSACYG00000007800 | -2.2221888 | 2.31E-15 | 1.13E-12 |
| HSPA2 | -1.072147928 | 5.95E-15 | 2.80E-12 |
| RRAD | -1.294474384 | 7.50E-15 | 3.33E-12 |
| ENSACYG00000014576 | 6.252514906 | 7.57E-15 | 3.33E-12 |
| SRSF5 | -1.582237015 | 9.76E-15 | 4.16E-12 |
| FSIP1 | 2.278172621 | 1.47E-14 | 6.04E-12 |
| ST6GALNAC1 | 2.164273093 | 7.22E-14 | 2.89E-11 |
| CITED4 | -1.249362893 | 1.01E-13 | 3.90E-11 |
| ENSACYG00000007716 | 4.290252805 | 1.03E-13 | 3.90E-11 |
| TNFAIP2 | 1.21345486 | 1.25E-13 | 4.58E-11 |
| ENSACYG00000003191 | 3.256022692 | 1.33E-13 | 4.73E-11 |
| SHQ1 | -1.457936056 | 1.63E-13 | 5.66E-11 |
| HSP90AA1 | -1.015067647 | 8.86E-13 | 2.99E-10 |
| ENC1 | -1.290660525 | 9.05E-13 | 2.99E-10 |
| CREB5 | 1.14513956 | 2.59E-12 | 8.33E-10 |
| FAAH2 | -1.184134502 | 2.97E-12 | 9.34E-10 |
| MAP1LC3C | -1.432055932 | 4.32E-12 | 1.33E-09 |
| DUSP4 | -1.335636982 | 4.88E-12 | 1.47E-09 |
| LOC118244434 | 1.080351876 | 5.43E-12 | 1.59E-09 |

| | | | |
|---|---|---|---|
| ATP2B1 | -1.04326864 | 7.84E-12 | 2.25E-09 |
| LOC118259586 | 3.69922664 | 8.63E-12 | 2.42E-09 |
| ING1 | -1.438405111 | 9.52E-12 | 2.62E-09 |
| IL12RB2 | 2.59592289 | 1.14E-11 | 3.08E-09 |
| LOC118250194 | 1.302368927 | 1.24E-11 | 3.28E-09 |
| FASN | -1.011409185 | 1.61E-11 | 4.17E-09 |
| TCIM | -1.989711736 | 1.75E-11 | 4.43E-09 |
| KLF2 | -1.249259299 | 2.61E-11 | 6.51E-09 |
| LOC118260896 | -1.316698577 | 3.31E-11 | 8.09E-09 |
| DDX60 | 1.686597891 | 3.49E-11 | 8.37E-09 |
| BTG1 | -1.077831805 | 4.32E-11 | 1.02E-08 |
| DNAH17 | 3.092624528 | 4.80E-11 | 1.11E-08 |
| NEU3 | -1.946778268 | 5.09E-11 | 1.16E-08 |
| COL9A3 | 3.08487153 | 5.62E-11 | 1.26E-08 |
| ENSACYG00000008104 | -1.43977034 | 5.73E-11 | 1.26E-08 |
| NR0B1 | -1.663653787 | 7.49E-11 | 1.62E-08 |
| MARS2 | -1.290585705 | 8.41E-11 | 1.79E-08 |
| SLC16A10 | 2.387959825 | 8.79E-11 | 1.84E-08 |
| CUNH3orf52 | -1.408583777 | 1.42E-10 | 2.93E-08 |
| VCAN | -0.936519302 | 1.45E-10 | 2.95E-08 |
| PTX3 | -0.853272376 | 1.59E-10 | 3.17E-08 |
| ID2 | -1.454905827 | 1.72E-10 | 3.39E-08 |
| PLEKHA6 | 1.701525256 | 1.77E-10 | 3.44E-08 |
| RIOX1 | -1.090113776 | 1.95E-10 | 3.74E-08 |
| METTL18 | -1.119667826 | 2.16E-10 | 4.07E-08 |
| SPRY2 | -1.313198245 | 2.71E-10 | 5.04E-08 |
| RBM20 | 3.491956132 | 3.56E-10 | 6.52E-08 |
| LINGO1 | 0.919570232 | 5.40E-10 | 9.77E-08 |
| ENSACYG00000017295 | 2.632321833 | 6.48E-10 | 1.16E-07 |
| MITD1 | 1.025077977 | 7.98E-10 | 1.41E-07 |
| FANCF | -1.171218516 | 8.79E-10 | 1.53E-07 |
| SLC35E4 | -0.954045479 | 9.59E-10 | 1.64E-07 |
| CCKAR | 4.071263531 | 1.15E-09 | 1.95E-07 |
| FBXO48 | -1.282634229 | 1.21E-09 | 2.00E-07 |
| GTF2B | -0.980863724 | 1.21E-09 | 2.00E-07 |
| ENSACYG00000005816 | 4.908638144 | 1.33E-09 | 2.17E-07 |
| LOC118258706 | 1.551283193 | 1.37E-09 | 2.20E-07 |
| IRF7 | 1.745901208 | 1.45E-09 | 2.22E-07 |
| SLC2A9 | 7.797230023 | 1.44E-09 | 2.22E-07 |
| IL6 | 1.844550235 | 1.41E-09 | 2.22E-07 |
| ENSACYG00000008294 | 4.275266889 | 1.42E-09 | 2.22E-07 |
| LOC118254002 | 1.688382627 | 1.54E-09 | 2.33E-07 |
| TBXAS1 | 2.818796526 | 1.66E-09 | 2.48E-07 |
| LOC118254070 | -0.852998203 | 1.78E-09 | 2.64E-07 |
| ZFAND2B | -1.230402071 | 2.26E-09 | 3.32E-07 |
| TRUB1 | -1.119258098 | 2.66E-09 | 3.81E-07 |
| NELFA | -0.924509813 | 2.65E-09 | 3.81E-07 |
| DUS3L | -1.24552966 | 2.81E-09 | 3.99E-07 |

| | | | |
|---|---|---|---|
| SASS6 | -0.98428245 | 3.30E-09 | 4.63E-07 |
| ENSACYG00000001324 | -1.237713592 | 3.40E-09 | 4.73E-07 |
| R3HDM1 | 0.880811897 | 4.10E-09 | 5.64E-07 |
| RRBP1 | -0.864458522 | 4.15E-09 | 5.65E-07 |
| ENSACYG00000012255 | 3.554320128 | 4.62E-09 | 6.22E-07 |
| CCN1 | -1.086904006 | 5.34E-09 | 7.05E-07 |
| SLC38A2 | 0.975729567 | 5.29E-09 | 7.05E-07 |
| DUSP1 | -1.083200374 | 6.57E-09 | 8.59E-07 |
| LOC118248062 | 3.03903724 | 8.56E-09 | 1.11E-06 |
| MYH9 | -1.055902423 | 8.98E-09 | 1.15E-06 |
| G0S2 | -1.109953144 | 9.35E-09 | 1.19E-06 |
| VCAM1 | 1.790847269 | 1.05E-08 | 1.32E-06 |
| SLC26A4 | 6.376011231 | 1.12E-08 | 1.40E-06 |
| LOC118251676 | 3.447902462 | 1.13E-08 | 1.40E-06 |
| TPM1 | 2.402101785 | 1.15E-08 | 1.41E-06 |
| TPR | -0.950697258 | 1.21E-08 | 1.46E-06 |
| HAUS8 | -0.864383898 | 1.21E-08 | 1.46E-06 |
| DNPEP | -1.066736825 | 1.26E-08 | 1.49E-06 |
| MED18 | -1.197063088 | 1.26E-08 | 1.49E-06 |
| HEXB | -0.813078015 | 1.38E-08 | 1.60E-06 |
| LOC118251878 | 1.456187298 | 1.38E-08 | 1.60E-06 |
| ENSACYG00000008278 | 2.938877762 | 1.54E-08 | 1.77E-06 |
| NEMP2 | 1.061732056 | 1.56E-08 | 1.77E-06 |
| CUNH2orf49 | -0.907704957 | 1.68E-08 | 1.90E-06 |
| SYNE1 | -0.980365761 | 1.80E-08 | 2.01E-06 |
| ESS2 | -1.111893384 | 1.96E-08 | 2.18E-06 |
| MED13 | 1.090369967 | 2.02E-08 | 2.22E-06 |
| RASL11B | -0.899780172 | 2.08E-08 | 2.25E-06 |
| ENSACYG00000012368 | 3.125181819 | 2.07E-08 | 2.25E-06 |
| MCL1 | -0.943125585 | 2.13E-08 | 2.29E-06 |
| SNAPC3 | -1.03563998 | 2.20E-08 | 2.34E-06 |
| MYC | -0.892925534 | 2.39E-08 | 2.52E-06 |
| ARL4A | -0.961099269 | 2.57E-08 | 2.70E-06 |
| NECTIN3 | -0.792077121 | 2.61E-08 | 2.72E-06 |
| SLC5A1 | 3.993031358 | 2.67E-08 | 2.75E-06 |
| MMAA | -0.913567628 | 3.06E-08 | 3.13E-06 |
| HIPK2 | 0.888853789 | 3.12E-08 | 3.17E-06 |
| CCNL2 | -1.061472207 | 3.90E-08 | 3.93E-06 |
| LOC118244242 | -0.755677579 | 4.25E-08 | 4.25E-06 |
| CLDN1 | -0.825540386 | 4.78E-08 | 4.74E-06 |
| ZBTB25 | -0.974410821 | 5.39E-08 | 5.31E-06 |
| WEE1 | -1.164838979 | 5.43E-08 | 5.31E-06 |
| PPP2R2B | 0.838024139 | 5.72E-08 | 5.56E-06 |
| TENT5A | -0.875170139 | 5.89E-08 | 5.67E-06 |
| SMTN | -0.836298296 | 6.28E-08 | 6.01E-06 |
| ARL10 | -1.156946133 | 6.50E-08 | 6.18E-06 |
| ENSACYG00000009359 | 2.011367472 | 6.86E-08 | 6.45E-06 |
| TXLNB | 3.414940487 | 6.89E-08 | 6.45E-06 |

| | | | |
|---|---|---|---|
| ENSACYG00000000041 | 2.6435625 | 7.27E-08 | 6.76E-06 |
| EXOSC3 | -1.235916813 | 7.77E-08 | 7.18E-06 |
| TRA2B | -0.908413209 | 8.74E-08 | 8.01E-06 |
| ATF4 | -0.924615219 | 8.86E-08 | 8.07E-06 |
| IL1RL1 | 1.614160155 | 9.01E-08 | 8.15E-06 |
| ENSACYG00000013288 | -1.391271802 | 9.51E-08 | 8.54E-06 |
| LIPT1 | -0.984761858 | 1.05E-07 | 9.31E-06 |
| LOC118258158 | 1.070987532 | 1.05E-07 | 9.31E-06 |
| LOC118249052 | -0.805846326 | 1.07E-07 | 9.38E-06 |
| BMPR1B | 4.494371119 | 1.11E-07 | 9.70E-06 |
| LOC118248085 | -1.110420319 | 1.13E-07 | 9.83E-06 |
| CCM2 | 0.755094907 | 1.17E-07 | 9.93E-06 |
| DND1 | -3.273289986 | 1.17E-07 | 9.93E-06 |
| RND2 | -1.089125135 | 1.15E-07 | 9.93E-06 |
| PAX1 | 1.134763224 | 1.17E-07 | 9.93E-06 |
| CCNB2 | -0.927169767 | 1.18E-07 | 9.95E-06 |
| PARS2 | -0.915264627 | 1.23E-07 | 1.03E-05 |
| ANAPC7 | -0.921617429 | 1.28E-07 | 1.06E-05 |
| EXOSC9 | -0.932195397 | 1.33E-07 | 1.09E-05 |
| LOC118253381 | -1.654932298 | 1.33E-07 | 1.09E-05 |
| WSB1 | -0.765716622 | 1.36E-07 | 1.11E-05 |
| WDR89 | -0.887090585 | 1.40E-07 | 1.13E-05 |
| ASPHD2 | 1.059771392 | 1.43E-07 | 1.15E-05 |
| ENSACYG00000008802 | 7.083971698 | 1.52E-07 | 1.22E-05 |
| NLRC5 | 1.769350073 | 1.59E-07 | 1.26E-05 |
| POC1A | -0.997006512 | 1.60E-07 | 1.26E-05 |
| DYNC1H1 | -0.854031071 | 1.62E-07 | 1.27E-05 |
| POP1 | -0.960294536 | 1.66E-07 | 1.29E-05 |
| FLNB | -0.854381119 | 1.69E-07 | 1.31E-05 |
| PLEC | -1.06066991 | 1.87E-07 | 1.44E-05 |
| ENSACYG00000002531 | 4.452858297 | 1.90E-07 | 1.46E-05 |
| LOC118245596 | 7.092582392 | 1.98E-07 | 1.51E-05 |
| PABPC1 | 0.74800618 | 2.09E-07 | 1.58E-05 |
| PUS3 | -0.839919272 | 2.13E-07 | 1.61E-05 |
| TLN1 | -1.227764504 | 2.25E-07 | 1.69E-05 |
| FERMT1 | -0.793829415 | 2.39E-07 | 1.78E-05 |
| ENSACYG00000017004 | 7.011333312 | 2.48E-07 | 1.84E-05 |
| GCH1 | 1.002391677 | 2.52E-07 | 1.86E-05 |
| TGDS | -0.871534306 | 2.78E-07 | 2.04E-05 |
| NOC3L | -0.809976362 | 2.85E-07 | 2.08E-05 |
| NCOA7 | 0.904600837 | 2.95E-07 | 2.14E-05 |
| LOC118247374 | -1.049362264 | 3.33E-07 | 2.40E-05 |
| CDC20 | -0.800666466 | 3.42E-07 | 2.46E-05 |
| LOC118261382 | 1.247185742 | 3.46E-07 | 2.47E-05 |
| CXCR4 | -0.856779646 | 3.52E-07 | 2.50E-05 |
| OGT | -0.817270643 | 3.76E-07 | 2.63E-05 |
| ENSACYG00000009286 | 2.508441777 | 3.76E-07 | 2.63E-05 |
| ENSACYG00000012852 | 1.837357737 | 3.73E-07 | 2.63E-05 |

| | | | |
|---|---|---|---|
| ARMC6 | -0.733491202 | 3.87E-07 | 2.69E-05 |
| EPSTI1 | 1.762138054 | 3.93E-07 | 2.71E-05 |
| ENSACYG00000000249 | -1.006917773 | 3.99E-07 | 2.74E-05 |
| ALG11 | -0.845949625 | 4.16E-07 | 2.84E-05 |
| TRAF7 | 0.822751166 | 4.17E-07 | 2.84E-05 |
| PHOSPHO1 | 1.129145206 | 4.35E-07 | 2.93E-05 |
| MYCN | -1.066468717 | 4.37E-07 | 2.93E-05 |
| RNF217 | 0.911279599 | 4.36E-07 | 2.93E-05 |
| SLC9A4 | 6.92936322 | 4.41E-07 | 2.93E-05 |
| KLHL42 | -0.797332517 | 4.42E-07 | 2.93E-05 |
| PIGL | -0.796261666 | 4.54E-07 | 2.99E-05 |
| RPP25L | -0.771593225 | 4.66E-07 | 3.06E-05 |
| SRSF7 | -0.82592492 | 4.73E-07 | 3.09E-05 |
| TMEM200B | 1.073437235 | 4.96E-07 | 3.23E-05 |
| ENSACYG00000011789 | 5.116057588 | 5.00E-07 | 3.24E-05 |
| ATP2A2 | -0.727762152 | 5.11E-07 | 3.29E-05 |
| ZBTB20 | 0.912955108 | 5.26E-07 | 3.37E-05 |
| GOT2 | -0.671124793 | 5.36E-07 | 3.42E-05 |
| POLR1F | -1.069375365 | 5.70E-07 | 3.62E-05 |
| LOC118244925 | -0.952798602 | 5.96E-07 | 3.77E-05 |
| WIF1 | -0.815473635 | 6.42E-07 | 4.03E-05 |
| SETD6 | -1.047046099 | 6.60E-07 | 4.13E-05 |
| DYNAP | 4.073014401 | 6.77E-07 | 4.22E-05 |
| INPP5B | -1.162274219 | 7.00E-07 | 4.34E-05 |
| CX3CL1 | 1.939813044 | 7.17E-07 | 4.41E-05 |
| FBXO42 | 0.880471227 | 7.18E-07 | 4.41E-05 |
| CLK1 | -0.89144568 | 7.68E-07 | 4.69E-05 |
| PIM3 | -0.703341269 | 7.79E-07 | 4.74E-05 |
| SURF2 | -1.035464848 | 8.00E-07 | 4.85E-05 |
| MAP3K8 | 1.077754107 | 8.19E-07 | 4.94E-05 |
| LONRF3 | 0.731613904 | 8.77E-07 | 5.26E-05 |
| LOC118253911 | -0.845608339 | 9.37E-07 | 5.60E-05 |
| ATM | -0.82493099 | 9.52E-07 | 5.66E-05 |
| JMJD4 | 0.718561255 | 9.68E-07 | 5.73E-05 |
| LOC118248082 | -0.904416384 | 9.85E-07 | 5.81E-05 |
| LOC118254130 | 1.391697946 | 1.01E-06 | 5.90E-05 |
| PKP4 | 0.847441798 | 1.03E-06 | 6.01E-05 |
| BTG2 | -0.999699575 | 1.07E-06 | 6.25E-05 |
| MTG2 | -0.96681488 | 1.10E-06 | 6.36E-05 |
| HSF1 | 0.884756713 | 1.13E-06 | 6.52E-05 |
| SCNN1B | 0.684176826 | 1.16E-06 | 6.63E-05 |
| LOC118244481 | 2.115945447 | 1.26E-06 | 7.21E-05 |
| ENSACYG00000000109 | 1.916095771 | 1.31E-06 | 7.46E-05 |
| MYBL1 | 0.828317566 | 1.33E-06 | 7.52E-05 |
| DGKH | 0.800113915 | 1.44E-06 | 8.12E-05 |
| CWC25 | -1.21363937 | 1.45E-06 | 8.12E-05 |
| TRIM45 | -0.9787944 | 1.46E-06 | 8.15E-05 |
| ENSACYG00000004093 | 3.397086649 | 1.53E-06 | 8.53E-05 |

| | | | |
|---|---|---|---|
| LOC118253948 | 0.816152101 | 1.58E-06 | 8.74E-05 |
| LMBRD1 | -0.73086765 | 1.60E-06 | 8.81E-05 |
| DOK1 | -1.033069502 | 1.60E-06 | 8.81E-05 |
| E2F6 | -1.005766608 | 1.62E-06 | 8.86E-05 |
| CMYA5 | 3.365914847 | 1.63E-06 | 8.91E-05 |
| ENSACYG00000007266 | 2.602613758 | 1.67E-06 | 9.10E-05 |
| HERPUD1 | -0.747666229 | 1.69E-06 | 9.17E-05 |
| ADM | -0.97699209 | 1.82E-06 | 9.82E-05 |
| SLC16A12 | 3.522122914 | 1.96E-06 | 0.000104634 |
| LOC118261525 | 6.680507627 | 1.96E-06 | 0.000104634 |
| RUBCNL | -0.902166764 | 1.99E-06 | 0.000105424 |
| OGFR | 0.762863693 | 1.99E-06 | 0.000105424 |
| UTP23 | -0.887940824 | 2.23E-06 | 0.000117894 |
| KLHDC1 | -0.882203126 | 2.33E-06 | 0.000122752 |
| LOC118247601 | 6.627722545 | 2.36E-06 | 0.000123823 |
| LRRC47 | -0.937035309 | 2.37E-06 | 0.000123911 |
| DDX49 | -0.786737853 | 2.51E-06 | 0.00013021 |
| ATP5F1A | -0.754237952 | 2.63E-06 | 0.000135935 |
| LOC118251879 | 1.392034441 | 2.67E-06 | 0.000137177 |
| SDR16C5 | -0.746705121 | 2.67E-06 | 0.000137177 |
| ANKRD37 | -0.726028123 | 2.80E-06 | 0.000143104 |
| ZNF143 | -0.837150234 | 2.99E-06 | 0.000152178 |
| ZNFX1 | -0.65411607 | 3.15E-06 | 0.000159749 |
| NES | -1.205426175 | 3.18E-06 | 0.000160732 |
| ANAPC10 | -1.155238545 | 3.29E-06 | 0.000165558 |
| ATXN7 | 0.927624662 | 3.36E-06 | 0.000168883 |
| LOC118259027 | -0.928018269 | 3.40E-06 | 0.000169661 |
| FAM53A | -0.715286518 | 3.41E-06 | 0.000169661 |
| SLC48A1 | 0.675994487 | 3.73E-06 | 0.000185015 |
| CNOT10 | -0.792394127 | 3.87E-06 | 0.00019068 |
| MED27 | -0.736891508 | 3.87E-06 | 0.00019068 |
| LOC118247555 | -0.715443409 | 3.94E-06 | 0.000193502 |
| METTL21A | -0.919231573 | 4.02E-06 | 0.000196681 |
| ATG10 | 0.842633025 | 4.05E-06 | 0.000197262 |
| TKT | -0.657973246 | 4.10E-06 | 0.000198891 |
| AKAP10 | 0.80482146 | 4.14E-06 | 0.000200173 |
| ENSACYG00000011963 | -1.049804387 | 4.15E-06 | 0.000200173 |
| SLC34A2 | 1.736195424 | 4.22E-06 | 0.000202386 |
| MBTPS1 | 0.829753652 | 4.26E-06 | 0.000203531 |
| POMGNT2 | 0.673430773 | 4.28E-06 | 0.000203752 |
| LOC118252548 | -1.021354336 | 4.35E-06 | 0.000206554 |
| LOC118251921 | -1.054889039 | 4.39E-06 | 0.000207709 |
| MYH10 | -0.658155848 | 4.41E-06 | 0.000207709 |
| MTA3 | -1.01770599 | 4.46E-06 | 0.000209524 |
| PRKCD | -0.694449713 | 4.58E-06 | 0.000214408 |
| ACTR5 | -1.074077064 | 4.64E-06 | 0.000216459 |
| LOC118260289 | -0.789108077 | 4.82E-06 | 0.000223988 |
| BAMBI | -0.810476031 | 4.84E-06 | 0.000224169 |

| | | | |
|---|---|---|---|
| ERICH1 | -0.922975817 | 4.90E-06 | 0.000225971 |
| CD83 | 2.439266654 | 5.31E-06 | 0.000244139 |
| NAT10 | -0.815660689 | 5.36E-06 | 0.000245533 |
| ZBTB8B | -1.057009176 | 5.37E-06 | 0.000245533 |
| CCNL1 | -0.689897221 | 5.45E-06 | 0.000248319 |
| ENSACYG00000007129 | 3.333535918 | 6.21E-06 | 0.00028176 |
| TOP3B | -0.902007316 | 6.28E-06 | 0.000283987 |
| TPRG1L | 0.639348807 | 6.33E-06 | 0.000285114 |
| ASTE1 | -1.156726176 | 6.46E-06 | 0.000289874 |
| MOB3C | -0.91374161 | 6.63E-06 | 0.000296681 |
| AIP | -0.774463877 | 6.79E-06 | 0.000301847 |
| EDAR | 6.591255428 | 6.82E-06 | 0.000301847 |
| PPP1R3C | -0.703154457 | 6.84E-06 | 0.000301847 |
| LOC118258728 | -0.89152534 | 6.83E-06 | 0.000301847 |
| GXYLT1 | 1.239018573 | 6.95E-06 | 0.000305891 |
| MGAT3 | 0.843327761 | 7.01E-06 | 0.000307642 |
| DRAM1 | 0.862555334 | 7.05E-06 | 0.000308076 |
| CCDC141 | 3.785691695 | 7.41E-06 | 0.000323019 |
| NR4A3 | 0.921645654 | 7.48E-06 | 0.000324712 |
| UNC50 | -0.935618623 | 7.51E-06 | 0.000324905 |
| KLHL10 | 3.237019792 | 7.66E-06 | 0.000330662 |
| STK17B | -0.843655383 | 7.76E-06 | 0.000333667 |
| FBLN5 | 2.349278213 | 7.84E-06 | 0.000334977 |
| CUNH5orf51 | -2.051158434 | 7.82E-06 | 0.000334977 |
| PRPF38A | -0.750951481 | 7.96E-06 | 0.000339055 |
| BICD1 | -6.24631199 | 8.15E-06 | 0.000345801 |
| CUBN | 0.826515429 | 8.24E-06 | 0.000348843 |
| CKAP2L | -0.694732428 | 8.36E-06 | 0.000352615 |
| PIP5K1A | 0.821612502 | 8.50E-06 | 0.000356965 |
| ST6GALNAC6 | 1.050660762 | 8.52E-06 | 0.000356965 |
| ENSACYG00000001459 | 4.285340565 | 8.59E-06 | 0.00035905 |
| CACNG3 | 0.662739191 | 8.66E-06 | 0.000360688 |
| MTF1 | 0.626285751 | 8.73E-06 | 0.000362612 |
| DACT1 | -1.012491459 | 9.04E-06 | 0.000373983 |
| BBS12 | -1.026174973 | 9.09E-06 | 0.000375086 |
| ALKBH4 | -1.018771987 | 9.18E-06 | 0.000377392 |
| ABHD17B | -1.026943677 | 9.24E-06 | 0.000378772 |
| ILVBL | -0.921848386 | 9.29E-06 | 0.000379593 |
| LOC118244680 | -1.884122954 | 9.37E-06 | 0.000381822 |
| RC3H2 | 0.657731936 | 9.58E-06 | 0.000389163 |
| UTP6 | -0.662390294 | 9.62E-06 | 0.000389734 |
| TADA1 | -0.761717147 | 9.73E-06 | 0.000392811 |
| ENSACYG00000001091 | 0.610235468 | 9.95E-06 | 0.000400393 |
| LEO1 | -0.807654507 | 1.04E-05 | 0.000416162 |
| TSPAN5 | 0.798805498 | 1.04E-05 | 0.000416162 |
| ABHD17A | 0.641783554 | 1.05E-05 | 0.000417592 |
| UBAP1 | 0.690805783 | 1.08E-05 | 0.000427262 |
| TUBB4B | -0.749829853 | 1.08E-05 | 0.000427262 |

| | | | |
|---|---|---|---|
| LOC118260317 | -1.103959218 | 1.08E-05 | 0.000427262 |
| IFI30 | -0.847841895 | 1.09E-05 | 0.00043066 |
| HTR1B | 0.730787254 | 1.10E-05 | 0.00043066 |
| SIMC1 | -1.44159215 | 1.10E-05 | 0.00043066 |
| LOC118246392 | -0.647431193 | 1.11E-05 | 0.000433194 |
| HHIPL2 | 3.421815264 | 1.18E-05 | 0.000457802 |
| EIF4G2 | 0.759010403 | 1.21E-05 | 0.000468617 |
| PSTK | -0.826830832 | 1.23E-05 | 0.000477175 |
| UBC | -0.714307168 | 1.25E-05 | 0.000482114 |
| PDIA3 | -0.599629461 | 1.28E-05 | 0.000492251 |
| RGS3 | -0.880537112 | 1.28E-05 | 0.000492251 |
| CCT6A | -0.627408422 | 1.29E-05 | 0.000495474 |
| SPOUT1 | -0.778919904 | 1.34E-05 | 0.000511149 |
| HAUS2 | -1.099998664 | 1.36E-05 | 0.00051748 |
| SAXO1 | 0.742552158 | 1.46E-05 | 0.000555445 |
| ENSACYG00000001051 | 0.705497004 | 1.48E-05 | 0.000559395 |
| CASTOR1 | -0.578284777 | 1.48E-05 | 0.000559395 |
| LOC118243550 | 3.929151293 | 1.52E-05 | 0.000573201 |
| HMGCL | -0.642805757 | 1.54E-05 | 0.000578727 |
| ENSACYG00000001520 | 4.199139957 | 1.58E-05 | 0.000590053 |
| FNBP1L | -0.834132974 | 1.59E-05 | 0.000590053 |
| LOC118256259 | 0.635797664 | 1.58E-05 | 0.000590053 |
| LATS2 | 0.896620032 | 1.60E-05 | 0.000594264 |
| LOC118260046 | 0.629400592 | 1.66E-05 | 0.000612531 |
| EDC3 | -0.656028261 | 1.66E-05 | 0.000612531 |
| RHOQ | 0.578948213 | 1.68E-05 | 0.000618754 |
| WDR74 | -0.747416796 | 1.69E-05 | 0.000618785 |
| LOC118260593 | -1.017878939 | 1.73E-05 | 0.000633905 |
| ARL15 | 0.697447958 | 1.79E-05 | 0.000650581 |
| CITED2 | -0.957856486 | 1.79E-05 | 0.000650581 |
| LOC118243364 | -0.77983434 | 1.84E-05 | 0.000666724 |
| SQLE | -0.661594709 | 1.85E-05 | 0.000667579 |
| TLCD3A | 0.642613397 | 1.86E-05 | 0.000669734 |
| LOC118260291 | -1.21225064 | 1.90E-05 | 0.000684638 |
| LPAR2 | -0.79596046 | 1.91E-05 | 0.000685283 |
| MFAP5 | 0.886984896 | 1.92E-05 | 0.000687841 |
| TULP1 | 5.2464326 | 2.01E-05 | 0.000718927 |
| TRIM8 | 0.684981012 | 2.06E-05 | 0.000733171 |
| TCFL5 | 2.813858608 | 2.20E-05 | 0.000779563 |
| SMARCC2 | -1.4525534 | 2.22E-05 | 0.000784569 |
| EXOSC8 | -0.814092061 | 2.23E-05 | 0.000787331 |
| ENSACYG00000003770 | 6.24002163 | 2.24E-05 | 0.000789133 |
| ADORA2B | 1.133670946 | 2.29E-05 | 0.000804563 |
| CTSL | -0.615420546 | 2.35E-05 | 0.000819868 |
| ENSACYG00000015548 | 1.701178907 | 2.34E-05 | 0.000819868 |
| RBM41 | -0.922992465 | 2.40E-05 | 0.000834538 |
| SLC25A28 | 0.707182076 | 2.52E-05 | 0.000875643 |
| SH3BGRL2 | 0.745432379 | 2.55E-05 | 0.000880936 |

| | | | |
|---|---|---|---|
| AFAP1 | -0.572400493 | 2.55E-05 | 0.000880936 |
| GPAT4 | 0.649446633 | 2.62E-05 | 0.000904006 |
| ZBTB8OS | -0.73950974 | 2.64E-05 | 0.000905979 |
| CUNH4orf48 | 0.612001313 | 2.66E-05 | 0.000911589 |
| VAV3 | -0.834358379 | 2.69E-05 | 0.000921154 |
| LOC118251182 | -0.727897611 | 2.77E-05 | 0.000946184 |
| LOC118261260 | 0.647425261 | 2.82E-05 | 0.000960472 |
| CENPF | -0.903158775 | 2.92E-05 | 0.000990019 |
| CDC42SE2 | 0.559417497 | 2.97E-05 | 0.001002857 |
| SLC11A2 | 0.69225311 | 2.97E-05 | 0.001002857 |
| MRPS22 | -0.83780253 | 3.05E-05 | 0.001026729 |
| PABPC1L | -1.237293517 | 3.07E-05 | 0.001030584 |
| P4HB | -0.589818047 | 3.10E-05 | 0.001037307 |
| CASP8 | -0.69676408 | 3.12E-05 | 0.00104084 |
| MRPL32 | -0.787783443 | 3.12E-05 | 0.00104084 |
| ENSACYG00000016778 | 2.904743919 | 3.15E-05 | 0.001046269 |
| B4GALT2 | 0.703621922 | 3.17E-05 | 0.0010526 |
| CSNK1G2 | 0.585801781 | 3.21E-05 | 0.001061479 |
| KTN1 | -0.688062913 | 3.26E-05 | 0.001074953 |
| NLE1 | -0.664648288 | 3.29E-05 | 0.001084304 |
| ENSACYG00000004655 | 2.719901556 | 3.33E-05 | 0.00109147 |
| SUOX | -0.730016073 | 3.33E-05 | 0.00109147 |
| TRIM54 | -0.666045495 | 3.36E-05 | 0.001098928 |
| ENSACYG00000002826 | 1.444761487 | 3.47E-05 | 0.001127163 |
| AKTIP | -0.728179635 | 3.46E-05 | 0.001127163 |
| MED4 | -0.792130765 | 3.52E-05 | 0.001140435 |
| SPAG16 | -1.087482204 | 3.56E-05 | 0.0011479 |
| TBX19 | 2.188538267 | 3.55E-05 | 0.0011479 |
| LOC118243824 | 4.402458635 | 3.61E-05 | 0.001160869 |
| EXOSC4 | -0.778629228 | 3.61E-05 | 0.001160869 |
| GMCL1 | 0.717835806 | 3.71E-05 | 0.001187889 |
| SGK1 | -0.669979141 | 3.75E-05 | 0.001199365 |
| LOC118251862 | 1.189677661 | 3.80E-05 | 0.001210845 |
| FASTKD3 | -0.940703785 | 3.87E-05 | 0.001232531 |
| MRM1 | -0.730054078 | 3.96E-05 | 0.001256546 |
| ENSACYG00000010865 | 3.820112046 | 4.04E-05 | 0.00127821 |
| FBXO33 | 0.816273962 | 4.07E-05 | 0.001282187 |
| GYS2 | 6.119032688 | 4.08E-05 | 0.001282187 |
| GATAD2A | 0.64805902 | 4.07E-05 | 0.001282187 |
| TAF7L | -0.696994163 | 4.10E-05 | 0.001284964 |
| PRLHR | 3.242154591 | 4.25E-05 | 0.001328788 |
| PTPN14 | 0.69732254 | 4.26E-05 | 0.001328788 |
| YEATS4 | -0.723714833 | 4.28E-05 | 0.001332909 |
| RBM48 | -0.767928755 | 4.37E-05 | 0.001357074 |
| SIAH1 | -0.67829425 | 4.42E-05 | 0.001371309 |
| PDHA1 | -0.639898033 | 4.49E-05 | 0.001387624 |
| SLC25A29 | -0.671816274 | 4.54E-05 | 0.001401929 |
| TMCC2 | 1.096562443 | 4.62E-05 | 0.001422839 |

| | | | |
|---|---|---|---|
| LOC118261676 | -0.663635667 | 4.74E-05 | 0.001455136 |
| SERTAD2 | 0.82500163 | 4.79E-05 | 0.001465573 |
| CTSK | -0.548603151 | 4.80E-05 | 0.001465573 |
| SRPX2 | -0.593499597 | 4.85E-05 | 0.001479183 |
| PRNP | 0.601082333 | 4.95E-05 | 0.001505415 |
| ACAT1 | -0.601844436 | 5.00E-05 | 0.001518127 |
| YBX3 | 0.619708905 | 5.07E-05 | 0.001534643 |
| CH25H | 0.890155256 | 5.09E-05 | 0.001538397 |
| LOC118251550 | 0.568970601 | 5.19E-05 | 0.001565289 |
| ME1 | -1.086252686 | 5.22E-05 | 0.001567598 |
| DNASE1L3 | 3.729830923 | 5.21E-05 | 0.001567598 |
| HMOX2 | -0.567698996 | 5.26E-05 | 0.001568268 |
| TNIP2 | -0.968879158 | 5.25E-05 | 0.001568268 |
| ENSACYG00000016081 | 1.364479016 | 5.25E-05 | 0.001568268 |
| HIPK1 | 0.649530202 | 5.35E-05 | 0.001589317 |
| C1QTNF12 | 0.602091543 | 5.46E-05 | 0.001620112 |
| TNFSF13B | 1.953200693 | 5.55E-05 | 0.001643805 |
| TGIF1 | -1.219993779 | 5.58E-05 | 0.001647921 |
| ENSACYG00000009966 | 0.802272977 | 5.69E-05 | 0.001673225 |
| MTHFR | 0.581370297 | 5.70E-05 | 0.001673225 |
| CXCR5 | 2.944047405 | 5.70E-05 | 0.001673225 |
| SLC5A7 | 0.812560708 | 5.73E-05 | 0.001678367 |
| LTO1 | -0.713370347 | 5.79E-05 | 0.001691729 |
| NAE1 | -0.639777317 | 5.84E-05 | 0.001700248 |
| ENSACYG00000004268 | 5.592818313 | 5.85E-05 | 0.001700248 |
| TIAL1 | -0.578547186 | 5.89E-05 | 0.001708504 |
| ENSACYG00000006242 | 0.87694933 | 5.96E-05 | 0.001725019 |
| DMP1 | 3.878746965 | 5.99E-05 | 0.001725345 |
| ENSACYG00000001849 | 0.67548638 | 5.98E-05 | 0.001725345 |
| CNOT7 | 0.533277038 | 6.05E-05 | 0.001741492 |
| XRCC5 | -0.779701162 | 6.11E-05 | 0.001745614 |
| MEOX2 | -0.667092465 | 6.09E-05 | 0.001745614 |
| PLEKHF1 | -0.833526908 | 6.10E-05 | 0.001745614 |
| HPSE | -0.655735253 | 6.20E-05 | 0.001763552 |
| ZWILCH | -0.633641914 | 6.19E-05 | 0.001763552 |
| ANXA6 | -0.830187903 | 6.27E-05 | 0.001781058 |
| CUNH1orf109 | -0.791454761 | 6.43E-05 | 0.001818747 |
| NBN | -0.627401018 | 6.42E-05 | 0.001818747 |
| RFC4 | -0.777393905 | 6.49E-05 | 0.00182963 |
| GLMN | -0.671503926 | 6.54E-05 | 0.001840245 |
| SPTBN1 | -0.666939988 | 6.61E-05 | 0.001855494 |
| HDX | -0.75549689 | 6.66E-05 | 0.001865703 |
| TAX1BP3 | 0.597955322 | 6.73E-05 | 0.001882062 |
| PRTG | 2.054020524 | 6.79E-05 | 0.00189384 |
| URB1 | -0.879792809 | 6.83E-05 | 0.0019015 |
| LOC118261526 | 5.611832357 | 6.97E-05 | 0.00193449 |
| RBM7 | -0.721068336 | 6.96E-05 | 0.00193449 |
| BMPR2 | 0.593182784 | 7.02E-05 | 0.001942458 |

| | | | |
|---|---|---|---|
| AFF2 | 0.772664967 | 7.03E-05 | 0.001943 |
| CSTF1 | -0.671833591 | 7.38E-05 | 0.002034669 |
| EIF2B1 | -0.610728526 | 7.43E-05 | 0.002043274 |
| CDC37L1 | -0.70954265 | 7.49E-05 | 0.002050969 |
| LOC118259344 | -0.738636485 | 7.49E-05 | 0.002050969 |
| SCG3 | -0.739415535 | 7.74E-05 | 0.002114141 |
| ITGB1BP1 | -0.706772174 | 7.75E-05 | 0.002114141 |
| DUSP10 | -0.624322488 | 7.82E-05 | 0.002127907 |
| IGFBP7 | -0.620687373 | 7.94E-05 | 0.002152602 |
| MBNL2 | 0.613201587 | 7.93E-05 | 0.002152602 |
| THNSL2 | -0.799469252 | 8.10E-05 | 0.002187187 |
| FGF9 | 0.713587955 | 8.11E-05 | 0.002187187 |
| ZCCHC2 | 0.735851479 | 8.12E-05 | 0.002187187 |
| LOC118257380 | -0.737061011 | 8.22E-05 | 0.002210262 |
| DYSF | -0.682084862 | 8.40E-05 | 0.002253429 |
| CFL2 | 0.591784502 | 8.42E-05 | 0.002253993 |
| LOC118245549 | -1.902091725 | 8.46E-05 | 0.002261611 |
| KAT6B | -0.83634461 | 8.52E-05 | 0.00227203 |
| EAPP | -0.66397705 | 9.16E-05 | 0.00243941 |
| SC5D | 0.563872208 | 9.34E-05 | 0.00247619 |
| LOC118257919 | 5.984608151 | 9.33E-05 | 0.00247619 |
| GMNN | -0.749577467 | 9.37E-05 | 0.002478834 |
| NIM1K | 2.36467093 | 9.56E-05 | 0.002523568 |
| EXOC1 | -0.697504066 | 9.69E-05 | 0.002552201 |
| CREB3L1 | 0.966071903 | 9.74E-05 | 0.0025622 |
| ABCG2 | 3.340175697 | 9.79E-05 | 0.002563897 |
| MGLL | 0.844067336 | 9.77E-05 | 0.002563897 |
| NATD1 | 0.8860576 | 0.000100296 | 0.002621983 |
| TPP1 | -0.548053161 | 0.000101491 | 0.002647986 |
| ENSACYG00000013245 | 1.179075791 | 0.000102011 | 0.002656305 |
| MICU1 | -0.538650788 | 0.000102669 | 0.002668188 |
| UST | 0.571820074 | 0.0001035 | 0.0026845 |
| ENSACYG00000017148 | 5.518920504 | 0.000104369 | 0.002701726 |
| NEB | 1.703342533 | 0.000105386 | 0.002722725 |
| FN1 | -0.595774231 | 0.000106564 | 0.002743118 |
| EPRS1 | -0.575987506 | 0.000106591 | 0.002743118 |
| CDK7 | -0.643056931 | 0.000108668 | 0.002785691 |
| LOC118252592 | -0.724772032 | 0.000108584 | 0.002785691 |
| THAP12 | -0.972231953 | 0.000109727 | 0.002807391 |
| ENSACYG00000000083 | 2.507231261 | 0.000111024 | 0.002824153 |
| ST6GALNAC2 | -0.763206321 | 0.000110907 | 0.002824153 |
| LOC118260034 | -0.613440361 | 0.000110705 | 0.002824153 |
| DMAP1 | -0.789695465 | 0.000111467 | 0.002829966 |
| PMS1 | -0.675454978 | 0.000111886 | 0.002833929 |
| MTFR2 | -0.775296021 | 0.000112052 | 0.002833929 |
| RAD17 | -0.730587217 | 0.000112441 | 0.002838331 |
| SELE | 5.597246114 | 0.000113009 | 0.00284647 |
| APPBP2 | -0.653314028 | 0.000113626 | 0.00284647 |

| | | | |
|---|---|---|---|
| NHP2 | -0.739106678 | 0.000113566 | 0.00284647 |
| LOC118248865 | 5.908234277 | 0.000113595 | 0.00284647 |
| LOC118255061 | 0.751254632 | 0.000114581 | 0.002864947 |
| LOC118255340 | 0.879670797 | 0.000115972 | 0.00289425 |
| LOC118257056 | -0.985508954 | 0.000116243 | 0.002895539 |
| COBL | -1.240212151 | 0.000117528 | 0.002922031 |
| PRRG3 | 1.101662913 | 0.000117924 | 0.002926385 |
| RNF19B | 1.238824155 | 0.000118242 | 0.002928754 |
| TTI2 | -0.684693483 | 0.000118629 | 0.002932837 |
| MRPL46 | -0.656996806 | 0.000119715 | 0.002948646 |
| CCDC88A | -0.774857391 | 0.000119672 | 0.002948646 |
| ENSACYG00000007688 | -0.906931227 | 0.000120488 | 0.00296135 |
| RNF144A | 0.686880761 | 0.000120679 | 0.00296135 |
| TNK2 | -0.95366246 | 0.00012202 | 0.002977886 |
| CCNI | 0.589774562 | 0.000121795 | 0.002977886 |
| SERPINE2 | -0.576148852 | 0.00012203 | 0.002977886 |
| DNAJC16 | 0.676890118 | 0.000122394 | 0.002981253 |
| EVL | 0.57695267 | 0.000123545 | 0.003003761 |
| LOC118245354 | 0.70765823 | 0.000124548 | 0.003022583 |
| SMC2 | -0.639482295 | 0.000124956 | 0.003026908 |
| ENSACYG00000016847 | -0.88102821 | 0.000127091 | 0.003072993 |
| TSC22D3 | -0.791497373 | 0.000128772 | 0.00310794 |
| HIVEP2 | 0.862449358 | 0.000130455 | 0.003142812 |
| RAI2 | 0.94259211 | 0.000131731 | 0.003162502 |
| DNAJC5G | -0.769697821 | 0.000131751 | 0.003162502 |
| TUFT1 | -0.520938296 | 0.00013316 | 0.003190523 |
| PRPF4B | -0.664211731 | 0.000134341 | 0.00321298 |
| RBM17 | -0.592022159 | 0.00013806 | 0.003290016 |
| CSPG4 | -0.830825573 | 0.000137969 | 0.003290016 |
| AP5Z1 | -0.81313534 | 0.000141518 | 0.003364964 |
| HINFP | -0.868855841 | 0.000141715 | 0.003364964 |
| EFNA1 | -1.172593947 | 0.000144662 | 0.003428764 |
| CTSH | -0.516459065 | 0.000145272 | 0.00343707 |
| CUNH2orf50 | 1.766997217 | 0.000147998 | 0.003495292 |
| COL6A3 | -0.597345467 | 0.000150553 | 0.003549291 |
| SAMSN1 | 5.885751251 | 0.000151833 | 0.003573086 |
| RPUSD1 | 0.675760717 | 0.00015385 | 0.003614113 |
| OLFM1 | 0.821055133 | 0.000155856 | 0.003654724 |
| ZFYVE19 | -1.039237684 | 0.00015739 | 0.003684144 |
| CPE | -0.882418583 | 0.000160837 | 0.003758186 |
| ENSACYG00000006869 | 0.636510763 | 0.000161743 | 0.003770653 |
| VPS33A | -0.587689777 | 0.000162166 | 0.003770653 |
| DCAF5 | 0.636605671 | 0.000162228 | 0.003770653 |
| SRGAP2 | 0.63532288 | 0.000162647 | 0.003773749 |
| MAPKAP1 | 0.584317188 | 0.000163005 | 0.003775434 |
| LOC118247959 | -0.645008026 | 0.000163797 | 0.003787128 |
| HSPD1 | -0.643147168 | 0.000164704 | 0.003788197 |
| VASH2 | 0.870254188 | 0.000164162 | 0.003788197 |

| | | | |
|---|---|---|---|
| SLC30A4 | 0.796619981 | 0.000164458 | 0.003788197 |
| CTNS | -0.629780709 | 0.000167721 | 0.003844179 |
| LOC118245892 | -0.90387024 | 0.000167437 | 0.003844179 |
| RBBP6 | 0.580041551 | 0.0001683 | 0.003850784 |
| VLDLR | -0.733206222 | 0.000170112 | 0.003885499 |
| NUP93 | -0.597411287 | 0.000172104 | 0.00392422 |
| ENSACYG00000008373 | 1.541268004 | 0.000173018 | 0.003938239 |
| NSUN5 | -0.563780703 | 0.000174857 | 0.003973251 |
| NHEJ1 | -2.071973714 | 0.000175514 | 0.003981344 |
| LMO7 | -0.613524128 | 0.000176682 | 0.004000953 |
| ST3GAL1 | 0.551945686 | 0.000177129 | 0.004004216 |
| IRF8 | 1.097135429 | 0.00018108 | 0.004086516 |
| LOC118255699 | -0.957917994 | 0.000182537 | 0.004112384 |
| PIAS2 | 0.553188229 | 0.000185508 | 0.004172187 |
| GPR162 | 0.627988785 | 0.000192714 | 0.004319546 |
| ADHFE1 | 0.54380821 | 0.000192623 | 0.004319546 |
| DCSTAMP | -0.789818473 | 0.000193834 | 0.004329933 |
| R3HDM2 | 0.695059567 | 0.000193599 | 0.004329933 |
| MKKS | 0.655008318 | 0.000195651 | 0.004363156 |
| SELENOO | -0.699183709 | 0.000197611 | 0.004392019 |
| FAM219A | 0.622580401 | 0.000197474 | 0.004392019 |
| CBX8 | -0.72563813 | 0.000198602 | 0.00440663 |
| TPX2 | -0.549395031 | 0.000199026 | 0.004408616 |
| EIF3A | -0.599512739 | 0.000200348 | 0.004415682 |
| RCHY1 | -0.542336754 | 0.000200146 | 0.004415682 |
| LOC118259472 | -0.658960942 | 0.000199733 | 0.004415682 |
| ENSACYG00000003165 | 3.879709475 | 0.000203477 | 0.004477168 |
| LOC118244561 | -0.622431248 | 0.000205107 | 0.00450553 |
| YRDC | -0.645616105 | 0.000205525 | 0.00450721 |
| CCNI2 | 0.528302822 | 0.000206676 | 0.004524946 |
| CD40 | -0.836317779 | 0.000210521 | 0.004580056 |
| KATNA1 | -0.695454035 | 0.000210581 | 0.004580056 |
| MYO5B | 2.540875821 | 0.000210439 | 0.004580056 |
| CEP44 | -0.678148928 | 0.000210297 | 0.004580056 |
| LOC118253960 | 0.743146197 | 0.000213032 | 0.00462573 |
| CCDC71L | -0.882369778 | 0.00021484 | 0.004657341 |
| CNOT4 | 0.688111014 | 0.000220238 | 0.004758719 |
| FER1L6 | 5.820352293 | 0.000220178 | 0.004758719 |
| SGMS2 | 0.504829249 | 0.0002223 | 0.004795426 |
| LRRC8B | 0.772661574 | 0.000223712 | 0.004818027 |
| ARHGAP10 | -0.619996 | 0.000224295 | 0.004822712 |
| TARDBP | -0.573285548 | 0.000228692 | 0.004909245 |
| ENSACYG00000005007 | 0.702254525 | 0.00023031 | 0.004935969 |
| DKK2 | 0.670476995 | 0.000232623 | 0.004977453 |
| TP53RK | -0.549521516 | 0.000237529 | 0.005074209 |
| WDR75 | -0.65746501 | 0.000240253 | 0.005116842 |
| HGH1 | -0.638105424 | 0.0002403 | 0.005116842 |
| DNAJC17 | -0.719049863 | 0.000244157 | 0.005182255 |

| | | | |
|---|---|---|---|
| VCPKMT | -0.928032686 | 0.000244139 | 0.005182255 |
| UVRAG | 0.664236354 | 0.000245018 | 0.005192181 |
| HEXD | -0.886142631 | 0.00024669 | 0.005219236 |
| LOC118260553 | -0.731454454 | 0.000250034 | 0.005281511 |
| ZDHHC3 | 0.613908155 | 0.000254508 | 0.005367428 |
| POLD2 | -0.857049806 | 0.000256891 | 0.005409056 |
| PAQR3 | -0.570088623 | 0.000258485 | 0.005433951 |
| SPRING1 | 0.61311135 | 0.000259647 | 0.005447518 |
| CUNH12orf4 | -0.605798487 | 0.000259956 | 0.005447518 |
| CCT8 | -0.527754931 | 0.000261607 | 0.005456143 |
| POMT2 | 0.809922252 | 0.000260913 | 0.005456143 |
| LOC118245314 | -0.663070951 | 0.000261267 | 0.005456143 |
| FGFR2 | 0.719237755 | 0.000262556 | 0.005467303 |
| ENSACYG00000000051 | 5.772230898 | 0.000265281 | 0.005515339 |
| MMP2 | -0.501179174 | 0.000265892 | 0.005519339 |
| ENSACYG00000000843 | 5.778008077 | 0.000268735 | 0.005569614 |
| ETFBKMT | -0.898468648 | 0.000269825 | 0.005583436 |
| ADGRL3 | 1.161291356 | 0.000271476 | 0.005608799 |
| CMTM7 | 0.620172761 | 0.000271953 | 0.005609877 |
| CHM | -0.623657836 | 0.000273145 | 0.005612798 |
| TRAPPC12 | -0.559824352 | 0.00027337 | 0.005612798 |
| RGP1 | -0.646164538 | 0.000273016 | 0.005612798 |
| ENSACYG00000005978 | 1.054726806 | 0.000275269 | 0.005643021 |
| LOC118257862 | -0.950630422 | 0.000277579 | 0.005681552 |
| PLK2 | -0.656400813 | 0.000281318 | 0.00574916 |
| LOC118253380 | -0.559095753 | 0.000282307 | 0.005760468 |
| DIABLO | -0.5933467 | 0.000284952 | 0.005805455 |
| MED16 | 0.834878775 | 0.000290106 | 0.005901351 |
| DDX28 | -0.730702393 | 0.000292523 | 0.00594137 |
| RPP38 | -0.686381277 | 0.000293524 | 0.005952538 |
| ENSACYG00000005870 | 2.992632605 | 0.000294949 | 0.005972257 |
| MED7 | -0.640270971 | 0.000301022 | 0.006085898 |
| TMEM179B | -0.744091919 | 0.000301573 | 0.006087715 |
| LRRC8C | -0.580223408 | 0.000302848 | 0.006094814 |
| MEX3C | 0.603058416 | 0.000302431 | 0.006094814 |
| MGAT4B | 0.68868942 | 0.000303526 | 0.006099161 |
| SAXO2 | 1.266025727 | 0.000304781 | 0.006115078 |
| AGAP3 | 0.80653259 | 0.000306558 | 0.0061414 |
| MRPL19 | -0.616188982 | 0.0003079 | 0.006156576 |
| BRD8 | -0.811009661 | 0.000309181 | 0.006156576 |
| FAM110B | -0.747571492 | 0.000308854 | 0.006156576 |
| SLC22A23 | 0.73506827 | 0.000308256 | 0.006156576 |
| TLN2 | -0.992446217 | 0.000310634 | 0.006176196 |
| TTC26 | -0.643507394 | 0.000312289 | 0.00619975 |
| USP10 | 0.533451228 | 0.000313102 | 0.006206559 |
| DUS2 | -0.721012095 | 0.000314386 | 0.00622268 |
| PTP4A1 | 0.569313466 | 0.000314863 | 0.006222791 |
| EPB41L2 | -0.501603675 | 0.000318483 | 0.00627554 |

| | | | |
|---|---|---|---|
| LOC118246781 | 5.776775137 | 0.000318334 | 0.00627554 |
| TAF8 | -0.963834821 | 0.00031923 | 0.006280877 |
| ENSACYG00000006528 | 0.90289057 | 0.00032113 | 0.006308864 |
| ZFAND2A | -0.629020958 | 0.000323589 | 0.006347725 |
| IL22RA2 | 4.839302396 | 0.000324174 | 0.006349772 |
| ENSACYG00000000669 | 0.842371322 | 0.000327814 | 0.006377222 |
| YTHDF1 | 0.614150659 | 0.000327093 | 0.006377222 |
| TACC2 | 0.676751115 | 0.000327991 | 0.006377222 |
| ENSACYG00000007530 | 1.693949073 | 0.000327484 | 0.006377222 |
| PACSIN2 | -0.569326851 | 0.000327919 | 0.006377222 |
| CCNY | 0.65945913 | 0.000330686 | 0.006410735 |
| ENSACYG00000004254 | 4.656301664 | 0.000330382 | 0.006410735 |
| SORBS2 | 0.856595673 | 0.00033225 | 0.006431617 |
| CD276 | -0.509810014 | 0.000335399 | 0.006483063 |
| ENSACYG00000012635 | 3.835716589 | 0.00033899 | 0.0065429 |
| SIL1 | -0.593661368 | 0.000342366 | 0.006598419 |
| KPNA2 | -0.55194254 | 0.000348354 | 0.00670403 |
| MADCAM1 | 5.679546605 | 0.0003491 | 0.006708621 |
| CUNH16orf70 | -0.621908596 | 0.000350395 | 0.006723705 |
| ZNF704 | 0.539386973 | 0.000356061 | 0.006812631 |
| LOC118244782 | 1.350402883 | 0.000355623 | 0.006812631 |
| NRBP2 | 0.756186724 | 0.000359344 | 0.006865493 |
| NFATC3 | 0.76736878 | 0.000360116 | 0.006870308 |
| PRDM1 | 0.839179524 | 0.000363688 | 0.006928434 |
| AKAP17A | -0.601832071 | 0.000367312 | 0.006987394 |
| FNDC11 | 5.235161218 | 0.000368992 | 0.007009252 |
| PDXP | -0.586272644 | 0.000370585 | 0.007029393 |
| BTBD7 | 0.544208675 | 0.000372859 | 0.007062383 |
| SYT16 | 0.906559726 | 0.000373955 | 0.007072995 |
| MID1IP1 | -0.784694049 | 0.000377307 | 0.007116016 |
| GPX3 | 0.577599118 | 0.000376899 | 0.007116016 |
| LOC118258430 | 4.065431761 | 0.000383978 | 0.007231495 |
| RGCC | -0.632435626 | 0.000386401 | 0.007266767 |
| LOC118254681 | -0.768268523 | 0.000390437 | 0.007325653 |
| FGF18 | 2.381351024 | 0.000390642 | 0.007325653 |
| SEPTIN11 | -0.608342261 | 0.000394916 | 0.007384809 |
| MASP2 | 5.199900169 | 0.000394915 | 0.007384809 |
| PCDH17 | 0.954321114 | 0.00039724 | 0.007417764 |
| LOC118260312 | 5.203158461 | 0.000399756 | 0.007443698 |
| DDX6 | -0.553632071 | 0.000399715 | 0.007443698 |
| PARP9 | 0.525102535 | 0.000401563 | 0.007466813 |
| ST7L | -0.524725161 | 0.000403359 | 0.00748966 |
| VCL | -0.592090524 | 0.000405717 | 0.00752286 |
| CTSS | -0.55820435 | 0.000408737 | 0.007568226 |
| LOC118247783 | 5.654160177 | 0.000422103 | 0.007802605 |
| TTL | 0.628632417 | 0.000422624 | 0.007802605 |
| KCTD2 | 0.775944718 | 0.000423168 | 0.007802605 |
| BCL2 | 0.667750455 | 0.000425706 | 0.007838461 |

| | | | |
|---|---|---|---|
| FAM177A1 | 0.612177447 | 0.000427783 | 0.007854792 |
| ZNF639 | -0.64570522 | 0.000427355 | 0.007854792 |
| ARHGAP35 | 0.845325755 | 0.000430416 | 0.007892163 |
| ZNF830 | -0.656790483 | 0.000431416 | 0.007899514 |
| LOC118245664 | 2.209559364 | 0.000434631 | 0.007947371 |
| SMARCA1 | -0.576155501 | 0.000436525 | 0.007970961 |
| TFPI2 | -1.429678714 | 0.000443886 | 0.008079899 |
| ENSACYG00000016016 | 1.130203763 | 0.000444327 | 0.008079899 |
| PREB | -0.673567571 | 0.000443869 | 0.008079899 |
| B3GALNT2 | 0.589028198 | 0.000445542 | 0.008090853 |
| GABRA4 | 1.048107023 | 0.000449262 | 0.00814719 |
| SELPLG | 5.173983799 | 0.000450512 | 0.008158658 |
| NFATC1 | 0.931054709 | 0.000453302 | 0.008164375 |
| ENSACYG00000008594 | -0.75606771 | 0.000452871 | 0.008164375 |
| ENSACYG00000013730 | -0.76390651 | 0.000452018 | 0.008164375 |
| SLC41A2 | 0.602648263 | 0.000453107 | 0.008164375 |
| SYNE2 | -0.788656795 | 0.000454179 | 0.008169031 |
| FGF2 | 0.680438599 | 0.000455427 | 0.008180341 |
| DAGLB | -0.727731815 | 0.000457691 | 0.008201662 |
| FKBP11 | -0.740290486 | 0.000457857 | 0.008201662 |
| ENSACYG00000005318 | -0.59477981 | 0.000459125 | 0.008202126 |
| ENSACYG00000016331 | 1.027512875 | 0.000458986 | 0.008202126 |
| LOC118253067 | -0.713108063 | 0.000461041 | 0.00822522 |
| KCNE1 | 0.92129349 | 0.000462688 | 0.008243464 |
| BNIP1 | -0.675096294 | 0.000466025 | 0.00829172 |
| TMEM204 | 0.777194178 | 0.000472314 | 0.008392318 |
| TUBE1 | -0.648609245 | 0.000479402 | 0.008502184 |
| LOC118258757 | -0.503979527 | 0.000479785 | 0.008502184 |
| LOC118256212 | -0.937457433 | 0.000482584 | 0.008540309 |
| N6AMT1 | -0.678110018 | 0.000486029 | 0.008566833 |
| NFATC2 | 0.566545357 | 0.000485464 | 0.008566833 |
| ZNF652 | 0.616182332 | 0.000485927 | 0.008566833 |
| FOXL1 | 2.406656603 | 0.00048699 | 0.008572318 |
| CCT2 | -0.492202942 | 0.000489351 | 0.008602414 |
| ABHD5 | 0.661745077 | 0.000491541 | 0.008628426 |
| MAP3K3 | 0.957858308 | 0.000492138 | 0.008628426 |
| ENSACYG00000009513 | 2.369370458 | 0.00049369 | 0.008644154 |
| PTHLH | -0.600689125 | 0.000498894 | 0.008723709 |
| CENPE | -0.617083453 | 0.000500922 | 0.008747584 |
| ZDHHC14 | 0.657521861 | 0.000509812 | 0.008891074 |
| LIF | 0.60990564 | 0.000512633 | 0.008928464 |
| AIF1L | 0.550259681 | 0.000514143 | 0.00894297 |
| ENSACYG00000010664 | 1.258774035 | 0.000515093 | 0.008947708 |
| ALAS1 | -0.476250252 | 0.000518537 | 0.008995696 |
| MBNL1 | 0.659250331 | 0.000520443 | 0.009016918 |
| CUNH1orf52 | -0.846559181 | 0.000522597 | 0.009042366 |
| ENSACYG00000005373 | 3.581259164 | 0.000529613 | 0.009151767 |
| SLC35A1 | -0.670964802 | 0.000530971 | 0.009163239 |

| | | | |
|---|---|---|---|
| ENSACYG00000009389 | 1.546091427 | 0.000534411 | 0.009210575 |
| ZDHHC7 | 0.712308611 | 0.000537849 | 0.00925241 |
| NUP85 | -0.491680743 | 0.00053824 | 0.00925241 |
| LOC118256438 | 1.172131077 | 0.000539666 | 0.00926485 |
| LOC118245864 | 0.661480596 | 0.000540629 | 0.009269329 |
| LOC118251023 | -0.716803775 | 0.000542758 | 0.00929376 |
| ENSACYG00000002908 | 0.746756415 | 0.000551312 | 0.009428009 |
| ACAT2 | -0.506408977 | 0.000557209 | 0.00951652 |
| ZNF335 | 0.92183911 | 0.000558638 | 0.009528605 |
| XRN2 | -0.477634031 | 0.000562911 | 0.009589095 |
| CD109 | -0.767313591 | 0.000566177 | 0.009632313 |
| SCLT1 | -0.61699844 | 0.000569993 | 0.009647491 |
| TMEM267 | 0.492118789 | 0.000569855 | 0.009647491 |
| EFNB1 | 0.492338961 | 0.000569923 | 0.009647491 |
| CEP57 | -0.756222002 | 0.000567845 | 0.009647491 |
| CHRAC1 | -0.609089037 | 0.000573148 | 0.009688477 |
| LRFN5 | 0.695952215 | 0.00057451 | 0.009699072 |
| ATXN1L | 0.763694849 | 0.000575362 | 0.009701067 |
| ZNF831 | 2.150257985 | 0.000577836 | 0.009717949 |
| GFPT2 | -0.494712326 | 0.000577442 | 0.009717949 |
| TMEM18 | -0.695480515 | 0.000579466 | 0.009732964 |
| TAB3 | 0.684976409 | 0.000580458 | 0.009737234 |
| GPBP1L1 | 0.623641168 | 0.000587781 | 0.009847573 |
| RBM5 | -0.473853804 | 0.000592898 | 0.00992071 |
| TMLHE | 0.724444954 | 0.000596125 | 0.009940215 |
| N4BP2L1 | 0.603220692 | 0.000596672 | 0.009940215 |
| KDM4C | -0.57490819 | 0.000596244 | 0.009940215 |
| LOC118257087 | -0.755487387 | 0.000597075 | 0.009940215 |
| BRF2 | -1.021685557 | 0.000604226 | 0.010046586 |
| KIF11 | -0.750814842 | 0.000605979 | 0.010063056 |
| LPAR3 | 5.083463792 | 0.000607502 | 0.01006589 |
| ENSACYG00000010040 | 0.603164788 | 0.000607674 | 0.01006589 |
| LOC118243356 | 0.484433935 | 0.00061225 | 0.01012897 |
| ATF3 | 0.49851386 | 0.000614149 | 0.010147683 |
| LCLAT1 | 0.675573694 | 0.000617112 | 0.010183895 |
| PUM3 | -0.510464519 | 0.000620772 | 0.010231492 |
| ANKRD33B | 5.122353913 | 0.000621882 | 0.010237015 |
| ARL5A | 0.612396682 | 0.000624692 | 0.010267111 |
| ENSACYG00000010064 | -3.820381332 | 0.000625266 | 0.010267111 |
| UCHL5 | -0.596297333 | 0.000627667 | 0.010293733 |
| AOPEP | -0.579910339 | 0.000647921 | 0.010612728 |
| PDP2 | -0.564386037 | 0.000655416 | 0.010691636 |
| CKAP5 | -0.766863296 | 0.000653654 | 0.010691636 |
| DUS1L | -0.654241069 | 0.000654987 | 0.010691636 |
| ENSACYG00000015618 | 0.524870566 | 0.000655978 | 0.010691636 |
| GHITM | -0.502245167 | 0.000658614 | 0.010696381 |
| CPSF4 | 0.468420408 | 0.0006587 | 0.010696381 |
| PLTP | -0.597414933 | 0.000657351 | 0.010696381 |

| | | | |
|---|---|---|---|
| ENSACYG00000002600 | 0.709938822 | 0.000659913 | 0.010702914 |
| ARPC4 | -0.619120412 | 0.000662819 | 0.010736856 |
| SLC35F3 | 0.747785479 | 0.000667604 | 0.010798368 |
| ZNF319 | 0.676735298 | 0.000668252 | 0.010798368 |
| ENSACYG00000008946 | 3.916204769 | 0.000670417 | 0.010820099 |
| STXBP6 | 0.640294903 | 0.000676826 | 0.010905322 |
| PAPSS1 | -0.491892726 | 0.000678175 | 0.010905322 |
| PPM1B | 0.545630533 | 0.000677875 | 0.010905322 |
| RAP2C | 0.594723583 | 0.0006837 | 0.010980795 |
| LOC118244824 | -0.627743491 | 0.000688858 | 0.011050188 |
| ENSACYG00000005176 | 1.284448644 | 0.000692444 | 0.011088593 |
| SPIN1 | 0.553304712 | 0.000692932 | 0.011088593 |
| XRCC3 | -0.843233887 | 0.000694569 | 0.011101334 |
| GOLGB1 | -0.499446032 | 0.000696388 | 0.011116953 |
| LOC118251874 | -1.123149196 | 0.000701251 | 0.011167566 |
| NSUN6 | -0.816291548 | 0.000701137 | 0.011167566 |
| TSEN15 | -0.755088385 | 0.000708996 | 0.011213287 |
| LOC118253620 | -0.57926497 | 0.000709854 | 0.011213287 |
| IMPACT | -0.650103695 | 0.000709841 | 0.011213287 |
| CKS2 | -0.710049265 | 0.000710067 | 0.011213287 |
| FNDC7 | 3.336027606 | 0.00070799 | 0.011213287 |
| PUF60 | -0.484323335 | 0.000708644 | 0.011213287 |
| SLC25A25 | 0.736566825 | 0.000707133 | 0.011213287 |
| BBS2 | -0.49351549 | 0.000711661 | 0.011225035 |
| ENSACYG00000000170 | 2.22482295 | 0.000713079 | 0.011233967 |
| MOGS | -0.837110025 | 0.00071411 | 0.0112368 |
| TMEM38A | 0.481726002 | 0.000716442 | 0.011260088 |
| ENSACYG00000006103 | 0.598582186 | 0.000721454 | 0.011325365 |
| AGO2 | 0.719022839 | 0.000724501 | 0.011359689 |
| LTN1 | -0.643833835 | 0.00072903 | 0.011390116 |
| DCBLD2 | 0.573639242 | 0.000728878 | 0.011390116 |
| NAB1 | 0.524826775 | 0.000727623 | 0.011390116 |
| LRMDA | 0.518551671 | 0.000732073 | 0.011410661 |
| STAMBP | -0.509107763 | 0.000731393 | 0.011410661 |
| POLR1A | -0.646504484 | 0.000733382 | 0.011417582 |
| ENSACYG00000012047 | 1.077486031 | 0.000736817 | 0.011457554 |
| LOC118248965 | -0.880618936 | 0.000741826 | 0.011521868 |
| ENSACYG00000009906 | 0.520792927 | 0.000744647 | 0.011538526 |
| ATG13 | 0.813391215 | 0.000744505 | 0.011538526 |
| ADAMTS12 | 0.632278538 | 0.000748902 | 0.011590862 |
| P2RX5 | 1.035284162 | 0.000756098 | 0.011674866 |
| LOC118254393 | -0.519453812 | 0.000755469 | 0.011674866 |
| WDR45B | 0.518845345 | 0.000759085 | 0.011680046 |
| TMPRSS2 | 5.508750093 | 0.000759088 | 0.011680046 |
| SRGN | -0.671606505 | 0.000758148 | 0.011680046 |
| IFT57 | -0.652876569 | 0.000760512 | 0.011688342 |
| RNF186 | 0.733213474 | 0.000774374 | 0.011887546 |
| ARL4C | -0.561192747 | 0.000781558 | 0.011983892 |

| | | | |
|---|---|---|---|
| PCGF6 | -0.827508534 | 0.000786433 | 0.012044652 |
| ENSACYG00000001336 | 2.12251559 | 0.000788312 | 0.012059441 |
| PTCHD1 | 0.794074802 | 0.000790331 | 0.012076334 |
| HADHB | -0.491719316 | 0.000792997 | 0.012103065 |
| SLC52A3 | -0.676138346 | 0.000798048 | 0.012166082 |
| SPTB | -0.756331152 | 0.000801006 | 0.012193926 |
| MYBL2 | 0.577852129 | 0.000801722 | 0.012193926 |
| DUSP6 | -0.501921963 | 0.000802857 | 0.012196517 |
| NAA25 | -0.479250428 | 0.00080374 | 0.012196517 |
| SPSB3 | -0.583815187 | 0.000805488 | 0.012209008 |
| ENSACYG00000007059 | 0.97796631 | 0.000809025 | 0.012234538 |
| LOC118256771 | 4.467989192 | 0.000808737 | 0.012234538 |
| SLC40A1 | 0.667891463 | 0.000809987 | 0.012235061 |
| ENTPD5 | 0.758033975 | 0.000816986 | 0.012312606 |
| PHYHD1 | 1.245231725 | 0.00081686 | 0.012312606 |
| ABCB6 | -0.836699141 | 0.000820622 | 0.012325199 |
| ALS2CL | 0.741303145 | 0.000820136 | 0.012325199 |
| ENSACYG00000010518 | 2.151393964 | 0.000820398 | 0.012325199 |
| CTSZ | -0.556152111 | 0.000824968 | 0.012370616 |
| ANKRD49 | -0.5990997 | 0.00082552 | 0.012370616 |
| CLK4 | -0.60319734 | 0.000830626 | 0.012433025 |
| MRM3 | -0.632910747 | 0.000834839 | 0.012481931 |
| PRMT7 | -0.676757388 | 0.000837561 | 0.012508459 |
| RASA2 | -0.844794286 | 0.000845194 | 0.012608186 |
| CTHRC1 | -0.56258917 | 0.000846318 | 0.012610705 |
| HIPK3 | 0.622136184 | 0.000848031 | 0.012621984 |
| LOC118255547 | 4.451382364 | 0.000851259 | 0.012640107 |
| CUNH3orf14 | 0.515832996 | 0.000852121 | 0.012640107 |
| RPS6 | -0.542642917 | 0.000852064 | 0.012640107 |
| TTYH2 | 0.652348178 | 0.000856491 | 0.012689662 |
| GEMIN5 | -0.607732742 | 0.000857384 | 0.012689662 |
| UBE4A | 0.788083583 | 0.000859732 | 0.012710171 |
| TARBP1 | -0.782679386 | 0.000860831 | 0.012712186 |
| PRIM1 | -0.600396755 | 0.000880102 | 0.012982249 |
| ID4 | -0.769011704 | 0.000881702 | 0.012991322 |
| RPF1 | -0.673092795 | 0.000888935 | 0.013068733 |
| VGLL3 | 0.486049441 | 0.000888232 | 0.013068733 |
| SRPX | -0.617177989 | 0.000891572 | 0.013092751 |
| FBXW5 | 0.811155102 | 0.000892552 | 0.013092751 |
| VPS37D | -0.734062875 | 0.000901776 | 0.013213366 |
| PLD6 | 5.023146149 | 0.000904609 | 0.013240179 |
| STS | 0.601245405 | 0.000905644 | 0.01324066 |
| ENSACYG00000012501 | 2.384479048 | 0.000908498 | 0.013253026 |
| CFAP92 | 4.961397409 | 0.000907969 | 0.013253026 |
| DDX47 | -0.483881348 | 0.000916375 | 0.013353181 |
| RASD1 | 1.948052723 | 0.000918954 | 0.013376 |
| MRPL2 | -0.612655403 | 0.000928745 | 0.013503626 |
| GTF3C5 | -0.610497362 | 0.000931437 | 0.013527866 |

| | | | |
|---|---|---|---|
| RAD52 | -0.687371352 | 0.00093412 | 0.013551929 |
| THG1L | -0.55745579 | 0.000943142 | 0.01366779 |
| RMDN3 | 0.740438092 | 0.000952426 | 0.013787198 |
| MEIS1 | 0.886151481 | 0.000953677 | 0.013790195 |
| GLIS1 | 0.84148522 | 0.0009678 | 0.013948576 |
| EZR | -0.571089518 | 0.000966792 | 0.013948576 |
| LOC118248843 | -0.553825317 | 0.000966896 | 0.013948576 |
| CUNH21orf91 | -0.567267887 | 0.000971582 | 0.013987818 |
| LOC118253250 | -0.554594211 | 0.000980264 | 0.014097439 |
| B4GALT3 | 0.687028522 | 0.000982165 | 0.01410941 |
| PLB1 | 1.700295067 | 0.000985701 | 0.014125087 |
| MTCH2 | -0.695902852 | 0.000989552 | 0.014125087 |
| ENSACYG00000002341 | -2.660909032 | 0.000989676 | 0.014125087 |
| LOC118247123 | 2.454066719 | 0.000988075 | 0.014125087 |
| FOXM1 | 0.877858151 | 0.000987569 | 0.014125087 |
| PAK1IP1 | -0.663945931 | 0.00098546 | 0.014125087 |
| CUNH2orf69 | -0.570880881 | 0.000992546 | 0.01415075 |
| NFKB2 | -0.554914429 | 0.000994897 | 0.014153697 |
| GRM7 | 0.587903155 | 0.000994613 | 0.014153697 |
| MMD | 0.535552032 | 0.000997381 | 0.01417376 |
| NDUFB9 | -0.654916935 | 0.001005952 | 0.014272872 |
| SLC29A3 | -0.80351248 | 0.001006517 | 0.014272872 |
| SCN8A | 0.606153909 | 0.001009483 | 0.014299564 |
| S1PR3 | 0.687825081 | 0.00101172 | 0.014315889 |
| GSG1 | 0.539321342 | 0.001018221 | 0.014392461 |
| MIA3 | -0.572064888 | 0.001020898 | 0.014414855 |
| UNC13B | 0.896697088 | 0.001027343 | 0.014467775 |
| ZNF521 | 0.480906295 | 0.001027933 | 0.014467775 |
| FNDC3A | 0.500617529 | 0.001025775 | 0.014467775 |
| PGBD5 | 0.815694999 | 0.001036108 | 0.014567303 |
| TMEM200A | 0.732880755 | 0.001042365 | 0.01463969 |
| ENSACYG00000004415 | 2.62384456 | 0.001051003 | 0.014745317 |
| GLIS3 | 0.501959562 | 0.001052356 | 0.014748618 |
| LOC118253833 | 2.233775145 | 0.001064557 | 0.014903804 |
| EXOSC7 | -0.582232858 | 0.001069734 | 0.014960415 |
| CUNH5orf24 | 0.452626488 | 0.001079928 | 0.015087 |
| GGT5 | 3.110555842 | 0.00108108 | 0.015087129 |
| RARS1 | -0.475879676 | 0.001093747 | 0.015247776 |
| BACE1 | 0.856754217 | 0.001098536 | 0.015298394 |
| CBFB | 0.467885618 | 0.00110112 | 0.015302094 |
| ENSACYG00000015590 | 3.122250684 | 0.001100793 | 0.015302094 |
| BLCAP | 0.996428377 | 0.001105144 | 0.015341864 |
| CACUL1 | 0.534027879 | 0.001112189 | 0.015423439 |
| FAM214A | 0.46831828 | 0.001115614 | 0.015454713 |
| STAT1 | 0.566851192 | 0.001123442 | 0.015546839 |
| NCLN | -0.492088193 | 0.00113155 | 0.015642641 |
| ENSACYG00000000629 | 0.797782377 | 0.001140933 | 0.01575585 |
| SMYD4 | -0.562426747 | 0.00114442 | 0.015787502 |

| | | | |
|---|---|---|---|
| GPR27 | 0.503371255 | 0.001151893 | 0.015874 |
| SYNM | -0.66207539 | 0.001155276 | 0.015885181 |
| PEAK1 | 0.463774607 | 0.001156314 | 0.015885181 |
| ICMT | 0.500086245 | 0.001155035 | 0.015885181 |
| DPP7 | -0.465917326 | 0.001164227 | 0.015962542 |
| INSIG1 | -0.483199923 | 0.001164364 | 0.015962542 |
| SCAMP4 | 0.752333839 | 0.001170104 | 0.016024599 |
| ECHS1 | -0.585410838 | 0.001172962 | 0.016047088 |
| DARS2 | -0.718044574 | 0.001177509 | 0.016092626 |
| EVI5L | 0.60943843 | 0.001195294 | 0.016318793 |
| ENSACYG00000006346 | 0.483865566 | 0.001205863 | 0.016446084 |
| HHAT | 0.518628569 | 0.001207214 | 0.016447509 |
| ROCK2 | -0.450881396 | 0.001217515 | 0.016570755 |
| ISY1 | -0.534320435 | 0.001220933 | 0.016600158 |
| EPAS1 | 0.763123594 | 0.001223589 | 0.016619163 |
| PTGS2 | -0.822925924 | 0.001228685 | 0.016671229 |
| HCRTR2 | 5.384345791 | 0.001236989 | 0.016766665 |
| ATR | -0.902515948 | 0.001246011 | 0.016871629 |
| MRPS10 | -0.64402296 | 0.001250914 | 0.016920661 |
| YARS2 | -0.569979496 | 0.001254549 | 0.016952456 |
| ENSACYG00000016817 | 3.880079818 | 0.001257098 | 0.016969542 |
| LOC118251000 | -0.697271747 | 0.001260654 | 0.017000155 |
| SUSD3 | 0.499143298 | 0.001267128 | 0.017070029 |
| ENSACYG00000004126 | 3.771591077 | 0.001273608 | 0.01712238 |
| DR1 | 0.479642834 | 0.001272993 | 0.01712238 |
| TKFC | -0.792622573 | 0.001278123 | 0.017165601 |
| PFN2 | 0.471210111 | 0.001283867 | 0.017225221 |
| ENSACYG00000000445 | 1.608980042 | 0.001290034 | 0.01729038 |
| CHRDL1 | 0.453787964 | 0.001292413 | 0.017304706 |
| GPRC5A | 5.383530647 | 0.001293752 | 0.017305076 |
| EXTL3 | 0.648028928 | 0.001299858 | 0.017369149 |
| DAP | 0.475676826 | 0.001302606 | 0.017388279 |
| ARRDC3 | -0.654261957 | 0.001304936 | 0.017401787 |
| AIMP1 | -0.563735431 | 0.001306603 | 0.017406433 |
| TRPC7 | 5.377329379 | 0.001317215 | 0.017530115 |
| CHD4 | -0.597758147 | 0.001321311 | 0.017549246 |
| FAP | -0.527893425 | 0.001321066 | 0.017549246 |
| ENSACYG00000006979 | 4.937651154 | 0.001323617 | 0.017562198 |
| LOC118246339 | 2.060417971 | 0.0013386 | 0.017743174 |
| DDX10 | -0.643809688 | 0.001340867 | 0.017755386 |
| RUFY1 | -0.534301443 | 0.001342752 | 0.017762539 |
| ENOX2 | 0.585861689 | 0.00134629 | 0.017791508 |
| TRIB2 | -0.912847106 | 0.001347878 | 0.017794691 |
| LOC118246505 | 1.61093871 | 0.001350594 | 0.017794947 |
| HSP90B1 | -0.44386302 | 0.001349978 | 0.017794947 |
| NMRK2 | -0.638978361 | 0.001355561 | 0.017842582 |
| KLB | 3.808116773 | 0.001368743 | 0.017998157 |
| ALG3 | -0.73075289 | 0.001375212 | 0.018065227 |

| | | | |
|---|---|---|---|
| TCF7 | 1.292420077 | 0.001378045 | 0.018084438 |
| PICK1 | -0.699917813 | 0.001379642 | 0.018087423 |
| BDNF | -0.493358235 | 0.001384423 | 0.018132093 |
| EXT1 | 0.611019702 | 0.001395553 | 0.018259747 |
| NR1H3 | 0.624918013 | 0.001404699 | 0.018361218 |
| RNF111 | 0.671866576 | 0.001415315 | 0.018481692 |
| IDS | -0.509329926 | 0.001417469 | 0.018491526 |
| IKBIP | -0.535243015 | 0.001420277 | 0.018503256 |
| ATP8B2 | 0.91212736 | 0.001421605 | 0.018503256 |
| ENSACYG00000015774 | 0.631116893 | 0.001422573 | 0.018503256 |
| SGO1 | -0.659193688 | 0.001451962 | 0.018866936 |
| ENSACYG00000004515 | 5.42612661 | 0.001455293 | 0.018873059 |
| PWWP2A | -0.478068319 | 0.001455259 | 0.018873059 |
| PSMB3 | -0.568554546 | 0.001456909 | 0.018875474 |
| KLHL31 | 1.40971134 | 0.001463029 | 0.018884569 |
| LOC118253663 | -0.53073572 | 0.001463332 | 0.018884569 |
| WBP1 | 0.815216962 | 0.001461079 | 0.018884569 |
| GPR89B | -0.519477906 | 0.001462631 | 0.018884569 |
| CDC42EP2 | -0.847434254 | 0.001470122 | 0.018947451 |
| CCT5 | -0.443631217 | 0.001471075 | 0.018947451 |
| CUNH19orf47 | 0.593069527 | 0.001481856 | 0.019067701 |
| MYOF | -0.584791941 | 0.001483975 | 0.019076374 |
| FKBP4 | -0.430110322 | 0.001486323 | 0.019087968 |
| TMEM140 | 0.555426273 | 0.001488751 | 0.01910057 |
| TOB2 | -0.724674029 | 0.001493543 | 0.019143445 |
| ENSACYG00000006681 | 4.220332326 | 0.001504733 | 0.019268171 |
| CCNB3 | -0.929278914 | 0.001510491 | 0.019323162 |
| LOC118260588 | -0.976398903 | 0.001512464 | 0.019329677 |
| CDCA7L | 0.475129756 | 0.001528638 | 0.019512654 |
| GMPPB | -0.618451541 | 0.001529738 | 0.019512654 |
| LOC118244251 | 3.075727747 | 0.001534327 | 0.019552308 |
| ENSACYG00000006881 | 1.394733211 | 0.001542794 | 0.019641244 |
| MTFR1 | 0.516873873 | 0.00154958 | 0.019708632 |
| CDC5L | -0.563847612 | 0.001555796 | 0.019768636 |
| PITPNM2 | 0.844803996 | 0.001561527 | 0.019815941 |
| CGREF1 | -0.699162781 | 0.00156252 | 0.019815941 |
| TUB | 0.530957739 | 0.001575342 | 0.019942166 |
| TMTC1 | 0.602500972 | 0.001575495 | 0.019942166 |
| LYAR | -0.645156875 | 0.001578895 | 0.019966068 |
| SRP72 | -0.422200198 | 0.001584847 | 0.020022158 |
| RGL1 | 0.62396604 | 0.001602349 | 0.020105082 |
| ING2 | -0.694668382 | 0.001601316 | 0.020105082 |
| RNF130 | 0.440493048 | 0.001593391 | 0.020105082 |
| DOT1L | 0.460579295 | 0.001597912 | 0.020105082 |
| BCL9L | 0.836744509 | 0.00159529 | 0.020105082 |
| RBSN | 0.63155016 | 0.001603594 | 0.020105082 |
| LOC118261520 | 2.246702969 | 0.001601776 | 0.020105082 |
| VPS45 | -0.489688549 | 0.001603304 | 0.020105082 |

| | | | |
|---|---|---|---|
| CCT4 | -0.474416463 | 0.001620979 | 0.020303764 |
| GUCY1A2 | 0.679693711 | 0.001622571 | 0.020304443 |
| DYNC2H1 | -0.672991391 | 0.001627311 | 0.020344464 |
| ARHGEF4 | -0.631114183 | 0.0016337 | 0.020405019 |
| ZNF362 | 0.685027062 | 0.001636946 | 0.020426233 |
| SRSF3 | -0.481004287 | 0.001641791 | 0.020448044 |
| NAT9 | -0.59373009 | 0.001641716 | 0.020448044 |
| LOC118250417 | -0.497463856 | 0.001654655 | 0.020588832 |
| ATP5PO | -0.552596778 | 0.001657868 | 0.020609394 |
| XXYLT1 | 0.481245434 | 0.001667143 | 0.020705193 |
| CCDC47 | -0.446670191 | 0.001670355 | 0.020725595 |
| EFNB2 | 0.743058419 | 0.001675051 | 0.020744858 |
| XPO6 | 0.774487504 | 0.001673541 | 0.020744858 |
| EXOSC2 | -0.503661162 | 0.00168141 | 0.020804098 |
| ENSACYG00000003391 | 0.572434942 | 0.001683006 | 0.020804349 |
| FAM102A | 0.621610786 | 0.001686598 | 0.020829249 |
| PTGDR | -0.940467404 | 0.001694783 | 0.020910769 |
| PIM1 | -0.86617884 | 0.00170164 | 0.020975773 |
| SMPD2 | -0.727786847 | 0.001711977 | 0.02108351 |
| TPMT | -0.598271672 | 0.001719023 | 0.021146842 |
| TTC30B | -0.674348831 | 0.001720482 | 0.021146842 |
| PAIP1 | -0.524855454 | 0.001721925 | 0.021146842 |
| ENSACYG00000015447 | 2.18396992 | 0.001726025 | 0.021177494 |
| DCTN1 | -0.902701166 | 0.001727959 | 0.021181533 |
| POMGNT1 | 0.678151928 | 0.001729941 | 0.021186158 |
| SQOR | -0.459125593 | 0.001741416 | 0.021306923 |
| LOC118244463 | -0.450181164 | 0.001756528 | 0.021432236 |
| GNAI1 | -0.455085135 | 0.001756478 | 0.021432236 |
| YWHAH | -0.445723612 | 0.001754337 | 0.021432236 |
| GPATCH4 | -0.753265059 | 0.001763708 | 0.021499976 |
| CCPG1 | -0.545155191 | 0.001765894 | 0.021506759 |
| LOC118251084 | -0.51211121 | 0.001772922 | 0.021552593 |
| GABRR2 | 4.823151033 | 0.001772226 | 0.021552593 |
| IPMK | 0.687912869 | 0.001781919 | 0.021622143 |
| ENSACYG00000004338 | 3.59354444 | 0.001780319 | 0.021622143 |
| TAP2 | -0.598311876 | 0.001784218 | 0.021630156 |
| HRAS | 0.462981549 | 0.001786464 | 0.021637526 |
| LYPD6 | 1.191671154 | 0.001793613 | 0.021688545 |
| MPP3 | -0.984216916 | 0.001793962 | 0.021688545 |
| EIF4E3 | 0.436490678 | 0.001800045 | 0.021742174 |
| GTF3C1 | -0.521147997 | 0.00181832 | 0.02194283 |
| REXO5 | -0.749187805 | 0.001823124 | 0.021980714 |
| TMEM185A | 0.530402189 | 0.001834442 | 0.022096995 |
| SPTAN1 | -0.610623039 | 0.001841378 | 0.02216032 |
| LOC118247154 | 2.176127352 | 0.001849933 | 0.022243003 |
| LOC118247056 | -0.454677893 | 0.001869596 | 0.022458974 |
| VPS37C | 0.535737051 | 0.001871421 | 0.02246046 |
| LOC118248278 | -0.703268108 | 0.00187742 | 0.022511987 |

| | | | |
|---|---|---|---|
| ERI2 | 0.77938153 | 0.00188327 | 0.022553041 |
| ADGRD1 | 0.560960627 | 0.00188426 | 0.022553041 |
| ENSACYG00000005690 | 4.239362558 | 0.001888949 | 0.022568381 |
| LOC118256513 | -0.551153767 | 0.001888961 | 0.022568381 |
| TRMT10C | -0.579602105 | 0.001893386 | 0.022580376 |
| ENSACYG00000004717 | 0.518513296 | 0.001892319 | 0.022580376 |
| PPWD1 | 0.451186499 | 0.001917332 | 0.022845325 |
| ENSACYG00000005147 | 3.697055032 | 0.001924729 | 0.022912777 |
| INTS12 | -0.806497676 | 0.001944501 | 0.023127296 |
| ENSACYG00000008470 | 1.94125499 | 0.001961558 | 0.023309169 |
| JDP2 | -0.653911334 | 0.001965965 | 0.023340533 |
| YLPM1 | -1.059698637 | 0.001968721 | 0.023352245 |
| LOC118243466 | 4.223950727 | 0.001976765 | 0.023426616 |
| KCNS1 | 3.473449114 | 0.001988225 | 0.023541295 |
| PRELID2 | 0.451797252 | 0.001994152 | 0.023590316 |
| MAP2K5 | 0.432034933 | 0.002007848 | 0.023716272 |
| IST1 | -0.497147954 | 0.002008392 | 0.023716272 |
| DUSP12 | -0.601519465 | 0.002016005 | 0.023784894 |
| MYNN | -0.73808236 | 0.002024363 | 0.023862174 |
| RPS6KA5 | -0.578437384 | 0.002029917 | 0.023887974 |
| FARP2 | 0.599450741 | 0.00203198 | 0.023887974 |
| TBC1D16 | 0.587574548 | 0.002030201 | 0.023887974 |
| MLST8 | -0.465893383 | 0.002033954 | 0.023889912 |
| LOC118255688 | 0.493490092 | 0.002044241 | 0.023989392 |
| LOC118252466 | 4.210819041 | 0.002050183 | 0.02402085 |
| NOP14 | -0.638575655 | 0.00205056 | 0.02402085 |
| RAB11FIP3 | 0.920208527 | 0.002054069 | 0.02404062 |
| FBRSL1 | 0.715951537 | 0.002070428 | 0.024210618 |
| HMMR | -0.572572039 | 0.00207413 | 0.024232447 |
| RNASEH2B | -0.492382129 | 0.002077112 | 0.024245828 |
| DNMBP | -0.754238414 | 0.002081189 | 0.024271959 |
| ENSACYG00000002918 | -1.316491878 | 0.002086502 | 0.024312449 |
| MARCHF1 | 0.691529704 | 0.002113753 | 0.024598417 |
| RNF114 | 0.440510041 | 0.002114771 | 0.024598417 |
| BAZ1A | -0.628352381 | 0.002124774 | 0.024673138 |
| PAQR5 | 0.538755969 | 0.002124932 | 0.024673138 |
| ENSACYG00000003333 | -0.721492378 | 0.002138169 | 0.024783236 |
| NAA35 | -0.436322454 | 0.002137417 | 0.024783236 |
| CLIP1 | -0.443165955 | 0.002142266 | 0.024808938 |
| NLN | -0.453610442 | 0.002157105 | 0.024937967 |
| LOC118258134 | 0.814721257 | 0.002157185 | 0.024937967 |
| PGM2 | -0.453328841 | 0.002168461 | 0.025046387 |
| METTL5 | -0.57018592 | 0.00217302 | 0.02507711 |
| HSP90AB1 | -1.02694157 | 0.002181783 | 0.025112374 |
| ENSACYG00000008861 | 0.594828984 | 0.002181319 | 0.025112374 |
| PEPD | -0.483062187 | 0.002181682 | 0.025112374 |
| PER2 | 0.698851352 | 0.002187712 | 0.025158683 |
| RAB19 | -0.753521095 | 0.002190226 | 0.025165674 |

| | | | |
|---|---|---|---|
| LOC118260427 | -0.531329125 | 0.002202105 | 0.025280171 |
| ENSACYG00000010584 | 0.703250963 | 0.002216078 | 0.025418476 |
| NFKBIE | 0.468855098 | 0.002222293 | 0.025445547 |
| ENSACYG00000009408 | 3.747696542 | 0.002221919 | 0.025445547 |
| DOP1B | -0.62507145 | 0.002242446 | 0.025654047 |
| SNX17 | 0.422562712 | 0.002265642 | 0.025896975 |
| PSMC2 | -0.476526356 | 0.002269319 | 0.025916564 |
| CIITA | 3.079775331 | 0.002277125 | 0.02594258 |
| PIGM | -0.641528379 | 0.002277492 | 0.02594258 |
| ACER3 | 0.516280553 | 0.002276061 | 0.02594258 |
| ENSACYG00000006849 | 0.89569441 | 0.002294478 | 0.026113538 |
| SMC4 | -0.52853723 | 0.002300133 | 0.026145385 |
| SRPRA | 0.466891031 | 0.002301238 | 0.026145385 |
| DAP3 | -0.4211491 | 0.002316014 | 0.026262768 |
| GJC2 | 0.674051611 | 0.002317537 | 0.026262768 |
| ENSACYG00000011468 | 1.031020489 | 0.002314972 | 0.026262768 |
| NUF2 | -0.496767155 | 0.00232785 | 0.026357012 |
| IRF2 | -0.502095538 | 0.002341638 | 0.026464297 |
| ROPN1L | -0.592011696 | 0.002343339 | 0.026464297 |
| KIAA0319L | 0.779809171 | 0.002340748 | 0.026464297 |
| ENSACYG00000003889 | 1.968696169 | 0.002346511 | 0.026465897 |
| LOC118261652 | -0.490617038 | 0.00234749 | 0.026465897 |
| LOC118258353 | -0.646472862 | 0.002350195 | 0.02647378 |
| LOC118259110 | -0.589243888 | 0.002356314 | 0.026520081 |
| CABIN1 | -0.508093404 | 0.002360236 | 0.026541595 |
| LOC118245372 | -2.144069584 | 0.002367983 | 0.026583433 |
| ENSACYG00000003588 | 0.685708671 | 0.002366046 | 0.026583433 |
| A1CF | 4.724344993 | 0.002372366 | 0.026588 |
| ENSACYG00000003384 | -1.213220566 | 0.002373838 | 0.026588 |
| GALNT17 | -4.829657379 | 0.00237531 | 0.026588 |
| EP300 | 0.595943323 | 0.002376446 | 0.026588 |
| FZD8 | -1.141093402 | 0.002384197 | 0.026645567 |
| RNF20 | -0.460541693 | 0.002385628 | 0.026645567 |
| CDCA4 | -0.496845114 | 0.002404265 | 0.026763441 |
| LSM1 | -0.547996174 | 0.002401933 | 0.026763441 |
| TMEM9 | -0.516959239 | 0.00240429 | 0.026763441 |
| ENSACYG00000014400 | -0.996284895 | 0.002399236 | 0.026763441 |
| MSH2 | -0.503860569 | 0.002407312 | 0.026774499 |
| DNAAF4 | -0.560719923 | 0.002409535 | 0.026776663 |
| GADD45G | -0.690363259 | 0.002413017 | 0.02679281 |
| RILP | -0.776988136 | 0.002415062 | 0.026792977 |
| ENSACYG00000011163 | -1.801120594 | 0.002422783 | 0.026833543 |
| ASPM | -0.741095513 | 0.002421002 | 0.026833543 |
| AHRR | 0.628790864 | 0.002429185 | 0.026881899 |
| ASCC3 | -0.523361586 | 0.002431429 | 0.026884192 |
| P3H3 | -0.712800903 | 0.002434 | 0.026890095 |
| B3GNT7 | -0.688574347 | 0.002446664 | 0.026983407 |
| MRPS35 | -0.545914937 | 0.002448578 | 0.026983407 |

| | | | |
|---|---|---|---|
| POLR3H | -0.519311373 | 0.002447953 | 0.026983407 |
| GDAP2 | 0.465806203 | 0.002464568 | 0.027120572 |
| MAP1A | -0.671317588 | 0.002465133 | 0.027120572 |
| MFSD10 | -0.511714686 | 0.002469422 | 0.02714514 |
| MGAT5 | 0.592547898 | 0.002482818 | 0.027269688 |
| GABPB1 | -0.541159719 | 0.002487909 | 0.027302893 |
| ACVR2A | 0.52633316 | 0.002506421 | 0.0274832 |
| PRRT1B | 0.443739084 | 0.002509337 | 0.02749234 |
| AP2A2 | -0.472744195 | 0.0025131 | 0.027510735 |
| POC5 | -0.483044617 | 0.002517353 | 0.027534461 |
| SEC14L1 | 0.459126449 | 0.002523906 | 0.027560464 |
| MFSD5 | 0.485728007 | 0.002523777 | 0.027560464 |
| TLE5 | 0.570109812 | 0.002526727 | 0.027568471 |
| TMEM11 | -0.587661913 | 0.002545553 | 0.02775094 |
| BEND5 | -0.996149171 | 0.002548572 | 0.027760931 |
| CDKN1C | -0.62599093 | 0.002556645 | 0.027825907 |
| LCT | 2.315859256 | 0.002579888 | 0.028032664 |
| UBB | -0.528754718 | 0.002579819 | 0.028032664 |
| GLB1 | 0.725064302 | 0.002598031 | 0.028206579 |
| RASSF6 | 0.468873125 | 0.00261779 | 0.028304723 |
| CHN1 | 0.431885548 | 0.002610165 | 0.028304723 |
| RTKN2 | 0.659327351 | 0.002614321 | 0.028304723 |
| BTD | -0.537605902 | 0.002616613 | 0.028304723 |
| SEC14L2 | -0.574625274 | 0.002613086 | 0.028304723 |
| YJU2 | -0.549334462 | 0.002620351 | 0.028309222 |
| TLCD2 | -0.612606026 | 0.002632092 | 0.02841282 |
| BABAM2 | 0.429144134 | 0.002642372 | 0.028500486 |
| CHUK | -0.950142564 | 0.002649258 | 0.028551433 |
| TTC4 | -0.494994385 | 0.002679731 | 0.028845523 |
| PPIL1 | -0.674859729 | 0.002680916 | 0.028845523 |
| CTNND2 | 0.616605467 | 0.002685208 | 0.028868174 |
| PCDH19 | 0.786659298 | 0.002693143 | 0.028929917 |
| TRIM46 | -1.662483176 | 0.00270365 | 0.02901918 |
| NUDT13 | -0.898610229 | 0.002719635 | 0.029167031 |
| CCDC80 | -0.4103546 | 0.002726038 | 0.029188282 |
| LOC118243257 | 0.446340565 | 0.002724494 | 0.029188282 |
| TMEM132B | 4.71770758 | 0.002739823 | 0.029288379 |
| LOC118244827 | 0.562834232 | 0.002738417 | 0.029288379 |
| SPRY3 | 0.583173274 | 0.002744074 | 0.029302573 |
| ATP6AP1 | -0.514001366 | 0.002747256 | 0.029302573 |
| CUNH16orf87 | -0.624463901 | 0.00274781 | 0.029302573 |
| ALG1 | -0.51833114 | 0.002790771 | 0.029736687 |
| SNX31 | 4.747299195 | 0.002815663 | 0.029879894 |
| ENSACYG00000001739 | 0.629370454 | 0.002817354 | 0.029879894 |
| AIDA | 0.431685117 | 0.00281429 | 0.029879894 |
| LOC118245898 | 3.05141426 | 0.002812675 | 0.029879894 |
| TTBK2 | 0.766971853 | 0.00281779 | 0.029879894 |
| SEPTIN9 | 0.781282056 | 0.002812699 | 0.029879894 |

| | | | |
|---|---|---|---|
| R3HCC1L | -0.777964293 | 0.00282426 | 0.029924465 |
| MDN1 | -0.594818695 | 0.002830865 | 0.029964687 |
| CDK11A | -0.535911225 | 0.002832596 | 0.029964687 |
| LOC118254088 | -0.843248715 | 0.00285091 | 0.030134277 |
| BMP4 | 0.545780996 | 0.002858766 | 0.030193142 |
| LOC118258128 | -0.619809181 | 0.002871413 | 0.030302477 |
| MVB12B | 0.55698371 | 0.00287603 | 0.030326955 |
| FLVCR2 | 0.56588621 | 0.002880421 | 0.030349014 |
| SHPRH | -0.61735175 | 0.002892027 | 0.030443199 |
| WNT5B | -0.538572805 | 0.002893972 | 0.030443199 |
| DMAC2L | -0.53899914 | 0.002906134 | 0.030546796 |
| B3GNT9 | 0.606238671 | 0.002926917 | 0.030740775 |
| GTF2H1 | 0.49311921 | 0.002939032 | 0.030843481 |
| GUSB | -0.437702326 | 0.002957083 | 0.031008268 |
| MRPS27 | -0.446979071 | 0.002965905 | 0.031076094 |
| UBN2 | 0.752709369 | 0.002971094 | 0.031105774 |
| ENSACYG00000012367 | 4.11691378 | 0.002978593 | 0.031159577 |
| IL11RA | 0.422201158 | 0.00298804 | 0.031233652 |
| FASTKD2 | -0.437787466 | 0.003009912 | 0.031425638 |
| NDUFAF5 | -0.589902828 | 0.003011167 | 0.031425638 |
| LIN52 | 0.47306508 | 0.003016419 | 0.031455578 |
| GRB10 | -0.499595125 | 0.00305032 | 0.031783995 |
| ARFGEF1 | -0.672566192 | 0.003052734 | 0.031784063 |
| RHOBTB2 | 0.732432725 | 0.003074834 | 0.031988937 |
| ENSACYG00000015280 | 0.674371432 | 0.003080538 | 0.032023047 |
| BTBD10 | -0.411462682 | 0.003083593 | 0.032029583 |
| TSGA10 | 2.965588391 | 0.003096215 | 0.032135406 |
| MEX3B | -0.588032017 | 0.003100902 | 0.032158764 |
| ENSACYG00000007005 | 1.746071542 | 0.003109655 | 0.032224223 |
| TIFA | -0.694516582 | 0.003126053 | 0.032368745 |
| PSPH | -0.63872403 | 0.003157961 | 0.032622379 |
| NBAS | -0.536548652 | 0.003153116 | 0.032622379 |
| RDM1 | -0.513138991 | 0.003156637 | 0.032622379 |
| OAF | 0.5330868 | 0.003179631 | 0.032820551 |
| DHX29 | -0.597195991 | 0.003186753 | 0.032868372 |
| BCAN | -0.598492503 | 0.003213181 | 0.033115078 |
| LOC118243167 | -0.566123823 | 0.003221778 | 0.033177783 |
| HIVEP3 | 0.580588115 | 0.003225326 | 0.033188432 |
| TMEM177 | -0.485111149 | 0.003232857 | 0.033216828 |
| ENSACYG00000008044 | -0.706485063 | 0.003233118 | 0.033216828 |
| RRS1 | -0.786882156 | 0.003249699 | 0.033353792 |
| NDUFA11 | -0.612170042 | 0.003251502 | 0.033353792 |
| IER5 | -1.111728351 | 0.003254836 | 0.033362071 |
| PDZD8 | 0.409753893 | 0.003264878 | 0.033439038 |
| PKIA | 0.461963098 | 0.003276958 | 0.033536746 |
| CPD | -0.555431102 | 0.003281705 | 0.033559306 |
| EFL1 | -0.474559477 | 0.003307519 | 0.033707143 |
| BMS1 | -0.545700121 | 0.003304672 | 0.033707143 |

| | | | |
|---|---|---|---|
| KIF20B | -0.514749291 | 0.003301666 | 0.033707143 |
| CCNJ | -0.601384146 | 0.003308927 | 0.033707143 |
| ENSACYG00000014743 | 0.653998516 | 0.003305789 | 0.033707143 |
| ADAR | 0.699951422 | 0.003316347 | 0.033726555 |
| LYSMD4 | -0.594305977 | 0.003315064 | 0.033726555 |
| LOC118248016 | -0.433734576 | 0.003318497 | 0.033726555 |
| CRTAP | -0.433258871 | 0.003328683 | 0.033804054 |
| LRRC36 | 0.802869713 | 0.003337203 | 0.033864532 |
| PDE4B | 0.492096972 | 0.003345776 | 0.033925448 |
| NTNG2 | 0.731466783 | 0.003362493 | 0.034068789 |
| EPB41L4A | 0.535030683 | 0.003367475 | 0.034093106 |
| SYNCRIP | -0.393622498 | 0.003380797 | 0.034201753 |
| ENSACYG00000011174 | 0.51933285 | 0.003390837 | 0.034277057 |
| STRADB | 0.551322706 | 0.003425815 | 0.034474662 |
| ANGPT1 | -0.419498364 | 0.003431276 | 0.034474662 |
| LOC118244586 | 3.643991691 | 0.003430639 | 0.034474662 |
| NIP7 | -0.52908857 | 0.003422264 | 0.034474662 |
| LOC118252907 | -0.575404973 | 0.003425585 | 0.034474662 |
| GFM1 | -0.443084936 | 0.003424548 | 0.034474662 |
| IGSF9B | 0.615956252 | 0.003428861 | 0.034474662 |
| TBCCD1 | -0.525421199 | 0.003422905 | 0.034474662 |
| SP1 | 0.770422348 | 0.003441291 | 0.034548992 |
| SLC2A8 | -0.428272734 | 0.00344749 | 0.034584921 |
| COLQ | 3.004538294 | 0.003455662 | 0.034640585 |
| POLH | -0.703259929 | 0.003467473 | 0.034706274 |
| SLAIN2 | 0.618562729 | 0.003465107 | 0.034706274 |
| ENSACYG00000005083 | 0.548632123 | 0.003470206 | 0.034707317 |
| TNFRSF21 | 0.648852897 | 0.003494159 | 0.034920428 |
| LOC118259705 | 0.528930413 | 0.003514181 | 0.035067999 |
| KIF4A | -0.50200867 | 0.003514237 | 0.035067999 |
| THOC2 | -0.429982087 | 0.003518922 | 0.035088227 |
| TSC22D2 | 0.569488226 | 0.003525709 | 0.035129364 |
| APRT | -0.582871783 | 0.003542855 | 0.035273582 |
| RPL24 | -0.501432297 | 0.003602635 | 0.035814752 |
| TMPRSS4 | -1.028776944 | 0.003600317 | 0.035814752 |
| GABPA | 0.608959283 | 0.003606439 | 0.035825593 |
| ENSACYG00000012379 | 1.065691939 | 0.003618121 | 0.035912092 |
| ENSACYG00000013189 | 4.034169463 | 0.003620587 | 0.035912092 |
| ARHGAP17 | 0.475202082 | 0.003628682 | 0.03596536 |
| SMAD4 | 0.45725768 | 0.003644279 | 0.036092856 |
| IL16 | 1.024760421 | 0.003648804 | 0.036110579 |
| TAGLN3 | 3.605864 | 0.003675701 | 0.03634952 |
| BTC | -0.890177876 | 0.00368308 | 0.036395223 |
| ENSACYG00000016135 | -0.771865836 | 0.003720061 | 0.036733171 |
| TET2 | 0.612403657 | 0.003731254 | 0.03681616 |
| CANT1 | -0.503382955 | 0.003734325 | 0.036818935 |
| ENSACYG00000009875 | 0.948715266 | 0.003738462 | 0.036832222 |
| WDR26 | -0.44578751 | 0.003749145 | 0.036909928 |

| | | | |
|---|---|---|---|
| PRRC2C | -0.621110794 | 0.00376683 | 0.037045683 |
| HAT1 | -0.458269684 | 0.003768547 | 0.037045683 |
| CBLB | -0.468947438 | 0.003787259 | 0.037174271 |
| TTC29 | 1.024764478 | 0.003786253 | 0.037174271 |
| SLC35B4 | 0.517694567 | 0.003795021 | 0.037222782 |
| FAM193A | -0.635830514 | 0.003801207 | 0.037255782 |
| CEP164 | 0.579180254 | 0.003809846 | 0.037285091 |
| LOC118255703 | -0.495325533 | 0.003807275 | 0.037285091 |
| GEMIN2 | -0.497288738 | 0.003813023 | 0.037288543 |
| MTHFD2 | -0.41936312 | 0.003823068 | 0.037359095 |
| C1S | 0.791137474 | 0.00383404 | 0.037438608 |
| DSE | 0.568025253 | 0.003836974 | 0.037439563 |
| ABI1 | 0.403984102 | 0.003856499 | 0.037586344 |
| ENSACYG00000016566 | 0.54418589 | 0.003857711 | 0.037586344 |
| MXD1 | 0.457164841 | 0.003868679 | 0.037665417 |
| ARHGEF10 | -0.486555265 | 0.003882575 | 0.037717257 |
| MRPL22 | -0.545785158 | 0.003878145 | 0.037717257 |
| SNAI2 | -0.69348292 | 0.003880234 | 0.037717257 |
| ASL | -0.460030229 | 0.003888741 | 0.037721643 |
| XYLB | -0.5769652 | 0.003885945 | 0.037721643 |
| PCBD2 | 0.922355446 | 0.003905379 | 0.037855221 |
| IFT20 | -0.506053315 | 0.0039256 | 0.037995432 |
| GATA2 | 1.673944349 | 0.003923621 | 0.037995432 |
| PTGER2 | -0.694159918 | 0.00393091 | 0.038015783 |
| TUBB6 | -0.397755586 | 0.003933462 | 0.038015783 |
| SLC7A3 | -0.547727756 | 0.003938599 | 0.038021637 |
| NFE2L1 | 0.656232986 | 0.003939827 | 0.038021637 |
| STK26 | 0.479559169 | 0.003943888 | 0.038033027 |
| ASF1A | -0.534110291 | 0.003963883 | 0.03817032 |
| CUNH18orf21 | -0.526821971 | 0.003963908 | 0.03817032 |
| CACTIN | 0.4801574 | 0.004002538 | 0.038514221 |
| SFT2D2 | 0.655865856 | 0.004016923 | 0.038596375 |
| MSRB3 | 0.525991193 | 0.004014766 | 0.038596375 |
| SNN | -0.62814102 | 0.004030125 | 0.038672768 |
| HTRA1 | -0.400719991 | 0.004030732 | 0.038672768 |
| FAM76B | -0.506784193 | 0.00405051 | 0.038834298 |
| SEPTIN6 | -0.421920845 | 0.004057592 | 0.038873968 |
| DDX46 | -0.476494647 | 0.004070785 | 0.038972081 |
| HS2ST1 | 0.492097288 | 0.004081613 | 0.039047435 |
| NUDT16L1 | -0.482509229 | 0.004095437 | 0.039151306 |
| OTOGL | 4.02520838 | 0.00410608 | 0.039224652 |
| TFCP2L1 | 1.771698782 | 0.004116215 | 0.039264643 |
| ENSACYG00000006221 | -0.713405293 | 0.004115661 | 0.039264643 |
| ENSACYG00000009714 | 0.530867511 | 0.004123589 | 0.039306582 |
| LOC118254226 | 2.900319087 | 0.0041295 | 0.039334527 |
| LOC118260146 | 0.573537479 | 0.004154002 | 0.039539386 |
| PHF14 | -0.555455252 | 0.004158816 | 0.039556692 |
| GLS | -0.378234443 | 0.004161927 | 0.039557786 |

| | | | |
|---|---|---|---|
| LOC118248563 | -0.534022002 | 0.004174043 | 0.03961886 |
| ARHGEF3 | -0.456470305 | 0.004174355 | 0.03961886 |
| CUNH10orf71 | 0.4613188 | 0.004191341 | 0.039740674 |
| ZBTB26 | 0.44972556 | 0.00419321 | 0.039740674 |
| CALCRL | -0.75251068 | 0.004198923 | 0.039766271 |
| IRF2BP2 | -0.569143908 | 0.004211376 | 0.039855616 |
| LOC118244275 | -0.541337798 | 0.004215503 | 0.039866096 |
| ARHGDIA | 0.390452508 | 0.004219199 | 0.039872486 |
| SMNDC1 | -0.490982201 | 0.004246732 | 0.040075312 |
| ENSACYG00000012312 | 1.942318719 | 0.004245871 | 0.040075312 |
| BRICD5 | 1.3217615 | 0.0042638 | 0.040207632 |
| ADCY5 | 0.711814533 | 0.004293713 | 0.040460811 |
| SPAG9 | -0.460032336 | 0.004323278 | 0.040710352 |
| VPS13A | -0.418922874 | 0.004327907 | 0.040724896 |
| DBNDD1 | -0.706077194 | 0.004342626 | 0.040834295 |
| KAZN | 0.643169856 | 0.004349536 | 0.040870157 |
| LOC118257323 | 3.995912588 | 0.004358149 | 0.040921965 |
| LOC118243739 | 3.166003986 | 0.004372442 | 0.041026995 |
| HHIPL1 | 0.58801319 | 0.004386071 | 0.041125649 |
| LOC118245540 | 0.394310234 | 0.004419099 | 0.041405923 |
| MAN1A2 | 0.56682443 | 0.004423021 | 0.04141328 |
| NECAP2 | 0.399503656 | 0.004426546 | 0.041416913 |
| ENSACYG00000007646 | 4.533112181 | 0.004431529 | 0.041434168 |
| WDR3 | -0.409311907 | 0.004437433 | 0.041460011 |
| KIFBP | -0.414721102 | 0.00449904 | 0.042005888 |
| CILK1 | 0.643448313 | 0.00450421 | 0.042024437 |
| LOC118248104 | -0.675823147 | 0.004513947 | 0.042085546 |
| COMMD1 | -0.650955728 | 0.004521417 | 0.042125441 |
| PML | 0.629748145 | 0.004531235 | 0.042187143 |
| GCNT3 | 4.094882459 | 0.004535107 | 0.042193431 |
| ENSACYG00000001024 | 3.504301367 | 0.004547339 | 0.042277446 |
| UBA6 | -0.647417815 | 0.004610377 | 0.042833357 |
| ARHGEF6 | 0.438667833 | 0.004622584 | 0.042916567 |
| MAP1B | -0.522580126 | 0.004646117 | 0.043044238 |
| TRIP13 | -0.557544453 | 0.00464461 | 0.043044238 |
| FICD | 0.608624723 | 0.004641509 | 0.043044238 |
| ENSACYG00000003157 | 3.505837685 | 0.004659247 | 0.043135607 |
| SMC5 | -0.55292884 | 0.004680954 | 0.043306201 |
| NTAQ1 | -0.520986136 | 0.004694911 | 0.043404913 |
| ENSACYG00000003901 | 0.51457165 | 0.004732782 | 0.043632814 |
| TMEM71 | 0.519250502 | 0.004727048 | 0.043632814 |
| PRKD3 | 0.717965683 | 0.004730361 | 0.043632814 |
| STC1 | -0.608719012 | 0.004723069 | 0.043632814 |
| CRAMP1 | 0.73383453 | 0.004737673 | 0.04364466 |
| RBPMS2 | 0.414151951 | 0.004740679 | 0.04364466 |
| SLC25A33 | -0.568550848 | 0.004748584 | 0.043686972 |
| STRIP2 | 1.233908712 | 0.004765601 | 0.043812999 |
| ARL11 | -0.473621325 | 0.004770123 | 0.043824052 |

| | | | |
|---|---|---|---|
| ASPG | -0.449836003 | 0.004788308 | 0.043960532 |
| PDSS1 | -0.431607151 | 0.004796721 | 0.044007163 |
| SERINC5 | 0.577041568 | 0.004805778 | 0.04402906 |
| FKBP7 | -0.502048442 | 0.004804088 | 0.04402906 |
| DDX20 | -0.433497923 | 0.004816246 | 0.044094367 |
| LOC118251281 | 0.479854113 | 0.004822105 | 0.044117414 |
| ADAM33 | -0.621777613 | 0.004840695 | 0.044256822 |
| ACAP3 | 0.531397651 | 0.00485635 | 0.04436923 |
| ENSACYG00000003677 | -0.810729603 | 0.004874883 | 0.044507745 |
| LOC118250227 | -0.476150142 | 0.004885729 | 0.044570314 |
| ZDHHC13 | -0.453318288 | 0.004890545 | 0.044570314 |
| CHRM4 | 0.868520793 | 0.004891864 | 0.044570314 |
| MSTO1 | -0.478796363 | 0.004900174 | 0.044615238 |
| TMEM184B | 0.504154989 | 0.004942864 | 0.044972909 |
| RND3 | -0.452629972 | 0.004955251 | 0.045054564 |
| TRMT2B | -0.75826861 | 0.004971078 | 0.045166262 |
| SDHAF2 | -0.511270232 | 0.004974379 | 0.045166262 |
| CCT3 | -0.407782374 | 0.004980515 | 0.045190901 |
| MRPS12 | -0.561630709 | 0.005000545 | 0.045310365 |
| STK25 | 0.716929535 | 0.00499734 | 0.045310365 |
| RNF38 | 0.473648299 | 0.005023232 | 0.045484714 |
| ENSACYG00000002590 | 0.608090444 | 0.005043886 | 0.045588082 |
| LHX6 | 0.701790381 | 0.00504846 | 0.045588082 |
| BRIX1 | -0.500444472 | 0.00504625 | 0.045588082 |
| HDLBP | 0.649613732 | 0.005045611 | 0.045588082 |
| EBI3 | -0.900003867 | 0.005060339 | 0.045638047 |
| YARS1 | -0.419808622 | 0.005060908 | 0.045638047 |
| STOML3 | 2.284000211 | 0.005070129 | 0.045689999 |
| TTC1 | -0.431858394 | 0.005079412 | 0.04572232 |
| RMND5A | -0.465481869 | 0.005080643 | 0.04572232 |
| ENSACYG00000016889 | 1.385280192 | 0.005087366 | 0.045751637 |
| LOC118247600 | -0.421373191 | 0.005093952 | 0.045779679 |
| ITGA2 | -0.532453304 | 0.005101723 | 0.045818331 |
| FRMD3 | 0.523610413 | 0.005115524 | 0.045911042 |
| DENND10 | -0.410752331 | 0.005128358 | 0.045970925 |
| RNF128 | -0.415227869 | 0.00512916 | 0.045970925 |
| TP53INP2 | 0.654749637 | 0.005143008 | 0.04606377 |
| MRPL18 | -0.541547207 | 0.005156703 | 0.046142161 |
| VILL | 1.351912074 | 0.005158751 | 0.046142161 |
| ING4 | -0.468566923 | 0.005170675 | 0.046193467 |
| SLC25A16 | 0.448996485 | 0.005171485 | 0.046193467 |
| FZD9 | 0.769726843 | 0.005191126 | 0.046337556 |
| UVSSA | -0.641923119 | 0.005247535 | 0.046809433 |
| DCAF10 | 0.461000537 | 0.005257112 | 0.046831576 |
| ING3 | -0.436673421 | 0.005254393 | 0.046831576 |
| LFNG | -0.568255199 | 0.005263247 | 0.046854607 |
| ABCB10 | 0.475176408 | 0.005297361 | 0.047126524 |
| IK | -0.443953154 | 0.005325795 | 0.047347568 |

| | | | |
|---|---|---|---|
| TCP1 | -0.423135297 | 0.005358039 | 0.047602173 |
| CASP2 | -0.891848103 | 0.005378891 | 0.047755292 |
| EVC | -0.638222319 | 0.005395798 | 0.0478732 |
| LRP1 | -0.465891386 | 0.00540966 | 0.04793177 |
| WDR73 | 0.446956865 | 0.005408197 | 0.04793177 |
| FADD | -0.496532813 | 0.005430514 | 0.048019861 |
| SHANK2 | 1.392798027 | 0.005428893 | 0.048019861 |
| RALBP1 | 0.394699309 | 0.005426002 | 0.048019861 |
| PHPT1 | -0.497556828 | 0.005459459 | 0.04824349 |
| LOC118245392 | 2.739148259 | 0.005465569 | 0.04826518 |
| RAE1 | -0.400849549 | 0.005474166 | 0.04830878 |
| ST6GAL1 | 0.608988859 | 0.005493606 | 0.048447954 |
| NOX3 | 0.826591953 | 0.005525765 | 0.048699027 |
| GPSM1 | 0.67425096 | 0.005533491 | 0.048734588 |
| CUNH5orf34 | -0.562944757 | 0.005576671 | 0.04908214 |
| MRPS11 | -0.507947093 | 0.005582869 | 0.049103959 |
| MRPL50 | -0.537728933 | 0.005586819 | 0.049105978 |
| ENSACYG00000003044 | 1.604343674 | 0.005600462 | 0.049160439 |
| PTCD3 | -0.504793543 | 0.005598608 | 0.049160439 |
| ENSACYG00000004263 | 1.867911119 | 0.005619215 | 0.049194216 |
| CAV1 | 0.484602933 | 0.005618444 | 0.049194216 |
| GCC2 | -0.419029424 | 0.005614095 | 0.049194216 |
| ZHX3 | 0.822495572 | 0.005608717 | 0.049194216 |
| PRDM4 | 0.476235781 | 0.005627445 | 0.049233617 |
| LOC118251891 | 0.421819219 | 0.005636166 | 0.049277262 |
| ZBTB6 | -0.86554822 | 0.005646554 | 0.04933541 |
| PDP1 | -0.621875607 | 0.005657561 | 0.049398888 |
| SRSF11 | -0.487989576 | 0.005672106 | 0.049493158 |
| TRAPPC2L | -0.507521602 | 0.005677196 | 0.049501358 |
| TMEM200C | 0.926736825 | 0.005680545 | 0.049501358 |
| RAB11FIP2 | 0.422412125 | 0.005698855 | 0.049593797 |
| ANKRD34B | 2.647281542 | 0.005699752 | 0.049593797 |
| CALB1 | 3.498203167 | 0.005702423 | 0.049593797 |
| ELOVL4 | -0.502044588 | 0.005726873 | 0.049773651 |
| LOC118249154 | 4.048936612 | 0.005735949 | 0.049819737 |
| ENSACYG00000002353 | -4.695020176 | 0.005751406 | 0.049895575 |
| ENSACYG00000005196 | -2.277470068 | 0.005776086 | 0.049895575 |
| FOXP1 | 0.431556238 | 0.005775097 | 0.049895575 |
| TUBGCP5 | -0.749777241 | 0.005762284 | 0.049895575 |
| LOC118258018 | -0.988384832 | 0.005760636 | 0.049895575 |
| FOXP4 | 0.589574223 | 0.005778695 | 0.049895575 |
| CEP19 | 0.482294511 | 0.005764385 | 0.049895575 |
| CHST11 | 1.506771891 | 0.005778427 | 0.049895575 |
| ESD | -0.514770258 | 0.005775315 | 0.049895575 |

**Supplementary Table 7: Differentially expressed genes in infected chicken swan endothelial cells**

| Gene | log2FoldChange | pvalue | padj |
|---|---|---|---|
| 05-Mar | -0.379765253 | 3.37E-05 | 0.000100296 |
| 08-Mar | 0.382333805 | 0.002251563 | 0.00505421 |
| 07-Sep | 0.661687094 | 4.99E-20 | 5.63E-19 |
| A4GALT | 2.323574318 | 0.00658573 | 0.01343236 |
| AACS | -1.640595921 | 3.63E-51 | 2.26E-49 |
| AADAC | -0.562717706 | 0.000116122 | 0.000319306 |
| AADAT | 1.095069165 | 5.36E-05 | 0.000155691 |
| AAK1 | 0.546650106 | 0.002156035 | 0.00485984 |
| AAMDC | 0.858629219 | 7.59E-13 | 5.01E-12 |
| AAMP | -0.365513848 | 7.41E-05 | 0.000210679 |
| AAR2 | 0.600694208 | 4.40E-10 | 2.24E-09 |
| AARS | -1.242947971 | 6.91E-49 | 4.00E-47 |
| AARS2 | -0.809262479 | 5.64E-06 | 1.86E-05 |
| AARSD1 | -0.927848376 | 3.88E-15 | 3.07E-14 |
| AASDH | 0.517896505 | 0.004548804 | 0.009590096 |
| AASDHPPT | -0.378673393 | 0.006990645 | 0.014173158 |
| AATF | -0.966146074 | 6.09E-11 | 3.36E-10 |
| ABCA12 | 5.417768261 | 1.21E-60 | 1.08E-58 |
| ABCA3 | -0.539782389 | 4.65E-07 | 1.76E-06 |
| ABCA4 | 1.538035167 | 0.002181352 | 0.004912831 |
| ABCA5 | -1.223128309 | 0.00470893 | 0.00990924 |
| ABCB1 | 1.316959519 | 0.001288206 | 0.003018843 |
| ABCB1.1 | -0.249310361 | 0.0086432 | 0.017142619 |
| ABCB11 | -1.487421278 | 2.26E-06 | 7.88E-06 |
| ABCB6 | -2.068871335 | 4.56E-34 | 1.20E-32 |
| ABCB8 | -0.646225934 | 0.000356019 | 0.000912848 |
| ABCB9 | -1.892172882 | 1.54E-06 | 5.49E-06 |
| ABCC1 | -0.734509379 | 3.88E-19 | 4.13E-18 |
| ABCC4 | -0.685616956 | 5.48E-05 | 0.00015902 |
| ABCC6 | 0.73675382 | 0.025351502 | 0.045339161 |
| ABCD2 | -0.669994234 | 1.33E-08 | 5.90E-08 |
| ABCD3 | -0.56771562 | 2.18E-08 | 9.49E-08 |
| ABCD4 | -0.415579847 | 0.000900559 | 0.002164144 |
| ABCE1 | -0.553886769 | 1.20E-07 | 4.84E-07 |
| ABCG2 | -1.379643602 | 7.28E-09 | 3.31E-08 |
| ABHD10 | -1.069059929 | 3.08E-06 | 1.06E-05 |
| ABHD12 | -0.319334147 | 0.00710313 | 0.014369064 |
| ABHD13 | 1.110171482 | 5.81E-35 | 1.61E-33 |
| ABHD14B | 1.146641263 | 2.31E-18 | 2.35E-17 |
| ABHD17A | 1.458201506 | 1.75E-63 | 1.78E-61 |
| ABHD17B | -1.182321062 | 2.34E-27 | 4.25E-26 |
| ABHD18 | 0.576020911 | 6.74E-07 | 2.51E-06 |
| ABHD2 | -0.352100898 | 0.017305699 | 0.032182112 |
| ABHD3 | 1.391723423 | 6.13E-09 | 2.80E-08 |
| ABHD4 | 0.844235307 | 2.20E-16 | 1.93E-15 |

| | | | |
|---|---|---|---|
| ABHD6 | 1.879265423 | 1.70E-52 | 1.12E-50 |
| ABI1 | 0.536082842 | 7.24E-05 | 0.000206009 |
| ABI2 | 0.600866278 | 0.010000672 | 0.019583509 |
| ABL2 | -1.464263224 | 1.45E-06 | 5.20E-06 |
| ABLIM2 | 1.170184839 | 1.92E-30 | 4.14E-29 |
| ABLIM3 | -1.750067693 | 0.013809973 | 0.02625247 |
| ABR | -0.451726453 | 0.000392571 | 0.001000154 |
| ABRA | -2.58896173 | 4.82E-05 | 0.000140956 |
| ABRACL | 0.278573997 | 0.010626724 | 0.020708018 |
| ABTB1 | 0.552608806 | 1.24E-08 | 5.53E-08 |
| ACAA1 | -1.573617008 | 1.24E-38 | 4.21E-37 |
| ACAA2 | -0.333236058 | 0.004816317 | 0.010103932 |
| ACAD11 | -1.118997999 | 5.22E-15 | 4.08E-14 |
| ACADL | -0.414243984 | 0.000393555 | 0.001002474 |
| ACADS | -0.6422367 | 9.41E-10 | 4.65E-09 |
| ACAP2 | -1.356943863 | 5.75E-35 | 1.59E-33 |
| ACAT1 | -0.724086548 | 7.23E-15 | 5.60E-14 |
| ACAT2 | -0.705113312 | 2.70E-11 | 1.55E-10 |
| ACBD3 | -0.987249429 | 1.21E-29 | 2.50E-28 |
| ACBD6 | 1.126735678 | 3.57E-29 | 7.17E-28 |
| ACBD7 | 0.922551391 | 1.88E-10 | 9.92E-10 |
| ACE | 3.767884996 | 0.000433412 | 0.001096805 |
| ACE2 | 2.924355086 | 0.007044395 | 0.014269362 |
| ACER3 | 0.600832623 | 5.26E-07 | 1.98E-06 |
| ACKR2 | -0.810184627 | 1.32E-07 | 5.31E-07 |
| ACLY | -1.225601853 | 2.10E-35 | 5.96E-34 |
| ACO1 | -1.420594899 | 1.38E-27 | 2.52E-26 |
| ACO2 | -0.780714189 | 2.84E-17 | 2.67E-16 |
| ACOT13 | 0.250270446 | 0.021942165 | 0.039855393 |
| ACOT7 | 0.674604837 | 5.51E-16 | 4.66E-15 |
| ACOT8 | -0.347630778 | 0.002365147 | 0.005296931 |
| ACOT9 | 0.538096211 | 1.17E-05 | 3.70E-05 |
| ACOX1 | -0.691797851 | 9.98E-11 | 5.40E-10 |
| ACOX3 | -0.811863278 | 7.63E-08 | 3.15E-07 |
| ACP1 | 0.249536298 | 0.024007992 | 0.043237927 |
| ACSBG2 | -1.200267201 | 3.68E-28 | 6.97E-27 |
| ACSL1 | -0.600822405 | 8.27E-11 | 4.51E-10 |
| ACSL3 | -0.87825964 | 3.53E-17 | 3.29E-16 |
| ACSL4 | -1.539291912 | 4.03E-46 | 2.04E-44 |
| ACSS2 | -0.736036867 | 5.30E-08 | 2.22E-07 |
| ACTA2 | 0.375827666 | 0.00056966 | 0.001413181 |
| ACTB | 0.254934929 | 0.009639326 | 0.018941408 |
| ACTC1 | 0.764071502 | 3.46E-07 | 1.33E-06 |
| ACTG2 | 0.42562429 | 0.001901064 | 0.004323116 |
| ACTL6A | -0.400156094 | 3.15E-05 | 9.42E-05 |
| ACTN4 | -0.338421916 | 0.007910414 | 0.015858164 |
| ACTR10 | -0.622388761 | 6.36E-11 | 3.50E-10 |
| ACTR1A | -0.400952264 | 3.05E-06 | 1.05E-05 |

| | | | |
|---|---|---|---|
| ACTR2 | 0.551394532 | 5.24E-12 | 3.20E-11 |
| ACTR5 | -0.855714905 | 4.24E-05 | 0.000124899 |
| ACTR6 | 1.010533211 | 5.27E-29 | 1.05E-27 |
| ACTR8 | 0.511706106 | 8.19E-09 | 3.71E-08 |
| ACVR1 | 0.907841033 | 5.88E-17 | 5.36E-16 |
| ACVR2A | 0.3461553 | 0.000358272 | 0.000918453 |
| ACYP1 | -0.409038635 | 0.022430221 | 0.040633288 |
| ACYP2 | 1.049276955 | 1.60E-11 | 9.35E-11 |
| ADA | 0.587525611 | 0.010804013 | 0.021017338 |
| ADAL | -0.65710331 | 3.23E-06 | 1.10E-05 |
| ADAM10 | -0.407862847 | 5.11E-07 | 1.93E-06 |
| ADAM12 | -0.669361116 | 1.68E-08 | 7.39E-08 |
| ADAM17 | 0.421215097 | 9.53E-05 | 0.000266074 |
| ADAM20 | 1.798004421 | 0.000368505 | 0.000943261 |
| ADAM20.1 | 3.776483733 | 1.95E-08 | 8.53E-08 |
| ADAM33 | -0.218280868 | 0.012507344 | 0.024058847 |
| ADAM8 | -0.541299129 | 1.70E-06 | 6.05E-06 |
| ADAM9 | -0.842032397 | 1.10E-15 | 9.07E-15 |
| ADAMTS1 | -1.436066432 | 1.24E-37 | 4.07E-36 |
| ADAMTS12 | 0.433671043 | 2.62E-05 | 7.91E-05 |
| ADAMTS14 | -0.457523617 | 0.00810116 | 0.016197553 |
| ADAMTS15 | 2.334889203 | 0.001442199 | 0.003354272 |
| ADAMTS3 | -0.897570567 | 0.000143089 | 0.000388121 |
| ADAMTS4 | -1.024147736 | 7.79E-14 | 5.57E-13 |
| ADAMTS6 | 0.336071261 | 0.014897848 | 0.028116016 |
| ADAMTSL1 | 2.572555153 | 0.008578203 | 0.01703857 |
| ADAMTSL2 | -0.884992883 | 1.33E-17 | 1.28E-16 |
| ADAT1 | 0.420953822 | 0.007953098 | 0.015924942 |
| ADCK1 | -1.557578173 | 8.39E-12 | 5.05E-11 |
| ADCK2 | -0.858458193 | 0.000196601 | 0.000523447 |
| ADD1 | -0.702216627 | 3.69E-14 | 2.72E-13 |
| ADGRA2 | 1.430353812 | 0.000612011 | 0.001510523 |
| ADGRA3 | -0.458874348 | 0.005171645 | 0.010781135 |
| ADGRD1 | 0.676112535 | 0.00999624 | 0.019579808 |
| ADGRG1 | -0.414295323 | 0.004494617 | 0.009487628 |
| ADGRG7 | 1.63187495 | 2.04E-10 | 1.07E-09 |
| ADGRL2 | -1.324738687 | 2.70E-12 | 1.70E-11 |
| ADH1C | 1.302627467 | 0.019412915 | 0.035714718 |
| ADH4 | -0.507738568 | 8.78E-07 | 3.23E-06 |
| ADI1 | 1.177501302 | 8.39E-21 | 9.87E-20 |
| ADIPOR1 | 0.30725307 | 0.000785336 | 0.001904761 |
| ADIPOR2 | 0.378491807 | 0.001901717 | 0.004323878 |
| ADNP2 | -0.659380373 | 0.000654524 | 0.00160699 |
| ADORA2B | 1.163488969 | 1.30E-15 | 1.06E-14 |
| ADPGK | -0.43887706 | 5.22E-07 | 1.97E-06 |
| ADPRM | -1.293892548 | 1.17E-13 | 8.28E-13 |
| ADRA1D | -4.634825348 | 0.001954914 | 0.004434444 |
| ADRM1 | 0.323661904 | 7.28E-05 | 0.00020711 |

| | | | |
|---|---|---|---|
| ADSL | -0.290673839 | 0.011245446 | 0.021788752 |
| ADTRP | 1.000075755 | 0.000241713 | 0.000636329 |
| AEBP1 | -1.183951318 | 2.57E-43 | 1.11E-41 |
| AEBP2 | -0.70241136 | 4.75E-11 | 2.64E-10 |
| AFAP1 | -0.580347877 | 4.23E-09 | 1.96E-08 |
| AFDN | -1.180735805 | 4.07E-11 | 2.29E-10 |
| AFF2 | 0.690756119 | 0.001980314 | 0.004490562 |
| AFF4 | -0.517045706 | 0.000453474 | 0.001143529 |
| AFG3L2 | -0.971745087 | 7.28E-31 | 1.60E-29 |
| AFMID | 0.384624501 | 0.00423414 | 0.008997879 |
| AFTPH | -1.425417583 | 1.36E-22 | 1.81E-21 |
| AGAP1 | 0.743670772 | 5.69E-05 | 0.000164614 |
| AGAP3 | 0.22613641 | 0.013361802 | 0.025514697 |
| AGBL5 | -1.191009796 | 1.15E-05 | 3.64E-05 |
| AGL | -0.929015814 | 1.83E-07 | 7.24E-07 |
| AGMAT | 2.029915337 | 0.001805984 | 0.004127615 |
| AGO2 | -1.395696898 | 0.000173523 | 0.000465376 |
| AGO3 | 0.835898755 | 7.40E-22 | 9.30E-21 |
| AGO4 | 0.423868335 | 9.58E-05 | 0.000267448 |
| AGPAT2 | 0.593282693 | 1.96E-09 | 9.41E-09 |
| AGPAT3 | 0.40254349 | 0.004347626 | 0.009205941 |
| AGPAT4 | -1.233741756 | 2.70E-16 | 2.34E-15 |
| AGPS | 0.319737633 | 0.001141458 | 0.002704323 |
| AGT | 2.185094266 | 0.011006115 | 0.021379894 |
| AGTRAP | 0.871614311 | 1.35E-10 | 7.19E-10 |
| AGXT2 | 4.300176273 | 7.58E-05 | 0.000214734 |
| AHCTF1 | -0.667436562 | 4.78E-07 | 1.81E-06 |
| AHCY | -1.37985998 | 2.00E-28 | 3.84E-27 |
| AHCYL2 | -1.009581892 | 9.43E-22 | 1.18E-20 |
| AHDC1 | 1.707848478 | 3.79E-12 | 2.35E-11 |
| AHI1 | 0.294461332 | 0.026664191 | 0.047443341 |
| AHSA1 | 0.317695592 | 0.00013637 | 0.000370707 |
| AHSA2 | 0.730255432 | 1.21E-14 | 9.24E-14 |
| AIDA | 1.012621977 | 1.02E-35 | 2.96E-34 |
| AIFM1 | -0.28680821 | 0.00259668 | 0.005774554 |
| AIFM2 | -0.645081166 | 4.03E-06 | 1.36E-05 |
| AIG1 | 0.386572755 | 0.000102901 | 0.000285667 |
| AIMP1 | -0.450563621 | 0.000281814 | 0.000733993 |
| AIMP2 | -0.270770686 | 0.00206425 | 0.00466688 |
| AK1 | 0.405429591 | 0.016425549 | 0.030688119 |
| AK3 | 1.054663392 | 3.24E-16 | 2.79E-15 |
| AK4 | 1.050523856 | 1.78E-30 | 3.84E-29 |
| AK5 | 1.222559586 | 0.016528586 | 0.030862882 |
| AK6 | 0.900390136 | 4.68E-17 | 4.31E-16 |
| AKAP1 | -0.515628642 | 2.14E-06 | 7.52E-06 |
| AKAP12 | -1.000329733 | 0.000415868 | 0.00105496 |
| AKAP14 | 5.883021008 | 0.000330541 | 0.000852184 |
| AKAP17A | -0.646306682 | 8.26E-07 | 3.05E-06 |

| | | | |
|---|---|---|---|
| AKAP6 | 0.745182901 | 0.02179585 | 0.039605508 |
| AKAP7 | 0.553042733 | 0.001278761 | 0.002998262 |
| AKAP8 | -0.616882264 | 0.001245314 | 0.002930963 |
| AKAP8L | 0.288371728 | 0.009961187 | 0.019523065 |
| AKAP9 | -1.076551896 | 6.88E-15 | 5.35E-14 |
| AKR1B10 | -0.542095851 | 4.05E-07 | 1.55E-06 |
| AKR7L | -0.77034471 | 1.68E-14 | 1.27E-13 |
| AKT1 | 0.443084105 | 5.28E-07 | 1.99E-06 |
| AKT2 | 0.620293666 | 0.001215439 | 0.002866605 |
| AKT3 | 0.467182864 | 0.026661735 | 0.047443341 |
| AKTIP | -1.345364908 | 3.04E-24 | 4.46E-23 |
| ALAD | 1.166973017 | 6.79E-32 | 1.59E-30 |
| ALAS1 | -1.848807037 | 6.37E-82 | 1.35E-79 |
| ALDH18A1 | -0.552458721 | 9.71E-11 | 5.26E-10 |
| ALDH1A3 | -0.350544627 | 0.00171809 | 0.003940647 |
| ALDH1L2 | -1.271090208 | 5.93E-07 | 2.22E-06 |
| ALDH3A2 | -0.237694198 | 0.012453914 | 0.023959462 |
| ALDH4A1 | -0.595822357 | 1.36E-06 | 4.88E-06 |
| ALDH9A1 | -0.62558577 | 1.28E-09 | 6.23E-09 |
| ALDOC | -0.619132451 | 3.28E-13 | 2.25E-12 |
| ALG14 | 0.472429031 | 0.004083744 | 0.008727399 |
| ALG2 | -0.55714161 | 5.96E-08 | 2.48E-07 |
| ALG3 | -0.57155747 | 0.000861086 | 0.002075157 |
| ALG5 | -0.205235716 | 0.023295248 | 0.04203784 |
| ALG6 | -0.423519337 | 0.001626128 | 0.003745523 |
| ALG8 | -0.381548298 | 0.010016452 | 0.019611583 |
| ALKAL2 | -1.658037805 | 1.56E-09 | 7.54E-09 |
| ALKBH1 | -0.341180381 | 0.009243488 | 0.018216252 |
| ALKBH3 | 0.442587866 | 0.001778312 | 0.004067793 |
| ALKBH8 | -0.560397281 | 0.00570261 | 0.011777826 |
| ALOX5 | -1.021820912 | 0.000471735 | 0.001184304 |
| ALPK1 | 1.519313233 | 0.008048937 | 0.016102613 |
| ALPK3 | -1.286159763 | 0.00555317 | 0.01150239 |
| ALX4 | -2.155445178 | 4.82E-22 | 6.16E-21 |
| ALYREF | 0.568862591 | 2.60E-10 | 1.35E-09 |
| AMACR | -0.638210566 | 0.000376831 | 0.000962762 |
| AMBRA1 | -0.868641312 | 1.34E-07 | 5.40E-07 |
| AMD1 | -0.698582679 | 1.08E-14 | 8.23E-14 |
| AMDHD1 | -1.008555078 | 0.00044936 | 0.001134418 |
| AMDHD2 | -1.022348486 | 3.47E-10 | 1.78E-09 |
| AMER1 | -1.106283644 | 5.29E-05 | 0.000153739 |
| AMFR | -0.26865469 | 0.001491614 | 0.003461503 |
| AMH | 3.960956932 | 4.08E-32 | 9.67E-31 |
| AMIGO3 | -1.559423204 | 2.52E-06 | 8.77E-06 |
| AMN | 0.841505061 | 0.007928976 | 0.015886002 |
| AMOT | -0.817800724 | 2.46E-12 | 1.55E-11 |
| AMOTL1 | -0.429555972 | 0.007157302 | 0.014470036 |
| AMOTL2 | -0.697618104 | 1.13E-09 | 5.55E-09 |

| | | | |
|---|---|---|---|
| AMPD3 | -0.572720633 | 0.005774615 | 0.011910253 |
| AMY1A | 0.960524217 | 0.000424906 | 0.001076682 |
| ANAPC1 | -0.608584064 | 0.00014999 | 0.000406029 |
| ANAPC13 | 1.051681944 | 1.42E-17 | 1.36E-16 |
| ANAPC16 | 1.08084696 | 1.87E-37 | 6.03E-36 |
| ANAPC2 | -0.815712259 | 1.18E-10 | 6.33E-10 |
| ANAPC5 | -1.850609892 | 1.03E-23 | 1.46E-22 |
| ANAPC7 | -1.496150561 | 2.16E-36 | 6.57E-35 |
| ANGEL1 | 1.041849978 | 2.79E-24 | 4.10E-23 |
| ANGEL2 | -0.379378772 | 0.00088419 | 0.002126312 |
| ANGPT1 | 0.555725303 | 0.000123096 | 0.000337594 |
| ANGPT2 | 0.356933091 | 0.004388495 | 0.009285246 |
| ANGPTL4 | 2.222289741 | 1.49E-162 | 2.03E-159 |
| ANK1 | 3.524370741 | 3.90E-05 | 0.000115225 |
| ANK2 | 1.127374741 | 0.000135047 | 0.000367333 |
| ANK3 | -0.729531277 | 3.65E-06 | 1.24E-05 |
| ANKFY1 | -0.608771925 | 0.005584009 | 0.011555701 |
| ANKHD1 | -0.745150459 | 5.70E-13 | 3.80E-12 |
| ANKIB1 | -0.661190255 | 3.10E-06 | 1.06E-05 |
| ANKRD1 | -1.418788912 | 1.62E-55 | 1.17E-53 |
| ANKRD10 | -1.130972645 | 1.02E-19 | 1.13E-18 |
| ANKRD11 | 0.967417423 | 1.56E-34 | 4.17E-33 |
| ANKRD13D | 0.936004174 | 2.60E-25 | 4.15E-24 |
| ANKRD17 | -0.566218123 | 5.11E-06 | 1.70E-05 |
| ANKRD22 | -1.482875044 | 9.87E-07 | 3.60E-06 |
| ANKRD28 | -1.270229438 | 8.28E-45 | 3.87E-43 |
| ANKRD29 | 0.468336188 | 0.00172649 | 0.003956575 |
| ANKRD39 | 1.05333122 | 0.000159748 | 0.000430643 |
| ANKRD44 | 0.626297096 | 0.00140303 | 0.003268767 |
| ANKRD49 | -0.651973485 | 3.63E-07 | 1.39E-06 |
| ANKRD52 | -0.747836234 | 1.36E-08 | 6.03E-08 |
| ANKRD61 | 5.1263244 | 1.30E-26 | 2.24E-25 |
| ANKRD9 | -1.979860679 | 3.64E-23 | 4.97E-22 |
| ANKS6 | -0.583368703 | 0.001219954 | 0.002876255 |
| ANKZF1 | -0.737978441 | 7.76E-06 | 2.52E-05 |
| ANLN | -0.880849708 | 0.000266022 | 0.000695074 |
| ANO10 | -1.060689459 | 4.33E-11 | 2.42E-10 |
| ANO5 | -0.980167089 | 2.96E-28 | 5.62E-27 |
| ANO8 | 1.75472447 | 3.70E-35 | 1.03E-33 |
| ANP32B | 0.366037148 | 5.51E-05 | 0.000159814 |
| ANTXR1 | -0.576239673 | 0.003727969 | 0.008025111 |
| ANXA1 | -0.853069903 | 1.10E-16 | 9.88E-16 |
| ANXA11 | 0.286573912 | 0.000587646 | 0.00145488 |
| ANXA5 | 0.217913892 | 0.016069792 | 0.030097922 |
| ANXA6 | -0.88260411 | 7.34E-22 | 9.23E-21 |
| ANXA7 | -0.521023814 | 1.23E-06 | 4.45E-06 |
| AOPEP | 0.421464903 | 6.16E-06 | 2.02E-05 |
| AP1AR | -0.604169033 | 2.86E-06 | 9.87E-06 |

| | | | |
|---|---|---|---|
| AP1B1 | -1.181188437 | 1.97E-30 | 4.24E-29 |
| AP1G1 | -0.767536497 | 2.73E-08 | 1.17E-07 |
| AP1M1 | -0.201256254 | 0.021393147 | 0.038946645 |
| AP1S2 | 0.575211649 | 1.63E-11 | 9.54E-11 |
| AP2A2 | -1.326625225 | 3.65E-49 | 2.17E-47 |
| AP2B1 | -0.838714772 | 2.69E-06 | 9.30E-06 |
| AP2M1 | 0.527408163 | 1.29E-11 | 7.59E-11 |
| AP3B1 | -1.073634134 | 3.41E-23 | 4.68E-22 |
| AP3M1 | 0.600290991 | 4.03E-12 | 2.50E-11 |
| AP3M2 | -0.515049144 | 2.47E-09 | 1.18E-08 |
| AP3S1 | 0.723511312 | 8.43E-15 | 6.51E-14 |
| AP3S2 | 0.702826405 | 3.25E-10 | 1.68E-09 |
| AP4B1 | -1.488782901 | 6.50E-24 | 9.34E-23 |
| AP4E1 | -1.447001915 | 3.84E-19 | 4.10E-18 |
| AP4S1 | 1.047013549 | 1.53E-07 | 6.11E-07 |
| AP5M1 | 1.193645275 | 1.72E-37 | 5.58E-36 |
| AP5Z1 | -0.730481416 | 6.47E-06 | 2.12E-05 |
| APAF1 | -1.06569301 | 2.62E-09 | 1.24E-08 |
| APBB1IP | -1.213524042 | 0.025098432 | 0.044927944 |
| APBB3 | 2.913084811 | 0.022680964 | 0.041038294 |
| APC | 0.714610302 | 3.24E-08 | 1.38E-07 |
| APC2 | 5.428686201 | 4.08E-12 | 2.52E-11 |
| APCDD1L | -0.276005899 | 0.014333914 | 0.027149743 |
| APEH | -0.48642492 | 8.62E-05 | 0.000242567 |
| APH1A | 0.720864959 | 0.006883886 | 0.013977558 |
| API5 | -0.713070174 | 2.61E-13 | 1.79E-12 |
| APIP | 0.307464061 | 0.011017869 | 0.021393554 |
| APLF | -0.592964681 | 3.48E-06 | 1.18E-05 |
| APLP2 | -0.888403381 | 1.76E-23 | 2.46E-22 |
| APMAP | 0.583867041 | 9.13E-10 | 4.51E-09 |
| APOA1 | -0.339758964 | 0.004107885 | 0.008769338 |
| APOLD1 | -2.294244221 | 1.14E-16 | 1.02E-15 |
| APOO | -0.528713918 | 1.75E-06 | 6.20E-06 |
| APOOL | -0.574214944 | 0.000481827 | 0.001206966 |
| APP | -1.397787321 | 1.18E-58 | 9.62E-57 |
| APPBP2 | -1.28184783 | 1.47E-25 | 2.38E-24 |
| APPL1 | -1.198999501 | 4.76E-13 | 3.21E-12 |
| APPL2 | -0.781668891 | 9.36E-17 | 8.43E-16 |
| APRT | -0.334190574 | 0.003263048 | 0.007113287 |
| AQP1 | 1.316797867 | 1.18E-61 | 1.10E-59 |
| AQR | -0.649157306 | 1.48E-07 | 5.91E-07 |
| AR | -1.361177847 | 5.58E-31 | 1.24E-29 |
| ARAP2 | -0.265659572 | 0.007044069 | 0.014269362 |
| ARCN1 | -0.440070299 | 6.12E-09 | 2.80E-08 |
| ARF1 | 0.535807507 | 2.10E-09 | 1.01E-08 |
| ARF4 | 0.286479038 | 0.00026016 | 0.000680673 |
| ARF5 | 0.528691113 | 0.025963748 | 0.046300087 |
| ARF6 | 0.899168517 | 7.89E-29 | 1.55E-27 |

| | | | |
|---|---|---|---|
| ARFGAP2 | -0.968498121 | 1.30E-08 | 5.81E-08 |
| ARFGAP3 | 0.238574821 | 0.008584693 | 0.017044083 |
| ARFGEF1 | -0.365109575 | 0.020322569 | 0.037161894 |
| ARFGEF2 | -0.358900911 | 0.026713217 | 0.047524352 |
| ARFIP1 | 0.40166584 | 0.000913049 | 0.002192608 |
| ARFIP2 | 0.669637467 | 4.23E-11 | 2.37E-10 |
| ARFRP1 | -0.630386605 | 6.39E-13 | 4.24E-12 |
| ARG2 | -1.005497647 | 1.96E-09 | 9.42E-09 |
| ARGLU1 | 0.301298813 | 0.000182137 | 0.000487133 |
| ARHGAP1 | -0.863748199 | 1.25E-11 | 7.36E-11 |
| ARHGAP10 | -1.494783995 | 1.11E-23 | 1.58E-22 |
| ARHGAP11B | -0.3206914 | 0.015766601 | 0.029581605 |
| ARHGAP12 | -0.425340768 | 0.000659063 | 0.001616964 |
| ARHGAP21 | -0.473812958 | 4.52E-05 | 0.000132741 |
| ARHGAP24 | -0.991575948 | 9.55E-10 | 4.71E-09 |
| ARHGAP27 | 2.109584716 | 0.002440206 | 0.005447082 |
| ARHGAP29 | -1.06329987 | 1.14E-12 | 7.39E-12 |
| ARHGAP31 | -0.496775721 | 0.023580626 | 0.042524594 |
| ARHGAP5 | 0.563040762 | 3.86E-08 | 1.64E-07 |
| ARHGDIA | 0.732130122 | 1.37E-16 | 1.21E-15 |
| ARHGDIB | 1.054036411 | 3.75E-29 | 7.52E-28 |
| ARHGEF10L | -0.651383457 | 0.001992015 | 0.004514082 |
| ARHGEF11 | 1.142759512 | 7.11E-06 | 2.32E-05 |
| ARHGEF16 | -0.95879804 | 0.002980889 | 0.006553906 |
| ARHGEF18 | -1.137518498 | 4.21E-12 | 2.60E-11 |
| ARHGEF3 | -1.217981172 | 6.48E-16 | 5.44E-15 |
| ARHGEF33 | 4.220367527 | 5.39E-06 | 1.78E-05 |
| ARHGEF6 | 0.659174571 | 1.40E-11 | 8.24E-11 |
| ARID3B | 0.863141581 | 0.000348386 | 0.000894458 |
| ARID4A | -0.72279087 | 3.74E-07 | 1.43E-06 |
| ARID4B | -0.256865934 | 0.01647863 | 0.030778829 |
| ARID5A | -0.77747979 | 2.45E-06 | 8.53E-06 |
| ARID5B | 0.618632563 | 1.01E-07 | 4.11E-07 |
| ARIH2 | -0.653487749 | 1.01E-09 | 4.95E-09 |
| ARL13B | 0.509805655 | 2.46E-06 | 8.56E-06 |
| ARL14 | -3.325013746 | 4.31E-116 | 2.34E-113 |
| ARL14EP | 0.809990624 | 5.50E-18 | 5.41E-17 |
| ARL15 | 1.450217579 | 3.01E-15 | 2.40E-14 |
| ARL16 | -2.159726163 | 0.001128426 | 0.002675778 |
| ARL2BP | 0.314612968 | 0.012607148 | 0.024225454 |
| ARL3 | 0.684140488 | 1.57E-10 | 8.30E-10 |
| ARL4A | -1.732803375 | 6.02E-44 | 2.71E-42 |
| ARL4C | -0.295032698 | 0.001254659 | 0.002950179 |
| ARL5A | 0.632970607 | 5.94E-07 | 2.22E-06 |
| ARL6 | 0.335224264 | 0.005594569 | 0.011575793 |
| ARL6IP4 | 0.239425493 | 0.027291348 | 0.048445083 |
| ARL6IP5 | 1.004687886 | 3.28E-36 | 9.83E-35 |
| ARL8A | 0.414453129 | 1.06E-06 | 3.83E-06 |

| | | | |
|---|---|---|---|
| ARMC1 | 0.524448092 | 3.92E-08 | 1.66E-07 |
| ARMC6 | -1.013350053 | 2.43E-27 | 4.41E-26 |
| ARMC9 | -1.024024161 | 1.31E-06 | 4.72E-06 |
| ARNTL | -1.402730647 | 5.28E-16 | 4.48E-15 |
| ARNTL2 | 1.115107617 | 9.34E-32 | 2.16E-30 |
| ARPC1A | 0.914731212 | 2.10E-25 | 3.36E-24 |
| ARPC2 | 0.629029604 | 1.55E-15 | 1.26E-14 |
| ARPC5 | 0.788705353 | 2.89E-18 | 2.91E-17 |
| ARPC5L | -0.51912964 | 0.00019435 | 0.000517556 |
| ARPP19 | 1.063815981 | 1.52E-52 | 1.00E-50 |
| ARRDC2 | -0.798081669 | 4.49E-11 | 2.51E-10 |
| ARRDC3 | -0.511765262 | 0.000266867 | 0.000697013 |
| ARSA | -1.03984193 | 6.36E-20 | 7.12E-19 |
| ARSB | -1.236938038 | 1.17E-26 | 2.02E-25 |
| ARSH | 0.685300006 | 0.001111112 | 0.002637946 |
| ARSJ | 0.824635624 | 4.28E-20 | 4.84E-19 |
| ARSK | -0.763091752 | 1.73E-08 | 7.58E-08 |
| ARV1 | -1.505307825 | 1.64E-24 | 2.47E-23 |
| ARVCF | 0.463636351 | 0.000479868 | 0.001202723 |
| AS3MT | 0.346750898 | 0.004969116 | 0.010398804 |
| ASAH1 | -0.379778525 | 8.92E-07 | 3.27E-06 |
| ASAP2 | -0.671567515 | 0.009450495 | 0.018594547 |
| ASB1 | 0.85085025 | 8.01E-06 | 2.59E-05 |
| ASB12 | 1.436589849 | 0.006961299 | 0.014117871 |
| ASB13 | -0.867782077 | 0.000130039 | 0.000354847 |
| ASB15 | 5.600605151 | 0.00017881 | 0.000478988 |
| ASB3 | -0.751864348 | 1.09E-06 | 3.97E-06 |
| ASB5 | 1.112389334 | 2.67E-23 | 3.67E-22 |
| ASB6 | -0.795750994 | 2.87E-11 | 1.64E-10 |
| ASB7 | 0.503674751 | 0.000284883 | 0.000740651 |
| ASB8 | -0.870220184 | 8.65E-09 | 3.90E-08 |
| ASB9 | 0.593660444 | 6.26E-06 | 2.05E-05 |
| ASCC3 | -0.605705476 | 0.00026031 | 0.000680934 |
| ASCL3 | 1.446696598 | 0.017398511 | 0.032314916 |
| ASH1L | -0.362025569 | 0.022938993 | 0.041466523 |
| ASH2L | -0.434109835 | 4.21E-05 | 0.000124056 |
| ASL2 | -0.664199111 | 0.024304683 | 0.043720123 |
| ASMT | 1.859047546 | 0.021036619 | 0.038369528 |
| ASMTL | -0.299087972 | 0.010260417 | 0.020034418 |
| ASNS | -1.195598652 | 2.89E-25 | 4.60E-24 |
| ASPG | -1.053648148 | 7.35E-11 | 4.03E-10 |
| ASPH | -0.898503609 | 9.74E-09 | 4.38E-08 |
| ASPHD2 | 1.636623345 | 3.65E-08 | 1.55E-07 |
| ASPM | 1.290341609 | 3.34E-06 | 1.14E-05 |
| ASPN | 1.183059319 | 1.29E-36 | 4.00E-35 |
| ASRGL1 | -0.328396535 | 0.025506556 | 0.045580477 |
| ASS1 | 0.412323518 | 0.001577217 | 0.003642741 |
| ASXL1 | -1.227397083 | 5.97E-08 | 2.48E-07 |

| | | | |
|---|---|---|---|
| ATAD1 | -0.466207047 | 1.17E-05 | 3.70E-05 |
| ATAD2 | -0.758894637 | 7.33E-07 | 2.72E-06 |
| ATAD5 | -0.932214879 | 0.00021592 | 0.000571637 |
| ATCAY | 3.196602893 | 1.65E-17 | 1.57E-16 |
| ATE1 | 0.747066175 | 5.31E-14 | 3.87E-13 |
| ATF1 | 0.633701288 | 4.78E-13 | 3.22E-12 |
| ATF2 | 0.581351465 | 4.73E-08 | 1.99E-07 |
| ATF3 | 0.463578415 | 2.63E-05 | 7.95E-05 |
| ATF4 | -0.270775047 | 0.00418963 | 0.008918631 |
| ATF6 | -0.51493258 | 3.82E-08 | 1.62E-07 |
| ATF7IP | 0.424424708 | 0.000637652 | 0.00156925 |
| ATG10 | 0.976445459 | 3.09E-09 | 1.45E-08 |
| ATG101 | 0.22950425 | 0.02201179 | 0.039971175 |
| ATG12 | 0.73042289 | 4.19E-16 | 3.58E-15 |
| ATG13 | 0.948213438 | 2.11E-20 | 2.42E-19 |
| ATG16L1 | -0.359018994 | 0.010883942 | 0.021160713 |
| ATG2B | -1.373177358 | 4.25E-17 | 3.93E-16 |
| ATG3 | -0.661023766 | 7.89E-10 | 3.93E-09 |
| ATG4A | 0.706042605 | 3.54E-12 | 2.20E-11 |
| ATG4C | 0.381805685 | 0.000836622 | 0.002020499 |
| ATG7 | -0.414298176 | 0.00059051 | 0.001460905 |
| ATG9B | 0.453914092 | 0.000105958 | 0.000293556 |
| ATIC | -0.358917432 | 2.16E-05 | 6.58E-05 |
| ATL1 | -0.299841135 | 0.001290886 | 0.003024604 |
| ATOH8 | 0.877155053 | 2.74E-11 | 1.57E-10 |
| ATOX1 | 1.315423851 | 5.80E-26 | 9.64E-25 |
| ATP10D | 1.392016888 | 0.009889021 | 0.019390014 |
| ATP11B | 0.550123046 | 2.76E-09 | 1.30E-08 |
| ATP13A1 | -1.437262978 | 1.66E-47 | 8.97E-46 |
| ATP13A2 | 0.357766641 | 0.008848693 | 0.017505429 |
| ATP1A1 | -1.668070602 | 1.58E-100 | 5.66E-98 |
| ATP1B1 | -0.90185366 | 4.75E-20 | 5.36E-19 |
| ATP2A2 | -1.394112025 | 7.96E-42 | 3.17E-40 |
| ATP2A3 | -1.252662215 | 0.011994619 | 0.023124945 |
| ATP2B1 | -1.192631509 | 6.06E-34 | 1.58E-32 |
| ATP2B4 | -1.024589264 | 2.71E-08 | 1.17E-07 |
| ATP2C1 | 0.388975592 | 2.78E-05 | 8.36E-05 |
| ATP5A1Z | -0.967187531 | 5.13E-29 | 1.02E-27 |
| ATP5B | -0.625413157 | 7.26E-14 | 5.21E-13 |
| ATP5C1 | 0.240910402 | 0.028034516 | 0.049608794 |
| ATP5F1D | 1.218653932 | 2.26E-54 | 1.57E-52 |
| ATP5G3 | 0.382929105 | 0.00023433 | 0.00061761 |
| ATP5I | 0.931765199 | 2.85E-11 | 1.63E-10 |
| ATP5J2 | 0.834057472 | 4.91E-11 | 2.73E-10 |
| ATP5PF | 0.738443146 | 3.63E-12 | 2.26E-11 |
| ATP6AP1 | 0.323968634 | 0.001711529 | 0.00392718 |
| ATP6V0A1 | -0.649846875 | 3.42E-13 | 2.34E-12 |
| ATP6V0A2 | 0.4484617 | 3.04E-06 | 1.05E-05 |

| | | | |
|---|---|---|---|
| ATP6V0B | 0.386780562 | 1.26E-06 | 4.53E-06 |
| ATP6V0D1 | -0.258638099 | 0.008976915 | 0.017750106 |
| ATP6V0E1 | 1.207302042 | 1.34E-49 | 8.00E-48 |
| ATP6V0E2 | 0.769051258 | 2.88E-14 | 2.14E-13 |
| ATP6V1A | -0.487105416 | 1.17E-09 | 5.71E-09 |
| ATP6V1C1 | -0.715503921 | 2.17E-13 | 1.51E-12 |
| ATP6V1D | 0.290627651 | 0.000759957 | 0.001847824 |
| ATP6V1E1 | 0.271459798 | 0.00079442 | 0.001924734 |
| ATP6V1G1 | -0.325125309 | 0.005272807 | 0.010975191 |
| ATP6V1H | -1.16179862 | 1.43E-33 | 3.61E-32 |
| ATP7A | -1.037095145 | 1.75E-15 | 1.41E-14 |
| ATP8A1 | 1.11993279 | 0.014314274 | 0.027116949 |
| ATP8B2 | 0.738213019 | 0.003505333 | 0.00758791 |
| ATP9A | -0.583517121 | 0.016188143 | 0.030298711 |
| ATPSCKMT | -1.394741504 | 3.89E-17 | 3.61E-16 |
| ATRN | -1.401015118 | 1.20E-39 | 4.20E-38 |
| ATRNL1 | -0.688260656 | 0.000133389 | 0.000363257 |
| ATXN1 | 0.571146955 | 0.006395824 | 0.013076421 |
| ATXN2 | -1.287536685 | 1.47E-28 | 2.85E-27 |
| ATXN7L3 | 1.813523155 | 4.47E-17 | 4.13E-16 |
| AURKA | -1.484601225 | 1.74E-15 | 1.41E-14 |
| AURKAIP1 | -0.779804847 | 5.44E-12 | 3.31E-11 |
| AvBD5 | 1.17402758 | 0.00355465 | 0.007684873 |
| AVL9 | -1.356256413 | 1.30E-14 | 9.86E-14 |
| AXIN1 | 0.688292075 | 7.76E-08 | 3.20E-07 |
| AXL | -0.271179638 | 0.002276885 | 0.005106835 |
| AZI2 | 1.412814777 | 1.04E-51 | 6.73E-50 |
| B3GALNT1 | 0.45629735 | 6.38E-06 | 2.09E-05 |
| B3GALNT2 | 0.712748589 | 1.44E-11 | 8.45E-11 |
| B3GALT2 | 1.824051983 | 0.000451116 | 0.00113839 |
| B3GLCT | 0.483267236 | 0.001592751 | 0.003673624 |
| B3GNT4 | -0.863677419 | 0.027019211 | 0.047999624 |
| B3GNT7 | 1.314880221 | 2.49E-05 | 7.54E-05 |
| B3GNT9 | 1.503536958 | 4.30E-40 | 1.54E-38 |
| B4GALNT3 | -0.902752565 | 0.000747655 | 0.001819865 |
| B4GALT2 | 1.329106751 | 1.13E-42 | 4.74E-41 |
| B4GALT3 | 1.276427462 | 8.97E-07 | 3.29E-06 |
| B4GALT4 | 0.651759954 | 1.67E-09 | 8.07E-09 |
| B4GALT5 | 0.472482595 | 1.99E-06 | 7.01E-06 |
| B4GALT6 | 1.026345808 | 9.02E-41 | 3.37E-39 |
| B9D1 | 0.425350124 | 0.001826827 | 0.004170346 |
| BACE1 | 0.908092109 | 4.42E-18 | 4.38E-17 |
| BACE2 | -0.351661706 | 0.003190399 | 0.006972134 |
| BACH1 | -0.648351919 | 3.25E-12 | 2.04E-11 |
| BAG1 | -0.705958973 | 3.03E-11 | 1.72E-10 |
| BAG2 | 1.145106529 | 7.97E-31 | 1.75E-29 |
| BAG5 | 0.49069606 | 2.03E-06 | 7.14E-06 |
| BAHCC1 | 1.570144501 | 3.33E-25 | 5.26E-24 |

| | | | |
|---|---|---|---|
| BAHD1 | -0.587291599 | 0.013234433 | 0.025299917 |
| BAIAP2 | -1.983571424 | 1.20E-24 | 1.81E-23 |
| BAIAP2L1 | -0.898453463 | 4.65E-11 | 2.59E-10 |
| BAK1 | 0.737711509 | 7.01E-18 | 6.84E-17 |
| BAMBI | -1.391496835 | 9.60E-61 | 8.69E-59 |
| BAP1 | 0.621676139 | 1.34E-08 | 5.94E-08 |
| BARD1 | -1.26422016 | 2.80E-07 | 1.09E-06 |
| BARX1 | 0.835122974 | 4.06E-06 | 1.36E-05 |
| BASP1 | 0.62312716 | 2.52E-13 | 1.74E-12 |
| BAZ1B | 0.29720833 | 0.006432435 | 0.013139414 |
| BAZ2B | -0.471895448 | 0.000102842 | 0.000285563 |
| BBIP1 | 0.916005599 | 4.89E-12 | 3.00E-11 |
| BBS12 | -1.219885295 | 8.19E-19 | 8.58E-18 |
| BBS2 | -0.325205756 | 0.008675447 | 0.017199048 |
| BBS5 | -0.781087079 | 7.79E-13 | 5.13E-12 |
| BBS7 | -0.402260375 | 0.000377865 | 0.000965039 |
| BBX | 0.339134525 | 0.004188122 | 0.008916816 |
| BCAN | -1.363437624 | 4.42E-28 | 8.33E-27 |
| BCAP29 | -0.678597307 | 1.79E-10 | 9.46E-10 |
| BCAR1 | -1.436136738 | 1.25E-35 | 3.61E-34 |
| BCAS3 | -0.559371389 | 0.005263353 | 0.010957191 |
| BCAT1 | -0.703203173 | 3.03E-09 | 1.42E-08 |
| BCCIP | 0.474509802 | 2.72E-07 | 1.06E-06 |
| BCDIN3D | -0.912685955 | 3.24E-08 | 1.38E-07 |
| BCHE | -0.593727865 | 9.82E-05 | 0.000273654 |
| BCKDHB | -0.720839641 | 7.62E-06 | 2.47E-05 |
| BCL10 | 0.484516365 | 8.37E-09 | 3.78E-08 |
| BCL2 | 0.73543665 | 0.00094756 | 0.002271871 |
| BCL2L1 | 1.213194143 | 3.31E-46 | 1.70E-44 |
| BCL2L11 | 1.612436272 | 7.76E-36 | 2.27E-34 |
| BCL2L13 | 0.47030239 | 0.001866508 | 0.004250221 |
| BCL6 | 0.426110516 | 9.75E-06 | 3.12E-05 |
| BCL7B | 0.46261456 | 5.13E-05 | 0.000149429 |
| BCL9L | 1.814648411 | 3.69E-38 | 1.24E-36 |
| BCOR | 4.770697246 | 0.000000 | 0.000000 |
| BCS1L | -1.18375873 | 2.12E-17 | 2.00E-16 |
| BDH1A | -0.588687532 | 8.27E-09 | 3.74E-08 |
| BDKRB2 | 0.371956406 | 0.00076487 | 0.001859104 |
| BDNF | 1.33663291 | 6.13E-64 | 6.41E-62 |
| BEAN1 | 2.018206607 | 3.75E-12 | 2.33E-11 |
| BECN1 | 0.601843741 | 2.86E-10 | 1.48E-09 |
| BEND5 | -1.056896261 | 1.96E-15 | 1.58E-14 |
| BEND6 | -1.373161496 | 7.92E-08 | 3.26E-07 |
| BEND7 | 0.6685189 | 0.000254139 | 0.000666331 |
| BEST1 | 4.990638651 | 1.14E-30 | 2.48E-29 |
| BEST3 | 5.154514838 | 0.001254166 | 0.002949753 |
| BET1 | -1.055593222 | 2.11E-26 | 3.59E-25 |
| BET1L | 0.415376943 | 6.66E-05 | 0.000190534 |

| | | | |
|---|---|---|---|
| BF2 | -0.601143395 | 0.000259976 | 0.000680322 |
| BFSP1 | 3.184253609 | 0.010180483 | 0.019898349 |
| BGLAP | 2.763086849 | 0.01490581 | 0.028127137 |
| BHLHE23 | -1.487212759 | 0.001232248 | 0.002903226 |
| BHLHE40 | 0.584971698 | 2.71E-10 | 1.41E-09 |
| BICC1 | -0.534192081 | 1.74E-08 | 7.65E-08 |
| BICD2 | -0.372476262 | 8.71E-06 | 2.80E-05 |
| BID | -0.428253784 | 4.97E-05 | 0.000144966 |
| BIK | -3.47729137 | 1.10E-10 | 5.91E-10 |
| BIRC3 | 0.392865556 | 0.000630761 | 0.001553697 |
| BIRC5 | -0.68411037 | 0.013261256 | 0.025344065 |
| BIRC8 | -0.912616332 | 6.16E-07 | 2.30E-06 |
| BKJ | 2.158454064 | 3.10E-19 | 3.33E-18 |
| BLCAP | 1.44345999 | 1.36E-58 | 1.09E-56 |
| BLM | -1.307819482 | 9.25E-06 | 2.96E-05 |
| BLMH | -0.74163276 | 5.38E-14 | 3.91E-13 |
| BLOC1S2 | 0.529987457 | 2.85E-07 | 1.11E-06 |
| BLOC1S4 | 0.573657843 | 1.21E-10 | 6.47E-10 |
| BLOC1S5 | 0.411461717 | 1.33E-05 | 4.15E-05 |
| BLOC1S6 | 1.179876305 | 6.73E-49 | 3.91E-47 |
| BLVRA | 1.312233226 | 0.025709819 | 0.045910743 |
| BLZF1 | -0.548502389 | 2.11E-06 | 7.42E-06 |
| BMF | 1.602633164 | 7.20E-34 | 1.86E-32 |
| BMI1 | 1.053108841 | 1.34E-22 | 1.79E-21 |
| BMP10 | 3.577595846 | 6.11E-05 | 0.000176039 |
| BMP15 | -2.078910283 | 0.021591581 | 0.039281591 |
| BMP2K | -1.301527713 | 2.15E-13 | 1.49E-12 |
| BMPR1A | -1.100256902 | 2.16E-11 | 1.25E-10 |
| BMPR1B | 6.460158145 | 2.88E-05 | 8.64E-05 |
| BNIP1 | -1.121319583 | 4.63E-22 | 5.93E-21 |
| BNIP2 | 0.305843621 | 0.000956689 | 0.002292141 |
| BNIP3 | 0.686838039 | 5.44E-09 | 2.50E-08 |
| BNIP3L | 0.336450395 | 0.000427921 | 0.001083716 |
| BOC | -1.037943396 | 1.63E-16 | 1.44E-15 |
| BOK | 0.467278176 | 8.88E-05 | 0.000249134 |
| BORA | -0.893449868 | 0.010547043 | 0.020574478 |
| BORCS6 | -0.573646159 | 0.000282321 | 0.000735115 |
| BORCS7 | -0.575489749 | 0.000562844 | 0.001397291 |
| BORCS8 | 0.543129474 | 3.91E-06 | 1.32E-05 |
| BPNT1 | -0.30674673 | 0.016336801 | 0.030547503 |
| BRAP | -0.383494935 | 4.83E-05 | 0.000141287 |
| BRAT1 | -0.523102882 | 0.000994038 | 0.002377855 |
| BRCC3 | -0.513601635 | 1.33E-05 | 4.17E-05 |
| BRD1 | -0.312956635 | 0.019875065 | 0.036461314 |
| BRD2 | -0.364803572 | 0.001683229 | 0.003868443 |
| BRD3 | 0.396453714 | 0.002202698 | 0.004954434 |
| BRD8 | -1.026642278 | 1.48E-17 | 1.42E-16 |
| BRE | 0.749269888 | 2.67E-16 | 2.32E-15 |

| | | | |
|---|---|---|---|
| BRF1 | 0.56018455 | 0.000785977 | 0.001905976 |
| BRI3 | 0.81699708 | 3.21E-21 | 3.87E-20 |
| BRI3BP | 0.324529928 | 0.001810348 | 0.004135505 |
| BRICD5 | 1.139462376 | 5.04E-06 | 1.67E-05 |
| BRIX1 | -0.253937718 | 0.022610505 | 0.040921702 |
| BRPF1 | -0.807506378 | 3.23E-09 | 1.52E-08 |
| BRS3 | 3.468670378 | 0.002408791 | 0.00538403 |
| BRWD1 | -0.837144446 | 7.63E-13 | 5.03E-12 |
| BSDC1 | -0.431681892 | 9.04E-06 | 2.90E-05 |
| BST1 | 0.811757696 | 1.12E-12 | 7.27E-12 |
| BSX | 6.29706755 | 1.72E-26 | 2.93E-25 |
| BTAF1 | -0.946424835 | 8.99E-10 | 4.45E-09 |
| BTBD1 | 0.219828746 | 0.00722129 | 0.014579883 |
| BTBD10 | -0.775986559 | 6.78E-12 | 4.11E-11 |
| BTBD11 | -1.440730395 | 2.71E-17 | 2.55E-16 |
| BTBD2 | -1.082718285 | 2.64E-09 | 1.25E-08 |
| BTBD3 | -0.529670941 | 0.011768198 | 0.022714193 |
| BTBD6 | -0.789687696 | 1.07E-10 | 5.76E-10 |
| BTBD8 | -0.336305776 | 0.018877742 | 0.034819686 |
| BTD | -0.615429989 | 0.000680975 | 0.001668297 |
| BTF3 | 1.017318918 | 4.34E-34 | 1.14E-32 |
| BTG1 | -1.714443341 | 9.72E-91 | 2.75E-88 |
| BTG2 | -1.744231558 | 6.26E-87 | 1.61E-84 |
| BTG4 | 3.422438623 | 0.001898461 | 0.00431792 |
| BTN1A1 | -0.935867237 | 0.003999697 | 0.008557199 |
| BTRC | -1.017342917 | 4.87E-06 | 1.62E-05 |
| BUB1B | -1.068280238 | 0.012792369 | 0.024548056 |
| BUB3 | -0.302592116 | 0.003000799 | 0.006592349 |
| BUD13 | -0.786698778 | 3.05E-07 | 1.18E-06 |
| BUD31 | 0.461482192 | 1.50E-05 | 4.66E-05 |
| BYSL | -0.848464133 | 1.74E-17 | 1.66E-16 |
| BZFP1 | -1.536276946 | 2.57E-05 | 7.76E-05 |
| BZW1 | -0.326586287 | 8.44E-05 | 0.000237912 |
| BZW2 | 0.311903531 | 0.000723835 | 0.001765685 |
| C10orf2 | -0.878931342 | 3.93E-17 | 3.65E-16 |
| C10orf71 | 0.406794748 | 0.00042505 | 0.001076847 |
| C10orf88 | 0.328280858 | 0.00131932 | 0.003084844 |
| C11H16orf70 | -0.745118686 | 4.59E-10 | 2.33E-09 |
| C11H16ORF87 | -0.765303923 | 1.76E-07 | 6.98E-07 |
| C11H19orf12 | 0.862374948 | 7.66E-16 | 6.40E-15 |
| C12orf4 | 0.324573843 | 0.002912272 | 0.006415496 |
| C12orf43 | -0.672697055 | 0.009329986 | 0.018368072 |
| C12orf66 | 0.525937321 | 5.89E-07 | 2.21E-06 |
| C12orf75 | 1.877977214 | 3.77E-101 | 1.38E-98 |
| C13H5orf15 | 0.288954441 | 0.0047221 | 0.009933878 |
| C14H16orf45 | 0.638414466 | 0.004548025 | 0.00958994 |
| C14orf2 | 1.150229366 | 3.61E-24 | 5.26E-23 |
| C15H12orf49 | 0.496565482 | 8.26E-09 | 3.74E-08 |

| | | | |
|---|---|---|---|
| C15orf48 | 1.268862427 | 1.54E-07 | 6.17E-07 |
| C15orf61 | 0.58224955 | 6.60E-05 | 0.000188883 |
| C15orf62 | -1.040973372 | 5.19E-09 | 2.40E-08 |
| C16orf72 | -0.348739921 | 0.001069447 | 0.002546141 |
| C16orf89 | 1.256491227 | 0.003723154 | 0.008017285 |
| C17H9orf16 | 0.604895994 | 5.04E-12 | 3.09E-11 |
| C17orf58 | 2.055946017 | 0.000806379 | 0.001951621 |
| C17orf75 | -0.246270608 | 0.022156837 | 0.040213076 |
| C19orf44 | -1.884052454 | 1.82E-12 | 1.16E-11 |
| C1GALT1 | 1.072628024 | 7.83E-36 | 2.29E-34 |
| C1GALT1C1 | 0.332983902 | 0.000650772 | 0.001598933 |
| C1H12ORF57 | 1.189993209 | 4.59E-40 | 1.63E-38 |
| C1H12ORF73 | 0.757746369 | 2.12E-14 | 1.59E-13 |
| C1H21ORF91 | -0.347731174 | 0.000347682 | 0.000893325 |
| C1H2ORF49 | -1.08986513 | 2.01E-23 | 2.79E-22 |
| C1orf109 | -0.809621146 | 0.018292383 | 0.033841033 |
| C1orf115 | 1.148495933 | 0.000953162 | 0.002284497 |
| C1orf145 | 0.930060425 | 0.003050646 | 0.00669104 |
| C1orf174 | 0.410960115 | 0.000454681 | 0.001146338 |
| C1orf198 | 0.80004785 | 7.60E-14 | 5.44E-13 |
| C1orf21 | 0.667463741 | 7.76E-17 | 7.01E-16 |
| C1QTNF12 | 1.401693107 | 1.10E-31 | 2.53E-30 |
| C1QTNF5 | 0.760847319 | 0.00313313 | 0.006859779 |
| C1QTNF6 | -1.269841718 | 0.023073223 | 0.041692535 |
| C1QTNF7 | 3.308463498 | 0.00297047 | 0.00653311 |
| C1QTNF8 | 1.375735544 | 1.93E-05 | 5.92E-05 |
| C1R | -0.806323795 | 0.000288428 | 0.000749583 |
| C1S | 0.519697553 | 0.000232231 | 0.000612315 |
| C20H20ORF24 | 1.315479367 | 1.83E-58 | 1.44E-56 |
| C20orf194 | -0.364997165 | 0.004736458 | 0.009954837 |
| C22orf23 | -1.312659366 | 4.09E-07 | 1.56E-06 |
| C25H1ORF43 | 0.698304687 | 5.68E-17 | 5.18E-16 |
| C26H6orf89 | -0.338591528 | 0.006772541 | 0.013769989 |
| C2CD5 | -0.691064827 | 1.58E-07 | 6.29E-07 |
| C2H6orf52 | 0.270207294 | 0.007973074 | 0.015960492 |
| C2H8ORF22 | -1.212499383 | 0.000409288 | 0.001039627 |
| C2orf88 | 1.330489671 | 0.012851102 | 0.024657282 |
| C3H6orf120 | 0.541210779 | 5.47E-09 | 2.51E-08 |
| C3H6ORF203 | -0.933110285 | 2.09E-11 | 1.20E-10 |
| C3orf14 | 1.040957039 | 6.08E-18 | 5.97E-17 |
| C3orf18 | 0.981039373 | 5.71E-05 | 0.000165106 |
| C4 | 0.416570952 | 0.000677879 | 0.001661628 |
| C4BPA | -0.464462739 | 1.03E-08 | 4.61E-08 |
| C4orf46 | 0.605841286 | 0.000528925 | 0.001318626 |
| C5 | 1.913102593 | 0.000604716 | 0.001493061 |
| C5H11orf58 | 0.48238384 | 2.60E-06 | 9.02E-06 |
| C5H11orf74 | 0.665260146 | 0.014243837 | 0.027009241 |
| C5H14ORF169 | -1.06042429 | 4.32E-16 | 3.69E-15 |

| | | | |
|---|---|---|---|
| C5H15orf41 | 0.793902276 | 3.12E-06 | 1.07E-05 |
| C5H15orf57 | 0.749102189 | 1.64E-17 | 1.56E-16 |
| C5orf22 | 1.024343239 | 2.09E-12 | 1.32E-11 |
| C5orf24 | 1.104998313 | 9.77E-35 | 2.65E-33 |
| C6orf58 | -0.904592493 | 2.52E-09 | 1.20E-08 |
| C7H2ORF69 | -0.60242959 | 6.42E-05 | 0.000183957 |
| C7H2ORF76 | 1.155070303 | 2.03E-24 | 3.02E-23 |
| C7orf25 | -0.492932287 | 4.08E-05 | 0.000120212 |
| C7orf26 | -0.487954013 | 0.003281844 | 0.007150822 |
| C7orf73 | 0.946659802 | 1.48E-15 | 1.21E-14 |
| C8G | 1.1426209 | 2.22E-06 | 7.77E-06 |
| C9H2ORF82 | 3.148215318 | 0.015767189 | 0.029581605 |
| C9orf116 | 0.826572553 | 0.000639996 | 0.001574447 |
| C9orf64 | -0.264407045 | 0.011496184 | 0.022239701 |
| C9orf69 | 0.883240834 | 7.16E-11 | 3.93E-10 |
| C9orf72 | -0.606844699 | 2.18E-06 | 7.64E-06 |
| C9orf85 | 0.720395922 | 5.78E-13 | 3.86E-12 |
| CA2 | 0.916996921 | 1.80E-16 | 1.58E-15 |
| CA3B | 0.764038742 | 0.000248315 | 0.000652318 |
| CAB39 | -0.330847291 | 0.000100876 | 0.000280448 |
| CAB39L | 0.818976802 | 6.42E-15 | 5.00E-14 |
| CABIN1 | -0.660454262 | 0.016027449 | 0.030022753 |
| CABLES2 | -1.087573283 | 0.000180524 | 0.000483294 |
| CABP2 | 0.518485089 | 0.003686471 | 0.007947109 |
| CACFD1 | 0.712623929 | 1.91E-07 | 7.53E-07 |
| CACHD1 | -1.451051984 | 1.06E-09 | 5.18E-09 |
| CACNA2D1 | -0.686122235 | 5.23E-09 | 2.41E-08 |
| CACNG3 | 1.229647834 | 1.56E-10 | 8.26E-10 |
| CACTIN | 0.307025757 | 0.003532678 | 0.007641018 |
| CADM1 | -1.238615981 | 0.000393981 | 0.001003371 |
| CADM2 | 2.41519476 | 0.002376553 | 0.005318971 |
| CALCA | -2.036839274 | 3.04E-71 | 4.13E-69 |
| CALD1 | -0.264279821 | 0.006413831 | 0.013109292 |
| CALHM2 | -1.260671212 | 8.22E-14 | 5.86E-13 |
| CALHM6 | -1.220790564 | 0.004942234 | 0.010347328 |
| CALM1 | 0.448534346 | 5.22E-08 | 2.18E-07 |
| CALM2 | -0.806004948 | 2.69E-11 | 1.54E-10 |
| CALML4 | 0.88970428 | 2.52E-12 | 1.59E-11 |
| CAMK1 | 0.559114811 | 0.001531492 | 0.003545569 |
| CAMK1D | 0.959679671 | 0.000168207 | 0.00045201 |
| CAMK1G | 0.570051266 | 0.000346096 | 0.000889587 |
| CAMK2B | 1.131120484 | 3.07E-07 | 1.19E-06 |
| CAMK2D | 0.446340976 | 1.04E-07 | 4.25E-07 |
| CAMK2G | 0.669445801 | 3.83E-07 | 1.47E-06 |
| CAMK2N2 | 1.000623252 | 0.000721747 | 0.001761535 |
| CAMK4 | 2.251810367 | 6.60E-06 | 2.16E-05 |
| CAMKK1 | -0.754598313 | 8.56E-05 | 0.000240934 |
| CAMSAP1 | -0.860752395 | 2.19E-06 | 7.67E-06 |

| | | | |
|---|---|---|---|
| CAND1 | -0.924867327 | 3.58E-19 | 3.83E-18 |
| CANT1 | -0.674073957 | 1.81E-09 | 8.72E-09 |
| CAP2 | 0.843528804 | 0.007444549 | 0.014990565 |
| CAPG | -0.817807845 | 1.64E-07 | 6.53E-07 |
| CAPN10 | -0.770268662 | 8.74E-07 | 3.21E-06 |
| CAPN11 | -0.351344156 | 0.000301693 | 0.000782261 |
| CAPN13 | 3.396349951 | 1.26E-09 | 6.12E-09 |
| CAPN14 | 0.456334291 | 0.009337833 | 0.018380858 |
| CAPN2 | -0.700794795 | 1.64E-10 | 8.70E-10 |
| CAPN7 | -1.002534494 | 3.90E-05 | 0.000115225 |
| CAPN9 | 1.362401863 | 9.50E-44 | 4.23E-42 |
| CAPRIN1 | -0.393619001 | 2.80E-05 | 8.43E-05 |
| CAPZA1 | 0.848551987 | 2.11E-18 | 2.14E-17 |
| CAPZA2 | 0.789757269 | 1.02E-15 | 8.39E-15 |
| CAPZB | 0.477640176 | 1.91E-07 | 7.54E-07 |
| CARHSP1 | 1.425900568 | 1.51E-58 | 1.20E-56 |
| CARMIL3 | 1.479203857 | 4.06E-05 | 0.000119648 |
| CARNMT1 | 0.601955697 | 6.66E-09 | 3.04E-08 |
| CARS | -0.266485422 | 0.002290893 | 0.005136558 |
| CARS2 | -0.538619304 | 9.39E-05 | 0.000262418 |
| CASC3 | 0.461131573 | 0.004193825 | 0.008926162 |
| CASD1 | -0.458836243 | 1.06E-05 | 3.38E-05 |
| CASP2 | -0.493529602 | 0.003899778 | 0.008355262 |
| CASP3 | -0.594597456 | 1.06E-07 | 4.32E-07 |
| CASP6 | -0.442092874 | 0.000643489 | 0.001582181 |
| CASP8 | -0.546988215 | 7.49E-08 | 3.09E-07 |
| CASS4 | 2.971771348 | 0.000260683 | 0.000681778 |
| CAST | -0.446987001 | 5.11E-07 | 1.93E-06 |
| CASTOR1 | -0.785746939 | 0.000680004 | 0.001666236 |
| CAT | -1.561035202 | 2.73E-37 | 8.76E-36 |
| CATHL1 | 3.754220383 | 4.09E-09 | 1.90E-08 |
| CAV1 | 0.709913173 | 2.60E-11 | 1.49E-10 |
| CAV2 | 0.438960007 | 2.08E-07 | 8.18E-07 |
| CAV3 | 3.445389288 | 0.001608478 | 0.003707383 |
| CBFA2T2 | 1.292255677 | 1.67E-19 | 1.83E-18 |
| CBFB | 0.625055701 | 1.78E-11 | 1.03E-10 |
| CBL | -0.687752868 | 0.000332054 | 0.00085576 |
| CBLL1 | -0.377286055 | 0.005995082 | 0.012323828 |
| CBWD1 | 0.734232991 | 8.27E-11 | 4.51E-10 |
| CBX3 | 0.636803239 | 2.12E-12 | 1.34E-11 |
| CBX4 | -1.216546238 | 1.73E-16 | 1.53E-15 |
| CBX8 | -1.734654489 | 3.54E-10 | 1.82E-09 |
| CBY1 | 0.326274482 | 0.002477869 | 0.005525708 |
| CC2D1B | -0.381423029 | 0.000396994 | 0.001010288 |
| CCDC12 | 0.770317587 | 7.09E-15 | 5.50E-14 |
| CCDC125 | 3.22118543 | 0.004875046 | 0.010214525 |
| CCDC126 | 0.827712315 | 2.39E-22 | 3.13E-21 |
| CCDC137 | -0.425159525 | 0.004143662 | 0.008834613 |

| | | | |
|---|---|---|---|
| CCDC14 | -0.82059269 | 3.83E-05 | 0.000113361 |
| CCDC141 | 1.412064773 | 0.008914058 | 0.017630947 |
| CCDC149 | -0.628036305 | 0.011665383 | 0.022541354 |
| CCDC167 | 0.567735046 | 3.02E-07 | 1.17E-06 |
| CCDC18 | 0.514603558 | 0.003468729 | 0.007518254 |
| CCDC186 | 0.604898769 | 5.58E-08 | 2.33E-07 |
| CCDC191 | -0.886201011 | 0.005301071 | 0.011030642 |
| CCDC25 | -0.473575321 | 5.82E-06 | 1.92E-05 |
| CCDC28B | -0.69843691 | 0.006894405 | 0.013994736 |
| CCDC43 | 0.573395866 | 3.60E-07 | 1.38E-06 |
| CCDC47 | -0.29924243 | 0.000841572 | 0.002031731 |
| CCDC50 | 0.886206771 | 8.39E-11 | 4.58E-10 |
| CCDC58 | -0.687005888 | 8.12E-06 | 2.63E-05 |
| CCDC59 | 0.458743972 | 4.71E-05 | 0.000137872 |
| CCDC61 | -0.733929507 | 5.23E-07 | 1.97E-06 |
| CCDC77 | -0.74724468 | 0.009680765 | 0.019011842 |
| CCDC80 | -0.581582202 | 9.18E-15 | 7.05E-14 |
| CCDC82 | -0.495391702 | 0.000849014 | 0.002048244 |
| CCDC88A | -0.385811067 | 0.000256876 | 0.000672729 |
| CCDC88C | -0.29107129 | 0.002160593 | 0.004868499 |
| CCDC90B | -0.475723324 | 1.31E-05 | 4.11E-05 |
| CCDC91 | 0.343528221 | 0.016851898 | 0.031411272 |
| CCDC92 | 0.447036364 | 0.003229588 | 0.007047128 |
| CCDC93 | -0.332336836 | 0.002687866 | 0.005955897 |
| CCK | -0.265638523 | 0.014782432 | 0.02792534 |
| CCL17 | 3.377103485 | 6.01E-05 | 0.000173305 |
| CCL20 | 1.345668765 | 0.004729864 | 0.009943489 |
| CCL4 | 5.482060632 | 9.70E-67 | 1.15E-64 |
| CCL4.1 | 3.322639273 | 1.44E-226 | 3.92E-223 |
| CCM2 | 0.391633143 | 0.000590257 | 0.001460546 |
| CCN3 | -1.79540982 | 4.68E-30 | 9.88E-29 |
| CCNA2 | 0.464763638 | 0.0006471 | 0.001590772 |
| CCNB1 | -1.432238882 | 1.04E-23 | 1.47E-22 |
| CCNB3 | -0.993681573 | 0.000120386 | 0.00033043 |
| CCNC | 0.920808049 | 2.60E-31 | 5.87E-30 |
| CCND1 | 0.750703199 | 1.90E-15 | 1.53E-14 |
| CCND2 | 0.693621612 | 7.59E-14 | 5.43E-13 |
| CCND3 | 0.745352054 | 3.28E-08 | 1.40E-07 |
| CCNDBP1 | -0.373966903 | 8.25E-05 | 0.000232633 |
| CCNE1 | -0.558181617 | 5.69E-05 | 0.000164614 |
| CCNE2 | -1.428447132 | 3.24E-14 | 2.40E-13 |
| CCNF | -0.397503613 | 0.021199447 | 0.03863023 |
| CCNG1 | 1.055859463 | 4.10E-45 | 1.95E-43 |
| CCNG2 | -0.460484938 | 1.72E-05 | 5.30E-05 |
| CCNI | 1.698912972 | 5.06E-91 | 1.46E-88 |
| CCNI2 | 0.566167772 | 0.002430093 | 0.005428074 |
| CCNK | 0.640044149 | 1.06E-07 | 4.30E-07 |
| CCNL1 | -0.426212706 | 0.009116221 | 0.017999371 |

| | | | |
|---|---|---|---|
| CCNL2 | -0.768029779 | 2.60E-15 | 2.08E-14 |
| CCNO | -2.11013489 | 0.001079877 | 0.002570073 |
| CCNT1 | 1.199548519 | 3.84E-15 | 3.04E-14 |
| CCNY | 1.023389798 | 0.016156125 | 0.03024295 |
| CCP110 | -1.016502776 | 1.47E-07 | 5.88E-07 |
| CCPG1 | -0.637735433 | 3.32E-12 | 2.07E-11 |
| CCR5 | 3.132337345 | 0.003442861 | 0.007465758 |
| CCR6 | 1.747831687 | 0.01961249 | 0.036033113 |
| CCSER1 | -1.31914974 | 5.25E-13 | 3.52E-12 |
| CCT2 | -0.384035849 | 9.24E-07 | 3.38E-06 |
| CCT4 | -1.029932718 | 7.29E-29 | 1.44E-27 |
| CCT5 | -0.597364153 | 9.62E-09 | 4.33E-08 |
| CCT6A | -0.665456249 | 7.56E-18 | 7.36E-17 |
| CCT7 | -0.913591681 | 6.18E-25 | 9.61E-24 |
| CCT8 | -0.81985748 | 1.64E-15 | 1.33E-14 |
| CCZ1 | 0.47331546 | 1.16E-06 | 4.21E-06 |
| CD109 | -1.141513745 | 1.03E-08 | 4.62E-08 |
| CD151 | -0.542265114 | 9.18E-07 | 3.36E-06 |
| CD164 | -0.556969466 | 3.06E-11 | 1.74E-10 |
| CD164L2 | -0.912406828 | 0.004741039 | 0.009962925 |
| CD180 | 3.132822055 | 0.008379189 | 0.016699404 |
| CD200 | -1.504859273 | 0.008766281 | 0.017358871 |
| CD200R1 | 1.233190816 | 1.22E-11 | 7.21E-11 |
| CD2AP | 0.300608577 | 0.000962579 | 0.002305846 |
| CD320 | 0.689074987 | 4.96E-05 | 0.00014488 |
| CD36 | -1.170656566 | 1.30E-06 | 4.68E-06 |
| CD40 | -0.820231725 | 0.00130198 | 0.003048495 |
| CD44 | -1.478634374 | 2.29E-48 | 1.28E-46 |
| CD47 | -0.327390488 | 0.000262205 | 0.000685627 |
| CD81 | -0.486723974 | 5.31E-08 | 2.22E-07 |
| CD82 | -0.567070231 | 1.04E-09 | 5.09E-09 |
| CD83 | 1.264712733 | 0.004490619 | 0.009481927 |
| CD9 | -0.320317703 | 0.008518942 | 0.016930759 |
| CD99 | 0.207974037 | 0.020987614 | 0.038285284 |
| CD99L2 | 0.75177822 | 7.67E-23 | 1.03E-21 |
| CDA | 1.24609558 | 7.19E-14 | 5.17E-13 |
| CDADC1 | -0.438497545 | 0.002451575 | 0.00547156 |
| CDC14A | -0.743480417 | 2.46E-08 | 1.06E-07 |
| CDC14B | -0.964238488 | 6.04E-06 | 1.99E-05 |
| CDC20 | -1.388185497 | 5.51E-12 | 3.35E-11 |
| CDC23 | -0.728317726 | 2.59E-07 | 1.01E-06 |
| CDC25A | -0.455844339 | 0.005223852 | 0.010879956 |
| CDC25B | 1.630101885 | 1.77E-67 | 2.18E-65 |
| CDC26 | 0.350785817 | 0.024272439 | 0.0436679 |
| CDC27 | -0.398378479 | 0.000465189 | 0.001169814 |
| CDC2L1 | -0.55456006 | 3.01E-06 | 1.03E-05 |
| CDC34 | -0.82075578 | 7.17E-08 | 2.96E-07 |
| CDC37L1 | -1.061415809 | 1.43E-10 | 7.62E-10 |

| | | | |
|---|---|---|---|
| CDC42 | 0.787607904 | 1.84E-23 | 2.55E-22 |
| CDC42EP4 | -0.894730805 | 4.22E-18 | 4.19E-17 |
| CDC42SE1 | 1.203624405 | 0.002762173 | 0.006102643 |
| CDC42SE2 | 0.902273873 | 1.19E-16 | 1.07E-15 |
| CDC45 | -0.606465057 | 0.000289582 | 0.000752292 |
| CDC73 | -0.954153752 | 1.22E-20 | 1.41E-19 |
| CDCA4 | 0.535727904 | 3.28E-07 | 1.27E-06 |
| CDCA9 | -1.061153405 | 2.78E-07 | 1.08E-06 |
| CDH11 | 0.284732846 | 0.000456176 | 0.001149489 |
| CDH13 | -0.461151456 | 3.98E-10 | 2.03E-09 |
| CDH15 | -1.417648607 | 4.71E-06 | 1.57E-05 |
| CDH2 | -0.437346418 | 2.18E-06 | 7.64E-06 |
| CDH23 | 3.386526615 | 0.012098953 | 0.023319479 |
| CDH5 | 2.332829062 | 0.008849309 | 0.017505429 |
| CDH6 | 0.59550838 | 0.007192809 | 0.014533772 |
| CDHR1 | 6.794104839 | 2.84E-07 | 1.10E-06 |
| CDIP1 | 0.694031591 | 0.000129573 | 0.000353716 |
| CDK14 | 0.352154739 | 0.003754476 | 0.008073222 |
| CDK15 | -2.182578806 | 0.008460152 | 0.016831146 |
| CDK19 | 0.966787181 | 2.93E-07 | 1.14E-06 |
| CDK2 | 1.038894792 | 0.023570764 | 0.04251245 |
| CDK2AP1 | 1.704372231 | 3.71E-105 | 1.53E-102 |
| CDK5 | 0.807063449 | 7.40E-11 | 4.05E-10 |
| CDK5RAP1 | 0.392216731 | 2.80E-05 | 8.42E-05 |
| CDK5RAP3 | -0.94873009 | 2.75E-13 | 1.89E-12 |
| CDK7 | -1.290657396 | 3.92E-27 | 7.01E-26 |
| CDK9 | -0.501388531 | 5.59E-07 | 2.10E-06 |
| CDKAL1 | 0.459594737 | 9.16E-06 | 2.94E-05 |
| CDKL1 | 0.271727073 | 0.019930571 | 0.036548343 |
| CDKN1A | -0.297159938 | 0.002016831 | 0.004565751 |
| CDKN1B | 0.789806528 | 2.95E-14 | 2.19E-13 |
| CDKN2A | 1.596983239 | 6.73E-36 | 1.98E-34 |
| CDKN2B | 1.020580254 | 2.21E-24 | 3.28E-23 |
| CDKN3 | 0.66697034 | 0.000419274 | 0.001063004 |
| CDR2L | -1.085926237 | 0.013039872 | 0.024977162 |
| CDT1 | -1.493363492 | 9.72E-11 | 5.26E-10 |
| CDYL2 | 0.564215479 | 0.005440423 | 0.01129295 |
| CEBPB | -0.815054335 | 5.17E-15 | 4.05E-14 |
| CEBPD | -0.850877955 | 5.57E-05 | 0.000161445 |
| CEBPG | 0.560428542 | 8.88E-11 | 4.83E-10 |
| CEBPZOS | 1.006479171 | 1.06E-12 | 6.94E-12 |
| CECR1 | -1.047559376 | 1.79E-05 | 5.50E-05 |
| CELF1 | -0.610513765 | 8.01E-09 | 3.63E-08 |
| CELF2 | -0.334401098 | 0.017778195 | 0.032979559 |
| CEMIP | -0.495982285 | 0.011212846 | 0.021728686 |
| CENPC | -1.65811527 | 2.03E-10 | 1.07E-09 |
| CENPI | -0.967992091 | 0.013495716 | 0.025748705 |
| CENPL | -2.058005579 | 5.40E-05 | 0.000156813 |

| | | | |
|---|---|---|---|
| CENPN | -0.912400222 | 9.02E-07 | 3.30E-06 |
| CENPP | 1.3356417 | 1.10E-32 | 2.64E-31 |
| CENPT | -1.666194055 | 6.42E-20 | 7.17E-19 |
| CENPW | 0.297204214 | 0.012186167 | 0.023467608 |
| CEP120 | -0.789762541 | 1.53E-07 | 6.11E-07 |
| CEP162 | -0.797483036 | 7.55E-06 | 2.46E-05 |
| CEP192 | -1.003586931 | 5.84E-05 | 0.000168648 |
| CEP41 | -1.195607465 | 5.14E-16 | 4.36E-15 |
| CEP43 | -1.094637875 | 5.74E-16 | 4.84E-15 |
| CEP44 | -1.363273938 | 2.44E-23 | 3.37E-22 |
| CEP55 | -1.849094715 | 0.0011587 | 0.002741826 |
| CEP57 | -1.392651519 | 4.31E-12 | 2.66E-11 |
| CEP57L1 | -1.439177316 | 0.001608951 | 0.003707843 |
| CEP63 | -0.525223155 | 0.013269701 | 0.025353075 |
| CEP68 | -1.176557834 | 5.21E-19 | 5.53E-18 |
| CEP76 | -0.866604002 | 0.00855129 | 0.016987597 |
| CEP89 | -0.5609121 | 0.002597291 | 0.005774967 |
| CEP95 | -0.538688296 | 0.005542165 | 0.011483095 |
| CEPT1 | -0.568027223 | 1.73E-08 | 7.58E-08 |
| CERCAM | -1.00556587 | 9.33E-34 | 2.37E-32 |
| CERK | 0.866998619 | 4.25E-21 | 5.09E-20 |
| CERS1 | 0.793987491 | 0.011564176 | 0.022361685 |
| CERS5 | 0.263750322 | 0.007815034 | 0.015690096 |
| CERS6 | -1.511318611 | 9.92E-24 | 1.41E-22 |
| CERT1 | -1.824451851 | 2.52E-31 | 5.71E-30 |
| CETN1 | 0.921501999 | 8.35E-16 | 6.95E-15 |
| CETN2 | 0.859426547 | 3.97E-18 | 3.95E-17 |
| CETN3 | 1.212427774 | 2.49E-26 | 4.21E-25 |
| CETP | 2.001016169 | 6.27E-06 | 2.06E-05 |
| CFAP20 | 0.608618774 | 3.50E-10 | 1.80E-09 |
| CFAP36 | 0.900642277 | 7.16E-16 | 6.00E-15 |
| CFAP45 | -1.636734402 | 0.007317128 | 0.014758036 |
| CFAP58 | 2.247461151 | 0.002100049 | 0.004741505 |
| CFAP92 | 2.159364459 | 8.60E-13 | 5.63E-12 |
| CFAP94 | 1.525563864 | 0.022845424 | 0.041319361 |
| CFAP97 | -1.003535041 | 1.70E-15 | 1.38E-14 |
| CFAP99 | -2.137335053 | 0.000558361 | 0.001386923 |
| CFDP1 | 1.529309982 | 1.09E-32 | 2.62E-31 |
| CFI | 1.211276778 | 0.009660767 | 0.01897531 |
| CFL2 | 1.038306776 | 5.70E-35 | 1.59E-33 |
| CFLAR | -0.520635747 | 9.88E-07 | 3.60E-06 |
| CG-16 | 0.645792513 | 0.001579153 | 0.003645974 |
| CG-1B | 1.602899123 | 5.00E-79 | 9.57E-77 |
| CGGBP1 | -0.72863635 | 6.17E-06 | 2.03E-05 |
| CGRRF1 | -0.282344267 | 0.023662552 | 0.042661018 |
| CH25H | -1.088775565 | 4.43E-18 | 4.38E-17 |
| CHAC1 | -3.263384587 | 4.92E-126 | 3.35E-123 |
| CHAC2 | 0.582277092 | 1.08E-05 | 3.44E-05 |

| | | | |
|---|---|---|---|
| CHADL | -1.253931428 | 1.69E-07 | 6.73E-07 |
| CHAF1A | -0.822298526 | 7.90E-09 | 3.58E-08 |
| CHAMP1 | 0.433203578 | 0.001055591 | 0.002515133 |
| CHCHD1 | 0.384382576 | 0.001968061 | 0.004463521 |
| CHCHD2P9 | 0.225524093 | 0.023323018 | 0.042082306 |
| CHD2 | 0.478291451 | 5.34E-06 | 1.77E-05 |
| CHD5 | 0.605666097 | 0.006749562 | 0.01372787 |
| CHD7 | 0.318699625 | 0.00382738 | 0.008210504 |
| CHD9 | -0.383742784 | 0.016934267 | 0.0315475 |
| CHERP | -0.768759551 | 9.21E-15 | 7.07E-14 |
| CHGB | -0.961967128 | 3.06E-21 | 3.70E-20 |
| CHIC1 | -0.412633162 | 0.004291189 | 0.009099195 |
| CHIC2 | -1.514693361 | 2.13E-09 | 1.02E-08 |
| CHKA | -0.848850026 | 3.41E-14 | 2.52E-13 |
| CHM | -1.052968665 | 9.62E-18 | 9.30E-17 |
| CHMP1B | 0.258268917 | 0.013080607 | 0.025037546 |
| CHMP4B | -0.487000221 | 9.05E-11 | 4.92E-10 |
| CHMP5 | 0.265586498 | 0.001494082 | 0.003466639 |
| CHMP6 | 0.706794552 | 1.01E-12 | 6.61E-12 |
| CHN1 | 1.097172357 | 7.74E-25 | 1.19E-23 |
| CHORDC1 | -0.877127507 | 1.64E-15 | 1.33E-14 |
| CHPT1 | 1.204547641 | 8.85E-46 | 4.33E-44 |
| CHRAC1 | -0.545557124 | 1.58E-05 | 4.91E-05 |
| CHRDL2 | 1.378954481 | 0.002027865 | 0.004589965 |
| CHRM4 | 1.751709252 | 1.31E-17 | 1.26E-16 |
| CHRNA3 | 3.340854064 | 3.88E-07 | 1.48E-06 |
| CHRNA4 | 3.466320884 | 0.006118007 | 0.012546164 |
| CHRNA5 | 1.578660471 | 2.11E-09 | 1.01E-08 |
| CHST1 | -1.030171276 | 2.83E-06 | 9.76E-06 |
| CHST10 | 1.239247809 | 3.46E-41 | 1.33E-39 |
| CHST14 | 0.510203264 | 4.00E-06 | 1.35E-05 |
| CHST15 | 0.407069547 | 6.41E-05 | 0.0001839 |
| CHST2 | 1.028216397 | 1.18E-11 | 7.00E-11 |
| CHST3 | -0.785753724 | 1.65E-21 | 2.03E-20 |
| CHST9 | -1.838182483 | 2.70E-06 | 9.36E-06 |
| CHSY1 | 1.158455317 | 7.90E-41 | 2.97E-39 |
| CHSY3 | -1.030275883 | 0.000255612 | 0.000669806 |
| CHTF18 | -0.944851651 | 2.80E-08 | 1.20E-07 |
| CHUK | -0.373426695 | 3.79E-05 | 0.000112273 |
| CHURC1 | 1.084362196 | 7.90E-21 | 9.32E-20 |
| CIAO1 | -1.11741005 | 2.61E-23 | 3.60E-22 |
| CIAO2B | 0.330424352 | 0.005317872 | 0.011062189 |
| CIAPIN1 | 0.378679346 | 4.83E-06 | 1.61E-05 |
| CIB2 | 1.198304715 | 2.59E-10 | 1.35E-09 |
| CIB4 | 1.314510077 | 3.79E-60 | 3.28E-58 |
| CIDEA | -1.513494697 | 9.11E-34 | 2.33E-32 |
| CIDEC | -0.943881824 | 4.24E-07 | 1.61E-06 |
| CILK1 | 0.575695048 | 0.000107267 | 0.000296959 |

| | | | |
|---|---|---|---|
| CILP | -0.922428397 | 0.000186095 | 0.000497034 |
| CILP2 | -1.267714122 | 0.008537281 | 0.016962245 |
| CINP | -0.791431118 | 2.00E-06 | 7.05E-06 |
| CIPC | -0.920239073 | 0.008586894 | 0.017045868 |
| CISD1 | 0.343233148 | 1.91E-05 | 5.86E-05 |
| CIT | 2.164437891 | 9.04E-06 | 2.90E-05 |
| CITED2 | -1.150815607 | 0.000136468 | 0.000370901 |
| CITED4 | -0.733086863 | 4.09E-17 | 3.80E-16 |
| CKAP2L | -0.836180729 | 3.67E-10 | 1.88E-09 |
| CKAP4 | -0.848583814 | 4.80E-25 | 7.50E-24 |
| CKAP5 | -1.278145868 | 9.40E-16 | 7.78E-15 |
| CKB | -1.0417029 | 1.11E-29 | 2.30E-28 |
| CKLF | 0.835613619 | 5.21E-08 | 2.18E-07 |
| CKS1B | 1.2661563 | 6.76E-10 | 3.39E-09 |
| CKS2 | -2.511673891 | 0.003522566 | 0.007621573 |
| CLASP1 | -0.392675739 | 0.020266307 | 0.037088951 |
| CLASP2 | -0.410562218 | 2.19E-05 | 6.66E-05 |
| CLBA1 | 1.165587159 | 4.46E-33 | 1.10E-31 |
| CLCC1 | -0.772567741 | 3.29E-10 | 1.69E-09 |
| CLCF1 | 1.268449363 | 1.23E-11 | 7.28E-11 |
| CLCN5 | -0.560240302 | 7.60E-06 | 2.47E-05 |
| CLCN6 | -0.831248055 | 5.07E-05 | 0.000147858 |
| CLCN7 | -0.761328855 | 3.45E-17 | 3.22E-16 |
| CLDN1 | -0.613780424 | 0.006237808 | 0.012772566 |
| CLDN12 | -1.072851185 | 2.98E-14 | 2.21E-13 |
| CLDN20 | 3.655090127 | 1.65E-14 | 1.25E-13 |
| CLDN5 | -0.45265122 | 4.36E-06 | 1.46E-05 |
| CLDND1 | 0.75252734 | 2.03E-22 | 2.67E-21 |
| CLEC3A | 0.946012806 | 0.005123096 | 0.010686482 |
| CLEC3B | 1.410280306 | 0.000904039 | 0.002172124 |
| CLIC3 | 0.417481726 | 0.01333924 | 0.025478774 |
| CLIC4 | 0.494650326 | 1.84E-11 | 1.07E-10 |
| CLINT1 | -0.627919758 | 9.76E-14 | 6.94E-13 |
| CLIP1 | -0.636419139 | 3.72E-08 | 1.58E-07 |
| CLIP2 | -0.348154732 | 0.007597163 | 0.015282025 |
| CLIP4 | -0.638815581 | 0.02728047 | 0.048438426 |
| CLK3 | -0.972043276 | 1.03E-14 | 7.88E-14 |
| CLK4 | -1.118000534 | 6.84E-20 | 7.62E-19 |
| CLMN | 0.651643068 | 0.021986507 | 0.0399306 |
| CLN5 | 0.345307813 | 0.00287625 | 0.006342309 |
| CLN6 | 1.077461171 | 1.37E-34 | 3.67E-33 |
| CLN8 | 0.223124824 | 0.011695463 | 0.022586635 |
| CLP1 | -0.921680518 | 1.14E-13 | 8.03E-13 |
| CLPTM1L | 0.346913675 | 4.58E-06 | 1.53E-05 |
| CLPX | -1.0943364 | 6.44E-28 | 1.20E-26 |
| CLSPN | 1.268030702 | 8.96E-05 | 0.00025108 |
| CLSTN1 | -0.450363826 | 9.52E-06 | 3.05E-05 |
| CLSTN3 | -0.685716983 | 3.23E-08 | 1.38E-07 |

| | | | |
|---|---|---|---|
| CLTA | -0.305879327 | 0.011065387 | 0.021470483 |
| CLTB | 0.516837319 | 3.28E-10 | 1.69E-09 |
| CLTC | -1.554227678 | 8.79E-67 | 1.05E-64 |
| CLTCL1 | -0.638484911 | 4.89E-06 | 1.63E-05 |
| CLUAP1 | -0.396500839 | 0.001869945 | 0.004257335 |
| CLUH | -0.566513429 | 1.49E-05 | 4.62E-05 |
| CMAS | -0.424566087 | 0.004610629 | 0.009715918 |
| CMC2 | 0.783265872 | 9.22E-06 | 2.95E-05 |
| CMC4 | 0.921129515 | 2.44E-08 | 1.05E-07 |
| CMIP | 0.498924159 | 1.10E-05 | 3.48E-05 |
| CMKLR1 | 1.598138253 | 9.23E-16 | 7.65E-15 |
| CMPK1 | 0.256099758 | 0.003607312 | 0.007788815 |
| CMPK2 | 3.152569322 | 3.12E-55 | 2.22E-53 |
| CMSS1 | 0.447749021 | 0.000204926 | 0.000543802 |
| CMTM3 | 1.011948445 | 1.31E-41 | 5.14E-40 |
| CMTM4 | 1.09778414 | 1.31E-36 | 4.05E-35 |
| CMTM6 | -0.698040169 | 0.014571287 | 0.027557108 |
| CMTR2 | -1.052061077 | 0.013563228 | 0.025841238 |
| CNBP | 0.750106393 | 2.51E-18 | 2.54E-17 |
| CNDP2 | -1.007036243 | 4.75E-25 | 7.43E-24 |
| CNEP1R1 | -0.320940411 | 0.003779897 | 0.008120176 |
| CNIH1 | 0.901232051 | 2.47E-30 | 5.27E-29 |
| CNIH4 | 0.639327491 | 7.46E-09 | 3.39E-08 |
| CNKSR2 | -2.245820103 | 0.002674712 | 0.005934489 |
| CNN2 | 0.733173471 | 8.38E-16 | 6.97E-15 |
| CNN3 | 0.326758357 | 8.41E-05 | 0.000237225 |
| CNOT10 | -1.006673888 | 4.70E-15 | 3.69E-14 |
| CNOT11 | -0.323616377 | 0.000460518 | 0.001159741 |
| CNOT2 | -0.445505767 | 2.18E-05 | 6.63E-05 |
| CNOT4 | 0.417318731 | 0.001098935 | 0.002612908 |
| CNOT6 | -0.824275784 | 1.17E-06 | 4.22E-06 |
| CNOT7 | 0.851305429 | 5.49E-28 | 1.03E-26 |
| CNOT9 | 0.325223077 | 6.07E-05 | 0.000174871 |
| CNP | -0.887422932 | 4.80E-23 | 6.50E-22 |
| CNPPD1 | -0.837275729 | 3.64E-24 | 5.31E-23 |
| CNPY3 | 0.304295007 | 0.005734836 | 0.011833596 |
| CNR2 | 3.427445979 | 0.001623247 | 0.003739521 |
| CNRIP1 | -2.261245753 | 0.009701674 | 0.019047401 |
| CNST | -0.689807474 | 0.007216836 | 0.014575221 |
| CNTD1 | 0.475918085 | 0.005072776 | 0.010597782 |
| CNTNAP1 | -1.954624378 | 0.000787734 | 0.001909896 |
| COA5 | 1.217693116 | 2.21E-34 | 5.88E-33 |
| COA8 | 1.188365146 | 6.89E-41 | 2.60E-39 |
| COASY | -0.978281085 | 0.004685893 | 0.009863816 |
| COBL | -1.528906608 | 5.69E-22 | 7.24E-21 |
| COBLL1 | 0.30804964 | 0.004215508 | 0.008968099 |
| COCH | -2.066162602 | 4.93E-06 | 1.64E-05 |
| COG1 | -1.266676164 | 4.25E-21 | 5.09E-20 |

| | | | |
|---|---|---|---|
| COG2 | -1.096348302 | 3.37E-18 | 3.38E-17 |
| COG3 | -0.656415605 | 1.72E-06 | 6.12E-06 |
| COG4 | -0.44611437 | 1.13E-05 | 3.58E-05 |
| COG5 | -0.370231721 | 0.000536047 | 0.001334669 |
| COG7 | -0.818444247 | 1.62E-10 | 8.57E-10 |
| COL12A1 | -1.156766223 | 7.29E-40 | 2.57E-38 |
| COL18A1 | -1.136875537 | 6.70E-24 | 9.60E-23 |
| COL1A2 | -1.067397068 | 5.40E-36 | 1.60E-34 |
| COL28A1 | 3.19160321 | 0.004085002 | 0.008728714 |
| COL3A1 | -1.019605273 | 4.50E-42 | 1.84E-40 |
| COL4A1 | -1.110990529 | 3.27E-42 | 1.35E-40 |
| COL4A2 | -1.061941692 | 6.84E-36 | 2.01E-34 |
| COL5A1 | -0.372624248 | 5.64E-06 | 1.86E-05 |
| COL5A2 | -0.698983915 | 9.09E-21 | 1.07E-19 |
| COL6A1 | -1.144686781 | 1.51E-45 | 7.32E-44 |
| COL6A2 | -0.703897629 | 6.49E-19 | 6.85E-18 |
| COL6A3 | -1.301731596 | 4.32E-50 | 2.61E-48 |
| COL8A1 | -1.331246861 | 6.25E-25 | 9.70E-24 |
| COL8A2 | -0.704363491 | 5.93E-09 | 2.72E-08 |
| COL9A3 | 1.993675774 | 1.76E-05 | 5.43E-05 |
| COLEC10 | 0.925926431 | 4.10E-12 | 2.54E-11 |
| COLEC11 | 2.331878381 | 4.99E-12 | 3.06E-11 |
| COLEC12 | 1.05766737 | 0.00013261 | 0.000361427 |
| COLGALT1 | -1.494986292 | 6.93E-56 | 5.10E-54 |
| COMMD1 | 0.888610401 | 1.83E-12 | 1.17E-11 |
| COMMD2 | 0.535605837 | 5.61E-09 | 2.58E-08 |
| COMMD4 | -0.318621358 | 0.011713828 | 0.022618888 |
| COMMD7 | 0.687470257 | 1.75E-10 | 9.25E-10 |
| COMMD9 | 0.432856812 | 0.000133542 | 0.000363531 |
| COMT | 0.543852957 | 0.014796036 | 0.02794327 |
| COMTD1 | -0.24801177 | 0.01812158 | 0.033543308 |
| COP1 | -0.816685775 | 3.49E-08 | 1.49E-07 |
| COPA | -1.336447457 | 8.21E-57 | 6.24E-55 |
| COPB1 | -1.455216862 | 5.38E-63 | 5.37E-61 |
| COPB2 | -0.407454675 | 7.02E-07 | 2.61E-06 |
| COPE | -0.260042916 | 0.021546571 | 0.039204951 |
| COPG1 | -0.603822846 | 7.25E-14 | 5.21E-13 |
| COPG2 | -0.995185297 | 6.20E-25 | 9.63E-24 |
| COPS2 | 0.479280966 | 2.91E-08 | 1.25E-07 |
| COPS4 | -0.421112934 | 4.63E-05 | 0.000135745 |
| COPS7A | 0.555684642 | 1.52E-10 | 8.06E-10 |
| COPS7B | 0.419349941 | 0.000302339 | 0.000783786 |
| COPS9 | 1.082862358 | 4.30E-22 | 5.51E-21 |
| COQ2 | -0.86803184 | 4.72E-06 | 1.58E-05 |
| COQ3 | 0.828743612 | 5.42E-12 | 3.30E-11 |
| COQ4 | 0.584024328 | 0.003923251 | 0.008400581 |
| COQ5 | -0.291651688 | 0.020740562 | 0.037856217 |
| COQ6 | -0.713523004 | 1.37E-06 | 4.91E-06 |

| | | | |
|---|---|---|---|
| COQ7 | 0.285338452 | 0.014254635 | 0.027018407 |
| COQ8A | -0.355015261 | 0.024427688 | 0.043912331 |
| CORIN | 3.305244624 | 0.026857684 | 0.047743871 |
| CORO2B | 1.768047936 | 0.022630914 | 0.040953187 |
| CORO7 | 0.572363768 | 5.37E-07 | 2.02E-06 |
| COTL1 | 1.805905775 | 4.09E-109 | 1.79E-106 |
| COX10 | -0.590709537 | 0.00019323 | 0.000514774 |
| COX11 | -0.42073645 | 1.32E-05 | 4.14E-05 |
| COX14 | 0.489320327 | 0.00535443 | 0.011128044 |
| COX15 | -0.36865318 | 0.009945373 | 0.019494881 |
| COX16 | -0.731282481 | 0.000179651 | 0.000481147 |
| COX18 | -0.765838825 | 4.07E-09 | 1.89E-08 |
| COX19 | 0.458866862 | 8.13E-06 | 2.63E-05 |
| COX20 | -0.699788668 | 1.74E-09 | 8.41E-09 |
| COX5A | 0.705647478 | 1.63E-16 | 1.44E-15 |
| COX5B | 0.805145187 | 3.46E-07 | 1.33E-06 |
| COX6A1 | 0.997602622 | 1.05E-30 | 2.29E-29 |
| COX6C | 0.477897602 | 1.60E-05 | 4.94E-05 |
| COX7A2 | 0.301706176 | 0.022332133 | 0.04050959 |
| COX7A2L | 0.949078211 | 1.52E-21 | 1.88E-20 |
| COX7B | 0.392649775 | 0.001643424 | 0.003782156 |
| COX7C | 1.272246171 | 6.20E-27 | 1.09E-25 |
| CP | -0.679306589 | 5.59E-07 | 2.10E-06 |
| CPAMD8 | -2.265783184 | 1.56E-45 | 7.56E-44 |
| CPB1 | 0.967149153 | 0.000312847 | 0.000808715 |
| CPD | -1.778074795 | 5.09E-49 | 3.01E-47 |
| CPE | -0.594753873 | 6.61E-05 | 0.0001891 |
| CPEB1 | -1.032487674 | 1.75E-05 | 5.39E-05 |
| CPEB2 | -1.643464835 | 1.17E-06 | 4.24E-06 |
| CPED1 | 0.841898484 | 0.003073118 | 0.006735981 |
| CPLANE1 | -0.320885775 | 0.027632855 | 0.048974542 |
| CPLANE2 | 0.380886236 | 0.023612758 | 0.042576891 |
| CPN2 | 1.388219441 | 0.00421699 | 0.008969848 |
| CPNE2 | 0.378904424 | 0.001592258 | 0.00367311 |
| CPNE3 | -0.481562666 | 9.44E-07 | 3.45E-06 |
| CPNE8 | -0.38658418 | 0.000119699 | 0.000328744 |
| CPPED1 | 0.436368229 | 0.001539095 | 0.003560745 |
| CPQ | -0.435253362 | 0.00172123 | 0.003945853 |
| CPSF2 | -0.331378059 | 0.000330424 | 0.000852044 |
| CPSF3 | -0.329736649 | 0.000142453 | 0.000386549 |
| CPSF4 | 1.130861826 | 6.04E-35 | 1.67E-33 |
| CPT1A | -1.077711775 | 3.80E-13 | 2.59E-12 |
| CPT2 | -0.961398528 | 3.66E-18 | 3.67E-17 |
| CPZ | -1.354281433 | 4.61E-21 | 5.50E-20 |
| CR1L | -1.360860242 | 3.17E-18 | 3.19E-17 |
| CRABP-I | -1.619727167 | 6.25E-10 | 3.14E-09 |
| CRACD | 1.13510621 | 5.67E-06 | 1.87E-05 |
| CRACR2B | -1.21333185 | 0.000446809 | 0.001128814 |

| | | | |
|---|---|---|---|
| CRADD | 1.159396888 | 1.34E-11 | 7.92E-11 |
| CRAT | 0.276197576 | 0.024054168 | 0.043315349 |
| CRB2 | 1.221916015 | 0.001240081 | 0.002920163 |
| CRCP | 0.56035062 | 2.61E-07 | 1.02E-06 |
| CREB1 | 0.754254448 | 4.59E-06 | 1.53E-05 |
| CREB3 | -0.727102517 | 2.77E-06 | 9.59E-06 |
| CREB3L1 | 0.902183471 | 1.77E-24 | 2.65E-23 |
| CREB3L2 | 0.289841888 | 0.001663026 | 0.003825326 |
| CREB3L3 | 2.109604113 | 0.012311951 | 0.023702059 |
| CREBBP | 0.436595672 | 0.000212213 | 0.000562371 |
| CREBL2 | 0.69251953 | 6.77E-10 | 3.39E-09 |
| CREG1 | -0.350604108 | 0.000637015 | 0.001567966 |
| CRELD1 | 0.612416567 | 4.35E-13 | 2.94E-12 |
| CRELD2 | -0.384131459 | 0.000480498 | 0.001204079 |
| CREM | 0.530088064 | 9.46E-11 | 5.13E-10 |
| CRHBP | 1.668719459 | 4.99E-07 | 1.89E-06 |
| CRIM1 | -1.170044193 | 5.85E-32 | 1.37E-30 |
| CRIP1 | 1.718140967 | 3.42E-39 | 1.19E-37 |
| CRIP2 | 0.661936192 | 1.84E-06 | 6.51E-06 |
| CRLF1 | -0.362354768 | 0.000110987 | 0.000306612 |
| CRMP1 | -1.335890938 | 0.001408911 | 0.003281906 |
| CRNKL1 | -0.626200246 | 9.12E-12 | 5.47E-11 |
| CRTAM | 3.689130973 | 0.00322427 | 0.007037783 |
| CRTAP | -0.696960351 | 1.02E-12 | 6.64E-12 |
| CRY1 | -0.180992971 | 0.020634528 | 0.03768671 |
| CRY2 | -0.725940346 | 6.25E-10 | 3.14E-09 |
| CRYBA1 | 1.225113483 | 0.000140019 | 0.000380247 |
| CRYBB2 | 2.321213345 | 0.000127436 | 0.000348513 |
| CRYBB3 | 4.704510804 | 2.90E-12 | 1.82E-11 |
| CRYGN | 2.825323506 | 8.05E-14 | 5.75E-13 |
| CRYL1 | 0.321540029 | 0.027589094 | 0.048916119 |
| CRYZL1 | -0.507837719 | 4.02E-05 | 0.000118518 |
| CSDC2 | 2.265432955 | 4.03E-11 | 2.26E-10 |
| CSDE1 | 0.688787471 | 1.97E-17 | 1.87E-16 |
| CSE1L | -0.49626839 | 4.70E-08 | 1.98E-07 |
| CSF1R | -1.191283195 | 0.001299505 | 0.003043748 |
| CSF2 | 2.094246974 | 1.89E-07 | 7.49E-07 |
| CSF2RA | 0.898758608 | 1.44E-12 | 9.24E-12 |
| CSGALNACT2 | 0.283418142 | 0.00116587 | 0.002757833 |
| CSMD1 | 1.22173322 | 0.014743113 | 0.027858806 |
| CSNK1A1 | 0.745140734 | 1.56E-24 | 2.34E-23 |
| CSNK1D | -0.406313751 | 8.27E-07 | 3.05E-06 |
| CSNK2A2 | 1.294749601 | 2.71E-37 | 8.71E-36 |
| CSRNP1 | 0.215618992 | 0.010446103 | 0.020382348 |
| CSRP1 | 1.467723003 | 1.49E-40 | 5.45E-39 |
| CSRP2 | 0.795216671 | 0.000632061 | 0.001556617 |
| CSTA | 1.089847615 | 4.34E-15 | 3.42E-14 |
| CSTB | 0.414016012 | 0.001261131 | 0.002962545 |

| | | | |
|---|---|---|---|
| CSTF1 | -0.574520423 | 2.17E-07 | 8.51E-07 |
| CSTF2 | -0.689934736 | 2.37E-11 | 1.36E-10 |
| CSTF3 | 1.50139284 | 9.69E-25 | 1.48E-23 |
| CTAGE1 | 7.456539433 | 4.05E-09 | 1.89E-08 |
| CTBP1 | 0.379079718 | 1.71E-07 | 6.81E-07 |
| CTBP2 | 0.863283697 | 5.19E-20 | 5.84E-19 |
| CTBPL | 0.422188223 | 6.37E-07 | 2.38E-06 |
| CTDSPL | 1.492571529 | 7.32E-25 | 1.13E-23 |
| CTDSPL2 | 0.293939923 | 0.009747615 | 0.019126547 |
| CTH | -0.783102882 | 9.72E-12 | 5.80E-11 |
| CTHRC1 | 0.23996221 | 0.026268765 | 0.046807164 |
| CTNNA1 | -0.908022948 | 3.99E-23 | 5.44E-22 |
| CTNNB1 | -1.131832102 | 5.03E-40 | 1.78E-38 |
| CTNNBIP1 | 0.533945584 | 1.02E-09 | 5.01E-09 |
| CTNNBL1 | -0.388774769 | 0.00033443 | 0.000861558 |
| CTNND1 | -0.86609415 | 7.23E-18 | 7.05E-17 |
| CTPS1 | 0.462927264 | 4.11E-07 | 1.57E-06 |
| CTPS2 | -0.83516684 | 3.18E-05 | 9.50E-05 |
| CTR9 | -1.014844864 | 4.55E-17 | 4.20E-16 |
| CTRL | 4.11961115 | 0.006648397 | 0.013543916 |
| CTSC | -0.648637045 | 5.21E-14 | 3.80E-13 |
| CTSD | -1.048888597 | 8.93E-41 | 3.34E-39 |
| CTSEAL | 4.745247794 | 0.004917531 | 0.010300863 |
| CTSH | -0.852778725 | 1.20E-16 | 1.07E-15 |
| CTSK | -0.989261634 | 3.37E-24 | 4.93E-23 |
| CTSO | -0.743374768 | 2.39E-09 | 1.14E-08 |
| CTSS | -1.084244529 | 2.31E-18 | 2.35E-17 |
| CTSV | -0.704799062 | 2.36E-10 | 1.24E-09 |
| CTSZ | 0.190886346 | 0.009761241 | 0.019150518 |
| CTTNBP2NL | -1.035236417 | 5.38E-14 | 3.91E-13 |
| CTU2 | -0.317291822 | 0.004157954 | 0.008860915 |
| CUEDC2 | -0.811700944 | 1.48E-07 | 5.95E-07 |
| CUL1 | -0.581423449 | 1.38E-10 | 7.35E-10 |
| CUL2 | -0.231858755 | 0.011355988 | 0.021981 |
| CUL3 | -1.114779113 | 2.52E-22 | 3.29E-21 |
| CUL4A | -0.600127819 | 4.40E-07 | 1.67E-06 |
| CUX1 | 0.710940171 | 2.28E-13 | 1.58E-12 |
| CWC15 | 0.74598773 | 1.09E-15 | 8.98E-15 |
| CWC22 | -0.574538072 | 2.03E-07 | 7.99E-07 |
| CWH43 | 0.452059502 | 0.026935065 | 0.047862649 |
| CX3CL1 | 1.786691483 | 2.23E-24 | 3.30E-23 |
| CXCL12 | 0.968260812 | 9.56E-20 | 1.06E-18 |
| CXCL14 | 1.380138661 | 1.27E-63 | 1.31E-61 |
| CXCR1 | -2.334589513 | 1.55E-06 | 5.52E-06 |
| CXCR4 | -0.626105061 | 7.72E-09 | 3.50E-08 |
| CXCR7 | -1.698176608 | 4.12E-25 | 6.50E-24 |
| CXXC5 | 1.575247026 | 2.55E-24 | 3.77E-23 |
| CYB561A3 | 0.526300411 | 0.00021895 | 0.000578869 |

| | | | |
|---|---|---|---|
| CYB561D2 | -0.385939769 | 0.027653011 | 0.049003875 |
| CYB5A | 0.284154494 | 0.01522896 | 0.028689123 |
| CYB5B | 0.686119974 | 4.52E-12 | 2.78E-11 |
| CYB5D2 | -0.359135875 | 0.000322257 | 0.000831774 |
| CYB5R1 | 0.398560863 | 9.78E-06 | 3.13E-05 |
| CYB5R2 | 0.34026946 | 0.002467459 | 0.005503397 |
| CYB5R4 | -0.40321482 | 0.000753418 | 0.001832581 |
| CYCS | 0.701087533 | 3.65E-13 | 2.49E-12 |
| CYFIP1 | -1.229517944 | 2.46E-43 | 1.07E-41 |
| CYP11A1 | -1.173167791 | 1.20E-05 | 3.79E-05 |
| CYP1A1 | -1.308172925 | 0.000866969 | 0.002087854 |
| CYP1C1 | 1.376021821 | 0.010103944 | 0.019760116 |
| CYP24A1 | 0.597562948 | 0.000311077 | 0.000804445 |
| CYP2C23a | -2.300515797 | 6.82E-13 | 4.52E-12 |
| CYP2J21 | -0.849361775 | 4.76E-08 | 2.00E-07 |
| CYP2J22 | -1.050106948 | 4.50E-19 | 4.79E-18 |
| CYP2J23 | -0.937663304 | 2.52E-05 | 7.63E-05 |
| CYP2R1 | -0.547253693 | 4.72E-06 | 1.57E-05 |
| CYP2U1 | -1.595116617 | 5.46E-14 | 3.96E-13 |
| CYP2W1 | 1.051928059 | 0.016636457 | 0.031039495 |
| CYP3A5 | 0.70916448 | 0.027876608 | 0.049355069 |
| CYP4A22 | -0.944572093 | 0.000497346 | 0.001244463 |
| CYP4V2 | -2.171720805 | 4.43E-42 | 1.81E-40 |
| CYP51A1 | -1.232380804 | 1.38E-29 | 2.83E-28 |
| CYRIB | 0.42496254 | 8.33E-06 | 2.69E-05 |
| CYSLTR1 | -1.922295806 | 0.005459608 | 0.011331042 |
| CYSLTR2 | 2.449839724 | 0.01090368 | 0.021196055 |
| CYTH1 | 0.68292079 | 6.46E-13 | 4.29E-12 |
| CYTH3 | -1.175798695 | 8.18E-13 | 5.37E-12 |
| CYTL1 | 0.457263062 | 0.000600469 | 0.001483119 |
| CZIB | 0.455669328 | 6.67E-05 | 0.000190606 |
| DAAM1 | -0.767815415 | 0.006518046 | 0.013304292 |
| DAB2 | 0.746257764 | 1.23E-18 | 1.28E-17 |
| DACT1 | -1.141493729 | 5.14E-30 | 1.08E-28 |
| DAD1 | 1.031440949 | 1.29E-30 | 2.80E-29 |
| DAG1 | -0.64140971 | 0.000138202 | 0.000375463 |
| DAGLA | 0.772888232 | 0.007587741 | 0.015265332 |
| DAGLB | -0.38683651 | 0.009339975 | 0.018382413 |
| DALRD3 | -1.183679562 | 4.19E-15 | 3.31E-14 |
| DAO | 0.694926823 | 0.000819483 | 0.001981571 |
| DAP | 1.093909746 | 4.36E-27 | 7.73E-26 |
| DAP3 | -1.583186958 | 8.01E-37 | 2.52E-35 |
| DAPK1 | -0.791174099 | 9.04E-08 | 3.70E-07 |
| DAPL1 | 1.268980075 | 0.005957223 | 0.012251563 |
| DARS | -0.771737292 | 1.36E-15 | 1.11E-14 |
| DARS2 | -1.098455056 | 3.85E-07 | 1.47E-06 |
| DAZAP1 | 0.424979438 | 0.00153224 | 0.003546698 |
| DAZAP2 | 0.421028042 | 0.022889044 | 0.041392747 |

| | | | |
|---|---|---|---|
| DBF4 | -0.457587344 | 0.000115824 | 0.000318616 |
| DBI | 0.896793935 | 4.38E-13 | 2.96E-12 |
| DBN1 | -0.574514093 | 3.28E-05 | 9.77E-05 |
| DBNDD1 | 1.090990681 | 5.92E-06 | 1.95E-05 |
| DBNDD2 | 1.176294899 | 5.48E-42 | 2.22E-40 |
| DBT | 0.550941465 | 1.25E-05 | 3.93E-05 |
| DCAF1 | -0.781558541 | 3.80E-08 | 1.61E-07 |
| DCAF12L2 | 0.553888883 | 1.86E-09 | 8.95E-09 |
| DCAF13 | -0.510372171 | 3.36E-09 | 1.57E-08 |
| DCAF17 | 0.893578989 | 1.08E-12 | 7.02E-12 |
| DCAF5 | 1.277398072 | 1.25E-34 | 3.37E-33 |
| DCAF7 | -1.181607036 | 9.12E-17 | 8.22E-16 |
| DCBLD2 | 0.602696106 | 1.62E-14 | 1.22E-13 |
| DCDC2 | 0.787558266 | 0.016347327 | 0.03056298 |
| DCK | -0.693279046 | 3.00E-09 | 1.41E-08 |
| DCLK2 | -0.378063116 | 0.028104401 | 0.049719512 |
| DCLRE1C | -0.638689493 | 0.001950875 | 0.004426022 |
| DCP1B | -0.885707731 | 4.70E-08 | 1.98E-07 |
| DCP2 | 0.909704768 | 7.07E-13 | 4.69E-12 |
| DCSTAMP | -1.202015386 | 0.011462437 | 0.022177572 |
| DCTD | 0.952082051 | 3.31E-12 | 2.07E-11 |
| DCTN1 | -0.767144849 | 9.44E-07 | 3.45E-06 |
| DCTN2 | -0.39665665 | 8.26E-05 | 0.000232965 |
| DCTN3 | 0.647362142 | 3.11E-09 | 1.46E-08 |
| DCTN4 | -0.688853848 | 7.42E-10 | 3.70E-09 |
| DCTN6 | 0.699221261 | 3.34E-11 | 1.89E-10 |
| DCUN1D1 | -0.885932697 | 6.54E-18 | 6.41E-17 |
| DCUN1D2 | -0.89133028 | 1.54E-06 | 5.50E-06 |
| DCUN1D3 | -0.252989556 | 0.01528493 | 0.028778607 |
| DCUN1D4 | 0.429754029 | 2.74E-05 | 8.25E-05 |
| DCUN1D5 | -0.644925193 | 1.51E-09 | 7.30E-09 |
| DCX | 1.279246757 | 0.027475668 | 0.048734084 |
| DDA1 | 0.502152268 | 4.41E-09 | 2.04E-08 |
| DDAH1 | 0.330173435 | 0.000695359 | 0.001701408 |
| DDB1 | -1.15322638 | 5.88E-33 | 1.44E-31 |
| DDHD2 | -0.75139563 | 4.29E-07 | 1.63E-06 |
| DDO | -1.271590951 | 0.020959596 | 0.038243106 |
| DDOST | -0.498079971 | 5.05E-08 | 2.12E-07 |
| DDR2 | -0.629803654 | 3.43E-13 | 2.34E-12 |
| DDRGK1 | -0.280700223 | 0.005084521 | 0.010614161 |
| DDTL | 0.963362146 | 1.07E-26 | 1.85E-25 |
| DDX10 | -0.457528128 | 0.000104193 | 0.000289079 |
| DDX20 | -0.896814423 | 5.81E-10 | 2.93E-09 |
| DDX23 | -0.529324277 | 5.65E-08 | 2.36E-07 |
| DDX24 | -0.532568622 | 3.92E-05 | 0.000115939 |
| DDX28 | -1.161778393 | 4.20E-09 | 1.95E-08 |
| DDX3X | -1.272014952 | 2.87E-34 | 7.58E-33 |
| DDX41 | -0.812463543 | 1.30E-13 | 9.16E-13 |

| | | | |
|---|---|---|---|
| DDX42 | -1.387714745 | 9.47E-25 | 1.45E-23 |
| DDX47 | 0.235838071 | 0.011836039 | 0.022838649 |
| DDX49 | -1.211484681 | 2.40E-34 | 6.37E-33 |
| DDX5 | -0.676285428 | 7.69E-18 | 7.48E-17 |
| DDX51 | 0.322612453 | 0.001469521 | 0.003413728 |
| DDX52 | -0.505971832 | 3.25E-05 | 9.69E-05 |
| DDX54 | -0.642478349 | 5.66E-07 | 2.13E-06 |
| DDX55 | -0.753490015 | 9.11E-07 | 3.33E-06 |
| DECR1 | -0.689099343 | 1.07E-07 | 4.35E-07 |
| DECR2 | 0.432463543 | 2.70E-07 | 1.05E-06 |
| DEDD | 1.012603646 | 2.86E-17 | 2.68E-16 |
| DEF8 | -0.754409229 | 1.44E-06 | 5.16E-06 |
| DEGS1 | -0.388971674 | 9.22E-06 | 2.95E-05 |
| DEK | -0.524320484 | 2.77E-05 | 8.33E-05 |
| DENND1A | 0.512072285 | 0.000101107 | 0.000281031 |
| DENND2A | -0.321710307 | 0.02235605 | 0.040542153 |
| DENND2B | -0.359913335 | 0.017121966 | 0.031879694 |
| DENND3 | -1.714296263 | 8.90E-11 | 4.83E-10 |
| DENND4A | -0.532649617 | 0.006452113 | 0.013177629 |
| DENND4C | -0.585726156 | 0.007014744 | 0.014217775 |
| DENND6B | -1.036115165 | 3.59E-06 | 1.22E-05 |
| DEPDC1B | -1.703750849 | 0.012060472 | 0.023248608 |
| DEPDC6 | -1.424119651 | 0.010020813 | 0.019617296 |
| DEPDC7 | -1.134548839 | 2.49E-16 | 2.17E-15 |
| DERA | -0.276158698 | 0.022394506 | 0.040601057 |
| DERL2 | 0.777565579 | 1.11E-15 | 9.10E-15 |
| DERL3 | 0.569538345 | 1.95E-06 | 6.87E-06 |
| DESI1 | 0.431757489 | 8.01E-08 | 3.30E-07 |
| DESI2 | 0.794506999 | 5.67E-19 | 6.00E-18 |
| DEUP1 | 3.334080587 | 0.001352155 | 0.003158359 |
| DFFA | 0.404647971 | 0.000441142 | 0.001115122 |
| DFFB | 0.433258271 | 0.023958321 | 0.043154188 |
| DFNA5 | -0.536502239 | 2.33E-05 | 7.08E-05 |
| DGAT2 | -1.352085749 | 9.83E-05 | 0.000273791 |
| DGCR8 | 0.894854325 | 1.82E-15 | 1.47E-14 |
| DGKB | 0.626982083 | 3.01E-06 | 1.03E-05 |
| DGKD | 0.364670684 | 0.0034786 | 0.007536043 |
| DGKI | 1.072030579 | 0.016532714 | 0.030862882 |
| DGLUCY | 1.113465992 | 0.000519646 | 0.001297635 |
| DHCR24 | -1.723808305 | 4.01E-33 | 9.95E-32 |
| DHCR7 | -0.340186025 | 0.001497317 | 0.003472959 |
| DHDDS | -1.372577298 | 5.39E-46 | 2.67E-44 |
| DHFR | 0.423859352 | 0.000532133 | 0.001325894 |
| DHODH | -0.688467548 | 8.20E-11 | 4.48E-10 |
| DHRS11 | -0.69647015 | 0.000180241 | 0.000482632 |
| DHRS4 | 0.796916543 | 1.23E-10 | 6.61E-10 |
| DHRS7 | -0.811174674 | 2.12E-10 | 1.11E-09 |
| DHRS7B | -0.25067111 | 0.016216558 | 0.030347716 |

| | | | |
|---|---|---|---|
| DHTKD1 | -0.736251973 | 0.001039304 | 0.002480024 |
| DHX15 | -0.295062678 | 0.001187014 | 0.002804432 |
| DHX30 | -0.751692796 | 1.24E-12 | 8.03E-12 |
| DHX38 | -0.602657244 | 1.20E-09 | 5.86E-09 |
| DHX40 | -0.479771513 | 0.000159065 | 0.00042897 |
| DHX58 | 0.540213669 | 0.019073886 | 0.035143318 |
| DHX8 | -0.84899656 | 1.55E-11 | 9.09E-11 |
| DIABLO | 0.606904459 | 1.04E-09 | 5.09E-09 |
| DIAPH2 | -0.671292037 | 1.83E-11 | 1.06E-10 |
| DIEXF | -0.756447218 | 2.27E-08 | 9.84E-08 |
| DIMT1 | 0.388742327 | 0.002198748 | 0.004947091 |
| DIP2A | -0.621673707 | 2.68E-05 | 8.08E-05 |
| DIP2C | -0.644441943 | 0.002233796 | 0.005019302 |
| DIPK1B | 0.997283938 | 0.000242016 | 0.000637003 |
| DIPK2A | -1.813337549 | 8.62E-46 | 4.23E-44 |
| DIPK2B | 2.534501148 | 1.37E-06 | 4.92E-06 |
| DIS3L | 2.901543068 | 2.19E-226 | 4.96E-223 |
| DIS3L2 | -0.355498293 | 0.013758947 | 0.026177436 |
| DISP1 | -1.519176595 | 1.10E-15 | 9.06E-15 |
| DLAT | -0.818967149 | 2.28E-14 | 1.71E-13 |
| DLC1 | 0.475664877 | 2.92E-06 | 1.01E-05 |
| DLD | -0.778165358 | 1.30E-18 | 1.34E-17 |
| DLEC1 | 0.468203263 | 3.77E-05 | 0.000111839 |
| DLG5 | -0.671074042 | 0.001998057 | 0.00452551 |
| DLGAP2 | 0.968715335 | 0.000182133 | 0.000487133 |
| DLGAP4 | 0.438381782 | 3.74E-07 | 1.43E-06 |
| DLL1 | 1.695543385 | 3.50E-30 | 7.46E-29 |
| DLST | -0.457983348 | 0.000283003 | 0.00073661 |
| DLX5 | 1.468801129 | 6.42E-30 | 1.34E-28 |
| DMA | 0.610168931 | 2.14E-06 | 7.52E-06 |
| DMAP1 | -0.750054826 | 0.000249411 | 0.000655072 |
| DMB2 | 0.563244141 | 0.011017825 | 0.021393554 |
| DMD | -0.678915752 | 3.29E-05 | 9.80E-05 |
| DMP1 | 1.568274601 | 0.024659756 | 0.044288505 |
| DMRT2 | -2.290200638 | 6.35E-16 | 5.34E-15 |
| DMXL1 | -1.115790875 | 0.000190004 | 0.000506875 |
| DNA2 | -1.089343955 | 0.023861795 | 0.042991717 |
| DNAAF2 | -0.671582729 | 0.025513075 | 0.045586133 |
| DNAH10 | 1.556841712 | 2.83E-11 | 1.62E-10 |
| DNAH12 | 1.766886811 | 1.19E-07 | 4.80E-07 |
| DNAH5 | 2.683343516 | 0.017943333 | 0.033245063 |
| DNAI3 | 0.784426524 | 0.004874214 | 0.010214357 |
| DNAJA2 | -0.275618908 | 0.001116621 | 0.00265056 |
| DNAJA3 | 0.503937972 | 1.67E-08 | 7.36E-08 |
| DNAJA4 | -1.219170946 | 1.73E-47 | 9.30E-46 |
| DNAJB12 | 0.599238778 | 1.32E-11 | 7.76E-11 |
| DNAJB2 | 0.488189467 | 8.09E-07 | 2.99E-06 |
| DNAJB9 | -0.75091845 | 2.76E-10 | 1.43E-09 |

| | | | |
|---|---|---|---|
| DNAJC1 | 0.531039922 | 9.77E-07 | 3.56E-06 |
| DNAJC12 | 1.172547012 | 2.77E-17 | 2.60E-16 |
| DNAJC13 | -0.94274148 | 7.17E-09 | 3.27E-08 |
| DNAJC16 | 0.603516093 | 9.19E-07 | 3.36E-06 |
| DNAJC17 | -1.234544169 | 5.27E-12 | 3.22E-11 |
| DNAJC18 | 0.968784125 | 2.51E-22 | 3.27E-21 |
| DNAJC19 | 0.503483841 | 0.000111301 | 0.00030738 |
| DNAJC2 | 0.920273155 | 2.63E-19 | 2.84E-18 |
| DNAJC24 | 0.654286803 | 0.000192267 | 0.000512511 |
| DNAJC27 | -0.782986989 | 3.13E-06 | 1.07E-05 |
| DNAJC28 | -0.939646211 | 0.007193105 | 0.014533772 |
| DNAJC3 | -0.464440286 | 1.17E-06 | 4.23E-06 |
| DNAJC8 | 0.352970037 | 0.000230929 | 0.00060912 |
| DNAJC9 | -0.396318495 | 0.002197952 | 0.004946442 |
| DNAL1 | 0.7270048 | 4.71E-05 | 0.000138047 |
| DNAL4 | 0.29526707 | 0.003484837 | 0.007547149 |
| DNASE1L3 | 3.347295488 | 3.40E-09 | 1.59E-08 |
| DNASE2B | 2.182213196 | 0.000114971 | 0.000316332 |
| DND1 | -1.301321113 | 0.009970932 | 0.019536529 |
| DNER | -1.461485766 | 1.79E-41 | 6.97E-40 |
| DNLZ | 0.981403073 | 5.56E-14 | 4.03E-13 |
| DNMBP | -1.469432845 | 3.98E-13 | 2.70E-12 |
| DNMT3A | 0.51675945 | 3.62E-09 | 1.69E-08 |
| DNPEP | 0.555871853 | 1.39E-09 | 6.73E-09 |
| DOC2B | -0.385364407 | 0.015218868 | 0.028674086 |
| DOCK1 | -0.606041915 | 3.06E-05 | 9.17E-05 |
| DOCK10 | -0.576152106 | 0.00604428 | 0.012411822 |
| DOCK7 | -0.480588898 | 1.61E-05 | 4.98E-05 |
| DOHH | -0.442077151 | 8.73E-06 | 2.81E-05 |
| DOK1 | -1.213685103 | 5.46E-07 | 2.05E-06 |
| DOK2 | -0.874404547 | 1.88E-11 | 1.09E-10 |
| DOK4 | 2.114685323 | 1.80E-72 | 2.63E-70 |
| DOK7 | 0.878923785 | 2.59E-06 | 8.99E-06 |
| DOLPP1 | -0.264227366 | 0.02678434 | 0.047619719 |
| DOP1A | -0.990878558 | 7.01E-09 | 3.20E-08 |
| DOT1L | 0.68617372 | 1.70E-17 | 1.62E-16 |
| DPAGT1 | 1.256837306 | 2.44E-42 | 1.01E-40 |
| DPCD | 0.4308586 | 0.000910566 | 0.002187032 |
| DPH1 | -0.887998146 | 7.05E-12 | 4.27E-11 |
| DPH2 | -1.061021365 | 2.33E-08 | 1.01E-07 |
| DPH3P1 | -0.944019909 | 2.17E-15 | 1.74E-14 |
| DPH5 | -1.177566154 | 1.10E-15 | 9.05E-15 |
| DPH7 | 0.387422429 | 0.000380797 | 0.000972344 |
| DPP7 | -1.326404276 | 1.14E-62 | 1.11E-60 |
| DPP8 | -0.549722137 | 3.77E-05 | 0.000111687 |
| DPY30 | 0.362609396 | 0.002348003 | 0.005259403 |
| DPYD | -0.462418501 | 0.00666289 | 0.013569374 |
| DPYSL2 | -0.732720933 | 9.47E-17 | 8.52E-16 |

| | | | |
|---|---|---|---|
| DQX1 | -0.335086437 | 0.016322055 | 0.030524129 |
| DR1 | -0.286492351 | 0.002318182 | 0.005194332 |
| DRAM1 | 0.700622043 | 1.52E-06 | 5.41E-06 |
| DRAM2 | 1.486232556 | 2.37E-44 | 1.08E-42 |
| DRAP1 | 0.603200297 | 0.003019243 | 0.006629654 |
| DRG1 | -1.511858553 | 3.80E-41 | 1.45E-39 |
| DRG2 | 0.262041612 | 0.003999041 | 0.008557143 |
| DROSHA | -0.571469632 | 0.000554481 | 0.001377537 |
| DSCC1 | -1.848106768 | 3.28E-05 | 9.79E-05 |
| DSEL | 0.909290222 | 1.38E-05 | 4.30E-05 |
| DSP | 1.841860996 | 1.42E-07 | 5.72E-07 |
| DSTN | 0.859537514 | 3.66E-17 | 3.41E-16 |
| DSTYK | -0.68668289 | 0.000111328 | 0.00030738 |
| DTD1 | -0.419273729 | 0.010781388 | 0.02097933 |
| DTNBP1 | -0.78550496 | 4.66E-07 | 1.77E-06 |
| DTX2 | 0.565796843 | 1.94E-07 | 7.68E-07 |
| DTX4 | 1.253323946 | 8.23E-05 | 0.000232132 |
| DUS2 | -0.73709874 | 2.48E-09 | 1.18E-08 |
| DUSP1 | -0.316554714 | 8.95E-05 | 0.000250756 |
| DUSP10 | -0.9633993 | 3.80E-24 | 5.51E-23 |
| DUSP12 | -1.52452988 | 2.25E-15 | 1.81E-14 |
| DUSP14 | 0.319251701 | 0.001223746 | 0.002884197 |
| DUSP19 | 0.681086223 | 5.10E-15 | 4.00E-14 |
| DUSP23 | 1.248379176 | 1.25E-19 | 1.38E-18 |
| DUSP26 | 1.817894507 | 0.015059938 | 0.028402204 |
| DUSP4 | 0.291816636 | 0.002660459 | 0.005906724 |
| DUSP5 | -0.715726467 | 3.18E-11 | 1.80E-10 |
| DUSP6 | 0.482751006 | 1.15E-05 | 3.63E-05 |
| DUSP7 | -1.528803933 | 1.91E-40 | 6.89E-39 |
| DUT | 0.358485872 | 0.018377638 | 0.033961775 |
| DVL3 | -0.836155535 | 2.69E-10 | 1.40E-09 |
| DYL1 | 0.753283557 | 2.85E-15 | 2.27E-14 |
| DYNC1H1 | -0.29950703 | 0.009124412 | 0.018007697 |
| DYNC1I1 | 1.526349956 | 9.99E-41 | 3.70E-39 |
| DYNC1I2 | -0.734606492 | 2.07E-15 | 1.66E-14 |
| DYNC1LI1 | -0.295547869 | 0.000803756 | 0.001945966 |
| DYNC1LI2 | -0.249078961 | 0.004464795 | 0.009433467 |
| DYNC2H1 | -0.794070431 | 2.38E-05 | 7.24E-05 |
| DYNC2I1 | -1.484477584 | 2.20E-22 | 2.88E-21 |
| DYNC2I2 | -0.98693295 | 5.20E-08 | 2.18E-07 |
| DYNLL2 | 0.849834213 | 1.79E-21 | 2.19E-20 |
| DYNLRB1 | 1.243872951 | 5.89E-34 | 1.54E-32 |
| DYNLT2 | 0.690144352 | 0.000181187 | 0.000484877 |
| DYNLT3 | 0.901358488 | 1.82E-21 | 2.23E-20 |
| DYRK1A | -0.44576466 | 0.006968129 | 0.014129615 |
| DYRK2 | -0.349269479 | 0.005545551 | 0.01148836 |
| DYSF | 0.969197746 | 7.64E-16 | 6.38E-15 |
| DZANK1 | -0.946346183 | 0.000243253 | 0.000639764 |

| | | | |
|---|---|---|---|
| DZIP1L | -1.167854576 | 1.25E-05 | 3.95E-05 |
| E2F4 | 0.527315721 | 1.33E-05 | 4.17E-05 |
| E2F5 | 0.742945964 | 9.07E-15 | 6.98E-14 |
| E2F6 | -0.437670161 | 0.002311226 | 0.005180439 |
| E2F8 | -2.112242673 | 0.000156666 | 0.000423089 |
| E4F1 | 0.403292655 | 0.003778093 | 0.008118668 |
| EAF1 | -0.423234614 | 0.000952165 | 0.002282509 |
| EAF2 | 0.293449333 | 0.006194827 | 0.012690292 |
| EAPP | -0.625567811 | 2.65E-08 | 1.14E-07 |
| EARS2 | -0.883271493 | 6.08E-06 | 2.00E-05 |
| EBAG9 | 1.281470124 | 8.85E-34 | 2.28E-32 |
| EBF4 | 1.568196908 | 0.002541279 | 0.005660614 |
| EBNA1BP2 | 0.345308912 | 0.000651775 | 0.001601106 |
| ECD | 0.748379435 | 3.69E-16 | 3.17E-15 |
| ECHDC3 | -0.409833332 | 0.020539793 | 0.037533862 |
| ECHS1 | -1.377028587 | 8.28E-36 | 2.42E-34 |
| ECI1 | -0.254220764 | 0.004281509 | 0.009089352 |
| ECI2 | 0.26244558 | 0.006794018 | 0.01380746 |
| ECPAS | -0.641858948 | 0.000278662 | 0.000726563 |
| ECRG4 | 1.025050625 | 2.98E-06 | 1.03E-05 |
| ECSIT | -0.8223215 | 3.88E-13 | 2.64E-12 |
| ECT2 | -0.878723718 | 0.000512944 | 0.001282078 |
| EDA | 0.65523136 | 0.000343866 | 0.000884692 |
| EDA2R | -1.179580761 | 7.82E-13 | 5.14E-12 |
| EDC3 | -0.420083846 | 0.003048838 | 0.006689198 |
| EDEM1 | 0.393261379 | 5.89E-06 | 1.94E-05 |
| EDEM3 | -0.408824014 | 0.001244392 | 0.002929808 |
| EDF1 | 0.427117912 | 2.95E-07 | 1.14E-06 |
| EDNRA | 0.338647981 | 8.10E-05 | 0.00022871 |
| EDNRB | -2.792986921 | 0.008179258 | 0.016332081 |
| EDQM3 | -3.485026314 | 0.018996463 | 0.035019656 |
| EEA1 | -0.2858237 | 0.011086411 | 0.021508207 |
| EED | -0.841458225 | 6.05E-10 | 3.05E-09 |
| EEF1A1 | -0.234716975 | 0.000738973 | 0.001800344 |
| EEF1AKMT1 | 0.609952078 | 4.67E-09 | 2.16E-08 |
| EEF1AKNMT | -1.323933119 | 6.91E-15 | 5.37E-14 |
| EEF1B2 | 0.397616753 | 2.66E-05 | 8.02E-05 |
| EEF1D | 0.716699143 | 1.76E-18 | 1.81E-17 |
| EEF2 | -0.969094827 | 8.15E-33 | 1.97E-31 |
| EEF2K | 0.30798806 | 0.025361112 | 0.04535038 |
| EEFSEC | -0.60527946 | 0.000380871 | 0.000972352 |
| EEPD1 | -2.309768085 | 1.46E-43 | 6.44E-42 |
| EFCAB1 | 0.717226552 | 1.47E-06 | 5.26E-06 |
| EFCAB14 | 0.599291589 | 4.63E-08 | 1.95E-07 |
| EFCAB2 | 1.677405691 | 0.007119935 | 0.014398774 |
| EFCAB5 | 0.896259888 | 0.00044759 | 0.001130369 |
| EFEMP1 | -0.338024012 | 0.000167482 | 0.00045015 |
| EFHD2 | -0.974193362 | 1.48E-09 | 7.16E-09 |

| | | | |
|---|---|---|---|
| EFL1 | -0.342191002 | 0.00024579 | 0.000646185 |
| EFNA5 | 0.726938607 | 0.005783149 | 0.011924236 |
| EFNB1 | 1.277718351 | 3.45E-51 | 2.16E-49 |
| EFNB2 | -0.763487535 | 1.20E-12 | 7.80E-12 |
| EFR3A | 0.325042201 | 0.001247837 | 0.002936392 |
| EFR3B | -0.820930015 | 0.0053642 | 0.011144942 |
| EFTUD2 | -0.662246917 | 4.65E-13 | 3.14E-12 |
| EGF | 2.695633011 | 0.001399377 | 0.003260815 |
| EGFL7 | 0.592231944 | 0.002220118 | 0.004990636 |
| EGFLAM | -0.547296871 | 5.93E-09 | 2.72E-08 |
| EGFR | -1.206486781 | 5.83E-19 | 6.16E-18 |
| EGR1 | -1.055705846 | 1.14E-26 | 1.96E-25 |
| EGR2 | -1.477599958 | 1.02E-09 | 5.02E-09 |
| EGR3 | 1.690413953 | 9.08E-07 | 3.32E-06 |
| EGR4 | -0.863761844 | 5.28E-06 | 1.75E-05 |
| EHBP1 | 1.167040955 | 1.44E-43 | 6.38E-42 |
| EHD2 | -0.544885082 | 4.59E-10 | 2.33E-09 |
| EHD3 | 0.51023365 | 2.04E-11 | 1.18E-10 |
| EI24 | 1.412289334 | 3.37E-62 | 3.25E-60 |
| EIF1 | 1.470735946 | 1.53E-76 | 2.81E-74 |
| EIF1B | 0.434413243 | 6.13E-06 | 2.02E-05 |
| EIF2AK3 | -0.627977763 | 1.82E-09 | 8.74E-09 |
| EIF2AK4 | 0.695827823 | 1.51E-05 | 4.69E-05 |
| EIF2B1 | -1.183101031 | 1.60E-20 | 1.84E-19 |
| EIF2B2 | -0.372430896 | 3.94E-05 | 0.000116517 |
| EIF2B3 | -0.478060902 | 0.000494034 | 0.001236631 |
| EIF2B4 | -0.317992938 | 0.003715921 | 0.008002978 |
| EIF2B5 | -0.339325236 | 0.000133367 | 0.000363257 |
| EIF2S1 | -0.440200281 | 2.27E-06 | 7.92E-06 |
| EIF2S3 | -0.385318451 | 3.26E-05 | 9.74E-05 |
| EIF3A | -0.613830938 | 1.34E-11 | 7.90E-11 |
| EIF3D | -0.250481224 | 0.001000846 | 0.002392876 |
| EIF3I | 0.306164429 | 0.000653576 | 0.001604951 |
| EIF3J | 0.99476253 | 6.28E-31 | 1.39E-29 |
| EIF4A2 | -0.26337077 | 0.00094422 | 0.002264262 |
| EIF4A3 | -1.010430515 | 1.36E-30 | 2.95E-29 |
| EIF4B | -0.494905724 | 5.48E-05 | 0.000158909 |
| EIF4E2 | 0.404317781 | 9.53E-05 | 0.000266074 |
| EIF4E3 | 0.606868593 | 5.42E-06 | 1.79E-05 |
| EIF4EBP1 | 1.649230702 | 1.55E-70 | 2.06E-68 |
| EIF4EBP2 | 1.151961817 | 7.81E-34 | 2.02E-32 |
| EIF4G1 | -0.504539766 | 0.000454757 | 0.001146338 |
| EIF4G2 | 1.203202443 | 2.24E-41 | 8.69E-40 |
| EIF4G3 | 0.355421234 | 0.016530493 | 0.030862882 |
| EIF4H | 0.807886136 | 8.14E-25 | 1.25E-23 |
| EIF5 | 0.291987774 | 0.000127259 | 0.000348169 |
| EIF5A2 | 0.730157023 | 4.15E-16 | 3.55E-15 |
| EIF5B | -0.433306295 | 3.45E-06 | 1.17E-05 |

| | | | |
|---|---|---|---|
| ELAC2 | -0.633128438 | 5.09E-07 | 1.92E-06 |
| ELF1 | -0.434657959 | 5.53E-07 | 2.08E-06 |
| ELF3 | 2.765640045 | 3.31E-06 | 1.13E-05 |
| ELFN1 | 1.553500751 | 3.86E-39 | 1.33E-37 |
| ELK3 | 0.964675374 | 1.52E-29 | 3.11E-28 |
| ELK4 | 0.786117044 | 6.90E-05 | 0.000196847 |
| ELL | 1.025815582 | 3.14E-33 | 7.84E-32 |
| ELL2 | 2.001639486 | 5.01E-53 | 3.39E-51 |
| ELMO2 | 0.556685895 | 1.17E-10 | 6.30E-10 |
| ELMO3 | -0.837040123 | 4.55E-05 | 0.000133611 |
| ELMOD2 | 1.138394064 | 1.04E-13 | 7.37E-13 |
| ELOVL4 | -5.622832181 | 2.80E-05 | 8.42E-05 |
| ELOVL7 | -0.495976318 | 0.018386882 | 0.033974238 |
| ELP2 | -0.75866695 | 2.13E-10 | 1.12E-09 |
| ELP3 | -0.646816914 | 1.05E-07 | 4.26E-07 |
| ELP4 | 0.470371481 | 0.024874924 | 0.044598334 |
| ELP6 | -0.455716923 | 0.001632027 | 0.003758474 |
| EMC1 | -1.042477261 | 2.76E-38 | 9.32E-37 |
| EMC6 | 0.798236929 | 3.67E-17 | 3.42E-16 |
| EMC7 | -0.809707663 | 3.44E-15 | 2.73E-14 |
| EMC8 | -0.740110486 | 1.41E-13 | 9.92E-13 |
| EME1 | -0.665804235 | 0.008338396 | 0.01662298 |
| EMG1 | -0.545305758 | 2.06E-07 | 8.12E-07 |
| EMILIN2 | -0.779647095 | 3.93E-21 | 4.72E-20 |
| EML1 | -0.66536813 | 2.98E-06 | 1.03E-05 |
| EMP1 | 0.206859401 | 0.008800354 | 0.017421265 |
| EMP2 | 0.27821461 | 0.005726542 | 0.011818276 |
| EN1 | 1.018902796 | 7.41E-07 | 2.74E-06 |
| ENAH | 0.37840568 | 6.16E-05 | 0.00017713 |
| ENC1 | 0.336626894 | 0.015404327 | 0.028983335 |
| ENDOD1 | -1.131524221 | 2.18E-19 | 2.36E-18 |
| ENDOG | -0.684896343 | 1.47E-08 | 6.52E-08 |
| ENDOV | -1.050947979 | 6.40E-08 | 2.66E-07 |
| ENG | -1.644997624 | 1.30E-28 | 2.52E-27 |
| ENHO | 1.277796099 | 0.001022392 | 0.002441812 |
| ENKUR | -0.651980724 | 0.003178575 | 0.006950321 |
| ENO2 | -1.949689661 | 3.84E-18 | 3.83E-17 |
| ENOPH1 | -0.254440138 | 0.017395214 | 0.032313208 |
| ENOX1 | 0.689979301 | 0.000410385 | 0.001042025 |
| ENOX2 | 0.380389416 | 9.98E-06 | 3.19E-05 |
| ENPP4 | -1.045172811 | 2.21E-11 | 1.27E-10 |
| ENSA | 0.242837896 | 0.002772329 | 0.006123089 |
| ENSGALG00000000071 | -0.909446803 | 7.37E-06 | 2.40E-05 |
| ENSGALG00000000264 | -0.596230379 | 9.27E-05 | 0.000259319 |
| ENSGALG00000000433 | -0.493022484 | 1.02E-05 | 3.25E-05 |
| ENSGALG00000000497 | -1.757309032 | 4.27E-08 | 1.81E-07 |
| ENSGALG00000000544 | 1.080819143 | 8.87E-05 | 0.000249012 |
| ENSGALG00000000584 | 2.240900043 | 6.44E-42 | 2.57E-40 |

| | | | |
|---|---|---|---|
| ENSGALG00000000611 | -0.838674787 | 4.98E-16 | 4.23E-15 |
| ENSGALG00000000629 | 0.893336642 | 6.03E-17 | 5.49E-16 |
| ENSGALG00000000818 | -1.087424001 | 1.59E-27 | 2.91E-26 |
| ENSGALG00000000996 | 0.447198581 | 3.64E-06 | 1.23E-05 |
| ENSGALG00000001003 | 0.698902766 | 1.11E-11 | 6.58E-11 |
| ENSGALG00000001028 | 2.424170188 | 0.002092585 | 0.00472701 |
| ENSGALG00000001035 | 3.726501831 | 3.27E-22 | 4.23E-21 |
| ENSGALG00000001049 | 0.270133704 | 0.007019915 | 0.014226134 |
| ENSGALG00000001109 | 1.174236928 | 6.07E-28 | 1.13E-26 |
| ENSGALG00000001110 | -0.645035045 | 4.14E-05 | 0.000122115 |
| ENSGALG00000001289 | -0.909145934 | 6.88E-07 | 2.56E-06 |
| ENSGALG00000001304 | 0.381893074 | 0.00386505 | 0.008284776 |
| ENSGALG00000001317 | 1.281702665 | 1.06E-65 | 1.23E-63 |
| ENSGALG00000001476 | 1.442329631 | 1.80E-71 | 2.50E-69 |
| ENSGALG00000001483 | 0.556359857 | 8.93E-05 | 0.000250385 |
| ENSGALG00000001525 | 0.388548391 | 7.34E-06 | 2.39E-05 |
| ENSGALG00000001536 | -0.985065172 | 3.25E-06 | 1.11E-05 |
| ENSGALG00000001604 | -0.350251518 | 0.020741263 | 0.037856217 |
| ENSGALG00000001620 | 2.154229938 | 4.20E-45 | 1.99E-43 |
| ENSGALG00000001720 | 0.924292503 | 1.96E-24 | 2.92E-23 |
| ENSGALG00000001800 | -0.65341471 | 1.86E-07 | 7.35E-07 |
| ENSGALG00000001824 | -0.869231528 | 0.005684147 | 0.011741478 |
| ENSGALG00000001921 | -1.601722313 | 9.52E-08 | 3.88E-07 |
| ENSGALG00000001948 | 0.29585129 | 0.018473617 | 0.034129862 |
| ENSGALG00000001972 | -0.417779235 | 9.38E-05 | 0.000262194 |
| ENSGALG00000001983 | 1.331640804 | 0.003826032 | 0.008208908 |
| ENSGALG00000002043 | 0.757798486 | 1.83E-06 | 6.50E-06 |
| ENSGALG00000002046 | 1.608023032 | 6.91E-15 | 5.37E-14 |
| ENSGALG00000002067 | -0.727456099 | 1.04E-08 | 4.66E-08 |
| ENSGALG00000002099 | -0.632284438 | 7.15E-08 | 2.96E-07 |
| ENSGALG00000002108 | 2.587726613 | 3.76E-12 | 2.33E-11 |
| ENSGALG00000002193 | -1.552591066 | 3.13E-14 | 2.31E-13 |
| ENSGALG00000002326 | 0.420991703 | 0.001382639 | 0.003225129 |
| ENSGALG00000002389 | 1.773426468 | 2.18E-06 | 7.63E-06 |
| ENSGALG00000002399 | -0.737866459 | 3.03E-13 | 2.08E-12 |
| ENSGALG00000002470 | -1.135675484 | 0.000469646 | 0.001179495 |
| ENSGALG00000002638 | 1.750919513 | 2.32E-14 | 1.74E-13 |
| ENSGALG00000002957 | 6.776974775 | 5.85E-143 | 5.30E-140 |
| ENSGALG00000002971 | -0.678025436 | 0.000695022 | 0.001700889 |
| ENSGALG00000002988 | -0.814487257 | 2.31E-13 | 1.60E-12 |
| ENSGALG00000003022 | -1.193302609 | 0.003738866 | 0.008040928 |
| ENSGALG00000003025 | 1.039191507 | 0.009209744 | 0.018160284 |
| ENSGALG00000003058 | 5.408178158 | 0.000727304 | 0.001773822 |
| ENSGALG00000003074 | -1.048208811 | 2.94E-12 | 1.84E-11 |
| ENSGALG00000003149 | -0.563955462 | 0.000386716 | 0.000986349 |
| ENSGALG00000003235 | -1.133826419 | 6.43E-07 | 2.40E-06 |
| ENSGALG00000003283 | -0.423169301 | 0.002835189 | 0.006255825 |
| ENSGALG00000003345 | 1.279004164 | 0.013300747 | 0.025408819 |

| | | | |
|---|---|---|---|
| ENSGALG00000003348 | 0.826350714 | 7.55E-12 | 4.56E-11 |
| ENSGALG00000003409 | -1.25147454 | 1.34E-09 | 6.51E-09 |
| ENSGALG00000003412 | -0.454155566 | 0.000310337 | 0.000802835 |
| ENSGALG00000003419 | -0.743698506 | 0.001548887 | 0.00358096 |
| ENSGALG00000003464 | -1.703811495 | 0.001877362 | 0.004272791 |
| ENSGALG00000003575 | 0.634992615 | 3.89E-08 | 1.65E-07 |
| ENSGALG00000003605 | -0.527385701 | 0.00104309 | 0.00248775 |
| ENSGALG00000003644 | -1.446646455 | 0.02492279 | 0.044661217 |
| ENSGALG00000003684 | -0.983473783 | 0.017247813 | 0.032078855 |
| ENSGALG00000003696 | 0.581236335 | 1.47E-07 | 5.90E-07 |
| ENSGALG00000003747 | 0.251896979 | 0.002516783 | 0.005611568 |
| ENSGALG00000003805 | 0.294220852 | 0.025067794 | 0.044891329 |
| ENSGALG00000003836 | 2.023800273 | 0.013903103 | 0.02641473 |
| ENSGALG00000003861 | -1.668623526 | 3.14E-25 | 4.96E-24 |
| ENSGALG00000003920 | -0.457029208 | 0.00012013 | 0.000329792 |
| ENSGALG00000004070 | -0.45407945 | 0.003190603 | 0.006972134 |
| ENSGALG00000004113 | 0.613710337 | 0.000775199 | 0.001882191 |
| ENSGALG00000004127 | -2.383519106 | 1.75E-07 | 6.95E-07 |
| ENSGALG00000004192 | -0.681410519 | 0.001009009 | 0.002410697 |
| ENSGALG00000004231 | 0.340542186 | 0.008100641 | 0.016197553 |
| ENSGALG00000004243 | 1.86862897 | 0.002938584 | 0.006468217 |
| ENSGALG00000004322 | 0.397370828 | 0.000772721 | 0.001876844 |
| ENSGALG00000004329 | 1.4053309 | 7.99E-50 | 4.78E-48 |
| ENSGALG00000004394 | 1.906093473 | 0.000784662 | 0.001903468 |
| ENSGALG00000004400 | -0.806344603 | 2.41E-12 | 1.52E-11 |
| ENSGALG00000004440 | -0.937901993 | 9.08E-19 | 9.47E-18 |
| ENSGALG00000004441 | 3.839421171 | 8.62E-05 | 0.000242547 |
| ENSGALG00000004477 | -0.757146095 | 0.019874098 | 0.036461314 |
| ENSGALG00000004503 | 1.000697995 | 3.71E-12 | 2.30E-11 |
| ENSGALG00000004509 | -1.047635237 | 1.02E-19 | 1.13E-18 |
| ENSGALG00000004518 | -2.30405418 | 3.85E-46 | 1.96E-44 |
| ENSGALG00000004627 | -2.368389466 | 6.85E-25 | 1.06E-23 |
| ENSGALG00000004660 | 1.4441 | 0.000284688 | 0.000740288 |
| ENSGALG00000004721 | 1.786552245 | 1.26E-07 | 5.10E-07 |
| ENSGALG00000004725 | -0.332676253 | 0.000440497 | 0.001113697 |
| ENSGALG00000004734 | -1.5699466 | 0.019177043 | 0.035309451 |
| ENSGALG00000004747 | 1.555060807 | 0.002824687 | 0.006234677 |
| ENSGALG00000004799 | -1.236660546 | 1.21E-15 | 9.90E-15 |
| ENSGALG00000004816 | -0.929928135 | 1.32E-18 | 1.36E-17 |
| ENSGALG00000004825 | 1.166609078 | 1.41E-41 | 5.52E-40 |
| ENSGALG00000004878 | 0.878394118 | 0.000654859 | 0.001607521 |
| ENSGALG00000004881 | -0.757611962 | 5.93E-18 | 5.83E-17 |
| ENSGALG00000004928 | -0.63222059 | 5.71E-05 | 0.000165146 |
| ENSGALG00000004955 | 0.465660953 | 0.002951855 | 0.006495323 |
| ENSGALG00000004975 | -0.352601534 | 0.012925135 | 0.02477834 |
| ENSGALG00000005012 | -0.728755647 | 3.89E-06 | 1.31E-05 |
| ENSGALG00000005020 | 0.439068861 | 0.01015867 | 0.019861427 |
| ENSGALG00000005037 | -0.707712856 | 0.0250914 | 0.044927191 |

| | | | |
|---|---|---|---|
| ENSGALG00000005076 | 1.063473267 | 0.000310051 | 0.000802248 |
| ENSGALG00000005093 | -1.024670568 | 9.57E-08 | 3.90E-07 |
| ENSGALG00000005180 | -1.224671587 | 3.92E-14 | 2.88E-13 |
| ENSGALG00000005181 | -0.76339141 | 2.13E-18 | 2.17E-17 |
| ENSGALG00000005204 | 0.396906281 | 0.000202253 | 0.000537339 |
| ENSGALG00000005291 | -1.31543219 | 1.87E-06 | 6.61E-06 |
| ENSGALG00000005298 | 1.802385555 | 0.00529408 | 0.011017782 |
| ENSGALG00000005350 | -0.999519489 | 2.76E-05 | 8.32E-05 |
| ENSGALG00000005464 | 0.657564134 | 6.33E-09 | 2.89E-08 |
| ENSGALG00000005475 | 0.757709832 | 1.66E-12 | 1.06E-11 |
| ENSGALG00000005648 | -1.601293784 | 5.47E-10 | 2.76E-09 |
| ENSGALG00000005675 | 0.573214577 | 1.34E-09 | 6.51E-09 |
| ENSGALG00000005682 | -0.99292084 | 0.000243474 | 0.00064022 |
| ENSGALG00000005775 | 0.887523495 | 9.90E-05 | 0.000275561 |
| ENSGALG00000005776 | 0.86042899 | 0.000622046 | 0.001533063 |
| ENSGALG00000005778 | 0.594058457 | 0.002713998 | 0.006007925 |
| ENSGALG00000005924 | 1.042672418 | 4.19E-25 | 6.60E-24 |
| ENSGALG00000005929 | 1.439044323 | 0.000104552 | 0.000289955 |
| ENSGALG00000005958 | 1.804596107 | 3.73E-06 | 1.26E-05 |
| ENSGALG00000005964 | 0.868582126 | 0.025764812 | 0.045993607 |
| ENSGALG00000006083 | -0.726044868 | 0.001398719 | 0.003259841 |
| ENSGALG00000006092 | -0.680974525 | 3.46E-12 | 2.16E-11 |
| ENSGALG00000006104 | -0.467020161 | 0.021056431 | 0.038400511 |
| ENSGALG00000006123 | -2.576416831 | 9.98E-46 | 4.86E-44 |
| ENSGALG00000006140 | -0.63944164 | 0.00106853 | 0.002544405 |
| ENSGALG00000006143 | -0.625826936 | 7.91E-11 | 4.33E-10 |
| ENSGALG00000006203 | 1.842797235 | 4.35E-10 | 2.21E-09 |
| ENSGALG00000006272 | 0.82405172 | 0.001711038 | 0.003927123 |
| ENSGALG00000006292 | 0.512070183 | 6.56E-06 | 2.15E-05 |
| ENSGALG00000006325 | -0.287880025 | 0.022445935 | 0.040656336 |
| ENSGALG00000006394 | 4.573929689 | 5.13E-05 | 0.000149346 |
| ENSGALG00000006407 | -0.757058666 | 3.19E-14 | 2.36E-13 |
| ENSGALG00000006422 | -0.555127515 | 0.000298862 | 0.000775364 |
| ENSGALG00000006423 | 0.652117264 | 0.003475871 | 0.007531331 |
| ENSGALG00000006430 | -1.26561888 | 5.14E-12 | 3.14E-11 |
| ENSGALG00000006465 | -0.464071634 | 7.70E-05 | 0.000218 |
| ENSGALG00000006705 | 0.240300992 | 0.015615134 | 0.029339356 |
| ENSGALG00000006724 | 0.392200495 | 7.43E-05 | 0.000211037 |
| ENSGALG00000006838 | -1.047537552 | 1.24E-12 | 8.04E-12 |
| ENSGALG00000006864 | 3.239491779 | 5.18E-06 | 1.72E-05 |
| ENSGALG00000006899 | 0.442711598 | 0.00070951 | 0.001733223 |
| ENSGALG00000006900 | 1.93848217 | 0.026741428 | 0.04756209 |
| ENSGALG00000006976 | -0.525074583 | 0.005981342 | 0.012299305 |
| ENSGALG00000007080 | -1.175598292 | 0.000129818 | 0.000354314 |
| ENSGALG00000007127 | -0.602699704 | 7.23E-11 | 3.97E-10 |
| ENSGALG00000007131 | 0.714154368 | 4.43E-12 | 2.72E-11 |
| ENSGALG00000007171 | 1.687132898 | 0.014560122 | 0.027539825 |
| ENSGALG00000007195 | 0.559141942 | 2.46E-07 | 9.61E-07 |

| | | | |
|---|---|---|---|
| ENSGALG00000007248 | 0.509497543 | 7.33E-05 | 0.000208405 |
| ENSGALG00000007395 | 2.532277583 | 1.65E-14 | 1.25E-13 |
| ENSGALG00000007522 | -2.614023754 | 0.006036232 | 0.012400916 |
| ENSGALG00000007536 | -0.406694338 | 0.008796439 | 0.017416053 |
| ENSGALG00000007550 | -1.068707388 | 2.31E-10 | 1.21E-09 |
| ENSGALG00000007575 | -1.033115074 | 1.21E-08 | 5.42E-08 |
| ENSGALG00000007596 | 2.746520668 | 1.56E-08 | 6.86E-08 |
| ENSGALG00000007645 | 1.713560826 | 0.009329099 | 0.018368072 |
| ENSGALG00000007683 | 0.757557248 | 1.50E-10 | 7.98E-10 |
| ENSGALG00000007699 | 0.710024041 | 3.46E-09 | 1.62E-08 |
| ENSGALG00000007710 | -1.356032854 | 1.62E-08 | 7.13E-08 |
| ENSGALG00000007728 | -1.12101267 | 2.82E-06 | 9.74E-06 |
| ENSGALG00000007841 | 2.473361363 | 8.92E-43 | 3.76E-41 |
| ENSGALG00000008032 | -0.879552302 | 0.013621787 | 0.025945533 |
| ENSGALG00000008054 | -0.247831054 | 0.024795714 | 0.044485658 |
| ENSGALG00000008077 | 0.745518506 | 0.004924174 | 0.010311103 |
| ENSGALG00000008314 | 0.696410712 | 1.33E-06 | 4.79E-06 |
| ENSGALG00000008404 | -1.137660414 | 2.04E-20 | 2.35E-19 |
| ENSGALG00000008469 | -0.955782486 | 1.63E-23 | 2.28E-22 |
| ENSGALG00000008485 | 0.271055296 | 0.01774203 | 0.032916963 |
| ENSGALG00000008597 | 0.791459554 | 1.33E-19 | 1.46E-18 |
| ENSGALG00000008689 | 1.008759475 | 7.21E-06 | 2.35E-05 |
| ENSGALG00000008725 | -0.251888705 | 0.026909255 | 0.047829292 |
| ENSGALG00000008727 | 0.849191687 | 3.37E-05 | 0.000100478 |
| ENSGALG00000008803 | -0.836235123 | 0.000309201 | 0.000800353 |
| ENSGALG00000009006 | -0.381004012 | 0.000565221 | 0.001402938 |
| ENSGALG00000009128 | -1.496282226 | 4.96E-13 | 3.34E-12 |
| ENSGALG00000009149 | 0.480936229 | 7.61E-05 | 0.000215584 |
| ENSGALG00000009272 | -0.475046066 | 0.004081758 | 0.008724524 |
| ENSGALG00000009292 | -1.282728705 | 8.69E-06 | 2.80E-05 |
| ENSGALG00000009298 | 0.675442976 | 3.23E-09 | 1.52E-08 |
| ENSGALG00000009332 | -0.425301068 | 0.008187425 | 0.016345987 |
| ENSGALG00000009387 | 1.674898522 | 5.94E-19 | 6.27E-18 |
| ENSGALG00000009409 | -0.631848959 | 0.003147644 | 0.006889336 |
| ENSGALG00000009479 | 6.633651788 | 6.22E-07 | 2.32E-06 |
| ENSGALG00000009504 | -1.018295534 | 5.36E-14 | 3.90E-13 |
| ENSGALG00000009538 | -0.483070011 | 2.92E-05 | 8.76E-05 |
| ENSGALG00000009608 | -0.955865416 | 0.000169473 | 0.000455322 |
| ENSGALG00000009628 | -1.552883323 | 1.04E-07 | 4.23E-07 |
| ENSGALG00000009650 | 0.344344363 | 0.000266636 | 0.000696544 |
| ENSGALG00000009792 | -0.377605498 | 0.008213087 | 0.016392405 |
| ENSGALG00000009879 | 0.717598413 | 2.75E-17 | 2.59E-16 |
| ENSGALG00000010030 | -0.886055284 | 2.39E-07 | 9.33E-07 |
| ENSGALG00000010032 | 2.497963803 | 1.34E-23 | 1.90E-22 |
| ENSGALG00000010074 | -1.158018398 | 1.08E-18 | 1.12E-17 |
| ENSGALG00000010108 | 0.827302229 | 4.76E-16 | 4.05E-15 |
| ENSGALG00000010189 | -0.820332955 | 1.38E-11 | 8.12E-11 |
| ENSGALG00000010203 | -0.991230538 | 2.71E-06 | 9.39E-06 |

| | | | |
|---|---|---|---|
| ENSGALG00000010217 | -1.409506912 | 2.69E-11 | 1.54E-10 |
| ENSGALG00000010226 | -0.472861327 | 0.009963068 | 0.019523937 |
| ENSGALG00000010268 | 0.759649368 | 1.14E-12 | 7.38E-12 |
| ENSGALG00000010302 | -0.85249522 | 2.78E-16 | 2.40E-15 |
| ENSGALG00000010303 | 1.536318448 | 0.000203771 | 0.000541053 |
| ENSGALG00000010313 | 0.57732401 | 6.19E-05 | 0.000177891 |
| ENSGALG00000010335 | 1.005767233 | 1.22E-29 | 2.50E-28 |
| ENSGALG00000010433 | -0.718706936 | 0.001380029 | 0.003220146 |
| ENSGALG00000010457 | 1.557238319 | 0.003175999 | 0.006946923 |
| ENSGALG00000010635 | 3.886406363 | 0.004086737 | 0.008729677 |
| ENSGALG00000010758 | 0.819600443 | 3.36E-17 | 3.15E-16 |
| ENSGALG00000010854 | -0.413932861 | 0.008269523 | 0.016500199 |
| ENSGALG00000010901 | 0.945641447 | 0.024150811 | 0.043477858 |
| ENSGALG00000010963 | -0.920995109 | 0.007306643 | 0.014739076 |
| ENSGALG00000010979 | -0.503801457 | 3.07E-05 | 9.19E-05 |
| ENSGALG00000011164 | 1.011263281 | 5.53E-35 | 1.54E-33 |
| ENSGALG00000011166 | -0.883217342 | 0.01496156 | 0.028228419 |
| ENSGALG00000011285 | -0.473341405 | 1.64E-05 | 5.07E-05 |
| ENSGALG00000011324 | -0.48424708 | 8.19E-08 | 3.37E-07 |
| ENSGALG00000011459 | -0.848818583 | 0.006307019 | 0.012902619 |
| ENSGALG00000011528 | -0.313894576 | 0.015879891 | 0.029762753 |
| ENSGALG00000011608 | -0.788622285 | 0.00149483 | 0.003467781 |
| ENSGALG00000011687 | -0.731873918 | 1.10E-13 | 7.77E-13 |
| ENSGALG00000011796 | -2.168249692 | 0.001784379 | 0.004080983 |
| ENSGALG00000011799 | 0.769859731 | 0.026080281 | 0.046483501 |
| ENSGALG00000011805 | 0.430738906 | 7.48E-05 | 0.00021232 |
| ENSGALG00000011812 | 0.297908432 | 0.00428281 | 0.009089352 |
| ENSGALG00000011889 | -1.055309554 | 1.66E-11 | 9.69E-11 |
| ENSGALG00000012021 | 1.192037658 | 2.94E-11 | 1.67E-10 |
| ENSGALG00000012055 | -0.971554692 | 1.50E-07 | 5.99E-07 |
| ENSGALG00000012085 | -0.608638737 | 0.000294794 | 0.000765103 |
| ENSGALG00000012103 | 0.60908788 | 0.000145839 | 0.000395421 |
| ENSGALG00000012121 | 0.66726842 | 4.46E-15 | 3.51E-14 |
| ENSGALG00000012187 | 0.981195135 | 4.96E-09 | 2.29E-08 |
| ENSGALG00000012207 | -1.187972988 | 4.31E-14 | 3.16E-13 |
| ENSGALG00000012259 | -0.647251043 | 4.50E-10 | 2.28E-09 |
| ENSGALG00000012269 | -0.715308998 | 7.90E-06 | 2.56E-05 |
| ENSGALG00000012394 | -1.892139175 | 1.51E-53 | 1.03E-51 |
| ENSGALG00000012591 | 0.517064338 | 1.72E-09 | 8.31E-09 |
| ENSGALG00000012621 | -1.003092694 | 1.26E-08 | 5.64E-08 |
| ENSGALG00000012766 | 0.881227038 | 0.010693848 | 0.020823894 |
| ENSGALG00000013057 | 0.271262518 | 0.013560246 | 0.025839178 |
| ENSGALG00000013155 | 0.879433352 | 0.005994239 | 0.012323828 |
| ENSGALG00000013207 | -0.532186571 | 0.008715865 | 0.017266585 |
| ENSGALG00000013239 | 0.499044877 | 0.000829499 | 0.002003956 |
| ENSGALG00000013247 | 1.606953021 | 8.00E-06 | 2.59E-05 |
| ENSGALG00000013430 | 0.547955361 | 0.000105803 | 0.000293185 |
| ENSGALG00000013565 | -1.140567462 | 7.62E-08 | 3.14E-07 |

| | | | |
|---|---|---|---|
| ENSGALG00000013624 | -1.929232787 | 0.023326081 | 0.042082306 |
| ENSGALG00000013765 | -0.449005617 | 1.75E-06 | 6.22E-06 |
| ENSGALG00000013777 | -0.347031794 | 0.000373636 | 0.000955317 |
| ENSGALG00000013804 | 0.805182019 | 6.83E-19 | 7.19E-18 |
| ENSGALG00000013912 | 3.795825366 | 0.013148518 | 0.025156906 |
| ENSGALG00000013943 | 0.399207804 | 3.28E-06 | 1.12E-05 |
| ENSGALG00000013950 | -0.556473637 | 1.07E-05 | 3.40E-05 |
| ENSGALG00000014066 | -0.519757386 | 8.03E-06 | 2.60E-05 |
| ENSGALG00000014126 | 1.850344844 | 0.009704273 | 0.019049751 |
| ENSGALG00000014262 | -0.554287211 | 0.015935237 | 0.029854133 |
| ENSGALG00000014347 | 0.493891965 | 0.00428939 | 0.009098213 |
| ENSGALG00000014551 | -0.569861336 | 0.008205323 | 0.016379313 |
| ENSGALG00000014589 | 1.820496118 | 2.46E-44 | 1.12E-42 |
| ENSGALG00000014680 | -1.438967505 | 1.22E-13 | 8.61E-13 |
| ENSGALG00000014719 | 0.864039587 | 8.32E-07 | 3.06E-06 |
| ENSGALG00000014721 | -0.517751134 | 0.002526119 | 0.005631459 |
| ENSGALG00000014750 | 1.340777036 | 0.000649257 | 0.001595497 |
| ENSGALG00000014857 | 1.030807635 | 3.77E-24 | 5.48E-23 |
| ENSGALG00000014902 | -0.645353338 | 5.25E-09 | 2.42E-08 |
| ENSGALG00000014960 | -1.136521313 | 1.42E-06 | 5.10E-06 |
| ENSGALG00000015059 | -1.735814994 | 2.50E-17 | 2.36E-16 |
| ENSGALG00000015140 | -1.20604303 | 3.91E-14 | 2.88E-13 |
| ENSGALG00000015142 | -0.526573962 | 0.00070733 | 0.001728209 |
| ENSGALG00000015157 | -1.01697089 | 4.02E-16 | 3.44E-15 |
| ENSGALG00000015183 | 0.716954976 | 0.007350879 | 0.014817314 |
| ENSGALG00000015234 | 0.70306444 | 0.000739608 | 0.001801569 |
| ENSGALG00000015240 | -1.21068247 | 3.73E-07 | 1.43E-06 |
| ENSGALG00000015249 | -1.282926203 | 8.62E-09 | 3.89E-08 |
| ENSGALG00000015297 | -1.473714963 | 2.32E-27 | 4.22E-26 |
| ENSGALG00000015324 | -1.478711015 | 0.000846876 | 0.00204381 |
| ENSGALG00000015342 | -1.980982934 | 4.48E-25 | 7.01E-24 |
| ENSGALG00000015349 | -1.044172532 | 5.58E-09 | 2.56E-08 |
| ENSGALG00000015352 | -1.261202159 | 8.34E-07 | 3.07E-06 |
| ENSGALG00000015391 | -0.755310157 | 1.64E-12 | 1.06E-11 |
| ENSGALG00000015428 | 1.548711251 | 0.001269461 | 0.002978327 |
| ENSGALG00000015576 | -0.730503041 | 0.000377002 | 0.000963016 |
| ENSGALG00000015652 | -0.355908454 | 0.002661111 | 0.005907206 |
| ENSGALG00000015667 | -1.263873449 | 5.49E-08 | 2.29E-07 |
| ENSGALG00000015684 | -0.728099636 | 2.59E-10 | 1.35E-09 |
| ENSGALG00000015687 | -1.731804059 | 9.53E-11 | 5.16E-10 |
| ENSGALG00000015705 | 0.478659996 | 0.015490765 | 0.02912581 |
| ENSGALG00000015822 | 0.699546 | 1.58E-07 | 6.31E-07 |
| ENSGALG00000016011 | 6.247117428 | 7.94E-05 | 0.000224457 |
| ENSGALG00000016196 | -2.027612911 | 1.73E-12 | 1.11E-11 |
| ENSGALG00000016217 | -0.374705822 | 0.013178043 | 0.025202751 |
| ENSGALG00000016236 | -0.359701174 | 0.000612912 | 0.001512197 |
| ENSGALG00000016285 | -0.434007346 | 0.00034828 | 0.000894356 |
| ENSGALG00000016292 | 0.740396253 | 0.022375466 | 0.04057195 |

| | | | |
|---|---|---|---|
| ENSGALG00000016327 | 1.334443782 | 1.08E-41 | 4.25E-40 |
| ENSGALG00000016499 | 4.707734463 | 0.004694573 | 0.009880556 |
| ENSGALG00000016508 | -1.395154386 | 2.89E-10 | 1.50E-09 |
| ENSGALG00000016514 | 0.723360431 | 1.66E-13 | 1.16E-12 |
| ENSGALG00000016524 | -0.677285081 | 4.48E-06 | 1.50E-05 |
| ENSGALG00000016556 | -1.016895116 | 0.024383371 | 0.043850064 |
| ENSGALG00000016651 | -0.854210193 | 0.007329084 | 0.014779957 |
| ENSGALG00000016702 | -1.626517608 | 3.36E-10 | 1.73E-09 |
| ENSGALG00000016722 | 3.213340504 | 0.009656862 | 0.018973124 |
| ENSGALG00000016760 | 0.429413291 | 0.00294668 | 0.006484986 |
| ENSGALG00000016808 | -0.959151357 | 3.20E-06 | 1.09E-05 |
| ENSGALG00000016815 | 3.427860626 | 1.10E-31 | 2.54E-30 |
| ENSGALG00000016934 | -0.592876084 | 0.002053652 | 0.004644466 |
| ENSGALG00000016964 | 1.427884975 | 0.006473542 | 0.013215437 |
| ENSGALG00000016971 | -1.34270338 | 2.57E-28 | 4.91E-27 |
| ENSGALG00000017031 | -1.202412747 | 0.01823563 | 0.033745225 |
| ENSGALG00000017072 | 1.339462435 | 1.45E-47 | 7.90E-46 |
| ENSGALG00000017220 | -0.448454658 | 0.000334076 | 0.000860807 |
| ENSGALG00000017250 | 1.942447743 | 0.005052187 | 0.010556392 |
| ENSGALG00000017276 | 2.548744315 | 0.0061075 | 0.012530286 |
| ENSGALG00000017290 | 0.694453629 | 8.61E-07 | 3.17E-06 |
| ENSGALG00000017334 | -2.257471195 | 9.11E-11 | 4.95E-10 |
| ENSGALG00000017336 | -1.205539381 | 2.45E-18 | 2.48E-17 |
| ENSGALG00000017376 | -0.976156912 | 0.000183186 | 0.000489841 |
| ENSGALG00000017398 | -0.304753529 | 0.000569836 | 0.001413359 |
| ENSGALG00000019077 | -1.636362221 | 0.00138372 | 0.003227097 |
| ENSGALG00000019147 | 4.790164849 | 2.69E-07 | 1.05E-06 |
| ENSGALG00000019262 | 2.84026221 | 0.000148663 | 0.000402515 |
| ENSGALG00000019286 | -0.292291631 | 0.01720545 | 0.032008826 |
| ENSGALG00000019554 | 2.72696551 | 8.10E-12 | 4.88E-11 |
| ENSGALG00000020342 | -0.657335121 | 2.80E-08 | 1.20E-07 |
| ENSGALG00000020458 | 1.118666533 | 2.51E-05 | 7.60E-05 |
| ENSGALG00000020488 | 0.519071628 | 1.90E-08 | 8.29E-08 |
| ENSGALG00000020561 | -1.794958747 | 0.002134818 | 0.004814409 |
| ENSGALG00000020626 | 1.050116395 | 1.86E-28 | 3.58E-27 |
| ENSGALG00000020775 | 0.789408665 | 6.72E-05 | 0.000192101 |
| ENSGALG00000020921 | 0.752864543 | 0.002544719 | 0.005667347 |
| ENSGALG00000021171 | 1.512333683 | 5.48E-08 | 2.29E-07 |
| ENSGALG00000021174 | 0.498703446 | 0.001516708 | 0.003514936 |
| ENSGALG00000021286 | 2.7058896 | 0.00812466 | 0.016234987 |
| ENSGALG00000021301 | -1.380719713 | 9.85E-06 | 3.15E-05 |
| ENSGALG00000021589 | -2.773152871 | 0.000554081 | 0.001376835 |
| ENSGALG00000021611 | -0.547422339 | 8.62E-08 | 3.53E-07 |
| ENSGALG00000021665 | -0.371712681 | 0.020171019 | 0.03693446 |
| ENSGALG00000021676 | 1.981928139 | 2.21E-94 | 7.14E-92 |
| ENSGALG00000021692 | 0.568207687 | 2.02E-08 | 8.84E-08 |
| ENSGALG00000022804 | 0.438970632 | 0.000710868 | 0.00173623 |
| ENSGALG00000022891 | -0.718488331 | 4.48E-08 | 1.89E-07 |

| | | | |
|---|---|---|---|
| ENSGALG00000023632 | -0.379083574 | 0.011667408 | 0.022542063 |
| ENSGALG00000023661 | -0.8539316 | 7.85E-13 | 5.16E-12 |
| ENSGALG00000023709 | 1.651955497 | 0.000829236 | 0.002003729 |
| ENSGALG00000023763 | 1.459808524 | 0.011309116 | 0.021899631 |
| ENSGALG00000023819 | -0.556806419 | 1.50E-11 | 8.82E-11 |
| ENSGALG00000023909 | 2.00499835 | 5.48E-42 | 2.22E-40 |
| ENSGALG00000024077 | -0.613941745 | 0.024871036 | 0.044598334 |
| ENSGALG00000024295 | 0.317443309 | 0.021382419 | 0.03893233 |
| ENSGALG00000024335 | 0.742076673 | 2.30E-08 | 9.98E-08 |
| ENSGALG00000024379 | 4.753413385 | 0.003883047 | 0.008322041 |
| ENSGALG00000025800 | 1.724932745 | 6.38E-10 | 3.20E-09 |
| ENSGALG00000025948 | 1.529773978 | 0.000306176 | 0.000792825 |
| ENSGALG00000026152 | -0.999906806 | 0.000111333 | 0.00030738 |
| ENSGALG00000026154 | 0.550384026 | 0.000232867 | 0.000613873 |
| ENSGALG00000026161 | 1.23729576 | 0.004580888 | 0.009656238 |
| ENSGALG00000026278 | 0.328513089 | 0.022355821 | 0.040542153 |
| ENSGALG00000026301 | -0.957018445 | 0.013750339 | 0.026164721 |
| ENSGALG00000026422 | 2.293177464 | 6.51E-08 | 2.70E-07 |
| ENSGALG00000026426 | -1.283376168 | 2.47E-35 | 7.01E-34 |
| ENSGALG00000026449 | 0.354616374 | 0.010952972 | 0.021282745 |
| ENSGALG00000026553 | 5.014751225 | 1.10E-19 | 1.22E-18 |
| ENSGALG00000026607 | 0.854680807 | 1.73E-13 | 1.21E-12 |
| ENSGALG00000026816 | 1.650863838 | 9.69E-33 | 2.34E-31 |
| ENSGALG00000026875 | 3.533755289 | 0.012442713 | 0.023944691 |
| ENSGALG00000026896 | -1.844710425 | 2.81E-36 | 8.49E-35 |
| ENSGALG00000026928 | -1.880870988 | 5.39E-41 | 2.05E-39 |
| ENSGALG00000026970 | 1.814317699 | 1.89E-104 | 7.57E-102 |
| ENSGALG00000027132 | -1.164208218 | 3.14E-08 | 1.34E-07 |
| ENSGALG00000027147 | 3.221880928 | 0.00021253 | 0.000563102 |
| ENSGALG00000027159 | -0.910735438 | 2.83E-19 | 3.04E-18 |
| ENSGALG00000027170 | -0.698344016 | 0.015490579 | 0.02912581 |
| ENSGALG00000027173 | 0.795885418 | 1.22E-13 | 8.62E-13 |
| ENSGALG00000027397 | 1.162018439 | 0.000302818 | 0.000784877 |
| ENSGALG00000027418 | 1.211292432 | 9.51E-05 | 0.000265607 |
| ENSGALG00000027520 | 1.638015409 | 0.005715913 | 0.011799922 |
| ENSGALG00000027540 | 1.018933226 | 4.09E-24 | 5.91E-23 |
| ENSGALG00000027620 | 0.455394688 | 0.013348897 | 0.025493636 |
| ENSGALG00000027645 | 0.949435475 | 0.000722828 | 0.001763857 |
| ENSGALG00000027696 | -1.388627914 | 3.95E-15 | 3.12E-14 |
| ENSGALG00000027742 | 7.296264602 | 2.03E-16 | 1.78E-15 |
| ENSGALG00000027765 | 4.848931969 | 0.00025496 | 0.000668353 |
| ENSGALG00000027887 | 1.013807045 | 1.08E-05 | 3.44E-05 |
| ENSGALG00000027899 | -0.831009532 | 0.000991713 | 0.00237271 |
| ENSGALG00000027983 | -0.313480558 | 0.000214186 | 0.000567379 |
| ENSGALG00000028009 | 0.524017628 | 4.45E-09 | 2.06E-08 |
| ENSGALG00000028016 | 1.909409155 | 0.004085761 | 0.008728964 |
| ENSGALG00000028056 | 0.333852088 | 0.000768517 | 0.001867302 |
| ENSGALG00000028135 | 0.913419554 | 1.66E-11 | 9.68E-11 |

| | | | |
|---|---|---|---|
| ENSGALG00000028174 | 0.652168684 | 5.24E-13 | 3.52E-12 |
| ENSGALG00000028222 | 0.649556892 | 0.016954637 | 0.031576792 |
| ENSGALG00000028267 | 0.323862075 | 0.007933464 | 0.015892651 |
| ENSGALG00000028301 | -0.816280491 | 4.79E-07 | 1.81E-06 |
| ENSGALG00000028327 | 0.640119711 | 0.022496255 | 0.040736623 |
| ENSGALG00000028340 | 0.999324751 | 4.58E-16 | 3.90E-15 |
| ENSGALG00000028341 | 0.951529958 | 0.010231124 | 0.019985834 |
| ENSGALG00000028357 | 3.025984422 | 0.017371081 | 0.0322751 |
| ENSGALG00000028466 | 1.953928206 | 1.81E-74 | 3.04E-72 |
| ENSGALG00000028496 | 3.620732643 | 0.000391445 | 0.000997472 |
| ENSGALG00000028530 | 0.629688827 | 2.48E-09 | 1.18E-08 |
| ENSGALG00000028551 | 0.572461979 | 3.38E-06 | 1.15E-05 |
| ENSGALG00000028573 | 0.312699851 | 0.013953472 | 0.026503018 |
| ENSGALG00000028612 | 0.790862281 | 0.026539753 | 0.047246669 |
| ENSGALG00000028898 | -0.346291997 | 0.010575171 | 0.020619383 |
| ENSGALG00000028985 | -0.841734158 | 9.13E-06 | 2.93E-05 |
| ENSGALG00000029012 | 1.189998356 | 2.63E-05 | 7.95E-05 |
| ENSGALG00000029066 | -0.423569382 | 1.79E-05 | 5.50E-05 |
| ENSGALG00000029168 | 2.307239548 | 0.004230003 | 0.008994712 |
| ENSGALG00000029316 | 0.509845789 | 0.024918492 | 0.044661217 |
| ENSGALG00000029324 | -0.399701832 | 0.011388268 | 0.022040344 |
| ENSGALG00000029330 | 0.758570226 | 8.02E-10 | 3.98E-09 |
| ENSGALG00000029346 | -0.958706682 | 5.42E-06 | 1.79E-05 |
| ENSGALG00000029360 | -1.190890062 | 2.36E-22 | 3.09E-21 |
| ENSGALG00000029381 | -1.117893341 | 6.48E-05 | 0.000185671 |
| ENSGALG00000029569 | -2.560423771 | 0.008582875 | 0.017044083 |
| ENSGALG00000029674 | -0.808865809 | 7.80E-11 | 4.27E-10 |
| ENSGALG00000029691 | 0.583363377 | 1.06E-10 | 5.71E-10 |
| ENSGALG00000029862 | 2.651479796 | 8.22E-26 | 1.35E-24 |
| ENSGALG00000029918 | -0.876538992 | 0.000154778 | 0.000418156 |
| ENSGALG00000030007 | 0.736377979 | 0.00700677 | 0.014203731 |
| ENSGALG00000030064 | 0.622995525 | 2.28E-13 | 1.58E-12 |
| ENSGALG00000030086 | 0.722497247 | 5.55E-17 | 5.07E-16 |
| ENSGALG00000030099 | -1.113572877 | 0.006742014 | 0.013716128 |
| ENSGALG00000030139 | 1.044607995 | 1.36E-26 | 2.34E-25 |
| ENSGALG00000030153 | 0.924318447 | 4.78E-06 | 1.60E-05 |
| ENSGALG00000030164 | -0.676555394 | 1.45E-05 | 4.51E-05 |
| ENSGALG00000030245 | -1.05268902 | 0.00705344 | 0.014281298 |
| ENSGALG00000030350 | -1.610961769 | 2.07E-07 | 8.15E-07 |
| ENSGALG00000030384 | -0.272310833 | 0.00172547 | 0.003954905 |
| ENSGALG00000030466 | -1.319310891 | 1.52E-26 | 2.60E-25 |
| ENSGALG00000030487 | 0.573034163 | 2.06E-05 | 6.30E-05 |
| ENSGALG00000030543 | 1.513968259 | 1.76E-19 | 1.92E-18 |
| ENSGALG00000030552 | 0.338452313 | 0.004175163 | 0.008890618 |
| ENSGALG00000030587 | 0.529395455 | 1.66E-07 | 6.63E-07 |
| ENSGALG00000030707 | -0.831381 | 0.000140396 | 0.000381194 |
| ENSGALG00000030710 | 1.745514984 | 0.000345835 | 0.000889083 |
| ENSGALG00000030718 | -0.612576784 | 6.24E-07 | 2.33E-06 |

| | | | |
|---|---|---|---|
| ENSGALG00000030724 | 2.628247231 | 0.012631702 | 0.024260293 |
| ENSGALG00000030808 | 0.622699807 | 1.73E-08 | 7.60E-08 |
| ENSGALG00000030810 | 0.297039343 | 0.000201958 | 0.000536661 |
| ENSGALG00000031002 | -0.901944677 | 5.73E-07 | 2.15E-06 |
| ENSGALG00000031149 | -0.38302075 | 0.010845717 | 0.021092428 |
| ENSGALG00000031167 | -0.727961514 | 0.000779465 | 0.001891535 |
| ENSGALG00000031210 | -1.101837221 | 5.44E-05 | 0.000157942 |
| ENSGALG00000031262 | 1.646059457 | 0.000242527 | 0.0006381 |
| ENSGALG00000031296 | 0.396225073 | 7.12E-07 | 2.65E-06 |
| ENSGALG00000031313 | 1.283732298 | 5.32E-09 | 2.45E-08 |
| ENSGALG00000031355 | 1.172919179 | 0.001311007 | 0.003065933 |
| ENSGALG00000031430 | 2.519302042 | 1.34E-14 | 1.01E-13 |
| ENSGALG00000031518 | -2.495580325 | 2.84E-20 | 3.25E-19 |
| ENSGALG00000031524 | -0.915439788 | 0.003584066 | 0.007744773 |
| ENSGALG00000031594 | -1.715549984 | 0.0279269 | 0.049424795 |
| ENSGALG00000031604 | 1.132679888 | 1.37E-10 | 7.29E-10 |
| ENSGALG00000031737 | 2.40032941 | 6.36E-11 | 3.50E-10 |
| ENSGALG00000031860 | 0.733287029 | 0.000114726 | 0.000315722 |
| ENSGALG00000031929 | 0.886210391 | 1.48E-16 | 1.31E-15 |
| ENSGALG00000032066 | 1.138273624 | 7.88E-05 | 0.000222777 |
| ENSGALG00000032206 | 0.74586581 | 0.000533482 | 0.001329012 |
| ENSGALG00000032250 | 1.286769038 | 2.22E-06 | 7.76E-06 |
| ENSGALG00000032428 | 0.469566259 | 0.00563077 | 0.011645379 |
| ENSGALG00000032534 | 1.608991141 | 0.001797817 | 0.004109641 |
| ENSGALG00000032799 | -0.789218257 | 0.010027683 | 0.01962792 |
| ENSGALG00000032803 | 2.73939339 | 6.38E-33 | 1.55E-31 |
| ENSGALG00000032808 | -1.160440096 | 6.08E-11 | 3.36E-10 |
| ENSGALG00000032819 | 1.123031266 | 3.98E-19 | 4.23E-18 |
| ENSGALG00000032903 | 1.965155475 | 4.53E-61 | 4.16E-59 |
| ENSGALG00000032915 | -2.061204532 | 5.12E-71 | 6.88E-69 |
| ENSGALG00000032916 | -0.675171909 | 0.004132668 | 0.008816703 |
| ENSGALG00000033014 | -1.097560893 | 1.24E-14 | 9.46E-14 |
| ENSGALG00000033143 | -0.588731411 | 4.58E-06 | 1.53E-05 |
| ENSGALG00000033303 | 1.100341712 | 3.95E-05 | 0.000116767 |
| ENSGALG00000033320 | -0.455142374 | 4.83E-06 | 1.61E-05 |
| ENSGALG00000033381 | -2.66865216 | 5.25E-08 | 2.20E-07 |
| ENSGALG00000033498 | -1.117752261 | 2.03E-21 | 2.48E-20 |
| ENSGALG00000033522 | -1.87990235 | 7.14E-86 | 1.80E-83 |
| ENSGALG00000033569 | 1.325592822 | 6.29E-43 | 2.66E-41 |
| ENSGALG00000033570 | -0.667341825 | 0.00264141 | 0.005868265 |
| ENSGALG00000033597 | -1.693952864 | 0.006042856 | 0.012410774 |
| ENSGALG00000033647 | -0.764896838 | 5.14E-06 | 1.71E-05 |
| ENSGALG00000033705 | -0.718007901 | 4.40E-16 | 3.76E-15 |
| ENSGALG00000033780 | 0.818198803 | 0.000369667 | 0.000946058 |
| ENSGALG00000033919 | 1.045192867 | 1.65E-05 | 5.09E-05 |
| ENSGALG00000034013 | 0.714239001 | 1.08E-05 | 3.43E-05 |
| ENSGALG00000034050 | -0.265405297 | 0.007050953 | 0.014280517 |
| ENSGALG00000034054 | 4.883141807 | 2.09E-16 | 1.83E-15 |

| | | | |
|---|---|---|---|
| ENSGALG00000034204 | 5.586553632 | 6.92E-23 | 9.32E-22 |
| ENSGALG00000034218 | 0.215565106 | 0.024715611 | 0.044377091 |
| ENSGALG00000034232 | -1.253105908 | 0.011319795 | 0.021914066 |
| ENSGALG00000034264 | -0.516518429 | 0.000345704 | 0.000888917 |
| ENSGALG00000034282 | 2.075521429 | 4.52E-15 | 3.55E-14 |
| ENSGALG00000034345 | 0.327234545 | 0.00841067 | 0.016749864 |
| ENSGALG00000034349 | 6.059084704 | 1.34E-05 | 4.19E-05 |
| ENSGALG00000034354 | 0.509843748 | 6.52E-08 | 2.70E-07 |
| ENSGALG00000034358 | -1.812750156 | 3.41E-43 | 1.46E-41 |
| ENSGALG00000034397 | 1.934035255 | 7.09E-23 | 9.55E-22 |
| ENSGALG00000034511 | -1.242280296 | 3.24E-06 | 1.11E-05 |
| ENSGALG00000034512 | -1.606079925 | 0.001597849 | 0.003684758 |
| ENSGALG00000034527 | 0.542433766 | 8.31E-07 | 3.06E-06 |
| ENSGALG00000034572 | -1.656791527 | 3.75E-13 | 2.56E-12 |
| ENSGALG00000034584 | 0.846459876 | 1.41E-12 | 9.10E-12 |
| ENSGALG00000034591 | 0.731468675 | 3.38E-12 | 2.11E-11 |
| ENSGALG00000034735 | -1.400742875 | 9.39E-44 | 4.20E-42 |
| ENSGALG00000034761 | 7.769658642 | 7.14E-09 | 3.25E-08 |
| ENSGALG00000034772 | -1.025695529 | 1.97E-05 | 6.05E-05 |
| ENSGALG00000034806 | 1.222514252 | 0.00428711 | 0.009094803 |
| ENSGALG00000034903 | -0.813462384 | 9.21E-05 | 0.00025789 |
| ENSGALG00000034939 | -0.695427007 | 0.000464668 | 0.001168919 |
| ENSGALG00000035029 | -0.549716601 | 0.001432181 | 0.003332684 |
| ENSGALG00000035047 | 0.883691334 | 2.98E-08 | 1.28E-07 |
| ENSGALG00000035049 | -1.213900271 | 9.46E-05 | 0.000264211 |
| ENSGALG00000035185 | 2.100671213 | 0.021118163 | 0.03850276 |
| ENSGALG00000035206 | 0.40278329 | 3.05E-06 | 1.05E-05 |
| ENSGALG00000035237 | 0.220236745 | 0.010247188 | 0.020014338 |
| ENSGALG00000035244 | 0.419248428 | 0.000289251 | 0.000751576 |
| ENSGALG00000035246 | 2.001575822 | 0.024740124 | 0.04440707 |
| ENSGALG00000035447 | -1.33656538 | 7.65E-24 | 1.10E-22 |
| ENSGALG00000035453 | 1.076207399 | 2.99E-12 | 1.88E-11 |
| ENSGALG00000035675 | 0.541800763 | 0.001578929 | 0.003645974 |
| ENSGALG00000035749 | -1.16295481 | 0.002042947 | 0.004623333 |
| ENSGALG00000035769 | -0.875652507 | 4.03E-09 | 1.87E-08 |
| ENSGALG00000035780 | -1.021994134 | 1.17E-15 | 9.62E-15 |
| ENSGALG00000035818 | 1.617451593 | 0.016496938 | 0.030808792 |
| ENSGALG00000035825 | -2.871673041 | 0.008396342 | 0.016723781 |
| ENSGALG00000035879 | -1.43132528 | 0.005934756 | 0.012210901 |
| ENSGALG00000035912 | 0.991767472 | 1.26E-07 | 5.07E-07 |
| ENSGALG00000035960 | 1.702019376 | 0.002283654 | 0.005121173 |
| ENSGALG00000036171 | -1.19163142 | 0.008318787 | 0.016588756 |
| ENSGALG00000036263 | 0.707182612 | 0.013788863 | 0.026223343 |
| ENSGALG00000036286 | 0.753361699 | 4.19E-09 | 1.94E-08 |
| ENSGALG00000036310 | 1.428586171 | 1.92E-65 | 2.17E-63 |
| ENSGALG00000036466 | -0.351249187 | 0.003573739 | 0.007724564 |
| ENSGALG00000036550 | 0.751149442 | 2.27E-11 | 1.31E-10 |
| ENSGALG00000036763 | 1.243240238 | 0.015768005 | 0.029581605 |

| | | | |
|---|---|---|---|
| ENSGALG00000036846 | 3.209364546 | 0.000463548 | 0.001166769 |
| ENSGALG00000036908 | 1.105757584 | 0.001913961 | 0.004348806 |
| ENSGALG00000036935 | 0.25444394 | 0.018853815 | 0.034780273 |
| ENSGALG00000037051 | -0.660659376 | 7.47E-08 | 3.08E-07 |
| ENSGALG00000037112 | 1.375769509 | 1.18E-05 | 3.73E-05 |
| ENSGALG00000037160 | 1.105940428 | 0.000208664 | 0.000553377 |
| ENSGALG00000037166 | 0.919211594 | 5.08E-26 | 8.48E-25 |
| ENSGALG00000037186 | 1.062554984 | 3.58E-11 | 2.02E-10 |
| ENSGALG00000037200 | 0.780075425 | 0.025339664 | 0.045329917 |
| ENSGALG00000037214 | -1.935790501 | 1.67E-12 | 1.07E-11 |
| ENSGALG00000037273 | 1.397171611 | 1.07E-08 | 4.79E-08 |
| ENSGALG00000037314 | 0.948561042 | 1.31E-08 | 5.82E-08 |
| ENSGALG00000037441 | 1.10758375 | 5.69E-45 | 2.67E-43 |
| ENSGALG00000037678 | 2.457178591 | 0.000127413 | 0.000348513 |
| ENSGALG00000037699 | -0.924091705 | 2.32E-13 | 1.61E-12 |
| ENSGALG00000037723 | -0.555727894 | 1.32E-06 | 4.73E-06 |
| ENSGALG00000037737 | 0.39405131 | 0.003198588 | 0.006987336 |
| ENSGALG00000037816 | -1.823345555 | 7.59E-15 | 5.87E-14 |
| ENSGALG00000037860 | 1.288759365 | 0.007790866 | 0.015648508 |
| ENSGALG00000037941 | -0.683929733 | 1.06E-05 | 3.37E-05 |
| ENSGALG00000037957 | 1.416578303 | 1.80E-05 | 5.53E-05 |
| ENSGALG00000037997 | 3.21256347 | 0.001994284 | 0.004517718 |
| ENSGALG00000038143 | 0.611304059 | 2.54E-10 | 1.33E-09 |
| ENSGALG00000038160 | -0.95671348 | 4.00E-12 | 2.48E-11 |
| ENSGALG00000038254 | 2.303358903 | 0.000701581 | 0.001715397 |
| ENSGALG00000038432 | 3.660081588 | 0.020786836 | 0.037934301 |
| ENSGALG00000038528 | 0.435503059 | 1.03E-05 | 3.28E-05 |
| ENSGALG00000038550 | 0.645425744 | 0.000773769 | 0.001879056 |
| ENSGALG00000038574 | -2.933949056 | 0.025068068 | 0.044891329 |
| ENSGALG00000038579 | -2.465279118 | 0.00048059 | 0.001204088 |
| ENSGALG00000038584 | 2.692674975 | 0.004371856 | 0.009253652 |
| ENSGALG00000038652 | 0.449991014 | 0.000332011 | 0.00085576 |
| ENSGALG00000038671 | 1.536360113 | 5.49E-18 | 5.41E-17 |
| ENSGALG00000038672 | 0.634848757 | 2.01E-11 | 1.16E-10 |
| ENSGALG00000038728 | 1.168155029 | 0.002227238 | 0.005005393 |
| ENSGALG00000038783 | 0.313895784 | 0.000219174 | 0.00057935 |
| ENSGALG00000038810 | -1.416927692 | 1.85E-15 | 1.49E-14 |
| ENSGALG00000038820 | 0.612661648 | 7.60E-05 | 0.000215171 |
| ENSGALG00000038853 | -1.587069417 | 2.63E-07 | 1.02E-06 |
| ENSGALG00000038896 | -1.219505467 | 6.41E-14 | 4.63E-13 |
| ENSGALG00000038923 | 0.609425376 | 0.007745509 | 0.015564305 |
| ENSGALG00000038972 | -0.485354812 | 0.012764206 | 0.02450093 |
| ENSGALG00000038978 | 0.242770153 | 0.016807978 | 0.031338002 |
| ENSGALG00000038984 | 2.675139921 | 1.20E-09 | 5.85E-09 |
| ENSGALG00000039023 | -0.463938927 | 0.00130794 | 0.003060341 |
| ENSGALG00000039072 | -0.710353908 | 0.018254193 | 0.033774978 |
| ENSGALG00000039156 | -4.499436058 | 0.003182187 | 0.006957099 |
| ENSGALG00000039164 | 1.156875807 | 2.84E-27 | 5.12E-26 |

| | | | |
|---|---|---|---|
| ENSGALG00000039186 | -0.50789624 | 0.000466697 | 0.001172957 |
| ENSGALG00000039334 | 2.395516134 | 0.006805871 | 0.013829481 |
| ENSGALG00000039468 | 1.264160732 | 2.61E-56 | 1.96E-54 |
| ENSGALG00000039525 | 0.708136612 | 8.44E-06 | 2.72E-05 |
| ENSGALG00000039554 | 3.607726529 | 0.002722309 | 0.006025342 |
| ENSGALG00000039590 | -0.580205862 | 0.000397775 | 0.001012086 |
| ENSGALG00000039611 | -1.531622899 | 1.87E-06 | 6.63E-06 |
| ENSGALG00000039648 | 1.087755874 | 8.85E-05 | 0.000248536 |
| ENSGALG00000039708 | 1.242897863 | 0.003346997 | 0.007276453 |
| ENSGALG00000039733 | -0.652142002 | 0.000539815 | 0.001343311 |
| ENSGALG00000039755 | -0.69821532 | 2.89E-05 | 8.67E-05 |
| ENSGALG00000039964 | -0.851928055 | 6.28E-14 | 4.54E-13 |
| ENSGALG00000039995 | 0.548876038 | 6.84E-09 | 3.12E-08 |
| ENSGALG00000039999 | -0.992139065 | 7.98E-05 | 0.000225397 |
| ENSGALG00000040026 | 5.319192685 | 0 | 0 |
| ENSGALG00000040050 | 5.08523938 | 5.78E-27 | 1.01E-25 |
| ENSGALG00000040120 | 1.241861778 | 2.17E-08 | 9.44E-08 |
| ENSGALG00000040131 | 1.120394698 | 3.23E-11 | 1.83E-10 |
| ENSGALG00000040182 | 0.818456199 | 0.007920573 | 0.015873846 |
| ENSGALG00000040185 | -0.47563672 | 0.020283517 | 0.03710546 |
| ENSGALG00000040269 | 1.247635111 | 0.003410777 | 0.007401654 |
| ENSGALG00000040291 | 0.611937484 | 0.00084086 | 0.002030372 |
| ENSGALG00000040371 | 0.916351673 | 3.02E-33 | 7.55E-32 |
| ENSGALG00000040434 | 0.644710581 | 2.57E-08 | 1.11E-07 |
| ENSGALG00000040489 | -1.833895128 | 0.007296987 | 0.014722394 |
| ENSGALG00000040546 | -1.57259702 | 2.79E-11 | 1.59E-10 |
| ENSGALG00000040646 | 0.917886251 | 1.42E-15 | 1.16E-14 |
| ENSGALG00000040663 | -0.739535771 | 1.42E-07 | 5.69E-07 |
| ENSGALG00000040713 | 0.386086374 | 9.14E-06 | 2.93E-05 |
| ENSGALG00000040770 | 0.99219986 | 4.52E-07 | 1.72E-06 |
| ENSGALG00000040783 | -1.125974164 | 1.17E-20 | 1.36E-19 |
| ENSGALG00000040808 | -0.713989561 | 0.001333086 | 0.003115425 |
| ENSGALG00000040828 | 0.375503872 | 7.64E-06 | 2.48E-05 |
| ENSGALG00000040833 | -1.345112826 | 1.60E-08 | 7.04E-08 |
| ENSGALG00000040869 | 1.602303559 | 1.57E-08 | 6.92E-08 |
| ENSGALG00000040886 | -1.283515012 | 2.47E-08 | 1.07E-07 |
| ENSGALG00000040995 | 2.501327867 | 7.30E-07 | 2.71E-06 |
| ENSGALG00000041030 | 1.224650335 | 4.80E-05 | 0.000140653 |
| ENSGALG00000041052 | 0.901421625 | 0.006893438 | 0.013994736 |
| ENSGALG00000041054 | -0.801513733 | 0.00685893 | 0.013931048 |
| ENSGALG00000041081 | 1.145894899 | 4.06E-18 | 4.02E-17 |
| ENSGALG00000041181 | -0.775831561 | 2.40E-08 | 1.04E-07 |
| ENSGALG00000041205 | -0.958754347 | 0.000648966 | 0.001595072 |
| ENSGALG00000041221 | 1.04409577 | 8.65E-28 | 1.60E-26 |
| ENSGALG00000041225 | 1.052746004 | 0.004374765 | 0.009257638 |
| ENSGALG00000041338 | 1.038469361 | 2.28E-05 | 6.94E-05 |
| ENSGALG00000041372 | 2.497310119 | 0.004141319 | 0.008831001 |
| ENSGALG00000041422 | 1.020932141 | 1.23E-27 | 2.26E-26 |

| | | | |
|---|---|---|---|
| ENSGALG00000041423 | -0.397538507 | 0.000795286 | 0.001926489 |
| ENSGALG00000041435 | 0.998873152 | 2.01E-09 | 9.62E-09 |
| ENSGALG00000041442 | 2.528263448 | 1.99E-06 | 7.01E-06 |
| ENSGALG00000041463 | 0.353462149 | 5.63E-05 | 0.000163123 |
| ENSGALG00000041514 | 0.915123026 | 6.22E-09 | 2.84E-08 |
| ENSGALG00000041524 | 1.831957746 | 0.019826604 | 0.036396975 |
| ENSGALG00000041592 | 1.0450408 | 0.010333665 | 0.020171646 |
| ENSGALG00000041858 | 1.038934051 | 0.000139333 | 0.00037846 |
| ENSGALG00000041897 | 0.980158764 | 0.009027693 | 0.017837535 |
| ENSGALG00000041973 | -0.630777182 | 0.003858876 | 0.008274151 |
| ENSGALG00000042001 | 1.320453584 | 8.50E-16 | 7.06E-15 |
| ENSGALG00000042073 | 0.822186806 | 0.009027286 | 0.017837535 |
| ENSGALG00000042129 | 0.470042864 | 0.000363582 | 0.000931539 |
| ENSGALG00000042254 | -0.225764519 | 0.009997342 | 0.019579808 |
| ENSGALG00000042275 | 0.789385881 | 0.001380965 | 0.003221777 |
| ENSGALG00000042325 | 0.62235481 | 3.61E-10 | 1.85E-09 |
| ENSGALG00000042491 | -0.452716775 | 0.000107571 | 0.000297721 |
| ENSGALG00000042503 | 0.546766187 | 5.96E-09 | 2.73E-08 |
| ENSGALG00000042570 | 3.245290605 | 2.48E-16 | 2.16E-15 |
| ENSGALG00000042572 | -1.229379946 | 5.95E-16 | 5.01E-15 |
| ENSGALG00000042612 | -0.907821822 | 2.74E-09 | 1.30E-08 |
| ENSGALG00000042647 | 1.84104306 | 3.17E-08 | 1.36E-07 |
| ENSGALG00000042733 | 0.742202959 | 4.22E-07 | 1.61E-06 |
| ENSGALG00000042748 | -1.162403345 | 0.004525182 | 0.009546217 |
| ENSGALG00000042797 | 1.503030824 | 1.10E-62 | 1.08E-60 |
| ENSGALG00000042928 | -0.758298649 | 0.002650358 | 0.005885258 |
| ENSGALG00000042929 | -0.328769728 | 0.024763008 | 0.044432846 |
| ENSGALG00000042963 | 1.120546486 | 0.000444201 | 0.001122644 |
| ENSGALG00000042984 | 0.300372563 | 0.012545361 | 0.024114913 |
| ENSGALG00000043074 | -0.506373383 | 0.00015452 | 0.000417543 |
| ENSGALG00000043126 | -1.070773464 | 4.95E-22 | 6.31E-21 |
| ENSGALG00000043130 | 0.328830792 | 0.010576932 | 0.020619859 |
| ENSGALG00000043137 | 0.529567601 | 0.008126537 | 0.016235618 |
| ENSGALG00000043198 | 1.331794977 | 8.77E-07 | 3.22E-06 |
| ENSGALG00000043243 | 0.336555606 | 0.008432119 | 0.016789436 |
| ENSGALG00000043245 | 4.692159213 | 8.48E-12 | 5.10E-11 |
| ENSGALG00000043256 | 1.101914204 | 0.012539212 | 0.024110175 |
| ENSGALG00000043257 | 2.377132654 | 8.25E-16 | 6.87E-15 |
| ENSGALG00000043329 | 1.398397651 | 2.55E-34 | 6.75E-33 |
| ENSGALG00000043336 | 0.257719 | 0.027481785 | 0.048738573 |
| ENSGALG00000043381 | 1.185931213 | 1.37E-46 | 7.10E-45 |
| ENSGALG00000043421 | -0.802455674 | 1.31E-10 | 7.01E-10 |
| ENSGALG00000043593 | -0.768894986 | 2.07E-10 | 1.08E-09 |
| ENSGALG00000043654 | 1.779976477 | 9.27E-14 | 6.60E-13 |
| ENSGALG00000043660 | 1.376704712 | 0.000112468 | 0.000310201 |
| ENSGALG00000043676 | 0.468206314 | 3.50E-05 | 0.000104088 |
| ENSGALG00000043736 | 0.992453741 | 0.005006019 | 0.010469582 |
| ENSGALG00000043758 | -1.243197616 | 1.17E-40 | 4.30E-39 |

| | | | |
|---|---|---|---|
| ENSGALG00000043765 | 0.550870568 | 8.58E-06 | 2.76E-05 |
| ENSGALG00000043772 | 1.377664968 | 0.002527386 | 0.00563336 |
| ENSGALG00000043838 | -0.688069732 | 0.00804623 | 0.016099567 |
| ENSGALG00000043903 | 1.102301903 | 1.96E-07 | 7.75E-07 |
| ENSGALG00000043909 | 1.214273907 | 6.40E-26 | 1.06E-24 |
| ENSGALG00000043916 | 2.637146388 | 0.004487881 | 0.009477825 |
| ENSGALG00000043926 | 0.50016346 | 0.003494875 | 0.007566477 |
| ENSGALG00000043930 | -1.472327214 | 6.98E-22 | 8.81E-21 |
| ENSGALG00000044002 | 2.926520126 | 6.40E-05 | 0.000183746 |
| ENSGALG00000044064 | -0.575674357 | 0.000264802 | 0.00069215 |
| ENSGALG00000044069 | -0.495818341 | 0.011132674 | 0.021588051 |
| ENSGALG00000044098 | -0.930693096 | 0.000239548 | 0.000630994 |
| ENSGALG00000044224 | -1.498515628 | 0.001188751 | 0.002808047 |
| ENSGALG00000044241 | 0.994795563 | 1.25E-06 | 4.49E-06 |
| ENSGALG00000044260 | -1.464860661 | 1.36E-19 | 1.49E-18 |
| ENSGALG00000044261 | 3.942620002 | 0.009485079 | 0.018659892 |
| ENSGALG00000044278 | 0.458407288 | 0.002822613 | 0.006231112 |
| ENSGALG00000044404 | -2.158040335 | 1.40E-06 | 5.02E-06 |
| ENSGALG00000044433 | 1.383632656 | 4.66E-59 | 3.86E-57 |
| ENSGALG00000044466 | 0.782688409 | 7.97E-16 | 6.65E-15 |
| ENSGALG00000044521 | 0.289195235 | 0.00645811 | 0.013187895 |
| ENSGALG00000044569 | 1.873367043 | 2.16E-05 | 6.58E-05 |
| ENSGALG00000044628 | -0.526048143 | 2.17E-05 | 6.61E-05 |
| ENSGALG00000044632 | -1.238800179 | 0.001269602 | 0.002978327 |
| ENSGALG00000044678 | 1.097109691 | 3.03E-07 | 1.17E-06 |
| ENSGALG00000044733 | 4.0454599 | 3.25E-06 | 1.11E-05 |
| ENSGALG00000044739 | 1.152499337 | 7.25E-09 | 3.30E-08 |
| ENSGALG00000044768 | 1.081399965 | 2.74E-07 | 1.06E-06 |
| ENSGALG00000044811 | 0.705269607 | 3.70E-07 | 1.42E-06 |
| ENSGALG00000044816 | 3.380242233 | 0.019811592 | 0.036379243 |
| ENSGALG00000044914 | 2.849315023 | 2.17E-05 | 6.61E-05 |
| ENSGALG00000045021 | -1.548705823 | 1.86E-11 | 1.08E-10 |
| ENSGALG00000045030 | 1.82661167 | 0.002643299 | 0.005870543 |
| ENSGALG00000045045 | -1.11170726 | 0.008329272 | 0.016607227 |
| ENSGALG00000045093 | 1.003110199 | 3.27E-19 | 3.51E-18 |
| ENSGALG00000045132 | 0.263794274 | 0.018610987 | 0.034355634 |
| ENSGALG00000045168 | -0.991392053 | 7.35E-08 | 3.04E-07 |
| ENSGALG00000045199 | 1.515923744 | 2.77E-42 | 1.14E-40 |
| ENSGALG00000045228 | 0.556067637 | 0.002730325 | 0.006041116 |
| ENSGALG00000045305 | 0.666090893 | 7.23E-06 | 2.36E-05 |
| ENSGALG00000045350 | -0.940496219 | 8.85E-06 | 2.84E-05 |
| ENSGALG00000045478 | 1.319106178 | 2.23E-22 | 2.92E-21 |
| ENSGALG00000045534 | 0.47375577 | 0.006282137 | 0.012855586 |
| ENSGALG00000045584 | 1.227488645 | 1.54E-53 | 1.05E-51 |
| ENSGALG00000045602 | -1.38288185 | 9.99E-05 | 0.00027785 |
| ENSGALG00000045606 | -0.348673031 | 0.021525531 | 0.039171914 |
| ENSGALG00000045611 | -0.697556597 | 1.07E-11 | 6.34E-11 |
| ENSGALG00000045636 | 2.199933131 | 0.027887671 | 0.049368225 |

| | | | |
|---|---|---|---|
| ENSGALG00000045671 | -0.954025695 | 1.37E-18 | 1.41E-17 |
| ENSGALG00000045726 | -1.004880008 | 5.08E-27 | 8.98E-26 |
| ENSGALG00000045738 | 0.702575024 | 3.50E-05 | 0.000104136 |
| ENSGALG00000045796 | 6.039081378 | 1.73E-05 | 5.32E-05 |
| ENSGALG00000045842 | -0.943468077 | 0.005237075 | 0.010905826 |
| ENSGALG00000045873 | 2.064138623 | 0.000464523 | 0.001168789 |
| ENSGALG00000045907 | 1.071623704 | 0.014337626 | 0.027152991 |
| ENSGALG00000045953 | 0.515370156 | 0.001206301 | 0.002847029 |
| ENSGALG00000046098 | 0.911022167 | 8.71E-07 | 3.20E-06 |
| ENSGALG00000046141 | 3.721091723 | 0.017184563 | 0.03198748 |
| ENSGALG00000046160 | 0.518728666 | 0.000249912 | 0.000656007 |
| ENSGALG00000046235 | -0.553493549 | 0.005258246 | 0.010948236 |
| ENSGALG00000046258 | 0.498007473 | 2.68E-06 | 9.29E-06 |
| ENSGALG00000046371 | 0.356598162 | 0.00118475 | 0.002800056 |
| ENSGALG00000046412 | 1.917000841 | 2.66E-06 | 9.21E-06 |
| ENSGALG00000046444 | 2.02448032 | 0.016579907 | 0.030938235 |
| ENSGALG00000046517 | 1.712430121 | 0.018356183 | 0.033935968 |
| ENSGALG00000046593 | 0.654765717 | 0.010547599 | 0.020574478 |
| ENSGALG00000046646 | -0.85469505 | 0.000157994 | 0.000426251 |
| ENSGALG00000046681 | 1.355300937 | 3.99E-06 | 1.34E-05 |
| ENSGALG00000046714 | -0.786640259 | 0.000105048 | 0.000291213 |
| ENSGALG00000046739 | 1.835019164 | 4.00E-07 | 1.53E-06 |
| ENSGALG00000046740 | 0.373386466 | 0.003335399 | 0.007254979 |
| ENSGALG00000046743 | 0.720293728 | 0.013940216 | 0.026481539 |
| ENSGALG00000046817 | -0.49351042 | 0.011022893 | 0.021400251 |
| ENSGALG00000046828 | 1.053703376 | 1.39E-24 | 2.10E-23 |
| ENSGALG00000046843 | 1.041651262 | 4.22E-22 | 5.42E-21 |
| ENSGALG00000046864 | 1.539234422 | 0.013511736 | 0.025768418 |
| ENSGALG00000046889 | 1.589064575 | 2.37E-05 | 7.21E-05 |
| ENSGALG00000046939 | -1.182486694 | 6.57E-05 | 0.00018812 |
| ENSGALG00000046950 | 1.06575605 | 0.009554826 | 0.018780795 |
| ENSGALG00000046958 | 1.312337139 | 7.77E-10 | 3.87E-09 |
| ENSGALG00000047027 | -2.29606446 | 0.000414058 | 0.001050762 |
| ENSGALG00000047031 | -1.692710004 | 0.001102782 | 0.002619998 |
| ENSGALG00000047034 | 1.182533154 | 1.33E-14 | 1.01E-13 |
| ENSGALG00000047043 | 0.707424292 | 1.67E-05 | 5.17E-05 |
| ENSGALG00000047126 | 0.738412882 | 0.001604526 | 0.003699529 |
| ENSGALG00000047142 | -0.929838506 | 0.000526906 | 0.00131409 |
| ENSGALG00000047176 | 1.153278357 | 0.000174314 | 0.000467405 |
| ENSGALG00000047178 | 2.830501777 | 0.025925436 | 0.046237833 |
| ENSGALG00000047187 | -0.375752613 | 0.005582749 | 0.011554854 |
| ENSGALG00000047190 | -0.977984187 | 1.31E-07 | 5.28E-07 |
| ENSGALG00000047196 | 1.291485421 | 5.65E-25 | 8.80E-24 |
| ENSGALG00000047208 | -4.128247066 | 0.000396248 | 0.001008578 |
| ENSGALG00000047223 | -0.971656171 | 2.97E-05 | 8.90E-05 |
| ENSGALG00000047233 | 0.634633757 | 1.46E-07 | 5.86E-07 |
| ENSGALG00000047313 | 1.315940315 | 9.36E-08 | 3.82E-07 |
| ENSGALG00000047321 | -1.556641043 | 9.92E-07 | 3.61E-06 |

| | | | |
|---|---|---|---|
| ENSGALG00000047323 | 0.753030724 | 0.000364116 | 0.000932731 |
| ENSGALG00000047341 | 4.926619834 | 0.000292412 | 0.000759064 |
| ENSGALG00000047357 | 2.054896903 | 6.03E-06 | 1.98E-05 |
| ENSGALG00000047376 | 4.205184615 | 7.76E-14 | 5.55E-13 |
| ENSGALG00000047380 | 1.215511227 | 5.78E-13 | 3.86E-12 |
| ENSGALG00000047383 | -1.792643866 | 6.49E-65 | 7.17E-63 |
| ENSGALG00000047461 | -0.649481724 | 1.79E-07 | 7.08E-07 |
| ENSGALG00000047462 | 0.34558223 | 0.016943104 | 0.031559637 |
| ENSGALG00000047510 | 1.352299471 | 2.16E-06 | 7.58E-06 |
| ENSGALG00000047526 | 2.912018835 | 4.55E-05 | 0.000133611 |
| ENSGALG00000047536 | 1.641110694 | 3.44E-60 | 3.00E-58 |
| ENSGALG00000047566 | 0.788948532 | 2.64E-05 | 7.95E-05 |
| ENSGALG00000047574 | -1.169005079 | 0.006753233 | 0.013732787 |
| ENSGALG00000047627 | 0.664412774 | 0.000588279 | 0.001456078 |
| ENSGALG00000047664 | -0.865528756 | 1.56E-09 | 7.57E-09 |
| ENSGALG00000047673 | -1.212713111 | 3.72E-08 | 1.58E-07 |
| ENSGALG00000047695 | -0.804539491 | 7.48E-10 | 3.73E-09 |
| ENSGALG00000047704 | 1.215221492 | 7.37E-15 | 5.70E-14 |
| ENSGALG00000047761 | 1.55138587 | 1.47E-11 | 8.60E-11 |
| ENSGALG00000047797 | -0.672022297 | 0.001681499 | 0.0038652 |
| ENSGALG00000047813 | -0.782723843 | 1.05E-05 | 3.36E-05 |
| ENSGALG00000047828 | 1.579144581 | 3.14E-08 | 1.34E-07 |
| ENSGALG00000047847 | 1.064799407 | 1.72E-07 | 6.81E-07 |
| ENSGALG00000047857 | 2.299752916 | 1.86E-09 | 8.95E-09 |
| ENSGALG00000047878 | 2.369776106 | 0.027448502 | 0.048692253 |
| ENSGALG00000047879 | -0.6280919 | 0.011058726 | 0.021463688 |
| ENSGALG00000047924 | 0.270924782 | 0.001008918 | 0.002410697 |
| ENSGALG00000047956 | 0.438878121 | 0.000511624 | 0.001279247 |
| ENSGALG00000047967 | 4.241659847 | 0.005402015 | 0.011220078 |
| ENSGALG00000048034 | -0.43137231 | 2.61E-06 | 9.06E-06 |
| ENSGALG00000048048 | 0.706634282 | 0.008499383 | 0.016901772 |
| ENSGALG00000048099 | 1.959372336 | 0.002710445 | 0.006001705 |
| ENSGALG00000048170 | 1.370249313 | 0.000104693 | 0.000290287 |
| ENSGALG00000048205 | -1.40122027 | 1.45E-43 | 6.38E-42 |
| ENSGALG00000048222 | 0.733866743 | 0.009527819 | 0.018733129 |
| ENSGALG00000048239 | 1.089136723 | 1.74E-09 | 8.38E-09 |
| ENSGALG00000048280 | 0.48569168 | 0.004220292 | 0.008975468 |
| ENSGALG00000048286 | 1.156762913 | 9.09E-18 | 8.80E-17 |
| ENSGALG00000048291 | 1.060987452 | 0.000413151 | 0.001048851 |
| ENSGALG00000048310 | 2.834387161 | 0.00143095 | 0.003330388 |
| ENSGALG00000048321 | -1.587849304 | 1.09E-08 | 4.89E-08 |
| ENSGALG00000048337 | 4.136611393 | 2.90E-09 | 1.37E-08 |
| ENSGALG00000048343 | 1.207065971 | 0.011791144 | 0.02275525 |
| ENSGALG00000048399 | 1.932066572 | 2.80E-05 | 8.42E-05 |
| ENSGALG00000048446 | 2.905818547 | 0.011061876 | 0.021466735 |
| ENSGALG00000048452 | 3.377913106 | 0.024750222 | 0.044415767 |
| ENSGALG00000048497 | 1.320475825 | 0.002672767 | 0.005932111 |
| ENSGALG00000048534 | -1.290076292 | 0.003938296 | 0.008431141 |

| | | | |
|---|---|---|---|
| ENSGALG00000048542 | 1.043565526 | 1.62E-23 | 2.27E-22 |
| ENSGALG00000048556 | 0.702884413 | 3.36E-05 | 0.00010014 |
| ENSGALG00000048563 | 0.364029696 | 0.001518599 | 0.003518718 |
| ENSGALG00000048612 | 1.282445887 | 9.59E-37 | 3.00E-35 |
| ENSGALG00000048625 | 0.991014017 | 0.013816449 | 0.026261107 |
| ENSGALG00000048671 | 0.731669705 | 1.30E-06 | 4.67E-06 |
| ENSGALG00000048697 | 2.559547492 | 6.90E-05 | 0.00019688 |
| ENSGALG00000048698 | 0.325983418 | 0.009731879 | 0.019101183 |
| ENSGALG00000048712 | 0.769510751 | 0.016258154 | 0.030417183 |
| ENSGALG00000048769 | -0.98760632 | 0.002555622 | 0.005689765 |
| ENSGALG00000048771 | -0.828024067 | 0.012539349 | 0.024110175 |
| ENSGALG00000048776 | -0.816327285 | 0.006243063 | 0.0127814 |
| ENSGALG00000048802 | 0.732063339 | 0.018336177 | 0.03391282 |
| ENSGALG00000048814 | 3.315113406 | 1.93E-11 | 1.12E-10 |
| ENSGALG00000048869 | -1.858794909 | 3.11E-13 | 2.13E-12 |
| ENSGALG00000048883 | -1.105057521 | 1.03E-06 | 3.74E-06 |
| ENSGALG00000048887 | 1.996154568 | 2.46E-06 | 8.55E-06 |
| ENSGALG00000048900 | 3.587721199 | 3.03E-08 | 1.30E-07 |
| ENSGALG00000048920 | 1.28353765 | 1.63E-23 | 2.28E-22 |
| ENSGALG00000048925 | 2.412216887 | 0.00126043 | 0.002961866 |
| ENSGALG00000048936 | 0.407602901 | 0.008064343 | 0.016131059 |
| ENSGALG00000048962 | 1.324048384 | 0.009045543 | 0.017870206 |
| ENSGALG00000048981 | 0.864698152 | 0.000143057 | 0.00038811 |
| ENSGALG00000049122 | 0.826344824 | 6.99E-10 | 3.50E-09 |
| ENSGALG00000049131 | 1.450576601 | 0.001720284 | 0.003944348 |
| ENSGALG00000049136 | 0.539721449 | 0.002959586 | 0.006511282 |
| ENSGALG00000049152 | 1.494317448 | 0.005091147 | 0.01062473 |
| ENSGALG00000049191 | 3.395573358 | 0.00040292 | 0.001024793 |
| ENSGALG00000049217 | 1.027248439 | 0.002639173 | 0.005864254 |
| ENSGALG00000049219 | 3.351144989 | 0.018008535 | 0.033356774 |
| ENSGALG00000049257 | 1.971321511 | 2.79E-07 | 1.08E-06 |
| ENSGALG00000049308 | 2.115549647 | 0.001566394 | 0.003620206 |
| ENSGALG00000049315 | 4.242967082 | 0.003314961 | 0.007218352 |
| ENSGALG00000049324 | 0.41485171 | 0.017785425 | 0.032988468 |
| ENSGALG00000049326 | 2.213655549 | 0.001743118 | 0.003992386 |
| ENSGALG00000049343 | 0.922200795 | 4.27E-10 | 2.17E-09 |
| ENSGALG00000049347 | 0.542031448 | 0.01428878 | 0.027075575 |
| ENSGALG00000049348 | 0.644615454 | 0.002367661 | 0.005300816 |
| ENSGALG00000049364 | 0.972415522 | 1.24E-06 | 4.48E-06 |
| ENSGALG00000049372 | 0.693866617 | 3.50E-09 | 1.64E-08 |
| ENSGALG00000049374 | 1.391476506 | 0.000100869 | 0.000280448 |
| ENSGALG00000049379 | -3.979505865 | 0.000641109 | 0.0015769 |
| ENSGALG00000049395 | 0.677993001 | 2.35E-05 | 7.14E-05 |
| ENSGALG00000049397 | 1.444581899 | 0.012911588 | 0.024759355 |
| ENSGALG00000049402 | 1.288894054 | 0.006786554 | 0.013794354 |
| ENSGALG00000049420 | -1.481547213 | 3.10E-17 | 2.90E-16 |
| ENSGALG00000049473 | -1.003456456 | 0.000186169 | 0.000497036 |
| ENSGALG00000049484 | 1.098081156 | 2.45E-05 | 7.43E-05 |

| | | | |
|---|---|---|---|
| ENSGALG00000049579 | 1.165562417 | 0.003760465 | 0.008083542 |
| ENSGALG00000049587 | -1.096383051 | 0.004781116 | 0.010042487 |
| ENSGALG00000049588 | 0.79699031 | 1.12E-13 | 7.91E-13 |
| ENSGALG00000049609 | -0.526249238 | 0.005761453 | 0.011886714 |
| ENSGALG00000049684 | -1.669374134 | 8.19E-24 | 1.17E-22 |
| ENSGALG00000049711 | 0.678809772 | 5.91E-08 | 2.46E-07 |
| ENSGALG00000049735 | 1.70577812 | 9.59E-05 | 0.000267681 |
| ENSGALG00000049751 | 0.574937548 | 4.62E-10 | 2.34E-09 |
| ENSGALG00000049774 | -0.541320029 | 0.011595333 | 0.022415555 |
| ENSGALG00000049776 | 1.035458087 | 9.23E-07 | 3.38E-06 |
| ENSGALG00000049780 | -1.134284388 | 1.30E-05 | 4.09E-05 |
| ENSGALG00000049799 | 5.271079778 | 0.001127656 | 0.002674421 |
| ENSGALG00000049810 | 1.187536413 | 8.22E-05 | 0.000231963 |
| ENSGALG00000049811 | 1.203100154 | 0.001365142 | 0.00318705 |
| ENSGALG00000049843 | -0.633932631 | 0.009397345 | 0.018492646 |
| ENSGALG00000049851 | 0.687766677 | 1.05E-07 | 4.28E-07 |
| ENSGALG00000049870 | 2.38449226 | 0.007880473 | 0.01580047 |
| ENSGALG00000049871 | -1.508545809 | 0.001249342 | 0.002938915 |
| ENSGALG00000049875 | -2.070429959 | 9.95E-12 | 5.94E-11 |
| ENSGALG00000049949 | 0.657550449 | 0.00370899 | 0.007990584 |
| ENSGALG00000049966 | 0.588042691 | 0.024873624 | 0.044598334 |
| ENSGALG00000050005 | 0.956388699 | 0.027630261 | 0.048974542 |
| ENSGALG00000050009 | 0.607886395 | 0.001580274 | 0.003647942 |
| ENSGALG00000050026 | 2.330291224 | 0.000268276 | 0.000700425 |
| ENSGALG00000050091 | 1.236895925 | 4.04E-07 | 1.54E-06 |
| ENSGALG00000050127 | -1.17063668 | 1.21E-06 | 4.36E-06 |
| ENSGALG00000050178 | -0.444115541 | 0.000256467 | 0.000671786 |
| ENSGALG00000050180 | 1.582476159 | 0.000183461 | 0.000490479 |
| ENSGALG00000050186 | 1.140245122 | 4.12E-07 | 1.57E-06 |
| ENSGALG00000050205 | 0.546290131 | 0.014832272 | 0.028003923 |
| ENSGALG00000050225 | 2.556564724 | 0.022931428 | 0.041458362 |
| ENSGALG00000050235 | -0.739555208 | 0.00047451 | 0.00119061 |
| ENSGALG00000050268 | 4.120531307 | 9.19E-13 | 6.01E-12 |
| ENSGALG00000050269 | 0.427894438 | 0.005632105 | 0.011646367 |
| ENSGALG00000050281 | -3.985041357 | 0.014185229 | 0.026913126 |
| ENSGALG00000050329 | 1.972623462 | 0.02039606 | 0.037291263 |
| ENSGALG00000050332 | -1.046967907 | 3.12E-05 | 9.34E-05 |
| ENSGALG00000050342 | 2.10521465 | 9.35E-07 | 3.42E-06 |
| ENSGALG00000050374 | -0.484865792 | 0.006215721 | 0.012729258 |
| ENSGALG00000050406 | -1.679294017 | 5.29E-18 | 5.22E-17 |
| ENSGALG00000050425 | 0.927966965 | 3.94E-09 | 1.83E-08 |
| ENSGALG00000050441 | 1.832028806 | 0.005040735 | 0.010535701 |
| ENSGALG00000050442 | 3.254453811 | 5.19E-32 | 1.22E-30 |
| ENSGALG00000050454 | 2.924909197 | 1.24E-09 | 6.02E-09 |
| ENSGALG00000050548 | 1.215994275 | 0.00380188 | 0.008162241 |
| ENSGALG00000050564 | 0.658941888 | 0.02675418 | 0.04757854 |
| ENSGALG00000050596 | 1.278879026 | 4.99E-17 | 4.58E-16 |
| ENSGALG00000050597 | 1.440041275 | 0.015133376 | 0.02852487 |

| | | | |
|---|---|---|---|
| ENSGALG00000050608 | 1.167467235 | 0.000113396 | 0.000312443 |
| ENSGALG00000050627 | 0.724516887 | 0.016398642 | 0.03064206 |
| ENSGALG00000050656 | -1.077505705 | 0.007096504 | 0.014357798 |
| ENSGALG00000050668 | 0.586296105 | 0.015675761 | 0.02942886 |
| ENSGALG00000050742 | 0.454102959 | 0.000136365 | 0.000370707 |
| ENSGALG00000050796 | -0.649809132 | 0.001641312 | 0.003778576 |
| ENSGALG00000050799 | 1.146025362 | 1.65E-06 | 5.87E-06 |
| ENSGALG00000050814 | -1.480762744 | 3.61E-16 | 3.10E-15 |
| ENSGALG00000050834 | 1.1951409 | 1.91E-12 | 1.22E-11 |
| ENSGALG00000050856 | 1.500208974 | 0.003473207 | 0.007526758 |
| ENSGALG00000050860 | 2.890241614 | 0.001648717 | 0.003793695 |
| ENSGALG00000050918 | 1.920047819 | 0.004655856 | 0.009803625 |
| ENSGALG00000050931 | -0.56645 | 0.000106708 | 0.000295574 |
| ENSGALG00000050946 | 2.32269331 | 0.00033915 | 0.000873219 |
| ENSGALG00000051009 | 1.999130061 | 0.002533134 | 0.005645247 |
| ENSGALG00000051033 | 2.165655055 | 1.39E-16 | 1.23E-15 |
| ENSGALG00000051034 | 2.244155334 | 0.026512213 | 0.047203825 |
| ENSGALG00000051043 | -0.495385305 | 0.025711637 | 0.045910743 |
| ENSGALG00000051046 | 0.848168209 | 0.007271279 | 0.01467863 |
| ENSGALG00000051178 | 1.373976395 | 0.003335518 | 0.007254979 |
| ENSGALG00000051184 | 1.024235009 | 0.002697989 | 0.005975405 |
| ENSGALG00000051197 | 1.668378532 | 0.004106976 | 0.008768773 |
| ENSGALG00000051202 | 1.625726934 | 6.00E-16 | 5.05E-15 |
| ENSGALG00000051209 | 1.441692683 | 0.024732613 | 0.044401753 |
| ENSGALG00000051237 | 1.04660447 | 9.94E-08 | 4.05E-07 |
| ENSGALG00000051258 | 0.3797831 | 0.012381677 | 0.023830608 |
| ENSGALG00000051259 | -0.939967152 | 0.000446903 | 0.001128844 |
| ENSGALG00000051266 | 0.664262359 | 1.31E-07 | 5.26E-07 |
| ENSGALG00000051275 | 0.951823236 | 2.87E-07 | 1.11E-06 |
| ENSGALG00000051283 | 2.050768951 | 0.014342295 | 0.027158049 |
| ENSGALG00000051325 | -0.710376988 | 3.02E-05 | 9.05E-05 |
| ENSGALG00000051355 | 3.287334259 | 0.000126969 | 0.000347515 |
| ENSGALG00000051358 | -1.1686647 | 7.11E-05 | 0.000202506 |
| ENSGALG00000051361 | -1.184818083 | 0.002737111 | 0.006053175 |
| ENSGALG00000051372 | 0.462514662 | 0.009635506 | 0.01893664 |
| ENSGALG00000051398 | 1.292164707 | 7.02E-32 | 1.64E-30 |
| ENSGALG00000051428 | 1.0370673 | 5.85E-13 | 3.90E-12 |
| ENSGALG00000051534 | 1.445649711 | 6.49E-28 | 1.21E-26 |
| ENSGALG00000051598 | 1.013037468 | 1.12E-17 | 1.09E-16 |
| ENSGALG00000051612 | 1.508110563 | 1.52E-13 | 1.07E-12 |
| ENSGALG00000051624 | 1.78083856 | 9.65E-12 | 5.77E-11 |
| ENSGALG00000051644 | -0.850431011 | 0.021094147 | 0.038464132 |
| ENSGALG00000051776 | 2.701808655 | 5.37E-05 | 0.000155909 |
| ENSGALG00000051806 | 1.590414955 | 0.007920381 | 0.015873846 |
| ENSGALG00000051832 | 4.719883076 | 1.41E-08 | 6.28E-08 |
| ENSGALG00000051843 | 2.54336192 | 0.000602834 | 0.001488683 |
| ENSGALG00000051869 | 2.251163887 | 0.000242142 | 0.00063721 |
| ENSGALG00000051879 | 1.459154256 | 0.008501982 | 0.016904468 |

| | | | |
|---|---|---|---|
| ENSGALG00000051922 | 0.813940409 | 9.23E-12 | 5.53E-11 |
| ENSGALG00000051926 | 1.28881106 | 3.31E-11 | 1.87E-10 |
| ENSGALG00000051944 | 1.509391191 | 0.011044239 | 0.021438631 |
| ENSGALG00000051984 | -1.373101863 | 9.17E-08 | 3.75E-07 |
| ENSGALG00000052012 | -0.810695229 | 0.000110757 | 0.000306039 |
| ENSGALG00000052022 | 0.840974846 | 3.45E-06 | 1.17E-05 |
| ENSGALG00000052037 | -0.636679551 | 0.007053306 | 0.014281298 |
| ENSGALG00000052043 | 0.463934501 | 0.003351196 | 0.007282783 |
| ENSGALG00000052056 | 0.72058312 | 2.12E-09 | 1.01E-08 |
| ENSGALG00000052154 | -1.869522251 | 1.02E-60 | 9.18E-59 |
| ENSGALG00000052168 | 1.850244436 | 0.002568139 | 0.005715759 |
| ENSGALG00000052183 | -1.342800975 | 0.00101687 | 0.00242905 |
| ENSGALG00000052199 | 0.811859647 | 0.013695322 | 0.026074634 |
| ENSGALG00000052202 | -0.740463576 | 0.000112632 | 0.00031059 |
| ENSGALG00000052210 | -0.420074234 | 0.000915726 | 0.002198647 |
| ENSGALG00000052213 | 3.286320827 | 9.04E-122 | 5.59E-119 |
| ENSGALG00000052233 | 1.135004575 | 3.08E-05 | 9.21E-05 |
| ENSGALG00000052240 | 0.968548442 | 0.000107274 | 0.000296959 |
| ENSGALG00000052244 | 0.886271305 | 0.000123496 | 0.000338624 |
| ENSGALG00000052265 | 4.696855534 | 2.99E-05 | 8.96E-05 |
| ENSGALG00000052302 | -0.613409738 | 0.009296606 | 0.018312967 |
| ENSGALG00000052318 | 1.85400788 | 3.53E-15 | 2.80E-14 |
| ENSGALG00000052326 | -1.58880112 | 2.00E-39 | 6.95E-38 |
| ENSGALG00000052348 | -1.658934679 | 0.00018545 | 0.000495407 |
| ENSGALG00000052368 | 1.987277445 | 0.004119911 | 0.008792249 |
| ENSGALG00000052374 | 3.307383638 | 0.017794817 | 0.033001386 |
| ENSGALG00000052375 | -2.408761047 | 0.001386482 | 0.00323243 |
| ENSGALG00000052396 | -1.673566962 | 3.07E-16 | 2.65E-15 |
| ENSGALG00000052411 | 1.299195797 | 0.017630767 | 0.032723936 |
| ENSGALG00000052441 | 1.919437536 | 0.002253099 | 0.005056823 |
| ENSGALG00000052449 | 1.406496884 | 0.005042847 | 0.010538496 |
| ENSGALG00000052452 | 1.165052021 | 0.000171784 | 0.000461258 |
| ENSGALG00000052502 | 1.707180603 | 0.000805754 | 0.001950454 |
| ENSGALG00000052538 | -0.803940084 | 0.027298211 | 0.048450939 |
| ENSGALG00000052541 | 2.050571071 | 5.41E-21 | 6.42E-20 |
| ENSGALG00000052568 | 1.393880643 | 1.40E-12 | 9.01E-12 |
| ENSGALG00000052586 | 2.339843889 | 0.00185912 | 0.004235527 |
| ENSGALG00000052647 | 0.371134405 | 0.000131286 | 0.000358034 |
| ENSGALG00000052652 | 1.71557116 | 0.014229947 | 0.026986666 |
| ENSGALG00000052672 | 1.468856575 | 9.36E-47 | 4.91E-45 |
| ENSGALG00000052682 | -1.163857322 | 0.000186156 | 0.000497036 |
| ENSGALG00000052732 | 3.93038805 | 0.001605006 | 0.003700006 |
| ENSGALG00000052741 | 4.51042587 | 0.002379831 | 0.00532543 |
| ENSGALG00000052766 | 1.154645632 | 0.015074085 | 0.02842494 |
| ENSGALG00000052786 | 1.288689341 | 0.000187758 | 0.000501081 |
| ENSGALG00000052797 | 0.907469087 | 0.00463317 | 0.009760393 |
| ENSGALG00000052799 | 0.805786478 | 0.014668024 | 0.027728481 |
| ENSGALG00000052882 | 1.071963292 | 4.28E-12 | 2.64E-11 |

| | | | |
|---|---|---|---|
| ENSGALG00000052907 | 2.758678548 | 2.65E-05 | 7.99E-05 |
| ENSGALG00000052918 | 2.668696266 | 3.51E-08 | 1.49E-07 |
| ENSGALG00000052919 | 1.533171713 | 0.007337793 | 0.014794047 |
| ENSGALG00000052920 | 2.327689525 | 8.46E-06 | 2.73E-05 |
| ENSGALG00000052938 | 1.056375432 | 3.96E-09 | 1.84E-08 |
| ENSGALG00000052946 | 1.34453988 | 0.001847895 | 0.004212072 |
| ENSGALG00000052949 | 3.626824307 | 8.02E-06 | 2.59E-05 |
| ENSGALG00000052971 | 0.367168439 | 0.008526677 | 0.016943653 |
| ENSGALG00000052976 | 0.63889392 | 0.006654186 | 0.013553679 |
| ENSGALG00000052983 | -2.052933786 | 0.000249764 | 0.000655805 |
| ENSGALG00000053019 | 0.881715362 | 1.06E-05 | 3.39E-05 |
| ENSGALG00000053025 | -0.907834057 | 2.58E-08 | 1.11E-07 |
| ENSGALG00000053035 | -1.049746272 | 0.026934301 | 0.047862649 |
| ENSGALG00000053041 | 0.361128659 | 0.000686605 | 0.001680905 |
| ENSGALG00000053046 | 1.690693063 | 0.001419951 | 0.003305922 |
| ENSGALG00000053077 | 1.060827516 | 1.68E-21 | 2.07E-20 |
| ENSGALG00000053107 | 0.891685995 | 4.78E-08 | 2.01E-07 |
| ENSGALG00000053132 | 0.667114561 | 0.027402259 | 0.048616564 |
| ENSGALG00000053194 | -0.360052334 | 0.002070378 | 0.004679956 |
| ENSGALG00000053218 | 0.842639108 | 0.000192653 | 0.000513438 |
| ENSGALG00000053243 | -0.603134252 | 0.012979164 | 0.024871394 |
| ENSGALG00000053245 | 4.954603273 | 3.10E-06 | 1.06E-05 |
| ENSGALG00000053274 | 1.529801619 | 2.62E-10 | 1.37E-09 |
| ENSGALG00000053332 | 0.804843107 | 3.77E-09 | 1.76E-08 |
| ENSGALG00000053338 | 2.100448382 | 0.012614336 | 0.024233788 |
| ENSGALG00000053371 | -1.601819284 | 1.52E-25 | 2.45E-24 |
| ENSGALG00000053374 | 1.364706746 | 0.018592325 | 0.034330509 |
| ENSGALG00000053417 | 1.231277174 | 0.001840986 | 0.004199142 |
| ENSGALG00000053430 | -5.734159483 | 1.49E-05 | 4.63E-05 |
| ENSGALG00000053510 | -0.275794472 | 0.025876283 | 0.046162283 |
| ENSGALG00000053555 | 1.570074989 | 0.019492165 | 0.035850814 |
| ENSGALG00000053585 | 2.068881563 | 0.000955438 | 0.002289548 |
| ENSGALG00000053659 | -1.923721806 | 2.83E-06 | 9.79E-06 |
| ENSGALG00000053663 | -0.554765214 | 5.04E-05 | 0.000146984 |
| ENSGALG00000053665 | 1.085933607 | 1.45E-08 | 6.44E-08 |
| ENSGALG00000053675 | 0.520996982 | 8.76E-05 | 0.00024622 |
| ENSGALG00000053765 | 0.563910548 | 3.93E-08 | 1.67E-07 |
| ENSGALG00000053766 | 1.220004355 | 2.95E-11 | 1.68E-10 |
| ENSGALG00000053770 | 3.785557417 | 0.002999386 | 0.00659092 |
| ENSGALG00000053778 | 2.446990941 | 0.015263063 | 0.028741417 |
| ENSGALG00000053784 | -1.327589184 | 0.00391913 | 0.008395401 |
| ENSGALG00000053860 | -1.17889863 | 1.89E-13 | 1.32E-12 |
| ENSGALG00000053875 | -0.767760633 | 8.86E-09 | 4.00E-08 |
| ENSGALG00000053878 | -1.066297643 | 0.006941612 | 0.014082148 |
| ENSGALG00000053889 | 0.571418526 | 0.000473289 | 0.001187767 |
| ENSGALG00000053912 | 1.708204494 | 0.004866556 | 0.010203027 |
| ENSGALG00000053950 | 1.940702861 | 0.01185141 | 0.022865063 |
| ENSGALG00000053980 | 0.969121887 | 0.002407139 | 0.005381222 |

| | | | |
|---|---|---|---|
| ENSGALG00000053990 | -1.03225458 | 8.65E-05 | 0.000243215 |
| ENSGALG00000054026 | 1.183252722 | 0.000455003 | 0.001146745 |
| ENSGALG00000054044 | -0.460284702 | 0.001161811 | 0.00274871 |
| ENSGALG00000054048 | 1.993662813 | 2.51E-88 | 6.57E-86 |
| ENSGALG00000054057 | 0.59199151 | 3.06E-06 | 1.05E-05 |
| ENSGALG00000054059 | 0.554628876 | 0.003669589 | 0.007915736 |
| ENSGALG00000054070 | 1.493656937 | 5.00E-15 | 3.92E-14 |
| ENSGALG00000054102 | 1.594305615 | 3.81E-05 | 0.000112945 |
| ENSGALG00000054145 | -0.552771325 | 0.000593214 | 0.001466794 |
| ENSGALG00000054164 | 1.544390124 | 8.24E-16 | 6.87E-15 |
| ENSGALG00000054173 | -0.745034805 | 0.005074947 | 0.010598015 |
| ENSGALG00000054244 | -0.425936832 | 0.00379934 | 0.008158077 |
| ENSGALG00000054370 | 1.36163025 | 0.024614929 | 0.044225528 |
| ENSGALG00000054387 | 1.469036425 | 9.23E-07 | 3.37E-06 |
| ENSGALG00000054424 | 0.97065821 | 0.001271923 | 0.002983257 |
| ENSGALG00000054505 | 1.103911031 | 0.020509625 | 0.037483773 |
| ENSGALG00000054512 | 1.564991714 | 0.000317853 | 0.000821031 |
| ENSGALG00000054546 | 2.876938204 | 0.014472653 | 0.02737819 |
| ENSGALG00000054550 | 2.483944649 | 0.000144962 | 0.000393121 |
| ENSGALG00000054599 | 0.825894485 | 2.51E-11 | 1.44E-10 |
| ENSGALG00000054600 | 1.118091556 | 1.58E-05 | 4.89E-05 |
| ENSGALG00000054624 | -1.961210163 | 0.005027508 | 0.010511289 |
| ENSGALG00000054634 | 1.220560794 | 6.32E-08 | 2.63E-07 |
| ENSGALG00000054642 | 1.312171301 | 0.011009996 | 0.021384377 |
| ENSGALG00000054657 | 1.089806858 | 2.41E-06 | 8.40E-06 |
| ENSGALG00000054689 | 3.736737613 | 1.43E-06 | 5.11E-06 |
| ENSGALG00000054713 | 0.487411125 | 0.021197919 | 0.03863023 |
| ENSGALG00000054732 | 2.383044226 | 0.000161921 | 0.000435808 |
| ENSGALG00000054746 | 0.664093751 | 4.74E-08 | 1.99E-07 |
| ENSGALG00000054753 | -1.374049226 | 1.10E-06 | 4.00E-06 |
| ENSGALG00000054754 | 2.097905335 | 0.004433741 | 0.009372226 |
| ENSGALG00000054765 | 0.458075011 | 0.025022855 | 0.044825466 |
| ENSGALG00000054787 | 2.444886565 | 0.00322755 | 0.007043811 |
| ENSGALG00000054809 | 1.719859597 | 0.006691127 | 0.013622798 |
| ENSGALG00000054915 | -0.598996925 | 0.014190877 | 0.026920083 |
| ENSGALG00000054937 | 3.116242651 | 6.55E-26 | 1.08E-24 |
| ENSGALG00000054956 | 1.005736658 | 9.70E-10 | 4.77E-09 |
| ENSGALG00000054981 | 2.827164367 | 0.00301234 | 0.006615564 |
| ENSGALG00000055002 | 1.117861066 | 0.001904748 | 0.004330045 |
| ENSGALG00000055025 | -2.894621452 | 0.005780106 | 0.011919769 |
| ENSGALG00000055026 | 4.647350088 | 3.54E-12 | 2.21E-11 |
| ENSGALG00000055028 | 0.907579158 | 0.002397228 | 0.005361713 |
| ENSGALG00000055035 | 0.85662859 | 0.003111177 | 0.00681391 |
| ENSGALG00000055056 | 2.583719545 | 1.09E-24 | 1.65E-23 |
| ENSGALG00000055058 | -3.118968015 | 6.29E-131 | 4.50E-128 |
| ENSGALG00000055092 | 1.242541727 | 0.001983695 | 0.004496728 |
| ENSGALG00000055094 | -0.693785461 | 4.16E-11 | 2.33E-10 |
| ENSGALG00000055127 | 0.968395615 | 0.002537822 | 0.005653839 |

| | | | |
|---|---|---|---|
| ENTPD4 | 0.357030662 | 2.27E-05 | 6.92E-05 |
| ENTPD5 | 1.076135788 | 1.08E-27 | 1.99E-26 |
| ENTPD6 | 0.664136688 | 4.93E-11 | 2.74E-10 |
| ENY2 | 0.678912479 | 1.04E-07 | 4.23E-07 |
| EOGT | 0.552339949 | 1.72E-07 | 6.81E-07 |
| EOMES | -2.760147076 | 3.12E-21 | 3.77E-20 |
| EP300 | 0.368612208 | 0.003336267 | 0.007255446 |
| EPB41 | -1.290462095 | 0.001501567 | 0.003481627 |
| EPB41L5 | -1.523571935 | 0.001396213 | 0.003254558 |
| EPC2 | -1.213538656 | 0.000527895 | 0.001316299 |
| EPDR1 | 0.937120802 | 9.24E-30 | 1.92E-28 |
| EPGN | -0.871209805 | 7.94E-05 | 0.000224349 |
| EPHA2 | -0.337915848 | 5.72E-06 | 1.89E-05 |
| EPHA3 | 0.508164034 | 4.01E-09 | 1.87E-08 |
| EPHB1 | 0.265046421 | 0.024167394 | 0.04350195 |
| EPHB3 | -1.358559788 | 9.05E-34 | 2.32E-32 |
| EPHX2 | -0.4867912 | 0.001100141 | 0.002615095 |
| EPM2A | 0.567139545 | 0.00011643 | 0.000320088 |
| EPRS | -1.058777419 | 4.21E-33 | 1.04E-31 |
| EPS15L1 | -1.087257265 | 2.48E-11 | 1.42E-10 |
| ERAL1 | -0.637236599 | 4.21E-05 | 0.000124056 |
| ERAP1 | -0.874272078 | 8.36E-08 | 3.43E-07 |
| ERBB2 | -0.544705657 | 0.002995778 | 0.006585575 |
| ERBB3 | 6.687134051 | 2.11E-09 | 1.01E-08 |
| ERBIN | 0.477364131 | 0.002126819 | 0.004797963 |
| ERCC3 | -0.693839051 | 1.78E-08 | 7.80E-08 |
| ERCC4 | -1.209181853 | 1.68E-13 | 1.17E-12 |
| ERCC5 | -0.32645969 | 0.02784769 | 0.049316718 |
| ERCC6L | -1.443421651 | 0.000620642 | 0.00153016 |
| ERG | 1.135515799 | 4.38E-27 | 7.77E-26 |
| ERGIC1 | 0.31796406 | 1.29E-05 | 4.05E-05 |
| ERGIC2 | 1.315493922 | 8.32E-73 | 1.23E-70 |
| ERGIC3 | -0.401863972 | 4.26E-05 | 0.000125291 |
| ERH | 0.552664571 | 7.17E-07 | 2.66E-06 |
| ERI1 | 0.618524787 | 2.21E-12 | 1.39E-11 |
| ERICH1 | -0.858763858 | 2.26E-06 | 7.88E-06 |
| ERLIN1 | 0.231907981 | 0.005930735 | 0.012204474 |
| ERLIN2 | 0.511661378 | 2.22E-09 | 1.06E-08 |
| ERMP1 | -0.754563697 | 0.004312485 | 0.009140073 |
| ERN1 | 0.254649939 | 0.016647931 | 0.03105664 |
| ERO1A | -0.52033994 | 1.15E-05 | 3.64E-05 |
| ERRFI1 | 0.817307423 | 8.93E-19 | 9.32E-18 |
| ESCO1 | -0.868273995 | 2.04E-09 | 9.75E-09 |
| ESCO2 | -1.378657444 | 9.47E-07 | 3.46E-06 |
| ESRRG | 0.770310532 | 2.32E-07 | 9.07E-07 |
| ESS2 | -0.610990045 | 2.47E-09 | 1.17E-08 |
| ETF1 | 0.431108165 | 8.28E-08 | 3.40E-07 |
| ETNK1 | -0.556856344 | 0.010297001 | 0.020102965 |

| | | | |
|---|---|---|---|
| ETNPPL | 1.70268866 | 0.015156981 | 0.028561443 |
| ETV1 | 0.50603795 | 1.62E-06 | 5.76E-06 |
| ETV3 | 0.563652407 | 0.017842922 | 0.033081571 |
| ETV5 | -1.098346543 | 2.79E-10 | 1.45E-09 |
| ETV6 | 0.290455744 | 0.007356145 | 0.01482573 |
| EVC | -0.47447179 | 0.005488971 | 0.011383291 |
| EVI5 | 0.483235966 | 2.49E-09 | 1.18E-08 |
| EVL | 1.227679784 | 1.18E-25 | 1.93E-24 |
| EWSR1 | 0.313814902 | 0.00023525 | 0.000619914 |
| EXD2 | -0.309709005 | 0.014726515 | 0.027831312 |
| EXFABP | 1.183188612 | 0.001108575 | 0.002632842 |
| EXO1 | -2.539741662 | 4.86E-07 | 1.84E-06 |
| EXOC1 | -0.841861594 | 2.15E-17 | 2.03E-16 |
| EXOC2 | -0.465387425 | 0.001994281 | 0.004517718 |
| EXOC3 | -0.906542125 | 1.27E-21 | 1.56E-20 |
| EXOC6 | -0.350569221 | 0.021207836 | 0.038640335 |
| EXOC8 | -1.168524884 | 3.42E-12 | 2.13E-11 |
| EXOSC1 | -0.4370552 | 0.003468502 | 0.007518254 |
| EXOSC10 | -1.05300071 | 8.17E-21 | 9.62E-20 |
| EXOSC3 | -1.535469885 | 5.51E-20 | 6.19E-19 |
| EXOSC8 | -0.86943637 | 1.99E-10 | 1.05E-09 |
| EXOSC9 | -1.27702011 | 3.19E-22 | 4.13E-21 |
| EXT1 | 0.530033854 | 4.15E-10 | 2.12E-09 |
| EXT2 | -0.833168379 | 1.27E-19 | 1.40E-18 |
| EXTL2 | 0.530099673 | 0.003310823 | 0.007210497 |
| EXTL3 | 1.071704683 | 5.40E-29 | 1.07E-27 |
| EYA3 | 0.529065238 | 7.91E-09 | 3.58E-08 |
| EYA4 | 0.732325018 | 6.19E-06 | 2.03E-05 |
| EZH2 | -0.255699987 | 0.009980051 | 0.019551579 |
| EZR | -1.935417047 | 2.18E-81 | 4.42E-79 |
| F10 | -0.770600163 | 0.000152308 | 0.000412056 |
| F13A1 | 1.216077197 | 1.28E-06 | 4.60E-06 |
| F13B | 3.829276542 | 0.000664656 | 0.001630393 |
| F2RL1 | 0.404898085 | 1.87E-06 | 6.63E-06 |
| F2RL2 | 0.59666194 | 8.34E-06 | 2.69E-05 |
| F2RL3 | -1.648727785 | 5.58E-05 | 0.000161587 |
| F3 | 0.296196496 | 0.005847749 | 0.012048294 |
| F8 | -1.994743526 | 2.04E-10 | 1.07E-09 |
| FA2H | -1.519734837 | 0.013528782 | 0.025797307 |
| FAAH2 | -0.370392504 | 0.003087211 | 0.006763599 |
| FAAP24 | -0.779829651 | 1.29E-06 | 4.65E-06 |
| FABP6 | 1.219211685 | 3.48E-10 | 1.79E-09 |
| FABP7 | 0.36659784 | 0.000204774 | 0.000543505 |
| FAF1 | -0.385866496 | 0.002251524 | 0.00505421 |
| FAF2 | -0.325830834 | 4.14E-05 | 0.000121996 |
| FAH | 0.86014743 | 1.31E-10 | 6.99E-10 |
| FAHD1 | -0.567712628 | 1.53E-07 | 6.11E-07 |
| FAHD2A | 0.556710424 | 3.18E-10 | 1.64E-09 |

| | | | |
|---|---|---|---|
| FAIM | 1.150233639 | 3.72E-18 | 3.72E-17 |
| FAM102A | -1.559733175 | 7.24E-07 | 2.69E-06 |
| FAM102B | -0.793642878 | 7.79E-13 | 5.13E-12 |
| FAM103A1 | 0.797660888 | 2.19E-18 | 2.23E-17 |
| FAM105A | 0.514414147 | 1.42E-08 | 6.30E-08 |
| FAM107B | -1.266144652 | 1.80E-36 | 5.51E-35 |
| FAM110B | -0.569855051 | 0.002196183 | 0.004943776 |
| FAM114A1 | -0.475987309 | 4.72E-06 | 1.57E-05 |
| FAM114A2 | -0.98555399 | 1.15E-18 | 1.20E-17 |
| FAM117A | -0.782446592 | 1.50E-08 | 6.65E-08 |
| FAM118B | -0.780293663 | 5.82E-14 | 4.22E-13 |
| FAM120B | -1.08303596 | 4.35E-12 | 2.68E-11 |
| FAM122A | 1.313135429 | 5.56E-49 | 3.24E-47 |
| FAM126A | -0.396009665 | 1.32E-05 | 4.14E-05 |
| FAM129A | -0.452680731 | 0.009821406 | 0.019262994 |
| FAM133B | 0.535427158 | 6.51E-06 | 2.13E-05 |
| FAM135A | -0.33452935 | 0.004071666 | 0.008704322 |
| FAM13A | -0.508398716 | 0.002748486 | 0.006075365 |
| FAM13B | 0.991006382 | 1.14E-15 | 9.34E-15 |
| FAM13C | -1.08687608 | 1.28E-05 | 4.01E-05 |
| FAM149A | -1.320586545 | 1.43E-05 | 4.45E-05 |
| FAM149B1 | 1.275298446 | 7.37E-29 | 1.45E-27 |
| FAM160A1 | -0.396089458 | 0.003799017 | 0.008158077 |
| FAM160B1 | -0.364776176 | 0.002735275 | 0.006050098 |
| FAM161A | 0.517295407 | 0.00781945 | 0.015696643 |
| FAM162A | 0.593806759 | 1.99E-06 | 7.03E-06 |
| FAM168A | 0.675982992 | 1.49E-06 | 5.32E-06 |
| FAM168B | 0.695628709 | 5.00E-12 | 3.06E-11 |
| FAM171B | 1.177609242 | 4.90E-05 | 0.000143081 |
| FAM172A | 0.870551707 | 1.18E-12 | 7.63E-12 |
| FAM173A | -0.341801446 | 0.021600648 | 0.039292828 |
| FAM174A | -0.49940052 | 6.59E-05 | 0.000188758 |
| FAM175A | -0.379520912 | 0.010697365 | 0.020827758 |
| FAM175B | -0.435985156 | 0.000596596 | 0.001474888 |
| FAM177A1 | 0.864675414 | 2.82E-28 | 5.36E-27 |
| FAM179A | 3.01641461 | 5.42E-05 | 0.000157337 |
| FAM180B | 2.57988003 | 5.26E-10 | 2.65E-09 |
| FAM184A | -1.575078325 | 1.85E-07 | 7.31E-07 |
| FAM185A | -1.260204503 | 1.15E-17 | 1.11E-16 |
| FAM189A2 | -0.819329475 | 0.001697396 | 0.003897129 |
| FAM192A | 0.728828271 | 4.63E-12 | 2.85E-11 |
| FAM193B | -1.220292355 | 6.50E-09 | 2.97E-08 |
| FAM204A | 0.876345704 | 6.05E-21 | 7.16E-20 |
| FAM20A | 1.008704907 | 4.35E-11 | 2.43E-10 |
| FAM20B | 0.321962343 | 0.008304859 | 0.016568273 |
| FAM20C | 1.265846782 | 1.04E-13 | 7.39E-13 |
| FAM210B | 0.515100857 | 0.002673608 | 0.005933008 |
| FAM214A | -0.40201448 | 0.000157889 | 0.000426054 |

| | | | |
|---|---|---|---|
| FAM214B | -0.694513976 | 0.00175343 | 0.004014254 |
| FAM217B | 0.824821321 | 8.45E-11 | 4.61E-10 |
| FAM219A | 0.928898795 | 7.99E-07 | 2.95E-06 |
| FAM221A | 1.127105871 | 0.005432392 | 0.011278001 |
| FAM222B | 1.391992275 | 4.46E-14 | 3.27E-13 |
| FAM234A | -0.386958276 | 3.84E-05 | 0.000113602 |
| FAM234B | -1.106754381 | 2.80E-08 | 1.20E-07 |
| FAM241A | 0.907668657 | 0.000788891 | 0.001912361 |
| FAM32A | 0.792307938 | 5.14E-13 | 3.46E-12 |
| FAM3C | 0.32231391 | 0.006167287 | 0.012641498 |
| FAM3D | -1.454181735 | 0.002891277 | 0.006374411 |
| FAM43A | 0.985152841 | 9.98E-06 | 3.19E-05 |
| FAM45A | 0.609094639 | 8.78E-11 | 4.78E-10 |
| FAM46A | 0.28807254 | 0.010733771 | 0.020892655 |
| FAM46C | -0.963831437 | 0.026589765 | 0.047329502 |
| FAM53A | -1.286563164 | 3.41E-31 | 7.64E-30 |
| FAM53B | 1.808234802 | 5.30E-07 | 2.00E-06 |
| FAM60A | 0.843239288 | 2.69E-28 | 5.13E-27 |
| FAM76B | -1.592685139 | 5.00E-17 | 4.58E-16 |
| FAM81A | -1.092788075 | 9.09E-06 | 2.92E-05 |
| FAM83B | -0.64915986 | 1.14E-06 | 4.13E-06 |
| FAM83G | -1.608702165 | 3.63E-07 | 1.39E-06 |
| FAM83H | -1.01234651 | 1.05E-05 | 3.33E-05 |
| FAM89A | 0.890388055 | 6.66E-17 | 6.05E-16 |
| FAM8A1 | -0.436314936 | 5.71E-05 | 0.000165089 |
| FAM96A | 0.47143423 | 0.000147228 | 0.000398868 |
| FAM98A | -1.099673593 | 1.43E-14 | 1.08E-13 |
| FAN1 | -0.963614545 | 2.73E-08 | 1.17E-07 |
| FANCA | 0.78149222 | 2.38E-09 | 1.13E-08 |
| FANCD2 | 1.407757742 | 1.38E-22 | 1.83E-21 |
| FANCL | 0.911779728 | 7.08E-10 | 3.54E-09 |
| FANCM | -0.813679572 | 0.000666123 | 0.001633697 |
| FAP | -0.40468959 | 1.91E-06 | 6.76E-06 |
| FAR1 | -0.50878419 | 1.31E-07 | 5.28E-07 |
| FARP1 | 0.666857587 | 1.36E-11 | 8.03E-11 |
| FARP2 | -0.339275422 | 0.01228701 | 0.023658456 |
| FARSA | -0.534743382 | 3.54E-07 | 1.36E-06 |
| FARSB | 0.557982322 | 9.17E-08 | 3.75E-07 |
| FAS | -0.794091097 | 1.12E-09 | 5.48E-09 |
| FASN | -1.449809396 | 8.74E-13 | 5.72E-12 |
| FASTK | 1.668991392 | 2.74E-34 | 7.24E-33 |
| FASTKD1 | -0.684033163 | 0.000368099 | 0.0009424 |
| FASTKD2 | -0.534579002 | 4.52E-06 | 1.51E-05 |
| FASTKD3 | -0.639111992 | 0.00150892 | 0.00349808 |
| FAT1 | -0.839705715 | 4.30E-12 | 2.65E-11 |
| FAU | 0.382086343 | 0.00019998 | 0.000531611 |
| FBLN1 | -1.236518245 | 5.30E-45 | 2.49E-43 |
| FBLN2 | -2.55637577 | 3.77E-140 | 3.01E-137 |

| | | | |
|---|---|---|---|
| FBN1 | -2.312322744 | 2.05E-65 | 2.30E-63 |
| FBN2 | 0.818684223 | 1.99E-08 | 8.69E-08 |
| FBXL15 | -1.736378763 | 3.49E-27 | 6.24E-26 |
| FBXL2 | 0.969178914 | 3.48E-08 | 1.48E-07 |
| FBXL22 | 2.91751344 | 2.92E-14 | 2.17E-13 |
| FBXL3 | -1.063964374 | 2.95E-10 | 1.53E-09 |
| FBXL4 | -0.807054082 | 0.001572494 | 0.003633069 |
| FBXL5 | -0.245827942 | 0.009151628 | 0.018056165 |
| FBXL7 | -0.762270306 | 0.000191954 | 0.000511777 |
| FBXO11 | -0.302962231 | 0.003675966 | 0.007928233 |
| FBXO18 | -1.186505292 | 2.74E-13 | 1.88E-12 |
| FBXO2 | -1.355666211 | 3.14E-12 | 1.97E-11 |
| FBXO21 | -0.638098968 | 1.73E-05 | 5.34E-05 |
| FBXO22 | -0.60247768 | 3.74E-13 | 2.55E-12 |
| FBXO30 | -1.448373588 | 1.85E-25 | 2.99E-24 |
| FBXO31 | -1.977969198 | 1.40E-18 | 1.44E-17 |
| FBXO32 | -2.001139507 | 1.63E-26 | 2.79E-25 |
| FBXO33 | -0.457355438 | 0.000217753 | 0.000576041 |
| FBXO34 | -0.411125819 | 0.002318188 | 0.005194332 |
| FBXO4 | 0.491181673 | 0.012891711 | 0.024728215 |
| FBXO44 | -1.770022068 | 1.80E-40 | 6.53E-39 |
| FBXO48 | -1.224572338 | 2.17E-18 | 2.21E-17 |
| FBXO5 | 0.639608962 | 0.027661623 | 0.049012747 |
| FBXO7 | 0.243145395 | 0.012706406 | 0.024396874 |
| FBXO8 | 0.465851197 | 0.000515208 | 0.001287262 |
| FBXO9 | 0.736743187 | 1.12E-15 | 9.24E-15 |
| FBXW11 | 0.343910807 | 0.001687675 | 0.003876776 |
| FBXW2 | 0.588670715 | 3.55E-11 | 2.00E-10 |
| FBXW5 | 0.490488498 | 1.65E-07 | 6.59E-07 |
| FBXW7 | -0.42385282 | 0.001294288 | 0.003032052 |
| FBXW8 | 0.492765721 | 0.002347323 | 0.005258747 |
| FCHSD1 | -0.631881806 | 0.00013271 | 0.000361628 |
| FCHSD2 | -1.018291228 | 0.004491219 | 0.009481927 |
| FDFT1 | -0.439829177 | 1.44E-05 | 4.48E-05 |
| FDPS | -0.770932518 | 4.78E-11 | 2.66E-10 |
| FDXR | -0.508387488 | 0.000158651 | 0.000427938 |
| FEM1A | -0.697102642 | 2.91E-06 | 1.00E-05 |
| FEM1B | 1.048403378 | 8.81E-26 | 1.45E-24 |
| FEM1C | -0.619391153 | 0.00055945 | 0.001389374 |
| FEN1 | -0.371878654 | 0.01404096 | 0.026658014 |
| FER | -0.381971281 | 0.019980824 | 0.036625672 |
| FER1L6 | 2.095240791 | 2.26E-05 | 6.87E-05 |
| FERMT1 | -0.715857224 | 4.56E-05 | 0.000133682 |
| FERMT2 | 0.758742364 | 1.91E-18 | 1.95E-17 |
| FEZ1 | 0.570630759 | 0.000523927 | 0.001307606 |
| FGD3 | 0.28312179 | 0.004774755 | 0.010030675 |
| FGD6 | -0.386736535 | 0.003923403 | 0.008400581 |
| FGF10 | 0.493508779 | 1.28E-06 | 4.63E-06 |

| | | | |
|---|---|---|---|
| FGF12 | 1.371622173 | 0.000187588 | 0.000500726 |
| FGF14 | 1.13060619 | 0.026658992 | 0.047443341 |
| FGF16 | 0.826003105 | 0.014621995 | 0.027645313 |
| FGF18 | 3.114628927 | 4.74E-16 | 4.03E-15 |
| FGF19 | 2.606559638 | 3.07E-25 | 4.88E-24 |
| FGF7 | 1.873971604 | 1.29E-14 | 9.81E-14 |
| FGF8 | 4.427179081 | 1.21E-05 | 3.83E-05 |
| FGF9 | 1.601138845 | 0.00205777 | 0.004653004 |
| FGFR1 | 0.865439795 | 1.04E-21 | 1.29E-20 |
| FGFR1OP2 | 0.54463499 | 2.67E-08 | 1.15E-07 |
| FGFR2 | 0.573730427 | 9.39E-08 | 3.83E-07 |
| FGGY | 1.203026515 | 5.56E-20 | 6.24E-19 |
| FH | -0.393037174 | 1.37E-05 | 4.28E-05 |
| FHL1 | 0.312498206 | 0.005556821 | 0.0115082 |
| FHL2 | 0.917032367 | 2.21E-11 | 1.27E-10 |
| FHL3 | 0.59096761 | 1.44E-11 | 8.43E-11 |
| FHL5 | 1.125974613 | 2.80E-35 | 7.87E-34 |
| FHOD1 | -0.66719573 | 9.09E-12 | 5.45E-11 |
| FIBIN | 0.291907502 | 0.003242324 | 0.007072648 |
| FIG4 | -1.533885699 | 3.40E-12 | 2.12E-11 |
| FIGNL1 | -0.700341335 | 0.001857287 | 0.00423206 |
| FIGNL2 | 1.363004003 | 0.003897699 | 0.008352125 |
| FILIP1 | -0.517840777 | 0.000777972 | 0.001888249 |
| FJX1 | 0.986179961 | 0.000290154 | 0.000753635 |
| FKBP10 | -0.779143735 | 4.18E-15 | 3.30E-14 |
| FKBP11 | -0.736353638 | 2.26E-07 | 8.87E-07 |
| FKBP14 | 0.758988857 | 3.21E-18 | 3.23E-17 |
| FKBP15 | -0.36213605 | 0.005927592 | 0.012199852 |
| FKBP1A | 1.358116055 | 1.85E-64 | 1.98E-62 |
| FKBP3 | 1.144292197 | 4.85E-45 | 2.29E-43 |
| FKBP4 | -0.615034464 | 1.65E-11 | 9.65E-11 |
| FKBP5 | 0.550928477 | 1.07E-09 | 5.22E-09 |
| FKBP8 | -0.209633477 | 0.019611922 | 0.036033113 |
| FKTN | 0.745510239 | 1.45E-07 | 5.81E-07 |
| FLAD1 | -1.067984419 | 1.23E-12 | 7.96E-12 |
| FLI1 | -1.619926514 | 9.06E-14 | 6.45E-13 |
| FLII | -0.940962033 | 6.49E-20 | 7.25E-19 |
| FLOT2 | 0.305475502 | 0.001837873 | 0.004192744 |
| FLRT2 | -1.007371141 | 8.20E-15 | 6.33E-14 |
| FLRT3 | -1.084872461 | 1.02E-06 | 3.71E-06 |
| FLVCR2 | -1.116586481 | 7.32E-05 | 0.000208084 |
| FMNL2 | -0.609526093 | 0.00141546 | 0.003296032 |
| FN1 | -1.780669934 | 3.29E-74 | 5.27E-72 |
| FN3K | 2.308584364 | 0.001931987 | 0.004387502 |
| FNDC3A | 1.197322536 | 1.50E-28 | 2.90E-27 |
| FNDC4 | 0.965423816 | 1.55E-07 | 6.18E-07 |
| FNDC5 | 1.789116422 | 7.33E-30 | 1.53E-28 |
| FNIP2 | -0.629587798 | 2.62E-08 | 1.13E-07 |

| | | | |
|---|---|---|---|
| FOCAD | -0.94505194 | 1.47E-08 | 6.52E-08 |
| FOLH1 | -0.275022725 | 0.0039216 | 0.008399368 |
| FOPNL | -0.469224868 | 3.57E-05 | 0.000106213 |
| FOS | -1.022940546 | 3.45E-19 | 3.70E-18 |
| FOSL2 | 0.571601034 | 0.009189222 | 0.018125076 |
| FOXJ1 | -2.069580316 | 1.04E-37 | 3.44E-36 |
| FOXK2 | -0.410677389 | 0.000966059 | 0.002313775 |
| FOXL3 | 5.807522268 | 8.25E-07 | 3.04E-06 |
| FOXM1 | 0.683518864 | 0.017203977 | 0.032008826 |
| FOXP1 | 0.525781506 | 6.88E-05 | 0.000196318 |
| FOXP4 | 0.563912128 | 0.000926095 | 0.002222759 |
| FPGS | -1.464773022 | 3.12E-08 | 1.33E-07 |
| FPGT | -0.568968002 | 6.07E-05 | 0.000174904 |
| FRA10AC1 | 0.545774909 | 1.81E-08 | 7.92E-08 |
| FRMD3 | 1.466413005 | 0.013683365 | 0.02605552 |
| FRMD4A | 0.858165183 | 0.018323539 | 0.033894059 |
| FRMD4B | -0.812857469 | 1.88E-05 | 5.77E-05 |
| FRMD6 | -0.434759782 | 0.000591199 | 0.001462077 |
| FRS2 | 1.159431197 | 4.06E-18 | 4.02E-17 |
| FRS3 | 1.15489471 | 2.05E-11 | 1.18E-10 |
| FRZB | 0.85027445 | 2.10E-22 | 2.77E-21 |
| FSBP | -1.761118785 | 1.30E-26 | 2.24E-25 |
| FSCN1 | 0.635919241 | 6.91E-09 | 3.15E-08 |
| FSHR | 1.796131105 | 0.019550954 | 0.035949212 |
| FSIP1 | 5.41481585 | 5.59E-160 | 6.33E-157 |
| FST | -1.279980936 | 8.82E-32 | 2.05E-30 |
| FSTL1 | -0.68908811 | 3.14E-09 | 1.47E-08 |
| FSTL3 | 0.717093748 | 3.93E-09 | 1.83E-08 |
| FTH1 | 1.941393587 | 4.82E-153 | 4.67E-150 |
| FTL | 0.929303402 | 6.54E-18 | 6.40E-17 |
| FTSJ3 | -0.839902819 | 2.21E-10 | 1.16E-09 |
| FUBP1 | -0.678916212 | 3.85E-10 | 1.97E-09 |
| FUBP3 | 0.6737252 | 5.24E-11 | 2.90E-10 |
| FUCA2 | -0.343847156 | 0.000352957 | 0.000905851 |
| FUNDC1 | 0.736894587 | 2.83E-14 | 2.11E-13 |
| FUNDC2 | 0.594002524 | 0.000279268 | 0.000727725 |
| FURIN | 0.424992721 | 0.010654392 | 0.02075301 |
| FUS | -0.372428089 | 4.64E-05 | 0.000135921 |
| FUT8 | 0.654599485 | 7.68E-08 | 3.16E-07 |
| FXN | 0.924041211 | 3.61E-07 | 1.38E-06 |
| FXR1 | 0.211796678 | 0.022038057 | 0.040013528 |
| FXYD2 | 2.324479751 | 0.000246801 | 0.000648593 |
| FXYD6 | 0.372054677 | 3.32E-05 | 9.91E-05 |
| FYB | 5.471716348 | 7.11E-08 | 2.94E-07 |
| FYN | 0.401115455 | 9.56E-07 | 3.49E-06 |
| FYTTD1 | 0.480687645 | 5.08E-09 | 2.34E-08 |
| FZD1 | 0.44059663 | 3.37E-06 | 1.14E-05 |
| FZD2 | -0.352805633 | 0.015259291 | 0.028738297 |

| | | | |
|---|---|---|---|
| FZD7 | -0.73732365 | 2.02E-06 | 7.11E-06 |
| FZD9 | 1.369870755 | 2.23E-25 | 3.57E-24 |
| G0S2 | 0.510742362 | 0.000433311 | 0.001096753 |
| G2E3 | -0.971945055 | 3.02E-11 | 1.72E-10 |
| G3BP1 | -0.519459264 | 4.78E-09 | 2.21E-08 |
| G3BP2 | -0.368770863 | 8.42E-06 | 2.72E-05 |
| GAB1 | -1.014337885 | 5.86E-09 | 2.69E-08 |
| GABARAPL1 | -0.360305765 | 0.020314268 | 0.037151713 |
| GABARAPL2 | 0.609788808 | 1.23E-09 | 6.00E-09 |
| GABRA4 | 0.963785938 | 2.99E-05 | 8.95E-05 |
| GABRG1 | 0.491429939 | 0.007534349 | 0.015162403 |
| GABRG3 | 1.807337402 | 0.000526053 | 0.001312429 |
| GABRR1 | 4.757148104 | 0.004094817 | 0.008745562 |
| GABRR2 | 2.570864787 | 1.03E-06 | 3.73E-06 |
| GADD45A | -0.850354528 | 9.73E-15 | 7.46E-14 |
| GADD45G | -0.710883352 | 3.63E-05 | 0.00010777 |
| GAL3ST1 | 4.124247588 | 0.000609301 | 0.001504107 |
| GALC | -0.694467026 | 3.47E-16 | 2.98E-15 |
| GALK1 | -0.408262758 | 0.012445024 | 0.023945748 |
| GALNS | -0.758421854 | 1.02E-14 | 7.79E-14 |
| GALNT1 | 0.561483277 | 1.43E-11 | 8.40E-11 |
| GALNT11 | 0.983706578 | 1.76E-27 | 3.22E-26 |
| GALNT12 | -0.925904069 | 1.03E-05 | 3.30E-05 |
| GALNT15 | -0.953790755 | 6.51E-05 | 0.000186671 |
| GALNT16 | -0.729284244 | 1.61E-11 | 9.40E-11 |
| GALNT17 | 1.079215681 | 6.09E-42 | 2.45E-40 |
| GALNT6 | -1.632972385 | 0.001942167 | 0.004407002 |
| GALNT7 | -0.340766489 | 0.022419939 | 0.040625491 |
| GAPDH | 0.669844572 | 7.27E-12 | 4.39E-11 |
| GAPVD1 | -0.640414169 | 1.55E-08 | 6.84E-08 |
| GAR1 | 1.303225088 | 8.10E-07 | 2.99E-06 |
| GARNL3 | 0.704012581 | 7.73E-13 | 5.09E-12 |
| GARS | -0.842065813 | 4.98E-23 | 6.74E-22 |
| GAS2L3 | -1.325170922 | 0.014019637 | 0.026621249 |
| GAS6 | -0.382264586 | 0.000469449 | 0.001179218 |
| GAS7 | -0.401477663 | 0.025396163 | 0.045401111 |
| GAS8 | -0.602383706 | 1.36E-05 | 4.25E-05 |
| GATA2 | 0.514512908 | 0.022714977 | 0.041088896 |
| GATA6 | 1.15306218 | 3.16E-26 | 5.32E-25 |
| GATAD1 | 0.293730234 | 0.005819349 | 0.011993418 |
| GATAD2A | -0.704215089 | 1.11E-10 | 5.97E-10 |
| GATB | -1.177800829 | 4.12E-06 | 1.38E-05 |
| GATD3AL1 | 0.432806661 | 0.000125163 | 0.000342912 |
| GATSL2 | 1.527779429 | 1.05E-69 | 1.35E-67 |
| GBA2 | -1.079824538 | 4.82E-07 | 1.83E-06 |
| GBF1 | -0.449924696 | 0.000320441 | 0.000827399 |
| GCC1 | -1.30600032 | 2.36E-22 | 3.09E-21 |
| GCC2 | -0.575701545 | 1.08E-05 | 3.43E-05 |

| | | | |
|---|---|---|---|
| GCDH | -1.046770692 | 4.84E-24 | 6.98E-23 |
| GCHFR | 1.702230778 | 1.36E-65 | 1.55E-63 |
| GCK | 0.991044716 | 1.39E-09 | 6.76E-09 |
| GCLC | 0.554952124 | 4.33E-12 | 2.67E-11 |
| GCNT1 | -4.448193022 | 0.003724352 | 0.008018595 |
| GCSH | 0.329870545 | 0.013712733 | 0.026100471 |
| GDA | 0.55121843 | 0.008439848 | 0.016798127 |
| GDAP1L1 | 2.360043668 | 8.50E-15 | 6.56E-14 |
| GDE1 | 0.44367728 | 0.008387918 | 0.016709451 |
| GDF9 | -2.863215405 | 7.71E-08 | 3.18E-07 |
| GDPD4 | 4.882151194 | 1.69E-07 | 6.74E-07 |
| GEM | -1.076286807 | 3.70E-29 | 7.43E-28 |
| GEMIN5 | -0.978330405 | 1.39E-16 | 1.23E-15 |
| GEMIN6 | -0.266998298 | 0.015316022 | 0.028829161 |
| GEN1 | 0.409740574 | 0.011729943 | 0.022643571 |
| GFM1 | -1.17440457 | 3.68E-24 | 5.37E-23 |
| GFOD1 | 0.247004942 | 0.019358761 | 0.035624733 |
| GFOD2 | 1.313459567 | 6.61E-44 | 2.96E-42 |
| GFPT1 | -0.921955972 | 2.18E-16 | 1.91E-15 |
| GFPT2 | -1.505825868 | 1.76E-35 | 5.03E-34 |
| GGACT | 0.821238026 | 2.03E-09 | 9.70E-09 |
| GGNBP2 | -0.98374092 | 3.59E-19 | 3.84E-18 |
| GGPS1 | -0.512446417 | 3.65E-05 | 0.000108321 |
| GHDC | -1.479641085 | 4.34E-13 | 2.94E-12 |
| GHITM | -0.602418744 | 1.03E-09 | 5.07E-09 |
| GHR | 1.606281552 | 5.59E-33 | 1.37E-31 |
| GHRL | 3.140338424 | 4.09E-16 | 3.50E-15 |
| GID4 | 1.006119497 | 9.11E-16 | 7.56E-15 |
| GID8 | 0.692914056 | 5.00E-17 | 4.58E-16 |
| GIGYF2 | -0.631598061 | 0.000255228 | 0.000668927 |
| GIN1 | -1.210548679 | 5.30E-06 | 1.76E-05 |
| GINM1 | 0.481486851 | 2.73E-06 | 9.45E-06 |
| GINS3 | -0.651704474 | 5.75E-08 | 2.39E-07 |
| GIPC2 | 0.596836666 | 4.89E-05 | 0.000142867 |
| GIT2 | 0.443813761 | 3.08E-05 | 9.20E-05 |
| GJA1 | -0.436893959 | 3.68E-06 | 1.24E-05 |
| GJA9 | 3.857605353 | 0.004835991 | 0.010140511 |
| GJB1 | 0.837674655 | 0.02585288 | 0.046132644 |
| GJC2 | 0.767797486 | 2.15E-13 | 1.50E-12 |
| GJC3 | 2.146097571 | 1.20E-117 | 7.09E-115 |
| GJD4 | 2.896771743 | 0.000808054 | 0.001954978 |
| GK5 | 0.754699675 | 1.86E-13 | 1.30E-12 |
| GKAP1 | -0.550718019 | 0.005563332 | 0.011519928 |
| GLA | -1.018224746 | 3.30E-13 | 2.26E-12 |
| GLB1 | -1.155515909 | 1.62E-36 | 5.01E-35 |
| GLB1L | -0.395395564 | 0.000203487 | 0.000540405 |
| GLCCI1 | -0.821311289 | 3.90E-05 | 0.000115225 |
| GLCE | -0.978152787 | 1.06E-05 | 3.38E-05 |

| | | | |
|---|---|---|---|
| GLDC | -0.656011197 | 0.001260623 | 0.002961866 |
| GLE1 | -0.439229439 | 0.000112981 | 0.000311426 |
| GLG1 | -1.69933388 | 7.80E-85 | 1.86E-82 |
| GLI2 | 0.443978725 | 0.0045405 | 0.009575559 |
| GLI3 | 0.413307465 | 0.004873442 | 0.010214314 |
| GLIPR1L | -0.313978956 | 0.000698473 | 0.001708411 |
| GLIPR2 | 0.749438473 | 2.37E-20 | 2.72E-19 |
| GLIS1 | -1.457101471 | 1.27E-15 | 1.04E-14 |
| GLIS3 | -1.675540193 | 6.16E-15 | 4.80E-14 |
| GLMN | -1.142514506 | 6.83E-17 | 6.19E-16 |
| GLO1 | 0.894622076 | 2.80E-20 | 3.21E-19 |
| GLOD4 | -0.512016711 | 2.73E-08 | 1.17E-07 |
| GLP1R | 1.622963684 | 0.000113956 | 0.000313752 |
| GLP2R | 1.536010565 | 0.000829625 | 0.002003956 |
| GLRA1 | 4.645974162 | 0.005148116 | 0.010733731 |
| GLRX | 0.669609997 | 2.99E-06 | 1.03E-05 |
| GLRX5 | 0.427366311 | 1.27E-05 | 3.99E-05 |
| GLS2 | -0.394455518 | 0.000347756 | 0.000893331 |
| GLT8D1 | -0.505900321 | 2.07E-07 | 8.15E-07 |
| GLT8D2 | 0.970466868 | 4.44E-11 | 2.48E-10 |
| GLUL | -1.384220094 | 0.003455975 | 0.007493 |
| GLYR1 | -0.213737627 | 0.023666389 | 0.042662275 |
| GMDS | 1.253612826 | 7.30E-50 | 4.39E-48 |
| GMEB2 | -0.517535479 | 0.001656356 | 0.003810628 |
| GMIP | 1.605492849 | 0.000723837 | 0.001765685 |
| GMNN | -1.332809521 | 2.46E-13 | 1.69E-12 |
| GMPS | -0.717528395 | 5.70E-11 | 3.15E-10 |
| GNA11 | 0.811606065 | 0.002052144 | 0.004641826 |
| GNAI1 | -0.69389524 | 1.00E-11 | 5.97E-11 |
| GNAI2 | 0.458375384 | 8.66E-06 | 2.79E-05 |
| GNAI3 | -0.354170018 | 1.58E-05 | 4.90E-05 |
| GNAS | 0.860777186 | 3.63E-17 | 3.38E-16 |
| GNB1L | -0.367688967 | 0.013056128 | 0.024997731 |
| GNB2L1 | 0.185972982 | 0.016116302 | 0.03017256 |
| GNB4 | -0.561101476 | 0.000326736 | 0.000842695 |
| GNG10 | 1.313511557 | 3.97E-32 | 9.43E-31 |
| GNG12 | 1.335584423 | 1.28E-61 | 1.19E-59 |
| GNG2 | 1.125724847 | 3.04E-36 | 9.15E-35 |
| GNG5 | 0.431103111 | 5.33E-05 | 0.00015498 |
| GNL2 | -0.541511332 | 4.73E-09 | 2.19E-08 |
| GNLY | 2.025947271 | 7.41E-97 | 2.46E-94 |
| GNPDA1 | 0.65764562 | 3.16E-09 | 1.48E-08 |
| GNPDA2 | 0.283850994 | 0.025516485 | 0.045586233 |
| GNPNAT1 | 0.773336244 | 6.09E-22 | 7.75E-21 |
| GnRH-I | 2.465535692 | 0.021685384 | 0.039420588 |
| GNS | -0.732663968 | 1.60E-20 | 1.84E-19 |
| GOLGA4 | -0.416648421 | 0.001769376 | 0.004048713 |
| GOLGA5 | -0.96923677 | 1.61E-25 | 2.59E-24 |

| | | | |
|---|---|---|---|
| GOLGA7 | 0.765792723 | 3.64E-21 | 4.39E-20 |
| GOLGB1 | -0.556098373 | 5.24E-06 | 1.74E-05 |
| GOLIM4 | 0.466129326 | 7.72E-07 | 2.85E-06 |
| GOLT1B | 0.6557533 | 3.61E-11 | 2.03E-10 |
| GORAB | 0.268288969 | 0.001747539 | 0.00400144 |
| GOSR1 | -0.781885165 | 4.17E-14 | 3.06E-13 |
| GOSR2 | 0.581295096 | 2.85E-11 | 1.63E-10 |
| GOT1 | -0.874986699 | 6.47E-24 | 9.30E-23 |
| GOT2 | -0.594011489 | 2.61E-08 | 1.12E-07 |
| GP1BB | 0.997988041 | 0.000286443 | 0.000744565 |
| GPAT3 | -0.628927639 | 6.46E-13 | 4.29E-12 |
| GPATCH11 | 0.535931512 | 5.85E-06 | 1.93E-05 |
| GPATCH2 | -0.843022672 | 2.13E-06 | 7.48E-06 |
| GPATCH3 | -0.47781543 | 3.61E-05 | 0.000107384 |
| GPATCH4 | -0.350670566 | 0.015653019 | 0.029398347 |
| GPC1 | -0.461948267 | 4.85E-05 | 0.000141714 |
| GPC3 | 1.124782103 | 0.013771462 | 0.026193913 |
| GPC4 | -0.630179563 | 1.80E-09 | 8.65E-09 |
| GPCPD1 | -0.690542697 | 3.91E-06 | 1.32E-05 |
| GPD1L | 0.430439933 | 0.008445229 | 0.016806377 |
| GPD2 | 0.339830017 | 0.003178435 | 0.006950321 |
| GPHN | -0.365750324 | 0.026325491 | 0.046902094 |
| GPI | -0.84580457 | 4.37E-12 | 2.69E-11 |
| GPN2 | 0.737453871 | 1.43E-12 | 9.21E-12 |
| GPN3 | 0.355877465 | 0.000873064 | 0.002101414 |
| GPR1 | 0.713411069 | 1.26E-11 | 7.46E-11 |
| GPR107 | -0.424000476 | 3.71E-06 | 1.25E-05 |
| GPR119 | 1.789299395 | 0.002554994 | 0.005689298 |
| GPR135 | -2.545242676 | 0.019732846 | 0.036249337 |
| GPR137B | 0.669865572 | 1.20E-07 | 4.84E-07 |
| GPR137C | -1.080855321 | 7.03E-10 | 3.52E-09 |
| GPR143 | 1.28729607 | 0.000469835 | 0.001179753 |
| GPR146 | -1.086033463 | 5.15E-10 | 2.60E-09 |
| GPR149 | 1.323968015 | 0.019575685 | 0.035984949 |
| GPR153 | 0.982572348 | 0.010079898 | 0.019715926 |
| GPR157 | 1.171537503 | 6.60E-10 | 3.31E-09 |
| GPR162 | 1.782005135 | 1.34E-58 | 1.09E-56 |
| GPR171 | 2.041608745 | 3.08E-21 | 3.72E-20 |
| GPR174 | 2.412940991 | 0.024561035 | 0.044140366 |
| GPR180 | 0.431472496 | 4.45E-05 | 0.000130692 |
| GPR27 | 1.178638995 | 3.40E-16 | 2.93E-15 |
| GPR34 | 2.727661828 | 1.79E-07 | 7.09E-07 |
| GPR37 | -0.587986779 | 0.008433011 | 0.016789436 |
| GPR39 | 2.303953127 | 0.000126962 | 0.000347515 |
| GPR68 | -1.101553118 | 5.52E-26 | 9.19E-25 |
| GPR75 | 0.663206254 | 0.003296021 | 0.00717941 |
| GPR82 | 2.195652412 | 0.004726444 | 0.009939938 |
| GPR89A | -0.848297591 | 3.31E-09 | 1.55E-08 |

| | | | |
|---|---|---|---|
| GPRC5B | 1.56321173 | 0.000998493 | 0.002388091 |
| GPRC5C | 1.146013824 | 0.000369955 | 0.000946616 |
| GPRC6A | 7.811247601 | 3.96E-10 | 2.02E-09 |
| GPRIN3 | -3.319634123 | 0.027619383 | 0.048963434 |
| GPS1 | -0.587462907 | 2.06E-08 | 8.96E-08 |
| GPSM1 | 1.067522071 | 9.52E-06 | 3.05E-05 |
| GPT2 | -2.099077072 | 4.47E-32 | 1.05E-30 |
| GPX1 | 0.923840941 | 2.47E-24 | 3.66E-23 |
| GPX3 | 1.214479482 | 1.09E-43 | 4.83E-42 |
| GPX4 | 0.630651643 | 3.98E-11 | 2.23E-10 |
| GPX7 | 0.595592619 | 7.44E-05 | 0.00021116 |
| GRAMD1C | -1.01039508 | 1.45E-07 | 5.81E-07 |
| GRAMD2B | 0.447718506 | 0.000119 | 0.000326954 |
| GRAMD4 | 0.468802452 | 0.001433758 | 0.003335781 |
| GRB10 | -0.312294207 | 0.006728931 | 0.013693611 |
| GRB2 | 0.588442588 | 1.31E-05 | 4.12E-05 |
| GREM1 | 5.162368693 | 1.40E-10 | 7.44E-10 |
| GRHL3 | -1.229405436 | 4.12E-05 | 0.000121381 |
| GRHPR | -0.625134304 | 0.001143526 | 0.002708278 |
| GRIA2 | -1.204664974 | 0.000228421 | 0.000602854 |
| GRIA4 | 2.696981518 | 0.004483222 | 0.009470927 |
| GRID1 | -0.796838056 | 0.000157747 | 0.000425924 |
| GRIK1 | 0.913105306 | 0.015664913 | 0.02941662 |
| GRIN2A | -1.516608164 | 0.019073767 | 0.035143318 |
| GRIP2 | 0.768810365 | 0.008847679 | 0.017505429 |
| GRK3 | 2.492019953 | 1.54E-34 | 4.14E-33 |
| GRK4 | -0.481493907 | 0.002367171 | 0.00530059 |
| GRK5 | -1.502572657 | 0.008996512 | 0.017786268 |
| GRK7 | 2.178939898 | 2.83E-08 | 1.22E-07 |
| GRM5 | 1.85048629 | 0.007709664 | 0.015494567 |
| GRN | -2.054328269 | 3.80E-75 | 6.79E-73 |
| GRP | -2.413660554 | 0.000451185 | 0.00113839 |
| GRPEL1 | -0.318479998 | 0.000861778 | 0.002076087 |
| GRSF1 | -0.95587208 | 2.59E-18 | 2.62E-17 |
| GSK3B | 0.583025946 | 2.40E-10 | 1.25E-09 |
| GSN | -1.096223561 | 1.46E-22 | 1.93E-21 |
| GSPT2 | -0.693001924 | 2.35E-16 | 2.06E-15 |
| GSR | 0.796567798 | 3.41E-24 | 5.00E-23 |
| GSS | -1.252980728 | 9.52E-23 | 1.27E-21 |
| GSTA2 | 0.676928561 | 8.15E-06 | 2.63E-05 |
| GSTCD | -1.317076923 | 2.29E-09 | 1.09E-08 |
| GSTM2 | 0.34871621 | 0.003002159 | 0.00659427 |
| GSTO2 | -0.516524953 | 0.00012836 | 0.000350828 |
| GSTT1 | 0.401602986 | 1.56E-05 | 4.84E-05 |
| GSTZ1 | 0.709117861 | 4.83E-05 | 0.000141242 |
| GTF2B | -2.119878074 | 1.31E-85 | 3.23E-83 |
| GTF2F2 | -0.256359639 | 0.024417172 | 0.043899233 |
| GTF2H1 | 0.525982052 | 3.36E-05 | 0.000100085 |

| | | | |
|---|---|---|---|
| GTF2H5 | 0.425239296 | 3.73E-05 | 0.000110762 |
| GTF2IRD1 | 0.656974278 | 6.41E-08 | 2.66E-07 |
| GTF3A | 0.989063138 | 6.23E-27 | 1.09E-25 |
| GTF3C1 | -0.897524029 | 2.17E-06 | 7.59E-06 |
| GTF3C3 | -0.826446438 | 7.48E-07 | 2.77E-06 |
| GTF3C4 | -1.595317666 | 1.02E-14 | 7.79E-14 |
| GTF3C5 | -1.174210497 | 5.34E-14 | 3.89E-13 |
| GTF3C6 | 1.13089693 | 1.11E-20 | 1.29E-19 |
| GTPBP1 | -0.32029927 | 0.001125843 | 0.002670586 |
| GTPBP2 | -1.299973763 | 6.02E-07 | 2.25E-06 |
| GTPBP4 | -0.3409864 | 0.000504453 | 0.001261782 |
| GTPBP8 | -0.778282482 | 1.11E-13 | 7.86E-13 |
| GTSE1 | -1.260386706 | 0.000600471 | 0.001483119 |
| GUCY1B1 | -0.493795378 | 0.001109936 | 0.002635614 |
| GUF1 | -0.532960516 | 9.22E-05 | 0.000257914 |
| GUK1 | 0.560880016 | 4.62E-05 | 0.000135558 |
| GULP1 | 0.635554316 | 2.40E-16 | 2.10E-15 |
| GUSB | -1.455432079 | 3.03E-40 | 1.09E-38 |
| GXYLT2 | 1.727306786 | 6.25E-72 | 8.76E-70 |
| GYPC | 1.027153948 | 0.000125645 | 0.00034403 |
| GYS2 | 4.746873117 | 4.00E-27 | 7.12E-26 |
| GZF1 | -0.605351838 | 8.97E-07 | 3.29E-06 |
| H2AFY2 | -0.837137973 | 0.000146553 | 0.000397198 |
| H2AFZ | -0.632520064 | 8.57E-07 | 3.15E-06 |
| H2A-VIII | -1.070609391 | 6.38E-05 | 0.000183045 |
| H2A-VIII.1 | -0.681238573 | 3.72E-07 | 1.42E-06 |
| H2A-VIII.2 | -0.342750001 | 0.01752873 | 0.032543437 |
| H2AZ2 | 0.438778978 | 6.62E-05 | 0.000189335 |
| H3F3B | -1.347388049 | 2.70E-35 | 7.62E-34 |
| H3F3C | 0.90986482 | 2.78E-24 | 4.09E-23 |
| HABP2 | 1.812504364 | 0.0038611 | 0.008277615 |
| HACD1 | 1.418469487 | 3.45E-38 | 1.16E-36 |
| HACL1 | -0.930971532 | 1.56E-10 | 8.30E-10 |
| HADH | -0.351658778 | 0.001072716 | 0.002553477 |
| HADHA | -0.991455689 | 1.30E-26 | 2.24E-25 |
| HADHB | -0.695110577 | 5.13E-16 | 4.36E-15 |
| HAGH | 1.151105782 | 4.26E-30 | 9.03E-29 |
| HAGHL | -0.760791696 | 2.72E-10 | 1.41E-09 |
| HAND2 | 0.371539188 | 0.021157815 | 0.03856988 |
| HARBI1 | -1.105479791 | 3.52E-09 | 1.65E-08 |
| HARS | -0.712428678 | 2.04E-16 | 1.79E-15 |
| HAT1 | -0.682880138 | 3.28E-10 | 1.69E-09 |
| HAUS1 | 0.475998664 | 0.006000255 | 0.012330733 |
| HAUS3 | -1.154080938 | 1.08E-05 | 3.44E-05 |
| HAUS8 | -0.591348463 | 5.12E-05 | 0.000149113 |
| HAVCR1 | 1.452937044 | 0.000355422 | 0.000911662 |
| HAX1 | -0.674765492 | 6.20E-07 | 2.32E-06 |
| HBP1 | -1.069807746 | 3.74E-33 | 9.31E-32 |

| | | | |
|---|---|---|---|
| HCCS | 0.31537395 | 0.000713417 | 0.001742142 |
| HCFC2 | -0.581901616 | 4.54E-05 | 0.000133231 |
| HCN1 | 1.52705568 | 1.03E-08 | 4.63E-08 |
| HDAC10 | -0.898532394 | 0.002152667 | 0.004853052 |
| HDAC11 | 0.346810092 | 0.025424378 | 0.045445574 |
| HDAC3 | -0.41551797 | 0.000259021 | 0.000678084 |
| HDAC7 | -0.48824583 | 0.000101497 | 0.000282059 |
| HDC | 2.917917324 | 0.00508859 | 0.010621025 |
| HDDC2 | 0.398958328 | 0.001006605 | 0.002406222 |
| HDGF | 1.333993309 | 1.11E-42 | 4.66E-41 |
| HDGFL3 | 0.924455216 | 3.05E-10 | 1.57E-09 |
| HDHD2 | -0.469066698 | 0.000119251 | 0.000327579 |
| HDLBP | 0.905367574 | 4.23E-28 | 7.98E-27 |
| HDX | -1.009289242 | 1.93E-13 | 1.35E-12 |
| HEATR5B | -0.714895532 | 0.0001236 | 0.000338842 |
| HECTD1 | -1.02843389 | 6.80E-22 | 8.59E-21 |
| HELLS | -0.757717882 | 3.65E-06 | 1.24E-05 |
| HELQ | -0.559540526 | 0.003215167 | 0.007019042 |
| HELZ2 | 0.840472643 | 0.010765013 | 0.020950466 |
| HEMK1 | -1.422617258 | 8.88E-13 | 5.81E-12 |
| HENMT1 | 0.503794703 | 0.003994631 | 0.008549052 |
| HEPH | -0.859372347 | 2.10E-06 | 7.38E-06 |
| HEPHL1 | 1.58183479 | 0.015727918 | 0.029522699 |
| HERC1 | 0.456567174 | 0.000878904 | 0.002113972 |
| HERC2 | -0.625758391 | 1.39E-05 | 4.34E-05 |
| HERC3 | -0.574960554 | 6.19E-07 | 2.31E-06 |
| HERPUD1 | -1.827321231 | 4.00E-14 | 2.94E-13 |
| HERPUD2 | -0.317283522 | 0.000214486 | 0.000568063 |
| HES1 | -1.162063781 | 0.000391375 | 0.000997472 |
| HES4 | 0.749036325 | 1.37E-10 | 7.32E-10 |
| HES6 | 0.973271356 | 3.70E-06 | 1.25E-05 |
| HESX1 | -1.737085429 | 6.94E-07 | 2.58E-06 |
| HEXA | -0.503479225 | 1.04E-08 | 4.66E-08 |
| HEXB | -0.886524464 | 1.21E-19 | 1.34E-18 |
| HEXD | -0.994130683 | 0.000218505 | 0.000577918 |
| HEXIM1 | 0.54260319 | 0.001053774 | 0.002511466 |
| HEY2 | 0.542353845 | 5.54E-06 | 1.83E-05 |
| HGFAC | 1.128571613 | 0.004155131 | 0.008856288 |
| HGS | -0.569192292 | 3.28E-10 | 1.69E-09 |
| HGSNAT | -0.329843276 | 0.007513392 | 0.015122468 |
| HHAT | 0.744671418 | 0.000533846 | 0.001329675 |
| HHEX | -0.677538066 | 6.24E-08 | 2.59E-07 |
| HHLA2 | 0.519220386 | 3.44E-06 | 1.17E-05 |
| HIBCH | 0.738160649 | 9.09E-15 | 6.99E-14 |
| HIC1 | 0.835515783 | 3.21E-20 | 3.66E-19 |
| HIC2 | 0.463515629 | 9.23E-05 | 0.000258296 |
| HIF1A | -0.589163063 | 2.30E-11 | 1.32E-10 |
| HIGD1C | -0.249797213 | 0.009885098 | 0.019385117 |

| | | | |
|---|---|---|---|
| HIKESHI | 0.85726013 | 6.56E-17 | 5.96E-16 |
| HINFP | -0.740410753 | 1.70E-05 | 5.23E-05 |
| HINT3 | 0.744796941 | 9.40E-13 | 6.14E-12 |
| HINTW | 1.177344491 | 1.73E-34 | 4.63E-33 |
| HIP1 | -0.372083741 | 0.000460586 | 0.001159741 |
| HIP1R | -0.686517854 | 1.80E-05 | 5.53E-05 |
| HIPK1 | 0.348248152 | 0.017203638 | 0.032008826 |
| HIPK2 | 0.808369739 | 1.85E-14 | 1.40E-13 |
| HIST1H3H | -0.87350967 | 0.000413713 | 0.001050082 |
| HIST1H46 | 0.39676077 | 0.001133975 | 0.002687532 |
| HIVEP1 | -0.473636434 | 0.004322928 | 0.009159352 |
| HIVEP3 | 1.577889639 | 2.65E-13 | 1.82E-12 |
| HJURP | -1.057640677 | 0.001620391 | 0.003733573 |
| HK1 | -0.945376022 | 5.39E-25 | 8.41E-24 |
| HK2 | -1.593056286 | 3.12E-11 | 1.77E-10 |
| HKDC1 | 3.235892727 | 0.000324447 | 0.000836949 |
| HLCS | 0.356293264 | 0.003774418 | 0.008112254 |
| HLX | -1.436256485 | 0.008172497 | 0.016320979 |
| HM13 | 0.502114836 | 2.35E-10 | 1.23E-09 |
| HMBOX1 | 0.537109302 | 0.000298919 | 0.000775364 |
| HMBS | -0.255251049 | 0.025309662 | 0.045288169 |
| HMG20A | -0.403030207 | 5.90E-05 | 0.000170387 |
| HMG20B | -0.743567375 | 6.23E-14 | 4.51E-13 |
| HMGA1 | 1.239463165 | 0.003246226 | 0.007080023 |
| HMGA2 | 1.670354612 | 4.30E-23 | 5.83E-22 |
| HMGB1 | 0.361048424 | 0.001263407 | 0.003161588 |
| HMGB2 | -0.987647626 | 3.14E-06 | 1.08E-05 |
| HMGCL | -0.789483676 | 1.05E-11 | 6.22E-11 |
| HMGCR | -0.958195492 | 6.69E-31 | 1.48E-29 |
| HMGCS1 | -1.280313596 | 5.02E-51 | 3.11E-49 |
| HMGN1 | 0.440821484 | 9.50E-06 | 3.04E-05 |
| HMGN2P46 | 1.14316153 | 2.35E-23 | 3.26E-22 |
| HMGN4 | 0.759712134 | 1.05E-15 | 8.67E-15 |
| HMOX1 | 1.468693563 | 5.03E-46 | 2.51E-44 |
| HMOX2 | -1.097504697 | 7.93E-33 | 1.92E-31 |
| HN1 | 0.287159788 | 0.004824976 | 0.010120535 |
| HN1L | 0.728078184 | 4.04E-11 | 2.27E-10 |
| HNMT | 0.923103305 | 3.26E-06 | 1.11E-05 |
| HNRNPA0 | -1.059929434 | 0.022592021 | 0.040893694 |
| HNRNPA2B1 | 0.488916677 | 7.93E-10 | 3.94E-09 |
| HNRNPA3 | 0.355212746 | 0.00044945 | 0.001134432 |
| HNRNPD | 0.722537256 | 1.55E-06 | 5.52E-06 |
| HNRNPDL | 0.231424027 | 0.017335804 | 0.032224871 |
| HNRNPH2 | -0.601974721 | 5.80E-15 | 4.53E-14 |
| HNRNPH3 | 0.341624328 | 1.20E-05 | 3.79E-05 |
| HNRNPKL | 0.231289305 | 0.014416223 | 0.027282832 |
| HNRNPM | -0.397928252 | 1.01E-05 | 3.22E-05 |
| HNRNPU | -0.24828644 | 0.003652883 | 0.007882202 |

| | | | |
|---|---|---|---|
| HOGA1 | 2.307009274 | 0.004306859 | 0.009129574 |
| HOMER2 | -0.800900697 | 3.82E-06 | 1.29E-05 |
| HOOK3 | -0.298737003 | 0.020591988 | 0.03761407 |
| HOPX | -1.125335648 | 1.12E-28 | 2.18E-27 |
| HOXA10 | 0.854036344 | 7.62E-06 | 2.47E-05 |
| HOXA3 | 0.67477237 | 0.006392296 | 0.013071173 |
| HOXA7 | 1.706209654 | 5.41E-62 | 5.14E-60 |
| HOXA9 | 1.188366669 | 1.53E-08 | 6.77E-08 |
| HOXD8 | 1.225369934 | 6.47E-07 | 2.41E-06 |
| HP1BP3 | -0.612077406 | 7.98E-07 | 2.95E-06 |
| HPCAL1 | 0.498071265 | 1.79E-08 | 7.86E-08 |
| HPD | 0.722396413 | 0.002733107 | 0.006046287 |
| HPDL | -1.27931675 | 1.67E-07 | 6.64E-07 |
| HPF1 | 0.705361778 | 4.07E-14 | 2.99E-13 |
| HPGDS | -0.494280317 | 5.60E-06 | 1.85E-05 |
| HPRT1 | 1.327614391 | 1.19E-59 | 1.02E-57 |
| HPS5 | -1.435804137 | 5.64E-26 | 9.39E-25 |
| H-RAS | 1.46169995 | 1.08E-78 | 2.03E-76 |
| HS1BP3 | 0.496222281 | 3.84E-05 | 0.000113546 |
| HS2ST1 | 0.800781375 | 1.73E-12 | 1.11E-11 |
| HS3ST1 | -2.014316795 | 8.77E-08 | 3.59E-07 |
| HS3ST3A1 | 1.116246483 | 0.004917768 | 0.010300863 |
| HS6ST1 | 0.66792303 | 5.27E-12 | 3.22E-11 |
| HSBP1 | 0.642750687 | 5.71E-09 | 2.62E-08 |
| HSBP1L1 | 0.877162063 | 2.83E-09 | 1.33E-08 |
| HSCB | 0.903269584 | 3.87E-16 | 3.32E-15 |
| HSD11B1L | -1.11203825 | 0.008680377 | 0.017203803 |
| HSD17B10 | 0.293219579 | 0.0026153 | 0.00581406 |
| HSD17B12 | -0.584594849 | 9.97E-12 | 5.95E-11 |
| HSD17B4 | -0.641029132 | 9.56E-10 | 4.71E-09 |
| HSD17B7 | -0.863551508 | 4.68E-21 | 5.58E-20 |
| HSD3B1 | 4.936915775 | 0.002203498 | 0.004955318 |
| HSDL1 | -0.538122238 | 9.18E-06 | 2.94E-05 |
| HSDL2 | -0.396022976 | 0.001050334 | 0.002503707 |
| HSF1 | 1.589566936 | 2.81E-09 | 1.33E-08 |
| HSF2 | -0.786671492 | 1.73E-18 | 1.77E-17 |
| HSF3 | -0.944422701 | 5.80E-19 | 6.14E-18 |
| HSP90AA1 | -0.356365985 | 1.52E-06 | 5.42E-06 |
| HSP90AB1 | -0.980398633 | 1.52E-21 | 1.88E-20 |
| HSP90B1 | -1.140885358 | 1.23E-27 | 2.26E-26 |
| HSPA12A | -0.672487887 | 2.25E-06 | 7.86E-06 |
| HSPA13 | -0.681929552 | 5.40E-09 | 2.48E-08 |
| HSPA14 | -0.492432496 | 5.33E-06 | 1.76E-05 |
| HSPA2 | -1.034264547 | 2.90E-38 | 9.77E-37 |
| HSPA4 | -1.709910497 | 5.75E-82 | 1.24E-79 |
| HSPA4L | -1.201367258 | 8.96E-39 | 3.07E-37 |
| HSPA5 | -1.012678412 | 9.50E-20 | 1.06E-18 |
| HSPA8 | -1.086549475 | 3.33E-48 | 1.85E-46 |

| | | | |
|---|---|---|---|
| HSPA9 | -0.710886774 | 5.29E-20 | 5.96E-19 |
| HSPB1 | 0.808636314 | 1.11E-21 | 1.38E-20 |
| HSPB11 | 0.631067317 | 1.46E-05 | 4.54E-05 |
| HSPB3 | 5.048965578 | 0.001301803 | 0.003048495 |
| HSPB7 | 0.970012219 | 5.35E-19 | 5.67E-18 |
| HSPB9 | -0.418319168 | 7.93E-06 | 2.57E-05 |
| HSPD1 | -0.551793762 | 3.06E-13 | 2.10E-12 |
| HSPG2 | -0.383671372 | 0.00035596 | 0.000912848 |
| HSPH1 | -0.708623766 | 3.10E-14 | 2.30E-13 |
| HTATSF1 | -0.691264282 | 4.64E-07 | 1.76E-06 |
| HTR2A | 1.012605413 | 3.41E-29 | 6.87E-28 |
| HTR2B | 0.949358154 | 2.27E-11 | 1.30E-10 |
| HTR4 | 1.224875977 | 1.28E-06 | 4.59E-06 |
| HTR7L | 5.87436372 | 3.48E-22 | 4.48E-21 |
| HTRA1 | -1.999864603 | 1.59E-111 | 7.69E-109 |
| HTRA3 | -1.500456299 | 4.18E-42 | 1.72E-40 |
| HYAL1 | -0.963450339 | 2.23E-14 | 1.67E-13 |
| HYAL2 | -1.049770016 | 2.16E-09 | 1.03E-08 |
| HYLS1 | -0.617887484 | 0.005881458 | 0.012114074 |
| HYOU1 | -0.807559266 | 2.49E-16 | 2.17E-15 |
| HYPK | 1.357357766 | 1.65E-31 | 3.76E-30 |
| IAH1 | 0.443212695 | 1.18E-05 | 3.72E-05 |
| IARS1 | -0.774956714 | 1.35E-17 | 1.30E-16 |
| IARS2 | -1.102743853 | 7.48E-17 | 6.76E-16 |
| IBA57 | -0.259434826 | 0.014462517 | 0.027362824 |
| ICE2 | -0.833059696 | 1.05E-05 | 3.36E-05 |
| ICMT | 0.274262239 | 0.005075227 | 0.010598015 |
| ID1 | -0.270868978 | 0.010038419 | 0.019646104 |
| ID2 | 1.024300294 | 6.86E-35 | 1.88E-33 |
| ID3 | 1.307452006 | 7.31E-15 | 5.66E-14 |
| ID4 | 0.937274323 | 9.53E-20 | 1.06E-18 |
| IDE | -1.223411577 | 9.53E-21 | 1.12E-19 |
| IDH1 | -0.741408528 | 4.59E-17 | 4.23E-16 |
| IDH2 | 0.25585917 | 0.015142955 | 0.028538968 |
| IDNK | 0.518467215 | 2.64E-05 | 7.95E-05 |
| IER3IP1 | 0.487990761 | 5.29E-05 | 0.000153875 |
| IER5L | 3.149742242 | 0.013168351 | 0.02518776 |
| IFFO1 | -0.792841402 | 1.31E-06 | 4.70E-06 |
| IFFO2 | 0.785375507 | 2.81E-10 | 1.46E-09 |
| IFI27L2 | 1.227751926 | 0.009001651 | 0.01779384 |
| IFI30 | -0.47775811 | 0.006922545 | 0.014045564 |
| IFI35 | 0.448309198 | 0.000438776 | 0.001109551 |
| IFI6 | 6.482424662 | 5.80E-37 | 1.83E-35 |
| IFIT5 | 4.175636248 | 3.62E-106 | 1.54E-103 |
| IFNAR1 | -1.158501109 | 7.78E-26 | 1.28E-24 |
| IFNAR2 | 1.203450418 | 7.53E-39 | 2.59E-37 |
| IFNGR2 | 1.309090335 | 1.64E-53 | 1.11E-51 |
| IFNL3A | 7.244697901 | 1.49E-08 | 6.61E-08 |

| | | | |
|---|---|---|---|
| IFRD1 | -1.672684332 | 3.34E-56 | 2.50E-54 |
| IFT140 | -0.529201605 | 0.002370144 | 0.005305501 |
| IFT172 | -0.453236849 | 0.010342024 | 0.020185065 |
| IFT20 | -0.331647156 | 0.004324959 | 0.009162226 |
| IFT22 | -0.659190329 | 4.57E-05 | 0.000133896 |
| IFT46 | 0.699763437 | 5.88E-14 | 4.26E-13 |
| IFT52 | 0.637636043 | 2.80E-12 | 1.76E-11 |
| IFT57 | -0.662839329 | 3.32E-06 | 1.13E-05 |
| IFT80 | 0.286619552 | 0.022275112 | 0.040411548 |
| IFT81 | -0.577966924 | 1.27E-05 | 3.99E-05 |
| IGDCC3 | 0.550172188 | 1.37E-05 | 4.28E-05 |
| IGF2BP3 | -1.101873321 | 8.48E-11 | 4.62E-10 |
| IGF2R | -1.692623464 | 5.77E-28 | 1.08E-26 |
| IGFBP4 | 2.205250061 | 5.65E-62 | 5.33E-60 |
| IGFBP7 | -1.102642051 | 7.20E-17 | 6.52E-16 |
| IGSF1 | 1.092105021 | 0.00067537 | 0.001655778 |
| IGSF3 | -0.300068446 | 0.005200478 | 0.010834595 |
| IK | -0.312658824 | 0.000509946 | 0.001275287 |
| IKBKB | -0.581768002 | 2.32E-06 | 8.09E-06 |
| IKBKE | 1.22897621 | 7.66E-06 | 2.49E-05 |
| IKZF2 | 0.677380363 | 0.002159148 | 0.00486605 |
| IKZF5 | 0.84241319 | 6.96E-14 | 5.01E-13 |
| IL10RA | 2.947688292 | 3.28E-22 | 4.24E-21 |
| IL10RB | 0.961921056 | 7.93E-22 | 9.92E-21 |
| IL11RA | 1.619228481 | 1.62E-16 | 1.43E-15 |
| IL12A | 4.915167565 | 1.44E-21 | 1.78E-20 |
| IL13RA1 | -2.088710642 | 1.11E-38 | 3.80E-37 |
| IL13RA2 | -0.480714221 | 0.00166994 | 0.003839929 |
| IL15 | 1.298532954 | 1.65E-15 | 1.33E-14 |
| IL16 | -0.993637865 | 1.18E-06 | 4.27E-06 |
| IL17RA | 0.323027726 | 0.000655275 | 0.001608252 |
| IL18 | 0.880595733 | 2.86E-10 | 1.48E-09 |
| IL18R1 | 2.115486554 | 5.02E-08 | 2.11E-07 |
| IL18RAP | 3.74718982 | 2.47E-09 | 1.17E-08 |
| IL19 | 1.451009385 | 0.004212349 | 0.008962781 |
| IL1R1 | 2.071452796 | 8.43E-99 | 2.86E-96 |
| IL1R2 | 0.514363685 | 7.18E-06 | 2.34E-05 |
| IL1RAP | 0.234917639 | 0.027916791 | 0.049413339 |
| IL1RL2 | 1.455396956 | 3.58E-38 | 1.20E-36 |
| IL20RA | 2.226824975 | 0.000791159 | 0.001917516 |
| IL22RA2 | 3.73204565 | 0.000191579 | 0.000510878 |
| IL2RB | -3.374200142 | 0.028230362 | 0.049916363 |
| IL31RA | 0.510975426 | 5.97E-05 | 0.000172268 |
| IL6 | 1.196259673 | 1.39E-41 | 5.48E-40 |
| IL6ST | -0.329259783 | 0.012568736 | 0.024156428 |
| IL7 | 1.993352596 | 0.013225684 | 0.025288447 |
| IL7R | 2.042324474 | 0.002757778 | 0.006093923 |
| IL8 | 3.641498059 | 0 | 0 |

| | | | |
|---|---|---|---|
| IL8L1 | 3.076241962 | 6.42E-90 | 1.75E-87 |
| ILDR2 | -1.73244614 | 1.49E-06 | 5.30E-06 |
| ILF2 | 0.451295449 | 2.96E-09 | 1.39E-08 |
| ILVBL | -1.631040894 | 4.96E-27 | 8.78E-26 |
| IMMP1L | 0.571540758 | 0.000597537 | 0.001476678 |
| IMMT | -0.241003582 | 0.004730327 | 0.009943489 |
| IMP4 | -0.351312635 | 0.000406698 | 0.001033433 |
| IMPA2 | 0.717062547 | 0.000991607 | 0.00237271 |
| IMPDH2 | -0.479933069 | 2.74E-09 | 1.30E-08 |
| IMPG2 | 1.309764121 | 0.001087019 | 0.002586165 |
| INAFM2 | 1.179680653 | 6.50E-18 | 6.37E-17 |
| INCENP | 0.408322852 | 0.001843547 | 0.004203571 |
| ING1 | -1.23740788 | 7.66E-18 | 7.45E-17 |
| ING2 | -1.592942748 | 1.47E-17 | 1.41E-16 |
| ING3 | -1.450958534 | 2.78E-37 | 8.92E-36 |
| ING4 | 0.327853393 | 0.001426358 | 0.00332027 |
| INHBB | 1.524121898 | 4.22E-06 | 1.42E-05 |
| INIP | 0.627273187 | 1.50E-05 | 4.66E-05 |
| INO80 | -0.931986738 | 1.34E-09 | 6.53E-09 |
| INO80C | 0.38962891 | 0.008825952 | 0.017466853 |
| INPP5A | 1.353465575 | 2.53E-42 | 1.05E-40 |
| INPP5D | 2.513384941 | 0.000146832 | 0.000397875 |
| INPP5F | 0.921415795 | 1.24E-15 | 1.01E-14 |
| INSIG1 | -0.277245209 | 0.000281836 | 0.000733993 |
| INSIG2 | 0.728758093 | 3.64E-15 | 2.88E-14 |
| INSR | 0.947444626 | 1.32E-06 | 4.73E-06 |
| INSYN2A | -0.548882975 | 0.016431185 | 0.03069443 |
| INTS1 | -1.614823175 | 6.04E-19 | 6.37E-18 |
| INTS10 | -0.82933289 | 5.49E-15 | 4.29E-14 |
| INTS11 | -0.582845383 | 6.13E-07 | 2.29E-06 |
| INTS12 | -1.380188919 | 7.20E-22 | 9.08E-21 |
| INTS13 | 0.825085429 | 1.07E-23 | 1.52E-22 |
| INTS2 | -0.495257053 | 7.07E-06 | 2.31E-05 |
| INTS4 | -0.635964854 | 7.36E-07 | 2.73E-06 |
| INTS6 | -1.166907036 | 6.13E-10 | 3.08E-09 |
| INTS6L | -0.658933954 | 0.00013353 | 0.000363531 |
| INTS7 | -1.059208175 | 4.14E-11 | 2.32E-10 |
| INTS8 | -0.539603393 | 3.30E-07 | 1.27E-06 |
| INTS9 | -0.846338964 | 2.36E-09 | 1.12E-08 |
| IP6K2 | 0.585891558 | 3.38E-11 | 1.91E-10 |
| IPMK | 0.541089548 | 0.000197567 | 0.000525643 |
| IPO11 | -0.427919639 | 0.000165382 | 0.00044486 |
| IPO13 | -1.120031159 | 1.92E-12 | 1.22E-11 |
| IPO5 | -1.052721334 | 2.02E-24 | 3.01E-23 |
| IPO7 | 0.263723906 | 0.001332932 | 0.003115425 |
| IPO9 | -0.786964395 | 4.59E-08 | 1.94E-07 |
| IPP | 0.227167878 | 0.017372324 | 0.0322751 |
| IPPK | -0.359426818 | 0.003689819 | 0.007953065 |

| | | | |
|---|---|---|---|
| IQCC | -1.106653014 | 0.002220304 | 0.004990636 |
| IQCD | -0.670581927 | 0.001546875 | 0.003576918 |
| IQCG | 1.369203744 | 0.004056638 | 0.00867356 |
| IQCK | 0.962502547 | 2.03E-08 | 8.88E-08 |
| IQGAP1 | -0.384342916 | 3.18E-05 | 9.51E-05 |
| IQGAP2 | -0.615462969 | 1.21E-05 | 3.82E-05 |
| IQSEC1 | -0.322020566 | 0.002590312 | 0.005762278 |
| IQUB | 1.480360672 | 2.90E-06 | 1.00E-05 |
| IRAG1 | 0.256500871 | 0.02310896 | 0.041740464 |
| IRAK2 | 0.965811227 | 2.43E-23 | 3.36E-22 |
| IRAK4 | 0.294676924 | 0.016284926 | 0.030463077 |
| IREB2 | -1.076898934 | 6.63E-22 | 8.40E-21 |
| IRF1 | 2.994316533 | 1.34E-81 | 2.80E-79 |
| IRF2 | -0.287649138 | 0.004868578 | 0.010205693 |
| IRF2BPL | -0.90551255 | 1.36E-05 | 4.26E-05 |
| IRF6 | -0.375745845 | 0.011889261 | 0.022934833 |
| IRF7 | 1.266332158 | 2.85E-16 | 2.47E-15 |
| IRS1 | -1.595817838 | 7.23E-14 | 5.20E-13 |
| IRS4 | -1.279610265 | 2.58E-15 | 2.06E-14 |
| ISCA1 | 1.122451256 | 6.86E-30 | 1.43E-28 |
| ISCU | 0.45415795 | 6.29E-05 | 0.000180785 |
| ISG20 | -0.754343025 | 1.90E-06 | 6.73E-06 |
| ISG20L2 | -1.172584979 | 5.86E-13 | 3.90E-12 |
| ISL1 | -3.892047224 | 0.006114829 | 0.012543428 |
| ISLR | -1.195290679 | 3.08E-14 | 2.28E-13 |
| ISM1 | -0.824169876 | 0.0041602 | 0.008862922 |
| IST1 | -0.300837663 | 0.000199539 | 0.000530646 |
| ITCH | -0.562851956 | 0.002050177 | 0.004638921 |
| ITFG2 | -0.296587042 | 0.021407752 | 0.038968015 |
| ITGA11 | -0.691143607 | 1.42E-13 | 9.98E-13 |
| ITGA2 | -0.99125522 | 4.50E-05 | 0.000132299 |
| ITGA3 | -1.582406999 | 3.75E-24 | 5.45E-23 |
| ITGA4 | -0.94371762 | 1.32E-05 | 4.13E-05 |
| ITGA6 | -0.965630838 | 1.17E-23 | 1.66E-22 |
| ITGA8 | -0.584171955 | 1.24E-07 | 5.02E-07 |
| ITGA9 | -0.8859089 | 3.49E-10 | 1.79E-09 |
| ITGAV | -1.231003692 | 2.37E-41 | 9.18E-40 |
| ITGB1 | -0.87261604 | 2.60E-27 | 4.70E-26 |
| ITGB1BP1 | -1.16612835 | 1.39E-16 | 1.23E-15 |
| ITGB1BP2 | 2.175563761 | 0.000846373 | 0.002042959 |
| ITGB1BP3 | 2.135220131 | 2.18E-142 | 1.85E-139 |
| ITGB3 | -0.870108902 | 3.99E-22 | 5.13E-21 |
| ITGB5 | -0.282914842 | 0.001811489 | 0.004137415 |
| ITIH5 | -0.787959007 | 0.003162616 | 0.006918762 |
| ITM2B | -0.387218199 | 3.65E-06 | 1.24E-05 |
| ITM2C | 1.477349991 | 1.97E-05 | 6.04E-05 |
| ITPA | -0.547313976 | 2.70E-05 | 8.15E-05 |
| ITPK1 | -1.227637617 | 3.98E-29 | 7.94E-28 |

| | | | |
|---|---|---|---|
| ITPKA | -1.716879689 | 1.18E-07 | 4.78E-07 |
| ITPKB | -0.43716987 | 0.008102504 | 0.016197857 |
| ITPR1 | 0.382031137 | 0.000984129 | 0.002355809 |
| ITSN1 | 1.029962388 | 2.90E-37 | 9.25E-36 |
| IVD | -0.522154817 | 1.08E-06 | 3.92E-06 |
| IVNS1ABP | -0.230783681 | 0.020573595 | 0.037590577 |
| IWS1 | -0.238174189 | 0.019865981 | 0.03645449 |
| JADE2 | 0.633405275 | 0.018006424 | 0.033356774 |
| JAG1 | -0.875523041 | 5.84E-05 | 0.000168648 |
| JAG2 | -0.799709282 | 1.64E-06 | 5.83E-06 |
| JAGN1 | -0.619491903 | 8.66E-07 | 3.18E-06 |
| JAK1 | -0.54317645 | 2.58E-09 | 1.22E-08 |
| JAK2 | -0.60993916 | 7.80E-05 | 0.000220705 |
| JAM2 | 0.852147098 | 0.01299552 | 0.024899227 |
| JAZF1 | 0.661854263 | 0.002002404 | 0.0045346 |
| JCAD | 1.396375328 | 1.39E-24 | 2.10E-23 |
| JCHAIN | 1.320714219 | 0.013795776 | 0.02623282 |
| JHY | 4.462822629 | 2.35E-13 | 1.62E-12 |
| JMJD4 | 1.262213138 | 1.92E-43 | 8.37E-42 |
| JMJD6 | -0.805062127 | 9.57E-21 | 1.12E-19 |
| JMJD7 | -0.528503551 | 0.018340504 | 0.033916208 |
| JMJD8 | 0.74414383 | 7.05E-15 | 5.47E-14 |
| JMY | -1.194683314 | 1.25E-11 | 7.37E-11 |
| JOSD1 | 0.577725359 | 0.003152141 | 0.006898068 |
| JPH3 | 2.60000731 | 5.63E-05 | 0.000162971 |
| JUN | -2.33184747 | 1.21E-74 | 2.08E-72 |
| JUND | -1.390595543 | 0.000108083 | 0.000299077 |
| JUP | -1.209769911 | 1.23E-20 | 1.42E-19 |
| K123 | 1.158528598 | 1.04E-18 | 1.08E-17 |
| KALRN | -0.482276654 | 0.003404548 | 0.00739093 |
| KANK1 | -0.439810339 | 0.013258155 | 0.025341702 |
| KANK3 | 0.487931081 | 7.90E-06 | 2.56E-05 |
| KANSL1L | -1.165085164 | 8.58E-23 | 1.15E-21 |
| KANSL2 | -1.005759812 | 1.13E-11 | 6.69E-11 |
| KARS | -0.542892898 | 3.00E-08 | 1.29E-07 |
| KAT2A | -1.11771584 | 3.36E-18 | 3.37E-17 |
| KAT2B | 0.992474005 | 1.76E-20 | 2.03E-19 |
| KAT6A | 0.408446115 | 0.00171538 | 0.003935096 |
| KAT7 | -0.59504312 | 1.94E-07 | 7.65E-07 |
| KATNA1 | -1.19000189 | 1.04E-09 | 5.10E-09 |
| KATNB1 | -0.958521471 | 2.93E-10 | 1.52E-09 |
| KATNBL1 | -0.662777682 | 1.04E-11 | 6.17E-11 |
| KATNIP | -0.849642614 | 0.000129133 | 0.000352658 |
| KAZALD1 | 0.827312563 | 0.001683479 | 0.003868443 |
| KBP | 2.738378084 | 0.005309056 | 0.011045567 |
| KBTBD12 | 3.2358827 | 0.024341771 | 0.043781044 |
| KBTBD13 | 3.398900714 | 0.003317147 | 0.007220799 |
| KBTBD2 | -1.367885836 | 1.12E-26 | 1.94E-25 |

| | | | |
|---|---|---|---|
| KCNAB1 | 4.217563958 | 2.18E-08 | 9.48E-08 |
| KCNB1 | 3.581430264 | 1.76E-08 | 7.70E-08 |
| KCNJ15 | 3.068249754 | 1.62E-06 | 5.75E-06 |
| KCNJ16 | 1.61765513 | 6.67E-05 | 0.000190693 |
| KCNJ4 | 1.860353603 | 0.000305499 | 0.000791373 |
| KCNJ5 | 0.612024879 | 5.11E-12 | 3.13E-11 |
| KCNK1 | -0.224924049 | 0.00864258 | 0.017142619 |
| KCNK10 | 0.916525412 | 0.004232299 | 0.008996287 |
| KCNK15 | -0.674585613 | 7.45E-05 | 0.000211356 |
| KCNK2 | 1.341712966 | 0.024410784 | 0.043893554 |
| KCNK3 | 1.377256924 | 0.019859343 | 0.03644723 |
| KCNMB1 | 1.48215156 | 3.19E-31 | 7.17E-30 |
| KCNN2 | -1.349576635 | 0.001941952 | 0.004407002 |
| KCNS1 | 4.188114893 | 2.64E-09 | 1.25E-08 |
| KCTD12 | -1.002605468 | 0.004393499 | 0.009294388 |
| KCTD15 | 1.525338112 | 2.20E-19 | 2.38E-18 |
| KCTD2 | 0.363086144 | 1.50E-05 | 4.67E-05 |
| KCTD20 | -1.120978204 | 9.59E-26 | 1.57E-24 |
| KCTD21 | 1.755856338 | 0.003710306 | 0.007992153 |
| KCTD3 | 0.448775478 | 1.28E-05 | 4.01E-05 |
| KCTD9 | 0.361068251 | 8.31E-06 | 2.68E-05 |
| KDELR2 | 0.935505502 | 2.26E-35 | 6.42E-34 |
| KDELR3 | 0.643234578 | 8.32E-12 | 5.01E-11 |
| KDM1A | -0.735039854 | 4.05E-10 | 2.06E-09 |
| KDM3A | -0.987040662 | 1.52E-08 | 6.72E-08 |
| KDM3B | -0.581439195 | 6.85E-06 | 2.24E-05 |
| KDM4B | -0.811043628 | 8.58E-08 | 3.52E-07 |
| KDM4C | -0.561798579 | 0.001385182 | 0.003229952 |
| KDM5A | -0.668249867 | 0.000430153 | 0.001088961 |
| KDM5B | -0.618203041 | 3.67E-06 | 1.24E-05 |
| KDM7A | -0.27105381 | 0.023879925 | 0.04301868 |
| KDR | -5.356167864 | 9.03E-05 | 0.000252898 |
| KERA | 0.622236906 | 0.011535589 | 0.022312754 |
| KHDRBS1 | -1.037507251 | 6.00E-20 | 6.72E-19 |
| KIAA0100 | -0.5203699 | 0.008612102 | 0.017090917 |
| KIAA0232 | 0.680455377 | 1.61E-07 | 6.41E-07 |
| KIAA0319 | 1.962576097 | 0.001266918 | 0.002974084 |
| KIAA0319L | 0.284970032 | 0.002770157 | 0.006119288 |
| KIAA0825 | -1.834113925 | 0.000153262 | 0.00041439 |
| KIAA0895 | 1.005841576 | 3.13E-14 | 2.31E-13 |
| KIAA0907 | -0.23995213 | 0.026019003 | 0.046392534 |
| KIAA0930 | -0.277836315 | 0.003138046 | 0.006869435 |
| KIAA1143 | 0.935136127 | 2.54E-27 | 4.61E-26 |
| KIAA1191 | -0.290828573 | 0.001500904 | 0.003480683 |
| KIAA1217 | -0.537580911 | 0.012101966 | 0.023321979 |
| KIAA1328 | 0.753389309 | 7.09E-05 | 0.000201846 |
| KIAA1522 | 0.921938814 | 2.45E-13 | 1.69E-12 |
| KIAA1549 | -0.401199456 | 0.010621823 | 0.020701437 |

| | | | |
|---|---|---|---|
| KIAA1671 | 1.376631652 | 3.01E-69 | 3.82E-67 |
| KIAA2013 | -0.41183216 | 0.000483918 | 0.001211756 |
| KIAA2026 | 0.46397322 | 9.64E-05 | 0.00026908 |
| KIDINS220 | -1.614653885 | 2.36E-33 | 5.92E-32 |
| KIF11 | -1.22225198 | 4.37E-06 | 1.46E-05 |
| KIF14 | 2.675875141 | 4.49E-17 | 4.15E-16 |
| KIF15 | -1.242246188 | 0.008436465 | 0.016793853 |
| KIF18A | -0.524535907 | 0.015831439 | 0.029687828 |
| KIF1A | -1.747858732 | 2.05E-08 | 8.94E-08 |
| KIF1B | -1.17176484 | 1.12E-27 | 2.06E-26 |
| KIF1BP | -0.867912726 | 2.33E-16 | 2.04E-15 |
| KIF20A | -0.756894704 | 0.000452479 | 0.001141442 |
| KIF23 | -1.165085615 | 2.51E-05 | 7.60E-05 |
| KIF26A | -0.841467079 | 2.32E-11 | 1.33E-10 |
| KIF2A | 0.63942954 | 1.27E-10 | 6.81E-10 |
| KIF2C | -1.152459003 | 0.00070319 | 0.001719022 |
| KIF3A | -0.619662359 | 1.42E-06 | 5.08E-06 |
| KIF4B | 0.762464953 | 0.000414179 | 0.001050871 |
| KIF5B | -0.823519783 | 1.19E-21 | 1.47E-20 |
| KIF7 | -0.824020175 | 3.23E-10 | 1.67E-09 |
| KIF9 | 0.913458532 | 0.004789232 | 0.010056685 |
| KIFAP3 | -1.163303137 | 1.67E-27 | 3.04E-26 |
| KIFC1 | -0.778997052 | 8.55E-10 | 4.24E-09 |
| KIFC3 | 0.519145744 | 3.74E-05 | 0.000111048 |
| KIN | -0.839055046 | 1.18E-06 | 4.28E-06 |
| KIRREL1 | 0.895949117 | 1.36E-17 | 1.31E-16 |
| KL | 4.100410518 | 2.09E-07 | 8.21E-07 |
| KLF10 | 0.37854261 | 1.58E-05 | 4.89E-05 |
| KLF11 | -0.79430487 | 2.92E-14 | 2.17E-13 |
| KLF13 | 0.850934705 | 2.48E-19 | 2.68E-18 |
| KLF2 | -2.204717767 | 1.11E-111 | 5.59E-109 |
| KLF3 | -1.36486246 | 1.75E-21 | 2.15E-20 |
| KLF5 | -0.41466316 | 0.001460936 | 0.00339669 |
| KLF9 | 1.560768832 | 2.87E-25 | 4.57E-24 |
| KLHDC1 | -0.48996947 | 0.004356547 | 0.009223394 |
| KLHDC2 | -0.458678864 | 3.45E-06 | 1.17E-05 |
| KLHL17 | 0.982428639 | 0.017660076 | 0.032773861 |
| KLHL18 | -0.873574032 | 3.61E-10 | 1.85E-09 |
| KLHL20 | -0.706812941 | 1.43E-10 | 7.60E-10 |
| KLHL21 | -1.223462732 | 3.82E-24 | 5.53E-23 |
| KLHL24 | -0.834153743 | 2.60E-16 | 2.26E-15 |
| KLHL28 | -1.096634502 | 3.81E-18 | 3.81E-17 |
| KLHL36 | 1.369018842 | 5.34E-47 | 2.82E-45 |
| KLHL40 | -1.17583333 | 2.84E-14 | 2.11E-13 |
| KLHL42 | -0.671355689 | 6.99E-08 | 2.89E-07 |
| KLHL5 | 1.473237175 | 6.11E-67 | 7.42E-65 |
| KLHL7 | 0.514082648 | 4.13E-08 | 1.75E-07 |
| KMT2E | -0.379591099 | 0.001875653 | 0.004269617 |

| | | | |
|---|---|---|---|
| KMT5B | 0.597030191 | 1.91E-09 | 9.19E-09 |
| KNG1 | -1.998745075 | 0.002198095 | 0.004946442 |
| KPNA1 | -1.274853723 | 3.46E-17 | 3.23E-16 |
| KPNA2 | -1.835421235 | 7.30E-06 | 2.38E-05 |
| KPNA3 | -0.313486861 | 0.002073524 | 0.004686288 |
| KPNA4 | -0.681284446 | 3.91E-16 | 3.36E-15 |
| KPNA5 | -0.357694333 | 0.003824535 | 0.008206991 |
| KPNA6 | -0.61475171 | 3.12E-11 | 1.77E-10 |
| KPNB1 | -0.484440914 | 2.42E-06 | 8.43E-06 |
| KRAS | 0.800810535 | 1.09E-20 | 1.27E-19 |
| KRIT1 | 0.319979671 | 0.002211565 | 0.004972637 |
| KRT12 | 2.774903723 | 0.00715682 | 0.014470036 |
| KRT13 | 3.880860394 | 8.63E-19 | 9.02E-18 |
| KRT14 | 0.839748145 | 5.38E-16 | 4.56E-15 |
| KRT17 | 0.848446103 | 7.21E-12 | 4.36E-11 |
| KRT18 | -1.517158517 | 1.72E-10 | 9.08E-10 |
| KRT222 | 3.93423051 | 0.003589765 | 0.007755855 |
| KRT23 | -0.705063487 | 0.001438638 | 0.003346563 |
| KRT40 | 2.929580173 | 8.02E-05 | 0.000226511 |
| KRT7 | -1.305667829 | 8.56E-25 | 1.31E-23 |
| KRT75 | -2.454012556 | 3.56E-22 | 4.59E-21 |
| KRTAP10-4 | 0.923687372 | 2.94E-09 | 1.39E-08 |
| KTN1 | -1.322346305 | 3.86E-36 | 1.15E-34 |
| KXD1 | 0.910638547 | 2.19E-16 | 1.92E-15 |
| KYAT3 | -0.428158903 | 0.002726427 | 0.006033474 |
| L2HGDH | 0.335037033 | 0.014249263 | 0.027011993 |
| L3MBTL2 | -0.455044405 | 0.000764453 | 0.001858422 |
| LACTB | 0.708223125 | 3.23E-06 | 1.10E-05 |
| LACTB2 | -0.451457963 | 0.000679603 | 0.001665555 |
| LACTBL1 | 1.503510227 | 1.59E-09 | 7.68E-09 |
| LAMA1 | 0.882875359 | 0.025177899 | 0.045064261 |
| LAMA3 | -0.81472994 | 0.000240648 | 0.000633647 |
| LAMA5 | -0.322247481 | 0.014799984 | 0.027946844 |
| LAMB1 | -0.862888363 | 4.03E-13 | 2.74E-12 |
| LAMB2 | -1.419740217 | 6.37E-22 | 8.09E-21 |
| LAMB3 | -0.963694661 | 1.35E-06 | 4.85E-06 |
| LAMC1 | -1.250410803 | 6.35E-20 | 7.11E-19 |
| LAMP1 | -0.244073293 | 0.002680714 | 0.005942956 |
| LAMP2 | -0.364973983 | 5.28E-05 | 0.000153665 |
| LAMTOR1 | 0.900405734 | 0.007428951 | 0.014963589 |
| LAMTOR2 | 0.911269909 | 1.22E-11 | 7.24E-11 |
| LAMTOR5 | 1.245553515 | 1.57E-31 | 3.58E-30 |
| LANCL1 | -0.539054455 | 1.64E-08 | 7.24E-08 |
| LANCL2 | -0.322989941 | 0.001879209 | 0.004276279 |
| LANCL3 | -0.870815258 | 0.002191568 | 0.004934204 |
| LARP1 | -0.281729955 | 0.020059247 | 0.036749601 |
| LARP6 | -0.661071746 | 0.004210921 | 0.008961145 |
| LARP7 | -0.413100502 | 0.02581419 | 0.046069652 |

| | | | |
|---|---|---|---|
| LARS1 | -0.427263494 | 1.57E-05 | 4.88E-05 |
| LARS2 | 0.430153743 | 0.00045804 | 0.001153757 |
| LAS1L | -0.326425305 | 0.000409793 | 0.001040714 |
| LASP1 | 1.020813215 | 1.04E-18 | 1.08E-17 |
| LAT2 | 0.966326456 | 0.006620704 | 0.013495592 |
| LATS1 | -0.29361266 | 0.016755747 | 0.031249193 |
| LATS2 | 0.375395809 | 0.001202072 | 0.002838035 |
| LBFABP | 3.621865987 | 0.00068818 | 0.001684449 |
| LBH | 0.738686023 | 8.57E-21 | 1.01E-19 |
| LBR | -0.861241157 | 6.74E-17 | 6.12E-16 |
| LCA5 | -0.534802887 | 0.00146655 | 0.003408575 |
| LCLAT1 | 0.947747517 | 3.16E-29 | 6.39E-28 |
| LCORL | -0.724805512 | 0.000208697 | 0.000553377 |
| LCP1 | 2.260649224 | 0.01288483 | 0.024718505 |
| LCT | 2.8588423 | 3.70E-06 | 1.25E-05 |
| LDAH | 0.767358218 | 7.57E-13 | 5.00E-12 |
| LDB2 | 0.781949703 | 2.13E-06 | 7.48E-06 |
| LDHA | -0.897649269 | 4.83E-21 | 5.75E-20 |
| LEAP2 | 6.240952075 | 5.46E-33 | 1.34E-31 |
| LEF1 | -1.468442754 | 9.99E-06 | 3.19E-05 |
| LEMD2 | -0.266531386 | 0.019059184 | 0.035125753 |
| LEMD3 | 0.740182585 | 2.62E-19 | 2.83E-18 |
| LEO1 | -1.20307602 | 1.02E-16 | 9.18E-16 |
| LEPROT | 0.779421022 | 9.03E-24 | 1.29E-22 |
| LEPROTL1 | 0.451687991 | 2.99E-06 | 1.03E-05 |
| LETM1 | -0.243723738 | 0.014247496 | 0.027011993 |
| LETM2 | -0.672372754 | 1.43E-06 | 5.13E-06 |
| LFNG | -0.564232836 | 3.67E-11 | 2.06E-10 |
| LGALS2 | 1.100194984 | 4.93E-29 | 9.82E-28 |
| LGALS3 | 1.585765488 | 2.76E-64 | 2.93E-62 |
| LGALSL | 0.818510934 | 1.13E-07 | 4.58E-07 |
| LGI1 | -0.883492692 | 0.000108332 | 0.000299704 |
| LGMN | -0.372360558 | 2.41E-07 | 9.43E-07 |
| LGR4 | -0.455068409 | 0.025487042 | 0.045551595 |
| LGR5 | 1.777587217 | 0.005635402 | 0.01165121 |
| LHFPL2 | 0.242186761 | 0.013431624 | 0.025644422 |
| LHFPL5 | 0.962801875 | 1.66E-22 | 2.19E-21 |
| LHFPL6 | 1.291462896 | 8.29E-48 | 4.52E-46 |
| LHX2 | 1.738898602 | 1.16E-13 | 8.22E-13 |
| LIAS | -0.598992629 | 7.20E-07 | 2.67E-06 |
| LIG1 | -1.508647321 | 8.17E-06 | 2.64E-05 |
| LIG3 | -1.202744915 | 6.66E-28 | 1.24E-26 |
| LIG4 | -1.052541059 | 3.79E-13 | 2.58E-12 |
| LIMA1 | 0.259284004 | 0.009097256 | 0.017964536 |
| LIMK1 | -1.31619321 | 6.17E-06 | 2.03E-05 |
| LIMK2 | -0.393168946 | 0.003328692 | 0.007243609 |
| LIMS1 | -0.194744077 | 0.025097399 | 0.044927944 |
| LIN52 | 0.326476957 | 0.000537599 | 0.001338287 |

| | | | |
|---|---|---|---|
| LIN54 | -1.10323249 | 4.08E-07 | 1.56E-06 |
| LIN9 | -1.146078734 | 4.00E-08 | 1.69E-07 |
| LINGO1 | 1.525475755 | 8.10E-62 | 7.59E-60 |
| LINS1 | -1.447253707 | 0.001026315 | 0.002450321 |
| LIPT1 | -0.633942607 | 1.38E-08 | 6.12E-08 |
| LIX1 | 4.446602172 | 3.19E-08 | 1.36E-07 |
| LLPH | 0.58787587 | 1.49E-05 | 4.63E-05 |
| LMAN1L | -0.579871725 | 4.33E-11 | 2.42E-10 |
| LMAN2 | 0.204576584 | 0.013458042 | 0.025687643 |
| LMAN2L | -0.302114926 | 0.017590921 | 0.032654438 |
| LMBR1 | 0.329952908 | 0.013042786 | 0.024979224 |
| LMBRD1 | -1.006222212 | 1.63E-18 | 1.68E-17 |
| LMBRD2 | 0.381739186 | 0.000590963 | 0.001461759 |
| LMF2 | -0.793504967 | 0.001220814 | 0.002877784 |
| LMLN | -1.378745751 | 0.000252492 | 0.000662269 |
| LMNB1 | -0.730530628 | 1.79E-05 | 5.51E-05 |
| LMO7 | -0.371562015 | 4.10E-06 | 1.38E-05 |
| LMOD1 | 0.553528065 | 1.66E-06 | 5.88E-06 |
| LNP1 | 1.604438243 | 5.09E-08 | 2.14E-07 |
| LNPEP | -0.696759517 | 6.39E-05 | 0.000183335 |
| LNX1 | 1.70297905 | 7.14E-08 | 2.95E-07 |
| LONP1 | -1.194490209 | 3.23E-11 | 1.83E-10 |
| LONP2 | -1.451796405 | 6.67E-29 | 1.32E-27 |
| LONRF1 | -0.804564047 | 8.21E-14 | 5.86E-13 |
| LONRF2 | -0.868452232 | 7.35E-11 | 4.03E-10 |
| LONRF3 | 0.568211192 | 1.52E-10 | 8.07E-10 |
| LOX | -0.675706018 | 8.36E-06 | 2.70E-05 |
| LOXL1 | 0.476480911 | 0.000298301 | 0.000774056 |
| LOXL2 | -1.159435384 | 2.39E-41 | 9.23E-40 |
| LOXL3 | -1.884218173 | 6.89E-59 | 5.68E-57 |
| LOXL4 | -2.769576139 | 0.001055682 | 0.002515133 |
| LPAR1 | 1.140189175 | 0.001254998 | 0.002950179 |
| LPAR3 | 3.554631777 | 3.48E-14 | 2.57E-13 |
| LPAR4 | 1.147975098 | 2.12E-09 | 1.01E-08 |
| LPAR6 | -0.55888853 | 0.021919996 | 0.039820447 |
| LPCAT1 | -0.681910703 | 7.31E-06 | 2.38E-05 |
| LPCAT2 | -0.441473761 | 1.90E-06 | 6.73E-06 |
| LPCAT3 | 0.675792106 | 4.36E-11 | 2.44E-10 |
| LPGAT1 | 0.525961956 | 2.83E-07 | 1.10E-06 |
| LPIN1 | -0.979781423 | 1.27E-07 | 5.14E-07 |
| LPL | -1.128838699 | 1.26E-27 | 2.31E-26 |
| LPP | 0.518338136 | 2.11E-05 | 6.46E-05 |
| LRAT | -2.107566278 | 0.000751484 | 0.001828359 |
| LRATD1 | -1.931444962 | 1.39E-07 | 5.57E-07 |
| LRCH1 | -1.31916314 | 0.000259566 | 0.000679381 |
| LRCH2 | -0.64012055 | 0.000539772 | 0.001343311 |
| LRIF1 | -0.776543839 | 1.52E-09 | 7.37E-09 |
| LRIG2 | -1.060700598 | 8.26E-09 | 3.74E-08 |

| | | | |
|---|---|---|---|
| LRIG3 | -1.890577231 | 4.28E-26 | 7.18E-25 |
| LRIT1 | 4.75716826 | 0.004796614 | 0.01007037 |
| LRIT2 | 5.159961272 | 4.09E-05 | 0.000120604 |
| LRMP | -0.305513215 | 0.011105917 | 0.021542975 |
| LRP1 | -1.159245423 | 1.66E-15 | 1.35E-14 |
| LRP2BP | 0.878940578 | 9.64E-06 | 3.08E-05 |
| LRP4 | -0.823437178 | 0.002433046 | 0.005432883 |
| LRP5 | -0.559342437 | 0.000463796 | 0.001167175 |
| LRP6 | 0.618082382 | 3.75E-05 | 0.000111325 |
| LRPPRC | -0.727114684 | 1.07E-13 | 7.60E-13 |
| LRRC1 | -0.28757458 | 0.01310702 | 0.025083452 |
| LRRC15 | -1.063911962 | 7.00E-10 | 3.51E-09 |
| LRRC17 | 0.615167547 | 6.02E-08 | 2.50E-07 |
| LRRC2 | 1.567589265 | 0.00628575 | 0.012861044 |
| LRRC20 | 0.525959375 | 2.24E-05 | 6.82E-05 |
| LRRC34 | -1.054275811 | 8.88E-05 | 0.00024904 |
| LRRC38 | 1.959600756 | 1.12E-64 | 1.22E-62 |
| LRRC39 | 2.898426107 | 0.000741533 | 0.001805611 |
| LRRC41 | -0.954778158 | 7.58E-05 | 0.00021468 |
| LRRC47 | -0.810585413 | 1.18E-09 | 5.75E-09 |
| LRRC49 | -0.42710112 | 0.000375848 | 0.00096043 |
| LRRC59 | -0.413549186 | 4.26E-05 | 0.000125291 |
| LRRC6 | 0.714722749 | 0.000686592 | 0.001680905 |
| LRRC8A | -1.543503646 | 6.86E-29 | 1.35E-27 |
| LRRC8C | -1.121716791 | 4.21E-14 | 3.08E-13 |
| LRRC8D | -0.746304991 | 1.71E-10 | 9.05E-10 |
| LRRFIP2 | -0.838717093 | 3.11E-15 | 2.47E-14 |
| LRRK2 | -0.562597758 | 0.000338798 | 0.000872479 |
| LRRN2 | 3.772425328 | 4.63E-06 | 1.55E-05 |
| LRRN4 | -1.216457386 | 1.08E-25 | 1.77E-24 |
| LRSAM1 | 0.628845033 | 5.79E-15 | 4.52E-14 |
| LRTM2 | 4.130025291 | 2.33E-08 | 1.01E-07 |
| LRWD1 | -0.440687807 | 0.000172663 | 0.000463434 |
| LSAMP | 0.596515432 | 0.007474033 | 0.015047705 |
| LSM12 | 1.108338916 | 9.28E-33 | 2.24E-31 |
| LSM14A | -0.514628113 | 6.72E-10 | 3.37E-09 |
| LSM14B | -0.709930604 | 0.003736103 | 0.008037596 |
| LSM5 | 0.880578691 | 5.00E-14 | 3.65E-13 |
| LSM6 | 0.580248906 | 1.44E-05 | 4.49E-05 |
| LSM8 | -0.588940518 | 9.19E-05 | 0.000257333 |
| LSMEM1 | 1.77189267 | 3.59E-45 | 1.72E-43 |
| LSP1P1 | 0.456619484 | 9.55E-07 | 3.48E-06 |
| LSS | -0.806231938 | 1.72E-15 | 1.39E-14 |
| LTA4H | -0.445387106 | 5.39E-08 | 2.25E-07 |
| LTBP1 | -1.462405174 | 1.80E-73 | 2.71E-71 |
| LTBP2 | -1.54243962 | 2.64E-08 | 1.14E-07 |
| LTN1 | -0.946639861 | 8.24E-08 | 3.39E-07 |
| LUC7L2 | -0.527493069 | 9.70E-06 | 3.10E-05 |

| | | | |
|---|---|---|---|
| LUC7L3 | 0.958518453 | 2.60E-19 | 2.80E-18 |
| LUM | 0.826393991 | 6.69E-17 | 6.07E-16 |
| LUZP1 | -1.068186841 | 4.81E-13 | 3.24E-12 |
| LY6E | 1.71555571 | 6.84E-05 | 0.000195224 |
| LY75 | -0.928380382 | 1.13E-12 | 7.36E-12 |
| LY86 | 3.859923865 | 1.51E-06 | 5.38E-06 |
| LY96 | 4.275016951 | 0.000250234 | 0.000656725 |
| LYG2 | 2.85809152 | 0.003193031 | 0.006976319 |
| LYPD1 | 1.272670961 | 7.98E-53 | 5.37E-51 |
| LYPLA1 | 0.396739118 | 7.93E-07 | 2.93E-06 |
| LYPLA2 | -0.609942172 | 3.74E-09 | 1.74E-08 |
| LYPLAL1 | -0.497820141 | 8.83E-05 | 0.000248015 |
| LYRM1 | 0.439867847 | 0.000635655 | 0.001565186 |
| LYRM2 | -0.419989447 | 0.018907474 | 0.034865063 |
| LYRM4 | 0.53549468 | 3.44E-06 | 1.17E-05 |
| LYRM7 | 0.48152441 | 0.001572161 | 0.003632916 |
| LYSMD2 | -0.463310358 | 4.31E-07 | 1.64E-06 |
| LYSMD3 | -0.658259584 | 1.48E-09 | 7.16E-09 |
| LYSMD4 | -0.611444195 | 9.99E-05 | 0.000277972 |
| LYVE1 | 1.897440385 | 0.001285901 | 0.003013961 |
| LZIC | 0.586067393 | 6.11E-11 | 3.37E-10 |
| LZTR1 | -0.299207134 | 0.016376256 | 0.030604438 |
| LZTS3 | 1.268221289 | 1.59E-05 | 4.93E-05 |
| M6PR | 0.222605187 | 0.020639294 | 0.037690351 |
| MAB21L2 | -0.601874893 | 0.006471627 | 0.013213512 |
| MACF1 | 0.285073704 | 0.011622766 | 0.022462197 |
| MACIR | 0.54341205 | 0.000152893 | 0.000413474 |
| MAD1L1 | -1.30839224 | 6.07E-13 | 4.04E-12 |
| MAD2L1 | -0.527463669 | 7.89E-05 | 0.000222999 |
| MAD2L1BP | -1.117656377 | 7.34E-09 | 3.34E-08 |
| MADD | -0.46689584 | 0.014373329 | 0.027209232 |
| MADPRT1 | 0.939737455 | 0.004650056 | 0.009794446 |
| MAEA | -0.770793023 | 3.79E-14 | 2.78E-13 |
| MAF1 | 0.801024777 | 0.000467054 | 0.001173635 |
| MAFB | 0.477504463 | 3.30E-06 | 1.13E-05 |
| MAFK | 0.338894177 | 0.000517803 | 0.00129327 |
| MAGI1 | -0.887814554 | 7.62E-08 | 3.14E-07 |
| MAGI2 | 0.705160899 | 4.59E-10 | 2.33E-09 |
| MAGI3 | -1.460764969 | 9.13E-09 | 4.11E-08 |
| MAGT1 | -1.187594374 | 4.65E-30 | 9.82E-29 |
| MAIP1 | 1.008763388 | 2.81E-35 | 7.90E-34 |
| MAL | 0.849217599 | 0.000113963 | 0.000313752 |
| MALL | 0.571881036 | 5.98E-05 | 0.000172383 |
| MALRD1 | 0.750039857 | 0.000659045 | 0.001616964 |
| MALSU1 | 0.379707023 | 4.28E-05 | 0.000125737 |
| MAMDC2 | 0.513012557 | 0.000160009 | 0.000431173 |
| MAML2 | 1.063236118 | 2.06E-17 | 1.95E-16 |
| MAMLD1 | -1.639064848 | 1.14E-34 | 3.08E-33 |

| | | | |
|---|---|---|---|
| MAN1A1 | -0.724068713 | 2.84E-05 | 8.53E-05 |
| MAN1A2 | 0.479471451 | 6.55E-05 | 0.000187619 |
| MAN1B1 | 0.751767683 | 1.28E-15 | 1.05E-14 |
| MAN1C1 | 0.503015619 | 6.76E-07 | 2.52E-06 |
| MAN2A1 | -1.248267219 | 1.39E-12 | 8.97E-12 |
| MAN2B2 | -0.771645158 | 4.33E-13 | 2.93E-12 |
| MAN2C1 | -0.797275084 | 2.73E-07 | 1.06E-06 |
| MANBA | -0.437140796 | 0.000185246 | 0.00049496 |
| MANBAL4 | 1.083181148 | 3.64E-13 | 2.49E-12 |
| MANEA | 0.853239131 | 6.59E-16 | 5.53E-15 |
| MANSC1 | 0.589984416 | 0.001183711 | 0.002798087 |
| MANSC4 | 2.724210349 | 0.009243488 | 0.018216252 |
| MAP10 | -1.494463001 | 6.23E-05 | 0.000179245 |
| MAP1A | -0.682217929 | 2.75E-06 | 9.53E-06 |
| MAP1B | -1.165054059 | 5.53E-28 | 1.04E-26 |
| MAP1LC3A | 0.721879861 | 8.77E-16 | 7.28E-15 |
| MAP1LC3B | 0.53579328 | 3.98E-13 | 2.71E-12 |
| MAP2K2 | 0.489469835 | 0.003213125 | 0.007016836 |
| MAP2K3 | 0.386398448 | 1.32E-05 | 4.14E-05 |
| MAP2K4 | -0.423367054 | 5.52E-05 | 0.000160077 |
| MAP2K5 | 0.841721698 | 1.13E-19 | 1.25E-18 |
| MAP3K1 | -0.57298423 | 0.004609195 | 0.009714402 |
| MAP3K21 | -0.90064037 | 0.025865799 | 0.046149637 |
| MAP3K5 | 0.426294461 | 7.55E-06 | 2.46E-05 |
| MAP3K7 | 0.868968159 | 6.99E-28 | 1.30E-26 |
| MAP4K5 | 0.21205128 | 0.016574206 | 0.030931842 |
| MAP7 | 0.713070888 | 3.78E-11 | 2.12E-10 |
| MAP7D2 | 2.430115949 | 0.015049111 | 0.028385723 |
| MAPK13 | -0.911147503 | 2.52E-05 | 7.64E-05 |
| MAPK14 | 0.646396432 | 0.000418825 | 0.001062201 |
| MAPK1IP1L | 0.335066673 | 0.000276139 | 0.000720539 |
| MAPK4 | 1.251225019 | 3.57E-48 | 1.97E-46 |
| MAPK8 | -0.859926807 | 6.21E-11 | 3.42E-10 |
| MAPK8IP1 | -0.515716336 | 0.0002478 | 0.00065109 |
| MAPK9 | 0.30738905 | 0.003948354 | 0.008451343 |
| MAPKAP1 | 0.373173924 | 0.000885344 | 0.00212871 |
| MAPKAPK2 | 0.816057516 | 0.001525315 | 0.003532472 |
| MAPKAPK3 | 1.26452139 | 9.68E-27 | 1.68E-25 |
| MAPKAPK5 | 0.278284454 | 0.002924908 | 0.006441242 |
| MAPRE3 | 0.822340929 | 2.24E-12 | 1.42E-11 |
| MARCHF11 | 1.957673796 | 0.015232732 | 0.028692253 |
| MARCHF2 | 0.50291429 | 0.000579086 | 0.001435392 |
| MARCHF4 | 1.897068568 | 9.05E-05 | 0.000253584 |
| MARCHF6 | -0.431952688 | 1.88E-05 | 5.77E-05 |
| MARCKSL1 | 0.627180894 | 0.000933669 | 0.002240147 |
| MARF1 | 0.384581906 | 0.006684971 | 0.013612303 |
| MARS | -0.822950424 | 1.23E-13 | 8.66E-13 |
| MARS2 | -0.761400882 | 6.36E-06 | 2.09E-05 |

| | | | |
|---|---|---|---|
| MARVELD2 | -1.113781474 | 1.87E-08 | 8.18E-08 |
| MAS1 | 4.088309676 | 0.016295445 | 0.030478559 |
| MASTL | -1.05215217 | 0.000580649 | 0.001438867 |
| MAT1A | -0.945724717 | 5.97E-20 | 6.69E-19 |
| MAT2B | 0.386506402 | 6.60E-05 | 0.000188883 |
| MATN2 | 0.590278054 | 0.024621668 | 0.044231788 |
| MATN4 | 4.134692895 | 2.92E-10 | 1.51E-09 |
| MAVS | -1.384394937 | 6.68E-18 | 6.54E-17 |
| MAX | 1.53485826 | 0.01467597 | 0.027739644 |
| MB | 3.234099097 | 1.46E-12 | 9.40E-12 |
| MB21D2 | -1.422158498 | 1.18E-06 | 4.27E-06 |
| MBD5 | 0.458215787 | 0.005482272 | 0.011371134 |
| MBLAC2 | -0.983651762 | 0.001302412 | 0.003048981 |
| MBNL1 | 1.072458869 | 4.27E-38 | 1.43E-36 |
| MBNL2 | 0.955175082 | 1.16E-22 | 1.55E-21 |
| MBNL3 | 1.147846513 | 1.15E-31 | 2.63E-30 |
| MBOAT1 | 0.99376481 | 0.001520727 | 0.003523048 |
| MBP | 0.772090153 | 1.20E-11 | 7.08E-11 |
| MBTD1 | 0.461704677 | 0.001990949 | 0.004512418 |
| MBTPS1 | 1.247671889 | 9.68E-56 | 7.04E-54 |
| MBTPS2 | -0.662010874 | 0.003075928 | 0.006741054 |
| MCAM | -0.933615522 | 0.004519218 | 0.009535115 |
| MCAT | -0.678161391 | 0.000122282 | 0.000335499 |
| MCC | -0.597168117 | 0.001095356 | 0.002605544 |
| MCCC1 | -0.872640591 | 0.000229454 | 0.000605462 |
| MCCC2 | 0.429229496 | 4.19E-05 | 0.000123305 |
| MCCC2L | -1.467580046 | 3.11E-10 | 1.61E-09 |
| MCF2L2 | 1.545420302 | 1.87E-05 | 5.73E-05 |
| MCFD2 | 0.546403031 | 2.26E-08 | 9.80E-08 |
| MCL1 | -0.5085629 | 3.41E-09 | 1.60E-08 |
| MCM10 | -1.296793171 | 0.000245942 | 0.00064646 |
| MCM2 | -1.265044857 | 3.82E-21 | 4.60E-20 |
| MCM3 | -1.198475972 | 7.50E-05 | 0.000212703 |
| MCM4 | -1.384097307 | 1.92E-21 | 2.35E-20 |
| MCM5 | 2.243939721 | 1.32E-82 | 2.94E-80 |
| MCM6 | -1.267835422 | 1.31E-17 | 1.26E-16 |
| MCM8 | 0.310050471 | 0.012623596 | 0.02424815 |
| MCM9 | -1.720516553 | 7.16E-07 | 2.66E-06 |
| MCMBP | -0.642326213 | 1.75E-12 | 1.12E-11 |
| MCOLN1 | 0.410657568 | 7.12E-05 | 0.0002027 |
| MCOLN2 | 2.082505192 | 0.000549136 | 0.001365255 |
| MCOLN3 | 0.989044502 | 5.77E-07 | 2.16E-06 |
| MCPH1 | -0.670612007 | 0.012313143 | 0.023702059 |
| MCRIP1 | 0.727268467 | 5.23E-15 | 4.09E-14 |
| MCRIP2 | 1.083216207 | 2.40E-12 | 1.51E-11 |
| MCRS1 | 0.560891391 | 7.11E-10 | 3.55E-09 |
| MCTS1 | 0.912967031 | 3.93E-21 | 4.73E-20 |
| MCU | -1.626862368 | 1.53E-25 | 2.48E-24 |

| | | | |
|---|---|---|---|
| MCUB | 0.592852076 | 0.001730779 | 0.003965737 |
| MCUR1 | 0.552480877 | 4.57E-09 | 2.11E-08 |
| MDFIC | 0.639443415 | 1.39E-12 | 8.95E-12 |
| MDGA1 | 3.192767415 | 0.010840678 | 0.021085647 |
| MDH1 | -0.646564179 | 5.13E-08 | 2.15E-07 |
| MDH2 | -0.232673295 | 0.007873242 | 0.015788302 |
| MDM1 | -0.414689341 | 0.014748234 | 0.027864609 |
| MDM2 | -0.947112866 | 2.52E-20 | 2.89E-19 |
| MDM4 | 0.964772518 | 1.19E-11 | 7.08E-11 |
| ME1 | -0.696987293 | 4.28E-18 | 4.24E-17 |
| MEA1 | 0.886262989 | 8.99E-09 | 4.05E-08 |
| MECOM | 0.760800168 | 0.000807781 | 0.001954665 |
| MED10 | 0.389975634 | 0.000418879 | 0.001062201 |
| MED12L | 1.201925333 | 0.00012893 | 0.000352173 |
| MED14 | -0.995887622 | 8.49E-13 | 5.57E-12 |
| MED16 | -1.059396416 | 2.14E-10 | 1.12E-09 |
| MED17 | -1.142377646 | 7.98E-32 | 1.86E-30 |
| MED18 | -0.377502877 | 0.020152231 | 0.036910002 |
| MED19 | 1.071817207 | 5.06E-26 | 8.46E-25 |
| MED21 | 1.253168584 | 2.13E-36 | 6.52E-35 |
| MED22 | 0.461544342 | 0.003351516 | 0.007282783 |
| MED23 | -0.593426231 | 3.74E-08 | 1.59E-07 |
| MED24 | -0.813200733 | 6.41E-11 | 3.53E-10 |
| MED27 | -0.964536638 | 4.64E-13 | 3.14E-12 |
| MED28 | 0.754125767 | 1.95E-19 | 2.11E-18 |
| MED4 | -0.9652794 | 5.26E-09 | 2.42E-08 |
| MED7 | -1.141384702 | 1.45E-26 | 2.48E-25 |
| MED8 | -0.897811017 | 8.16E-13 | 5.36E-12 |
| MED9 | 1.131852171 | 8.13E-38 | 2.69E-36 |
| MEF2A | 0.35078599 | 0.000386717 | 0.000986349 |
| MEF2D | 0.67032904 | 1.13E-06 | 4.09E-06 |
| MEGF11 | 2.819735354 | 8.39E-36 | 2.44E-34 |
| MEGF9 | 1.922470028 | 2.23E-13 | 1.55E-12 |
| MEIS1 | 0.781504582 | 2.58E-09 | 1.22E-08 |
| MEIS2 | 1.0076339 | 0.001916395 | 0.004353608 |
| MELK | -1.293568736 | 0.019830199 | 0.036398659 |
| MELTF | 1.563385096 | 1.18E-06 | 4.27E-06 |
| MEMO1 | 1.400031104 | 1.75E-70 | 2.31E-68 |
| MEOX2 | -0.454731777 | 0.016113503 | 0.030171475 |
| MEPE | 3.347796352 | 0.002752211 | 0.006082611 |
| MERTK | 0.845395509 | 5.65E-17 | 5.16E-16 |
| MESDC2 | 0.409235509 | 1.36E-07 | 5.48E-07 |
| MET | -0.584803683 | 3.98E-07 | 1.52E-06 |
| METAP1 | -0.259657355 | 0.008610124 | 0.017089488 |
| METAP2 | 0.245242327 | 0.006091327 | 0.012500878 |
| METRNL | 0.514927332 | 2.23E-08 | 9.66E-08 |
| METTL11B | 1.115620134 | 0.018991316 | 0.035014917 |
| METTL14 | -0.295830331 | 0.001328593 | 0.003105991 |

| | | | |
|---|---|---|---|
| METTL15P1 | -0.402055105 | 0.021682697 | 0.039420588 |
| METTL16 | -0.590116761 | 1.51E-05 | 4.69E-05 |
| METTL18 | -0.581245646 | 6.43E-07 | 2.40E-06 |
| METTL21A | -0.769116362 | 4.76E-08 | 2.00E-07 |
| METTL21C | -0.753362277 | 0.013717467 | 0.026105826 |
| METTL22 | 1.218786732 | 5.30E-18 | 5.23E-17 |
| METTL25 | -0.762641546 | 0.001361864 | 0.003179944 |
| METTL3 | 1.82048456 | 4.24E-12 | 2.62E-11 |
| METTL4 | -1.219781331 | 6.11E-10 | 3.07E-09 |
| METTL6 | 1.013251597 | 4.27E-27 | 7.60E-26 |
| METTL7A | -0.389096763 | 0.002438354 | 0.005443841 |
| METTL9 | 0.919892652 | 5.63E-21 | 6.67E-20 |
| MFAP1 | -1.139421757 | 1.04E-21 | 1.29E-20 |
| MFAP3 | 0.306238683 | 0.000304661 | 0.000789504 |
| MFAP3L | 1.026853101 | 0.000395193 | 0.001006268 |
| MFAP5 | 0.665110505 | 6.93E-14 | 4.99E-13 |
| MFGE8 | -0.629582522 | 2.01E-13 | 1.40E-12 |
| MFN2 | -0.513853953 | 1.38E-05 | 4.32E-05 |
| MFSD1 | -0.454504054 | 3.83E-05 | 0.000113399 |
| MFSD11 | 0.70281397 | 9.81E-17 | 8.81E-16 |
| MFSD12 | -1.026444677 | 1.47E-08 | 6.51E-08 |
| MFSD13A | 0.281443383 | 0.017879804 | 0.03314543 |
| MFSD14A | -0.859958594 | 1.89E-21 | 2.31E-20 |
| MFSD5 | -0.491776105 | 0.001118534 | 0.002654638 |
| MFSD8 | -0.645478024 | 0.000172611 | 0.000463387 |
| MGAT2 | -0.660764023 | 7.33E-08 | 3.03E-07 |
| MGAT3 | 1.261015193 | 1.47E-25 | 2.39E-24 |
| MGAT4C | 1.696951891 | 3.69E-05 | 0.000109603 |
| MGLL | 1.431541772 | 6.83E-33 | 1.66E-31 |
| MGP | 0.550384394 | 0.02714384 | 0.048208424 |
| MGRN1 | -0.273684374 | 0.012757372 | 0.024491271 |
| MGST1 | 0.570574721 | 8.28E-08 | 3.40E-07 |
| MGST3 | 1.114394225 | 6.13E-34 | 1.59E-32 |
| MIA3 | -0.852497273 | 6.46E-17 | 5.88E-16 |
| MIB1 | -0.243381952 | 0.017330466 | 0.032219354 |
| MIB2 | -0.740406714 | 8.76E-08 | 3.59E-07 |
| MICAL1 | -0.709932796 | 0.006340444 | 0.012969046 |
| MICOS13 | 0.6751011 | 2.32E-08 | 1.01E-07 |
| MICU1 | -0.918233803 | 5.35E-12 | 3.26E-11 |
| MICU2 | -0.49657337 | 0.000513464 | 0.001283142 |
| MID2 | 1.510462176 | 9.94E-08 | 4.05E-07 |
| MIDN | 0.822354421 | 5.09E-05 | 0.00014821 |
| MIEF2 | -1.403932266 | 2.90E-11 | 1.65E-10 |
| MIEN1 | 0.858792326 | 2.06E-25 | 3.31E-24 |
| MIF | 0.919438199 | 2.90E-24 | 4.27E-23 |
| MIIP | -0.986587637 | 7.39E-07 | 2.74E-06 |
| MINA | -0.791311324 | 4.45E-08 | 1.88E-07 |
| MINDY2 | 0.261192605 | 0.007175808 | 0.014504364 |

| | | | |
|---|---|---|---|
| MINDY3 | 1.064824223 | 4.87E-30 | 1.03E-28 |
| MINOS1 | 1.36776997 | 1.53E-37 | 5.00E-36 |
| MINPP1 | -1.783639487 | 3.27E-20 | 3.73E-19 |
| MIOS | -0.499132563 | 0.000736355 | 0.001794932 |
| MIPEP | -0.789950112 | 9.07E-10 | 4.48E-09 |
| MIS18A | -0.713369389 | 0.000309537 | 0.000801071 |
| MIS18BP1 | -0.724815179 | 0.024235671 | 0.043607525 |
| MITF | -0.58782431 | 2.30E-06 | 8.04E-06 |
| MKKS | 0.575044233 | 3.08E-09 | 1.45E-08 |
| MKLN1 | -0.326251939 | 0.000910532 | 0.002187032 |
| MKRN2 | 0.432847042 | 6.29E-05 | 0.000180759 |
| MKX | 0.227703318 | 0.024694332 | 0.044344743 |
| MLEC | 0.408146556 | 6.10E-06 | 2.01E-05 |
| MLF2 | 0.562965371 | 1.69E-13 | 1.19E-12 |
| MLLT1 | -0.585756294 | 0.015814605 | 0.029661391 |
| MLLT6 | 1.39869686 | 1.14E-11 | 6.75E-11 |
| MLST8 | 0.991278082 | 6.52E-25 | 1.01E-23 |
| MLX | 1.12635689 | 5.10E-52 | 3.30E-50 |
| MLYCD | -1.035003741 | 5.32E-06 | 1.76E-05 |
| MMAA | -0.447351233 | 0.021496048 | 0.039123498 |
| MMACHC | 0.692134624 | 0.00017547 | 0.000470226 |
| MMADHC | 0.56806977 | 2.13E-11 | 1.23E-10 |
| MMD | 1.260988041 | 7.71E-61 | 7.03E-59 |
| MME | 1.711159605 | 6.12E-05 | 0.000176292 |
| MMGT1 | 1.06271385 | 3.96E-45 | 1.89E-43 |
| MMP1 | -4.503240268 | 3.54E-170 | 6.02E-167 |
| MMP15 | 0.746346668 | 4.74E-09 | 2.19E-08 |
| MMP2 | -0.485874013 | 9.48E-10 | 4.68E-09 |
| MMP23B | 0.895221743 | 3.27E-16 | 2.81E-15 |
| MMP24 | 1.033598896 | 9.52E-12 | 5.70E-11 |
| MMP27 | -0.643701167 | 0.000719697 | 0.001756845 |
| MMP28 | -0.607180445 | 0.004136213 | 0.008821497 |
| MMP9 | -2.13536006 | 1.13E-19 | 1.25E-18 |
| MMR1L3 | 2.316029669 | 0.007796452 | 0.015657413 |
| MMR1L4 | 3.058773679 | 0.000577001 | 0.00143061 |
| MMS19 | -0.230248994 | 0.011298263 | 0.021881731 |
| MMS22L | -0.840336827 | 0.023101378 | 0.041737861 |
| MMUT | -0.823397275 | 4.24E-06 | 1.42E-05 |
| MNAT1 | -0.964924697 | 8.71E-15 | 6.70E-14 |
| MNS1 | -0.948088186 | 0.000586213 | 0.001452124 |
| MNT | -1.665245562 | 8.98E-21 | 1.05E-19 |
| MOB1A | 1.100511035 | 7.46E-13 | 4.93E-12 |
| MOB2 | 0.597045214 | 4.76E-10 | 2.41E-09 |
| MOB3A | -0.431790971 | 0.004270441 | 0.00906935 |
| MOB3C | 0.770812181 | 0.001043076 | 0.00248775 |
| MOB4 | 0.421638378 | 5.70E-07 | 2.14E-06 |
| MOCS2 | 0.346010907 | 0.01564189 | 0.029385566 |
| MOCS3 | -0.747669615 | 8.41E-07 | 3.09E-06 |

| | | | |
|---|---|---|---|
| MOGS | -1.212319342 | 1.52E-05 | 4.73E-05 |
| MON2 | -0.962695456 | 1.93E-15 | 1.55E-14 |
| MORC2 | 1.066172183 | 1.76E-12 | 1.13E-11 |
| MORF4L2 | 0.419893418 | 3.23E-05 | 9.64E-05 |
| MORN2 | 1.077635003 | 0.000159633 | 0.000430417 |
| MORN4 | 0.963366706 | 1.78E-30 | 3.85E-29 |
| MOSPD1 | 0.485434998 | 8.90E-07 | 3.26E-06 |
| MPC1L | 1.306117909 | 1.10E-23 | 1.56E-22 |
| MPC2 | 0.621966113 | 5.74E-11 | 3.17E-10 |
| MPDZ | -0.49422999 | 0.019962441 | 0.036601849 |
| MPG | -0.693987428 | 8.86E-06 | 2.84E-05 |
| MPHOSPH6 | 0.384865238 | 0.00126777 | 0.002975572 |
| MPI | -1.012032679 | 2.23E-13 | 1.55E-12 |
| MPP1 | 0.332845474 | 0.004835895 | 0.010140511 |
| MPP5 | -0.856375595 | 9.57E-10 | 4.71E-09 |
| MPP7 | 0.880274915 | 0.001540061 | 0.003562374 |
| MPPED1 | 0.59728264 | 0.001777218 | 0.004065973 |
| MPRIP | 0.439204549 | 0.01131855 | 0.021914066 |
| MPV17 | 0.757115133 | 1.08E-08 | 4.85E-08 |
| MPV17L2 | 0.692099106 | 2.20E-08 | 9.57E-08 |
| MRAS | 1.469657411 | 1.40E-58 | 1.12E-56 |
| MRC2 | -1.028230075 | 1.32E-05 | 4.14E-05 |
| MRE11 | -0.371384116 | 0.015766336 | 0.029581605 |
| MREG | 1.33637835 | 1.06E-11 | 6.30E-11 |
| MRGBP | 0.563054868 | 6.25E-11 | 3.44E-10 |
| MRM1 | -1.967234654 | 2.15E-30 | 4.61E-29 |
| MRM2 | -0.367891634 | 0.015452279 | 0.02906551 |
| MRPL10 | -0.941697948 | 1.26E-16 | 1.13E-15 |
| MRPL12 | 0.404371737 | 0.00146392 | 0.003403044 |
| MRPL13 | 0.897260236 | 1.14E-15 | 9.32E-15 |
| MRPL16 | 0.261391077 | 0.020259753 | 0.03708195 |
| MRPL20 | 0.573598189 | 8.11E-07 | 2.99E-06 |
| MRPL21 | 0.60847241 | 3.65E-06 | 1.23E-05 |
| MRPL22 | -0.626043524 | 1.81E-09 | 8.73E-09 |
| MRPL23 | 0.383941087 | 0.001129332 | 0.002677461 |
| MRPL24 | 0.634357671 | 3.64E-10 | 1.86E-09 |
| MRPL27 | 0.33998569 | 0.004797775 | 0.010071251 |
| MRPL3 | 0.266943801 | 0.014121007 | 0.026798761 |
| MRPL33 | 1.144617694 | 7.62E-30 | 1.59E-28 |
| MRPL37 | -1.076067741 | 2.04E-12 | 1.30E-11 |
| MRPL38 | -0.373638595 | 0.000347117 | 0.000892043 |
| MRPL39 | -1.205644773 | 3.69E-24 | 5.37E-23 |
| MRPL4 | 0.96318618 | 0.000142039 | 0.000385502 |
| MRPL40 | 0.263743333 | 0.017491427 | 0.032478617 |
| MRPL41 | -0.60449829 | 6.53E-06 | 2.14E-05 |
| MRPL43 | 0.690186505 | 1.31E-09 | 6.37E-09 |
| MRPL46 | -0.637755795 | 1.68E-08 | 7.40E-08 |
| MRPL51 | 0.43428574 | 0.001273194 | 0.002985722 |

| | | | |
|---|---|---|---|
| MRPL54 | 0.600580731 | 6.77E-06 | 2.22E-05 |
| MRPL9 | 0.291301848 | 0.005082718 | 0.010612029 |
| MRPS11 | 0.473165331 | 2.54E-05 | 7.67E-05 |
| MRPS14 | 1.347020963 | 1.04E-10 | 5.61E-10 |
| MRPS16 | 0.402400259 | 8.85E-05 | 0.00024844 |
| MRPS17 | 0.427370465 | 6.99E-05 | 0.000199232 |
| MRPS18A | 0.54116703 | 1.71E-07 | 6.79E-07 |
| MRPS18C | 0.323781515 | 0.011978453 | 0.023097054 |
| MRPS2 | 0.703582973 | 1.77E-11 | 1.03E-10 |
| MRPS21 | 0.873434048 | 7.76E-10 | 3.86E-09 |
| MRPS22 | -0.511810406 | 1.35E-05 | 4.22E-05 |
| MRPS24 | 0.720018738 | 2.60E-06 | 9.03E-06 |
| MRPS25 | 0.561628198 | 2.22E-08 | 9.65E-08 |
| MRPS26 | 0.423007 | 1.84E-06 | 6.53E-06 |
| MRPS27 | -0.33096381 | 0.016370078 | 0.030597098 |
| MRPS30 | -0.844133465 | 9.98E-13 | 6.51E-12 |
| MRPS31 | 0.685013252 | 2.06E-09 | 9.88E-09 |
| MRPS33 | 0.558953399 | 2.35E-08 | 1.02E-07 |
| MRPS34 | 0.397759595 | 7.57E-05 | 0.000214623 |
| MRPS35 | -1.154828129 | 6.40E-22 | 8.11E-21 |
| MRPS36 | 0.935645358 | 1.95E-14 | 1.47E-13 |
| MRPS5 | -0.249093123 | 0.024838059 | 0.044549869 |
| MRPS6 | 0.759508628 | 2.80E-13 | 1.93E-12 |
| MRS2 | -0.445326281 | 0.001935035 | 0.004393018 |
| MRTFA | -0.331713651 | 0.001939574 | 0.004402587 |
| MRTO4 | 0.489217661 | 4.05E-07 | 1.55E-06 |
| MSGN1 | 4.763455922 | 0.005472895 | 0.011353417 |
| MSH2 | -1.193734782 | 3.07E-16 | 2.65E-15 |
| MSH6 | -1.188478369 | 2.06E-12 | 1.31E-11 |
| MSI2 | 0.784338962 | 0.000463243 | 0.001166216 |
| MSL2 | 0.817084057 | 0.009007389 | 0.017802593 |
| MSL3 | -0.329347453 | 0.006194416 | 0.012690292 |
| MSN | -0.898476364 | 2.97E-22 | 3.86E-21 |
| MSRA | 0.504814055 | 0.000900353 | 0.00216403 |
| MSRB3 | 1.62432708 | 2.11E-92 | 6.68E-90 |
| MST1 | -1.120796966 | 0.002400298 | 0.005366812 |
| MST1R | -0.474929883 | 0.009906547 | 0.019421576 |
| MSTO1 | -0.438946401 | 0.013483667 | 0.025731016 |
| MSX1 | 0.243302532 | 0.006898258 | 0.014000466 |
| MT3 | 1.182730765 | 2.37E-55 | 1.70E-53 |
| MT4 | 0.824652548 | 8.91E-19 | 9.30E-18 |
| MTA3 | -1.340548369 | 7.76E-20 | 8.63E-19 |
| MTAP | 0.529171841 | 0.000766526 | 0.001862798 |
| MTCH2 | -0.497616785 | 3.79E-09 | 1.77E-08 |
| MTERF2 | 1.333489207 | 2.08E-06 | 7.33E-06 |
| MTERF3 | -0.806593619 | 1.22E-10 | 6.57E-10 |
| MTF1 | 0.389291452 | 3.91E-05 | 0.000115499 |
| MTFMT | -0.718804291 | 3.42E-08 | 1.46E-07 |

| | | | |
|---|---|---|---|
| MTFP1 | 1.133098461 | 2.03E-16 | 1.79E-15 |
| MTFR1 | 1.113613736 | 1.55E-37 | 5.05E-36 |
| MTFR1L | 0.69204389 | 4.92E-16 | 4.18E-15 |
| MTFR2 | -0.978130178 | 7.70E-06 | 2.50E-05 |
| MTHFD1 | -1.015576757 | 1.10E-19 | 1.22E-18 |
| MTHFD1L | -0.967950486 | 6.83E-11 | 3.75E-10 |
| MTHFD2 | -0.801903316 | 1.93E-21 | 2.36E-20 |
| MTHFD2L | 0.386230607 | 0.024923141 | 0.044661217 |
| MTHFR | 1.043565598 | 3.11E-26 | 5.25E-25 |
| MTHFS | -0.503379251 | 0.019750582 | 0.036272113 |
| MTIF2 | 0.339981673 | 0.004315855 | 0.009145791 |
| MTM1 | -0.615359387 | 3.18E-06 | 1.09E-05 |
| MTMR1 | -0.396634511 | 0.002191412 | 0.004934204 |
| MTMR10 | 0.557265729 | 0.000108441 | 0.000299946 |
| MTMR12 | -0.993960962 | 8.83E-05 | 0.000247987 |
| MTMR14 | 0.238418477 | 0.025788981 | 0.046030706 |
| MTMR3 | -0.83590284 | 7.25E-08 | 3.00E-07 |
| MTMR4 | -0.785312029 | 2.88E-06 | 9.92E-06 |
| MTMR6 | -1.072660204 | 1.45E-19 | 1.59E-18 |
| MTMR7 | 0.731301624 | 0.015080355 | 0.028432818 |
| MTNR1B | 1.887065474 | 1.16E-08 | 5.17E-08 |
| MTOR | -0.914414257 | 2.51E-06 | 8.74E-06 |
| MTPAP | 0.289210815 | 0.002135698 | 0.004815594 |
| MTPN | 1.140645935 | 3.90E-52 | 2.55E-50 |
| MTRF1L | -0.79854754 | 5.24E-05 | 0.000152418 |
| MTRR | -0.379966833 | 0.01269503 | 0.024378474 |
| MTSS2 | 0.671962257 | 0.018350425 | 0.033929938 |
| MTTP | 5.176627315 | 7.90E-09 | 3.58E-08 |
| MTUS1 | -0.640443053 | 2.72E-06 | 9.42E-06 |
| MTX2 | 0.384574631 | 0.001739629 | 0.003985342 |
| MUL1 | -0.613974 | 1.33E-05 | 4.16E-05 |
| MUM1L1 | -0.939599879 | 3.66E-07 | 1.40E-06 |
| MVB12B | -0.456957309 | 0.014270554 | 0.027044809 |
| MVD | -0.381101555 | 0.005569894 | 0.011530003 |
| MVK | -0.923237736 | 1.40E-11 | 8.22E-11 |
| MVP | -1.299224348 | 2.90E-37 | 9.25E-36 |
| MX1 | 1.865048557 | 0.010040512 | 0.019647372 |
| MXD1 | -0.536361778 | 1.42E-05 | 4.43E-05 |
| MXD4 | 0.962962736 | 3.26E-30 | 6.95E-29 |
| MYBBP1A | -0.967551435 | 1.06E-13 | 7.51E-13 |
| MYBL2 | 1.184574388 | 5.44E-23 | 7.35E-22 |
| MYBPC1 | 1.936933736 | 0.000146021 | 0.000395834 |
| MYBPC3 | 2.093936496 | 7.69E-18 | 7.48E-17 |
| MYC | -0.572605226 | 2.95E-10 | 1.53E-09 |
| MYCBP2 | -0.520943943 | 0.00303806 | 0.006668817 |
| MYCL | -0.942133139 | 0.00373227 | 0.008031826 |
| MYD88 | 0.498599518 | 4.74E-08 | 1.99E-07 |
| MYDGF | 0.404224859 | 0.000161732 | 0.000435385 |

| | | | |
|---|---|---|---|
| MYF6 | 4.960480488 | 0.005953137 | 0.012245011 |
| MYG1 | -1.29420151 | 1.87E-24 | 2.79E-23 |
| MYH10 | -1.249615611 | 3.39E-30 | 7.22E-29 |
| MYH15 | -1.528740924 | 1.36E-36 | 4.22E-35 |
| MYH7 | 3.640679887 | 0.001480117 | 0.003437755 |
| MYH9 | -1.173993526 | 4.58E-41 | 1.75E-39 |
| MYL12A | 1.181379498 | 1.68E-48 | 9.49E-47 |
| MYL12B | 1.03541532 | 1.93E-35 | 5.49E-34 |
| MYL3 | 0.966312809 | 0.005378161 | 0.011172239 |
| MYL9 | 1.61968561 | 2.02E-60 | 1.77E-58 |
| MYLIP | -0.370041033 | 0.000201304 | 0.000535027 |
| MYLK | -0.481016129 | 6.10E-06 | 2.01E-05 |
| MYO18B | 4.772972544 | 0.000621823 | 0.001532791 |
| MYO1C | -0.185750725 | 0.023807757 | 0.042900043 |
| MYO1E | 0.232407778 | 0.006878798 | 0.013969314 |
| MYO1G | 6.791878187 | 5.34E-06 | 1.77E-05 |
| MYO1H | 1.567488025 | 0.000230653 | 0.000608507 |
| MYO3B | 2.740801576 | 0.007927863 | 0.015886002 |
| MYO5A | 0.3657424 | 0.004729783 | 0.009943489 |
| MYO5C | 1.124645744 | 0.002573841 | 0.005727513 |
| MYO6 | -0.379670598 | 0.000448562 | 0.001132613 |
| MYO9A | 0.800546983 | 2.08E-05 | 6.35E-05 |
| MYOD1 | 2.151386588 | 1.35E-07 | 5.42E-07 |
| MYOM2 | 1.915304843 | 0.027355019 | 0.048539088 |
| MYOM3 | -0.477090841 | 0.017918653 | 0.033203863 |
| MYSM1 | -0.502745127 | 0.010797646 | 0.02100796 |
| MZT1 | 0.568329232 | 6.92E-07 | 2.57E-06 |
| N4BP1 | -0.873543204 | 1.54E-17 | 1.47E-16 |
| N4BP2L2 | -1.566586736 | 3.24E-26 | 5.46E-25 |
| N4BP3 | -1.268934347 | 1.70E-18 | 1.75E-17 |
| N6AMT1 | 0.479369793 | 1.03E-05 | 3.29E-05 |
| NAA20 | 1.100681982 | 2.56E-27 | 4.63E-26 |
| NAA25 | -0.675331432 | 1.59E-09 | 7.69E-09 |
| NAA40 | 0.365624893 | 0.000216153 | 0.000572141 |
| NAA50 | 0.201249863 | 0.015309505 | 0.028820886 |
| NAB1 | 0.634157595 | 1.47E-05 | 4.58E-05 |
| NABP1 | -0.907067485 | 4.04E-20 | 4.57E-19 |
| NAE1 | -0.525460975 | 4.86E-06 | 1.62E-05 |
| NAF1 | -1.021551188 | 1.02E-14 | 7.80E-14 |
| NAGA | -1.396338729 | 2.06E-32 | 4.91E-31 |
| NAGLU | -2.462981942 | 1.79E-29 | 3.65E-28 |
| NAIF1 | 2.03154918 | 1.66E-99 | 5.78E-97 |
| NAMPTP1 | -0.486512926 | 1.97E-06 | 6.96E-06 |
| NANOS1 | -0.911847269 | 9.41E-16 | 7.79E-15 |
| NAP1L4 | 0.317460303 | 0.001467675 | 0.003410606 |
| NAPG | -0.484641542 | 3.07E-07 | 1.19E-06 |
| NARFL | -0.870891038 | 1.87E-11 | 1.08E-10 |
| NARS | 0.459816136 | 4.73E-08 | 1.99E-07 |

| | | | |
|---|---|---|---|
| NARS2 | -0.560192039 | 0.000512201 | 0.001280456 |
| NASP | -0.921281863 | 6.22E-11 | 3.43E-10 |
| NAT | 1.5916708 | 2.84E-06 | 9.79E-06 |
| NAT10 | -0.79768333 | 8.91E-10 | 4.41E-09 |
| NAT9 | 0.616641755 | 4.71E-10 | 2.38E-09 |
| NATD1 | 1.005163716 | 5.26E-19 | 5.57E-18 |
| NAV3 | -0.645982806 | 1.76E-08 | 7.70E-08 |
| NBEAL2 | 1.164359858 | 0.002235559 | 0.005020774 |
| NBL1 | 1.687240854 | 1.22E-16 | 1.09E-15 |
| NBR1 | -0.623431832 | 0.000858759 | 0.002070649 |
| NCALD | 0.797000147 | 0.007060377 | 0.014291086 |
| NCAM1 | 0.674833831 | 0.000729394 | 0.001778601 |
| NCAPD2 | -1.725105168 | 2.12E-20 | 2.44E-19 |
| NCAPD3 | -1.087490128 | 0.000875618 | 0.002106816 |
| NCAPG | -1.386387643 | 0.000618086 | 0.001524688 |
| NCAPG2 | -0.953283576 | 0.007674776 | 0.015433578 |
| NCAPH | -0.542994788 | 0.015403204 | 0.028983335 |
| NCBP1 | -0.536840777 | 5.01E-09 | 2.31E-08 |
| NCBP2 | -0.211882069 | 0.011618415 | 0.022456982 |
| NCBP3 | -0.871439035 | 5.13E-07 | 1.94E-06 |
| NCDN | -1.187975553 | 1.16E-21 | 1.44E-20 |
| NCF1C | 0.655643333 | 0.012541131 | 0.024110191 |
| NCF2 | 2.017075444 | 2.07E-06 | 7.28E-06 |
| NCK1 | -0.25443304 | 0.004949082 | 0.010360068 |
| NCKAP1 | -0.17293645 | 0.022903779 | 0.041413884 |
| NCL | -0.460863782 | 2.16E-07 | 8.48E-07 |
| NCLN | -1.026138044 | 2.45E-23 | 3.38E-22 |
| NCOA1 | -0.67447178 | 0.018362032 | 0.03393755 |
| NCOA2 | -1.146960005 | 0.000264278 | 0.000690916 |
| NCOA3 | -0.71887573 | 3.06E-05 | 9.16E-05 |
| NCOA4 | -0.620207065 | 4.18E-10 | 2.13E-09 |
| NCOA6 | -0.296218669 | 0.024210417 | 0.043573621 |
| NCOA7 | 0.533967754 | 8.81E-06 | 2.83E-05 |
| NCS1 | 1.863721645 | 1.16E-82 | 2.63E-80 |
| NCSTN | -1.049373714 | 2.37E-19 | 2.56E-18 |
| NDC1 | -0.681361495 | 9.01E-09 | 4.06E-08 |
| NDC80 | -1.19166968 | 4.72E-06 | 1.57E-05 |
| NDEL1 | -0.333421017 | 4.02E-05 | 0.000118555 |
| NDFIP1 | 1.09340018 | 1.93E-28 | 3.72E-27 |
| NDNF | -1.125139234 | 1.76E-22 | 2.33E-21 |
| NDOR1 | 0.859400905 | 1.98E-22 | 2.62E-21 |
| NDP | 0.88879816 | 4.92E-06 | 1.64E-05 |
| NDRG1 | -0.857760688 | 7.93E-22 | 9.92E-21 |
| NDRG3 | 0.453602811 | 2.70E-07 | 1.05E-06 |
| NDST1 | 0.597350769 | 2.40E-05 | 7.28E-05 |
| NDUFA1 | 1.359521275 | 2.81E-29 | 5.70E-28 |
| NDUFA10 | 0.799613402 | 5.03E-22 | 6.41E-21 |
| NDUFA12 | 0.807511669 | 6.20E-13 | 4.12E-12 |

| | | | |
|---|---|---|---|
| NDUFA13 | 0.683311968 | 2.01E-11 | 1.16E-10 |
| NDUFA4 | 1.04731017 | 2.09E-21 | 2.54E-20 |
| NDUFA9 | -0.8137821 | 1.97E-09 | 9.47E-09 |
| NDUFAB1 | 0.510439263 | 4.50E-08 | 1.90E-07 |
| NDUFAF1 | 0.623577504 | 6.09E-14 | 4.41E-13 |
| NDUFAF7 | -0.539107731 | 1.56E-05 | 4.85E-05 |
| NDUFAF8 | 0.734490089 | 3.71E-06 | 1.25E-05 |
| NDUFB1 | 0.619283999 | 2.98E-07 | 1.15E-06 |
| NDUFB10 | 1.06223863 | 5.40E-30 | 1.13E-28 |
| NDUFB2 | 0.991077456 | 8.27E-19 | 8.66E-18 |
| NDUFB3 | 1.154454297 | 1.62E-18 | 1.67E-17 |
| NDUFB4 | 1.221856917 | 2.62E-28 | 5.00E-27 |
| NDUFB5 | 0.920163014 | 3.13E-22 | 4.05E-21 |
| NDUFB8 | 0.643138038 | 3.75E-12 | 2.33E-11 |
| NDUFS1 | -1.157789012 | 1.14E-25 | 1.87E-24 |
| NDUFS2 | -0.331381323 | 0.00016111 | 0.000433882 |
| NDUFS3 | 0.51662172 | 1.18E-08 | 5.29E-08 |
| NDUFS4 | 0.459971867 | 0.000374775 | 0.000957867 |
| NDUFS5 | 1.081315513 | 5.31E-21 | 6.30E-20 |
| NDUFS6 | 0.987261988 | 1.02E-11 | 6.10E-11 |
| NDUFS7 | 1.013116551 | 5.31E-27 | 9.36E-26 |
| NDUFS8 | 0.291192711 | 0.00424077 | 0.00901056 |
| NDUFV2 | -0.424910501 | 0.00058543 | 0.001450449 |
| NDUFV3 | 0.449899474 | 0.000132355 | 0.000360803 |
| NECAB2 | -1.417779268 | 2.04E-36 | 6.24E-35 |
| NECAP1 | 0.551716642 | 3.61E-10 | 1.85E-09 |
| NECTIN3 | -1.574077946 | 1.76E-36 | 5.40E-35 |
| NEDD8 | 1.006707637 | 2.92E-06 | 1.01E-05 |
| NEFH | -2.908384647 | 0.000404947 | 0.001029563 |
| NEGR1 | 0.736682769 | 0.000211008 | 0.000559287 |
| NEIL1 | -0.593783182 | 0.003077587 | 0.006743601 |
| NEK2 | -1.06041488 | 0.000979074 | 0.002344534 |
| NEK3 | -1.053862569 | 1.15E-05 | 3.63E-05 |
| NEK4 | 0.836927225 | 3.15E-15 | 2.50E-14 |
| NEK6 | 0.591171818 | 2.62E-14 | 1.95E-13 |
| NELFB | -0.383537121 | 0.000127118 | 0.000347853 |
| NELFCD | 0.545302978 | 9.42E-11 | 5.11E-10 |
| NELFE | -0.878044862 | 8.88E-17 | 8.01E-16 |
| NELL1 | -0.902125176 | 1.99E-10 | 1.05E-09 |
| NELL2 | -2.107056298 | 0.001239054 | 0.00291825 |
| NEMF | 0.269955038 | 0.005916175 | 0.012181708 |
| NEMP1 | -0.743381995 | 0.000586351 | 0.001452202 |
| NEMP2 | 1.241629128 | 3.12E-25 | 4.94E-24 |
| NENF | 0.967620604 | 2.08E-14 | 1.57E-13 |
| NEO1 | -0.688545064 | 4.78E-07 | 1.81E-06 |
| NEPRO | -1.266250921 | 7.51E-16 | 6.28E-15 |
| NET1 | -1.546612969 | 9.73E-57 | 7.34E-55 |
| NEURL1 | 0.587290537 | 0.001666502 | 0.003832672 |

| | | | |
|---|---|---|---|
| NEUROD1 | -1.665151899 | 8.88E-05 | 0.000249088 |
| NF2 | 1.392058429 | 1.60E-73 | 2.45E-71 |
| NFATC1 | -0.450149806 | 1.31E-05 | 4.12E-05 |
| NFATC2 | -0.412761076 | 0.024989445 | 0.044774128 |
| NFE2L2 | 0.227024616 | 0.007399553 | 0.014911005 |
| NFIA | 1.356640399 | 1.72E-24 | 2.57E-23 |
| NFIB | 0.966912876 | 0.007840474 | 0.015734199 |
| NFIC | 0.459518334 | 0.009263881 | 0.018253794 |
| NFIL3 | -1.871115886 | 7.45E-69 | 9.38E-67 |
| NFIX | 1.092799673 | 5.42E-13 | 3.63E-12 |
| NFKB1 | -0.478390716 | 1.42E-05 | 4.42E-05 |
| NFKBIA | 2.604179722 | 6.52E-110 | 2.95E-107 |
| NFKBIZ | -1.76565785 | 8.13E-46 | 4.00E-44 |
| NFRKB | -0.770239389 | 3.25E-08 | 1.38E-07 |
| NFS1 | -0.240918172 | 0.007025638 | 0.01423561 |
| NFXL1 | -0.852363194 | 2.07E-06 | 7.29E-06 |
| NFYA | -0.712168166 | 2.96E-06 | 1.02E-05 |
| NFYB | -0.754942149 | 1.87E-11 | 1.08E-10 |
| NFYC | 1.225406835 | 4.93E-46 | 2.47E-44 |
| NGLY1 | -1.589285679 | 2.22E-44 | 1.02E-42 |
| NHEJ1 | -0.770710353 | 0.012124352 | 0.023361807 |
| NHLRC2 | 0.319954992 | 0.008038695 | 0.016086859 |
| NIBAN2 | 0.383541303 | 3.61E-05 | 0.000107302 |
| NICN1 | 0.990597879 | 7.05E-06 | 2.30E-05 |
| NID1 | -2.058275068 | 0.008109004 | 0.016206086 |
| NIF3L1 | -0.553740487 | 2.11E-05 | 6.43E-05 |
| NIFK | -0.628915267 | 3.79E-09 | 1.77E-08 |
| NIN | -1.279691962 | 3.27E-15 | 2.60E-14 |
| NINJ1 | 1.104068615 | 3.16E-45 | 1.52E-43 |
| NINJ2 | 1.16979816 | 5.33E-43 | 2.27E-41 |
| NIP7 | -0.66326189 | 1.67E-07 | 6.66E-07 |
| NIPA2 | 0.500162299 | 1.09E-08 | 4.87E-08 |
| NIPAL3 | 0.515483989 | 3.27E-10 | 1.69E-09 |
| NIPSNAP1 | -0.442279828 | 0.001587813 | 0.003664102 |
| NIPSNAP2 | 0.345910179 | 0.002926052 | 0.006442717 |
| NIT1 | -0.790174398 | 2.48E-08 | 1.07E-07 |
| NKAPL | 0.668967285 | 1.77E-07 | 7.01E-07 |
| NKIRAS1 | 0.521777594 | 0.003819216 | 0.00819687 |
| NKIRAS2 | 0.274530852 | 0.005346063 | 0.011112356 |
| NKRF | -0.387764966 | 0.000157862 | 0.000426054 |
| NKX2-5 | 7.142336498 | 2.49E-07 | 9.72E-07 |
| NKX3-2 | 1.185470894 | 1.43E-18 | 1.47E-17 |
| NLGN4Y | 1.751546051 | 1.73E-10 | 9.15E-10 |
| NLRC5 | 0.891579636 | 0.000141016 | 0.000382802 |
| NMB | -1.867475273 | 4.65E-06 | 1.55E-05 |
| NMD3 | -0.57411622 | 2.09E-08 | 9.09E-08 |
| NME1 | -1.127897971 | 0.000540168 | 0.001343943 |
| NME2 | 0.434777008 | 2.63E-05 | 7.95E-05 |

| | | | |
|---|---|---|---|
| NME3 | -0.421215983 | 0.001302751 | 0.003049249 |
| NME5 | 1.492663469 | 9.10E-05 | 0.000254734 |
| NME6 | -0.524172995 | 0.009658966 | 0.018974514 |
| NMNAT2 | 1.931889059 | 0.022942093 | 0.041466614 |
| NMRAL1 | -0.731465045 | 4.65E-12 | 2.86E-11 |
| NMRK1 | 0.877177416 | 2.07E-15 | 1.66E-14 |
| NMT1 | -0.42454746 | 9.18E-06 | 2.94E-05 |
| NNF1 | 0.513572305 | 0.011970687 | 0.023085355 |
| NNT | -0.898389522 | 4.36E-09 | 2.02E-08 |
| NOB1 | -0.815084009 | 1.76E-15 | 1.42E-14 |
| NOC2L | -0.900077203 | 5.22E-17 | 4.77E-16 |
| NOC3L | -1.009035547 | 1.16E-29 | 2.39E-28 |
| NOC4L | -0.774133271 | 2.74E-15 | 2.19E-14 |
| NOCT | -1.3812154 | 1.21E-24 | 1.83E-23 |
| NOG | -2.803711832 | 5.17E-06 | 1.72E-05 |
| NOG2 | 1.167226603 | 0.001932283 | 0.004387502 |
| NOL10 | -0.900047429 | 4.92E-16 | 4.18E-15 |
| NOL6 | -0.862608696 | 6.59E-16 | 5.53E-15 |
| NOL7 | -0.464780784 | 6.14E-05 | 0.000176522 |
| NOL8 | -0.331693036 | 0.003511582 | 0.007599016 |
| NOLC1 | -0.776335737 | 5.43E-10 | 2.74E-09 |
| NOM1 | -0.86440162 | 2.89E-10 | 1.50E-09 |
| NONO | -0.536699103 | 5.43E-11 | 3.01E-10 |
| NOP14 | -0.688551302 | 1.02E-07 | 4.16E-07 |
| NOP16 | 0.275482047 | 0.009149509 | 0.018054606 |
| NOP2 | -0.342032768 | 0.000343049 | 0.000882758 |
| NOP53 | -0.361733008 | 0.000345627 | 0.000888885 |
| NOP56 | -0.272511946 | 0.000730687 | 0.001781435 |
| NOTCH2 | -1.25368456 | 1.78E-30 | 3.84E-29 |
| NOV | -1.485079308 | 2.17E-54 | 1.52E-52 |
| NOVA1 | -1.213786842 | 2.51E-10 | 1.31E-09 |
| NOX1 | 2.657238372 | 0.000166288 | 0.000447119 |
| NOX4 | 0.823748271 | 2.17E-11 | 1.25E-10 |
| NOXO1 | -0.860464224 | 1.47E-08 | 6.50E-08 |
| NPAS2 | 0.496017677 | 0.008105717 | 0.016201899 |
| NPC1 | -1.271811797 | 3.20E-23 | 4.38E-22 |
| NPC2 | 0.328741974 | 5.58E-05 | 0.000161547 |
| NPEPL1 | -0.467112982 | 3.88E-07 | 1.49E-06 |
| NPHP1 | 0.22621089 | 0.01913871 | 0.035248421 |
| NPM2 | 1.321222884 | 3.96E-13 | 2.69E-12 |
| NPM3 | 0.713230875 | 1.38E-17 | 1.33E-16 |
| NPNT | -0.720395285 | 1.17E-10 | 6.30E-10 |
| NPPA | 1.805048706 | 4.53E-33 | 1.11E-31 |
| NPPC | 4.337941044 | 0.001846131 | 0.004208757 |
| NPRL3 | -0.480669523 | 0.005636161 | 0.01165121 |
| NPTX2 | -1.527684089 | 8.31E-29 | 1.63E-27 |
| NPY | -1.343999479 | 3.93E-15 | 3.11E-14 |
| NQO2 | 0.63652231 | 8.26E-10 | 4.10E-09 |

| | | | |
|---|---|---|---|
| NR1D2 | -0.723962206 | 3.55E-11 | 2.00E-10 |
| NR1H3 | 0.288542925 | 0.000323628 | 0.000834995 |
| NR2C1 | -0.640933999 | 3.17E-06 | 1.08E-05 |
| NR2C2 | 0.954181848 | 9.65E-12 | 5.77E-11 |
| NR2F2 | 0.815211167 | 3.32E-17 | 3.11E-16 |
| NR4A2 | -1.862393241 | 3.06E-36 | 9.21E-35 |
| NR4A3 | 0.334909699 | 0.000772033 | 0.001875509 |
| NRAP | 2.73733906 | 1.05E-20 | 1.22E-19 |
| NRBF2 | -0.635165481 | 3.00E-07 | 1.16E-06 |
| NRBP1 | -0.630138758 | 1.13E-12 | 7.36E-12 |
| NRDC | -0.576115077 | 1.04E-09 | 5.09E-09 |
| NREP | 1.303723506 | 3.90E-29 | 7.80E-28 |
| NRG4 | 1.353966503 | 7.18E-06 | 2.34E-05 |
| NRIP1 | -0.405488455 | 0.000159846 | 0.00043082 |
| NRIP3 | 2.292992364 | 7.91E-11 | 4.33E-10 |
| NRK | 1.047233248 | 2.38E-15 | 1.91E-14 |
| NRP1 | -0.376159694 | 0.000267741 | 0.000699162 |
| NRP2 | -0.377088705 | 2.84E-06 | 9.79E-06 |
| NRSN1 | 0.758751328 | 1.75E-19 | 1.92E-18 |
| NRXN1 | 1.316944718 | 0.006039998 | 0.012406779 |
| NRXN3 | 0.853905648 | 4.06E-13 | 2.76E-12 |
| NSA2 | 0.723809817 | 1.84E-12 | 1.17E-11 |
| NSD3 | 0.650586412 | 4.00E-10 | 2.04E-09 |
| NSF | -0.823322168 | 2.67E-16 | 2.31E-15 |
| NSG1 | -0.43172538 | 0.005916983 | 0.012181708 |
| NSL1 | 0.586978404 | 2.16E-06 | 7.58E-06 |
| NSMCE1 | -1.029159452 | 1.15E-11 | 6.80E-11 |
| NSMCE2 | 0.525972095 | 7.15E-06 | 2.33E-05 |
| NSMCE3 | 0.362299357 | 0.001308405 | 0.003060903 |
| NSMF | 0.752305216 | 7.30E-09 | 3.32E-08 |
| NSUN2 | -0.494601868 | 1.98E-06 | 7.00E-06 |
| NSUN4 | -0.776249828 | 3.93E-08 | 1.67E-07 |
| NSUN5 | -0.69757431 | 5.58E-10 | 2.82E-09 |
| NT5C2 | -0.266844307 | 0.004410734 | 0.009326493 |
| NT5C3B | 0.446045047 | 0.000699734 | 0.001711187 |
| NT5DC2 | 1.400721683 | 5.32E-72 | 7.54E-70 |
| NT5DC3 | -0.967443216 | 7.21E-06 | 2.35E-05 |
| NT5DC4 | 2.83576722 | 0.000112343 | 0.000309917 |
| NTAN1 | 0.636944909 | 1.30E-07 | 5.24E-07 |
| NTAQ1 | -0.543162161 | 0.000741173 | 0.001805057 |
| NTF3 | -1.167886005 | 1.47E-06 | 5.26E-06 |
| NTHL1 | -0.735903445 | 1.77E-07 | 7.03E-07 |
| NTN4 | 0.554699532 | 5.44E-12 | 3.31E-11 |
| NTNG2 | 1.80507661 | 1.63E-15 | 1.33E-14 |
| NTS | -0.741397611 | 0.009324243 | 0.018364745 |
| NUAK1 | 0.244658206 | 0.002430932 | 0.005429055 |
| NUAK2 | 2.414807375 | 2.22E-06 | 7.76E-06 |
| NUB1 | 0.369038375 | 0.000283265 | 0.000737009 |

| | | | |
|---|---|---|---|
| NUBP2 | -0.338201076 | 0.000426798 | 0.001081072 |
| NUBPL | 0.954324828 | 4.87E-20 | 5.49E-19 |
| NUCB2 | -0.307626941 | 0.001486853 | 0.003451632 |
| NUCKS1 | 0.481728755 | 1.93E-06 | 6.83E-06 |
| NUDCD2 | 0.569601065 | 1.27E-12 | 8.19E-12 |
| NUDCD3 | -0.412746008 | 7.83E-06 | 2.54E-05 |
| NUDT13 | -0.726626436 | 0.011114261 | 0.021556084 |
| NUDT14 | 0.373099104 | 0.002410812 | 0.005387661 |
| NUDT16L1 | -0.555350102 | 7.05E-09 | 3.21E-08 |
| NUDT21 | 0.399596401 | 4.51E-06 | 1.51E-05 |
| NUDT3 | 0.999559859 | 0.001409626 | 0.003283008 |
| NUDT5 | 0.856068298 | 1.21E-07 | 4.88E-07 |
| NUDT6 | 0.193483317 | 0.014099299 | 0.0267613 |
| NUDT9 | -0.476981959 | 0.001634632 | 0.003763835 |
| NUF2 | -1.142555751 | 1.45E-08 | 6.43E-08 |
| NUMA1 | 0.489829285 | 3.55E-05 | 0.00010559 |
| NUMB | -1.014358674 | 2.24E-21 | 2.72E-20 |
| NUP107 | -0.356205068 | 0.00082613 | 0.001996579 |
| NUP133 | -1.442138465 | 7.46E-27 | 1.30E-25 |
| NUP153 | -0.710685365 | 1.02E-06 | 3.71E-06 |
| NUP155 | -1.307154805 | 7.16E-14 | 5.15E-13 |
| NUP188 | -0.776482276 | 7.57E-10 | 3.77E-09 |
| NUP205 | -0.852634737 | 1.10E-07 | 4.45E-07 |
| NUP214 | -1.14904514 | 7.68E-15 | 5.94E-14 |
| NUP35 | -0.797989096 | 4.29E-10 | 2.18E-09 |
| NUP37 | -0.285535983 | 0.004510497 | 0.00951967 |
| NUP42 | -0.291262619 | 0.004514347 | 0.009526317 |
| NUP43 | 0.242599136 | 0.021654277 | 0.039374573 |
| NUP54 | -0.448100422 | 2.14E-05 | 6.55E-05 |
| NUP62 | -0.479429511 | 3.72E-05 | 0.000110364 |
| NUP85 | -0.378849092 | 7.51E-05 | 0.000213052 |
| NUP88 | -0.64479352 | 7.99E-11 | 4.37E-10 |
| NUP93 | 0.731218156 | 5.92E-14 | 4.29E-13 |
| NUP98 | -0.838502412 | 2.45E-15 | 1.96E-14 |
| NUS1 | 0.435987916 | 5.25E-06 | 1.74E-05 |
| NUSAP1 | -0.790932878 | 0.006131031 | 0.012569075 |
| NVL | -0.510503878 | 0.000553723 | 0.001376409 |
| NWD1 | -0.441013717 | 4.95E-06 | 1.65E-05 |
| NWD2 | 1.250302741 | 0.004620175 | 0.009734525 |
| NXNL2 | -1.874040808 | 4.32E-08 | 1.83E-07 |
| NXT2 | 0.473268828 | 6.18E-06 | 2.03E-05 |
| OAF | 0.91282456 | 3.65E-26 | 6.12E-25 |
| OARD1 | -0.73991844 | 2.52E-12 | 1.59E-11 |
| OASL | 3.136266675 | 5.21E-75 | 9.19E-73 |
| OAT | -0.211056213 | 0.010259901 | 0.020034418 |
| OAZ1 | 1.101367936 | 4.40E-33 | 1.09E-31 |
| OAZ2 | 0.911194185 | 9.14E-19 | 9.52E-18 |
| ODAD2 | 0.931060363 | 0.001583228 | 0.00365414 |

| | | | |
|---|---|---|---|
| ODC1 | -0.189997198 | 0.015343734 | 0.028877324 |
| ODF2 | 1.313278895 | 1.28E-34 | 3.45E-33 |
| OFD1 | -0.49741516 | 0.00729729 | 0.014722394 |
| OGDH | -0.50155159 | 0.00013732 | 0.000373141 |
| OGDHL | 2.710222565 | 0.006202933 | 0.012704982 |
| OGFOD1 | -0.787668394 | 1.02E-10 | 5.52E-10 |
| OGFR | -0.671199813 | 9.92E-05 | 0.00027612 |
| OGFRL1 | -0.367199635 | 0.018897226 | 0.034850895 |
| OGG1 | -1.105219176 | 0.000878253 | 0.002112781 |
| OGT | -0.379730677 | 0.000279158 | 0.0007277 |
| OIP5 | -0.863129957 | 1.85E-05 | 5.69E-05 |
| OLA1 | 0.51703388 | 6.13E-09 | 2.81E-08 |
| OLFM1 | 2.767356556 | 9.18E-92 | 2.77E-89 |
| OLFM2 | 1.245343967 | 0.003648513 | 0.007874022 |
| OLFM3 | 0.892135771 | 1.51E-13 | 1.06E-12 |
| OLFML2A | -0.477834009 | 0.005658296 | 0.011691633 |
| OLFML2B | -0.77051402 | 5.52E-16 | 4.67E-15 |
| OLFML3 | -0.375872651 | 0.002116361 | 0.004775956 |
| OPA1 | -0.502640481 | 0.000475961 | 0.001193371 |
| OPHN1 | 0.794423408 | 6.61E-06 | 2.16E-05 |
| OPN5L1 | 1.82903656 | 9.00E-07 | 3.30E-06 |
| OPRD1 | 3.839468672 | 1.03E-22 | 1.38E-21 |
| ORAI1 | -0.419554848 | 0.000464747 | 0.001168919 |
| ORAI3 | -0.313411446 | 0.004437301 | 0.009378292 |
| ORAOV1 | 1.12431677 | 5.21E-23 | 7.05E-22 |
| ORC1 | -1.343944273 | 0.000310563 | 0.000803269 |
| ORC2 | -1.217023094 | 2.58E-09 | 1.22E-08 |
| ORC3 | -0.317187796 | 0.024655242 | 0.044286249 |
| ORC6 | -1.104347535 | 2.49E-18 | 2.52E-17 |
| ORMDL2 | 0.445731369 | 3.53E-07 | 1.36E-06 |
| OSBP | -0.260280408 | 0.019823404 | 0.036396016 |
| OSBPL10 | -1.194950639 | 3.53E-06 | 1.20E-05 |
| OSBPL11 | -0.355340399 | 0.001227705 | 0.002893024 |
| OSBPL1A | -0.332081729 | 0.008127365 | 0.016235618 |
| OSBPL2 | 0.273733174 | 0.015903467 | 0.029802829 |
| OSBPL5 | -0.523578524 | 1.27E-05 | 3.98E-05 |
| OSBPL9 | -0.425464827 | 0.000204608 | 0.000543171 |
| OSER1 | -0.812492024 | 1.21E-13 | 8.57E-13 |
| OSGIN2 | -1.109836563 | 4.60E-10 | 2.33E-09 |
| OSR2 | -1.695661789 | 2.97E-05 | 8.92E-05 |
| OST4 | 1.765841407 | 0.000117612 | 0.000323273 |
| OSTC | 0.473337238 | 4.18E-08 | 1.77E-07 |
| OSTF1 | 0.770672258 | 4.97E-19 | 5.28E-18 |
| OSTM1 | -0.423323149 | 0.000344956 | 0.000887327 |
| OTOF | -2.647175352 | 0.018609722 | 0.034355634 |
| OTOGL | 2.104615098 | 0.000737743 | 0.001797991 |
| OTOS | 0.571675741 | 0.010044661 | 0.01965266 |
| OTUD4 | -0.352809084 | 0.014370725 | 0.027208093 |

| | | | |
|---|---|---|---|
| OTUD6A | 1.294187204 | 6.58E-64 | 6.83E-62 |
| OTULIN | -1.041117247 | 0.003737881 | 0.008040081 |
| OTX2 | 6.515764079 | 2.35E-06 | 8.18E-06 |
| OVCH2 | 2.702598149 | 0.00171164 | 0.00392718 |
| OVOL2 | 2.558003034 | 0.000208846 | 0.000553664 |
| OXCT1 | -0.92624628 | 5.33E-15 | 4.16E-14 |
| OXLD1 | -1.540405837 | 4.15E-09 | 1.93E-08 |
| OXNAD1 | -0.460937075 | 0.000134814 | 0.000366846 |
| OXSR1 | -0.4310989 | 0.000388916 | 0.000991587 |
| P2RX4 | -0.541021723 | 0.001808506 | 0.004132685 |
| P2RX5 | 2.171809661 | 2.20E-19 | 2.38E-18 |
| P2RY1 | 1.064009294 | 3.57E-07 | 1.37E-06 |
| P3H1 | -1.695278972 | 1.50E-90 | 4.16E-88 |
| P3H2 | -1.673040243 | 5.17E-17 | 4.73E-16 |
| P3H3 | -2.200274661 | 1.61E-50 | 9.79E-49 |
| P4HA1 | -0.700618176 | 3.08E-14 | 2.28E-13 |
| P4HA2 | -0.453528435 | 2.15E-06 | 7.56E-06 |
| P4HB | -0.983853738 | 2.08E-25 | 3.33E-24 |
| PABIR2 | 1.138877725 | 2.87E-31 | 6.46E-30 |
| PABPC1 | 1.313503362 | 3.11E-43 | 1.34E-41 |
| PABPC1L | -0.928207214 | 0.008498029 | 0.016901552 |
| PABPC4 | -0.626817193 | 0.025024694 | 0.044825466 |
| PACSIN2 | -1.196789338 | 1.22E-29 | 2.51E-28 |
| PACSIN3 | -0.721269402 | 2.87E-05 | 8.62E-05 |
| PADI3 | 1.102974828 | 0.021653138 | 0.039374573 |
| PAFAH1B1 | -0.761233182 | 4.56E-17 | 4.20E-16 |
| PAFAH1B2 | 0.26536608 | 0.000859526 | 0.002072132 |
| PAICS | -0.932704196 | 2.06E-18 | 2.10E-17 |
| PAIP1 | -0.576232619 | 2.65E-07 | 1.03E-06 |
| PAIP2B | 0.629699883 | 3.39E-11 | 1.92E-10 |
| PAK1IP1 | 0.419351505 | 7.97E-06 | 2.58E-05 |
| PAK2 | 0.896986376 | 8.07E-09 | 3.65E-08 |
| PAK3 | 1.346955239 | 2.32E-13 | 1.61E-12 |
| PALB2 | -0.892635393 | 2.91E-06 | 1.00E-05 |
| PALD1 | 0.595388481 | 3.85E-10 | 1.97E-09 |
| PAN2 | -0.531965624 | 0.003789329 | 0.008139154 |
| PANK1 | -1.11151817 | 1.16E-10 | 6.23E-10 |
| PANK2 | 0.548819977 | 3.11E-08 | 1.33E-07 |
| PANK3 | 0.604944519 | 1.34E-09 | 6.53E-09 |
| PANX1 | 0.629193291 | 0.001204693 | 0.002843728 |
| PANX3 | 1.27465288 | 1.22E-05 | 3.84E-05 |
| PAPD7 | -1.050605038 | 7.69E-15 | 5.94E-14 |
| PAPOLB | 0.49621573 | 1.17E-08 | 5.23E-08 |
| PAPPA2 | -0.503129662 | 6.43E-07 | 2.40E-06 |
| PAPSS1 | -1.044401137 | 2.91E-36 | 8.78E-35 |
| PAPSS2 | -1.363498324 | 5.98E-12 | 3.63E-11 |
| PAQR3 | -0.652823033 | 0.005335 | 0.011091056 |
| PAQR7 | -0.743676588 | 0.000278157 | 0.000725387 |

| | | | |
|---|---|---|---|
| PARD3 | -0.893760116 | 1.09E-12 | 7.12E-12 |
| PARD3B | 0.565195621 | 0.000115893 | 0.000318742 |
| PARD6A | -0.598994928 | 0.021320234 | 0.038829508 |
| PARD6B | -0.919718256 | 1.92E-06 | 6.79E-06 |
| PARG | 0.447815969 | 0.000320245 | 0.000827052 |
| PARN | -0.4070282 | 8.54E-05 | 0.000240562 |
| PARP1 | -0.847731057 | 3.50E-14 | 2.58E-13 |
| PARP16 | 0.580602293 | 3.09E-07 | 1.19E-06 |
| PARP4 | -0.322222302 | 0.006630665 | 0.01351387 |
| PARP6 | 1.177610038 | 1.24E-16 | 1.11E-15 |
| PARP8 | 0.555552588 | 0.002695418 | 0.005970684 |
| PARP9 | 0.520793424 | 0.005815413 | 0.011987123 |
| PARPBP | -0.755611793 | 0.000545114 | 0.001356 |
| PARS2 | -1.525653339 | 2.01E-17 | 1.91E-16 |
| PARVA | 0.583692646 | 8.68E-05 | 0.000243997 |
| PARVB | -0.313991885 | 0.001195719 | 0.002823525 |
| PARVG | 1.093636303 | 0.001213915 | 0.002863508 |
| PAWR | 0.466690325 | 0.001100865 | 0.002616273 |
| PAX1 | 5.231980333 | 0.000747346 | 0.001819439 |
| PAX6 | -2.804963359 | 0.008218264 | 0.01640033 |
| PAX7 | 3.813639798 | 3.17E-12 | 1.99E-11 |
| PAXIP1 | -0.797712381 | 1.76E-05 | 5.43E-05 |
| PBLD | 0.592837174 | 0.00400133 | 0.008559345 |
| PBRM1 | -0.494726463 | 1.06E-05 | 3.38E-05 |
| PBX3 | 0.450027132 | 0.00220274 | 0.004954434 |
| PBX4 | -0.790054948 | 1.97E-07 | 7.76E-07 |
| PBXIP1 | -1.477174758 | 1.98E-06 | 6.97E-06 |
| PCARE | 3.948827165 | 6.13E-05 | 0.000176437 |
| PCBD1 | 1.796039726 | 1.24E-46 | 6.46E-45 |
| PCBD2 | 0.835957356 | 1.33E-11 | 7.83E-11 |
| PCCA | -0.765551647 | 6.33E-06 | 2.08E-05 |
| PCCB | -0.396018327 | 0.000170306 | 0.000457471 |
| PCDH19 | -0.84207912 | 5.05E-07 | 1.91E-06 |
| PCDH7 | -1.014387871 | 0.022562327 | 0.040845385 |
| PCDH8 | -1.206083964 | 4.56E-15 | 3.59E-14 |
| PCDHGC3 | -0.214235489 | 0.017918635 | 0.033203863 |
| PCDHGC5 | -0.986934021 | 0.007854077 | 0.015752194 |
| PCF11 | 0.987667076 | 6.47E-23 | 8.74E-22 |
| PCGF2 | 0.769063012 | 0.000927955 | 0.00222683 |
| PCGF3 | 0.748909795 | 0.003049314 | 0.006689198 |
| PCGF5 | 0.707265695 | 9.43E-08 | 3.85E-07 |
| PCGF6 | -0.94279028 | 4.88E-13 | 3.29E-12 |
| PCID2 | 0.204492094 | 0.016092189 | 0.030135716 |
| PCM1 | -0.797366675 | 5.31E-09 | 2.44E-08 |
| PCMT1 | -0.374548098 | 0.00190887 | 0.00433869 |
| PCMTD1 | 0.790628682 | 9.67E-15 | 7.41E-14 |
| PCNA | -1.275077213 | 8.50E-37 | 2.66E-35 |
| PCNP | 0.734148615 | 6.74E-14 | 4.86E-13 |

| | | | |
|---|---|---|---|
| PCNT | -0.62112976 | 0.001588634 | 0.003665372 |
| PCNX1 | -0.266186945 | 0.014161816 | 0.026872456 |
| PCNX4 | -1.366219591 | 8.83E-07 | 3.24E-06 |
| PCSK5 | -1.141208141 | 2.02E-24 | 3.01E-23 |
| PCSK6 | -1.249445761 | 5.39E-13 | 3.61E-12 |
| PCSK7 | 0.364006064 | 7.35E-05 | 0.000208954 |
| PCYOX1 | -0.43289894 | 0.000358811 | 0.000919661 |
| PCYT1A | -0.601568366 | 1.04E-09 | 5.11E-09 |
| PCYT2 | 1.03715495 | 4.33E-23 | 5.87E-22 |
| PDAP1 | 0.560348362 | 0.000697575 | 0.001706522 |
| PDCD10 | -1.046584042 | 1.60E-28 | 3.10E-27 |
| PDCD11 | -0.915758401 | 3.85E-10 | 1.97E-09 |
| PDCD2 | 0.271139336 | 0.005628654 | 0.011642773 |
| PDCD2L | -1.098302734 | 6.99E-18 | 6.83E-17 |
| PDCD4 | -0.897490828 | 1.32E-14 | 1.00E-13 |
| PDCD5 | 0.688287922 | 4.33E-12 | 2.67E-11 |
| PDCD6 | 0.316249076 | 0.003362192 | 0.007304814 |
| PDCD6IP | -0.819658711 | 1.22E-21 | 1.51E-20 |
| PDCD7 | -0.851331173 | 3.16E-07 | 1.22E-06 |
| PDCL2 | 3.518170208 | 2.07E-05 | 6.34E-05 |
| PDCL3 | 0.585529508 | 1.16E-09 | 5.67E-09 |
| PDDC1 | -0.8169106 | 6.04E-08 | 2.51E-07 |
| PDE12 | -0.584185361 | 1.05E-06 | 3.83E-06 |
| PDE1A | -0.801539421 | 5.33E-06 | 1.76E-05 |
| PDE4B | 2.066178871 | 2.99E-163 | 4.52E-160 |
| PDE6D | 1.172224578 | 5.30E-42 | 2.16E-40 |
| PDE7B | 0.843546361 | 7.53E-15 | 5.82E-14 |
| PDE8A | 0.356319806 | 0.016849489 | 0.03141109 |
| PDE9A | -1.602322939 | 4.07E-23 | 5.54E-22 |
| PDF | -0.893535736 | 0.000258917 | 0.000677942 |
| PDGFA | 1.206812014 | 2.24E-51 | 1.41E-49 |
| PDGFB | 1.557545167 | 7.05E-37 | 2.22E-35 |
| PDGFC | 1.132412534 | 6.93E-06 | 2.26E-05 |
| PDGFD | 0.442104342 | 7.34E-07 | 2.72E-06 |
| PDGFRL | 0.411899511 | 7.64E-06 | 2.48E-05 |
| PDHA2 | -1.599024977 | 1.58E-41 | 6.18E-40 |
| PDHX | -1.021012306 | 2.79E-15 | 2.23E-14 |
| PDIA2 | 1.950435397 | 0.015651709 | 0.029398347 |
| PDIA3 | -0.3708516 | 1.33E-06 | 4.79E-06 |
| PDIA4 | -1.47977594 | 1.08E-39 | 3.77E-38 |
| PDIA5 | -0.504358611 | 1.56E-08 | 6.90E-08 |
| PDIA6 | -0.407793412 | 1.25E-05 | 3.93E-05 |
| PDK1 | 0.403318395 | 3.44E-06 | 1.17E-05 |
| PDK3 | -0.377440832 | 0.000203185 | 0.000539708 |
| PDK4 | -1.404353595 | 8.55E-06 | 2.75E-05 |
| PDLIM3 | 1.00935896 | 0.000127791 | 0.000349344 |
| PDLIM4 | 0.443901902 | 7.67E-05 | 0.000217048 |
| PDLIM7 | 0.73846061 | 2.20E-11 | 1.27E-10 |

| | | | |
|---|---|---|---|
| PDP1 | -1.380810749 | 5.31E-41 | 2.02E-39 |
| PDP2 | -2.132729132 | 1.79E-60 | 1.58E-58 |
| PDPK1 | 0.279783037 | 0.002463979 | 0.005496537 |
| PDPR | -0.658357516 | 0.008742216 | 0.01731374 |
| PDRG1 | 0.991945621 | 2.29E-12 | 1.44E-11 |
| PDS5A | -1.659146796 | 3.31E-46 | 1.70E-44 |
| PDS5B | -0.370256206 | 0.001859481 | 0.00423564 |
| PDSS2 | 0.297540685 | 0.005879582 | 0.012112046 |
| PDXK | 0.548405205 | 2.00E-05 | 6.11E-05 |
| PDXP | -0.684354922 | 1.65E-11 | 9.62E-11 |
| PDZD11 | 0.55002022 | 8.00E-08 | 3.29E-07 |
| PDZD3 | 1.20674044 | 8.60E-06 | 2.77E-05 |
| PDZD7 | 1.834474972 | 0.019422664 | 0.035727818 |
| PDZD8 | -0.684282769 | 9.38E-08 | 3.83E-07 |
| PDZRN3 | -1.941114174 | 1.44E-73 | 2.23E-71 |
| PDZRN4 | -0.957095846 | 0.002130365 | 0.004805166 |
| PECAM1 | 1.851216727 | 1.19E-33 | 3.01E-32 |
| PEF1 | 0.477020838 | 2.02E-08 | 8.83E-08 |
| PELI2 | -1.843951142 | 4.81E-17 | 4.42E-16 |
| PELI3 | -1.480798425 | 1.97E-63 | 1.98E-61 |
| PELO | -0.813888656 | 2.29E-14 | 1.72E-13 |
| PEMT | 1.389249076 | 2.41E-21 | 2.92E-20 |
| PENK | 0.951484884 | 0.000275922 | 0.000720109 |
| PEPD | -0.731385076 | 6.86E-12 | 4.15E-11 |
| PER3 | 0.377958872 | 0.004173279 | 0.008887999 |
| PERP1 | -1.912084144 | 1.32E-07 | 5.34E-07 |
| PERP2 | -1.130332105 | 2.07E-28 | 3.97E-27 |
| PES1 | -0.49762104 | 3.52E-08 | 1.50E-07 |
| PEX10 | 0.647397671 | 2.02E-12 | 1.28E-11 |
| PEX11G | -0.554265822 | 0.000135127 | 0.000367477 |
| PEX12 | -1.059717887 | 8.59E-11 | 4.68E-10 |
| PEX13 | -0.913637646 | 6.72E-14 | 4.85E-13 |
| PEX16 | -0.657824098 | 6.77E-11 | 3.72E-10 |
| PEX2 | 0.758972801 | 6.40E-14 | 4.62E-13 |
| PEX5 | -1.45662631 | 1.64E-15 | 1.33E-14 |
| PEX6 | -0.808156029 | 0.001212503 | 0.002860674 |
| PEX7 | 0.574788126 | 2.71E-08 | 1.17E-07 |
| PFDN2 | 0.593136712 | 4.35E-08 | 1.84E-07 |
| PFDN4 | 0.47666691 | 1.65E-05 | 5.10E-05 |
| PFDN5 | 0.872709954 | 3.03E-16 | 2.62E-15 |
| PFDN6 | 0.486444107 | 0.000490453 | 0.001227893 |
| PFKFB3 | -0.310958315 | 0.019242806 | 0.035425737 |
| PFKL | -1.143464804 | 6.66E-22 | 8.42E-21 |
| PFKM | -1.193348588 | 0.014453587 | 0.027349737 |
| PFKP | -1.0079084 | 4.80E-31 | 1.07E-29 |
| PFN2 | 0.81728927 | 2.17E-22 | 2.85E-21 |
| PGA3 | 4.735107562 | 0.004112617 | 0.008778061 |
| PGAM1 | 0.293378863 | 0.000861079 | 0.002075157 |

| | | | |
|---|---|---|---|
| PGAM5 | -0.259863624 | 0.01195437 | 0.02305716 |
| PGAP2 | 0.774727453 | 1.33E-09 | 6.48E-09 |
| PGAP6 | 0.877134471 | 5.19E-08 | 2.17E-07 |
| PGD | -0.331890535 | 0.000988948 | 0.002366926 |
| PGF | 1.547285679 | 1.90E-23 | 2.64E-22 |
| PGGHG | -1.28617781 | 3.57E-09 | 1.67E-08 |
| PGGT1B | 0.359674667 | 0.000566927 | 0.001406659 |
| PGLS | 0.368755933 | 0.000622483 | 0.001533864 |
| PGM1 | 0.253010419 | 0.011724449 | 0.022636181 |
| PGM2 | -0.543144837 | 5.90E-06 | 1.94E-05 |
| PGM2L1 | -1.23341493 | 7.24E-06 | 2.36E-05 |
| PGM3 | -0.3163844 | 0.002269013 | 0.00509086 |
| PGP | -0.6092265 | 6.88E-08 | 2.85E-07 |
| PGRMC1 | 0.185204025 | 0.019745494 | 0.036267669 |
| PGRMC2 | -0.28375012 | 0.01157332 | 0.022376184 |
| PGS1 | 0.500899808 | 1.12E-05 | 3.55E-05 |
| PHACTR2 | 1.142140815 | 3.86E-06 | 1.30E-05 |
| PHACTR3 | 2.25398342 | 4.28E-17 | 3.96E-16 |
| PHB | -0.366936466 | 0.001268884 | 0.00297767 |
| PHB2 | -0.206929515 | 0.018070101 | 0.033457133 |
| PHC2 | 0.484000037 | 0.000562342 | 0.001396302 |
| PHETA1 | -1.614505514 | 8.80E-38 | 2.91E-36 |
| PHETA2 | 1.296978607 | 3.30E-19 | 3.54E-18 |
| PHEX | -0.90102214 | 4.38E-06 | 1.47E-05 |
| PHF10 | 0.463668182 | 4.25E-07 | 1.62E-06 |
| PHF12 | 0.492811487 | 5.95E-05 | 0.000171607 |
| PHF13 | -1.335407145 | 6.00E-13 | 3.99E-12 |
| PHF19 | 0.585379948 | 0.005319487 | 0.011062189 |
| PHF20 | 0.441997461 | 0.000102357 | 0.00028439 |
| PHF21A | 1.840913271 | 3.51E-81 | 7.02E-79 |
| PHF6 | -0.83543527 | 6.62E-13 | 4.39E-12 |
| PHKA2 | -1.237301166 | 9.52E-12 | 5.70E-11 |
| PHKG1 | 0.521581622 | 1.22E-08 | 5.45E-08 |
| PHLDA2 | 1.063745803 | 2.51E-35 | 7.09E-34 |
| PHLDB2 | -0.30110535 | 0.005361131 | 0.011140267 |
| PHLPP1 | -1.079772078 | 8.86E-11 | 4.82E-10 |
| PHOSPHO1 | 0.782567933 | 9.07E-06 | 2.91E-05 |
| PHYH | 0.303649821 | 0.025210601 | 0.045116852 |
| PHYKPL | 0.937963793 | 8.53E-10 | 4.23E-09 |
| PI4K2B | 0.627846679 | 3.68E-09 | 1.72E-08 |
| PI4KA | -0.71825947 | 1.10E-06 | 3.99E-06 |
| PI4KB | 1.278752668 | 2.90E-14 | 2.16E-13 |
| PIAS1 | -0.43298774 | 0.001080334 | 0.002570711 |
| PIAS3 | 1.82310306 | 0.01345635 | 0.025687643 |
| PIBF1 | -0.806350395 | 1.21E-05 | 3.81E-05 |
| PICK1 | -0.876296433 | 1.23E-13 | 8.67E-13 |
| PID1 | 0.55835758 | 1.16E-08 | 5.18E-08 |
| PIGB | 0.726705396 | 3.69E-07 | 1.41E-06 |

| | | | |
|---|---|---|---|
| PIGBOS1 | 0.958283351 | 1.96E-10 | 1.03E-09 |
| PIGC | -0.419669284 | 0.000809353 | 0.001957426 |
| PIGF | 0.466009292 | 7.06E-06 | 2.30E-05 |
| PIGM | -1.027991341 | 0.000321433 | 0.000829804 |
| PIGO | -0.879673722 | 0.011136483 | 0.021589941 |
| PIGQ | -0.531231412 | 0.025741606 | 0.045958218 |
| PIGR | 4.328242022 | 3.49E-06 | 1.19E-05 |
| PIGS | -1.320020045 | 2.62E-25 | 4.19E-24 |
| PIGT | -0.621891425 | 1.00E-09 | 4.92E-09 |
| PIGU | -0.463419605 | 0.000224327 | 0.000592278 |
| PIGW | -1.114464696 | 1.26E-06 | 4.55E-06 |
| PIGY | 0.706449074 | 1.02E-10 | 5.50E-10 |
| PIH1D2 | -0.977310136 | 0.000850035 | 0.002050342 |
| PIK3AP1 | -1.480117989 | 4.41E-05 | 0.000129562 |
| PIK3C3 | -0.670478026 | 1.72E-08 | 7.57E-08 |
| PIK3CA | -0.394285668 | 0.006616306 | 0.013488651 |
| PIK3CB | -0.765083718 | 7.40E-13 | 4.89E-12 |
| PIK3IP1 | 0.41496 | 0.008072668 | 0.016145336 |
| PIK3R4 | -0.632700468 | 5.78E-07 | 2.17E-06 |
| PIK3R5 | -0.435189406 | 0.010160662 | 0.019862465 |
| PIK3R6 | 1.030292909 | 0.000523509 | 0.001306802 |
| PIKFYVE | -0.854450654 | 0.002827106 | 0.006239004 |
| PIM3 | -1.522641931 | 1.23E-51 | 7.86E-50 |
| PIN1 | 0.617773481 | 1.83E-05 | 5.62E-05 |
| PIN4 | 0.456382388 | 0.000457145 | 0.001151718 |
| PINK1 | -1.513090351 | 7.98E-38 | 2.65E-36 |
| PIP4K2A | 0.460798025 | 1.51E-08 | 6.69E-08 |
| PIP5K1B | 1.275390834 | 2.73E-21 | 3.31E-20 |
| PISD | -1.12603237 | 9.58E-41 | 3.56E-39 |
| PITPNA | 0.466973241 | 1.98E-05 | 6.06E-05 |
| PITRM1 | -0.858927526 | 4.65E-19 | 4.95E-18 |
| PITX1 | 1.347009798 | 2.26E-36 | 6.86E-35 |
| PJVK | 0.869167267 | 0.007080051 | 0.014328776 |
| PKD1 | 0.760490909 | 0.005720531 | 0.011807663 |
| PKD2 | -0.948698499 | 3.84E-09 | 1.79E-08 |
| PKIA | 1.153127266 | 5.31E-43 | 2.26E-41 |
| PKIB | 0.973941765 | 0.00032245 | 0.000832113 |
| PKIG | 1.125081974 | 5.74E-28 | 1.08E-26 |
| PKLR | -0.852395336 | 7.11E-26 | 1.17E-24 |
| PKN1 | -0.99053349 | 0.000572507 | 0.001419726 |
| PKN2 | -0.982675615 | 1.49E-10 | 7.91E-10 |
| PKNOX2 | 0.978981544 | 5.98E-06 | 1.97E-05 |
| PKP4 | 0.817262809 | 1.42E-12 | 9.14E-12 |
| PLA2G12A | 0.532071746 | 0.003257723 | 0.007103958 |
| PLA2G15 | 0.587409908 | 1.75E-05 | 5.39E-05 |
| PLA2G4A | -0.682590272 | 1.14E-14 | 8.66E-14 |
| PLA2G6 | -1.698192977 | 2.90E-20 | 3.31E-19 |
| PLA2G7 | 0.469544934 | 6.94E-08 | 2.87E-07 |

| | | | |
|---|---|---|---|
| PLA2R1 | -1.348067844 | 3.44E-17 | 3.21E-16 |
| PLAA | -1.118123017 | 4.13E-23 | 5.62E-22 |
| PLAGL1 | 0.470505587 | 0.002171007 | 0.004891154 |
| PLAU | -0.939084781 | 1.10E-08 | 4.92E-08 |
| PLBD2 | -0.712703286 | 2.54E-10 | 1.33E-09 |
| PLCD4 | 2.110552764 | 0.010564039 | 0.020600634 |
| PLCE1 | -0.689005632 | 0.008698541 | 0.017237289 |
| PLCH1 | 2.400213161 | 0.013158866 | 0.025173161 |
| PLCL2 | -2.296219269 | 3.02E-10 | 1.56E-09 |
| PLCXD1 | 0.60156891 | 0.001743291 | 0.003992386 |
| PLEK2 | -0.503680873 | 2.56E-07 | 9.98E-07 |
| PLEKHA1 | -0.490095619 | 0.000252736 | 0.000662779 |
| PLEKHA3 | 1.110992974 | 1.59E-29 | 3.25E-28 |
| PLEKHA5 | -0.651514726 | 0.000367946 | 0.000942186 |
| PLEKHA8 | 0.492712755 | 0.015814898 | 0.029661391 |
| PLEKHB2 | 1.156579686 | 1.29E-52 | 8.59E-51 |
| PLEKHF1 | -0.373364014 | 9.13E-06 | 2.93E-05 |
| PLEKHF2 | -1.271891257 | 2.87E-11 | 1.64E-10 |
| PLEKHG1 | -0.947652588 | 0.008627516 | 0.017119007 |
| PLEKHO1 | -0.613798505 | 4.80E-08 | 2.02E-07 |
| PLEKHS1 | 1.369539254 | 0.004667919 | 0.009827503 |
| PLIN2 | -0.813125345 | 8.30E-22 | 1.04E-20 |
| PLIN4 | -1.299798998 | 9.06E-34 | 2.32E-32 |
| PLK1 | -1.575941998 | 1.54E-08 | 6.82E-08 |
| PLK2 | -0.593977484 | 6.34E-13 | 4.21E-12 |
| PLK3 | -2.30703876 | 9.95E-83 | 2.29E-80 |
| PLLP | 1.763559161 | 0.001688835 | 0.003878785 |
| PLOD1 | -1.47571879 | 1.29E-63 | 1.32E-61 |
| PLOD2 | 0.173359377 | 0.025913146 | 0.046221978 |
| PLPP1 | 0.431445972 | 0.000372526 | 0.000952838 |
| PLPP2 | 1.155613198 | 1.52E-12 | 9.78E-12 |
| PLPP3 | 0.261767593 | 0.004404604 | 0.009316429 |
| PLPP5 | -0.689902263 | 0.000546975 | 0.001360132 |
| PLPP6 | -0.680431977 | 1.16E-05 | 3.68E-05 |
| PLRG1 | -1.458129541 | 1.03E-44 | 4.78E-43 |
| PLS3 | -0.178946042 | 0.015583015 | 0.029283055 |
| PLSCR1 | -0.841579631 | 2.30E-13 | 1.59E-12 |
| PLTP | -1.633913012 | 1.58E-25 | 2.56E-24 |
| PLXDC1 | 0.580930739 | 0.000396075 | 0.001008325 |
| PLXDC2 | -0.26422306 | 0.003214363 | 0.007018413 |
| PLXNA1 | -0.614480974 | 2.05E-07 | 8.05E-07 |
| PLXNB2 | -0.809516274 | 3.69E-13 | 2.52E-12 |
| PMAIP1 | 3.94801275 | 2.93E-24 | 4.30E-23 |
| PMEPA1 | 1.069073595 | 1.47E-33 | 3.70E-32 |
| PMM2 | -0.228512716 | 0.025711232 | 0.045910743 |
| PMP22 | 0.628162095 | 7.21E-13 | 4.77E-12 |
| PMPCA | -0.523916366 | 2.45E-06 | 8.54E-06 |
| PMPCB | -0.393702891 | 0.001058604 | 0.002521209 |

| | | | |
|---|---|---|---|
| PMS1 | -0.987423321 | 2.97E-14 | 2.20E-13 |
| PMS2 | -1.547354481 | 6.58E-27 | 1.15E-25 |
| PMVK | 0.387554866 | 0.018484049 | 0.034144495 |
| PNAT10 | 1.849326318 | 0.00403281 | 0.008625326 |
| PNAT3 | 1.070795221 | 4.09E-06 | 1.37E-05 |
| PNISR | 0.748592608 | 7.43E-14 | 5.33E-13 |
| PNLDC1 | 1.536268181 | 0.001835885 | 0.004188911 |
| PNN | -0.307918364 | 0.003844838 | 0.008246652 |
| PNO1 | -0.451250683 | 0.000118045 | 0.000324396 |
| PNPLA2 | -0.825612477 | 0.003316991 | 0.007220799 |
| PNPLA3 | 1.125009899 | 9.15E-34 | 2.33E-32 |
| PNPLA4 | -0.605847863 | 0.008706103 | 0.01724976 |
| PNPLA7 | 0.362072748 | 0.007945631 | 0.01591468 |
| PNPLA8 | -1.850770621 | 6.73E-63 | 6.67E-61 |
| PNPT1 | -0.464626176 | 0.000798464 | 0.001933843 |
| PNRC1 | 1.599709731 | 3.67E-91 | 1.08E-88 |
| PODN | -0.473258378 | 7.48E-07 | 2.77E-06 |
| POFUT1 | -1.126625871 | 2.68E-36 | 8.13E-35 |
| POFUT2 | 0.756549405 | 1.91E-15 | 1.54E-14 |
| POGLUT3 | 1.320055281 | 9.16E-53 | 6.13E-51 |
| POLA1 | -0.633166896 | 0.00019758 | 0.000525643 |
| POLDIP2 | -0.571573944 | 1.69E-07 | 6.72E-07 |
| POLDIP3 | -0.571175599 | 1.49E-08 | 6.59E-08 |
| POLE | -1.748939048 | 0.000276895 | 0.000722234 |
| POLE2 | 0.557711686 | 3.16E-10 | 1.63E-09 |
| POLE4 | 0.682699856 | 0.003543247 | 0.007662659 |
| POLH | -0.324689862 | 0.020583912 | 0.037604373 |
| POLK | -0.813935491 | 2.08E-05 | 6.37E-05 |
| POLR1B | -0.550409426 | 2.00E-06 | 7.06E-06 |
| POLR1E | -0.262648994 | 0.019974196 | 0.036618461 |
| POLR2B | -0.667487449 | 5.20E-09 | 2.40E-08 |
| POLR2C | 0.35518089 | 7.32E-05 | 0.000208278 |
| POLR2E | 0.707389164 | 1.78E-17 | 1.69E-16 |
| POLR2F | 0.484545092 | 9.81E-05 | 0.000273268 |
| POLR2H | 0.834186322 | 1.38E-19 | 1.51E-18 |
| POLR2I | 0.614892715 | 9.33E-08 | 3.81E-07 |
| POLR2K | 0.531090827 | 2.81E-05 | 8.45E-05 |
| POLR2L | 0.305955585 | 0.022399551 | 0.040603376 |
| POLR2M | 0.753560855 | 4.83E-18 | 4.77E-17 |
| POLR3A | -0.378932412 | 0.001358182 | 0.003171893 |
| POLR3B | -0.914471713 | 7.70E-10 | 3.84E-09 |
| POLR3F | -0.449038548 | 0.000199593 | 0.000530685 |
| POLR3K | 0.516723609 | 0.019922477 | 0.03653843 |
| POLRMT | -0.953461385 | 9.70E-17 | 8.72E-16 |
| POMGNT2 | 0.791627549 | 7.28E-06 | 2.37E-05 |
| POMK | -0.901729662 | 6.31E-12 | 3.83E-11 |
| POMP | 0.703879999 | 8.08E-10 | 4.02E-09 |
| POMT2 | 0.487223779 | 0.016566573 | 0.030921844 |

| | | | |
|---|---|---|---|
| POP4 | -1.109834376 | 9.65E-17 | 8.68E-16 |
| POP5 | 0.565220726 | 2.53E-05 | 7.66E-05 |
| POR | -1.093373141 | 2.46E-24 | 3.64E-23 |
| PORCN | -1.701761127 | 1.29E-10 | 6.89E-10 |
| POSTN | -0.418579873 | 3.55E-07 | 1.37E-06 |
| POT1 | -0.720765165 | 3.25E-07 | 1.25E-06 |
| POU1F1 | 3.792388964 | 0.000792457 | 0.001920321 |
| POU2F1 | 0.319101474 | 0.012965525 | 0.024848761 |
| POU2F3 | 1.068376395 | 6.67E-05 | 0.000190693 |
| POU4F2 | -2.945578224 | 0.001686747 | 0.003875299 |
| Pou5f3 | 0.732227275 | 7.62E-06 | 2.48E-05 |
| PPA2 | 0.852557081 | 1.26E-10 | 6.73E-10 |
| PPARD | 0.297291344 | 0.000437045 | 0.001105381 |
| PPARG | -2.131784121 | 2.87E-15 | 2.29E-14 |
| PPAT | -0.631004228 | 4.07E-10 | 2.08E-09 |
| PPDPF | 1.677307878 | 4.55E-74 | 7.20E-72 |
| PPEF1 | 0.691125594 | 0.004372201 | 0.009253652 |
| PPFIA1 | -0.502796878 | 3.03E-08 | 1.30E-07 |
| PPFIBP1 | 0.640866343 | 2.10E-12 | 1.33E-11 |
| PPHLN1 | -0.639347307 | 2.74E-07 | 1.06E-06 |
| PPIA | 1.233721699 | 3.15E-65 | 3.51E-63 |
| PPIB | 0.663507139 | 4.75E-17 | 4.37E-16 |
| PPIC | 1.110167397 | 1.07E-37 | 3.53E-36 |
| PPID | -0.296165211 | 0.005622015 | 0.01163081 |
| PPIE | 0.493163421 | 1.40E-05 | 4.37E-05 |
| PPIF | -0.859286468 | 2.74E-09 | 1.29E-08 |
| PPIH | -0.580619543 | 9.66E-07 | 3.52E-06 |
| PPIL1 | -0.488011262 | 0.00011051 | 0.000305419 |
| PPIL2 | -0.735278957 | 1.45E-08 | 6.43E-08 |
| PPIL3 | -0.457302381 | 0.000738933 | 0.001800344 |
| PPIL4 | -0.345504965 | 0.019584069 | 0.035995493 |
| PPL | -1.234179461 | 1.30E-05 | 4.07E-05 |
| PPM1A | -0.829224527 | 3.64E-12 | 2.26E-11 |
| PPM1B | 0.600164591 | 5.39E-07 | 2.03E-06 |
| PPM1D | -0.672204829 | 1.47E-10 | 7.80E-10 |
| PPM1E | 0.394505358 | 0.000290703 | 0.000754916 |
| PPM1J | -0.931522885 | 0.024819152 | 0.044521831 |
| PPM1K | -0.649717436 | 1.46E-06 | 5.21E-06 |
| PPM1M | 0.364706681 | 0.000223196 | 0.000589406 |
| PPME1 | -0.396052966 | 2.23E-05 | 6.81E-05 |
| PPOX | -0.967008164 | 2.09E-05 | 6.40E-05 |
| PPP1CB | 0.36945658 | 5.90E-06 | 1.94E-05 |
| PPP1R12A | -0.252691992 | 0.003524342 | 0.007624201 |
| PPP1R12B | 0.927436 | 0.00338811 | 0.007357595 |
| PPP1R13B | -0.55588779 | 8.89E-08 | 3.64E-07 |
| PPP1R16B | -1.187100207 | 1.18E-05 | 3.72E-05 |
| PPP1R1C | 0.977490085 | 0.019304608 | 0.035534702 |
| PPP1R2 | 1.119540494 | 4.07E-46 | 2.06E-44 |

| | | | |
|---|---|---|---|
| PPP1R21 | -0.988811811 | 8.26E-13 | 5.42E-12 |
| PPP1R26 | -0.697239777 | 0.00306242 | 0.006713615 |
| PPP1R3B | -1.13069636 | 1.44E-15 | 1.17E-14 |
| PPP1R3E | 0.651655489 | 1.49E-17 | 1.42E-16 |
| PPP1R7 | 0.906442876 | 4.43E-21 | 5.30E-20 |
| PPP1R9B | -0.358561906 | 0.000184497 | 0.000493152 |
| PPP2CA | -0.285932809 | 0.000109863 | 0.000303754 |
| PPP2CB | 1.114045172 | 7.06E-49 | 4.07E-47 |
| PPP2R2A | 0.321616111 | 7.75E-05 | 0.000219131 |
| PPP2R2B | 1.648215976 | 1.01E-26 | 1.75E-25 |
| PPP2R2D | 0.372480441 | 1.47E-05 | 4.59E-05 |
| PPP2R3C | -0.300860299 | 0.008732428 | 0.017296876 |
| PPP2R5A | -0.324951762 | 0.003047404 | 0.006687168 |
| PPP2R5C | 0.337527615 | 0.000641988 | 0.001578777 |
| PPP2R5D | -0.701872599 | 1.88E-05 | 5.77E-05 |
| PPP2R5E | -0.450930127 | 0.000748245 | 0.001820975 |
| PPP3R1 | 0.430949186 | 2.47E-07 | 9.64E-07 |
| PPP4R2 | 0.597014414 | 3.99E-09 | 1.86E-08 |
| PPP4R3A | -0.860573824 | 1.85E-07 | 7.31E-07 |
| PPP6C | -0.285096432 | 0.004920979 | 0.010306001 |
| PPP6R2 | -0.612533342 | 2.96E-06 | 1.02E-05 |
| PPP6R3 | -0.939823844 | 5.43E-20 | 6.10E-19 |
| PPRC1 | -0.456974427 | 0.022688561 | 0.041046576 |
| PPT1 | -0.713798526 | 3.01E-14 | 2.23E-13 |
| PPTC7 | -1.079930964 | 5.77E-06 | 1.90E-05 |
| PQLC2 | -0.947004347 | 4.42E-10 | 2.24E-09 |
| PRAG1 | -0.616989299 | 0.012905267 | 0.024750724 |
| PRDM1 | -1.365264329 | 1.73E-07 | 6.86E-07 |
| PRDM4 | 0.367461867 | 0.00268146 | 0.005943641 |
| PRDX1 | 0.368931952 | 0.000686608 | 0.001680905 |
| PRDX4 | 0.418956505 | 8.23E-06 | 2.66E-05 |
| PRDX6 | 0.491938585 | 2.58E-09 | 1.22E-08 |
| PREB | -1.366991897 | 8.43E-08 | 3.46E-07 |
| PRELID1 | 1.673780114 | 8.89E-123 | 5.75E-120 |
| PRELID3A | 0.517163155 | 1.56E-10 | 8.29E-10 |
| PRELID3B | 1.344593637 | 1.44E-78 | 2.69E-76 |
| PREP | -0.713139787 | 9.14E-14 | 6.50E-13 |
| PRICKLE1 | -1.345429798 | 1.84E-11 | 1.07E-10 |
| PRICKLE2 | 1.095072724 | 2.97E-27 | 5.35E-26 |
| PRIM2 | -1.690615499 | 1.27E-40 | 4.65E-39 |
| PRKAA1 | -1.465762732 | 3.74E-36 | 1.12E-34 |
| PRKAB1 | 0.582451579 | 4.29E-13 | 2.90E-12 |
| PRKAG1 | 0.871209137 | 0.00612016 | 0.012548683 |
| PRKAR1A | -0.536101754 | 5.29E-09 | 2.43E-08 |
| PRKAR1B | 0.369875594 | 6.58E-05 | 0.000188372 |
| PRKCD | -1.426357406 | 7.37E-40 | 2.59E-38 |
| PRKD3 | 0.744960112 | 3.87E-13 | 2.63E-12 |
| PRKX | 0.329130031 | 0.001909303 | 0.004338949 |

| | | | |
|---|---|---|---|
| PRLH | 1.119453685 | 2.00E-14 | 1.50E-13 |
| PRLHR | 1.607180243 | 0.007799634 | 0.01566149 |
| PRMT7 | -0.981590899 | 3.91E-18 | 3.88E-17 |
| PRMT8 | 0.773868258 | 0.003371336 | 0.00732351 |
| PRMT9 | -0.420609631 | 3.46E-05 | 0.000103118 |
| PROC | 2.776769976 | 4.44E-06 | 1.49E-05 |
| PROM2 | 2.481160611 | 1.53E-05 | 4.77E-05 |
| PRORSD1P | -0.293756014 | 0.005639543 | 0.011656428 |
| PROSER1 | -0.819553829 | 1.23E-07 | 4.96E-07 |
| PROZ | 2.295895516 | 3.24E-06 | 1.11E-05 |
| PRPF3 | -0.441128114 | 6.54E-05 | 0.000187338 |
| PRPF38A | -0.81594389 | 5.00E-14 | 3.65E-13 |
| PRPF39 | -0.262890541 | 0.027253623 | 0.048397079 |
| PRPF4 | -0.892047583 | 3.07E-13 | 2.11E-12 |
| PRPF40A | -0.932082878 | 3.70E-20 | 4.20E-19 |
| PRPF4B | -0.305521756 | 0.013074133 | 0.025028679 |
| PRPF6 | -0.226234223 | 0.02422094 | 0.04358679 |
| PRPF8 | -1.11897002 | 4.29E-23 | 5.83E-22 |
| PRPSAP1 | 0.325263135 | 8.71E-05 | 0.000244822 |
| PRPSAP2 | -0.717888258 | 0.001482506 | 0.003442718 |
| PRR14L | -0.778233431 | 4.40E-07 | 1.67E-06 |
| PRR5 | 0.496386528 | 1.15E-05 | 3.63E-05 |
| PRR5L | -0.869696069 | 2.68E-05 | 8.09E-05 |
| PRR7 | 1.954729532 | 5.69E-24 | 8.18E-23 |
| PRRG1 | -0.271105502 | 0.018676491 | 0.034462513 |
| PRRG3 | 1.699068163 | 1.25E-12 | 8.07E-12 |
| PRRG4 | -0.574014173 | 0.000255905 | 0.000670443 |
| PRRX1 | 0.589397263 | 2.09E-08 | 9.09E-08 |
| PRRX2 | 0.979703834 | 7.67E-19 | 8.05E-18 |
| PRSS12 | 0.822389797 | 0.000113901 | 0.000313706 |
| PRSS23 | 1.092779612 | 4.64E-46 | 2.33E-44 |
| PRSS3 | 4.259634856 | 1.24E-06 | 4.47E-06 |
| PRTFDC1 | 0.91485919 | 4.39E-09 | 2.04E-08 |
| PRTG | 0.710991087 | 0.002305341 | 0.005168102 |
| PSAP | -0.967604782 | 1.74E-37 | 5.62E-36 |
| PSAT1 | -1.039372839 | 3.01E-31 | 6.76E-30 |
| PSEN1 | -0.849265818 | 1.26E-05 | 3.97E-05 |
| PSMA1 | 0.393975741 | 3.27E-06 | 1.11E-05 |
| PSMA4 | -0.258014028 | 0.003042179 | 0.00667678 |
| PSMB1 | -0.697298034 | 1.48E-08 | 6.55E-08 |
| PSMB4 | 0.551111355 | 4.77E-09 | 2.21E-08 |
| PSMB6 | 0.332055497 | 0.015419531 | 0.029007926 |
| PSMC2 | -0.668464671 | 2.99E-11 | 1.70E-10 |
| PSMC5 | -0.236155434 | 0.021300865 | 0.038804632 |
| PSMC6 | -0.354571241 | 0.000128414 | 0.000350904 |
| PSMD1 | -0.390283124 | 7.59E-07 | 2.81E-06 |
| PSMD10 | -0.441806943 | 0.000113459 | 0.000312553 |
| PSMD12 | -0.279588492 | 0.005183268 | 0.010803709 |

| | | | |
|---|---|---|---|
| PSMD14 | -0.416286441 | 5.08E-05 | 0.000148183 |
| PSMD2 | -0.947220469 | 1.77E-24 | 2.65E-23 |
| PSMD3 | -0.43600842 | 1.72E-06 | 6.10E-06 |
| PSMD5 | -0.196646554 | 0.026757678 | 0.04757854 |
| PSMD6 | -0.318425595 | 3.18E-05 | 9.51E-05 |
| PSMD7 | 0.295778286 | 0.00062344 | 0.001535943 |
| PSME3 | 0.424302469 | 4.98E-08 | 2.09E-07 |
| PSME4 | -0.833865144 | 1.13E-07 | 4.57E-07 |
| PSMF1 | -0.198474993 | 0.020283259 | 0.03710546 |
| PSMG2 | 0.264974803 | 0.0047527 | 0.009985885 |
| PSMG3 | 0.7481727 | 5.34E-14 | 3.89E-13 |
| PSMG4 | 0.864325892 | 1.73E-10 | 9.15E-10 |
| PSPC1 | -0.94768227 | 1.34E-16 | 1.19E-15 |
| PSPH | -0.907341654 | 8.10E-12 | 4.88E-11 |
| PTBP1 | -0.585605583 | 8.16E-10 | 4.06E-09 |
| PTBP2 | -1.194206637 | 1.88E-19 | 2.05E-18 |
| PTCD1 | -0.522997524 | 0.001000202 | 0.002391756 |
| PTCD3 | -0.889847876 | 5.55E-16 | 4.69E-15 |
| PTCHD1 | 1.200624514 | 2.56E-22 | 3.34E-21 |
| PTCHD4 | 3.87079465 | 4.90E-05 | 0.000143085 |
| PTDSS1 | -0.31412886 | 0.005564275 | 0.011520126 |
| PTEN | 0.520296507 | 2.69E-07 | 1.05E-06 |
| PTER | 1.016693853 | 1.73E-12 | 1.11E-11 |
| PTGDR | -2.267578981 | 4.52E-05 | 0.000132754 |
| PTGER2 | -2.098173449 | 1.53E-06 | 5.45E-06 |
| PTGER3 | 1.583082469 | 8.64E-05 | 0.000242902 |
| PTGES | -1.146092351 | 3.27E-15 | 2.60E-14 |
| PTGES3 | 1.065207954 | 1.49E-35 | 4.30E-34 |
| PTGES3L | 0.574598147 | 3.11E-05 | 9.31E-05 |
| PTGFR | -0.417547027 | 0.004952712 | 0.010366071 |
| PTGFRN | -1.148873624 | 0.001575754 | 0.003639981 |
| PTGS2 | -0.93086363 | 1.05E-28 | 2.06E-27 |
| PTK2 | 0.584519569 | 3.12E-12 | 1.95E-11 |
| PTK7 | -1.15064335 | 1.14E-38 | 3.88E-37 |
| PTN | 0.706606494 | 9.72E-08 | 3.96E-07 |
| PTP4A1 | 1.396438545 | 3.87E-62 | 3.70E-60 |
| PTP4A2 | 1.48215384 | 2.99E-89 | 7.96E-87 |
| PTP4A3 | 0.725211252 | 4.99E-08 | 2.10E-07 |
| PTPMT1 | 0.650182324 | 3.06E-08 | 1.31E-07 |
| PTPN12 | -0.285327601 | 0.00360728 | 0.007788815 |
| PTPN13 | -0.608198346 | 5.10E-07 | 1.93E-06 |
| PTPN20 | -1.59657868 | 0.006899581 | 0.01400106 |
| PTPN22 | 2.720542991 | 7.46E-05 | 0.00021164 |
| PTPN3 | -0.726821844 | 1.62E-06 | 5.75E-06 |
| PTPN5 | -2.184317951 | 1.65E-08 | 7.27E-08 |
| PTPN9 | 0.470014077 | 2.22E-06 | 7.78E-06 |
| PTPRA | 0.772686685 | 2.30E-18 | 2.33E-17 |
| PTPRC | 0.347574809 | 0.000465459 | 0.001170278 |

| | | | |
|---|---|---|---|
| PTPRF | -0.82036118 | 1.09E-05 | 3.46E-05 |
| PTPRM | -1.15238431 | 1.56E-16 | 1.38E-15 |
| PTPRN2 | 0.723446202 | 0.006184525 | 0.012673009 |
| PTPRQ | 0.314874476 | 0.005836839 | 0.012027639 |
| PTPRS | -0.634915096 | 9.57E-08 | 3.90E-07 |
| PTPRZ1 | 1.717302408 | 0.023216747 | 0.041907307 |
| PTRF | -0.230674481 | 0.006416874 | 0.013111569 |
| PTRHD1 | 0.626340669 | 9.23E-05 | 0.000258273 |
| PTS | 1.384539066 | 1.53E-13 | 1.07E-12 |
| PTX3 | 0.275276449 | 0.00867783 | 0.017201264 |
| PUDP | 0.882939401 | 4.79E-11 | 2.66E-10 |
| PUM3 | -0.458824192 | 1.25E-05 | 3.94E-05 |
| PURG | 1.26161958 | 0.000475041 | 0.001191503 |
| PUS1 | 0.758233614 | 4.80E-17 | 4.41E-16 |
| PUS10 | -0.838507522 | 9.78E-05 | 0.000272686 |
| PUS3 | -0.622116994 | 4.42E-05 | 0.000129851 |
| PUS7 | -0.722874158 | 1.74E-08 | 7.66E-08 |
| PWP1 | -0.698124747 | 2.41E-14 | 1.80E-13 |
| PWWP2A | -0.868178058 | 6.37E-07 | 2.38E-06 |
| PWWP2B | -0.536214268 | 1.14E-06 | 4.14E-06 |
| PXDN | -1.669008892 | 2.93E-45 | 1.41E-43 |
| PXMP4 | 0.649815991 | 8.30E-12 | 5.00E-11 |
| PYCR1 | -0.6734759 | 4.76E-15 | 3.74E-14 |
| PYCR3 | -0.420765918 | 0.002746732 | 0.006072476 |
| PYGB | -1.206512073 | 3.64E-09 | 1.70E-08 |
| PYGL | -1.072049752 | 2.60E-19 | 2.80E-18 |
| PYGO2 | 0.736497514 | 0.000422474 | 0.001070919 |
| PYROXD1 | -1.059948227 | 2.57E-06 | 8.93E-06 |
| QARS | -0.531364091 | 2.02E-10 | 1.06E-09 |
| QDPR | -0.377346591 | 0.000435522 | 0.001101939 |
| QPCT | 1.65260943 | 4.97E-80 | 9.79E-78 |
| QPCTL | 1.614069414 | 1.09E-79 | 2.11E-77 |
| QRICH1 | -0.372596018 | 0.00088637 | 0.002130801 |
| QRSL1 | -0.713916085 | 4.33E-08 | 1.83E-07 |
| QSER1 | 0.735911558 | 4.98E-07 | 1.88E-06 |
| QSOX1 | -1.148346072 | 4.67E-22 | 5.97E-21 |
| QSOX2 | -0.829083005 | 1.93E-07 | 7.62E-07 |
| QTRT2 | -0.272475547 | 0.004124321 | 0.008800279 |
| R3HCC1 | -0.568215275 | 0.000121927 | 0.000334591 |
| R3HDM1 | 1.09130757 | 1.23E-26 | 2.13E-25 |
| R3HDM4 | -0.939749245 | 1.85E-12 | 1.18E-11 |
| RAB11B | 1.16851119 | 6.42E-40 | 2.27E-38 |
| RAB11FIP2 | 1.452920813 | 5.16E-39 | 1.78E-37 |
| RAB11FIP3 | 0.951013011 | 7.97E-16 | 6.65E-15 |
| RAB11FIP4 | -1.314166904 | 7.01E-05 | 0.000199733 |
| RAB12 | 0.645979584 | 1.00E-11 | 5.97E-11 |
| RAB15 | 1.24107405 | 3.05E-07 | 1.18E-06 |
| RAB1B | 0.859068916 | 2.21E-32 | 5.27E-31 |

| | | | |
|---|---|---|---|
| RAB20 | -0.848949043 | 2.16E-07 | 8.46E-07 |
| RAB21 | 0.318671521 | 0.000370395 | 0.000947564 |
| RAB23 | -0.3363394 | 0.000423968 | 0.001074506 |
| RAB26 | 0.975731485 | 1.67E-05 | 5.14E-05 |
| RAB27A | 0.974232367 | 3.15E-23 | 4.32E-22 |
| RAB29 | 0.864156443 | 1.72E-14 | 1.30E-13 |
| RAB2A | 0.676460335 | 9.41E-13 | 6.15E-12 |
| RAB30 | -0.500210468 | 5.60E-07 | 2.11E-06 |
| RAB31 | 1.165999768 | 4.56E-52 | 2.96E-50 |
| RAB32 | -1.119159142 | 3.56E-17 | 3.32E-16 |
| RAB33B | 0.550550232 | 3.17E-10 | 1.63E-09 |
| RAB34 | 0.30536621 | 0.001180879 | 0.002791879 |
| RAB39A | 0.82351947 | 0.004134948 | 0.008820184 |
| RAB39B | 0.468621868 | 1.82E-05 | 5.60E-05 |
| RAB3GAP1 | -0.385626795 | 0.000316767 | 0.000818381 |
| RAB3GAP2 | -0.969540203 | 2.63E-21 | 3.19E-20 |
| RAB3IL1 | 0.826879437 | 4.23E-21 | 5.07E-20 |
| RAB43 | 0.872679395 | 1.62E-23 | 2.27E-22 |
| RAB4A | -0.384219007 | 0.001130681 | 0.002680191 |
| RAB5A | -0.706284084 | 2.29E-14 | 1.71E-13 |
| RAB5C | 0.806142262 | 1.04E-20 | 1.21E-19 |
| RAB6A | 0.254305528 | 0.004421973 | 0.009348804 |
| RAB7A | 0.387287155 | 0.000175217 | 0.000469641 |
| RAB7B | 1.386007084 | 1.02E-46 | 5.34E-45 |
| RAB8B | 0.46378567 | 0.00867414 | 0.017198967 |
| RAB9A | 0.300266657 | 0.000472405 | 0.001185766 |
| RAB9B | 0.801154707 | 7.73E-06 | 2.51E-05 |
| RABEP1 | 0.225383071 | 0.023781697 | 0.042858768 |
| RABEPK | 0.610687415 | 1.95E-12 | 1.24E-11 |
| RABGAP1L | 0.587831076 | 3.03E-08 | 1.30E-07 |
| RABGGTB | -0.533108287 | 0.001176457 | 0.002781907 |
| RABL6 | 0.628009501 | 3.72E-10 | 1.90E-09 |
| RAC1 | 0.653811214 | 2.01E-12 | 1.28E-11 |
| RAC2 | 0.884139728 | 4.61E-05 | 0.000135021 |
| RAC3 | 0.532960467 | 1.85E-09 | 8.90E-09 |
| RACGAP1 | -1.155787325 | 1.39E-05 | 4.33E-05 |
| RAD21 | -0.744102765 | 9.12E-16 | 7.57E-15 |
| RAD21L1 | 3.284802175 | 0.006412145 | 0.013107817 |
| RAD51 | -0.546128747 | 0.001643096 | 0.003782042 |
| RAD51B | 0.843265013 | 5.72E-07 | 2.15E-06 |
| RAD51C | -0.746708837 | 0.000189897 | 0.000506691 |
| RAD51D | -0.364646112 | 0.003805112 | 0.008167891 |
| RAD54B | -1.222282973 | 0.000148578 | 0.000402365 |
| RAD54L | -1.915418586 | 4.98E-13 | 3.35E-12 |
| RAD54L2 | 0.442986344 | 0.017245848 | 0.032078855 |
| RADX | 1.394776672 | 8.27E-10 | 4.10E-09 |
| RAF1 | -1.121687545 | 1.06E-29 | 2.19E-28 |
| RAI14 | -0.285101915 | 0.004163405 | 0.00886836 |

| | | | |
|---|---|---|---|
| RAI2 | 1.17994269 | 1.05E-14 | 8.04E-14 |
| RALA | 0.359515952 | 0.000170638 | 0.000458271 |
| RALB | 0.590379766 | 7.20E-12 | 4.35E-11 |
| RALBP1 | 0.84113995 | 6.20E-22 | 7.88E-21 |
| RALGAPB | -0.390092144 | 0.007207369 | 0.014560428 |
| RAMP1 | 0.520424253 | 6.02E-07 | 2.25E-06 |
| RAMP3 | 0.648381384 | 9.78E-05 | 0.000272759 |
| RAN | -0.216628759 | 0.018182527 | 0.03365154 |
| RANBP9 | -0.638646859 | 5.55E-13 | 3.71E-12 |
| RANGAP1 | -0.608094431 | 3.27E-10 | 1.68E-09 |
| RAP1B | 0.365822813 | 2.48E-06 | 8.63E-06 |
| RAP1GAP1 | 0.887148393 | 2.08E-05 | 6.37E-05 |
| RAP1GDS1 | -0.375369153 | 3.13E-06 | 1.07E-05 |
| RAP2A | 0.428998525 | 0.020726523 | 0.037844559 |
| RAP2C | 1.077118212 | 2.57E-32 | 6.12E-31 |
| RARA | 0.942170899 | 0.020312963 | 0.037151713 |
| RARB | 0.912817288 | 0.001158022 | 0.002740699 |
| RARS | 0.21939674 | 0.006092919 | 0.012502257 |
| RASA2 | -0.954806759 | 6.11E-14 | 4.42E-13 |
| RASA3 | -1.281113066 | 8.15E-31 | 1.78E-29 |
| RASAL2 | 0.36005539 | 0.00663298 | 0.013516561 |
| RASD1 | -1.215691891 | 0.000714047 | 0.001743368 |
| RASD2 | 2.676036352 | 9.77E-09 | 4.39E-08 |
| RASGEF1A | -0.78779094 | 0.003704994 | 0.00798324 |
| RASGEF1B | -1.500152391 | 4.23E-32 | 1.00E-30 |
| RASGEF1C | -4.552628365 | 0.002592813 | 0.005766898 |
| RASGRP1 | -0.897730671 | 1.80E-12 | 1.15E-11 |
| RASGRP3 | 2.794657504 | 3.37E-27 | 6.04E-26 |
| RASL11A | 1.328947301 | 2.59E-08 | 1.12E-07 |
| RASL11B | -1.404183025 | 3.14E-27 | 5.65E-26 |
| RASSF2 | 1.329431371 | 0.002904947 | 0.006401434 |
| RASSF3 | 0.548860698 | 3.41E-07 | 1.31E-06 |
| RASSF9 | -0.589314134 | 0.00498341 | 0.010425506 |
| RAVER2 | -0.476668632 | 0.002710747 | 0.006001705 |
| RB1 | -0.464681605 | 1.67E-06 | 5.93E-06 |
| RB1CC1 | -0.331309232 | 0.000191243 | 0.000510082 |
| RBBP4 | 0.349404481 | 1.18E-05 | 3.73E-05 |
| RBBP5 | -0.622255336 | 3.13E-06 | 1.07E-05 |
| RBBP6 | 0.602544265 | 2.80E-08 | 1.20E-07 |
| RBBP7 | -1.22416629 | 7.85E-12 | 4.74E-11 |
| RBBP8 | 0.504459582 | 2.55E-05 | 7.70E-05 |
| RBFA | -0.689649124 | 0.00022114 | 0.000584205 |
| RBFOX2 | 0.336750748 | 0.005503485 | 0.011409909 |
| RBIS | 0.762059744 | 2.28E-07 | 8.92E-07 |
| RBKS | 0.832444933 | 4.86E-07 | 1.84E-06 |
| RBL1 | -0.65951274 | 0.013557466 | 0.025839178 |
| RBL2 | -0.587436315 | 0.000276503 | 0.00072135 |
| RBM11 | 1.949581942 | 1.61E-05 | 4.98E-05 |

| | | | |
|---|---|---|---|
| RBM12 | -1.311332573 | 8.97E-21 | 1.05E-19 |
| RBM12B | -0.406194776 | 0.003593642 | 0.007762997 |
| RBM14 | -0.453649224 | 0.004789356 | 0.010056685 |
| RBM15B | -1.832284508 | 3.95E-44 | 1.79E-42 |
| RBM17 | -0.586269844 | 8.99E-09 | 4.05E-08 |
| RBM18 | 1.134628536 | 1.40E-44 | 6.52E-43 |
| RBM19 | 0.269302655 | 0.013653188 | 0.0260017 |
| RBM24 | 1.089463297 | 3.18E-15 | 2.52E-14 |
| RBM26 | -0.748319289 | 1.67E-09 | 8.06E-09 |
| RBM3 | 0.27226319 | 0.002891928 | 0.006374811 |
| RBM33 | -0.63821136 | 2.36E-05 | 7.19E-05 |
| RBM38 | 0.972878862 | 1.72E-07 | 6.81E-07 |
| RBM39 | -0.374751807 | 3.53E-05 | 0.000104919 |
| RBM41 | -0.84619324 | 0.000205378 | 0.000544895 |
| RBM43 | -1.037711245 | 6.69E-07 | 2.49E-06 |
| RBM45 | -0.446096612 | 4.94E-06 | 1.64E-05 |
| RBM48 | -0.597581885 | 3.44E-07 | 1.32E-06 |
| RBMX2 | 0.376190491 | 0.000220131 | 0.000581653 |
| RBP | 0.91365462 | 0.006545822 | 0.013358982 |
| RBP4A | -0.843287003 | 4.19E-07 | 1.60E-06 |
| RBP5 | 1.039739503 | 1.03E-06 | 3.74E-06 |
| RBP7 | 0.709261683 | 1.12E-05 | 3.55E-05 |
| RBPMS | 1.560139701 | 2.21E-51 | 1.41E-49 |
| RBSN | -0.380901045 | 0.012923073 | 0.024777882 |
| RBX1 | 0.849710649 | 3.88E-11 | 2.18E-10 |
| RC3H2 | 0.770679107 | 3.75E-06 | 1.27E-05 |
| RCAN1 | 1.471591438 | 7.95E-75 | 1.39E-72 |
| RCAN3 | 1.19825528 | 2.16E-51 | 1.38E-49 |
| RCBTB1 | 0.382633164 | 0.000243182 | 0.000639701 |
| RCC1 | 1.221675099 | 2.19E-06 | 7.68E-06 |
| RCC1L | -0.697992697 | 4.92E-08 | 2.07E-07 |
| RCHY1 | 0.64023199 | 5.12E-13 | 3.44E-12 |
| RCL1 | -1.314100386 | 1.28E-22 | 1.70E-21 |
| RCN2 | 1.089497878 | 3.48E-43 | 1.49E-41 |
| RCOR3 | 1.076878212 | 3.34E-15 | 2.65E-14 |
| RCSD1 | 2.016859188 | 1.10E-07 | 4.45E-07 |
| RD3 | 2.587292695 | 0.000531389 | 0.001324526 |
| RDH10 | -1.678192109 | 1.56E-32 | 3.71E-31 |
| RDX | 0.226423783 | 0.008457172 | 0.016827679 |
| RECK | -1.135609909 | 5.99E-18 | 5.89E-17 |
| RECQL | -1.057835383 | 2.56E-14 | 1.91E-13 |
| RECQL5 | -1.836560504 | 9.23E-23 | 1.24E-21 |
| REEP1 | 0.875139127 | 0.004812317 | 0.0100971 |
| REEP2 | 0.635372428 | 0.000598596 | 0.001479025 |
| REEP3 | 0.204566511 | 0.017478234 | 0.032458554 |
| REEP5 | 0.549702892 | 2.27E-10 | 1.19E-09 |
| REL | -1.108410376 | 8.82E-22 | 1.10E-20 |
| RELCH | -0.787340116 | 2.65E-06 | 9.19E-06 |

| | | | |
|---|---|---|---|
| RELN | -1.482179322 | 2.50E-05 | 7.58E-05 |
| REPS1 | -0.889376048 | 1.84E-12 | 1.17E-11 |
| RER1 | 0.785283902 | 1.50E-13 | 1.05E-12 |
| RERG | 1.253391187 | 3.54E-41 | 1.36E-39 |
| RERGL | 1.102890366 | 4.50E-28 | 8.47E-27 |
| RET | -0.768531174 | 2.40E-09 | 1.14E-08 |
| RETREG1 | -1.05126846 | 0.020031554 | 0.03670876 |
| REXO2 | 0.620240693 | 4.63E-10 | 2.35E-09 |
| RFC3 | -1.078443576 | 8.63E-12 | 5.18E-11 |
| RFC4 | -0.702409846 | 1.62E-09 | 7.83E-09 |
| RFC5 | 0.388412145 | 0.001134318 | 0.002687874 |
| RFKL | 1.260221199 | 3.57E-40 | 1.28E-38 |
| RFLNB | 0.842725952 | 4.30E-08 | 1.82E-07 |
| RFT1 | -1.271020714 | 6.60E-31 | 1.46E-29 |
| RFWD3 | -0.917254282 | 0.000197329 | 0.000525235 |
| RFX2 | -1.384439526 | 5.23E-36 | 1.55E-34 |
| RFX3 | -1.325246317 | 7.75E-19 | 8.13E-18 |
| RFX5 | 1.067105356 | 9.85E-16 | 8.14E-15 |
| RFX7 | 0.550584401 | 0.004329955 | 0.009171095 |
| RFXANK | 0.711853796 | 5.55E-13 | 3.71E-12 |
| RFXAP | -0.440813389 | 4.45E-07 | 1.69E-06 |
| RGCC | -0.310273551 | 0.001122921 | 0.00266412 |
| RGL1 | 1.416946338 | 6.11E-46 | 3.02E-44 |
| RGN | 0.7873453 | 0.000125186 | 0.000342912 |
| RGP1 | -0.72793291 | 1.27E-05 | 4.00E-05 |
| RGS10 | 0.963394654 | 6.55E-15 | 5.10E-14 |
| RGS11 | 1.049573081 | 0.009685823 | 0.019019027 |
| RGS2 | 1.683456073 | 1.01E-64 | 1.11E-62 |
| RGS20 | -0.448677683 | 0.000122346 | 0.000335606 |
| RHBDD1 | -0.986215499 | 1.35E-05 | 4.23E-05 |
| RHBDF1 | 0.360033087 | 0.002969618 | 0.006532295 |
| RHBDF2 | -0.873507203 | 2.15E-05 | 6.56E-05 |
| RHBG | -0.987566103 | 0.004725099 | 0.009938648 |
| RHCE | 0.491818427 | 0.004158928 | 0.008861601 |
| RHEB | 0.752486099 | 7.62E-21 | 9.00E-20 |
| RHOA | 1.082554063 | 1.64E-47 | 8.86E-46 |
| RHOB | -2.097532172 | 3.31E-82 | 7.26E-80 |
| RHOBTB2 | 0.734797377 | 5.67E-06 | 1.87E-05 |
| RHOF | 0.923705092 | 9.38E-15 | 7.20E-14 |
| RHOJ | 1.400195908 | 1.50E-17 | 1.43E-16 |
| RHOQ | 2.187264159 | 2.91E-134 | 2.20E-131 |
| RHOT1 | -0.201138741 | 0.019991442 | 0.036640195 |
| RHOT2 | 0.515553396 | 1.40E-06 | 5.00E-06 |
| RHPN1 | -2.116809419 | 1.25E-05 | 3.93E-05 |
| RHPN2 | -1.585673304 | 3.41E-33 | 8.49E-32 |
| RIC1 | -0.653033313 | 1.15E-06 | 4.18E-06 |
| RIC3 | -0.390171977 | 0.004275322 | 0.009078297 |
| RIF1 | -0.374817806 | 0.013226572 | 0.025288447 |

| | | | |
|---|---|---|---|
| RILP | -0.987027029 | 2.56E-05 | 7.74E-05 |
| RILPL1 | 0.797992267 | 8.21E-24 | 1.17E-22 |
| RILPL2 | 1.24825757 | 4.35E-22 | 5.58E-21 |
| RIMKLB | -0.874998729 | 5.46E-08 | 2.28E-07 |
| RIMS2 | 0.460767469 | 0.005508046 | 0.011417624 |
| RIMS3 | 2.050141279 | 0.000174442 | 0.000467655 |
| RIN2 | -0.977423163 | 2.12E-23 | 2.94E-22 |
| RIN3 | -0.798816401 | 0.000382961 | 0.000977503 |
| RINT1 | -1.816158092 | 1.46E-57 | 1.14E-55 |
| RIOK2 | -0.591493621 | 6.68E-07 | 2.49E-06 |
| RIOK3 | -1.164944698 | 5.34E-37 | 1.70E-35 |
| RIPK2 | -0.618567135 | 4.19E-07 | 1.60E-06 |
| RIPK4 | 0.843748338 | 0.007410581 | 0.0149288 |
| RIPOR3 | 0.703995008 | 3.10E-06 | 1.06E-05 |
| RIT1 | 1.037392738 | 7.33E-32 | 1.71E-30 |
| RIT2 | 3.427744816 | 7.14E-08 | 2.95E-07 |
| RLF | 0.544791384 | 1.87E-07 | 7.39E-07 |
| RLIM | -0.77400756 | 3.64E-13 | 2.49E-12 |
| RLN3 | -2.219436056 | 3.34E-11 | 1.89E-10 |
| RMDN1 | -0.54546071 | 0.002862325 | 0.006313651 |
| RMDN3 | -0.64916231 | 8.25E-06 | 2.66E-05 |
| RMI1 | -0.418534843 | 0.024879427 | 0.044600524 |
| RMND1 | 0.281844917 | 0.006259724 | 0.012811651 |
| RMND5A | -0.475374612 | 0.000129354 | 0.00035319 |
| RNASEL | -0.677425701 | 0.012523184 | 0.024085908 |
| RND3 | -0.533664493 | 2.07E-07 | 8.14E-07 |
| RNF10 | 0.686324391 | 4.94E-15 | 3.88E-14 |
| RNF103 | -0.437039962 | 2.49E-05 | 7.56E-05 |
| RNF11 | 1.088243779 | 2.15E-48 | 1.21E-46 |
| RNF111 | -0.678875618 | 7.63E-09 | 3.47E-08 |
| RNF113B | -0.51400411 | 2.61E-05 | 7.90E-05 |
| RNF114 | 0.959853354 | 1.44E-25 | 2.35E-24 |
| RNF121 | 0.629264512 | 3.79E-05 | 0.000112216 |
| RNF122 | 1.004902531 | 2.47E-09 | 1.17E-08 |
| RNF123 | -0.904027894 | 2.39E-08 | 1.04E-07 |
| RNF126 | 0.874387778 | 6.42E-21 | 7.59E-20 |
| RNF128 | -1.528826331 | 2.00E-13 | 1.40E-12 |
| RNF13 | 0.916316625 | 9.60E-25 | 1.47E-23 |
| RNF130 | 0.914570168 | 5.02E-34 | 1.31E-32 |
| RNF138 | 0.878367102 | 1.86E-11 | 1.08E-10 |
| RNF139 | -0.468549324 | 2.26E-06 | 7.89E-06 |
| RNF141 | 1.115574688 | 5.48E-37 | 1.74E-35 |
| RNF146 | -0.605856672 | 1.12E-09 | 5.49E-09 |
| RNF149 | -0.814184898 | 5.19E-12 | 3.17E-11 |
| RNF165 | 1.22525086 | 3.06E-22 | 3.96E-21 |
| RNF166 | 1.011243456 | 4.40E-16 | 3.76E-15 |
| RNF168 | -0.716038148 | 2.14E-06 | 7.52E-06 |
| RNF170 | 1.025575309 | 2.49E-05 | 7.56E-05 |

| | | | |
|---|---|---|---|
| RNF19A | -1.722803956 | 1.13E-31 | 2.60E-30 |
| RNF2 | 0.499751689 | 0.000316764 | 0.000818381 |
| RNF20 | -0.468430412 | 0.002910429 | 0.006412473 |
| RNF207 | 0.690257273 | 0.001345874 | 0.003144229 |
| RNF214 | -0.879413339 | 1.06E-13 | 7.54E-13 |
| RNF215 | -0.526805361 | 0.016885543 | 0.031469669 |
| RNF216 | 0.46365038 | 0.020471114 | 0.037423453 |
| RNF220 | 0.637108123 | 7.79E-12 | 4.70E-11 |
| RNF25 | -0.527548774 | 5.91E-05 | 0.000170396 |
| RNF34 | -0.531345134 | 2.98E-09 | 1.40E-08 |
| RNF38 | 1.303492747 | 6.10E-41 | 2.31E-39 |
| RNF5 | 0.906304461 | 1.66E-29 | 3.38E-28 |
| RNF6 | -0.971774343 | 2.92E-17 | 2.73E-16 |
| RNF7 | 1.115651227 | 5.98E-31 | 1.33E-29 |
| RNF8 | -0.606257608 | 5.87E-07 | 2.20E-06 |
| RNH1 | -0.754311507 | 7.73E-17 | 6.99E-16 |
| RNLS | 0.526693989 | 0.027790045 | 0.049227458 |
| RNMT | -0.476730241 | 5.33E-06 | 1.76E-05 |
| RNPEP | -1.213568188 | 1.03E-14 | 7.88E-14 |
| RNPEPL1 | -1.531367096 | 0.003508917 | 0.007594459 |
| RNPS1 | -0.24294273 | 0.005030638 | 0.010516216 |
| ROBO1 | -0.94552694 | 2.14E-08 | 9.29E-08 |
| ROGDI | 0.399060892 | 0.009525195 | 0.018730678 |
| ROMO1 | 1.072613702 | 4.49E-21 | 5.36E-20 |
| ROPN1L | 0.949429535 | 0.000800236 | 0.001937787 |
| ROR1 | -1.552862297 | 1.15E-23 | 1.63E-22 |
| ROR2 | -1.000592863 | 0.001720093 | 0.003944348 |
| RORB | 0.532210337 | 4.83E-05 | 0.000141242 |
| RP11-529K1.3 | 0.655759162 | 3.01E-15 | 2.40E-14 |
| RP1-27O5.3 | -1.081263524 | 9.60E-09 | 4.33E-08 |
| RP2 | 0.649892531 | 7.39E-16 | 6.19E-15 |
| RPA1 | -1.269494907 | 8.15E-35 | 2.23E-33 |
| RPA3 | -0.424026472 | 0.001254924 | 0.002950179 |
| RPAP1 | -0.593189551 | 2.66E-05 | 8.02E-05 |
| RPAP3 | -0.920580253 | 7.75E-13 | 5.11E-12 |
| RPF1 | -0.559578201 | 7.90E-07 | 2.92E-06 |
| RPF2 | -0.766971193 | 3.61E-09 | 1.69E-08 |
| RPGRIP1L | -1.011815733 | 0.00028423 | 0.000739237 |
| RPIA | 0.920084144 | 4.93E-13 | 3.32E-12 |
| RPL10A | 0.71980741 | 1.17E-11 | 6.92E-11 |
| RPL11 | 0.306486558 | 0.007220673 | 0.014579883 |
| RPL12 | 1.077993476 | 9.19E-32 | 2.13E-30 |
| RPL13 | 0.454117979 | 6.52E-06 | 2.13E-05 |
| RPL14 | 0.561876832 | 8.83E-07 | 3.24E-06 |
| RPL18A | 0.267281225 | 0.013802904 | 0.026242701 |
| RPL19 | 0.743780858 | 4.02E-16 | 3.44E-15 |
| RPL21 | 0.867660749 | 3.86E-18 | 3.85E-17 |
| RPL22 | 0.785277099 | 4.45E-15 | 3.51E-14 |

| | | | |
|---|---|---|---|
| RPL22L1 | 1.394182069 | 1.65E-34 | 4.42E-33 |
| RPL23 | 0.315276381 | 0.000703509 | 0.001719184 |
| RPL23A | 1.167770495 | 5.70E-31 | 1.27E-29 |
| RPL24 | 0.41975231 | 5.99E-05 | 0.000172601 |
| RPL26L1 | 0.334469835 | 0.00513333 | 0.010704544 |
| RPL27 | 0.748600628 | 7.13E-15 | 5.52E-14 |
| RPL27A | 0.603417603 | 1.71E-07 | 6.80E-07 |
| RPL29 | 0.936937221 | 1.98E-17 | 1.87E-16 |
| RPL3 | 0.336429088 | 4.38E-06 | 1.47E-05 |
| RPL30 | 0.923156813 | 1.81E-16 | 1.59E-15 |
| RPL31 | 0.637202296 | 7.59E-10 | 3.79E-09 |
| RPL32 | 0.909560956 | 1.78E-18 | 1.82E-17 |
| RPL34 | 0.565867564 | 3.40E-07 | 1.31E-06 |
| RPL35 | 0.390158335 | 0.003595277 | 0.007765295 |
| RPL35A | 0.671655832 | 9.78E-09 | 4.40E-08 |
| RPL36 | 0.365347119 | 0.003056345 | 0.006702459 |
| RPL36AL | 1.074456535 | 9.33E-18 | 9.02E-17 |
| RPL37 | 0.708943953 | 5.44E-09 | 2.50E-08 |
| RPL37A | 1.263023313 | 7.39E-31 | 1.63E-29 |
| RPL38 | 0.444477789 | 0.001266264 | 0.002973063 |
| RPL39L | 1.637840416 | 3.28E-83 | 7.69E-81 |
| RPL3L | 1.256024852 | 0.02177851 | 0.03957929 |
| RPL4 | -0.296783421 | 0.000166712 | 0.00044817 |
| RPL6 | 0.41707471 | 4.21E-07 | 1.61E-06 |
| RPL7 | 0.577168577 | 7.82E-09 | 3.55E-08 |
| RPL7A | 0.683779684 | 2.08E-12 | 1.32E-11 |
| RPL7L1 | 0.257434612 | 0.009819221 | 0.019261488 |
| RPL9 | 0.372291077 | 0.000173376 | 0.000465074 |
| RPLP0 | 1.329545862 | 6.34E-74 | 9.90E-72 |
| RPLP1 | 1.701770623 | 4.35E-59 | 3.63E-57 |
| RPLP2 | 0.971099166 | 1.21E-24 | 1.83E-23 |
| RPN1 | -0.394333621 | 2.70E-07 | 1.05E-06 |
| RPN2 | -0.678757869 | 1.22E-13 | 8.62E-13 |
| RPP25L | -0.497298707 | 1.44E-06 | 5.16E-06 |
| RPP30 | -0.330290233 | 0.002677869 | 0.005939556 |
| RPP38 | -1.087611793 | 5.36E-13 | 3.60E-12 |
| RPP40 | -1.870103117 | 1.10E-40 | 4.05E-39 |
| RPRD2 | 0.713132373 | 0.000111753 | 0.000308478 |
| RPS10 | 1.075958729 | 6.99E-35 | 1.91E-33 |
| RPS11 | 1.453278001 | 5.13E-76 | 9.29E-74 |
| RPS12 | 0.843892283 | 1.64E-14 | 1.24E-13 |
| RPS13 | 1.070060562 | 2.26E-28 | 4.34E-27 |
| RPS14 | 0.404785335 | 0.000237897 | 0.000626769 |
| RPS15 | 1.118343537 | 5.78E-27 | 1.01E-25 |
| RPS15A | 0.91633689 | 2.75E-17 | 2.59E-16 |
| RPS16 | 0.544994121 | 4.52E-07 | 1.72E-06 |
| RPS17 | 0.444070305 | 0.000218827 | 0.000578656 |
| RPS19 | 1.127931627 | 2.03E-30 | 4.37E-29 |

| | | | |
|---|---|---|---|
| RPS2 | 0.962588356 | 3.48E-27 | 6.23E-26 |
| RPS20 | 1.544390088 | 4.18E-70 | 5.46E-68 |
| RPS21 | 0.675337601 | 1.91E-08 | 8.37E-08 |
| RPS23 | 0.479940634 | 1.29E-05 | 4.05E-05 |
| RPS24 | 1.108867143 | 6.29E-26 | 1.04E-24 |
| RPS27A | 0.566860153 | 1.47E-08 | 6.52E-08 |
| RPS27L | 0.994820253 | 1.31E-20 | 1.52E-19 |
| RPS28 | 0.954230462 | 2.54E-20 | 2.91E-19 |
| RPS29 | 1.715095768 | 2.97E-58 | 2.34E-56 |
| RPS3 | 0.401507403 | 4.00E-05 | 0.000118059 |
| RPS3A | 0.935718712 | 5.01E-17 | 4.58E-16 |
| RPS4Y1 | 0.869154709 | 9.86E-25 | 1.50E-23 |
| RPS6 | 0.653188464 | 1.56E-16 | 1.38E-15 |
| RPS6KA1 | 0.600246767 | 0.000534427 | 0.001330878 |
| RPS6KA3 | -0.69668542 | 0.001692285 | 0.003886052 |
| RPS6KA6 | -0.947145756 | 7.76E-06 | 2.52E-05 |
| RPS6KB1 | 0.383998961 | 0.000347816 | 0.000893331 |
| RPS6KC1 | -0.897626739 | 4.54E-08 | 1.91E-07 |
| RPS6KL1 | 1.005948326 | 4.93E-14 | 3.60E-13 |
| RPS8 | 0.392602953 | 3.97E-05 | 0.000117145 |
| RPSAP58 | 1.338768632 | 9.54E-66 | 1.11E-63 |
| RPUSD1 | 0.774059979 | 3.40E-07 | 1.31E-06 |
| RPUSD2 | -0.255370046 | 0.023147159 | 0.04179835 |
| RPUSD4 | -0.99378444 | 4.47E-12 | 2.75E-11 |
| RRAD | -0.903813433 | 3.15E-27 | 5.66E-26 |
| RRAGC | -0.287920983 | 0.013762244 | 0.026180045 |
| RRAGD | 0.911127021 | 4.61E-31 | 1.03E-29 |
| RRAS | 0.799703531 | 5.07E-06 | 1.68E-05 |
| RRBP1 | -1.477679297 | 3.41E-48 | 1.89E-46 |
| RRH | 3.411578627 | 4.36E-06 | 1.46E-05 |
| RRM1 | -1.084126443 | 5.59E-14 | 4.05E-13 |
| RRM2 | -0.971281683 | 1.95E-05 | 5.96E-05 |
| RRN3 | -0.811565171 | 6.77E-14 | 4.88E-13 |
| RRP12 | -0.819497952 | 9.18E-11 | 4.98E-10 |
| RRP1B | -0.30296744 | 0.00315795 | 0.006909668 |
| RRP7A | -1.330747133 | 1.44E-22 | 1.92E-21 |
| RRS1 | -0.500909529 | 0.007338248 | 0.014794047 |
| RSAD2 | 1.003798704 | 0.000866843 | 0.002087854 |
| RSBN1 | 0.396775741 | 0.004232729 | 0.008996287 |
| RSBN1L | -0.647997617 | 0.008488133 | 0.016884341 |
| RSF1 | -0.657065273 | 0.000389487 | 0.000992857 |
| RSFR | 1.971667243 | 0.000225426 | 0.000595065 |
| RSL1D1 | -1.071639835 | 3.14E-20 | 3.58E-19 |
| RSL24D1 | 0.521911831 | 4.96E-06 | 1.65E-05 |
| RSPH3 | -4.526419632 | 0.002901098 | 0.006393988 |
| RSPH9 | 2.590133525 | 0.002580675 | 0.00574178 |
| RSPO3 | 1.096591605 | 2.72E-18 | 2.74E-17 |
| RSPO4 | 0.965893245 | 0.002936462 | 0.006464593 |

| | | | |
|---|---|---|---|
| RSPRY1 | -0.580924537 | 3.31E-07 | 1.28E-06 |
| RSRC1 | -0.523362236 | 5.25E-05 | 0.000152827 |
| RSRC2 | -0.799756034 | 1.26E-15 | 1.03E-14 |
| RSRP1 | 0.904113302 | 4.59E-19 | 4.88E-18 |
| RSU1 | 0.688091252 | 7.76E-14 | 5.55E-13 |
| RTCA | -0.435419121 | 0.000113116 | 0.000311734 |
| RTCB | -0.482716725 | 2.03E-08 | 8.84E-08 |
| RTF1 | -0.604478195 | 2.83E-08 | 1.22E-07 |
| RTKN2 | 0.750041759 | 0.026030646 | 0.046407206 |
| RTN4 | -0.308376822 | 0.000751548 | 0.001828359 |
| RTN4IP1 | -0.603285152 | 0.000823308 | 0.001990466 |
| RTTN | -0.858936417 | 0.001550128 | 0.00358322 |
| RUNDC1 | -0.377409967 | 0.000404588 | 0.001028842 |
| RUNDC3B | 0.380028001 | 0.013700262 | 0.026080387 |
| RUNX3 | -0.776366244 | 0.022541679 | 0.040813441 |
| RUVBL2 | -0.489262132 | 1.69E-05 | 5.20E-05 |
| RWDD1 | 0.446259252 | 8.27E-05 | 0.000233077 |
| RWDD2A | -1.061248494 | 1.45E-13 | 1.02E-12 |
| RWDD2B | -0.562837127 | 0.000102706 | 0.000285243 |
| RWDD3 | 0.399910002 | 0.002074182 | 0.004686997 |
| RWDD4 | 1.045671791 | 1.31E-23 | 1.85E-22 |
| RXFP1 | 2.132832968 | 0.000546038 | 0.001358052 |
| RYK | 0.600251564 | 7.03E-11 | 3.86E-10 |
| RYR3 | 1.24391239 | 9.81E-09 | 4.41E-08 |
| S100A1 | 1.007562939 | 0.00838102 | 0.016700604 |
| S100A10 | 0.536183423 | 2.37E-07 | 9.28E-07 |
| S100A11 | 1.300954172 | 1.03E-29 | 2.14E-28 |
| S100A4 | 0.529117701 | 5.69E-08 | 2.37E-07 |
| S100A6 | 0.464369914 | 8.66E-05 | 0.000243624 |
| S100B | 1.138651885 | 8.73E-06 | 2.81E-05 |
| S100Z | 2.571600446 | 6.41E-25 | 9.92E-24 |
| S1PR1 | -0.706738416 | 2.16E-06 | 7.57E-06 |
| S1PR3 | 1.416188812 | 0.007708928 | 0.015494567 |
| SAAL1 | 0.428489338 | 0.000474604 | 0.001190627 |
| SACM1L | -0.986530602 | 2.74E-25 | 4.37E-24 |
| SAG | 1.728271205 | 8.83E-05 | 0.000248031 |
| SALL4 | 1.974639552 | 5.45E-25 | 8.50E-24 |
| SAMD11 | 1.708017068 | 1.21E-36 | 3.78E-35 |
| SAMD4A | 0.772855272 | 1.55E-09 | 7.49E-09 |
| SAMM50 | 0.21076399 | 0.028183216 | 0.049845969 |
| SAP18 | 0.308794661 | 0.01279058 | 0.024548056 |
| SAP30BP | -0.970030861 | 8.09E-21 | 9.53E-20 |
| SAR1A | 0.822715511 | 1.06E-25 | 1.73E-24 |
| SAR1B | 0.37867989 | 0.000373838 | 0.000955654 |
| SARAF | -0.409787093 | 5.04E-05 | 0.000147061 |
| SARDH | -1.881390613 | 2.01E-07 | 7.91E-07 |
| SARNP | 0.485475807 | 1.68E-07 | 6.67E-07 |
| SARS | -1.067223072 | 5.67E-30 | 1.19E-28 |

| | | | |
|---|---|---|---|
| SART3 | -0.752716872 | 7.54E-07 | 2.79E-06 |
| SAT1 | 0.275134861 | 0.004283872 | 0.009089352 |
| SAV1 | 0.621436746 | 5.41E-12 | 3.30E-11 |
| SBDS | 0.519898361 | 1.03E-09 | 5.07E-09 |
| SBF2 | -0.513945689 | 0.002854125 | 0.006296586 |
| SBNO2 | 0.741543442 | 5.61E-07 | 2.11E-06 |
| SBSPON | -0.660059691 | 0.006749805 | 0.01372787 |
| SC5D | 0.556010142 | 1.94E-06 | 6.85E-06 |
| SCAF8 | -0.87050088 | 4.17E-07 | 1.59E-06 |
| SCAMP1 | -0.361246985 | 0.001046843 | 0.00249626 |
| SCAMP4 | 0.51535746 | 1.81E-08 | 7.91E-08 |
| SCAMP5 | 0.387121616 | 6.24E-05 | 0.000179318 |
| SCAP | -0.5080225 | 0.000162439 | 0.000437115 |
| SCAPER | -0.486260329 | 0.003729344 | 0.008026799 |
| SCARB2 | -0.345998274 | 0.000495589 | 0.001240296 |
| SCARF2 | -0.837277201 | 3.55E-18 | 3.56E-17 |
| SCCPDH | -0.619134395 | 2.10E-07 | 8.25E-07 |
| SCD5 | 1.012073023 | 5.71E-16 | 4.81E-15 |
| SCFD1 | -0.51603139 | 3.24E-06 | 1.11E-05 |
| SCG3 | -0.421900876 | 0.002535902 | 0.005650488 |
| SCG5 | 1.798280156 | 2.50E-07 | 9.75E-07 |
| SCLT1 | 0.436644013 | 0.000554097 | 0.001376835 |
| SCLY | -0.954861414 | 1.66E-09 | 8.04E-09 |
| SCN4B | 1.713249564 | 3.48E-11 | 1.96E-10 |
| SCN5A | 0.618818149 | 8.45E-06 | 2.72E-05 |
| SCN8A | 1.417718723 | 2.46E-33 | 6.17E-32 |
| SCNN1G | 1.424598691 | 4.45E-30 | 9.41E-29 |
| SCOC | 0.95841594 | 5.41E-26 | 9.03E-25 |
| SCP2 | 0.975136704 | 7.32E-22 | 9.22E-21 |
| SCPEP1 | -0.913578332 | 1.64E-18 | 1.69E-17 |
| SCRN3 | -0.365938154 | 0.003685047 | 0.007945297 |
| SCUBE3 | 2.829018127 | 0.01350487 | 0.025762553 |
| SCX | 3.966447269 | 0 | 0 |
| SCYL2 | -0.54281373 | 7.83E-05 | 0.000221372 |
| SCYL3 | -0.487187227 | 6.25E-07 | 2.33E-06 |
| SDC1 | -0.929573109 | 5.67E-27 | 9.96E-26 |
| SDC2 | 0.636009607 | 4.47E-12 | 2.75E-11 |
| SDC4 | 1.131644109 | 1.45E-33 | 3.65E-32 |
| SDCBP | 0.767978763 | 3.60E-20 | 4.09E-19 |
| SDCBP2 | 2.752688021 | 4.61E-15 | 3.62E-14 |
| SDE2 | -1.332869988 | 5.86E-44 | 2.65E-42 |
| SDF2 | -0.256639505 | 0.018031296 | 0.033389834 |
| SDF2L1 | -0.320685563 | 0.003759517 | 0.008082782 |
| SDF4 | -0.709075541 | 1.72E-14 | 1.30E-13 |
| SDHA | -1.154991793 | 9.21E-34 | 2.34E-32 |
| SDHB | 0.227793737 | 0.028197117 | 0.049864066 |
| SDHC | 0.839923334 | 0.000983301 | 0.00235424 |
| SDHD | 0.751880433 | 4.78E-18 | 4.72E-17 |

| | | | |
|---|---|---|---|
| SDR42E1 | -0.547901477 | 0.020047485 | 0.036733004 |
| SEC11C | 1.087984563 | 1.44E-24 | 2.16E-23 |
| SEC13 | 0.743910553 | 1.90E-21 | 2.32E-20 |
| SEC14L1 | 0.295048604 | 0.004407194 | 0.009320458 |
| SEC22A | 0.216279692 | 0.008645109 | 0.017143905 |
| SEC22B | -0.266841562 | 0.003276992 | 0.007141395 |
| SEC23A | 0.455006048 | 1.63E-08 | 7.20E-08 |
| SEC23B | -0.804597198 | 1.64E-18 | 1.69E-17 |
| SEC23IP | -0.469356668 | 2.64E-06 | 9.16E-06 |
| SEC24B | -1.087799053 | 8.48E-18 | 8.24E-17 |
| SEC24D | -0.49996779 | 2.07E-06 | 7.29E-06 |
| SEC31A | -0.304464158 | 0.000312042 | 0.000806785 |
| SEC31B | -1.392025959 | 1.44E-10 | 7.64E-10 |
| SEC61A2 | -0.580569111 | 8.33E-10 | 4.13E-09 |
| SEC61B | 0.590224719 | 4.14E-06 | 1.39E-05 |
| SEC61G | 0.94640753 | 2.61E-16 | 2.27E-15 |
| SEC63 | -0.394853105 | 8.84E-06 | 2.84E-05 |
| SECISBP2 | 0.624695387 | 3.37E-07 | 1.30E-06 |
| SEH1L | -0.32350275 | 0.001099028 | 0.002612908 |
| SEL1L | -1.179191446 | 3.45E-24 | 5.04E-23 |
| SELENBP1 | 0.383978857 | 0.020734733 | 0.037854466 |
| SELENOF | 0.635338252 | 2.88E-07 | 1.11E-06 |
| SELENOI | -0.599552271 | 3.84E-10 | 1.96E-09 |
| SELENOK | 0.930884256 | 1.20E-22 | 1.60E-21 |
| SELENOM | 1.041558718 | 4.31E-36 | 1.28E-34 |
| SELENOO | -1.614914064 | 8.37E-14 | 5.96E-13 |
| SELENOP1 | 1.378372171 | 8.38E-35 | 2.28E-33 |
| SELENOS | 0.813822885 | 1.54E-23 | 2.16E-22 |
| SELENOT | 0.349761407 | 0.000446806 | 0.001128814 |
| SELENOW | 1.93748902 | 2.03E-67 | 2.49E-65 |
| SEMA3A | 0.798576955 | 0.020221299 | 0.037016551 |
| SEMA3C | -0.986051585 | 4.72E-16 | 4.02E-15 |
| SEMA3D | -0.977760664 | 8.92E-24 | 1.27E-22 |
| SEMA4B | -1.313535186 | 1.90E-13 | 1.33E-12 |
| SENP2 | 1.020253787 | 9.33E-15 | 7.16E-14 |
| SENP5 | 0.605797515 | 3.51E-12 | 2.19E-11 |
| SENP6 | 0.367625468 | 0.002240631 | 0.005031333 |
| SEPHS2L | 0.741286989 | 2.88E-16 | 2.49E-15 |
| SEPSECS | -0.420067897 | 0.002691297 | 0.005962529 |
| SEPT2L | 0.721035592 | 4.68E-17 | 4.31E-16 |
| SEPTIN11 | -0.385116784 | 2.20E-05 | 6.69E-05 |
| SEPTIN9 | 0.739752008 | 1.67E-07 | 6.67E-07 |
| SERF2 | 1.346286001 | 5.78E-10 | 2.91E-09 |
| SERHL | -0.544683505 | 0.000104267 | 0.000289223 |
| SERINC1 | -0.652083046 | 1.16E-20 | 1.35E-19 |
| SERINC2 | -0.887152904 | 0.001249133 | 0.002938915 |
| SERINC4 | -3.104785714 | 0.013534178 | 0.025800358 |
| SERP1 | 1.51550238 | 1.87E-74 | 3.10E-72 |

| | | | |
|---|---|---|---|
| SERPINB1 | 1.699561288 | 1.02E-21 | 1.27E-20 |
| SERPINB10 | -1.136837037 | 8.92E-34 | 2.29E-32 |
| SERPINB10B | 1.08409107 | 7.26E-07 | 2.70E-06 |
| SERPINB5 | -0.987220071 | 0.001194683 | 0.00282157 |
| SERPINB6 | -0.263843255 | 0.010579418 | 0.020621749 |
| SERPINE2 | -0.59407952 | 6.71E-06 | 2.20E-05 |
| SERPINF1 | -0.274899373 | 0.000299887 | 0.000777726 |
| SERPINF2 | -1.090600801 | 1.74E-12 | 1.11E-11 |
| SERPINI1 | -1.625524438 | 1.07E-31 | 2.47E-30 |
| SERTAD2 | 1.808247457 | 9.30E-59 | 7.61E-57 |
| SERTM1 | 0.80780727 | 1.24E-06 | 4.48E-06 |
| SESN1 | -1.308290713 | 2.58E-31 | 5.84E-30 |
| SESTD1 | -0.327371222 | 0.000102436 | 0.000284551 |
| SET | 0.817264915 | 3.36E-19 | 3.60E-18 |
| SETBP1 | 0.7058322 | 0.001765101 | 0.004039612 |
| SETD3 | -0.213793676 | 0.021905928 | 0.03980021 |
| SETD6 | -0.46477941 | 0.000153777 | 0.0004157 |
| SETD7 | 0.781738947 | 1.32E-13 | 9.26E-13 |
| SETDB1 | -1.55013508 | 9.15E-08 | 3.74E-07 |
| SETDB2 | -0.694414097 | 8.98E-09 | 4.05E-08 |
| SF3A1 | -0.43027365 | 4.18E-06 | 1.40E-05 |
| SF3A2 | 0.683488954 | 6.06E-12 | 3.68E-11 |
| SF3B1 | -0.902735061 | 8.89E-18 | 8.62E-17 |
| SF3B3 | -0.515465553 | 5.79E-10 | 2.92E-09 |
| SF3B4 | 0.597331692 | 0.001245084 | 0.00293093 |
| SF3B5 | 1.397915796 | 3.78E-39 | 1.31E-37 |
| SF3B6 | 0.698899029 | 3.38E-12 | 2.11E-11 |
| SFMBT1 | -0.64657054 | 2.44E-05 | 7.40E-05 |
| SFPQ | -0.971101033 | 1.53E-29 | 3.13E-28 |
| SFR1 | -0.559417332 | 0.018089011 | 0.033487583 |
| SFT2D1 | 0.608536666 | 1.30E-10 | 6.98E-10 |
| SFT2D2 | 0.76995776 | 0.000192732 | 0.000513549 |
| SFXN1 | -0.372134584 | 0.000651942 | 0.001601227 |
| SFXN2 | -0.410529993 | 0.018632832 | 0.034391289 |
| SFXN3 | 0.364752788 | 0.000620154 | 0.001529232 |
| SFXN4 | 0.561301116 | 2.76E-06 | 9.55E-06 |
| SGCB | -0.218754268 | 0.009123608 | 0.018007697 |
| SGF29 | -0.572774043 | 1.32E-08 | 5.88E-08 |
| SGK1 | -1.369194541 | 2.73E-48 | 1.52E-46 |
| SGK3 | -0.249201295 | 0.01483575 | 0.028006599 |
| SGMS1 | 0.367363118 | 1.81E-05 | 5.57E-05 |
| SGMS2 | 0.591743858 | 4.80E-11 | 2.67E-10 |
| SGPL1 | -0.955248069 | 1.28E-18 | 1.33E-17 |
| SGPP1 | -1.047078759 | 1.44E-08 | 6.41E-08 |
| SGSM3 | -1.355605589 | 7.65E-09 | 3.47E-08 |
| SGTB | 0.26962143 | 0.003493297 | 0.007564265 |
| SH2D4A | -2.048595317 | 6.80E-19 | 7.17E-18 |
| SH3BGRL | 1.067118037 | 1.68E-42 | 7.04E-41 |

| | | | |
|---|---|---|---|
| SH3BGRL3 | 1.262982383 | 4.31E-59 | 3.62E-57 |
| SH3BP4 | -1.311889585 | 1.60E-15 | 1.30E-14 |
| SH3D19 | 0.362413859 | 0.001824034 | 0.004165369 |
| SH3GL1 | -0.402339905 | 2.79E-05 | 8.40E-05 |
| SH3GL3 | -0.437694096 | 0.024575099 | 0.044159803 |
| SH3GLB1 | 0.376723391 | 0.000588344 | 0.001456078 |
| SH3GLB2 | -0.581379186 | 5.51E-05 | 0.000159814 |
| SH3KBP1 | 0.410571927 | 1.05E-06 | 3.83E-06 |
| SH3PXD2A | 0.381785576 | 0.006245493 | 0.012784449 |
| SH3RF1 | -1.07219399 | 4.10E-05 | 0.000121009 |
| SH3RF3 | 0.841417181 | 3.35E-11 | 1.89E-10 |
| SHC3 | 1.056788642 | 0.000103807 | 0.000288064 |
| SHCBP1 | -1.680840906 | 2.59E-05 | 7.82E-05 |
| SHFM1 | 1.517673418 | 3.40E-55 | 2.41E-53 |
| SHISA2 | 0.939339599 | 7.25E-27 | 1.26E-25 |
| SHISA4 | 1.725953367 | 3.09E-51 | 1.94E-49 |
| SHISA5 | 0.726392459 | 4.39E-10 | 2.23E-09 |
| SHLD1 | -0.541534932 | 0.004303129 | 0.00912309 |
| SHLD3 | 0.344454117 | 0.017199674 | 0.032008826 |
| SHMT1 | -1.330101059 | 9.94E-05 | 0.000276546 |
| SHOC2 | 0.428035272 | 0.00036776 | 0.000941886 |
| SHQ1 | -0.969781952 | 3.67E-20 | 4.16E-19 |
| SHROOM2 | -0.93060207 | 9.01E-05 | 0.000252498 |
| SHROOM3 | 1.245386949 | 3.81E-18 | 3.80E-17 |
| SHROOM4 | 0.802360111 | 4.31E-08 | 1.82E-07 |
| SIAH1 | -1.815199321 | 1.12E-53 | 7.70E-52 |
| SIAH3 | 2.22628729 | 0.003383766 | 0.007349337 |
| SIK1 | -0.533592787 | 1.07E-08 | 4.79E-08 |
| SIK3 | -0.521226477 | 0.027829392 | 0.049290734 |
| SIKE1 | 0.512703609 | 8.76E-11 | 4.77E-10 |
| SIL1 | -0.892118034 | 8.54E-08 | 3.51E-07 |
| SIN3A | -0.934160607 | 1.09E-10 | 5.85E-10 |
| SIN3B | -0.604175364 | 0.000119761 | 0.000328846 |
| SIPA1L1 | -0.657307028 | 2.01E-09 | 9.64E-09 |
| SIPA1L2 | -1.037841822 | 6.23E-08 | 2.59E-07 |
| SIRT3 | 0.377634093 | 0.016919898 | 0.031526323 |
| SIRT6 | 0.613654694 | 2.88E-09 | 1.36E-08 |
| SIRT7 | 1.234761895 | 6.61E-07 | 2.46E-06 |
| SIX1 | 1.325781874 | 7.74E-29 | 1.52E-27 |
| SIX4 | -2.139102813 | 0.000313302 | 0.000809735 |
| SKA2 | 0.767954867 | 3.30E-12 | 2.06E-11 |
| SKI | 0.442656679 | 0.000436309 | 0.001103724 |
| SKIV2L2 | -0.473325407 | 1.51E-05 | 4.69E-05 |
| SKP1 | 0.573858514 | 6.19E-09 | 2.83E-08 |
| SKP2 | -0.741915633 | 9.80E-16 | 8.10E-15 |
| SLAIN2 | -0.690543212 | 9.45E-07 | 3.45E-06 |
| SLAMF8 | 1.611501692 | 0.027288017 | 0.048445083 |
| SLBP | -1.734464092 | 2.37E-06 | 8.27E-06 |

| | | | |
|---|---|---|---|
| SLC10A7 | 0.395844896 | 7.48E-06 | 2.43E-05 |
| SLC11A2 | 0.876655346 | 6.85E-10 | 3.43E-09 |
| SLC12A2 | -1.975172836 | 1.29E-28 | 2.51E-27 |
| SLC12A4 | -1.027556895 | 4.23E-24 | 6.11E-23 |
| SLC12A7 | -1.483986341 | 1.77E-48 | 9.95E-47 |
| SLC13A1 | 3.180489044 | 0.011247643 | 0.021789903 |
| SLC13A3 | -1.199350255 | 9.90E-05 | 0.000275656 |
| SLC13A4 | 1.62753877 | 1.79E-10 | 9.44E-10 |
| SLC13A5 | 2.292514116 | 0.003429525 | 0.007440399 |
| SLC15A4 | -1.390302658 | 8.01E-31 | 1.76E-29 |
| SLC16A1 | -1.02904491 | 5.85E-32 | 1.37E-30 |
| SLC16A10 | -0.932535956 | 2.43E-06 | 8.45E-06 |
| SLC16A3 | -0.745186172 | 8.53E-15 | 6.58E-14 |
| SLC16A5 | 2.449557654 | 3.47E-05 | 0.000103257 |
| SLC17A5 | -0.206661656 | 0.02317599 | 0.041839296 |
| SLC17A9 | 0.748995343 | 4.22E-05 | 0.000124166 |
| SLC18B1 | 0.645402932 | 6.68E-11 | 3.67E-10 |
| SLC19A2 | -0.879487771 | 0.020112731 | 0.03684262 |
| SLC1A4 | -0.955739974 | 6.50E-11 | 3.58E-10 |
| SLC1A7 | 1.298792913 | 0.014055758 | 0.026682384 |
| SLC20A2 | -0.284373358 | 0.004975676 | 0.010410928 |
| SLC22A3 | -2.68716955 | 4.80E-22 | 6.13E-21 |
| SLC23A2 | 0.379830976 | 0.017327532 | 0.032218305 |
| SLC24A3 | -0.472404783 | 0.005660947 | 0.011695332 |
| SLC25A1 | -0.247871883 | 0.002384089 | 0.00533408 |
| SLC25A11 | 0.738596446 | 6.83E-05 | 0.000195097 |
| SLC25A13 | -0.36929368 | 0.001310837 | 0.003065933 |
| SLC25A14 | 1.071662605 | 6.30E-35 | 1.74E-33 |
| SLC25A16 | 1.563948061 | 1.44E-60 | 1.28E-58 |
| SLC25A19 | -1.495582753 | 8.74E-06 | 2.81E-05 |
| SLC25A20 | 0.909057625 | 9.57E-25 | 1.46E-23 |
| SLC25A24 | -0.742993305 | 3.83E-18 | 3.82E-17 |
| SLC25A25 | 0.938421823 | 5.16E-21 | 6.13E-20 |
| SLC25A26 | 0.478595941 | 0.002079447 | 0.004698113 |
| SLC25A29 | -0.99033196 | 7.74E-06 | 2.51E-05 |
| SLC25A3 | -0.208487329 | 0.006785753 | 0.013794354 |
| SLC25A30 | -0.954303848 | 7.07E-06 | 2.31E-05 |
| SLC25A32 | 0.551612202 | 2.82E-06 | 9.73E-06 |
| SLC25A33 | -0.683631457 | 9.85E-10 | 4.85E-09 |
| SLC25A37 | 1.425414841 | 9.00E-35 | 2.45E-33 |
| SLC25A38 | 0.914242061 | 6.95E-07 | 2.59E-06 |
| SLC25A39 | 0.606509083 | 1.32E-09 | 6.42E-09 |
| SLC25A46 | -1.037805675 | 3.17E-16 | 2.74E-15 |
| SLC25A6 | -0.235196191 | 0.003394299 | 0.007369857 |
| SLC26A2 | -1.374146657 | 0.000222802 | 0.000588482 |
| SLC26A5 | 0.240156813 | 0.024492204 | 0.044022486 |
| SLC27A1 | -1.088622999 | 1.64E-05 | 5.08E-05 |
| SLC27A3 | -0.824721416 | 0.003736134 | 0.008037596 |

| | | | |
|---|---|---|---|
| SLC27A4 | -0.840711244 | 1.23E-15 | 1.01E-14 |
| SLC2A1 | -1.204769194 | 1.18E-35 | 3.43E-34 |
| SLC2A10 | -0.512594065 | 0.000110307 | 0.000304921 |
| SLC2A11 | -0.706779984 | 0.008967702 | 0.017734469 |
| SLC2A14 | -1.232738685 | 1.33E-52 | 8.83E-51 |
| SLC2A2 | 2.497198608 | 9.73E-07 | 3.55E-06 |
| SLC2A6 | -0.677683291 | 0.010219251 | 0.019965511 |
| SLC2A8 | -0.28980691 | 0.003332665 | 0.007251093 |
| SLC2A9 | 2.714186129 | 0.022155319 | 0.040213076 |
| SLC30A1 | 0.281041832 | 0.001468577 | 0.003412119 |
| SLC30A4 | 1.479726423 | 5.04E-60 | 4.34E-58 |
| SLC30A5 | -0.224803595 | 0.013864874 | 0.026349464 |
| SLC30A6 | -0.398963985 | 0.001096329 | 0.002607402 |
| SLC31A1 | 0.585955954 | 3.03E-12 | 1.90E-11 |
| SLC31A2 | -0.775520584 | 1.82E-06 | 6.43E-06 |
| SLC34A2 | 4.020433832 | 3.99E-27 | 7.11E-26 |
| SLC35B1 | 0.589446094 | 7.51E-13 | 4.96E-12 |
| SLC35B4 | 0.430623472 | 0.000251762 | 0.00066048 |
| SLC35C2 | 0.480678654 | 1.60E-05 | 4.95E-05 |
| SLC35E2B | 1.110459333 | 3.71E-36 | 1.11E-34 |
| SLC35E3 | 1.164341831 | 9.25E-51 | 5.72E-49 |
| SLC35E4 | -2.12422821 | 5.09E-15 | 3.99E-14 |
| SLC35F5 | -0.386779531 | 2.34E-05 | 7.13E-05 |
| SLC35G1 | 0.348402922 | 0.000809183 | 0.001957361 |
| SLC35G2 | 1.493294389 | 2.26E-54 | 1.57E-52 |
| SLC36A1 | -0.951052362 | 1.25E-08 | 5.55E-08 |
| SLC36A4 | 0.731799706 | 0.003439716 | 0.007460127 |
| SLC37A1 | -1.0723992 | 1.39E-11 | 8.16E-11 |
| SLC37A2 | 0.580382459 | 0.00226942 | 0.005090931 |
| SLC37A3 | 0.641731463 | 5.11E-16 | 4.33E-15 |
| SLC37A4 | -0.853052901 | 2.09E-07 | 8.20E-07 |
| SLC38A1 | -0.248816142 | 0.019601444 | 0.036022557 |
| SLC38A2 | -1.460165409 | 1.68E-74 | 2.86E-72 |
| SLC38A6 | 0.89843533 | 9.95E-12 | 5.94E-11 |
| SLC38A7 | -0.571774736 | 0.000219508 | 0.000580119 |
| SLC39A10 | 0.984581909 | 2.60E-16 | 2.26E-15 |
| SLC39A11 | -0.443300045 | 1.77E-05 | 5.45E-05 |
| SLC39A13 | 0.917125437 | 2.99E-34 | 7.88E-33 |
| SLC39A14 | 1.782192991 | 2.92E-11 | 1.67E-10 |
| SLC39A3 | -0.652202227 | 2.14E-05 | 6.55E-05 |
| SLC39A6 | -0.938204561 | 1.28E-14 | 9.72E-14 |
| SLC39A9 | 0.575797132 | 9.08E-10 | 4.49E-09 |
| SLC40A1 | 0.359119689 | 0.000308902 | 0.000799733 |
| SLC41A2 | 1.234153169 | 4.09E-64 | 4.31E-62 |
| SLC41A3 | 0.830668097 | 1.67E-11 | 9.71E-11 |
| SLC43A2 | -0.651437912 | 0.000339481 | 0.000873907 |
| SLC43A3 | -0.975688953 | 7.30E-19 | 7.67E-18 |
| SLC44A1 | 0.42691659 | 5.93E-05 | 0.00017093 |

| | | | |
|---|---|---|---|
| SLC44A2 | -1.001314145 | 0.023175895 | 0.041839296 |
| SLC45A3 | -1.602084157 | 1.82E-05 | 5.60E-05 |
| SLC46A1 | -3.390591323 | 0.005490784 | 0.011385315 |
| SLC46A2 | -0.348267038 | 0.006416268 | 0.013111569 |
| SLC46A3 | 3.91419571 | 3.97E-31 | 8.88E-30 |
| SLC47A1 | -0.796214882 | 0.001983689 | 0.004496728 |
| SLC48A1 | 0.845314516 | 5.66E-16 | 4.78E-15 |
| SLC49A3 | -0.816768894 | 4.98E-11 | 2.77E-10 |
| SLC49A4 | -0.411412302 | 0.000579142 | 0.001435392 |
| SLC4A10 | 2.193285254 | 0.003778597 | 0.008118668 |
| SLC4A1AP | -1.001925707 | 5.72E-11 | 3.16E-10 |
| SLC4A2 | 0.58157443 | 7.64E-05 | 0.000216231 |
| SLC4A7 | -0.742162476 | 7.09E-10 | 3.54E-09 |
| SLC5A1 | 2.088612642 | 3.28E-14 | 2.42E-13 |
| SLC5A6 | -0.785406469 | 8.51E-05 | 0.00023973 |
| SLC5A7 | 1.142473477 | 1.10E-07 | 4.46E-07 |
| SLC66A2 | 0.615899795 | 1.81E-11 | 1.05E-10 |
| SLC6A15 | -1.06732093 | 1.64E-05 | 5.08E-05 |
| SLC6A20 | 3.58803593 | 0.006423875 | 0.013123899 |
| SLC6A4 | -0.867663765 | 0.006023554 | 0.012376741 |
| SLC6A5 | 2.551769491 | 0.015045535 | 0.028382917 |
| SLC6A6 | -0.687712702 | 0.000467542 | 0.001174646 |
| SLC6A9 | -0.385947119 | 0.014575055 | 0.027560399 |
| SLC7A1 | -0.709814721 | 1.07E-05 | 3.40E-05 |
| SLC7A11 | -1.133003542 | 9.47E-22 | 1.18E-20 |
| SLC7A14 | 3.01673307 | 6.08E-05 | 0.00017501 |
| SLC7A3 | -1.404927535 | 1.31E-46 | 6.78E-45 |
| SLC7A5 | -0.544785676 | 5.75E-10 | 2.90E-09 |
| SLC7A6OS | -0.902872052 | 2.48E-16 | 2.16E-15 |
| SLC8A3 | 1.929283827 | 0.005127109 | 0.010693212 |
| SLC9A1 | -0.63017097 | 4.84E-07 | 1.83E-06 |
| SLC9A2 | 3.69902375 | 8.99E-07 | 3.29E-06 |
| SLC9A3R1 | -0.681042464 | 0.001147752 | 0.002717813 |
| SLC9A3R2 | -0.775371134 | 2.11E-07 | 8.29E-07 |
| SLC9A4 | 7.071816557 | 5.44E-07 | 2.05E-06 |
| SLC9A6 | -0.515481565 | 1.24E-05 | 3.90E-05 |
| SLC9A8 | 0.438934524 | 8.55E-05 | 0.000240697 |
| SLC9B2 | -0.387862488 | 0.0264359 | 0.047086457 |
| SLCO2A1 | -0.913925438 | 0.000618949 | 0.001526538 |
| SLCO4A1 | -1.795225891 | 1.16E-48 | 6.63E-47 |
| SLF2 | -0.860048988 | 9.69E-05 | 0.000270301 |
| SLITRK2 | -0.377933749 | 0.010727075 | 0.020882614 |
| SMAD1 | -0.424324319 | 0.005769889 | 0.011902312 |
| SMAD5 | 0.349842827 | 0.000522555 | 0.00130466 |
| SMAD6 | 1.232785724 | 8.96E-39 | 3.07E-37 |
| SMAD9 | 1.266627662 | 4.16E-11 | 2.33E-10 |
| SMAP1 | -0.534879537 | 1.61E-07 | 6.44E-07 |
| SMAP2 | 0.560881683 | 5.64E-11 | 3.12E-10 |

| | | | |
|---|---|---|---|
| SMARCA1 | -0.692727554 | 2.31E-05 | 7.03E-05 |
| SMARCA2 | -0.920503974 | 2.78E-27 | 5.02E-26 |
| SMARCA5 | -0.486116921 | 1.19E-08 | 5.29E-08 |
| SMARCAD1 | -0.675765676 | 2.54E-06 | 8.83E-06 |
| SMARCB1 | -0.570642769 | 1.40E-11 | 8.23E-11 |
| SMARCD1 | -0.452572479 | 0.00013294 | 0.000362181 |
| SMARCD2 | 0.409844221 | 4.18E-06 | 1.40E-05 |
| SMARCE1 | 1.109873816 | 7.54E-40 | 2.65E-38 |
| SMC6 | -0.320980499 | 0.020489593 | 0.037452199 |
| SMCO4 | 0.892525212 | 1.53E-23 | 2.16E-22 |
| SMDT1 | 1.109787618 | 1.31E-28 | 2.55E-27 |
| SMG6 | -0.788651895 | 2.90E-20 | 3.31E-19 |
| SMIM14 | 1.494478649 | 4.46E-46 | 2.24E-44 |
| SMIM15 | 0.835936571 | 2.58E-17 | 2.44E-16 |
| SMIM18 | 1.29337595 | 7.14E-14 | 5.14E-13 |
| SMIM19 | -0.509811343 | 3.29E-06 | 1.12E-05 |
| SMIM20 | 0.699495666 | 3.74E-12 | 2.32E-11 |
| SMIM29 | 0.35520348 | 0.001785066 | 0.004081866 |
| SMIM4 | 1.3984389 | 1.43E-32 | 3.43E-31 |
| SMIM5 | -2.116049399 | 0.006807375 | 0.01383047 |
| SMIM7 | 0.848830087 | 1.53E-17 | 1.46E-16 |
| SMIM8 | 0.932970431 | 3.42E-05 | 0.000101908 |
| SMKR1 | 0.985527037 | 0.000355039 | 0.000910852 |
| SMN | 0.464964034 | 7.02E-06 | 2.29E-05 |
| SMNDC1 | -0.769990328 | 7.97E-13 | 5.24E-12 |
| SMOC1 | -0.47934752 | 4.67E-06 | 1.56E-05 |
| SMOC2 | -0.815450718 | 1.63E-12 | 1.04E-11 |
| SMOX | 0.25510396 | 0.001007264 | 0.002407373 |
| SMPD1 | -1.60022687 | 4.54E-56 | 3.35E-54 |
| SMPD2 | -1.438255845 | 2.56E-16 | 2.23E-15 |
| SMPD3 | 2.844582023 | 0.001546079 | 0.003575687 |
| SMPD4 | -0.500727544 | 0.002679285 | 0.005940756 |
| SMPDL3B | -0.7537798 | 3.17E-06 | 1.08E-05 |
| SMU1 | -0.519395556 | 9.78E-09 | 4.40E-08 |
| SMURF1 | -0.353494001 | 0.013268896 | 0.025353075 |
| SMURF2 | -0.393321599 | 0.000165474 | 0.000445019 |
| SMYD4 | -1.097245976 | 1.79E-14 | 1.35E-13 |
| SMYD5 | -0.864943754 | 3.16E-12 | 1.98E-11 |
| SNAI1 | 1.170409301 | 2.40E-29 | 4.88E-28 |
| SNAP29 | -0.568811902 | 1.70E-10 | 9.00E-10 |
| SNAP47 | 0.490712502 | 5.40E-05 | 0.000156796 |
| SNAPC1 | -0.851309249 | 7.69E-13 | 5.07E-12 |
| SNAPC3 | -1.068572168 | 1.51E-07 | 6.04E-07 |
| SNAPC4 | -0.621966282 | 0.013053123 | 0.024995498 |
| SNAPC5 | 0.682354895 | 9.18E-11 | 4.98E-10 |
| SNAPIN | -0.410442895 | 0.005646494 | 0.011669019 |
| SNCAIP | -0.82277511 | 0.001535942 | 0.003554055 |
| SNF8 | 0.473785973 | 2.01E-07 | 7.91E-07 |

| | | | |
|---|---|---|---|
| SNIP1 | -0.6007046 | 1.33E-06 | 4.78E-06 |
| SNN | -1.080056835 | 1.80E-13 | 1.26E-12 |
| SNRK | -2.256596051 | 1.47E-44 | 6.81E-43 |
| SNRNP200 | -0.788303727 | 5.74E-05 | 0.000165793 |
| SNRNP25 | 0.606425138 | 1.07E-05 | 3.40E-05 |
| SNRNP27 | 0.83483905 | 1.47E-20 | 1.70E-19 |
| SNRNP35 | -1.236443334 | 9.73E-11 | 5.26E-10 |
| SNRNP48 | -1.073911351 | 1.27E-05 | 4.00E-05 |
| SNRPA1 | 0.433402562 | 0.000275161 | 0.000718262 |
| SNRPC | 1.012014797 | 4.25E-25 | 6.67E-24 |
| SNRPD1 | -0.388559709 | 0.0003589 | 0.000919714 |
| SNRPD3 | 0.599420828 | 3.06E-10 | 1.58E-09 |
| SNRPE | 0.948606222 | 1.09E-16 | 9.76E-16 |
| SNRPGP15 | 0.524298145 | 0.000216357 | 0.000572571 |
| SNRPN | 0.285062773 | 0.002176268 | 0.004902194 |
| SNTB2 | -1.415396692 | 4.22E-12 | 2.61E-11 |
| SNU13 | 0.432146311 | 5.29E-06 | 1.75E-05 |
| SNW1 | -0.654863129 | 2.94E-09 | 1.38E-08 |
| SNX1 | -0.243362968 | 0.006118009 | 0.012546164 |
| SNX10 | 0.289394825 | 0.01853701 | 0.034237674 |
| SNX11 | -0.9571487 | 3.41E-18 | 3.41E-17 |
| SNX12 | 0.574730992 | 1.37E-10 | 7.30E-10 |
| SNX14 | -0.832499628 | 1.20E-16 | 1.07E-15 |
| SNX16 | 0.311494576 | 0.00137179 | 0.003202022 |
| SNX17 | 0.826685853 | 1.02E-20 | 1.19E-19 |
| SNX18 | -1.416491667 | 4.23E-09 | 1.96E-08 |
| SNX2 | -0.317104207 | 0.00669431 | 0.013627236 |
| SNX21 | 0.376595628 | 0.001446856 | 0.003364527 |
| SNX22 | 1.616151773 | 0.00768226 | 0.015446342 |
| SNX24 | 0.730182436 | 1.12E-09 | 5.47E-09 |
| SNX27 | 0.250785428 | 0.008819468 | 0.017456561 |
| SNX3 | 0.634783157 | 1.61E-11 | 9.40E-11 |
| SNX32 | 0.28445487 | 0.005105234 | 0.010650857 |
| SNX33 | 0.353143901 | 0.007116419 | 0.014393804 |
| SNX5 | -0.516119437 | 7.99E-05 | 0.000225702 |
| SNX9 | -0.932656725 | 8.01E-23 | 1.08E-21 |
| SOAT1 | -0.348810381 | 0.027873417 | 0.049355069 |
| SOBP | 0.73680193 | 0.009066805 | 0.017907006 |
| SOCS1 | -1.387055611 | 5.00E-17 | 4.58E-16 |
| SOCS3 | -0.772098086 | 4.53E-11 | 2.53E-10 |
| SOCS4 | -1.342393864 | 2.17E-13 | 1.50E-12 |
| SOCS5 | -0.611062029 | 1.74E-09 | 8.39E-09 |
| SOCS6 | -0.602214318 | 1.15E-07 | 4.65E-07 |
| SOD1 | 0.286415922 | 0.00599827 | 0.012328517 |
| SOD2 | -0.32525477 | 0.002973674 | 0.0065391 |
| SOD3 | 0.553609178 | 2.61E-07 | 1.02E-06 |
| SORBS1 | -1.368637023 | 4.18E-09 | 1.94E-08 |
| SORCS2 | 3.161849678 | 0.003258946 | 0.007105486 |

| | | | |
|---|---|---|---|
| SORT1 | -1.100440492 | 0.021746423 | 0.039526261 |
| SOS2 | 0.342750611 | 0.010561761 | 0.020599146 |
| SOST | 2.438687474 | 0.007829846 | 0.015715191 |
| SOSTDC1 | 2.435464315 | 0.012126226 | 0.023362106 |
| SOX10 | 3.609829492 | 2.44E-10 | 1.28E-09 |
| SOX4 | 0.663071224 | 7.13E-13 | 4.72E-12 |
| SOX5 | -0.565384975 | 0.003549787 | 0.007675581 |
| SOX6 | 1.173129613 | 0.0147946 | 0.02794327 |
| SOX8 | 2.164413375 | 2.65E-101 | 1.03E-98 |
| SOX9 | -1.099173567 | 0.001522565 | 0.003526706 |
| SP3 | 0.30497235 | 0.003119568 | 0.006831185 |
| SP4 | -0.542476361 | 0.010693471 | 0.020823894 |
| SP9 | -2.361969298 | 0.000180933 | 0.000484292 |
| SPAG9 | -1.36926652 | 1.21E-24 | 1.83E-23 |
| SPART | -0.648539828 | 1.30E-08 | 5.77E-08 |
| SPAST | -0.950750252 | 4.02E-21 | 4.82E-20 |
| SPATA18 | -0.946887049 | 0.000666873 | 0.001635241 |
| SPATA20 | -0.940313094 | 0.001824889 | 0.004166621 |
| SPATA22 | 6.349825697 | 5.69E-08 | 2.37E-07 |
| SPATA5 | -0.707341405 | 6.86E-12 | 4.15E-11 |
| SPATA7 | -1.666726187 | 1.15E-30 | 2.50E-29 |
| SPATS2 | 0.248925114 | 0.009120294 | 0.018004798 |
| SPATS2L | -0.66206555 | 1.54E-06 | 5.48E-06 |
| SPC25 | -0.717558941 | 4.50E-05 | 0.000132169 |
| SPCS1 | 0.569787955 | 2.36E-08 | 1.02E-07 |
| SPCS2 | 0.372192648 | 0.016366337 | 0.030594313 |
| SPCS3 | 0.833356946 | 5.29E-24 | 7.62E-23 |
| SPDL1 | -0.448373721 | 0.006589743 | 0.013438528 |
| SPECC1L | 0.35741376 | 0.00608847 | 0.012496903 |
| SPEN | 0.407313638 | 2.56E-05 | 7.73E-05 |
| SPG11 | -0.378039222 | 0.020275148 | 0.037100137 |
| SPG21 | 0.275838437 | 0.00661606 | 0.013488651 |
| SPG7 | 0.7962927 | 9.78E-25 | 1.49E-23 |
| SPHK1 | 1.013251172 | 1.41E-17 | 1.35E-16 |
| SPI1 | 2.619939552 | 1.15E-160 | 1.43E-157 |
| SPIN1W | 1.326025705 | 4.05E-25 | 6.39E-24 |
| SPIN1Z | 0.749558923 | 5.94E-16 | 5.00E-15 |
| SPIRE1 | -0.363475105 | 0.002631937 | 0.005849131 |
| SPOCK2 | -0.957836298 | 0.015833362 | 0.029687828 |
| SPON1 | -0.937780696 | 1.16E-23 | 1.64E-22 |
| SPOPL | 0.263991071 | 0.015104628 | 0.028474632 |
| SPOUT1 | -0.498766882 | 4.87E-05 | 0.000142356 |
| SPP1 | 0.715426363 | 1.82E-10 | 9.60E-10 |
| SPP2 | 1.133559389 | 0.000406251 | 0.001032685 |
| SPPL2C | -0.997932377 | 5.14E-12 | 3.14E-11 |
| SPPL3 | 0.514628694 | 3.60E-07 | 1.38E-06 |
| SPR | -0.44851435 | 2.87E-05 | 8.61E-05 |
| SPRED2 | 1.07496525 | 2.56E-16 | 2.23E-15 |

| | | | |
|---|---|---|---|
| SPRY1 | -1.432541494 | 2.45E-55 | 1.76E-53 |
| SPRY2 | -1.119863202 | 2.69E-29 | 5.46E-28 |
| SPRY3 | 2.578852874 | 8.45E-28 | 1.57E-26 |
| SPRY4 | 2.673341455 | 0.018820235 | 0.034723041 |
| SPRYD7 | 0.301060267 | 0.002675314 | 0.005934857 |
| SPSB3 | -0.964421455 | 3.90E-17 | 3.62E-16 |
| SPSB4 | -1.006952781 | 8.90E-11 | 4.83E-10 |
| SPTAN1 | -1.764248531 | 7.61E-55 | 5.36E-53 |
| SPTBN1 | -1.477984523 | 1.95E-19 | 2.11E-18 |
| SPTLC1 | -1.223837592 | 1.50E-31 | 3.43E-30 |
| SPTLC2 | -0.930749346 | 5.30E-13 | 3.56E-12 |
| SPTLC3 | -1.220576856 | 1.80E-10 | 9.47E-10 |
| SPTSSA | 0.77524135 | 4.57E-18 | 4.52E-17 |
| SPTY2D1 | -1.081082503 | 7.81E-09 | 3.54E-08 |
| SQLE | -1.180814872 | 3.98E-29 | 7.94E-28 |
| SQSTM1 | -1.388836941 | 6.80E-63 | 6.69E-61 |
| SRA1 | -0.662291754 | 3.50E-07 | 1.34E-06 |
| SRBD1 | -0.348857948 | 0.00118664 | 0.002804036 |
| SRC | 0.356571974 | 0.007851545 | 0.015751764 |
| SREBF2 | -0.98308351 | 1.01E-32 | 2.43E-31 |
| SREK1IP1 | 0.513077773 | 0.006558989 | 0.013381833 |
| SRF | 1.343731741 | 7.07E-57 | 5.43E-55 |
| SRFBP1 | -0.351663768 | 0.009229358 | 0.018193682 |
| SRGAP1 | -0.364782027 | 0.001809343 | 0.004133902 |
| SRGAP3 | -0.307764048 | 0.026721902 | 0.047533581 |
| SRI | 0.369154248 | 0.000938378 | 0.00225065 |
| SRP68 | -0.857879199 | 2.07E-21 | 2.52E-20 |
| SRP72 | -0.323832279 | 0.000703405 | 0.001719184 |
| SRP9 | 1.334413756 | 1.53E-64 | 1.65E-62 |
| SRPK1 | 0.564774303 | 2.87E-09 | 1.36E-08 |
| SRPK2 | -0.735593896 | 0.003701042 | 0.007975989 |
| SRPRA | 0.518671666 | 3.38E-10 | 1.74E-09 |
| SRPRB | -0.531565406 | 1.37E-10 | 7.31E-10 |
| SRPX | -0.504287606 | 0.000123911 | 0.000339555 |
| SRPX2 | -1.199051599 | 1.10E-16 | 9.83E-16 |
| SRR | -0.940556026 | 0.00201069 | 0.004552607 |
| SRRD | -0.936934102 | 2.27E-08 | 9.83E-08 |
| SRRM1 | -0.67359242 | 7.70E-09 | 3.50E-08 |
| SRRM3 | 5.303457595 | 2.04E-07 | 8.03E-07 |
| SRSF1 | 0.901044139 | 1.52E-19 | 1.67E-18 |
| SRSF10 | 0.776727452 | 3.50E-23 | 4.79E-22 |
| SRSF2 | 0.364699615 | 2.36E-05 | 7.16E-05 |
| SRSF4 | 0.84065302 | 3.99E-06 | 1.34E-05 |
| SRSF5 | -0.887893936 | 3.90E-05 | 0.000115254 |
| SRSF5A | 0.654279121 | 1.88E-16 | 1.65E-15 |
| SRSF6 | 0.236263446 | 0.023069208 | 0.041690823 |
| SRXN1 | 0.986982137 | 2.47E-25 | 3.95E-24 |
| SS18 | 0.427629179 | 2.01E-07 | 7.91E-07 |

| | | | |
|---|---|---|---|
| SSBP1 | 0.548792383 | 3.18E-06 | 1.09E-05 |
| SSBP2 | 0.824756015 | 2.02E-17 | 1.91E-16 |
| SSBP4 | 2.341161715 | 2.25E-08 | 9.76E-08 |
| SSPN | 0.728846621 | 1.64E-08 | 7.24E-08 |
| SSR1 | 0.840069459 | 1.77E-23 | 2.46E-22 |
| SSR2 | -0.499830566 | 1.01E-07 | 4.11E-07 |
| SSTR3 | -2.039051025 | 0.000848305 | 0.002046895 |
| SSU72 | 0.821828424 | 1.56E-23 | 2.19E-22 |
| SSX2IP | 1.713533455 | 0.000133955 | 0.000364581 |
| ST13P5 | -0.516270342 | 7.02E-10 | 3.51E-09 |
| ST3GAL1 | 1.045992914 | 3.89E-26 | 6.52E-25 |
| ST3GAL2 | 0.659337937 | 1.38E-06 | 4.94E-06 |
| ST3GAL3 | 0.676606439 | 2.53E-11 | 1.45E-10 |
| ST3GAL4 | 0.871891476 | 7.69E-22 | 9.64E-21 |
| ST3GAL5 | 0.603253504 | 1.41E-05 | 4.41E-05 |
| ST6GAL1 | 0.897394226 | 0.02240176 | 0.040603376 |
| ST6GALNAC1 | 1.837658838 | 5.10E-05 | 0.000148681 |
| ST6GALNAC2 | 2.802400642 | 2.04E-12 | 1.29E-11 |
| ST6GALNAC4 | 1.288351966 | 1.58E-44 | 7.28E-43 |
| ST7 | -1.120559323 | 5.69E-11 | 3.15E-10 |
| ST7L | -0.952450531 | 1.83E-19 | 2.00E-18 |
| ST8SIA5 | 1.341911022 | 6.86E-18 | 6.71E-17 |
| STAMBP | -0.387991758 | 0.003857709 | 0.008272954 |
| STAP1 | 4.229248693 | 0.005207054 | 0.010846632 |
| STAR | 2.358807283 | 3.48E-11 | 1.97E-10 |
| STARD3 | 0.677198669 | 9.65E-05 | 0.000269183 |
| STARD3NL | -0.683143919 | 9.30E-10 | 4.59E-09 |
| STARD4 | 0.532763659 | 6.65E-09 | 3.04E-08 |
| STARD5 | -0.371065499 | 0.000215184 | 0.000569799 |
| STAT3 | -0.824232373 | 4.99E-21 | 5.94E-20 |
| STAU1 | 0.23577376 | 0.007438744 | 0.014981096 |
| STBD1 | -0.51047276 | 1.79E-05 | 5.50E-05 |
| STC2 | 0.592172497 | 4.99E-07 | 1.89E-06 |
| STEAP4 | 0.430642141 | 0.005022093 | 0.010501582 |
| STIL | -1.253666866 | 0.006734867 | 0.01370364 |
| STIM1 | -0.764809638 | 0.001841392 | 0.004199361 |
| STIM2 | -1.111768306 | 3.28E-28 | 6.23E-27 |
| STK11 | 0.766160242 | 3.56E-06 | 1.21E-05 |
| STK11IP | -1.167989951 | 2.44E-16 | 2.14E-15 |
| STK17B | -1.284592801 | 4.89E-35 | 1.37E-33 |
| STK24 | -0.857074991 | 3.69E-10 | 1.89E-09 |
| STK25 | 0.92453732 | 2.60E-24 | 3.83E-23 |
| STK26 | -0.433988036 | 4.97E-08 | 2.09E-07 |
| STK3 | 0.357464151 | 3.08E-05 | 9.21E-05 |
| STK38 | -1.068958796 | 6.68E-12 | 4.05E-11 |
| STK38L | 0.474051812 | 1.06E-06 | 3.84E-06 |
| STK39 | -0.442926728 | 0.017825233 | 0.033053284 |
| STK4 | 0.386723404 | 0.000281362 | 0.00073304 |

| | | | |
|---|---|---|---|
| STK40 | 1.210481567 | 1.71E-24 | 2.57E-23 |
| STMN1 | -0.307172881 | 0.026511694 | 0.047203825 |
| STMN4 | 3.684954219 | 0.01545631 | 0.02906907 |
| STN1 | -1.212391519 | 3.55E-05 | 0.000105648 |
| STRADB | 1.299565264 | 6.66E-32 | 1.56E-30 |
| STRAP | -0.508141269 | 2.73E-09 | 1.29E-08 |
| STRBP | -0.434734188 | 0.01039881 | 0.020292983 |
| STRIP1 | -1.093102019 | 1.76E-20 | 2.03E-19 |
| STRN | -0.966868063 | 5.39E-11 | 2.99E-10 |
| STRN3 | -0.514882459 | 0.000566178 | 0.001405056 |
| STT3A | -0.413701728 | 0.001206576 | 0.002847184 |
| STT3B | -0.680539495 | 2.24E-12 | 1.42E-11 |
| STX10 | 0.24549004 | 0.009274322 | 0.018271719 |
| STX16 | 0.791151897 | 8.83E-17 | 7.97E-16 |
| STX1A | 0.858633113 | 0.000206059 | 0.000546595 |
| STX3 | 0.884268284 | 0.02614365 | 0.046590336 |
| STX7 | 0.355473193 | 0.000386604 | 0.000986349 |
| STX8 | 0.276110277 | 0.004283844 | 0.009089352 |
| STXBP3 | -0.996288475 | 9.84E-17 | 8.84E-16 |
| STXBP6 | 0.631336737 | 7.30E-13 | 4.83E-12 |
| STYK1 | 1.107806959 | 0.004330495 | 0.009171095 |
| STYX | 0.461112153 | 0.002097235 | 0.004736725 |
| SUB1 | 0.803873278 | 2.14E-22 | 2.81E-21 |
| SUCLA2 | -0.270478094 | 0.00176019 | 0.004029051 |
| SUCLG2 | -0.813107696 | 7.84E-08 | 3.23E-07 |
| SUFU | 0.246912538 | 0.016254875 | 0.030415233 |
| SUGP1 | -1.10587816 | 7.90E-12 | 4.76E-11 |
| SULT1B | 0.513503767 | 0.001885444 | 0.004289747 |
| SULT1B1 | 0.38969754 | 0.001335467 | 0.003120452 |
| SULT4A1 | 1.086918039 | 1.38E-23 | 1.94E-22 |
| SULT6B1 | 0.836122314 | 0.001921374 | 0.00436419 |
| SUMF1 | -0.403091031 | 1.27E-05 | 4.00E-05 |
| SUMF2 | -1.43275926 | 7.10E-34 | 1.84E-32 |
| SUMO1 | 0.827440229 | 4.93E-24 | 7.11E-23 |
| SUMO2 | 1.911126082 | 2.47E-111 | 1.16E-108 |
| SUMO3 | 0.893133603 | 5.24E-21 | 6.22E-20 |
| SUN1 | -0.957704352 | 7.42E-14 | 5.33E-13 |
| SUPT3H | 0.637179637 | 0.002558783 | 0.00569587 |
| SUPT5H | -0.219549203 | 0.021348378 | 0.038875557 |
| SUPV3L1 | -0.337654449 | 0.014203075 | 0.026939464 |
| SURF1 | -0.782104994 | 2.10E-08 | 9.15E-08 |
| SURF2 | -0.977181549 | 5.03E-10 | 2.54E-09 |
| SURF4 | -0.281935608 | 0.00012848 | 0.000351016 |
| SURF6 | 0.66879311 | 1.43E-07 | 5.75E-07 |
| SUSD1 | 1.525201254 | 0.001306894 | 0.003058421 |
| SUSD5 | 3.175011776 | 1.14E-05 | 3.60E-05 |
| SUV39H1 | -1.541582611 | 0.001038757 | 0.002479155 |
| SVBP | 1.610663826 | 2.92E-101 | 1.10E-98 |

| | | | |
|---|---|---|---|
| SVIL | -0.447794578 | 3.17E-06 | 1.08E-05 |
| SVIP | 1.091737711 | 1.06E-32 | 2.54E-31 |
| SWI5 | 0.87623371 | 5.34E-08 | 2.23E-07 |
| SWT1 | 0.478803441 | 0.00546493 | 0.011340356 |
| SYAP1 | -0.307732451 | 0.00278974 | 0.006159542 |
| SYBU | -0.86808169 | 9.53E-10 | 4.70E-09 |
| SYDE2 | -0.274731169 | 0.013108281 | 0.025083452 |
| SYN3 | 1.3484068 | 0.001031852 | 0.002463106 |
| SYNC | 0.850225977 | 1.18E-14 | 8.94E-14 |
| SYNCRIP | -0.779958013 | 1.78E-22 | 2.36E-21 |
| SYNDIG1L | -1.037708332 | 0.011001783 | 0.021374533 |
| SYNE2 | -0.389485548 | 0.003290388 | 0.007168288 |
| SYNE3 | -0.879799035 | 0.000123725 | 0.000339114 |
| SYNGR1 | 1.518560903 | 2.77E-14 | 2.07E-13 |
| SYNGR3 | 1.445209075 | 1.07E-05 | 3.40E-05 |
| SYNJ2BP | 0.971275088 | 1.05E-14 | 8.00E-14 |
| SYNM | -0.622268774 | 1.81E-07 | 7.16E-07 |
| SYNPO | 0.608519605 | 0.000305075 | 0.000790427 |
| SYNRG | -1.046645397 | 9.73E-09 | 4.38E-08 |
| SYT11 | 0.314603848 | 0.004096027 | 0.008746772 |
| SYT14 | 2.277936678 | 0.00955397 | 0.018780795 |
| SYT8 | 1.115428978 | 0.008306741 | 0.016569595 |
| SYTL4 | 1.013998865 | 3.84E-19 | 4.10E-18 |
| SZRD1 | 1.14206738 | 4.33E-25 | 6.79E-24 |
| TAB1 | -1.04066268 | 1.96E-12 | 1.25E-11 |
| TAB2 | 0.395673547 | 0.007769525 | 0.015607949 |
| TAB3 | 1.288867105 | 1.01E-12 | 6.61E-12 |
| TACC1 | 1.193488317 | 1.02E-24 | 1.55E-23 |
| TACC3 | -1.030941554 | 5.38E-31 | 1.20E-29 |
| TACSTD2 | 4.90153909 | 1.16E-27 | 2.14E-26 |
| TADA1 | -0.968808881 | 4.15E-13 | 2.81E-12 |
| TAF12 | 0.64569627 | 1.14E-12 | 7.40E-12 |
| TAF13 | 0.71644795 | 6.17E-17 | 5.62E-16 |
| TAF1A | -1.205942592 | 3.20E-16 | 2.76E-15 |
| TAF1B | -0.568404952 | 8.38E-07 | 3.09E-06 |
| TAF3 | 0.734643007 | 3.01E-15 | 2.40E-14 |
| TAF4 | -0.914026351 | 0.000823986 | 0.001991752 |
| TAF4B | -0.976371173 | 0.003056929 | 0.006702658 |
| TAF7 | -1.029551712 | 2.88E-20 | 3.29E-19 |
| TAF9 | -0.374202574 | 0.00019372 | 0.00051598 |
| TAFA5 | 0.76060388 | 2.02E-05 | 6.19E-05 |
| TAGLN | 1.132841017 | 3.58E-26 | 6.03E-25 |
| TAGLN2 | 0.558198503 | 5.02E-05 | 0.000146501 |
| TAL1 | -2.102446083 | 0.014003728 | 0.026594756 |
| TAL2 | 5.278657676 | 0.000531576 | 0.00132475 |
| TALDO1 | -0.967566426 | 1.42E-23 | 2.00E-22 |
| TAMM41 | -0.581020122 | 7.09E-05 | 0.00020189 |
| TANC1 | 0.522251274 | 0.000184985 | 0.000494361 |

| | | | |
|---|---|---|---|
| TANGO2 | 1.063185701 | 2.69E-22 | 3.51E-21 |
| TANGO6 | -1.202151277 | 9.29E-11 | 5.04E-10 |
| TANK | 0.397562335 | 0.000114187 | 0.000314303 |
| TAOK1 | -0.974431199 | 6.42E-12 | 3.90E-11 |
| TAOK3 | 0.272524664 | 0.00498989 | 0.010437456 |
| TAP1 | -1.264386239 | 0.008516133 | 0.016927651 |
| TAPT1 | -0.6451703 | 4.81E-11 | 2.67E-10 |
| TARBP1 | -1.180647614 | 1.14E-09 | 5.58E-09 |
| TARDBP | -0.8932543 | 2.34E-18 | 2.37E-17 |
| TARP | 3.427589971 | 0.027512457 | 0.048786604 |
| TARS | -1.164938139 | 2.90E-35 | 8.12E-34 |
| TARSL2 | -1.327958302 | 6.11E-28 | 1.14E-26 |
| TASL | 3.793238475 | 0.000597467 | 0.001476678 |
| TASP1 | -0.570194485 | 6.52E-07 | 2.43E-06 |
| TATDN3 | 0.587442783 | 1.21E-06 | 4.37E-06 |
| TAX1BP3 | 1.170620297 | 1.50E-59 | 1.27E-57 |
| TAZ | 1.533701917 | 2.57E-66 | 3.02E-64 |
| TBC1D14 | -0.358042515 | 0.007091576 | 0.014349962 |
| TBC1D15 | -0.833826498 | 5.98E-13 | 3.99E-12 |
| TBC1D16 | -0.50269 | 0.022067313 | 0.040061295 |
| TBC1D2 | -0.581034541 | 0.007287454 | 0.014706915 |
| TBC1D20 | -0.991184303 | 2.33E-10 | 1.22E-09 |
| TBC1D22A | 0.280314928 | 0.005329998 | 0.011082351 |
| TBC1D22B | -0.42653787 | 0.0033203 | 0.007226504 |
| TBC1D23 | -0.690337401 | 6.21E-13 | 4.13E-12 |
| TBC1D24 | -0.712275994 | 4.67E-05 | 0.000136659 |
| TBC1D32 | -1.3051994 | 3.66E-06 | 1.24E-05 |
| TBC1D4 | -1.03738146 | 3.34E-06 | 1.14E-05 |
| TBC1D7 | -0.490796096 | 2.60E-06 | 9.02E-06 |
| TBC1D8B | -0.586366358 | 0.003340054 | 0.00726252 |
| TBCA | 0.802796633 | 1.03E-13 | 7.31E-13 |
| TBCC | -1.295064732 | 2.12E-09 | 1.01E-08 |
| TBCCD1 | -1.703525554 | 1.36E-15 | 1.11E-14 |
| TBCD | -0.987472149 | 1.04E-15 | 8.57E-15 |
| TBCK | -1.090648253 | 1.39E-09 | 6.73E-09 |
| TBK1 | -0.320529029 | 0.007497403 | 0.015092522 |
| TBL1X | 1.284187061 | 7.33E-56 | 5.36E-54 |
| TBL1XR1 | 1.014129718 | 1.71E-26 | 2.92E-25 |
| TBL2 | 0.242099586 | 0.009515183 | 0.018716407 |
| TBL3 | -0.501712888 | 4.42E-07 | 1.68E-06 |
| TBP | -0.240136406 | 0.011291092 | 0.02187096 |
| TBRG4 | -0.299016133 | 0.00093389 | 0.002240282 |
| TBX15 | -1.040910927 | 0.003189266 | 0.006971455 |
| TBX22 | 4.013863924 | 3.25E-07 | 1.25E-06 |
| TBX3 | 1.369592081 | 7.91E-09 | 3.58E-08 |
| TBXA2R | 0.727901338 | 0.019117208 | 0.035213591 |
| TCAIM | 0.595118063 | 1.33E-07 | 5.37E-07 |
| TCEA1 | -0.453982451 | 1.84E-05 | 5.66E-05 |

| | | | |
|---|---|---|---|
| TCEA2 | 0.503169458 | 5.83E-05 | 0.000168352 |
| TCEA3 | -0.453851807 | 0.01692058 | 0.031526323 |
| TCEANC2 | -0.835998791 | 5.29E-08 | 2.21E-07 |
| TCEB1 | 0.985646612 | 1.76E-23 | 2.45E-22 |
| TCEB3 | -0.251673769 | 0.019019805 | 0.035057932 |
| TCERG1 | -0.495410019 | 7.65E-07 | 2.83E-06 |
| TCF12 | 0.552134072 | 8.49E-12 | 5.10E-11 |
| TCF15 | 2.471596792 | 2.05E-12 | 1.30E-11 |
| TCF24 | 0.723074975 | 0.021626181 | 0.039334009 |
| TCF3 | 0.424086153 | 4.15E-06 | 1.39E-05 |
| TCIM | -1.28618593 | 3.82E-21 | 4.60E-20 |
| TCIRG1 | -0.821135874 | 1.35E-05 | 4.22E-05 |
| TCN2 | -0.754911019 | 9.07E-10 | 4.48E-09 |
| TCOF1 | -0.260740326 | 0.002458919 | 0.00548705 |
| TCP1 | 0.378276053 | 1.37E-07 | 5.49E-07 |
| TCP11L1 | 0.253406388 | 0.022238232 | 0.040350639 |
| TCP11X2 | -0.180894339 | 0.024742109 | 0.04440707 |
| TCTN1 | -0.662561715 | 5.16E-05 | 0.000150264 |
| TCTN2 | 0.352738192 | 0.00186443 | 0.004246201 |
| TDG | -0.574937419 | 2.18E-11 | 1.26E-10 |
| TDP1 | -0.553825256 | 7.09E-05 | 0.000202018 |
| TDP2 | -0.265806579 | 0.012608215 | 0.024225454 |
| TDRP | 0.682796607 | 0.001167236 | 0.002760583 |
| TEAD1 | 0.525847283 | 0.000587321 | 0.001454339 |
| TEAD3 | -0.623097008 | 4.55E-08 | 1.92E-07 |
| TEC | -1.243844657 | 6.36E-05 | 0.000182744 |
| TECPR2 | -0.713418332 | 2.19E-07 | 8.58E-07 |
| TECTA | 2.196497831 | 0.001153729 | 0.002731014 |
| TEK | 1.998093006 | 6.11E-06 | 2.01E-05 |
| TEKT1 | -1.461638263 | 0.00015011 | 0.000406272 |
| TEKT2 | 1.833812793 | 3.67E-06 | 1.24E-05 |
| TEKT5 | -1.848194719 | 0.001679348 | 0.003860909 |
| TENM1 | 2.173248114 | 0.019172493 | 0.035305856 |
| TENM3 | -0.911053811 | 4.99E-07 | 1.89E-06 |
| TENT4B | 0.387383295 | 0.00068109 | 0.001668297 |
| TENT5B | -0.902826928 | 0.011434065 | 0.022125827 |
| TERF2 | -0.855214295 | 4.52E-14 | 3.31E-13 |
| TERF2IP | 0.621100698 | 1.50E-12 | 9.64E-12 |
| TES | -0.337878032 | 0.00017629 | 0.00047233 |
| TEX10 | -0.524424631 | 3.66E-08 | 1.56E-07 |
| TEX12 | 3.849528607 | 8.62E-10 | 4.27E-09 |
| TEX264 | 0.551346359 | 2.50E-09 | 1.19E-08 |
| TEX30 | -1.155398102 | 1.75E-10 | 9.23E-10 |
| TFB1M | -0.642640684 | 0.000475819 | 0.001193234 |
| TFB2M | -0.91021818 | 2.47E-15 | 1.98E-14 |
| TFDP1 | 0.285904172 | 0.001141912 | 0.002704927 |
| TFG | -0.333982439 | 0.000372964 | 0.000953778 |
| TFIP11 | -0.740362598 | 2.01E-11 | 1.16E-10 |

| | | | |
|---|---|---|---|
| TFPI | 0.732146844 | 2.62E-09 | 1.24E-08 |
| TFRC | -0.974796723 | 2.52E-05 | 7.62E-05 |
| TGDS | -0.394205452 | 6.41E-05 | 0.000183762 |
| TGFB1 | 0.903452159 | 8.88E-08 | 3.64E-07 |
| TGFB1I1 | 1.312843615 | 4.33E-26 | 7.25E-25 |
| TGFB2 | 0.353778239 | 0.00026539 | 0.000693554 |
| TGFBI | -1.257750416 | 5.19E-49 | 3.05E-47 |
| TGFBR1 | -0.331248443 | 0.000197349 | 0.000525235 |
| TGFBR3 | -0.393142546 | 0.018028619 | 0.033389425 |
| TGFBRAP1 | -0.913451385 | 1.01E-06 | 3.67E-06 |
| TGIF1 | -0.562743581 | 3.22E-10 | 1.66E-09 |
| TGIF2 | 0.673144586 | 0.006087721 | 0.012496903 |
| TGM2 | -0.787669355 | 1.86E-05 | 5.71E-05 |
| TGM2.1 | 0.416239931 | 5.09E-06 | 1.69E-05 |
| TGS1 | -0.792030694 | 3.06E-06 | 1.05E-05 |
| THADA | -0.728283585 | 1.16E-07 | 4.70E-07 |
| THAP12 | 0.872173685 | 6.92E-05 | 0.000197175 |
| THAP4 | -0.466079812 | 4.98E-06 | 1.66E-05 |
| THAP5 | -1.408889396 | 6.86E-35 | 1.88E-33 |
| THAP9 | -0.735691877 | 1.78E-06 | 6.30E-06 |
| THBS1 | -0.627847036 | 1.76E-08 | 7.71E-08 |
| THBS2 | -1.612424241 | 5.39E-92 | 1.67E-89 |
| THEMIS2 | 2.350997827 | 1.05E-06 | 3.83E-06 |
| THG1L | -0.587447026 | 2.26E-06 | 7.88E-06 |
| THNSL1 | -0.51387627 | 0.01940189 | 0.035699267 |
| THNSL2 | 0.940873477 | 1.45E-25 | 2.35E-24 |
| THOC3 | -0.747548536 | 4.33E-10 | 2.20E-09 |
| THOC5 | -0.316701563 | 0.000341128 | 0.000877981 |
| THOC7 | 0.724415923 | 9.17E-13 | 6.00E-12 |
| THOP1 | -1.14551583 | 1.90E-19 | 2.06E-18 |
| THRAP3 | 0.3088661 | 0.000351534 | 0.00090237 |
| THRB | 0.833722675 | 9.80E-16 | 8.11E-15 |
| THRSP | 4.097073844 | 1.58E-19 | 1.73E-18 |
| THSD4 | -1.29873566 | 6.15E-08 | 2.56E-07 |
| THSD7B | -1.142262981 | 0.007580129 | 0.015252274 |
| THUMPD2 | 0.459173381 | 0.000515325 | 0.001287318 |
| THUMPD3 | 0.334347115 | 5.78E-05 | 0.000166946 |
| THY1 | 1.724086411 | 2.41E-72 | 3.49E-70 |
| THYN1 | 0.495354295 | 7.46E-05 | 0.000211599 |
| TIA1 | 0.654594618 | 3.96E-12 | 2.45E-11 |
| TIAM2 | -0.814355935 | 1.27E-12 | 8.22E-12 |
| TICRR | -1.080987933 | 1.17E-05 | 3.71E-05 |
| TIMM13 | 0.943086051 | 2.42E-12 | 1.53E-11 |
| TIMM17B | 0.41919028 | 1.49E-05 | 4.64E-05 |
| TIMM22 | 0.394091025 | 7.05E-05 | 0.000200945 |
| TIMM8A | 0.393830146 | 0.000173211 | 0.000464815 |
| TIMMDC1 | 0.584829464 | 1.53E-08 | 6.75E-08 |
| TIMP2 | 0.558283214 | 1.19E-12 | 7.74E-12 |

| | | | |
|---|---|---|---|
| TIMP3 | 0.705222037 | 7.27E-10 | 3.63E-09 |
| TIMP4 | 0.787625453 | 1.77E-14 | 1.33E-13 |
| TIPARP | 0.60226626 | 1.21E-13 | 8.52E-13 |
| TIPIN | -0.412263834 | 1.05E-05 | 3.36E-05 |
| TIPRL | -0.656689546 | 1.48E-11 | 8.69E-11 |
| TJP1 | -1.56795832 | 2.17E-21 | 2.64E-20 |
| TJP2 | -0.492335247 | 6.29E-05 | 0.000180785 |
| TJP3 | -0.75850471 | 0.005521961 | 0.011442979 |
| TK1 | -1.059863262 | 2.38E-13 | 1.64E-12 |
| TK2 | 1.230202441 | 1.67E-43 | 7.32E-42 |
| TKFC | -1.020353802 | 3.54E-15 | 2.81E-14 |
| TKT | -1.06522893 | 1.48E-25 | 2.40E-24 |
| TLCD1 | 1.195169567 | 2.05E-31 | 4.67E-30 |
| TLCD2 | 0.691294802 | 1.30E-07 | 5.25E-07 |
| TLE1 | -1.113044829 | 9.14E-15 | 7.02E-14 |
| TLE4 | -0.483485249 | 0.006724149 | 0.013685929 |
| TLE5 | 1.36875126 | 7.59E-22 | 9.53E-21 |
| TLK1 | -1.479304883 | 3.03E-16 | 2.62E-15 |
| TLN1 | -1.249448723 | 4.01E-28 | 7.58E-27 |
| TLR15 | 1.790068135 | 0.026481987 | 0.047162364 |
| TLR1A | -1.284700098 | 0.001255312 | 0.002950407 |
| TLR1B | -1.151709532 | 6.95E-17 | 6.29E-16 |
| TLR21 | -0.970009703 | 0.00552009 | 0.011440846 |
| TLR2A | 1.170271499 | 1.73E-12 | 1.11E-11 |
| TLR2B | 4.181354053 | 0.006553737 | 0.013373126 |
| TM2D1 | 1.197347793 | 4.87E-34 | 1.27E-32 |
| TM2D2 | 0.303013201 | 0.00480417 | 0.010081562 |
| TM2D3 | -1.2975687 | 7.34E-22 | 9.23E-21 |
| TM4SF1 | 0.508529983 | 0.026070343 | 0.046471883 |
| TM4SF18 | 0.484269331 | 0.010881034 | 0.021158085 |
| TM7SF3 | -1.472936234 | 1.26E-17 | 1.21E-16 |
| TM9SF2 | -0.484472489 | 4.25E-07 | 1.62E-06 |
| TM9SF2L | -0.425579325 | 1.98E-07 | 7.79E-07 |
| TM9SF4 | -0.707353258 | 7.19E-16 | 6.02E-15 |
| TMA16 | 0.411512751 | 0.003205154 | 0.007000555 |
| TMC6 | -1.278667802 | 1.37E-19 | 1.51E-18 |
| TMCC1 | 0.95437091 | 6.80E-22 | 8.59E-21 |
| TMCC2 | 1.52737322 | 1.24E-13 | 8.74E-13 |
| TMCC3 | -2.461265713 | 9.99E-11 | 5.40E-10 |
| TMCO1 | -0.218201454 | 0.018582994 | 0.034317942 |
| TMCO4 | -0.655020052 | 0.007753051 | 0.015577158 |
| TMCO6 | 0.376714834 | 0.011683129 | 0.022569228 |
| TMED10 | 0.471764117 | 3.29E-10 | 1.69E-09 |
| TMED5 | 0.584021875 | 1.76E-10 | 9.29E-10 |
| TMED6 | 2.691368518 | 0.001026286 | 0.002450321 |
| TMEM100 | 0.952607478 | 0.015847515 | 0.029710265 |
| TMEM101 | -0.819429703 | 3.24E-09 | 1.52E-08 |
| TMEM104 | -1.294410408 | 7.42E-17 | 6.71E-16 |

| | | | |
|---|---|---|---|
| TMEM106B | 0.289101867 | 0.000477904 | 0.001198022 |
| TMEM106C | -1.320946741 | 3.95E-17 | 3.66E-16 |
| TMEM11 | -0.309406984 | 0.002921782 | 0.006435402 |
| TMEM117 | 0.731526827 | 0.015567989 | 0.029258865 |
| TMEM119 | 0.329471604 | 0.00148616 | 0.003450613 |
| TMEM120A | -0.928023661 | 7.93E-10 | 3.94E-09 |
| TMEM120B | -0.579242102 | 9.34E-08 | 3.81E-07 |
| TMEM123 | 0.400057583 | 4.03E-06 | 1.36E-05 |
| TMEM127 | 0.855787008 | 0.004653706 | 0.009800616 |
| TMEM128 | 0.263809465 | 0.009209407 | 0.018160284 |
| TMEM131 | -1.626623211 | 1.92E-47 | 1.03E-45 |
| TMEM131L | -1.080075065 | 2.61E-15 | 2.09E-14 |
| TMEM132A | -2.067375365 | 1.12E-50 | 6.84E-49 |
| TMEM132C | -0.990810671 | 1.71E-05 | 5.27E-05 |
| TMEM136-1 | -0.955690156 | 7.70E-10 | 3.84E-09 |
| TMEM138 | 1.806483782 | 2.76E-85 | 6.70E-83 |
| TMEM140 | -0.35403336 | 0.026336751 | 0.046916005 |
| TMEM144 | -0.694989036 | 4.33E-09 | 2.01E-08 |
| TMEM14A | 0.997990471 | 5.41E-16 | 4.58E-15 |
| TMEM150C | -0.330719782 | 0.010150317 | 0.019847952 |
| TMEM154 | 1.417902066 | 0.004484098 | 0.009471307 |
| TMEM161A | -1.097879163 | 8.59E-05 | 0.000241691 |
| TMEM161B | -0.510033046 | 0.002642108 | 0.005868856 |
| TMEM167A | 0.764676911 | 2.61E-16 | 2.27E-15 |
| TMEM169 | 0.904132527 | 1.61E-11 | 9.40E-11 |
| TMEM170A | 0.338538438 | 0.006571397 | 0.013405137 |
| TMEM170B | 0.978856771 | 2.97E-24 | 4.37E-23 |
| TMEM171 | -0.985624988 | 9.69E-12 | 5.79E-11 |
| TMEM174 | 1.108076387 | 0.015546801 | 0.029223084 |
| TMEM175 | 0.487946847 | 2.86E-05 | 8.60E-05 |
| TMEM177 | -1.134317963 | 1.08E-10 | 5.83E-10 |
| TMEM178B | 0.671413044 | 0.023517974 | 0.042422868 |
| TMEM181 | -0.287986907 | 0.007975895 | 0.015963536 |
| TMEM183A | 0.829929162 | 3.39E-21 | 4.09E-20 |
| TMEM184B | 0.926271456 | 3.01E-26 | 5.09E-25 |
| TMEM184C | 0.498902967 | 8.78E-10 | 4.35E-09 |
| TMEM185A | 0.596871396 | 3.43E-11 | 1.94E-10 |
| TMEM186 | -0.744515145 | 2.69E-08 | 1.16E-07 |
| TMEM189 | 0.316276813 | 0.001262443 | 0.002964603 |
| TMEM19 | 0.617512729 | 7.55E-09 | 3.43E-08 |
| TMEM192 | -0.453304833 | 0.000217747 | 0.000576041 |
| TMEM196 | -1.069574734 | 2.93E-06 | 1.01E-05 |
| TMEM199 | -0.614906558 | 6.53E-07 | 2.43E-06 |
| TMEM200A | 0.534058842 | 7.88E-09 | 3.57E-08 |
| TMEM203 | 1.665588757 | 1.54E-50 | 9.38E-49 |
| TMEM204 | 1.309446937 | 8.78E-34 | 2.26E-32 |
| TMEM209 | -0.520147005 | 2.64E-06 | 9.17E-06 |
| TMEM214 | -0.598350937 | 7.04E-10 | 3.52E-09 |

| | | | |
|---|---|---|---|
| TMEM220 | 1.049478512 | 1.60E-06 | 5.69E-06 |
| TMEM223 | 1.010704534 | 4.22E-25 | 6.64E-24 |
| TMEM229B | 0.464126042 | 0.000453296 | 0.001143291 |
| TMEM230 | 0.792957938 | 1.83E-14 | 1.38E-13 |
| TMEM231 | -0.34464823 | 0.003574145 | 0.007724564 |
| TMEM233 | 2.448921038 | 1.24E-06 | 4.48E-06 |
| TMEM237 | 0.545878904 | 8.32E-06 | 2.68E-05 |
| TMEM240 | 1.901074884 | 0.009062295 | 0.017900699 |
| TMEM241 | 1.592797548 | 4.76E-57 | 3.68E-55 |
| TMEM242 | 0.780815647 | 2.34E-11 | 1.34E-10 |
| TMEM243 | 0.924200028 | 6.81E-05 | 0.000194566 |
| TMEM245 | -1.025408421 | 1.00E-06 | 3.65E-06 |
| TMEM248 | 1.047069288 | 2.53E-31 | 5.72E-30 |
| TMEM251 | 1.832713521 | 2.14E-74 | 3.46E-72 |
| TMEM254 | 0.785613701 | 2.38E-22 | 3.12E-21 |
| TMEM255A | 1.071480831 | 1.12E-15 | 9.21E-15 |
| TMEM255B | 1.606114845 | 0.0032372 | 0.007062603 |
| TMEM258 | 0.81517455 | 1.76E-13 | 1.23E-12 |
| TMEM26 | -0.271952642 | 0.007059472 | 0.014291086 |
| TMEM260 | -1.14191714 | 1.84E-08 | 8.05E-08 |
| TMEM263 | 0.889134335 | 2.74E-26 | 4.64E-25 |
| TMEM267 | 0.944350397 | 5.00E-11 | 2.78E-10 |
| TMEM268 | 0.875315324 | 7.80E-13 | 5.13E-12 |
| TMEM35A | 0.812289569 | 8.45E-05 | 0.000238193 |
| TMEM37 | -1.629141997 | 5.10E-11 | 2.83E-10 |
| TMEM39A | 0.811990444 | 1.93E-19 | 2.09E-18 |
| TMEM41B | 0.75392423 | 6.63E-23 | 8.94E-22 |
| TMEM43 | -0.348108114 | 3.45E-06 | 1.17E-05 |
| TMEM45A | 0.464132909 | 5.02E-05 | 0.000146501 |
| TMEM47 | -0.778102898 | 0.013531306 | 0.025798501 |
| TMEM50B | -0.238878701 | 0.013026925 | 0.024955879 |
| TMEM61 | 1.487413385 | 5.20E-27 | 9.17E-26 |
| TMEM63A | -0.690599702 | 0.000131 | 0.000357324 |
| TMEM63B | -0.978241145 | 7.54E-14 | 5.40E-13 |
| TMEM63C | 0.532185501 | 0.025345429 | 0.045334264 |
| TMEM65 | 0.263553305 | 0.006955286 | 0.014107783 |
| TMEM67 | -0.66500867 | 0.009166388 | 0.018082663 |
| TMEM68 | 0.341490376 | 0.000869677 | 0.002094003 |
| TMEM71 | 1.723330325 | 1.77E-23 | 2.46E-22 |
| TMEM80 | 0.780677543 | 1.58E-06 | 5.62E-06 |
| TMEM82 | 3.020444606 | 0.004232327 | 0.008996287 |
| TMEM86A | -1.017876246 | 9.72E-06 | 3.11E-05 |
| TMEM9 | -1.345084114 | 7.85E-06 | 2.54E-05 |
| TMEM97 | 0.461233795 | 1.61E-05 | 4.98E-05 |
| TMEM9B | 0.979105677 | 5.45E-17 | 4.98E-16 |
| TMF1 | -0.497150813 | 3.90E-08 | 1.65E-07 |
| TMLHE | -0.443072443 | 0.005196858 | 0.010828712 |
| TMOD3 | 0.187096113 | 0.028128805 | 0.049756209 |

| | | | |
|---|---|---|---|
| TMOD4 | 0.728918024 | 0.014887355 | 0.028100114 |
| TMPPE | -0.3366832 | 0.014407594 | 0.0272703 |
| TMPRSS9 | -1.43745091 | 0.01113392 | 0.021588051 |
| TMSB15B | 0.835657696 | 1.91E-17 | 1.81E-16 |
| TMSB4X | 1.008158054 | 4.34E-20 | 4.90E-19 |
| TMTC1 | 0.231737676 | 0.017888718 | 0.033157433 |
| TMTC2 | 0.457489181 | 0.001889676 | 0.004298657 |
| TMTC4 | 1.221376941 | 9.25E-23 | 1.24E-21 |
| TMX1 | 1.672713774 | 0.000335502 | 0.000864154 |
| TNFAIP2 | 1.242960202 | 3.94E-27 | 7.03E-26 |
| TNFAIP3 | 0.971787581 | 2.97E-12 | 1.87E-11 |
| TNFAIP8 | 0.919523669 | 4.22E-21 | 5.07E-20 |
| TNFAIP8L3 | 1.214468277 | 1.01E-28 | 1.97E-27 |
| TNFRSF10B | 0.927851361 | 4.27E-25 | 6.71E-24 |
| TNFRSF11A | -0.669974064 | 0.000157872 | 0.000426054 |
| TNFRSF11B | -0.765823257 | 8.28E-07 | 3.05E-06 |
| TNFRSF13C | -4.058175409 | 0.019095385 | 0.035178162 |
| TNFRSF19 | -1.396597461 | 0.002399675 | 0.005366301 |
| TNFRSF1A | -0.665336659 | 4.13E-11 | 2.32E-10 |
| TNFRSF21 | 1.458612255 | 2.12E-74 | 3.46E-72 |
| TNFRSF6B | -1.179579388 | 7.25E-41 | 2.73E-39 |
| TNFSF11 | -2.246909428 | 5.54E-14 | 4.02E-13 |
| TNFSF13B | 3.60216458 | 2.83E-14 | 2.11E-13 |
| TNFSF15 | 0.4130685 | 6.34E-05 | 0.000181933 |
| TNFSF8 | 3.891447313 | 7.83E-117 | 4.43E-114 |
| TNIK | 0.727887261 | 7.35E-11 | 4.03E-10 |
| TNIP1 | 0.73532326 | 3.26E-12 | 2.04E-11 |
| TNIP2 | 1.974028792 | 1.90E-34 | 5.06E-33 |
| TNIP3 | 0.995097852 | 4.21E-17 | 3.90E-16 |
| TNKS | -0.76489039 | 2.50E-08 | 1.08E-07 |
| TNKS2 | -0.78473275 | 2.15E-12 | 1.36E-11 |
| TNN | 1.625415574 | 2.21E-16 | 1.94E-15 |
| TNNT3 | 1.693349636 | 4.96E-05 | 0.000144938 |
| TNPO1 | -0.564079583 | 3.86E-05 | 0.000114286 |
| TNR | 1.791199792 | 0.007410177 | 0.0149288 |
| TNRC6A | 1.018160511 | 5.19E-11 | 2.87E-10 |
| TNRC6B | -0.525620895 | 0.000130809 | 0.000356877 |
| TNS1 | 0.281621788 | 0.008145888 | 0.01627023 |
| TNS3 | 0.538792295 | 1.38E-07 | 5.55E-07 |
| TNS4 | 0.847202121 | 0.015669563 | 0.029421287 |
| TOB1 | -0.865897183 | 7.68E-16 | 6.41E-15 |
| TOB2 | -1.273472708 | 4.98E-12 | 3.05E-11 |
| TOE1 | -0.958609027 | 1.86E-11 | 1.08E-10 |
| TOGARAM1 | -0.672105616 | 2.38E-05 | 7.24E-05 |
| TOM1 | -1.417911134 | 7.85E-35 | 2.15E-33 |
| TOM1L1 | 0.575411641 | 2.94E-06 | 1.01E-05 |
| TOM1L2 | -0.606626146 | 0.00025051 | 0.000657322 |
| TOMM20 | 0.305991481 | 0.003607994 | 0.00778905 |

| | | | |
|---|---|---|---|
| TOMM22 | 0.669267859 | 8.01E-12 | 4.83E-11 |
| TOMM40L | 1.425119842 | 1.00E-41 | 3.96E-40 |
| TOMM6 | 0.961312765 | 1.27E-12 | 8.18E-12 |
| TOMM7 | 0.816356591 | 4.85E-10 | 2.45E-09 |
| TOMM70 | -0.873947602 | 1.08E-20 | 1.25E-19 |
| TOP2A | 0.675675511 | 3.19E-05 | 9.53E-05 |
| TOP2B | -0.33650103 | 0.002679026 | 0.005940756 |
| TOP3A | 0.574759328 | 1.67E-05 | 5.16E-05 |
| TOP3B | -0.525331671 | 6.18E-06 | 2.03E-05 |
| TOPBP1 | -0.70186866 | 0.000249787 | 0.000655805 |
| TOR1B | -0.45683754 | 6.94E-06 | 2.27E-05 |
| TOR2A | -0.324141412 | 0.01355896 | 0.025839178 |
| TP53BP1 | -0.877347929 | 1.46E-07 | 5.86E-07 |
| TP53BP2 | -0.91238642 | 5.41E-27 | 9.52E-26 |
| TP53I3 | -0.497245729 | 0.007287251 | 0.014706915 |
| TP53INP1 | -0.259660169 | 0.017368271 | 0.0322751 |
| TP53RK | -0.69235162 | 2.80E-06 | 9.69E-06 |
| TP63 | 0.829514002 | 8.00E-07 | 2.95E-06 |
| TPBG | -1.795621546 | 4.69E-10 | 2.37E-09 |
| TPCN1 | 0.724574831 | 5.43E-13 | 3.64E-12 |
| TPCN3 | -0.529706993 | 0.000636313 | 0.001566523 |
| TPD52 | 0.361053367 | 0.001217001 | 0.00286979 |
| TPD52L1 | 1.027952888 | 0.000526912 | 0.00131409 |
| TPGS1 | -0.570019993 | 6.83E-05 | 0.000195068 |
| TPGS2 | 0.692485157 | 0.000106932 | 0.000296133 |
| TPI1 | -0.280579087 | 0.003411122 | 0.007401654 |
| TPM1 | -0.410996902 | 6.57E-05 | 0.00018812 |
| TPM2 | 1.320818305 | 2.09E-42 | 8.72E-41 |
| TPM3 | 0.307852001 | 0.000525422 | 0.001311094 |
| TPM4 | 0.470915641 | 0.000160888 | 0.000433372 |
| TPMT | 0.337095488 | 0.002684175 | 0.005948688 |
| TPP1 | -1.192795996 | 2.36E-40 | 8.51E-39 |
| TPP2 | -0.676483918 | 1.31E-09 | 6.40E-09 |
| TPPP3 | -1.443440909 | 7.42E-05 | 0.000210684 |
| TPR | -1.024205371 | 3.42E-20 | 3.89E-19 |
| TPRA1 | -1.21917907 | 1.15E-17 | 1.11E-16 |
| TPRG1L | 1.371762812 | 2.54E-72 | 3.63E-70 |
| TPRKB | -0.568473938 | 1.09E-07 | 4.42E-07 |
| TPST1 | 1.260057763 | 9.10E-41 | 3.39E-39 |
| TPST2 | 1.440214283 | 6.58E-11 | 3.62E-10 |
| TPT1 | 1.199633271 | 4.56E-47 | 2.42E-45 |
| TRA2B | -1.072445271 | 7.65E-21 | 9.03E-20 |
| TRABD | -0.820287951 | 6.87E-20 | 7.65E-19 |
| TRABD2B | 1.34133294 | 0.000112973 | 0.000311426 |
| TRAF1 | -0.468461635 | 0.007853756 | 0.015752194 |
| TRAF2 | -0.726748682 | 3.71E-05 | 0.000110248 |
| TRAF3IP1 | 0.627408717 | 4.55E-05 | 0.000133512 |
| TRAF4 | -0.986464139 | 0.010642465 | 0.020732748 |

| | | | |
|---|---|---|---|
| TRAF6 | -2.070124017 | 5.79E-42 | 2.33E-40 |
| TRAF7 | -0.448172217 | 2.96E-07 | 1.15E-06 |
| TRAFD1 | -0.619504957 | 2.59E-06 | 8.99E-06 |
| TRAIL-LIKE | -0.680629043 | 0.000305999 | 0.000792518 |
| TRAM1L1 | 0.260218626 | 0.00246245 | 0.005494027 |
| TRANK1 | -1.010713766 | 0.006831945 | 0.013878314 |
| TRAP1 | -0.717856223 | 3.82E-12 | 2.37E-11 |
| TRAPPC11 | -0.673558009 | 1.96E-09 | 9.39E-09 |
| TRAPPC12 | -0.747644459 | 6.54E-08 | 2.71E-07 |
| TRAPPC2 | 0.493160305 | 2.97E-06 | 1.02E-05 |
| TRAPPC2L | -0.655802311 | 2.41E-05 | 7.33E-05 |
| TRAPPC3 | 0.693619448 | 6.14E-16 | 5.17E-15 |
| TRAPPC5 | -0.884095736 | 1.62E-06 | 5.75E-06 |
| TRAPPC6A | -0.374088774 | 0.006644213 | 0.013537423 |
| TRAPPC9 | -0.908026297 | 0.000173333 | 0.00046505 |
| TRDMT1 | -0.927635011 | 2.11E-12 | 1.34E-11 |
| TRERF1 | 1.120211984 | 2.11E-14 | 1.58E-13 |
| TRIAP1 | 1.269480384 | 2.29E-47 | 1.22E-45 |
| TRIB1 | -1.662107512 | 1.19E-33 | 3.01E-32 |
| TRIB2 | 0.403514 | 1.21E-05 | 3.83E-05 |
| TRIM13 | -2.159736074 | 3.24E-29 | 6.54E-28 |
| TRIM23 | -0.779495386 | 0.000161187 | 0.000434005 |
| TRIM25 | -0.481945148 | 0.008584741 | 0.017044083 |
| TRIM28 | -0.393050131 | 0.00570385 | 0.011778597 |
| TRIM35 | 0.34990031 | 8.57E-05 | 0.00024122 |
| TRIM36 | 1.969933577 | 0.006169597 | 0.012644324 |
| TRIM37 | -0.750923173 | 8.69E-07 | 3.19E-06 |
| TRIM45 | -1.199945956 | 1.20E-10 | 6.47E-10 |
| TRIM54 | -0.921464957 | 9.20E-10 | 4.54E-09 |
| TRIM8 | 0.902868062 | 2.21E-20 | 2.54E-19 |
| TRIP12 | -0.825067251 | 7.85E-14 | 5.61E-13 |
| TRIP13 | -1.040440737 | 4.87E-05 | 0.000142314 |
| TRIT1 | -0.486430268 | 0.010636914 | 0.020724904 |
| TRMT10C | -1.065850279 | 9.77E-25 | 1.49E-23 |
| TRMT11 | 0.739566545 | 2.18E-10 | 1.14E-09 |
| TRMT12 | -0.483572493 | 0.017015673 | 0.031686125 |
| TRMT1L | -0.468694198 | 3.01E-05 | 9.01E-05 |
| TRMT2A | -0.727212402 | 3.24E-05 | 9.68E-05 |
| TRMT44 | -0.740006316 | 5.38E-05 | 0.000156307 |
| TRMT5 | -1.075465908 | 3.31E-11 | 1.87E-10 |
| TRMT6 | -0.348181476 | 0.008504855 | 0.016907706 |
| TRMT61A | -0.876834352 | 3.46E-20 | 3.94E-19 |
| TRMT61B | -0.362323045 | 0.004282508 | 0.009089352 |
| TRMT9B | -1.625743439 | 4.89E-22 | 6.24E-21 |
| TRNT1 | -0.277441248 | 0.004267601 | 0.00906615 |
| TRPC1 | 0.793977844 | 5.50E-18 | 5.41E-17 |
| TRPC3 | 1.983706021 | 1.87E-19 | 2.03E-18 |
| TRPC4 | 0.610355601 | 0.00010525 | 0.000291712 |

| | | | |
|---|---|---|---|
| TRPC7 | 5.230684609 | 4.23E-05 | 0.000124511 |
| TRPM6 | 1.67977733 | 9.78E-05 | 0.000272759 |
| TRPM7 | -0.406607365 | 0.00028312 | 0.000736772 |
| TRPS1 | -0.729242282 | 4.87E-06 | 1.62E-05 |
| TRPV1 | 4.520615956 | 2.09E-08 | 9.11E-08 |
| TRRAP | 0.60150985 | 6.97E-08 | 2.89E-07 |
| TSC22D1 | -1.334303795 | 9.77E-38 | 3.22E-36 |
| TSC22D2 | -0.664577481 | 2.02E-09 | 9.70E-09 |
| TSC22D4 | 0.916365002 | 1.14E-19 | 1.26E-18 |
| TSEN2 | -0.937386576 | 2.26E-11 | 1.30E-10 |
| TSEN54 | -0.698267757 | 8.73E-08 | 3.58E-07 |
| TSG101 | 0.794572951 | 8.67E-26 | 1.42E-24 |
| TSHZ1 | 1.52548195 | 1.01E-10 | 5.46E-10 |
| TSNAX | 0.320383519 | 0.000232125 | 0.000612155 |
| TSPAN12 | 0.357988566 | 0.004290056 | 0.009098213 |
| TSPAN14 | -0.547024522 | 8.96E-08 | 3.67E-07 |
| TSPAN15 | 1.753581932 | 0.005947556 | 0.012235384 |
| TSPAN2 | 0.513292777 | 1.24E-06 | 4.48E-06 |
| TSPAN3 | -0.650787127 | 3.43E-08 | 1.46E-07 |
| TSPAN5 | 0.402724722 | 0.01168573 | 0.022571045 |
| TSPAN6 | -0.38202692 | 0.000100649 | 0.000279931 |
| TSPAN8 | -1.231068453 | 0.00153506 | 0.003552619 |
| TSPO | 1.102874615 | 4.40E-33 | 1.09E-31 |
| TSR3 | -0.803242761 | 2.36E-07 | 9.24E-07 |
| TSSC1 | -0.963952104 | 2.19E-14 | 1.64E-13 |
| TTC1 | 0.293972363 | 0.003658128 | 0.007892267 |
| TTC13 | -0.592324504 | 4.58E-06 | 1.53E-05 |
| TTC17 | -0.581287603 | 1.32E-06 | 4.76E-06 |
| TTC21B | -0.653318141 | 7.93E-05 | 0.000224143 |
| TTC28 | 0.50612174 | 0.002872025 | 0.006334019 |
| TTC30B | -1.335307711 | 1.74E-11 | 1.01E-10 |
| TTC39C | -1.279880713 | 6.13E-07 | 2.29E-06 |
| TTC4 | -0.53183437 | 3.63E-06 | 1.23E-05 |
| TTC7A | -0.673372754 | 7.54E-05 | 0.000213747 |
| TTC7B | -0.528270315 | 0.00921599 | 0.018169965 |
| TTC9 | 1.004644665 | 1.74E-20 | 2.01E-19 |
| TTI1 | -1.139180048 | 6.27E-06 | 2.06E-05 |
| TTI2 | -2.076957117 | 1.38E-18 | 1.43E-17 |
| TTK | -0.636715827 | 0.005416002 | 0.011247411 |
| TTL | 0.934554563 | 2.32E-28 | 4.44E-27 |
| TTLL1 | 0.406725442 | 0.012132159 | 0.023370226 |
| TTLL4 | -0.689112361 | 0.000429359 | 0.001087154 |
| TTLL5 | 0.801875355 | 2.99E-13 | 2.06E-12 |
| TTLL9 | 0.994460601 | 0.000406567 | 0.001033295 |
| TTR | 1.110947228 | 2.72E-10 | 1.42E-09 |
| TTYH3 | 0.356561246 | 3.39E-05 | 0.000101005 |
| TUB | 0.93888291 | 1.37E-16 | 1.21E-15 |
| TUBA1A | -1.051361179 | 7.23E-47 | 3.81E-45 |

| | | | |
|---|---|---|---|
| TUBA1B | -0.817186896 | 2.73E-18 | 2.75E-17 |
| TUBA1C | -0.696958031 | 1.62E-09 | 7.83E-09 |
| TUBA3E | -1.388192518 | 1.39E-37 | 4.53E-36 |
| TUBA8B | 3.653444799 | 1.65E-05 | 5.11E-05 |
| TUBB | -0.671065378 | 8.55E-12 | 5.14E-11 |
| TUBB1 | 1.215982697 | 9.91E-09 | 4.45E-08 |
| TUBB2A | -0.784901464 | 1.88E-18 | 1.92E-17 |
| TUBB2B | -1.122617421 | 1.66E-40 | 6.05E-39 |
| TUBB3 | -1.42313441 | 9.20E-18 | 8.90E-17 |
| TUBB4B | -1.535058833 | 4.50E-73 | 6.72E-71 |
| TUBB6 | -1.136984628 | 6.84E-37 | 2.16E-35 |
| TUBD1 | 1.693109328 | 2.08E-26 | 3.54E-25 |
| TUBG1 | -0.208996148 | 0.022406339 | 0.040606261 |
| TUBGCP2 | -0.596810079 | 4.89E-07 | 1.85E-06 |
| TUBGCP3 | -1.256488149 | 3.88E-18 | 3.86E-17 |
| TUBGCP4 | 0.302075326 | 0.002384665 | 0.005334492 |
| TUBGCP5 | -0.730316495 | 0.002999664 | 0.00659092 |
| TUBGCP6 | -1.551421158 | 5.07E-11 | 2.81E-10 |
| TULP4 | 1.499838069 | 0.005318747 | 0.011062189 |
| TVP23A | 1.558339024 | 0.001262285 | 0.002964603 |
| TWF1 | 0.89008619 | 8.09E-30 | 1.68E-28 |
| TWIST1 | -1.081722399 | 9.98E-22 | 1.24E-20 |
| TWIST2 | 2.364243825 | 1.62E-09 | 7.83E-09 |
| TXLNA | -0.571090703 | 4.11E-08 | 1.74E-07 |
| TXLNG | -0.70806669 | 1.92E-05 | 5.88E-05 |
| TXN | 0.664747167 | 1.80E-11 | 1.04E-10 |
| TXN2 | 0.453821573 | 2.40E-06 | 8.38E-06 |
| TXNDC11 | -0.357336433 | 0.002099358 | 0.004740733 |
| TXNDC12 | 0.281253414 | 0.013484552 | 0.025731016 |
| TXNDC15 | -1.065872696 | 3.85E-24 | 5.57E-23 |
| TXNDC16 | -0.340911646 | 0.018647361 | 0.034413432 |
| TXNDC17 | 1.04135399 | 3.73E-23 | 5.09E-22 |
| TXNDC5 | -0.219989771 | 0.024123646 | 0.043434705 |
| TXNL4A | 0.636927281 | 2.06E-08 | 8.98E-08 |
| TXNL4B | -1.494202956 | 4.23E-11 | 2.37E-10 |
| TXNRD3 | -0.88879278 | 1.74E-18 | 1.79E-17 |
| TYR | 1.380222056 | 0.027006745 | 0.047983749 |
| TYSND1 | -1.179989041 | 4.11E-10 | 2.10E-09 |
| TYW3 | -0.420289264 | 0.017129462 | 0.031889283 |
| TYW5 | -1.027134934 | 1.24E-10 | 6.64E-10 |
| UAP1 | 0.301474226 | 0.001285573 | 0.003013714 |
| UBA2 | -1.360383292 | 3.26E-43 | 1.40E-41 |
| UBA3 | -0.24773275 | 0.017347417 | 0.032242048 |
| UBA5 | -0.436341031 | 0.005885291 | 0.012120133 |
| UBA52 | 0.916939886 | 7.31E-19 | 7.67E-18 |
| UBA6 | -0.761728025 | 2.56E-08 | 1.10E-07 |
| UBAC2 | 1.275445036 | 8.53E-42 | 3.39E-40 |
| UBAP1 | -1.150334752 | 5.59E-12 | 3.40E-11 |

| | | | |
|---|---|---|---|
| UBAP2 | -0.540867674 | 5.69E-05 | 0.000164614 |
| UBASH3B | 0.495880871 | 0.028091249 | 0.049702715 |
| UBE2A | 0.320533917 | 9.42E-05 | 0.000263116 |
| UBE2D1 | 1.244150794 | 1.27E-40 | 4.65E-39 |
| UBE2D2 | 1.083192104 | 7.29E-27 | 1.27E-25 |
| UBE2D3 | 0.527173261 | 3.69E-14 | 2.72E-13 |
| UBE2E3 | 1.312533981 | 1.30E-36 | 4.03E-35 |
| UBE2F | 1.029507221 | 1.81E-26 | 3.08E-25 |
| UBE2G2 | 0.255603448 | 0.011162791 | 0.021634772 |
| UBE2H | 1.155697294 | 7.34E-57 | 5.61E-55 |
| UBE2I | 0.770319931 | 4.13E-20 | 4.67E-19 |
| UBE2J2 | -0.619293339 | 4.83E-12 | 2.96E-11 |
| UBE2K | -0.306510714 | 0.005468194 | 0.011345396 |
| UBE2L3 | 0.679099732 | 4.41E-13 | 2.98E-12 |
| UBE2N | 0.951035751 | 6.74E-25 | 1.04E-23 |
| UBE2O | -0.51379086 | 0.01678387 | 0.031297347 |
| UBE2Q1 | 0.655943418 | 3.77E-14 | 2.78E-13 |
| UBE2Q2 | 0.770567196 | 2.47E-15 | 1.98E-14 |
| UBE2QL1 | -1.533695523 | 2.17E-09 | 1.04E-08 |
| UBE2R2 | 0.385994625 | 3.82E-06 | 1.29E-05 |
| UBE2S | 0.616071598 | 0.010208631 | 0.019950498 |
| UBE2T | -0.396809992 | 0.023271989 | 0.042001445 |
| UBE2V1 | 1.2091114 | 4.08E-56 | 3.03E-54 |
| UBE2V2 | 0.269611544 | 0.008342735 | 0.01662919 |
| UBE2W | 0.384567391 | 0.000160505 | 0.000432426 |
| UBE2Z | -0.62336038 | 2.65E-06 | 9.17E-06 |
| UBE3A | -0.811033456 | 5.53E-13 | 3.70E-12 |
| UBE3D | 0.621889865 | 3.67E-05 | 0.000109041 |
| UBE4A | -0.289137339 | 0.001101021 | 0.002616273 |
| UBE4B | -0.523515212 | 0.001040651 | 0.002482803 |
| UBFD1 | 0.38751205 | 0.000132061 | 0.000360076 |
| UBIAD1 | -0.682512672 | 2.02E-07 | 7.96E-07 |
| UBL3 | 1.260783314 | 1.58E-48 | 9.00E-47 |
| UBL5 | 0.865454143 | 8.58E-11 | 4.68E-10 |
| UBL7 | -0.334430196 | 0.00362679 | 0.007828383 |
| UBLCP1 | 0.390645643 | 0.000103107 | 0.00028618 |
| UBN1 | -0.563727755 | 4.00E-06 | 1.35E-05 |
| UBN2 | -0.628921161 | 0.005791931 | 0.011940533 |
| UBQLN1 | -0.962093745 | 1.74E-35 | 5.00E-34 |
| UBR1 | -0.388998961 | 0.020160723 | 0.036920581 |
| UBR2 | -0.513141291 | 3.78E-06 | 1.28E-05 |
| UBR7 | 0.876780757 | 5.28E-29 | 1.05E-27 |
| UBTD1 | 0.784774178 | 3.71E-11 | 2.08E-10 |
| UBTD2 | -0.999557468 | 1.19E-05 | 3.76E-05 |
| UBTF | 0.342458957 | 0.021315218 | 0.038825574 |
| UBXN10 | -2.02463709 | 9.38E-19 | 9.76E-18 |
| UBXN2A | 0.549012547 | 1.27E-11 | 7.52E-11 |
| UBXN2B | -0.623541187 | 5.51E-07 | 2.07E-06 |

| | | | |
|---|---|---|---|
| UBXN6 | -0.618935127 | 5.92E-10 | 2.98E-09 |
| UCHL1 | -0.590334388 | 0.014314605 | 0.027116949 |
| UCHL5 | 0.258238246 | 0.022468713 | 0.040692172 |
| UCK1 | 0.504252828 | 3.30E-11 | 1.87E-10 |
| UCKL1 | 0.646421149 | 5.47E-07 | 2.06E-06 |
| UFD1L | 0.355005915 | 0.000861752 | 0.002076087 |
| UFM1 | 0.715674971 | 4.74E-11 | 2.64E-10 |
| UGDH | -0.590479936 | 1.19E-10 | 6.42E-10 |
| UGGT1 | -0.713417922 | 2.25E-13 | 1.56E-12 |
| UGGT2 | -0.818986524 | 1.18E-10 | 6.32E-10 |
| UHMK1 | 0.414379532 | 0.000279205 | 0.0007277 |
| UHRF1BP1L | 0.412956821 | 0.0015262 | 0.003533921 |
| ULK3 | -0.555429647 | 9.26E-05 | 0.000258882 |
| UMAD1 | -1.745823568 | 1.66E-35 | 4.76E-34 |
| UMPS | -0.414871017 | 1.87E-06 | 6.62E-06 |
| UNC119 | 0.760070955 | 8.35E-07 | 3.07E-06 |
| UNC13D | -1.255632991 | 1.79E-10 | 9.43E-10 |
| UNC45A | -1.247972023 | 1.50E-09 | 7.25E-09 |
| UNC50 | -0.946512822 | 2.45E-18 | 2.48E-17 |
| UNC5B | -0.25164813 | 0.00348281 | 0.00754396 |
| UNC5C | -1.016754283 | 0.000920267 | 0.00220916 |
| UNC93B1 | -0.708942185 | 2.42E-05 | 7.36E-05 |
| UNG | -1.522903439 | 4.66E-14 | 3.41E-13 |
| UNK | -0.526619463 | 0.023106893 | 0.041740464 |
| UPF1 | -1.023589612 | 1.79E-19 | 1.95E-18 |
| UPF3B | 0.845851094 | 5.13E-13 | 3.45E-12 |
| UPK3A | 1.50117403 | 0.001829202 | 0.004174365 |
| UPP1 | 0.443999922 | 4.00E-05 | 0.000117982 |
| UPRT | 0.616720361 | 1.61E-13 | 1.13E-12 |
| UQCR10 | 0.726457468 | 3.00E-09 | 1.41E-08 |
| UQCR11 | 0.847267734 | 1.82E-10 | 9.59E-10 |
| UQCRB | 1.282099596 | 1.90E-35 | 5.43E-34 |
| UQCRC1 | -0.542909434 | 8.19E-08 | 3.37E-07 |
| UQCRC2 | -1.30745903 | 5.06E-38 | 1.69E-36 |
| UQCRFS1 | 0.389233708 | 0.000182024 | 0.000487022 |
| UQCRHL | 0.547353726 | 2.01E-06 | 7.08E-06 |
| UQCRQ | 1.413961796 | 1.45E-33 | 3.65E-32 |
| URB1 | -0.590664976 | 0.001514592 | 0.00351063 |
| URB2 | -1.785140996 | 1.00E-21 | 1.25E-20 |
| URI1 | 0.803919597 | 2.28E-23 | 3.15E-22 |
| URM1 | 1.766674959 | 1.55E-67 | 1.93E-65 |
| USB1 | 1.137583016 | 7.79E-29 | 1.53E-27 |
| USE1 | -0.36788453 | 0.002741806 | 0.006062572 |
| USH2A | 2.322996487 | 0.002051683 | 0.004641558 |
| USMG5 | 0.887625355 | 8.39E-13 | 5.50E-12 |
| USO1 | -0.985839388 | 3.19E-29 | 6.44E-28 |
| USP1 | -1.131814528 | 2.31E-27 | 4.21E-26 |
| USP10 | 0.221295114 | 0.004151414 | 0.008849752 |

| | | | |
|---|---|---|---|
| USP12-like | 0.743259127 | 5.20E-14 | 3.80E-13 |
| USP14 | -0.44068501 | 0.001058557 | 0.002521209 |
| USP16 | 0.691674107 | 1.30E-11 | 7.65E-11 |
| USP19 | -1.177242903 | 2.26E-18 | 2.30E-17 |
| USP20 | -0.576033817 | 0.000134913 | 0.00036704 |
| USP24 | -0.593289384 | 0.000155931 | 0.000421188 |
| USP25 | -0.647721328 | 6.76E-10 | 3.39E-09 |
| USP3 | -0.538342928 | 2.30E-07 | 9.01E-07 |
| USP31 | 0.732673438 | 0.000407051 | 0.001034138 |
| USP37 | -0.353621634 | 0.025326419 | 0.045312187 |
| USP38 | -0.62809785 | 6.94E-08 | 2.88E-07 |
| USP4 | -1.303758423 | 3.42E-37 | 1.09E-35 |
| USP45 | -0.443952986 | 1.60E-05 | 4.97E-05 |
| USP46 | 0.966725267 | 7.17E-19 | 7.54E-18 |
| USP47 | -1.114203997 | 1.19E-30 | 2.60E-29 |
| USP48 | -0.770613338 | 2.88E-10 | 1.50E-09 |
| USP5 | -0.980004616 | 2.74E-22 | 3.56E-21 |
| USP54 | -1.56848798 | 0.001236501 | 0.002912742 |
| USP7 | -0.859230366 | 3.79E-13 | 2.58E-12 |
| USP8 | 0.247665296 | 0.022428714 | 0.040633288 |
| USPL1 | 1.043048378 | 2.01E-19 | 2.18E-18 |
| UST | 0.782058473 | 6.49E-12 | 3.93E-11 |
| UTP15 | 0.20624485 | 0.02223857 | 0.040350639 |
| UTP18 | -1.092220805 | 3.78E-22 | 4.86E-21 |
| UTP3 | -0.779345523 | 0.000387588 | 0.000988385 |
| UTP4 | -0.649726044 | 2.64E-11 | 1.51E-10 |
| UTP6 | -1.201617224 | 3.96E-28 | 7.49E-27 |
| UTRN | -0.972158818 | 2.21E-10 | 1.16E-09 |
| UTS2 | 2.761021218 | 0.007176416 | 0.014504364 |
| UTS2R | 1.000805515 | 0.007973201 | 0.015960492 |
| UXS1 | 0.52869863 | 1.12E-08 | 5.02E-08 |
| VAMP1 | -1.705167097 | 5.38E-05 | 0.000156213 |
| VAMP3 | 0.788631142 | 6.94E-19 | 7.30E-18 |
| VAMP4 | -0.631781043 | 3.58E-10 | 1.83E-09 |
| VAMP7 | 0.255816866 | 0.007702081 | 0.015483905 |
| VAPB | 0.387359115 | 7.12E-05 | 0.0002027 |
| VASH1 | 1.248830848 | 7.90E-25 | 1.21E-23 |
| VASN | -0.533642991 | 9.56E-10 | 4.71E-09 |
| VAT1 | -0.622952445 | 0.010906564 | 0.021198631 |
| VAV2 | 0.704408517 | 2.02E-11 | 1.17E-10 |
| VAV3 | -0.835656907 | 0.000127463 | 0.000348517 |
| VBP1 | 0.462728351 | 2.45E-07 | 9.58E-07 |
| VCAN | -0.409245752 | 0.000152581 | 0.000412713 |
| VCL | -1.182636656 | 8.01E-48 | 4.39E-46 |
| VCP | -0.872159187 | 5.20E-28 | 9.78E-27 |
| VDAC1 | 0.353423127 | 5.34E-05 | 0.000155249 |
| VEGFA | 2.227886322 | 3.13E-188 | 6.09E-185 |
| VEGFC | -0.750701999 | 4.97E-07 | 1.88E-06 |

| | | | |
|---|---|---|---|
| VEZF1 | 0.573280811 | 0.000111851 | 0.000308687 |
| VEZT | 0.27464101 | 0.006045625 | 0.012412709 |
| VGLL1 | 4.386937905 | 8.99E-08 | 3.68E-07 |
| VGLL3 | 0.292226177 | 0.000481933 | 0.001207009 |
| VIM | -0.934776562 | 2.06E-25 | 3.31E-24 |
| VIP | -0.891024345 | 0.003270458 | 0.007128299 |
| VIRMA | -1.37962332 | 1.91E-07 | 7.54E-07 |
| VLDLR | -2.019235251 | 1.72E-43 | 7.51E-42 |
| VMA21 | 0.265328754 | 0.009329025 | 0.018368072 |
| VOPP1 | 1.200071584 | 3.12E-25 | 4.94E-24 |
| VPREB3 | 7.199272528 | 1.85E-07 | 7.33E-07 |
| VPS11 | -0.929454595 | 1.75E-19 | 1.92E-18 |
| VPS13A | -0.647692459 | 0.000869897 | 0.002094163 |
| VPS18 | -1.55360394 | 6.31E-36 | 1.86E-34 |
| VPS25 | 0.678317848 | 3.68E-11 | 2.07E-10 |
| VPS26A | -0.431879499 | 1.14E-05 | 3.62E-05 |
| VPS26B | 0.629414707 | 0.007952399 | 0.015924942 |
| VPS26C | -0.359196954 | 0.000776087 | 0.001884011 |
| VPS29 | 0.602019861 | 5.21E-11 | 2.89E-10 |
| VPS33B | 0.999347043 | 3.24E-40 | 1.16E-38 |
| VPS35 | -0.872845124 | 3.36E-22 | 4.33E-21 |
| VPS35L | 0.236830997 | 0.010079535 | 0.019715926 |
| VPS36 | -0.786148427 | 6.02E-09 | 2.76E-08 |
| VPS37A | 0.301428654 | 0.000857443 | 0.002067844 |
| VPS37B | -0.475723672 | 0.027676933 | 0.049033481 |
| VPS37C | -0.457089116 | 4.95E-06 | 1.65E-05 |
| VPS39 | -1.173652902 | 2.02E-22 | 2.67E-21 |
| VPS41 | 0.458994159 | 3.79E-06 | 1.28E-05 |
| VPS45 | -1.200464907 | 2.76E-17 | 2.59E-16 |
| VPS50 | -0.718649589 | 8.96E-12 | 5.37E-11 |
| VPS51 | -1.672506379 | 4.40E-36 | 1.30E-34 |
| VPS54 | -1.391916079 | 6.31E-26 | 1.05E-24 |
| VPS72 | 1.100721502 | 6.39E-42 | 2.56E-40 |
| VRK1 | -0.761917544 | 3.12E-06 | 1.07E-05 |
| VRK2 | -1.228363577 | 1.87E-18 | 1.91E-17 |
| VRTN | -2.4520657 | 8.51E-08 | 3.49E-07 |
| VSIG10 | 1.059763958 | 1.14E-12 | 7.40E-12 |
| VSIG4 | 1.461147881 | 0.005096103 | 0.01063344 |
| VSNL1 | 0.669845195 | 1.52E-07 | 6.10E-07 |
| VSTM5 | 1.399491373 | 0.005190625 | 0.010817383 |
| VSX2 | 3.162404225 | 0.002109558 | 0.004761393 |
| VTA1 | -0.455115552 | 2.60E-07 | 1.02E-06 |
| VTG1 | 3.972465324 | 5.66E-13 | 3.78E-12 |
| VTG2 | 2.837554574 | 1.66E-05 | 5.14E-05 |
| VTI1B | -0.772891288 | 5.85E-09 | 2.68E-08 |
| VTN | 2.768145815 | 0.023112301 | 0.041740951 |
| VWA1 | 0.364956062 | 0.001048973 | 0.002500901 |
| VWA3A | 2.621577575 | 5.30E-06 | 1.76E-05 |

| | | | |
|---|---|---|---|
| VWA8 | 0.284970346 | 0.001710026 | 0.003925464 |
| VWA9 | -0.720935403 | 9.93E-07 | 3.61E-06 |
| VWC2 | -2.95446363 | 8.88E-05 | 0.000249134 |
| VWC2L | 3.813666051 | 0.002416854 | 0.005399388 |
| VWF | 0.44365317 | 0.01292963 | 0.024783463 |
| VXN | 3.804566251 | 0.010929529 | 0.021240228 |
| WAC | -0.381340165 | 0.004458646 | 0.00942194 |
| WAPL | -0.320877144 | 0.00226234 | 0.005076726 |
| WAS | 1.627819146 | 2.68E-17 | 2.53E-16 |
| WASF2 | 0.879998171 | 0.000639166 | 0.00157269 |
| WASH1 | -0.918357159 | 2.05E-06 | 7.21E-06 |
| WASHC3 | 0.894696231 | 1.33E-28 | 2.58E-27 |
| WASHC4 | -1.12194015 | 3.56E-18 | 3.56E-17 |
| WASHC5 | -0.735009989 | 6.66E-12 | 4.04E-11 |
| WASL | 0.472849219 | 1.17E-05 | 3.70E-05 |
| WBP1 | 1.739929768 | 8.49E-18 | 8.24E-17 |
| WBP2 | -0.367719782 | 0.000400496 | 0.001018816 |
| WBP2NL | 0.812183973 | 1.48E-25 | 2.40E-24 |
| WBSCR22 | -1.056917907 | 8.04E-26 | 1.32E-24 |
| WDCP | -1.025366299 | 6.47E-12 | 3.93E-11 |
| WDHD1 | -0.76340495 | 0.027302326 | 0.048451914 |
| WDR1 | -0.325663225 | 0.000112015 | 0.000309076 |
| WDR11 | -0.833330808 | 1.62E-09 | 7.83E-09 |
| WDR12 | -0.317684615 | 0.001153437 | 0.0027308 |
| WDR18 | -0.71662993 | 3.42E-07 | 1.32E-06 |
| WDR24 | -0.3418218 | 0.003351361 | 0.007282783 |
| WDR26 | -1.276078469 | 2.52E-13 | 1.74E-12 |
| WDR3 | -0.499147372 | 1.54E-07 | 6.15E-07 |
| WDR31 | 1.073328911 | 0.002998064 | 0.006589535 |
| WDR35 | -1.019354076 | 8.39E-10 | 4.16E-09 |
| WDR37 | -0.685960224 | 0.000353261 | 0.000906462 |
| WDR4 | -0.328669344 | 0.000199152 | 0.000529721 |
| WDR41 | -0.74469032 | 0.000125374 | 0.000343357 |
| WDR43 | -0.507606596 | 4.73E-11 | 2.64E-10 |
| WDR45B | 0.197359234 | 0.025662401 | 0.045840893 |
| WDR47 | -0.520489384 | 1.92E-06 | 6.79E-06 |
| WDR53 | -0.849849767 | 8.86E-12 | 5.32E-11 |
| WDR5B | 0.505408752 | 5.14E-06 | 1.71E-05 |
| WDR6 | -1.450063053 | 0.0008746 | 0.002104738 |
| WDR7 | -1.111924567 | 1.73E-06 | 6.14E-06 |
| WDR70 | -0.508042384 | 2.99E-05 | 8.96E-05 |
| WDR74 | -0.471241772 | 0.000150562 | 0.000407415 |
| WDR75 | -1.002080905 | 2.32E-30 | 4.96E-29 |
| WDR77 | -0.755922741 | 1.17E-20 | 1.36E-19 |
| WDR81 | -1.016655436 | 3.85E-14 | 2.83E-13 |
| WDR89 | -0.712727181 | 2.10E-09 | 1.00E-08 |
| WDR90 | -1.715232869 | 0.003676868 | 0.007928922 |
| WDR91 | -0.557459875 | 2.52E-06 | 8.76E-06 |

| | | | |
|---|---|---|---|
| WDSUB1 | 0.310491889 | 0.013893852 | 0.026400844 |
| WEE1 | 0.389626736 | 0.000292188 | 0.000758627 |
| WFDC1 | 1.361520931 | 5.80E-05 | 0.000167429 |
| WFDC2 | 0.618114124 | 0.004527403 | 0.00954942 |
| WFS1 | -1.123398521 | 2.13E-31 | 4.84E-30 |
| WHAMM | -0.481853644 | 0.006359366 | 0.013005793 |
| WIPI1 | 1.211248565 | 4.18E-58 | 3.27E-56 |
| WIPI2 | -0.187552701 | 0.020961679 | 0.038243106 |
| WISP1 | 0.397654146 | 6.60E-05 | 0.000188833 |
| WLS | -0.537271328 | 8.27E-10 | 4.11E-09 |
| WNT5B | -0.472959133 | 4.88E-05 | 0.000142527 |
| WNT6 | 1.672173844 | 5.23E-08 | 2.19E-07 |
| WNT9A | 0.977886127 | 2.64E-14 | 1.97E-13 |
| WRAP73 | -0.818731873 | 3.61E-06 | 1.22E-05 |
| WRN | -0.506363773 | 0.002622656 | 0.005829458 |
| WSB2 | 1.061981688 | 3.78E-44 | 1.72E-42 |
| WSCD1 | 2.098810547 | 0.00010909 | 0.000301678 |
| WT1 | 1.889291815 | 5.79E-05 | 0.000167334 |
| WTAP | -1.068678679 | 1.08E-21 | 1.34E-20 |
| WWC2 | -0.250917007 | 0.01989969 | 0.036501563 |
| WWP1 | 0.823462721 | 6.82E-15 | 5.30E-14 |
| XBP1 | 0.533255884 | 2.37E-12 | 1.50E-11 |
| XDH | 3.009057198 | 1.42E-06 | 5.09E-06 |
| XIRP1 | -2.605199838 | 1.47E-48 | 8.42E-47 |
| XKR5 | -1.445656826 | 0.004051507 | 0.008663953 |
| XKR8 | -0.759739492 | 0.005713649 | 0.011797041 |
| XPA | 1.687618918 | 1.10E-69 | 1.42E-67 |
| XPNPEP1 | -0.743148711 | 1.66E-11 | 9.69E-11 |
| XPNPEP3 | 0.450076205 | 2.90E-05 | 8.71E-05 |
| XPO1 | 0.370636588 | 7.64E-05 | 0.000216338 |
| XPO4 | -0.468676865 | 0.007843026 | 0.015736996 |
| XPO7 | 0.346504552 | 0.001790869 | 0.004094448 |
| XPOT | 0.427145427 | 7.22E-08 | 2.99E-07 |
| XPR1 | 1.262755685 | 1.82E-40 | 6.59E-39 |
| XRCC2 | -0.843528839 | 5.18E-08 | 2.17E-07 |
| XRCC5 | -0.688519981 | 1.13E-08 | 5.03E-08 |
| XRCC6 | -0.324223523 | 0.002235053 | 0.005020465 |
| XRN1 | -0.438446103 | 0.002778021 | 0.006134665 |
| XXYLT1 | 1.035125237 | 3.57E-23 | 4.89E-22 |
| YAE1 | 0.892486404 | 2.42E-11 | 1.39E-10 |
| YAF2 | 0.615723406 | 2.33E-07 | 9.12E-07 |
| YAP1 | -0.371093331 | 0.00038565 | 0.000984181 |
| YARS | -1.433887511 | 1.00E-50 | 6.15E-49 |
| YARS2 | -1.404982701 | 6.41E-25 | 9.92E-24 |
| YBEY | 0.8272389 | 1.26E-24 | 1.90E-23 |
| YBX1 | 1.98616216 | 1.70E-154 | 1.78E-151 |
| YBX3 | 0.944753023 | 1.82E-29 | 3.70E-28 |
| YEATS2 | -0.451484011 | 0.004801778 | 0.010078099 |

| | | | |
|---|---|---|---|
| YES1 | 0.921462332 | 2.27E-13 | 1.57E-12 |
| YIPF1 | -0.349540503 | 0.001104963 | 0.002624723 |
| YIPF3 | 0.526887077 | 2.63E-08 | 1.14E-07 |
| YIPF4 | 0.436219546 | 2.03E-09 | 9.71E-09 |
| YIPF6 | -0.267781752 | 0.013511124 | 0.025768418 |
| YLPM1 | -0.788873292 | 0.010218763 | 0.019965511 |
| YPEL1 | 0.490387264 | 3.55E-06 | 1.20E-05 |
| YPEL2 | 0.451258359 | 0.00541878 | 0.01125146 |
| YPEL5 | 0.441266325 | 9.29E-09 | 4.19E-08 |
| YRDC | 0.745549536 | 1.08E-14 | 8.22E-14 |
| YTHDC1 | -0.4665368 | 4.36E-06 | 1.46E-05 |
| YTHDC2 | -1.071902574 | 5.90E-09 | 2.70E-08 |
| YTHDF1 | 0.771005485 | 1.94E-08 | 8.50E-08 |
| YTHDF2 | 1.17781201 | 2.91E-22 | 3.78E-21 |
| YTHDF3 | -0.507110321 | 4.06E-06 | 1.37E-05 |
| YWHAB | 0.171963873 | 0.023732963 | 0.042776613 |
| YWHAE | 0.411636059 | 7.49E-07 | 2.77E-06 |
| YWHAG | 1.076142197 | 1.65E-35 | 4.76E-34 |
| YWHAH | -0.374766191 | 6.99E-06 | 2.29E-05 |
| YWHAZ | -0.398335039 | 0.000240632 | 0.000633647 |
| YY2 | -0.896033973 | 9.67E-19 | 1.01E-17 |
| ZAR1 | -1.431067489 | 0.00223419 | 0.005019356 |
| ZBED4 | 0.590756136 | 1.04E-10 | 5.62E-10 |
| ZBTB1 | 0.687525578 | 1.77E-16 | 1.56E-15 |
| ZBTB10 | -0.807675692 | 2.65E-05 | 7.99E-05 |
| ZBTB11 | -0.623577994 | 0.025380508 | 0.045379093 |
| ZBTB14 | 0.743227542 | 0.008641066 | 0.017142619 |
| ZBTB18 | 0.412768076 | 0.012175198 | 0.023449808 |
| ZBTB2 | -0.324875776 | 0.007614012 | 0.015313651 |
| ZBTB20 | 0.447635638 | 0.019526262 | 0.035908666 |
| ZBTB21 | -0.50684299 | 0.00241561 | 0.005397496 |
| ZBTB24 | -0.357074079 | 0.01360017 | 0.025907991 |
| ZBTB25 | -0.98905703 | 7.69E-06 | 2.50E-05 |
| ZBTB26 | 0.9564455 | 7.78E-25 | 1.20E-23 |
| ZBTB33 | 0.909273946 | 7.10E-18 | 6.92E-17 |
| ZBTB34 | 1.763290635 | 3.50E-46 | 1.79E-44 |
| ZBTB41 | 0.36431927 | 0.003430334 | 0.007440967 |
| ZBTB44 | -2.003845892 | 1.23E-05 | 3.88E-05 |
| ZBTB45 | -0.85381372 | 6.74E-07 | 2.51E-06 |
| ZBTB46 | -0.699577267 | 0.011555907 | 0.022348875 |
| ZBTB48 | -0.685098938 | 5.68E-05 | 0.000164508 |
| ZBTB49 | -1.013025878 | 1.46E-11 | 8.55E-11 |
| ZBTB5 | -1.424972386 | 9.38E-07 | 3.43E-06 |
| ZBTB6 | 0.775466072 | 0.003409463 | 0.007400419 |
| ZBTB7A | -1.58301437 | 7.96E-18 | 7.73E-17 |
| ZBTB8A | -0.880757462 | 7.18E-11 | 3.94E-10 |
| ZC2HC1A | 0.434300466 | 0.011147109 | 0.02160746 |
| ZC2HC1C | -1.912986171 | 1.24E-10 | 6.67E-10 |

| | | | |
|---|---|---|---|
| ZC3H12A | -0.83233026 | 1.75E-10 | 9.23E-10 |
| ZC3H12B | 1.164018291 | 4.20E-11 | 2.35E-10 |
| ZC3H14 | -0.382520762 | 0.000780033 | 0.001892575 |
| ZC3H15 | 0.546400921 | 5.42E-10 | 2.74E-09 |
| ZC3H3 | -0.583886491 | 1.54E-05 | 4.79E-05 |
| ZC3HC1 | -1.230250284 | 9.92E-17 | 8.90E-16 |
| ZCCHC10 | 0.341919034 | 0.000755333 | 0.001836908 |
| ZCCHC14 | -0.773901328 | 6.26E-05 | 0.00017995 |
| ZCCHC17 | 0.84981809 | 6.29E-16 | 5.29E-15 |
| ZCCHC24 | 1.189686611 | 2.13E-26 | 3.62E-25 |
| ZCCHC4 | 0.561681039 | 2.21E-05 | 6.72E-05 |
| ZCCHC8 | -0.949977229 | 1.24E-08 | 5.53E-08 |
| ZCRB1 | -0.953932731 | 6.27E-15 | 4.89E-14 |
| ZDHHC13 | -0.648208234 | 4.98E-09 | 2.30E-08 |
| ZDHHC14 | 0.520595452 | 0.003104499 | 0.006800379 |
| ZDHHC15 | 0.763284597 | 0.019572895 | 0.035984687 |
| ZDHHC16 | -0.766426927 | 1.37E-06 | 4.91E-06 |
| ZDHHC17 | 1.588038328 | 3.52E-59 | 2.97E-57 |
| ZDHHC2 | 0.523861223 | 0.015907107 | 0.029805541 |
| ZDHHC20 | 0.560710744 | 7.00E-10 | 3.51E-09 |
| ZDHHC21 | 1.080971435 | 6.57E-20 | 7.33E-19 |
| ZDHHC3 | 1.668512344 | 1.97E-81 | 4.07E-79 |
| ZDHHC4 | 0.635883217 | 0.000148392 | 0.000401941 |
| ZDHHC5 | 0.596125051 | 3.19E-06 | 1.09E-05 |
| ZDHHC6 | 1.378485058 | 2.25E-71 | 3.09E-69 |
| ZDHHC7 | 1.22305938 | 5.26E-33 | 1.29E-31 |
| ZDHHC9 | 1.084520533 | 5.45E-49 | 3.19E-47 |
| ZEB1 | 0.277329001 | 0.00721119 | 0.014565983 |
| ZER1 | -1.009553742 | 1.06E-07 | 4.31E-07 |
| ZFAND1 | 0.377604518 | 0.005922906 | 0.012192055 |
| ZFAND2A | -0.5742476 | 1.66E-08 | 7.31E-08 |
| ZFAND3 | 0.713956175 | 8.62E-19 | 9.01E-18 |
| ZFAND4 | -0.646142858 | 1.49E-06 | 5.33E-06 |
| ZFAND5 | -0.453022788 | 1.60E-05 | 4.96E-05 |
| ZFAND6 | 0.442143648 | 5.04E-06 | 1.67E-05 |
| ZFAT | -1.323032837 | 1.08E-07 | 4.37E-07 |
| ZFP36L2 | -0.960648669 | 2.41E-07 | 9.43E-07 |
| ZFP64 | -0.528097444 | 0.00015383 | 0.000415761 |
| ZFP92 | 0.902272698 | 5.48E-12 | 3.34E-11 |
| ZFYVE1 | -0.650936223 | 5.67E-09 | 2.60E-08 |
| ZFYVE27 | -1.46060204 | 8.57E-16 | 7.12E-15 |
| ZFYVE9 | 0.61928577 | 0.008384211 | 0.016704514 |
| ZHX2 | -0.819501434 | 0.009736482 | 0.019107459 |
| ZMAT2 | 0.689171396 | 1.07E-14 | 8.16E-14 |
| ZMYM3 | -1.524700749 | 0.001377077 | 0.003213809 |
| ZMYND10 | 1.437942387 | 0.021163558 | 0.038575175 |
| ZMYND11 | -0.577752569 | 7.44E-05 | 0.000211349 |
| ZMYND19 | -0.417979288 | 1.09E-05 | 3.47E-05 |

| | | | |
|---|---|---|---|
| ZNF131 | -0.360987978 | 0.002100997 | 0.004742858 |
| ZNF142 | -0.558409383 | 0.004268796 | 0.009067272 |
| ZNF318 | -0.785609098 | 5.32E-05 | 0.000154736 |
| ZNF330 | 1.086893752 | 1.91E-18 | 1.95E-17 |
| ZNF346 | -0.369425068 | 0.020174557 | 0.036935961 |
| ZNF362 | 1.67997604 | 2.08E-07 | 8.18E-07 |
| ZNF365 | 2.018707011 | 1.69E-115 | 8.83E-113 |
| ZNF384 | 1.242752966 | 9.01E-18 | 8.73E-17 |
| ZNF395 | -1.478297408 | 0.00472009 | 0.009931186 |
| ZNF407 | -0.703155653 | 0.002122964 | 0.00479006 |
| ZNF410 | 0.33128932 | 0.001121874 | 0.002662101 |
| ZNF414 | -0.690243006 | 0.015496964 | 0.029133435 |
| ZNF423 | 0.512010104 | 0.008315831 | 0.016585294 |
| ZNF462 | -1.343830766 | 0.016705482 | 0.031159725 |
| ZNF467 | 1.277228527 | 3.84E-19 | 4.10E-18 |
| ZNF469 | 0.509585811 | 0.00028254 | 0.000735546 |
| ZNF503 | -0.765108366 | 0.000465645 | 0.001170529 |
| ZNF512B | 0.741412931 | 1.16E-05 | 3.67E-05 |
| ZNF513 | -0.520299734 | 0.000612848 | 0.001512197 |
| ZNF516 | -0.434042947 | 0.005074769 | 0.010598015 |
| ZNF518A | -0.448109994 | 0.003024325 | 0.006639739 |
| ZNF518B | -1.320631297 | 2.01E-17 | 1.91E-16 |
| ZNF521 | -0.830147161 | 3.77E-08 | 1.60E-07 |
| ZNF592 | -0.885633127 | 0.000164539 | 0.000442679 |
| ZNF593 | 0.707415562 | 2.95E-11 | 1.68E-10 |
| ZNF598 | 0.299630763 | 0.001828128 | 0.004172615 |
| ZNF609 | -0.469130235 | 0.000283937 | 0.000738617 |
| ZNF618 | 0.431555085 | 0.000498484 | 0.00124708 |
| ZNF622 | -0.394509078 | 1.21E-05 | 3.81E-05 |
| ZNF628 | 1.469067065 | 1.78E-38 | 6.03E-37 |
| ZNF639 | -0.833527389 | 1.48E-12 | 9.49E-12 |
| ZNF644 | 0.930634491 | 2.70E-23 | 3.71E-22 |
| ZNF652 | 0.958812222 | 4.04E-13 | 2.74E-12 |
| ZNF704 | 1.282465169 | 2.00E-31 | 4.57E-30 |
| ZNF711 | 0.292511352 | 0.001852554 | 0.004221983 |
| ZNF767 | -0.557553627 | 2.50E-06 | 8.70E-06 |
| ZNF821 | 1.731866683 | 6.37E-67 | 7.67E-65 |
| ZNF830 | -0.616842509 | 3.82E-09 | 1.78E-08 |
| ZNF839 | -1.279646816 | 1.29E-10 | 6.91E-10 |
| ZNHIT3 | -0.312584569 | 0.009524056 | 0.018730678 |
| ZNRF1 | 1.333358011 | 9.85E-05 | 0.000274345 |
| ZNRF3 | -0.582648549 | 0.015859014 | 0.029727723 |
| ZP4 | 3.635221581 | 0.018358772 | 0.033936139 |
| ZPLD1 | 1.15767562 | 1.33E-07 | 5.35E-07 |
| ZRANB1 | -0.253843541 | 0.027110976 | 0.048156349 |
| ZRANB2 | 0.704448963 | 4.38E-11 | 2.45E-10 |
| ZRANB3 | -1.095668519 | 0.017674978 | 0.032797037 |
| ZRSR2 | -0.653140711 | 6.87E-05 | 0.000195933 |

| | | | |
|---|---|---|---|
| ZSWIM3 | -1.640609166 | 2.32E-05 | 7.06E-05 |
| ZSWIM5 | -0.413293602 | 0.001487638 | 0.003452865 |
| ZSWIM7 | 0.951709708 | 2.31E-10 | 1.21E-09 |
| ZSWIM8 | -1.063216703 | 2.04E-05 | 6.23E-05 |
| ZUP1 | -1.104745632 | 9.71E-12 | 5.80E-11 |
| ZW10 | -0.567304829 | 4.68E-07 | 1.78E-06 |
| ZWILCH | -1.295144171 | 4.20E-30 | 8.92E-29 |
| ZYG11A | -1.177824658 | 3.26E-18 | 3.28E-17 |
| ZYX | 0.215105843 | 0.019348009 | 0.035609769 |
| ZZEF1 | -0.863260644 | 5.90E-08 | 2.46E-07 |
| ZZZ3 | -0.562182415 | 2.48E-05 | 7.52E-05 |

**Supplementary Table 8: Differentially expressed genes in infected duck endothelial cells**

| Gene name | log2FoldChange | pvalue | padj |
|---|---|---|---|
| BCOR | 4.810686 | 4.19E-106 | 4.57E-102 |
| PDE4B | 2.934439 | 1.90E-66 | 1.03E-62 |
| ENSAPLG00000031170 | 2.353503 | 2.90E-42 | 1.05E-38 |
| SOX8 | 2.215518 | 1.47E-41 | 3.99E-38 |
| MMP1 | -4.50723 | 1.64E-40 | 3.56E-37 |
| SCX | 2.310445 | 1.46E-38 | 2.65E-35 |
| RGS16 | 4.549672 | 2.30E-38 | 3.58E-35 |
| PCF11 | 2.449039 | 1.24E-37 | 1.69E-34 |
| ENSAPLG00000008725 | 2.601815 | 1.05E-36 | 1.27E-33 |
| DNAH10 | 4.816031 | 2.47E-36 | 2.70E-33 |
| HIC1 | 2.705134 | 3.04E-36 | 3.01E-33 |
| RBBP6 | 2.065838 | 1.75E-34 | 1.58E-31 |
| ENSAPLG00000016836 | 6.280844 | 2.31E-32 | 1.93E-29 |
| ENSAPLG00000007834 | 2.78189 | 5.52E-32 | 4.29E-29 |
| NR4A3 | 2.38368 | 1.16E-30 | 8.42E-28 |
| PRICKLE2 | 1.891648 | 1.38E-30 | 9.43E-28 |
| DIS3L | 1.795107 | 9.18E-30 | 5.88E-27 |
| ABCA12 | 5.66765 | 1.48E-28 | 8.96E-26 |
| TNFRSF21 | 1.777043 | 1.48E-26 | 8.46E-24 |
| ENSAPLG00000018309 | 1.670103 | 2.24E-26 | 1.22E-23 |
| ENSAPLG00000022613 | 4.854975 | 2.82E-26 | 1.46E-23 |
| HSPA5 | -1.7945 | 7.69E-26 | 3.81E-23 |
| LSMEM1 | 3.751869 | 9.63E-26 | 4.56E-23 |
| PDIA4 | -1.97121 | 1.31E-25 | 5.71E-23 |
| TIPARP | 1.805563 | 1.30E-25 | 5.71E-23 |
| ENSAPLG00000012692 | 2.332517 | 1.52E-24 | 6.35E-22 |
| FBLN1 | -1.76955 | 2.71E-24 | 1.09E-21 |
| ABHD5 | 1.604006 | 5.22E-24 | 2.03E-21 |
| CCN2 | -1.64509 | 5.43E-24 | 2.04E-21 |
| SPI1 | 2.528119 | 1.87E-23 | 6.80E-21 |
| HERPUD1 | -1.97792 | 5.23E-23 | 1.84E-20 |
| UPF3A | 3.692505 | 1.00E-22 | 3.41E-20 |
| ZNF821 | 1.71516 | 1.56E-22 | 5.15E-20 |
| SPHK1 | 1.637424 | 2.43E-22 | 7.78E-20 |
| ENSAPLG00000028586 | 8.516654 | 2.91E-22 | 9.06E-20 |
| CAMTA1 | 2.051929 | 4.93E-22 | 1.49E-19 |
| ENSAPLG00000013728 | 2.347574 | 5.46E-22 | 1.61E-19 |
| MCM5 | 2.092079 | 1.13E-21 | 3.24E-19 |
| ENSAPLG00000011493 | 3.474897 | 1.29E-21 | 3.60E-19 |
| RYR3 | 2.131532 | 2.01E-21 | 5.47E-19 |
| TRIM8 | 1.817933 | 7.57E-21 | 2.01E-18 |

| | | | |
|---|---|---|---|
| CCNI | 1.63774 | 1.48E-20 | 3.84E-18 |
| RBM20 | 7.226371 | 1.82E-20 | 4.61E-18 |
| ENSAPLG00000023444 | 1.98318 | 4.52E-20 | 1.12E-17 |
| PHGDH | -2.20133 | 6.42E-20 | 1.55E-17 |
| NAIF1 | 1.647313 | 9.08E-20 | 2.15E-17 |
| SERTAD2 | 2.232292 | 1.37E-19 | 3.17E-17 |
| CSTF3 | 1.95822 | 1.70E-19 | 3.86E-17 |
| RASGRP3 | 3.430584 | 2.41E-19 | 5.37E-17 |
| ENSAPLG00000013720 | 1.983787 | 3.17E-19 | 6.91E-17 |
| AKAP13 | 1.745062 | 7.54E-19 | 1.61E-16 |
| HSP90B1 | -1.46901 | 1.38E-18 | 2.90E-16 |
| ABHD17A | 1.760709 | 2.13E-18 | 4.34E-16 |
| ENSAPLG00000024296 | 1.735451 | 2.15E-18 | 4.34E-16 |
| HYOU1 | -1.38186 | 3.11E-18 | 6.16E-16 |
| CPZ | -1.95219 | 3.49E-18 | 6.79E-16 |
| POU2F1 | 1.598825 | 5.03E-18 | 9.44E-16 |
| ITGB4 | 4.387276 | 5.02E-18 | 9.44E-16 |
| AHDC1 | 2.846925 | 7.12E-18 | 1.31E-15 |
| ENSAPLG00000007205 | 1.827747 | 1.15E-17 | 2.07E-15 |
| NRAP | 3.989021 | 1.16E-17 | 2.07E-15 |
| ENSAPLG00000008620 | 6.82055 | 1.32E-17 | 2.32E-15 |
| CITED4 | -1.61848 | 1.64E-17 | 2.83E-15 |
| PRR7 | 2.068587 | 4.69E-17 | 7.98E-15 |
| HSD11B2 | 2.702323 | 5.14E-17 | 8.61E-15 |
| CREB5 | 1.39319 | 5.31E-17 | 8.76E-15 |
| ZNF628 | 2.237488 | 6.55E-17 | 1.06E-14 |
| FGGY | 1.746143 | 8.31E-17 | 1.32E-14 |
| SARS1 | -1.4909 | 8.39E-17 | 1.32E-14 |
| BCL2L11 | 2.594726 | 9.82E-17 | 1.53E-14 |
| ENSAPLG00000022960 | -1.73333 | 9.95E-17 | 1.53E-14 |
| GCLC | 1.273943 | 1.20E-16 | 1.81E-14 |
| PIGT | 1.443444 | 1.21E-16 | 1.81E-14 |
| ENSAPLG00000017312 | 3.002753 | 1.24E-16 | 1.83E-14 |
| PRRX1 | 1.823967 | 2.19E-16 | 3.18E-14 |
| MAGT1 | -1.63039 | 2.22E-16 | 3.18E-14 |
| ANAPC7 | -1.53106 | 2.78E-16 | 3.94E-14 |
| ENSAPLG00000006823 | 1.470652 | 4.20E-16 | 5.86E-14 |
| SAMD4A | 1.687709 | 4.42E-16 | 6.02E-14 |
| CD44 | -1.33815 | 4.38E-16 | 6.02E-14 |
| STK40 | 1.442348 | 4.71E-16 | 6.33E-14 |
| ASNS | -1.43873 | 5.31E-16 | 7.05E-14 |
| BCL6 | 1.74783 | 5.44E-16 | 7.14E-14 |
| ENSAPLG00000029778 | 1.440778 | 6.40E-16 | 8.30E-14 |
| ENSAPLG00000011089 | 1.41499 | 6.66E-16 | 8.53E-14 |
| FST | -1.53634 | 6.75E-16 | 8.55E-14 |

| | | | |
|---|---|---|---|
| KCNJ2 | 1.807775 | 7.67E-16 | 9.61E-14 |
| PENK | 2.088784 | 7.93E-16 | 9.82E-14 |
| PPP2R2B | 1.712104 | 9.68E-16 | 1.18E-13 |
| TMEM251 | 1.377445 | 1.25E-15 | 1.51E-13 |
| SLC35E3 | 1.277646 | 1.44E-15 | 1.73E-13 |
| FAM222B | 1.923048 | 1.77E-15 | 2.09E-13 |
| DSP | 4.366519 | 1.95E-15 | 2.29E-13 |
| HCN1 | 2.458019 | 2.49E-15 | 2.88E-13 |
| P4HB | -1.25625 | 2.69E-15 | 3.09E-13 |
| MAPK6 | 1.492345 | 3.05E-15 | 3.46E-13 |
| TET2 | 1.814368 | 5.51E-15 | 6.19E-13 |
| PDE8A | 1.713582 | 5.60E-15 | 6.23E-13 |
| PEPD | -1.42732 | 6.69E-15 | 7.36E-13 |
| COPB1 | -1.21658 | 7.55E-15 | 8.22E-13 |
| GYS2 | 8.38027 | 8.25E-15 | 8.90E-13 |
| UQCRC2 | -1.42094 | 8.62E-15 | 9.17E-13 |
| ARID1A | 1.89519 | 8.67E-15 | 9.17E-13 |
| GUSB | -1.43357 | 1.07E-14 | 1.12E-12 |
| ABHD6 | 2.349445 | 1.09E-14 | 1.13E-12 |
| HTR2A | 1.302675 | 1.23E-14 | 1.27E-12 |
| ANKRD11 | 1.541441 | 1.29E-14 | 1.31E-12 |
| NCKAP5 | 1.239249 | 1.40E-14 | 1.41E-12 |
| ENSAPLG00000011970 | 2.715789 | 1.85E-14 | 1.85E-12 |
| PSAT1 | -1.66829 | 2.10E-14 | 2.08E-12 |
| LINGO1 | 1.528429 | 2.23E-14 | 2.18E-12 |
| MAFB | 1.765966 | 2.36E-14 | 2.30E-12 |
| MECOM | 1.619938 | 2.67E-14 | 2.58E-12 |
| B4GALT2 | 1.563807 | 2.92E-14 | 2.79E-12 |
| ARSB | -1.44755 | 3.14E-14 | 2.97E-12 |
| SNTB2 | -1.64652 | 4.09E-14 | 3.84E-12 |
| SRF | 1.311632 | 4.34E-14 | 4.04E-12 |
| ZBTB34 | 1.39469 | 4.40E-14 | 4.07E-12 |
| TMCC1 | 1.347714 | 4.54E-14 | 4.09E-12 |
| SIX1 | 1.742615 | 4.55E-14 | 4.09E-12 |
| PREP | -1.21272 | 4.51E-14 | 4.09E-12 |
| ANGPTL4 | 1.553283 | 4.85E-14 | 4.33E-12 |
| KMT5B | 1.244823 | 4.96E-14 | 4.39E-12 |
| PCSK5 | -1.39302 | 5.00E-14 | 4.39E-12 |
| CRTAP | -1.15852 | 6.27E-14 | 5.46E-12 |
| HES6 | 2.238781 | 6.53E-14 | 5.64E-12 |
| TARS1 | -1.27035 | 7.29E-14 | 6.25E-12 |
| ELL2 | 1.570416 | 9.61E-14 | 8.18E-12 |
| GPR39 | 1.204097 | 1.01E-13 | 8.52E-12 |
| ZNF652 | 1.33209 | 1.05E-13 | 8.77E-12 |
| FBN1 | -1.92345 | 1.15E-13 | 9.56E-12 |

| | | | |
|---|---|---|---|
| SEMA3D | -1.38 | 1.24E-13 | 1.02E-11 |
| MIPEP | -1.38499 | 1.35E-13 | 1.10E-11 |
| DECR1 | -1.67862 | 1.47E-13 | 1.20E-11 |
| R3HDM1 | 1.52304 | 1.51E-13 | 1.22E-11 |
| ENSAPLG00000026952 | 1.518703 | 1.57E-13 | 1.26E-11 |
| ENSAPLG00000001129 | 1.306828 | 1.63E-13 | 1.29E-11 |
| AHCY | -1.49218 | 1.85E-13 | 1.46E-11 |
| AGO3 | 1.24288 | 2.02E-13 | 1.58E-11 |
| PLOD1 | -1.3778 | 2.22E-13 | 1.73E-11 |
| ST3GAL4 | 1.266506 | 2.30E-13 | 1.78E-11 |
| GLB1 | -1.36671 | 2.50E-13 | 1.92E-11 |
| ENSAPLG00000019985 | 1.698288 | 2.63E-13 | 2.00E-11 |
| MMP2 | -1.16016 | 2.88E-13 | 2.18E-11 |
| IL1R1 | 1.189979 | 3.03E-13 | 2.28E-11 |
| DPP7 | -1.3283 | 3.17E-13 | 2.36E-11 |
| CTSC | -1.1873 | 3.42E-13 | 2.54E-11 |
| RTN4RL2 | 1.651925 | 3.57E-13 | 2.62E-11 |
| KLF9 | 1.617019 | 4.06E-13 | 2.97E-11 |
| PTP4A1 | 1.14704 | 4.21E-13 | 3.06E-11 |
| DUSP16 | 1.373302 | 4.45E-13 | 3.21E-11 |
| SPSB1 | 1.301972 | 4.52E-13 | 3.24E-11 |
| CMIP | 1.241148 | 4.55E-13 | 3.24E-11 |
| P3H3 | -1.60448 | 4.85E-13 | 3.41E-11 |
| DUSP6 | 1.237996 | 4.83E-13 | 3.41E-11 |
| DLGAP2 | 2.044335 | 5.18E-13 | 3.62E-11 |
| BEST1 | 5.006478 | 5.24E-13 | 3.64E-11 |
| ENSAPLG00000003693 | 2.920012 | 5.38E-13 | 3.71E-11 |
| FBN2 | 1.723536 | 6.09E-13 | 4.14E-11 |
| SPRED2 | 1.399396 | 6.07E-13 | 4.14E-11 |
| HTR4 | 1.569229 | 6.23E-13 | 4.21E-11 |
| ENSAPLG00000010116 | 5.580268 | 6.67E-13 | 4.48E-11 |
| SMAD6 | 1.472805 | 8.37E-13 | 5.59E-11 |
| HSD17B4 | -1.27617 | 8.43E-13 | 5.60E-11 |
| FOS | 1.666095 | 8.72E-13 | 5.76E-11 |
| PRELID1 | 1.389188 | 9.44E-13 | 6.19E-11 |
| IP6K2 | 1.331802 | 1.04E-12 | 6.78E-11 |
| PHOSPHO1 | 2.135349 | 1.10E-12 | 7.11E-11 |
| CRELD2 | -1.51016 | 1.24E-12 | 7.97E-11 |
| ENSAPLG00000006052 | 5.028184 | 1.26E-12 | 8.07E-11 |
| EFNB1 | 1.280833 | 1.31E-12 | 8.34E-11 |
| ENSAPLG00000010314 | -1.81859 | 1.33E-12 | 8.40E-11 |
| HEXB | -1.25151 | 1.48E-12 | 9.31E-11 |
| APP | -1.13696 | 1.52E-12 | 9.50E-11 |
| PAICS | -1.18261 | 1.57E-12 | 9.80E-11 |
| DCAF5 | 1.282327 | 1.58E-12 | 9.80E-11 |

| | | | |
|---|---|---|---|
| KIAA0232 | 1.34288 | 1.90E-12 | 1.17E-10 |
| CYP24A1 | -1.44042 | 2.07E-12 | 1.27E-10 |
| PAX7 | 5.494022 | 2.12E-12 | 1.29E-10 |
| OPRD1 | 8.586176 | 2.52E-12 | 1.53E-10 |
| ENSAPLG00000015803 | 1.746749 | 3.06E-12 | 1.84E-10 |
| WNT2B | 2.257319 | 3.18E-12 | 1.90E-10 |
| GLI3 | 1.220048 | 3.24E-12 | 1.93E-10 |
| ENSAPLG00000015474 | -1.44229 | 3.68E-12 | 2.18E-10 |
| G0S2 | 1.937415 | 3.79E-12 | 2.23E-10 |
| VEGFA | 1.335876 | 3.80E-12 | 2.23E-10 |
| ENSAPLG00000009880 | 1.67683 | 3.91E-12 | 2.28E-10 |
| RHOQ | 1.603618 | 4.50E-12 | 2.61E-10 |
| LHX2 | 1.59003 | 4.57E-12 | 2.62E-10 |
| CDKN1B | 1.466906 | 4.55E-12 | 2.62E-10 |
| ENSAPLG00000008458 | 1.351327 | 4.85E-12 | 2.77E-10 |
| FRS2 | 1.44956 | 5.17E-12 | 2.94E-10 |
| GRK3 | 1.98258 | 5.77E-12 | 3.26E-10 |
| ENSAPLG00000022579 | 7.430762 | 5.81E-12 | 3.26E-10 |
| GLT8D1 | -1.18414 | 6.15E-12 | 3.44E-10 |
| STRADB | 1.372349 | 6.45E-12 | 3.58E-10 |
| NAB1 | 1.386411 | 6.49E-12 | 3.59E-10 |
| HMOX1 | 1.776531 | 6.53E-12 | 3.59E-10 |
| SHISA2 | 1.246787 | 8.23E-12 | 4.50E-10 |
| PHF21A | 1.49028 | 9.14E-12 | 4.98E-10 |
| PIK3R6 | 3.089578 | 9.35E-12 | 5.07E-10 |
| FAM214A | 1.218574 | 9.40E-12 | 5.07E-10 |
| RCL1 | -1.37611 | 1.06E-11 | 5.71E-10 |
| SLC34A2 | 2.704814 | 1.26E-11 | 6.72E-10 |
| GSN | -1.48655 | 1.28E-11 | 6.76E-10 |
| ENSAPLG00000008450 | -1.24372 | 1.28E-11 | 6.76E-10 |
| MEIS1 | 1.394996 | 1.28E-11 | 6.76E-10 |
| RNF165 | 1.430701 | 1.29E-11 | 6.78E-10 |
| SLC15A4 | -1.36679 | 1.37E-11 | 7.14E-10 |
| HSPB8 | 1.510295 | 1.39E-11 | 7.23E-10 |
| FOSL2 | 2.300961 | 1.62E-11 | 8.36E-10 |
| SPEN | 1.346661 | 1.74E-11 | 8.94E-10 |
| PDIA6 | -1.15429 | 1.79E-11 | 9.14E-10 |
| LPCAT2 | -1.21244 | 1.82E-11 | 9.29E-10 |
| ENSAPLG00000003290 | 3.381127 | 1.87E-11 | 9.46E-10 |
| RAB31 | 1.068927 | 1.95E-11 | 9.85E-10 |
| ELL | 1.352736 | 2.11E-11 | 1.06E-09 |
| GOT1 | -1.28127 | 2.13E-11 | 1.07E-09 |
| RFX5 | 1.224472 | 2.23E-11 | 1.11E-09 |
| NDUFS1 | -1.17362 | 2.25E-11 | 1.11E-09 |
| MEF2A | 1.198362 | 2.29E-11 | 1.13E-09 |

| | | | |
|---|---|---|---|
| ENSAPLG00000009135 | 1.482093 | 2.31E-11 | 1.13E-09 |
| STT3B | -1.09496 | 2.39E-11 | 1.17E-09 |
| ENSAPLG00000016465 | -1.5845 | 2.43E-11 | 1.18E-09 |
| GARS1 | -1.0965 | 2.57E-11 | 1.24E-09 |
| PDIA5 | -1.12264 | 2.70E-11 | 1.30E-09 |
| TM2D3 | -1.24887 | 2.70E-11 | 1.30E-09 |
| ENSAPLG00000013296 | 1.420731 | 2.82E-11 | 1.35E-09 |
| ALDH9A1 | -1.20123 | 2.98E-11 | 1.42E-09 |
| ENSAPLG00000022525 | 1.208154 | 2.99E-11 | 1.42E-09 |
| ATP5F1A | -1.20572 | 3.04E-11 | 1.43E-09 |
| SDF4 | -1.07712 | 3.22E-11 | 1.51E-09 |
| BMF | 1.826836 | 3.34E-11 | 1.56E-09 |
| ENOX1 | 1.296652 | 3.63E-11 | 1.69E-09 |
| PPARD | 1.111175 | 3.96E-11 | 1.84E-09 |
| GMDS | 1.102636 | 4.00E-11 | 1.85E-09 |
| PER2 | 1.123843 | 4.07E-11 | 1.87E-09 |
| FN1 | -1.19224 | 4.43E-11 | 2.03E-09 |
| CKAP4 | -1.13824 | 4.66E-11 | 2.12E-09 |
| SBNO2 | 1.763534 | 4.71E-11 | 2.14E-09 |
| MGAT4B | 1.216512 | 4.84E-11 | 2.19E-09 |
| COLEC11 | 4.335909 | 5.60E-11 | 2.52E-09 |
| TMEM203 | 1.392406 | 5.64E-11 | 2.53E-09 |
| YWHAG | 1.06906 | 6.55E-11 | 2.93E-09 |
| ENSAPLG00000000589 | -1.89071 | 6.62E-11 | 2.94E-09 |
| ENSAPLG00000015132 | -1.35599 | 6.80E-11 | 3.01E-09 |
| SDF2L1 | -1.56647 | 7.46E-11 | 3.29E-09 |
| DARS1 | -1.12452 | 7.91E-11 | 3.47E-09 |
| ENSAPLG00000031224 | -1.46066 | 8.16E-11 | 3.57E-09 |
| TRPV1 | 4.785972 | 8.22E-11 | 3.58E-09 |
| GFOD1 | 1.21008 | 8.98E-11 | 3.90E-09 |
| TRMT61A | -1.12888 | 9.07E-11 | 3.92E-09 |
| UTS2R | 2.882884 | 9.22E-11 | 3.97E-09 |
| SUCLG2 | -1.58696 | 9.35E-11 | 4.01E-09 |
| HEY1 | 1.506604 | 9.60E-11 | 4.10E-09 |
| ENSAPLG00000005553 | 3.520602 | 9.69E-11 | 4.12E-09 |
| LRRC61 | 2.690482 | 9.80E-11 | 4.15E-09 |
| ENSAPLG00000000660 | 1.812879 | 1.01E-10 | 4.25E-09 |
| NSMF | 1.297669 | 1.06E-10 | 4.46E-09 |
| ENSAPLG00000014035 | 3.590203 | 1.08E-10 | 4.53E-09 |
| ITGB1 | -1.04038 | 1.12E-10 | 4.67E-09 |
| LBR | -1.2294 | 1.26E-10 | 5.24E-09 |
| MDFI | 2.966063 | 1.29E-10 | 5.33E-09 |
| ENSAPLG00000007073 | 1.203418 | 1.33E-10 | 5.48E-09 |
| ZNF362 | 1.925684 | 1.33E-10 | 5.48E-09 |
| SPON1 | -1.18259 | 1.34E-10 | 5.49E-09 |

| | | | |
|---|---|---|---|
| AMACR | -1.68488 | 1.35E-10 | 5.51E-09 |
| MEGF9 | 1.847934 | 1.43E-10 | 5.82E-09 |
| IDH1 | -1.04705 | 1.48E-10 | 5.98E-09 |
| ENSAPLG00000024789 | -1.44166 | 1.59E-10 | 6.38E-09 |
| DES | 1.45144 | 1.59E-10 | 6.38E-09 |
| GPR162 | 1.3462 | 1.59E-10 | 6.38E-09 |
| SUMO2 | 1.367224 | 1.68E-10 | 6.72E-09 |
| ENSAPLG00000024440 | 1.075826 | 1.72E-10 | 6.82E-09 |
| TOR1B | -1.2429 | 1.76E-10 | 6.99E-09 |
| SLC25A37 | 1.74365 | 1.78E-10 | 7.04E-09 |
| GABRR2 | 5.151723 | 1.81E-10 | 7.13E-09 |
| SCPEP1 | -1.11476 | 1.91E-10 | 7.47E-09 |
| NFYC | 1.08502 | 1.96E-10 | 7.65E-09 |
| CYP4V2 | -1.7246 | 1.97E-10 | 7.65E-09 |
| PDZRN3 | -1.52735 | 2.01E-10 | 7.81E-09 |
| CERS6 | -1.26312 | 2.04E-10 | 7.88E-09 |
| RUNX1 | 1.251548 | 2.12E-10 | 8.15E-09 |
| FGF19 | 3.200464 | 2.17E-10 | 8.31E-09 |
| TACC2 | 1.174391 | 2.27E-10 | 8.68E-09 |
| PDGFA | 1.092633 | 2.33E-10 | 8.86E-09 |
| TPST1 | 1.127346 | 2.39E-10 | 9.05E-09 |
| YBX1 | 1.353157 | 2.42E-10 | 9.16E-09 |
| HIPK2 | 1.090639 | 2.50E-10 | 9.41E-09 |
| BAG5 | 1.169921 | 2.59E-10 | 9.74E-09 |
| PANX3 | 2.634459 | 2.70E-10 | 1.01E-08 |
| GDAP1L1 | 2.675983 | 2.82E-10 | 1.05E-08 |
| ATXN7 | 1.267922 | 2.85E-10 | 1.06E-08 |
| ECD | 1.251366 | 2.85E-10 | 1.06E-08 |
| ACO1 | -1.21571 | 2.89E-10 | 1.07E-08 |
| SCCPDH | -1.13015 | 3.02E-10 | 1.11E-08 |
| CDYL2 | 1.589222 | 3.16E-10 | 1.16E-08 |
| ENSAPLG00000027620 | 1.660157 | 3.16E-10 | 1.16E-08 |
| PMEPA1 | 1.324281 | 3.20E-10 | 1.17E-08 |
| KHDRBS1 | -1.10901 | 3.34E-10 | 1.21E-08 |
| APC | 1.29939 | 3.46E-10 | 1.25E-08 |
| TBCCD1 | -1.95949 | 3.48E-10 | 1.25E-08 |
| ENSAPLG00000007544 | 1.372245 | 3.51E-10 | 1.26E-08 |
| ERRFI1 | 1.412675 | 3.54E-10 | 1.27E-08 |
| TRERF1 | 1.443029 | 3.55E-10 | 1.27E-08 |
| GTF3C6 | 1.298418 | 3.63E-10 | 1.29E-08 |
| ENSAPLG00000002513 | -1.78753 | 3.65E-10 | 1.29E-08 |
| KIF14 | 4.279406 | 3.68E-10 | 1.30E-08 |
| PDIA3 | -1.24067 | 4.03E-10 | 1.42E-08 |
| BSX | 8.051357 | 4.02E-10 | 1.42E-08 |
| ENSAPLG00000027582 | 2.758299 | 4.10E-10 | 1.44E-08 |

| | | | |
|---|---|---|---|
| RNF38 | 1.176957 | 4.18E-10 | 1.46E-08 |
| DNAJC3 | -1.14667 | 4.40E-10 | 1.53E-08 |
| DOT1L | 1.284702 | 4.46E-10 | 1.55E-08 |
| SERPINI1 | -1.29222 | 4.51E-10 | 1.56E-08 |
| TAB3 | 1.375387 | 4.57E-10 | 1.57E-08 |
| HTRA1 | -1.23214 | 4.75E-10 | 1.63E-08 |
| MYO5C | 2.030945 | 4.75E-10 | 1.63E-08 |
| RPLP1 | 1.322761 | 4.88E-10 | 1.67E-08 |
| ENSAPLG00000010079 | -1.38142 | 5.00E-10 | 1.70E-08 |
| RASL11A | 3.398441 | 5.58E-10 | 1.89E-08 |
| PTEN | 1.062374 | 5.72E-10 | 1.93E-08 |
| RAB30 | 1.126871 | 5.83E-10 | 1.97E-08 |
| LANCL2 | -1.12939 | 6.15E-10 | 2.07E-08 |
| CHD7 | 1.173348 | 6.50E-10 | 2.18E-08 |
| JMJD4 | 1.305893 | 6.58E-10 | 2.20E-08 |
| EVL | 1.15524 | 6.60E-10 | 2.20E-08 |
| TKT | -1.09915 | 6.65E-10 | 2.21E-08 |
| LGMN | -1.01899 | 6.72E-10 | 2.22E-08 |
| SINHCAF | 1.015176 | 6.98E-10 | 2.30E-08 |
| ENSAPLG00000010843 | 5.364875 | 7.12E-10 | 2.34E-08 |
| ENSAPLG00000028209 | 3.009353 | 7.26E-10 | 2.38E-08 |
| SLC7A3 | -1.22909 | 7.79E-10 | 2.55E-08 |
| B3GNT9 | 1.511164 | 7.99E-10 | 2.60E-08 |
| NPEPL1 | -1.04044 | 7.98E-10 | 2.60E-08 |
| CTSH | -1.21328 | 8.12E-10 | 2.63E-08 |
| RNF126 | 1.017308 | 8.68E-10 | 2.81E-08 |
| TBCK | -1.44676 | 8.92E-10 | 2.88E-08 |
| SCN8A | 1.239641 | 8.98E-10 | 2.88E-08 |
| PLBD1 | -1.85124 | 9.69E-10 | 3.10E-08 |
| ATP1A1 | -0.98421 | 9.80E-10 | 3.13E-08 |
| WFS1 | -1.22587 | 9.92E-10 | 3.16E-08 |
| CPQ | -1.44181 | 1.01E-09 | 3.20E-08 |
| CCPG1 | -1.15366 | 1.05E-09 | 3.32E-08 |
| FSTL3 | 1.041923 | 1.07E-09 | 3.36E-08 |
| SLC43A3 | -1.33743 | 1.09E-09 | 3.44E-08 |
| ENSAPLG00000005477 | 1.487016 | 1.13E-09 | 3.53E-08 |
| ENSAPLG00000006627 | -1.07958 | 1.20E-09 | 3.73E-08 |
| HIPK3 | 1.211327 | 1.20E-09 | 3.73E-08 |
| UBE2D2 | 1.014843 | 1.20E-09 | 3.73E-08 |
| AARS1 | -1.03237 | 1.22E-09 | 3.79E-08 |
| ENSAPLG00000030885 | -1.22578 | 1.22E-09 | 3.79E-08 |
| TAF7 | -1.03975 | 1.24E-09 | 3.84E-08 |
| ATF3 | 1.407242 | 1.25E-09 | 3.84E-08 |
| XPA | 1.935231 | 1.32E-09 | 4.05E-08 |
| ACTL6A | -1.03825 | 1.39E-09 | 4.26E-08 |

| | | | |
|---|---|---|---|
| TNN | 1.829687 | 1.45E-09 | 4.41E-08 |
| SLC30A4 | 1.084553 | 1.50E-09 | 4.55E-08 |
| PRKX | 1.2244 | 1.67E-09 | 5.08E-08 |
| SUMF2 | -1.19268 | 1.68E-09 | 5.09E-08 |
| C18orf25 | 1.080385 | 1.75E-09 | 5.27E-08 |
| ENSAPLG00000029963 | 7.835947 | 1.76E-09 | 5.30E-08 |
| DNER | -1.07388 | 1.82E-09 | 5.47E-08 |
| ARHGEF33 | 7.833736 | 1.84E-09 | 5.49E-08 |
| ENSAPLG00000018355 | 1.840106 | 1.84E-09 | 5.49E-08 |
| TET1 | 1.18313 | 1.93E-09 | 5.76E-08 |
| MTNR1B | 2.002738 | 1.99E-09 | 5.90E-08 |
| ACAA2 | -1.13731 | 2.08E-09 | 6.15E-08 |
| MIDN | 1.95257 | 2.11E-09 | 6.22E-08 |
| UGDH | -1.07382 | 2.14E-09 | 6.29E-08 |
| RAI2 | 1.270443 | 2.14E-09 | 6.29E-08 |
| KPNA2 | -1.83746 | 2.21E-09 | 6.47E-08 |
| DDOST | -1.0628 | 2.34E-09 | 6.84E-08 |
| RHOBTB2 | 1.784743 | 2.44E-09 | 7.12E-08 |
| PSPH | -1.34239 | 2.59E-09 | 7.52E-08 |
| ENSAPLG00000017386 | -1.4097 | 2.60E-09 | 7.52E-08 |
| ELK3 | 1.056363 | 2.68E-09 | 7.71E-08 |
| SERPINE2 | -1.21179 | 2.67E-09 | 7.71E-08 |
| XPR1 | 1.079284 | 2.73E-09 | 7.84E-08 |
| ST3GAL1 | 0.976283 | 2.81E-09 | 8.05E-08 |
| RAB11FIP2 | 1.133487 | 3.13E-09 | 8.94E-08 |
| PPAT | -1.20024 | 3.15E-09 | 8.99E-08 |
| RIOX2 | -1.2807 | 3.39E-09 | 9.65E-08 |
| ATP2A2 | -1.27153 | 3.73E-09 | 1.06E-07 |
| ENSAPLG00000026694 | -1.11148 | 4.16E-09 | 1.18E-07 |
| TTL | 1.075837 | 4.31E-09 | 1.22E-07 |
| ACTR10 | -1.10303 | 4.38E-09 | 1.23E-07 |
| GPS1 | -0.97018 | 4.68E-09 | 1.31E-07 |
| ENSAPLG00000022307 | 4.339601 | 4.75E-09 | 1.33E-07 |
| ENSAPLG00000002309 | 1.959564 | 5.20E-09 | 1.45E-07 |
| ENSAPLG00000001088 | 5.206842 | 5.24E-09 | 1.46E-07 |
| CSRNP1 | 1.301524 | 5.28E-09 | 1.47E-07 |
| KCNB1 | 3.555418 | 5.32E-09 | 1.48E-07 |
| PXK | 1.401155 | 5.40E-09 | 1.49E-07 |
| PODN | -1.01033 | 5.78E-09 | 1.59E-07 |
| ENSAPLG00000015195 | -1.00046 | 5.79E-09 | 1.59E-07 |
| MNAT1 | -1.31686 | 5.81E-09 | 1.59E-07 |
| PTPN21 | 1.076727 | 6.09E-09 | 1.67E-07 |
| FEM1B | 1.313149 | 6.32E-09 | 1.73E-07 |
| KRT15 | 7.378309 | 6.53E-09 | 1.77E-07 |
| KLHL25 | 1.497813 | 6.50E-09 | 1.77E-07 |

| | | | |
|---|---|---|---|
| BCL9 | 1.507905 | 6.53E-09 | 1.77E-07 |
| MATN4 | 4.760661 | 6.74E-09 | 1.82E-07 |
| RPN2 | -0.99591 | 6.81E-09 | 1.84E-07 |
| BUD23 | -1.13571 | 6.96E-09 | 1.87E-07 |
| GALNT17 | 1.278567 | 7.06E-09 | 1.89E-07 |
| LPL | -1.01142 | 7.39E-09 | 1.97E-07 |
| DACT1 | -0.96773 | 7.38E-09 | 1.97E-07 |
| PTPN5 | -3.90601 | 7.44E-09 | 1.98E-07 |
| ANAPC5 | -1.15837 | 7.54E-09 | 2.00E-07 |
| PRELID3B | 0.959232 | 7.70E-09 | 2.04E-07 |
| SLC25A25 | 1.077891 | 8.20E-09 | 2.17E-07 |
| DNAAF5 | -1.86353 | 8.35E-09 | 2.20E-07 |
| PARP1 | -1.12574 | 8.34E-09 | 2.20E-07 |
| AIFM1 | -0.9606 | 8.38E-09 | 2.20E-07 |
| GLMN | -1.16417 | 8.40E-09 | 2.20E-07 |
| ENSAPLG00000017513 | -1.15695 | 8.56E-09 | 2.24E-07 |
| KIAA1522 | 1.688898 | 8.75E-09 | 2.28E-07 |
| SKP2 | -1.0626 | 8.87E-09 | 2.31E-07 |
| VWA8 | -1.82543 | 9.01E-09 | 2.34E-07 |
| SLC46A2 | -1.1905 | 9.06E-09 | 2.34E-07 |
| C1orf43 | 0.976981 | 9.26E-09 | 2.38E-07 |
| MYBPC3 | 2.193647 | 9.25E-09 | 2.38E-07 |
| CREBBP | 1.322856 | 9.44E-09 | 2.42E-07 |
| GAL3ST1 | 7.61192 | 9.46E-09 | 2.42E-07 |
| RRH | 3.064412 | 9.66E-09 | 2.47E-07 |
| PRKAR1A | -1.08304 | 9.92E-09 | 2.53E-07 |
| PDGFB | 1.525663 | 9.98E-09 | 2.54E-07 |
| MDH1 | -0.99644 | 1.01E-08 | 2.56E-07 |
| ENSAPLG00000018174 | -1.00667 | 1.01E-08 | 2.56E-07 |
| DUSP10 | 0.906331 | 1.03E-08 | 2.61E-07 |
| ENSAPLG00000011075 | -1.62589 | 1.08E-08 | 2.73E-07 |
| SOCS1 | -1.76212 | 1.11E-08 | 2.80E-07 |
| UBA2 | -1.03009 | 1.12E-08 | 2.82E-07 |
| MTFR1 | 1.254055 | 1.15E-08 | 2.88E-07 |
| NOC4L | -1.30107 | 1.16E-08 | 2.89E-07 |
| HTRA3 | -1.02099 | 1.16E-08 | 2.89E-07 |
| AK4 | 1.278805 | 1.19E-08 | 2.94E-07 |
| BAMBI | -1.13994 | 1.19E-08 | 2.94E-07 |
| TMEM204 | 1.464942 | 1.21E-08 | 3.00E-07 |
| ENSAPLG00000018141 | 1.067568 | 1.22E-08 | 3.01E-07 |
| MCM6 | -1.37081 | 1.28E-08 | 3.16E-07 |
| PIGG | -1.48142 | 1.28E-08 | 3.16E-07 |
| PAPPA | 1.747396 | 1.29E-08 | 3.16E-07 |
| ENSAPLG00000010658 | -0.9281 | 1.32E-08 | 3.24E-07 |
| NT5DC2 | 1.081797 | 1.34E-08 | 3.26E-07 |

| | | | |
|---|---|---|---|
| NCOR2 | 1.039002 | 1.35E-08 | 3.28E-07 |
| TARS3 | -1.27615 | 1.43E-08 | 3.48E-07 |
| CNOT7 | 0.895225 | 1.47E-08 | 3.58E-07 |
| EXOC1 | -1.0477 | 1.56E-08 | 3.76E-07 |
| DHRS7 | -1.31427 | 1.65E-08 | 3.98E-07 |
| MAN2B2 | -1.11865 | 1.83E-08 | 4.41E-07 |
| RPL39 | 1.409176 | 1.90E-08 | 4.56E-07 |
| ENSAPLG00000027783 | 1.119421 | 1.90E-08 | 4.57E-07 |
| CSNK1G2 | 1.044011 | 1.92E-08 | 4.60E-07 |
| SRRM3 | 7.198966 | 1.93E-08 | 4.60E-07 |
| YARS2 | -1.17636 | 1.94E-08 | 4.61E-07 |
| HES4 | 1.45325 | 1.96E-08 | 4.67E-07 |
| C1orf21 | 1.018647 | 2.00E-08 | 4.74E-07 |
| ECHS1 | -1.55878 | 2.05E-08 | 4.85E-07 |
| CASTOR2 | 1.161395 | 2.07E-08 | 4.90E-07 |
| MEF2D | 1.705494 | 2.08E-08 | 4.90E-07 |
| RLF | 1.033721 | 2.09E-08 | 4.92E-07 |
| YTHDF2 | 1.21883 | 2.11E-08 | 4.95E-07 |
| B3GNT7 | 2.823738 | 2.12E-08 | 4.97E-07 |
| ACADL | -0.99699 | 2.13E-08 | 4.99E-07 |
| HS6ST1 | 0.916413 | 2.15E-08 | 5.01E-07 |
| CMKLR1 | 2.013898 | 2.18E-08 | 5.07E-07 |
| GOT2 | -0.94619 | 2.22E-08 | 5.16E-07 |
| WDR33 | 1.029171 | 2.29E-08 | 5.30E-07 |
| SLC6A15 | -1.17756 | 2.33E-08 | 5.39E-07 |
| ENSAPLG00000027047 | -0.95223 | 2.35E-08 | 5.42E-07 |
| KDM7A | 1.00106 | 2.45E-08 | 5.65E-07 |
| AP2A2 | -0.89208 | 2.50E-08 | 5.74E-07 |
| ENSAPLG00000009852 | -1.3876 | 2.55E-08 | 5.84E-07 |
| ENSAPLG00000015291 | 1.54049 | 2.55E-08 | 5.84E-07 |
| ZBTB26 | 1.000098 | 2.56E-08 | 5.84E-07 |
| FAM126B | -1.24604 | 2.63E-08 | 6.00E-07 |
| KLHL5 | 1.199551 | 2.68E-08 | 6.08E-07 |
| GPR68 | -1.16365 | 2.69E-08 | 6.10E-07 |
| SLC35G2 | 1.013307 | 2.73E-08 | 6.17E-07 |
| ENSAPLG00000022050 | -1.38979 | 2.79E-08 | 6.31E-07 |
| MCM4 | -1.12387 | 2.82E-08 | 6.36E-07 |
| IRF1 | 2.458533 | 2.89E-08 | 6.50E-07 |
| ENSAPLG00000019156 | 1.716495 | 3.03E-08 | 6.80E-07 |
| ENSAPLG00000021952 | -1.38099 | 3.05E-08 | 6.84E-07 |
| ARHGAP12 | 0.940371 | 3.12E-08 | 6.97E-07 |
| DIPK2B | 3.780551 | 3.12E-08 | 6.97E-07 |
| NCF2 | 4.808054 | 3.42E-08 | 7.62E-07 |
| VIM | -0.96773 | 3.43E-08 | 7.63E-07 |
| ENSAPLG00000014831 | -1.01185 | 3.46E-08 | 7.67E-07 |

| | | | |
|---|---|---|---|
| GHITM | -0.9395 | 3.54E-08 | 7.83E-07 |
| UBE2E3 | 1.063118 | 3.56E-08 | 7.86E-07 |
| IGFBP7 | -1.66428 | 3.56E-08 | 7.86E-07 |
| MSH2 | -1.09261 | 3.70E-08 | 8.15E-07 |
| STOX2 | 1.178895 | 3.73E-08 | 8.19E-07 |
| CNDP2 | -0.91529 | 3.76E-08 | 8.24E-07 |
| EXOSC9 | -1.01706 | 3.78E-08 | 8.27E-07 |
| OLFML3 | -1.36191 | 3.80E-08 | 8.29E-07 |
| MTA3 | -1.14854 | 3.90E-08 | 8.49E-07 |
| ENSAPLG00000026806 | 4.223115 | 3.92E-08 | 8.52E-07 |
| SCARB2 | -1.00796 | 3.94E-08 | 8.56E-07 |
| PARP6 | 1.128217 | 3.97E-08 | 8.59E-07 |
| DUSP5 | 1.083977 | 4.03E-08 | 8.64E-07 |
| PKNOX2 | 1.372819 | 4.03E-08 | 8.64E-07 |
| CHRNA5 | 1.849625 | 4.01E-08 | 8.64E-07 |
| CSNK2A2 | 0.986912 | 4.00E-08 | 8.64E-07 |
| AZI2 | 1.035179 | 4.03E-08 | 8.64E-07 |
| ENSAPLG00000028920 | 7.058956 | 4.11E-08 | 8.79E-07 |
| PDE7B | 1.085694 | 4.37E-08 | 9.34E-07 |
| FBLN2 | -1.2421 | 4.41E-08 | 9.39E-07 |
| ARRDC3 | 1.182346 | 4.46E-08 | 9.49E-07 |
| THBS2 | -1.44249 | 4.50E-08 | 9.55E-07 |
| RASL11B | -1.48858 | 4.58E-08 | 9.71E-07 |
| YWHAQ | -0.85775 | 4.60E-08 | 9.72E-07 |
| UCHL1 | -1.34056 | 4.63E-08 | 9.74E-07 |
| ENSAPLG00000019314 | 1.450461 | 4.63E-08 | 9.74E-07 |
| CASTOR1 | -1.67153 | 4.62E-08 | 9.74E-07 |
| EIF4G2 | 1.278713 | 4.75E-08 | 9.96E-07 |
| SLC25A16 | 0.989932 | 4.84E-08 | 1.01E-06 |
| SLC30A1 | 1.058048 | 5.01E-08 | 1.05E-06 |
| ACAA1 | -1.21438 | 5.03E-08 | 1.05E-06 |
| P4HA1 | -0.93882 | 5.05E-08 | 1.05E-06 |
| PHIP | 1.013572 | 5.25E-08 | 1.09E-06 |
| EI24 | 0.979005 | 5.31E-08 | 1.10E-06 |
| MCU | -1.06351 | 5.33E-08 | 1.10E-06 |
| ENSAPLG00000015846 | 1.192228 | 5.41E-08 | 1.12E-06 |
| CDK13 | 1.00936 | 5.46E-08 | 1.13E-06 |
| ENSAPLG00000006980 | 2.327928 | 5.64E-08 | 1.16E-06 |
| FGFRL1 | 1.037491 | 5.82E-08 | 1.20E-06 |
| CYP2U1 | -1.4641 | 6.02E-08 | 1.23E-06 |
| NF2 | 0.917201 | 6.05E-08 | 1.24E-06 |
| MAT1A | -1.46421 | 6.09E-08 | 1.25E-06 |
| SH3D19 | 1.742366 | 6.21E-08 | 1.27E-06 |
| ASPH | -1.06536 | 6.35E-08 | 1.29E-06 |
| RAD54L2 | 1.195812 | 6.42E-08 | 1.30E-06 |

| | | | |
|---|---|---|---|
| CTH | -1.33409 | 6.47E-08 | 1.31E-06 |
| WDR20 | 1.008583 | 6.55E-08 | 1.33E-06 |
| ABCB6 | -1.21951 | 6.63E-08 | 1.34E-06 |
| SYNC | 0.980401 | 6.67E-08 | 1.34E-06 |
| ENSAPLG00000009942 | 6.985933 | 6.79E-08 | 1.37E-06 |
| MAS1 | 7.329796 | 6.85E-08 | 1.38E-06 |
| TPM1 | -1.03562 | 6.98E-08 | 1.40E-06 |
| PRIM2 | -1.28297 | 6.97E-08 | 1.40E-06 |
| TSPAN3 | -0.93162 | 7.18E-08 | 1.44E-06 |
| ENSAPLG00000016936 | 1.106911 | 7.21E-08 | 1.44E-06 |
| CA2 | 1.166853 | 7.31E-08 | 1.46E-06 |
| ENSAPLG00000007028 | -0.94216 | 7.33E-08 | 1.46E-06 |
| PTX3 | -1.60582 | 7.64E-08 | 1.51E-06 |
| HEXD | -1.55985 | 7.62E-08 | 1.51E-06 |
| DYSF | 1.184174 | 7.69E-08 | 1.52E-06 |
| HTR2B | 1.072081 | 7.83E-08 | 1.54E-06 |
| ENSAPLG00000020473 | -1.29369 | 7.85E-08 | 1.55E-06 |
| TRIB2 | 0.867807 | 7.95E-08 | 1.56E-06 |
| TUFT1 | 0.894881 | 8.18E-08 | 1.61E-06 |
| LONRF3 | 1.06042 | 8.42E-08 | 1.65E-06 |
| DEDD | 1.259279 | 8.45E-08 | 1.65E-06 |
| HMBOX1 | 1.011651 | 8.59E-08 | 1.68E-06 |
| ENSAPLG00000030007 | -1.23386 | 8.61E-08 | 1.68E-06 |
| ABHD13 | 0.890778 | 8.69E-08 | 1.69E-06 |
| INHBB | 2.357534 | 8.75E-08 | 1.70E-06 |
| ENSAPLG00000030664 | 2.544803 | 8.74E-08 | 1.70E-06 |
| PHF12 | 1.325582 | 9.30E-08 | 1.80E-06 |
| HRAS | 1.007233 | 9.37E-08 | 1.81E-06 |
| COMTD1 | -1.05317 | 9.61E-08 | 1.85E-06 |
| LONP2 | -1.03198 | 9.73E-08 | 1.87E-06 |
| ACSL3 | -0.8843 | 9.76E-08 | 1.87E-06 |
| RCAN3 | 0.886022 | 9.83E-08 | 1.88E-06 |
| MDM4 | 1.212153 | 9.94E-08 | 1.90E-06 |
| ENSAPLG00000017191 | 1.207365 | 9.94E-08 | 1.90E-06 |
| GNAI1 | -0.93157 | 1.02E-07 | 1.94E-06 |
| PIP4K2A | 0.951489 | 1.03E-07 | 1.97E-06 |
| MYG1 | -1.34253 | 1.05E-07 | 2.00E-06 |
| ENSAPLG00000021054 | -1.12798 | 1.07E-07 | 2.04E-06 |
| FSTL1 | -0.99389 | 1.08E-07 | 2.04E-06 |
| ENDOG | -1.01451 | 1.10E-07 | 2.08E-06 |
| CTSO | -1.20317 | 1.12E-07 | 2.11E-06 |
| AKR7L | -1.07158 | 1.12E-07 | 2.11E-06 |
| ENSAPLG00000005235 | 1.075018 | 1.13E-07 | 2.13E-06 |
| PLTP | -1.26737 | 1.13E-07 | 2.13E-06 |
| CCN1 | -0.94425 | 1.14E-07 | 2.14E-06 |

| | | | |
|---|---|---|---|
| ATP6V1H | -0.95522 | 1.16E-07 | 2.17E-06 |
| PFKP | -0.88395 | 1.19E-07 | 2.23E-06 |
| BCAP29 | -1.01263 | 1.20E-07 | 2.25E-06 |
| HELLS | -1.26107 | 1.25E-07 | 2.32E-06 |
| SMYD2 | -1.04305 | 1.27E-07 | 2.37E-06 |
| CXCL12 | 1.144468 | 1.28E-07 | 2.37E-06 |
| TNS1 | 1.35479 | 1.35E-07 | 2.50E-06 |
| FKBP5 | 1.058852 | 1.43E-07 | 2.65E-06 |
| ENSAPLG00000008364 | -0.87981 | 1.45E-07 | 2.68E-06 |
| VPS54 | -1.09384 | 1.48E-07 | 2.72E-06 |
| ERP29 | -1.03469 | 1.51E-07 | 2.77E-06 |
| WNT5B | -0.88 | 1.55E-07 | 2.84E-06 |
| NAE1 | -0.96293 | 1.57E-07 | 2.89E-06 |
| ENSAPLG00000018235 | 1.479785 | 1.60E-07 | 2.93E-06 |
| ATP5F1B | -0.89316 | 1.64E-07 | 2.99E-06 |
| SEPTIN9 | 1.40871 | 1.66E-07 | 3.02E-06 |
| SURF1 | -1.12697 | 1.66E-07 | 3.03E-06 |
| MYBL2 | 1.193528 | 1.68E-07 | 3.05E-06 |
| POLR3B | -1.0747 | 1.69E-07 | 3.07E-06 |
| NMD3 | -0.8972 | 1.71E-07 | 3.09E-06 |
| GSE1 | 1.499305 | 1.71E-07 | 3.09E-06 |
| PCCA | -1.09761 | 1.72E-07 | 3.11E-06 |
| CHSY1 | 1.424534 | 1.72E-07 | 3.11E-06 |
| BCHE | -1.42427 | 1.74E-07 | 3.14E-06 |
| RAP1GDS1 | -0.84284 | 1.77E-07 | 3.18E-06 |
| ENSAPLG00000003517 | 0.903545 | 1.78E-07 | 3.20E-06 |
| FOLH1 | -1.1648 | 1.83E-07 | 3.27E-06 |
| PAPSS1 | -0.87775 | 1.85E-07 | 3.31E-06 |
| MTF1 | 0.882125 | 1.86E-07 | 3.31E-06 |
| ENSAPLG00000017536 | 3.517483 | 1.85E-07 | 3.31E-06 |
| NUCB2 | -0.96206 | 1.90E-07 | 3.39E-06 |
| TAX1BP3 | 0.931912 | 1.91E-07 | 3.40E-06 |
| ENSAPLG00000020033 | 1.372992 | 1.92E-07 | 3.41E-06 |
| TMEM184B | 1.093545 | 1.95E-07 | 3.46E-06 |
| EHBP1 | 0.98956 | 1.97E-07 | 3.48E-06 |
| RHOU | 1.490485 | 1.97E-07 | 3.48E-06 |
| ENSAPLG00000019053 | 1.322291 | 1.98E-07 | 3.48E-06 |
| KXD1 | 1.082681 | 2.00E-07 | 3.52E-06 |
| MGAT2 | -0.99179 | 2.00E-07 | 3.52E-06 |
| OTOR | 7.189284 | 2.01E-07 | 3.53E-06 |
| SLC25A39 | 0.913294 | 2.02E-07 | 3.54E-06 |
| ENSAPLG00000028455 | 1.518277 | 2.05E-07 | 3.58E-06 |
| MAFK | 1.273174 | 2.07E-07 | 3.61E-06 |
| ENSAPLG00000021067 | 2.689363 | 2.10E-07 | 3.66E-06 |
| ENSAPLG00000003255 | -1.25299 | 2.14E-07 | 3.72E-06 |

| | | | |
|---|---|---|---|
| AGL | -1.15348 | 2.14E-07 | 3.72E-06 |
| DGKD | 0.901039 | 2.15E-07 | 3.74E-06 |
| SPRED1 | 0.890141 | 2.19E-07 | 3.80E-06 |
| SENP5 | 0.814167 | 2.21E-07 | 3.82E-06 |
| CDHR1 | 7.104399 | 2.23E-07 | 3.84E-06 |
| ENSAPLG00000007778 | -1.02837 | 2.23E-07 | 3.85E-06 |
| USP53 | 1.21498 | 2.24E-07 | 3.86E-06 |
| GATB | -1.78646 | 2.29E-07 | 3.93E-06 |
| POFUT1 | -1.06506 | 2.29E-07 | 3.93E-06 |
| RIC3 | -1.24155 | 2.31E-07 | 3.96E-06 |
| KCNS1 | 4.526742 | 2.38E-07 | 4.07E-06 |
| KANSL1 | 1.537603 | 2.41E-07 | 4.11E-06 |
| P3H1 | -1.0731 | 2.46E-07 | 4.20E-06 |
| MORC2 | 1.285283 | 2.47E-07 | 4.20E-06 |
| PI4KB | 1.219723 | 2.50E-07 | 4.24E-06 |
| WDR77 | -0.93442 | 2.50E-07 | 4.24E-06 |
| UBL3 | 0.98474 | 2.50E-07 | 4.24E-06 |
| WDR53 | -1.05558 | 2.51E-07 | 4.25E-06 |
| DLX5 | 1.220578 | 2.52E-07 | 4.26E-06 |
| FGFR1 | 0.86915 | 2.56E-07 | 4.31E-06 |
| WEE1 | 1.342896 | 2.57E-07 | 4.32E-06 |
| TRIM45 | -1.8778 | 2.58E-07 | 4.34E-06 |
| ENSAPLG00000007227 | 3.26385 | 2.62E-07 | 4.40E-06 |
| MCM2 | -1.25984 | 2.67E-07 | 4.47E-06 |
| ZPLD1 | 1.34412 | 2.67E-07 | 4.47E-06 |
| TRRAP | 1.107088 | 2.72E-07 | 4.55E-06 |
| TMEM260 | -1.31438 | 2.75E-07 | 4.59E-06 |
| ACAT1 | -0.95919 | 2.77E-07 | 4.61E-06 |
| SERINC1 | -0.85505 | 2.78E-07 | 4.61E-06 |
| MPI | -1.01573 | 2.78E-07 | 4.61E-06 |
| MLLT6 | 1.493835 | 2.86E-07 | 4.74E-06 |
| EP300 | 1.041331 | 2.87E-07 | 4.75E-06 |
| KCNJ4 | 4.052922 | 2.87E-07 | 4.75E-06 |
| CDC23 | -0.96718 | 2.88E-07 | 4.75E-06 |
| ACAT2 | -1.06491 | 2.88E-07 | 4.75E-06 |
| ENSAPLG00000016374 | 1.007931 | 3.00E-07 | 4.94E-06 |
| COLGALT1 | -1.17463 | 3.04E-07 | 5.00E-06 |
| TACC1 | 1.08106 | 3.05E-07 | 5.01E-06 |
| ENSAPLG00000018153 | 0.937401 | 3.07E-07 | 5.03E-06 |
| ENSAPLG00000008838 | 0.851701 | 3.09E-07 | 5.05E-06 |
| HSD17B12 | -0.86446 | 3.10E-07 | 5.06E-06 |
| ATAD1 | -0.9349 | 3.13E-07 | 5.10E-06 |
| APC2 | 7.051962 | 3.15E-07 | 5.13E-06 |
| FAM149B1 | 1.369512 | 3.15E-07 | 5.13E-06 |
| RAB3IL1 | 1.043085 | 3.20E-07 | 5.19E-06 |

| | | | |
|---|---|---|---|
| SDR42E1 | -1.54387 | 3.23E-07 | 5.23E-06 |
| NDC1 | -0.98497 | 3.23E-07 | 5.23E-06 |
| RTCA | -0.97718 | 3.22E-07 | 5.23E-06 |
| WIPI1 | 0.833873 | 3.28E-07 | 5.30E-06 |
| DYRK2 | 1.020367 | 3.34E-07 | 5.38E-06 |
| EMC7 | -0.96513 | 3.36E-07 | 5.40E-06 |
| TNFSF8 | 7.085821 | 3.38E-07 | 5.43E-06 |
| EXTL3 | 1.10499 | 3.51E-07 | 5.62E-06 |
| ZFP36L1 | 1.297999 | 3.51E-07 | 5.62E-06 |
| NNT | -1.31111 | 3.56E-07 | 5.70E-06 |
| SERPINF1 | -1.15686 | 3.63E-07 | 5.80E-06 |
| HP1BP3 | -0.9144 | 3.64E-07 | 5.80E-06 |
| ENSAPLG00000008343 | 1.88152 | 3.72E-07 | 5.92E-06 |
| SLC35A3 | -1.01268 | 3.75E-07 | 5.97E-06 |
| PCYT2 | 0.867384 | 3.80E-07 | 6.03E-06 |
| RNF182 | 2.256905 | 3.99E-07 | 6.32E-06 |
| DDX49 | -1.01257 | 3.99E-07 | 6.32E-06 |
| WWP1 | 1.153107 | 4.04E-07 | 6.39E-06 |
| ENSAPLG00000006443 | -0.86461 | 4.05E-07 | 6.39E-06 |
| SPTLC1 | -0.93492 | 4.07E-07 | 6.41E-06 |
| DAB2IP | 1.474468 | 4.08E-07 | 6.42E-06 |
| NKX3-2 | 1.477992 | 4.09E-07 | 6.42E-06 |
| PGM2 | -0.89216 | 4.15E-07 | 6.51E-06 |
| RSRP1 | 1.241557 | 4.23E-07 | 6.64E-06 |
| ENSAPLG00000013227 | 0.986428 | 4.25E-07 | 6.66E-06 |
| FAP | -0.96383 | 4.34E-07 | 6.79E-06 |
| TK2 | 1.006369 | 4.38E-07 | 6.83E-06 |
| PLAGL1 | 1.030352 | 4.39E-07 | 6.84E-06 |
| EIF4EBP1 | 1.13815 | 4.49E-07 | 6.99E-06 |
| PCMTD1 | 0.92915 | 4.51E-07 | 7.01E-06 |
| FIG4 | -1.28187 | 4.54E-07 | 7.05E-06 |
| HADHA | -0.80122 | 4.56E-07 | 7.07E-06 |
| WDR75 | -0.94505 | 4.80E-07 | 7.43E-06 |
| CTDSPL | 1.137636 | 4.81E-07 | 7.44E-06 |
| EBAG9 | 1.010105 | 4.89E-07 | 7.54E-06 |
| GPI | -0.89672 | 4.99E-07 | 7.68E-06 |
| ENSAPLG00000008579 | -0.91623 | 5.09E-07 | 7.82E-06 |
| HADH | -0.93208 | 5.17E-07 | 7.94E-06 |
| SKI | 1.272718 | 5.18E-07 | 7.94E-06 |
| RPL3L | 2.641828 | 5.19E-07 | 7.95E-06 |
| RAMP3 | 1.204367 | 5.23E-07 | 8.01E-06 |
| AGPAT4 | -1.06954 | 5.32E-07 | 8.13E-06 |
| DLD | -0.85974 | 5.37E-07 | 8.19E-06 |
| PTGER3 | 3.482524 | 5.54E-07 | 8.44E-06 |
| ENSAPLG00000026048 | -1.09906 | 5.71E-07 | 8.68E-06 |

| | | | |
|---|---|---|---|
| ENSAPLG00000001512 | 1.933583 | 5.87E-07 | 8.90E-06 |
| GFPT2 | -1.04098 | 5.86E-07 | 8.90E-06 |
| TXNRD3 | -0.91651 | 5.93E-07 | 8.99E-06 |
| ZCCHC24 | 0.951758 | 5.97E-07 | 9.04E-06 |
| ACTG2 | -1.19001 | 5.99E-07 | 9.05E-06 |
| TMEM250 | 1.143405 | 6.02E-07 | 9.09E-06 |
| PCYOX1L | -1.15507 | 6.06E-07 | 9.14E-06 |
| EGFLAM | -1.11741 | 6.15E-07 | 9.25E-06 |
| ENSAPLG00000013374 | 0.914096 | 6.23E-07 | 9.36E-06 |
| SYT17 | 0.953335 | 6.27E-07 | 9.40E-06 |
| WTIP | 1.07919 | 6.28E-07 | 9.41E-06 |
| KIN | -1.25656 | 6.30E-07 | 9.42E-06 |
| ENSAPLG00000016336 | 4.09568 | 6.51E-07 | 9.73E-06 |
| MCF2L2 | 3.005826 | 6.56E-07 | 9.79E-06 |
| MMUT | -0.97527 | 6.66E-07 | 9.93E-06 |
| ASPG | -1.20329 | 6.75E-07 | 1.00E-05 |
| USP4 | -0.98587 | 6.86E-07 | 1.02E-05 |
| FRS3 | 1.650909 | 6.85E-07 | 1.02E-05 |
| ENSAPLG00000004580 | 1.374434 | 6.89E-07 | 1.02E-05 |
| SMIM14 | 0.93703 | 6.88E-07 | 1.02E-05 |
| CTNNA1 | -0.79063 | 6.91E-07 | 1.02E-05 |
| RUNX2 | 1.335763 | 6.93E-07 | 1.02E-05 |
| PSAP | -1.04793 | 6.93E-07 | 1.02E-05 |
| LPAR4 | 1.461678 | 6.97E-07 | 1.03E-05 |
| ALDH18A1 | -0.84094 | 7.01E-07 | 1.03E-05 |
| CRB2 | 1.633466 | 7.06E-07 | 1.04E-05 |
| ZC3H12B | 1.182363 | 7.09E-07 | 1.04E-05 |
| SPG7 | 0.806375 | 7.12E-07 | 1.04E-05 |
| CBR4 | -1.0766 | 7.18E-07 | 1.05E-05 |
| ENSAPLG00000013478 | -0.95789 | 7.46E-07 | 1.09E-05 |
| JAM3 | -1.07084 | 7.54E-07 | 1.10E-05 |
| LTA4H | -0.82011 | 7.55E-07 | 1.10E-05 |
| SRPX2 | -1.15632 | 7.59E-07 | 1.10E-05 |
| AFAP1 | -0.85084 | 7.90E-07 | 1.15E-05 |
| KRT75 | -2.28053 | 7.89E-07 | 1.15E-05 |
| AGK | -1.06558 | 7.96E-07 | 1.15E-05 |
| NUMA1 | 1.233519 | 8.03E-07 | 1.16E-05 |
| ZDHHC13 | -0.99977 | 8.07E-07 | 1.16E-05 |
| UTP25 | -0.94794 | 8.07E-07 | 1.16E-05 |
| ABLIM2 | 1.100469 | 8.09E-07 | 1.17E-05 |
| IMPDH2 | -0.80691 | 8.20E-07 | 1.18E-05 |
| ANXA6 | -0.82704 | 8.23E-07 | 1.18E-05 |
| SHROOM3 | 1.182453 | 8.25E-07 | 1.18E-05 |
| ZFHX4 | 0.938002 | 8.38E-07 | 1.20E-05 |
| GRIK3 | 4.537488 | 8.67E-07 | 1.24E-05 |

| | | | |
|---|---|---|---|
| ENSAPLG00000024348 | -1.2155 | 8.82E-07 | 1.26E-05 |
| SEL1L | -1.22478 | 8.84E-07 | 1.26E-05 |
| ARNTL | -1.07438 | 8.87E-07 | 1.27E-05 |
| GXYLT2 | 0.964242 | 8.98E-07 | 1.28E-05 |
| ENSAPLG00000006772 | 2.499659 | 9.03E-07 | 1.28E-05 |
| YTHDF1 | 1.226615 | 9.16E-07 | 1.30E-05 |
| ABCC4 | -1.09247 | 9.25E-07 | 1.31E-05 |
| PSMD10 | -1.09041 | 9.31E-07 | 1.32E-05 |
| GALNT12 | -1.64947 | 9.65E-07 | 1.37E-05 |
| CAT | -1.05162 | 9.79E-07 | 1.38E-05 |
| ENSAPLG00000016255 | 0.811821 | 9.84E-07 | 1.39E-05 |
| FBRSL1 | 0.942027 | 9.90E-07 | 1.40E-05 |
| UBE2H | 0.990654 | 1.00E-06 | 1.41E-05 |
| ZDHHC3 | 0.886714 | 1.03E-06 | 1.45E-05 |
| AIMP2 | -0.9394 | 1.06E-06 | 1.48E-05 |
| FMN1 | 1.051137 | 1.06E-06 | 1.48E-05 |
| PEX13 | -1.05075 | 1.10E-06 | 1.54E-05 |
| PMS2 | -0.90529 | 1.11E-06 | 1.56E-05 |
| NUDT3 | 1.621045 | 1.13E-06 | 1.58E-05 |
| MSRB3 | 1.046385 | 1.13E-06 | 1.58E-05 |
| EZR | -0.77608 | 1.15E-06 | 1.60E-05 |
| TXNDC15 | -0.98671 | 1.15E-06 | 1.60E-05 |
| PLK2 | -0.79839 | 1.16E-06 | 1.61E-05 |
| IDH3A | -0.8144 | 1.16E-06 | 1.61E-05 |
| SIDT2 | 1.102888 | 1.16E-06 | 1.61E-05 |
| GMPS | -0.81361 | 1.17E-06 | 1.63E-05 |
| SUMO1 | 0.922385 | 1.18E-06 | 1.64E-05 |
| GJA1 | -0.85037 | 1.21E-06 | 1.67E-05 |
| FZD9 | 0.993975 | 1.24E-06 | 1.71E-05 |
| RHBDD1 | -1.52498 | 1.26E-06 | 1.74E-05 |
| GFM1 | -0.86617 | 1.27E-06 | 1.74E-05 |
| TRMT10A | -1.0954 | 1.27E-06 | 1.75E-05 |
| GGT7 | 3.323362 | 1.27E-06 | 1.75E-05 |
| PTP4A2 | 0.908005 | 1.28E-06 | 1.76E-05 |
| SPSB4 | -0.98715 | 1.29E-06 | 1.76E-05 |
| RWDD4 | 0.940792 | 1.30E-06 | 1.77E-05 |
| ENSAPLG00000009022 | 0.907177 | 1.31E-06 | 1.79E-05 |
| OGFOD1 | -0.95445 | 1.32E-06 | 1.79E-05 |
| DRG1 | -0.89962 | 1.35E-06 | 1.83E-05 |
| ALG12 | -0.7777 | 1.36E-06 | 1.85E-05 |
| PYROXD1 | -1.30048 | 1.39E-06 | 1.89E-05 |
| BLMH | -0.78559 | 1.41E-06 | 1.92E-05 |
| SHOC2 | 0.848084 | 1.43E-06 | 1.93E-05 |
| NCLN | -0.80922 | 1.43E-06 | 1.93E-05 |
| LMBRD1 | -0.88889 | 1.43E-06 | 1.93E-05 |

| | | | |
|---|---|---|---|
| MTUS2 | 2.867421 | 1.44E-06 | 1.95E-05 |
| SLC25A4 | -1.04671 | 1.44E-06 | 1.95E-05 |
| ENSAPLG00000029546 | -0.83319 | 1.46E-06 | 1.96E-05 |
| VAV2 | 0.932308 | 1.47E-06 | 1.97E-05 |
| MTTP | 6.848201 | 1.47E-06 | 1.98E-05 |
| USPL1 | 1.55018 | 1.48E-06 | 1.98E-05 |
| PRKCD | -0.87801 | 1.49E-06 | 1.99E-05 |
| VWF | 1.016526 | 1.49E-06 | 1.99E-05 |
| PLRG1 | -0.91069 | 1.48E-06 | 1.99E-05 |
| ODF2 | 1.035552 | 1.49E-06 | 1.99E-05 |
| ENSAPLG00000008168 | 1.4438 | 1.50E-06 | 2.00E-05 |
| CTNNBL1 | -0.8478 | 1.51E-06 | 2.01E-05 |
| C10orf71 | 1.006177 | 1.51E-06 | 2.01E-05 |
| IPO5 | -0.8223 | 1.52E-06 | 2.02E-05 |
| HEXIM1 | 1.321665 | 1.53E-06 | 2.03E-05 |
| CD109 | -2.13262 | 1.54E-06 | 2.04E-05 |
| ENSAPLG00000005387 | 1.048329 | 1.55E-06 | 2.06E-05 |
| PHF6 | -0.85037 | 1.58E-06 | 2.08E-05 |
| INTS14 | -0.96637 | 1.61E-06 | 2.13E-05 |
| AP5M1 | 0.926868 | 1.62E-06 | 2.13E-05 |
| AVL9 | -1.09567 | 1.62E-06 | 2.13E-05 |
| HSDL2 | -1.04045 | 1.62E-06 | 2.14E-05 |
| MTERF3 | -0.96803 | 1.63E-06 | 2.14E-05 |
| ARNTL2 | 0.961376 | 1.64E-06 | 2.16E-05 |
| ENSAPLG00000002228 | -0.98264 | 1.65E-06 | 2.16E-05 |
| DCP1A | 0.934907 | 1.65E-06 | 2.16E-05 |
| MYH10 | -0.86066 | 1.66E-06 | 2.17E-05 |
| BMPER | -1.1173 | 1.67E-06 | 2.19E-05 |
| RGL1 | 1.175118 | 1.69E-06 | 2.21E-05 |
| UNC50 | -0.98785 | 1.72E-06 | 2.24E-05 |
| PHLDA2 | 1.137312 | 1.76E-06 | 2.28E-05 |
| ADA2 | -1.83803 | 1.77E-06 | 2.30E-05 |
| SERPINB1 | 1.631446 | 1.78E-06 | 2.31E-05 |
| BDNF | 0.994638 | 1.81E-06 | 2.34E-05 |
| ENSAPLG00000026183 | -1.13443 | 1.81E-06 | 2.35E-05 |
| MRPS27 | -0.97661 | 1.83E-06 | 2.37E-05 |
| MAP2K3 | 0.885778 | 1.83E-06 | 2.37E-05 |
| ENSAPLG00000009366 | 1.104712 | 1.86E-06 | 2.40E-05 |
| PIP5K1B | 1.322112 | 1.87E-06 | 2.41E-05 |
| SEC61A2 | -0.83377 | 1.87E-06 | 2.41E-05 |
| TNRC6A | 1.206249 | 1.90E-06 | 2.44E-05 |
| COQ6 | -0.93683 | 1.90E-06 | 2.44E-05 |
| SLC17A5 | -1.03517 | 1.91E-06 | 2.45E-05 |
| ENSAPLG00000024724 | 1.083223 | 1.93E-06 | 2.47E-05 |
| ENSAPLG00000011828 | 0.968065 | 1.93E-06 | 2.48E-05 |

| | | | |
|---|---|---|---|
| PRDM4 | 0.922412 | 1.94E-06 | 2.48E-05 |
| FNIP1 | 1.262075 | 1.95E-06 | 2.50E-05 |
| CNOT4 | 0.994955 | 1.98E-06 | 2.52E-05 |
| ELP3 | -0.87878 | 2.01E-06 | 2.56E-05 |
| ENSAPLG00000013634 | -0.86844 | 2.04E-06 | 2.60E-05 |
| SIL1 | -1.21067 | 2.06E-06 | 2.61E-05 |
| HIVEP3 | 1.183418 | 2.06E-06 | 2.61E-05 |
| KLF13 | 1.220399 | 2.07E-06 | 2.62E-05 |
| ENSAPLG00000015183 | 0.991234 | 2.09E-06 | 2.65E-05 |
| ENSAPLG00000029052 | 1.00332 | 2.10E-06 | 2.66E-05 |
| ITGAV | -1.11976 | 2.11E-06 | 2.67E-05 |
| INSR | 1.688753 | 2.18E-06 | 2.75E-05 |
| GSS | -1.27252 | 2.18E-06 | 2.75E-05 |
| KRT12 | 6.387996 | 2.22E-06 | 2.79E-05 |
| COPG2 | -0.83925 | 2.22E-06 | 2.80E-05 |
| RD3 | 3.865547 | 2.24E-06 | 2.82E-05 |
| RRBP1 | -0.82684 | 2.26E-06 | 2.84E-05 |
| ENSAPLG00000009692 | 1.062581 | 2.27E-06 | 2.84E-05 |
| ENSAPLG00000026526 | -0.84513 | 2.27E-06 | 2.84E-05 |
| ERP44 | -0.90815 | 2.28E-06 | 2.85E-05 |
| RSL1D1 | -0.90992 | 2.30E-06 | 2.87E-05 |
| GRIA2 | -1.13677 | 2.31E-06 | 2.89E-05 |
| WT1 | 1.772425 | 2.33E-06 | 2.90E-05 |
| ANKRD12 | 1.175797 | 2.36E-06 | 2.94E-05 |
| ITGA4 | -1.26788 | 2.43E-06 | 3.02E-05 |
| ENSAPLG00000013064 | 0.960377 | 2.43E-06 | 3.02E-05 |
| EPHX1 | -1.28719 | 2.50E-06 | 3.10E-05 |
| PPIL4 | -1.23868 | 2.50E-06 | 3.10E-05 |
| ENSAPLG00000018690 | 0.990926 | 2.51E-06 | 3.11E-05 |
| RRAGD | 0.77978 | 2.52E-06 | 3.11E-05 |
| CAV3 | 6.71647 | 2.54E-06 | 3.13E-05 |
| ZWILCH | -0.97352 | 2.56E-06 | 3.16E-05 |
| SPPL2A | -0.747 | 2.58E-06 | 3.18E-05 |
| PHETA2 | 1.589535 | 2.60E-06 | 3.20E-05 |
| ZDHHC17 | 0.802549 | 2.65E-06 | 3.26E-05 |
| ENSAPLG00000002850 | -0.90855 | 2.66E-06 | 3.26E-05 |
| ZNF384 | 1.420644 | 2.71E-06 | 3.33E-05 |
| TOMM40L | 1.242896 | 2.74E-06 | 3.35E-05 |
| STAR | 4.357496 | 2.76E-06 | 3.37E-05 |
| OTX2 | 6.337214 | 2.76E-06 | 3.37E-05 |
| CCN3 | -1.58118 | 2.80E-06 | 3.42E-05 |
| LRPAP1 | -1.17431 | 2.80E-06 | 3.42E-05 |
| TMEM131L | -1.07722 | 2.82E-06 | 3.44E-05 |
| SALL4 | 1.614535 | 2.85E-06 | 3.47E-05 |
| ENSAPLG00000006104 | 1.652091 | 2.87E-06 | 3.49E-05 |

| | | | |
|---|---|---|---|
| GALNS | -0.79775 | 2.91E-06 | 3.54E-05 |
| CERT1 | -1.00458 | 2.99E-06 | 3.63E-05 |
| CYP51A1 | -0.94283 | 3.00E-06 | 3.63E-05 |
| ENSAPLG00000024644 | 6.297489 | 3.03E-06 | 3.67E-05 |
| RPL37 | 0.903622 | 3.06E-06 | 3.70E-05 |
| ENSAPLG00000028526 | -1.01042 | 3.06E-06 | 3.70E-05 |
| STC2 | 0.890649 | 3.07E-06 | 3.71E-05 |
| MAML1 | 1.841911 | 3.11E-06 | 3.75E-05 |
| ENSAPLG00000026428 | 1.11112 | 3.13E-06 | 3.77E-05 |
| ENSAPLG00000007015 | -0.81819 | 3.14E-06 | 3.78E-05 |
| GADL1 | 2.085491 | 3.15E-06 | 3.78E-05 |
| ACSS2 | -1.24459 | 3.16E-06 | 3.79E-05 |
| ENSAPLG00000011402 | -0.77934 | 3.17E-06 | 3.80E-05 |
| DLC1 | 0.884477 | 3.25E-06 | 3.89E-05 |
| ATG9B | 1.12057 | 3.31E-06 | 3.96E-05 |
| COG6 | -0.96371 | 3.32E-06 | 3.97E-05 |
| C1QTNF4 | -1.61831 | 3.33E-06 | 3.97E-05 |
| TAB1 | -0.91573 | 3.34E-06 | 3.98E-05 |
| RDH10 | -0.90466 | 3.36E-06 | 4.00E-05 |
| MSX1 | 0.994283 | 3.37E-06 | 4.00E-05 |
| TBC1D19 | -1.37956 | 3.43E-06 | 4.07E-05 |
| PRTG | 2.450934 | 3.58E-06 | 4.24E-05 |
| SLC25A46 | -0.95056 | 3.59E-06 | 4.25E-05 |
| EIF4A3 | -0.85067 | 3.63E-06 | 4.29E-05 |
| CERCAM | -0.86637 | 3.63E-06 | 4.29E-05 |
| GTF3C3 | -1.07198 | 3.68E-06 | 4.35E-05 |
| ITM2B | -0.83205 | 3.70E-06 | 4.36E-05 |
| ALG6 | -0.92444 | 3.70E-06 | 4.36E-05 |
| TNFAIP6 | -1.04933 | 3.71E-06 | 4.36E-05 |
| NIBAN2 | 0.969237 | 3.79E-06 | 4.45E-05 |
| FBXO30 | -1.10794 | 3.79E-06 | 4.45E-05 |
| FAM120B | -1.29423 | 3.81E-06 | 4.47E-05 |
| ENOPH1 | -0.93095 | 3.85E-06 | 4.51E-05 |
| IVD | -0.89645 | 3.97E-06 | 4.65E-05 |
| ENSAPLG00000003957 | 5.544549 | 3.99E-06 | 4.67E-05 |
| LRIT1 | 6.643794 | 4.01E-06 | 4.68E-05 |
| CIAO3 | -0.92241 | 4.00E-06 | 4.68E-05 |
| PLXND1 | 1.472683 | 4.04E-06 | 4.72E-05 |
| PLAA | -0.84768 | 4.12E-06 | 4.80E-05 |
| POC1A | -1.17793 | 4.13E-06 | 4.81E-05 |
| GSTCD | -1.27404 | 4.20E-06 | 4.89E-05 |
| MRPL3 | -1.05429 | 4.22E-06 | 4.90E-05 |
| ARID3B | 1.478305 | 4.23E-06 | 4.91E-05 |
| VPS35 | -0.81804 | 4.25E-06 | 4.93E-05 |
| TRMT2A | -1.0197 | 4.32E-06 | 5.00E-05 |

| | | | |
|---|---|---|---|
| TEKT5 | -1.70084 | 4.37E-06 | 5.06E-05 |
| API5 | -0.79695 | 4.41E-06 | 5.09E-05 |
| ENSAPLG00000018136 | -1.15435 | 4.41E-06 | 5.09E-05 |
| EED | -0.84069 | 4.43E-06 | 5.10E-05 |
| ERGIC2 | 0.700726 | 4.45E-06 | 5.12E-05 |
| ADAM10 | -0.76737 | 4.45E-06 | 5.12E-05 |
| RPS11 | 0.997041 | 4.51E-06 | 5.18E-05 |
| SLCO4A1 | -1.07219 | 4.53E-06 | 5.20E-05 |
| SERINC3 | -0.7238 | 4.53E-06 | 5.20E-05 |
| SCAF4 | 0.895717 | 4.54E-06 | 5.20E-05 |
| SATB1 | 1.072445 | 4.55E-06 | 5.21E-05 |
| GPBP1L1 | 0.773085 | 4.61E-06 | 5.27E-05 |
| PRELID3A | 0.801415 | 4.66E-06 | 5.31E-05 |
| ENSAPLG00000011334 | 2.395381 | 4.65E-06 | 5.31E-05 |
| MAEA | -0.85274 | 4.65E-06 | 5.31E-05 |
| SLC1A4 | -1.00319 | 4.69E-06 | 5.33E-05 |
| RASSF9 | -1.05076 | 4.75E-06 | 5.39E-05 |
| CAB39L | 0.903909 | 4.75E-06 | 5.39E-05 |
| OTUD6B | 0.738656 | 4.74E-06 | 5.39E-05 |
| SLC9A2 | 6.274938 | 4.77E-06 | 5.41E-05 |
| SLC25A33 | -0.8978 | 4.79E-06 | 5.43E-05 |
| GEMIN5 | -0.85162 | 4.83E-06 | 5.46E-05 |
| EXOSC3 | -1.19677 | 4.89E-06 | 5.52E-05 |
| APLP2 | -0.74229 | 4.90E-06 | 5.53E-05 |
| MTFR1L | 0.850708 | 4.90E-06 | 5.53E-05 |
| LANCL1 | -0.83194 | 4.95E-06 | 5.58E-05 |
| TCF7L2 | 1.153538 | 4.98E-06 | 5.60E-05 |
| ENSAPLG00000031160 | 1.002255 | 4.98E-06 | 5.60E-05 |
| NASP | -1.10422 | 5.06E-06 | 5.69E-05 |
| NRG1 | 1.439929 | 5.15E-06 | 5.78E-05 |
| TBX18 | 1.112352 | 5.24E-06 | 5.87E-05 |
| NPAS2 | 1.229225 | 5.36E-06 | 6.00E-05 |
| ENSAPLG00000025088 | 1.819655 | 5.41E-06 | 6.05E-05 |
| MMP23B | 0.89776 | 5.44E-06 | 6.08E-05 |
| LONRF1 | 0.866083 | 5.47E-06 | 6.10E-05 |
| TUBD1 | 1.272367 | 5.47E-06 | 6.10E-05 |
| EDEM2 | -0.83719 | 5.51E-06 | 6.14E-05 |
| C7orf26 | -0.84713 | 5.67E-06 | 6.31E-05 |
| CHRNA4 | 6.589601 | 5.68E-06 | 6.31E-05 |
| MICU1 | -1.1517 | 5.73E-06 | 6.36E-05 |
| JMJD7 | -1.61306 | 5.73E-06 | 6.36E-05 |
| SEH1L | -0.89633 | 5.75E-06 | 6.37E-05 |
| POLL | 1.019082 | 5.85E-06 | 6.47E-05 |
| SLC9A4 | 6.579105 | 5.90E-06 | 6.53E-05 |
| LRRC45 | -0.90838 | 5.92E-06 | 6.54E-05 |

| | | | |
|---|---|---|---|
| TNRC18 | 1.676193 | 6.03E-06 | 6.65E-05 |
| SULF1 | -1.62501 | 6.09E-06 | 6.70E-05 |
| NDNF | -0.75898 | 6.08E-06 | 6.70E-05 |
| ENSAPLG00000008403 | -1.57555 | 6.11E-06 | 6.72E-05 |
| HGSNAT | -0.8909 | 6.16E-06 | 6.77E-05 |
| FBXO41 | 4.894795 | 6.17E-06 | 6.78E-05 |
| CPLX4 | 6.597564 | 6.18E-06 | 6.78E-05 |
| ASPM | 3.033274 | 6.24E-06 | 6.84E-05 |
| FZD7 | 0.951057 | 6.26E-06 | 6.85E-05 |
| HELZ2 | 4.201798 | 6.26E-06 | 6.85E-05 |
| ENSAPLG00000003769 | -1.68233 | 6.33E-06 | 6.91E-05 |
| PGD | -0.88996 | 6.33E-06 | 6.91E-05 |
| RPA1 | -0.80414 | 6.40E-06 | 6.98E-05 |
| SIRT7 | 1.467996 | 6.41E-06 | 6.99E-05 |
| ENSAPLG00000012647 | 1.18822 | 6.51E-06 | 7.09E-05 |
| AKAP10 | 0.924646 | 6.58E-06 | 7.14E-05 |
| ZMPSTE24 | -0.76303 | 6.57E-06 | 7.14E-05 |
| DUSP1 | 1.128211 | 6.60E-06 | 7.16E-05 |
| ARMC6 | -0.743 | 6.64E-06 | 7.20E-05 |
| ENSAPLG00000030318 | 1.394812 | 6.65E-06 | 7.20E-05 |
| XKR6 | 1.365476 | 6.93E-06 | 7.49E-05 |
| CHST10 | 0.855498 | 6.98E-06 | 7.55E-05 |
| RTCB | -0.72016 | 7.02E-06 | 7.58E-05 |
| TALDO1 | -0.90371 | 7.04E-06 | 7.58E-05 |
| XRCC5 | -0.81691 | 7.04E-06 | 7.58E-05 |
| YARS1 | -0.7669 | 7.05E-06 | 7.59E-05 |
| ENSAPLG00000006258 | -0.87288 | 7.06E-06 | 7.59E-05 |
| TMEM138 | 1.163496 | 7.07E-06 | 7.60E-05 |
| OTOGL | 2.58034 | 7.14E-06 | 7.67E-05 |
| TMEM171 | -2.1344 | 7.23E-06 | 7.74E-05 |
| TNNT3 | 2.849326 | 7.23E-06 | 7.74E-05 |
| CCN5 | -2.50618 | 7.23E-06 | 7.74E-05 |
| ZNF469 | 1.306711 | 7.34E-06 | 7.85E-05 |
| ENSAPLG00000005283 | -1.07946 | 7.36E-06 | 7.86E-05 |
| ENSAPLG00000011759 | 3.708942 | 7.45E-06 | 7.95E-05 |
| ATXN1 | 1.399667 | 7.65E-06 | 8.16E-05 |
| EIF1 | 1.072811 | 7.67E-06 | 8.17E-05 |
| SLC38A9 | -1.16502 | 7.70E-06 | 8.19E-05 |
| RPL4 | -0.88503 | 7.86E-06 | 8.35E-05 |
| ENSAPLG00000021973 | 2.948388 | 7.92E-06 | 8.41E-05 |
| CMAS | -0.94198 | 8.01E-06 | 8.49E-05 |
| ORC3 | -0.99396 | 8.07E-06 | 8.55E-05 |
| PAIP1 | -0.85869 | 8.08E-06 | 8.56E-05 |
| CLK1 | 0.92732 | 8.15E-06 | 8.62E-05 |
| BYSL | -0.85897 | 8.18E-06 | 8.65E-05 |

| | | | |
|---|---|---|---|
| MTHFR | 0.784292 | 8.33E-06 | 8.79E-05 |
| LAMP1 | -0.78556 | 8.37E-06 | 8.83E-05 |
| RBBP7 | -0.88624 | 8.53E-06 | 8.98E-05 |
| ADD1 | -0.79267 | 8.54E-06 | 8.98E-05 |
| EEF1AKNMT | -0.9865 | 8.55E-06 | 8.99E-05 |
| PAK3 | 1.227012 | 8.61E-06 | 9.04E-05 |
| SLC13A1 | 6.581967 | 8.63E-06 | 9.06E-05 |
| PFKFB3 | 1.303768 | 8.66E-06 | 9.07E-05 |
| NR1H3 | 0.768538 | 8.66E-06 | 9.07E-05 |
| RECQL | -0.86762 | 8.68E-06 | 9.08E-05 |
| ATXN7L1 | 1.062263 | 8.79E-06 | 9.19E-05 |
| VASH1 | 1.045922 | 8.84E-06 | 9.23E-05 |
| TCF24 | 2.086152 | 8.84E-06 | 9.23E-05 |
| ABI1 | 0.796516 | 8.90E-06 | 9.28E-05 |
| ATF4 | 0.721636 | 8.95E-06 | 9.32E-05 |
| G3BP1 | -0.68791 | 8.98E-06 | 9.34E-05 |
| CORO1C | -0.83824 | 9.00E-06 | 9.35E-05 |
| SCUBE1 | -1.9828 | 9.02E-06 | 9.37E-05 |
| ENSAPLG00000007649 | -0.98452 | 9.24E-06 | 9.57E-05 |
| ENSAPLG00000009836 | 0.952874 | 9.23E-06 | 9.57E-05 |
| RPS24 | 0.876412 | 9.24E-06 | 9.57E-05 |
| TRIM66 | 2.946288 | 9.27E-06 | 9.59E-05 |
| ENSAPLG00000003990 | -1.40752 | 9.29E-06 | 9.60E-05 |
| MRPL39 | -0.90423 | 9.35E-06 | 9.65E-05 |
| TASOR | 0.875888 | 9.41E-06 | 9.71E-05 |
| ENSAPLG00000029441 | 1.187227 | 9.67E-06 | 9.96E-05 |
| LZTS3 | 1.778795 | 1.03E-05 | 0.000106 |
| ZFAND2A | 0.995378 | 1.05E-05 | 0.000108 |
| MRPS35 | -0.91837 | 1.05E-05 | 0.000108 |
| ADH5 | -0.78809 | 1.06E-05 | 0.000109 |
| NGF | -1.58192 | 1.06E-05 | 0.000109 |
| GREM1 | 5.388891 | 1.07E-05 | 0.000109 |
| WDR12 | -0.74919 | 1.07E-05 | 0.00011 |
| RFK | 0.930376 | 1.08E-05 | 0.000111 |
| ELK4 | 1.072804 | 1.09E-05 | 0.000111 |
| LIPG | -1.01457 | 1.09E-05 | 0.000111 |
| ENSAPLG00000007323 | 0.706082 | 1.09E-05 | 0.000112 |
| SH3BP4 | -1.07478 | 1.10E-05 | 0.000112 |
| DYNC1I2 | -0.79666 | 1.11E-05 | 0.000113 |
| NPPB | 1.104508 | 1.11E-05 | 0.000113 |
| LRRC59 | -0.78997 | 1.12E-05 | 0.000114 |
| ENSAPLG00000016403 | -0.81781 | 1.14E-05 | 0.000115 |
| LRRC38 | 1.743237 | 1.14E-05 | 0.000116 |
| ENSAPLG00000008997 | 2.301485 | 1.15E-05 | 0.000117 |
| ENSAPLG00000015422 | 2.835705 | 1.16E-05 | 0.000117 |

| | | | |
|---|---|---|---|
| RNH1 | -0.71031 | 1.16E-05 | 0.000118 |
| DENND2C | 0.912207 | 1.17E-05 | 0.000118 |
| TUBGCP2 | -0.90195 | 1.17E-05 | 0.000118 |
| ENSAPLG00000021975 | 0.933831 | 1.17E-05 | 0.000118 |
| SLC22A3 | -1.93654 | 1.19E-05 | 0.00012 |
| SLC2A10 | -1.01566 | 1.20E-05 | 0.00012 |
| GPC4 | -0.80787 | 1.21E-05 | 0.000121 |
| ENSAPLG00000004985 | -1.07481 | 1.21E-05 | 0.000122 |
| SLC29A4 | 2.015866 | 1.24E-05 | 0.000125 |
| NCSTN | -0.80925 | 1.24E-05 | 0.000125 |
| ALDH1L2 | -1.86192 | 1.25E-05 | 0.000126 |
| ACTA2 | -1.04069 | 1.26E-05 | 0.000126 |
| JARID2 | 1.017572 | 1.26E-05 | 0.000127 |
| GTPBP8 | -1.04706 | 1.27E-05 | 0.000127 |
| FGF4 | 6.52137 | 1.27E-05 | 0.000127 |
| IKBKB | -0.80346 | 1.27E-05 | 0.000127 |
| ACAD11 | -0.97678 | 1.28E-05 | 0.000128 |
| RNF19B | 1.184754 | 1.29E-05 | 0.000128 |
| CRHR2 | -0.81636 | 1.30E-05 | 0.000129 |
| SPAST | -0.69227 | 1.30E-05 | 0.00013 |
| ENSAPLG00000001044 | 1.966022 | 1.31E-05 | 0.00013 |
| TTC28 | 0.994711 | 1.31E-05 | 0.00013 |
| PIK3R5 | 1.436449 | 1.32E-05 | 0.000131 |
| ZFHX3 | 1.390728 | 1.32E-05 | 0.000131 |
| EPB41L2 | -1.14467 | 1.32E-05 | 0.000131 |
| EVC | -1.27628 | 1.33E-05 | 0.000132 |
| PTGES2 | -0.88326 | 1.35E-05 | 0.000134 |
| TPCN1 | 0.986727 | 1.37E-05 | 0.000135 |
| FBXL14 | 1.396502 | 1.37E-05 | 0.000135 |
| GCK | 1.114289 | 1.37E-05 | 0.000135 |
| ENSAPLG00000026953 | 4.293122 | 1.39E-05 | 0.000136 |
| SETD7 | 0.867311 | 1.40E-05 | 0.000137 |
| SDC1 | -0.72005 | 1.40E-05 | 0.000138 |
| ENSAPLG00000009313 | -0.82296 | 1.40E-05 | 0.000138 |
| ELFN1 | 0.882157 | 1.41E-05 | 0.000138 |
| ARID5B | 0.916635 | 1.41E-05 | 0.000138 |
| BRSK2 | 2.062967 | 1.41E-05 | 0.000138 |
| SETBP1 | 1.257559 | 1.43E-05 | 0.000139 |
| PKD2 | -1.12119 | 1.43E-05 | 0.00014 |
| NMT1 | -0.73744 | 1.45E-05 | 0.000141 |
| CCNB2 | -1.05567 | 1.48E-05 | 0.000144 |
| PPP2R3C | -0.83134 | 1.48E-05 | 0.000144 |
| ENSAPLG00000016555 | -0.7382 | 1.52E-05 | 0.000148 |
| ENSAPLG00000014266 | -0.84214 | 1.53E-05 | 0.000149 |
| ENSAPLG00000016418 | 6.470635 | 1.53E-05 | 0.000149 |

| | | | |
|---|---|---|---|
| ECM2 | -0.9592 | 1.54E-05 | 0.00015 |
| MMGT1 | 0.716323 | 1.55E-05 | 0.00015 |
| SDCBP2 | 4.736122 | 1.55E-05 | 0.00015 |
| CDK7 | -0.88996 | 1.56E-05 | 0.000151 |
| TNR | 5.2706 | 1.56E-05 | 0.000151 |
| ENSAPLG00000004897 | -1.44135 | 1.57E-05 | 0.000152 |
| ATF7IP | 0.868226 | 1.60E-05 | 0.000155 |
| TP53INP2 | 2.875474 | 1.60E-05 | 0.000155 |
| FNDC3A | 0.863428 | 1.61E-05 | 0.000155 |
| PSMA4 | -0.83152 | 1.61E-05 | 0.000156 |
| SEC23B | -0.70119 | 1.62E-05 | 0.000156 |
| ENSAPLG00000003696 | 6.467645 | 1.62E-05 | 0.000156 |
| UQCRC1 | -0.74799 | 1.63E-05 | 0.000157 |
| MTCH2 | -0.83947 | 1.64E-05 | 0.000157 |
| SAG | 6.195746 | 1.65E-05 | 0.000158 |
| NPY | -1.48082 | 1.66E-05 | 0.000159 |
| PARP3 | -1.50436 | 1.66E-05 | 0.000159 |
| COPB2 | -0.6889 | 1.68E-05 | 0.000161 |
| UTP18 | -0.77309 | 1.69E-05 | 0.000161 |
| CEP85L | 0.947977 | 1.68E-05 | 0.000161 |
| PURG | 1.856172 | 1.70E-05 | 0.000163 |
| USP5 | -0.86175 | 1.71E-05 | 0.000163 |
| FAM43A | 1.370385 | 1.71E-05 | 0.000163 |
| BCLAF3 | 0.768904 | 1.75E-05 | 0.000166 |
| PIK3IP1 | -0.81753 | 1.76E-05 | 0.000167 |
| HECW2 | 0.968263 | 1.76E-05 | 0.000167 |
| FUCA2 | -0.78982 | 1.77E-05 | 0.000168 |
| BUB3 | -0.78076 | 1.79E-05 | 0.00017 |
| LMAN1 | -0.77215 | 1.80E-05 | 0.000171 |
| C1GALT1 | 0.94407 | 1.83E-05 | 0.000173 |
| ZCCHC2 | 0.951401 | 1.87E-05 | 0.000177 |
| MAP3K13 | 1.07426 | 1.87E-05 | 0.000177 |
| SCFD1 | -0.76555 | 1.87E-05 | 0.000177 |
| SEC63 | -0.66831 | 1.89E-05 | 0.000179 |
| PPP2R2D | 0.678205 | 1.91E-05 | 0.00018 |
| CEMIP | 1.097054 | 1.92E-05 | 0.00018 |
| ARL14 | -2.10642 | 1.92E-05 | 0.000181 |
| ENSAPLG00000026807 | 1.125234 | 1.93E-05 | 0.000181 |
| UNG | -1.37655 | 1.93E-05 | 0.000182 |
| NOTUM | -1.01833 | 1.94E-05 | 0.000182 |
| ENSAPLG00000003866 | 1.669749 | 1.96E-05 | 0.000184 |
| CIT | 1.974614 | 1.96E-05 | 0.000184 |
| PROM2 | 4.2323 | 1.96E-05 | 0.000184 |
| DIPK2A | -1.23557 | 1.99E-05 | 0.000186 |
| PEX16 | -0.81492 | 2.00E-05 | 0.000186 |

| | | | |
|---|---|---|---|
| HS3ST3A1 | 1.807366 | 2.00E-05 | 0.000186 |
| FAM180B | 4.249611 | 2.03E-05 | 0.000189 |
| PITRM1 | -0.81534 | 2.04E-05 | 0.00019 |
| ENSAPLG00000030024 | -1.08551 | 2.06E-05 | 0.000192 |
| ENSAPLG00000006561 | 0.846006 | 2.07E-05 | 0.000192 |
| SLC5A1 | 2.578003 | 2.07E-05 | 0.000192 |
| ENSAPLG00000020029 | 1.110493 | 2.07E-05 | 0.000192 |
| SUMF1 | -0.86518 | 2.10E-05 | 0.000195 |
| HSDL1 | -0.84594 | 2.10E-05 | 0.000195 |
| AGAP1 | 0.91127 | 2.13E-05 | 0.000197 |
| PLK4 | 1.101782 | 2.13E-05 | 0.000197 |
| PLPP5 | -1.12675 | 2.13E-05 | 0.000197 |
| PANK3 | 0.78732 | 2.14E-05 | 0.000197 |
| ST3GAL3 | 0.891474 | 2.14E-05 | 0.000197 |
| ENSAPLG00000018551 | 0.811608 | 2.14E-05 | 0.000197 |
| ENSAPLG00000028904 | 1.191985 | 2.14E-05 | 0.000197 |
| ENSAPLG00000008352 | 1.756698 | 2.15E-05 | 0.000198 |
| CDC20 | -1.31463 | 2.17E-05 | 0.000199 |
| ABCE1 | -0.75481 | 2.19E-05 | 0.000201 |
| FAM114A1 | -0.79676 | 2.19E-05 | 0.000201 |
| KLHL40 | 0.913278 | 2.21E-05 | 0.000202 |
| SEMA3G | 1.096422 | 2.22E-05 | 0.000203 |
| PITX1 | 1.011481 | 2.22E-05 | 0.000203 |
| INPP5A | 0.818995 | 2.23E-05 | 0.000204 |
| MTHFD1 | -0.88747 | 2.24E-05 | 0.000204 |
| CAND1 | -0.87194 | 2.23E-05 | 0.000204 |
| ENSAPLG00000028020 | 0.970777 | 2.24E-05 | 0.000205 |
| PARN | -0.72575 | 2.28E-05 | 0.000208 |
| PRMT3 | -0.73318 | 2.29E-05 | 0.000208 |
| DUSP8 | 1.274387 | 2.31E-05 | 0.00021 |
| VSTM4 | -1.27876 | 2.31E-05 | 0.00021 |
| ENSAPLG00000008185 | -1.08571 | 2.33E-05 | 0.000212 |
| TUBGCP3 | -0.97226 | 2.33E-05 | 0.000212 |
| GLOD4 | -0.84157 | 2.39E-05 | 0.000217 |
| ATL1 | -0.71112 | 2.44E-05 | 0.000221 |
| ITPRID2 | 0.931117 | 2.44E-05 | 0.000221 |
| ZBTB18 | 0.934758 | 2.45E-05 | 0.000222 |
| RAB11FIP3 | 1.154311 | 2.48E-05 | 0.000224 |
| PTGES3 | 0.752059 | 2.48E-05 | 0.000224 |
| SH3PXD2A | 0.904303 | 2.51E-05 | 0.000226 |
| AKR1D1 | -1.34047 | 2.52E-05 | 0.000227 |
| MCM3 | -2.0165 | 2.51E-05 | 0.000227 |
| OAT | -0.71775 | 2.51E-05 | 0.000227 |
| FHOD1 | 1.003975 | 2.52E-05 | 0.000227 |
| XPNPEP1 | -0.73584 | 2.54E-05 | 0.000228 |

| | | | |
|---|---|---|---|
| RBPMS | 0.943934 | 2.56E-05 | 0.00023 |
| PLSCR1 | -0.93698 | 2.56E-05 | 0.00023 |
| RABL6 | 0.874783 | 2.60E-05 | 0.000233 |
| NSMCE1 | -1.00367 | 2.62E-05 | 0.000235 |
| TMEM41A | -1.0483 | 2.63E-05 | 0.000235 |
| CCT7 | -0.72811 | 2.64E-05 | 0.000237 |
| TLR6 | -1.21173 | 2.66E-05 | 0.000238 |
| PLA2G4A | -0.73789 | 2.66E-05 | 0.000238 |
| ORAI1 | -0.86332 | 2.67E-05 | 0.000238 |
| ALG5 | -0.77947 | 2.67E-05 | 0.000238 |
| HKDC1 | 4.573737 | 2.67E-05 | 0.000238 |
| ANGEL1 | 0.889539 | 2.70E-05 | 0.000241 |
| KCNC2 | 0.753165 | 2.72E-05 | 0.000242 |
| DDR2 | -0.68398 | 2.72E-05 | 0.000242 |
| VPS36 | -0.94636 | 2.73E-05 | 0.000242 |
| BACE1 | 0.747899 | 2.74E-05 | 0.000243 |
| MON1A | 1.00794 | 2.75E-05 | 0.000244 |
| FKBP10 | -0.72526 | 2.75E-05 | 0.000244 |
| JADE2 | 1.565753 | 2.76E-05 | 0.000245 |
| MB | 3.707927 | 2.77E-05 | 0.000245 |
| RASD2 | 5.197599 | 2.77E-05 | 0.000245 |
| ERCC4 | -0.92594 | 2.77E-05 | 0.000245 |
| LIG3 | -0.86168 | 2.78E-05 | 0.000246 |
| RTN4 | -0.73144 | 2.82E-05 | 0.000249 |
| SUCLA2 | -0.72302 | 2.83E-05 | 0.000249 |
| FASTKD1 | -1.67621 | 2.86E-05 | 0.000252 |
| ENSAPLG00000012949 | -1.41228 | 2.87E-05 | 0.000252 |
| ADAMTSL3 | -0.67323 | 2.88E-05 | 0.000253 |
| ERBB4 | 1.249799 | 2.89E-05 | 0.000254 |
| MEMO1 | 0.774508 | 2.91E-05 | 0.000255 |
| SETD5 | 0.959012 | 2.95E-05 | 0.000259 |
| ENSAPLG00000004084 | -0.84412 | 2.96E-05 | 0.000259 |
| MANEA | 0.732985 | 2.96E-05 | 0.000259 |
| TXNDC5 | -0.78447 | 2.96E-05 | 0.000259 |
| AKTIP | -0.72801 | 2.96E-05 | 0.000259 |
| HIPK1 | 1.081345 | 2.97E-05 | 0.000259 |
| ENSAPLG00000001060 | 3.735088 | 2.98E-05 | 0.00026 |
| WBP2NL | 0.784287 | 3.02E-05 | 0.000263 |
| TYW5 | -0.98952 | 3.03E-05 | 0.000264 |
| ENSAPLG00000016881 | 0.935459 | 3.05E-05 | 0.000266 |
| HACD1 | 0.829322 | 3.08E-05 | 0.000268 |
| SRPX | -1.09973 | 3.08E-05 | 0.000268 |
| TSHZ1 | 1.17978 | 3.14E-05 | 0.000273 |
| MED27 | -0.77625 | 3.14E-05 | 0.000273 |
| PNLDC1 | 3.706432 | 3.19E-05 | 0.000277 |

| | | | |
|---|---|---|---|
| NUP88 | -0.83444 | 3.23E-05 | 0.00028 |
| ZCCHC10 | 0.776331 | 3.24E-05 | 0.000281 |
| ATRN | -1.01782 | 3.25E-05 | 0.000281 |
| ENSAPLG00000008933 | -0.8207 | 3.25E-05 | 0.000281 |
| ENSAPLG00000023864 | 1.709042 | 3.28E-05 | 0.000283 |
| SSR2 | -0.74894 | 3.28E-05 | 0.000283 |
| SREK1 | -0.95414 | 3.30E-05 | 0.000284 |
| SETD3 | -0.74805 | 3.31E-05 | 0.000284 |
| KAT6A | 0.835009 | 3.31E-05 | 0.000284 |
| EXT2 | -0.89297 | 3.30E-05 | 0.000284 |
| KIAA0895 | 0.802932 | 3.30E-05 | 0.000284 |
| GUCY1B1 | -0.93417 | 3.36E-05 | 0.000289 |
| ENSAPLG00000011675 | 1.076716 | 3.37E-05 | 0.000289 |
| FAM76A | 0.868888 | 3.39E-05 | 0.000291 |
| CCNE1 | -0.84479 | 3.42E-05 | 0.000293 |
| POP4 | -1.02932 | 3.46E-05 | 0.000297 |
| ENSAPLG00000018678 | 1.013715 | 3.52E-05 | 0.000301 |
| IARS2 | -0.77377 | 3.53E-05 | 0.000302 |
| ENSAPLG00000002847 | -0.69116 | 3.54E-05 | 0.000302 |
| CKB | -0.86642 | 3.58E-05 | 0.000306 |
| OVOL2 | 1.949735 | 3.58E-05 | 0.000306 |
| TENT5A | 0.680862 | 3.59E-05 | 0.000306 |
| PTCHD1 | 0.918558 | 3.66E-05 | 0.000311 |
| PPT1 | -0.90764 | 3.67E-05 | 0.000313 |
| QDPR | -0.82109 | 3.68E-05 | 0.000313 |
| MYLIP | 0.748758 | 3.68E-05 | 0.000313 |
| ZNF644 | 0.65013 | 3.74E-05 | 0.000317 |
| ENSAPLG00000024538 | 1.049442 | 3.75E-05 | 0.000318 |
| SMPD1 | -0.75335 | 3.79E-05 | 0.000321 |
| ACSS3 | -1.54769 | 3.80E-05 | 0.000322 |
| SH3RF3 | 0.98598 | 3.82E-05 | 0.000323 |
| RPP30 | -0.76658 | 3.83E-05 | 0.000324 |
| MYO5A | 0.903017 | 3.85E-05 | 0.000325 |
| ENSAPLG00000000857 | -1.7198 | 3.85E-05 | 0.000325 |
| ARHGAP21 | 0.926433 | 3.87E-05 | 0.000326 |
| CALCB | -1.28092 | 3.89E-05 | 0.000328 |
| SLC39A6 | -0.74883 | 3.89E-05 | 0.000328 |
| CAMSAP2 | 0.878169 | 3.91E-05 | 0.000329 |
| MYOCD | 1.119789 | 3.91E-05 | 0.000329 |
| ENSAPLG00000016352 | -0.80456 | 3.92E-05 | 0.000329 |
| MMD | 0.807483 | 3.95E-05 | 0.000332 |
| TBL1XR1 | 0.650969 | 3.99E-05 | 0.000335 |
| MAP7 | 0.738061 | 4.00E-05 | 0.000336 |
| PRKAG1 | 1.551573 | 4.02E-05 | 0.000337 |
| DNAJC17 | -0.92443 | 4.04E-05 | 0.000338 |

| | | | |
|---|---|---|---|
| HK1 | -0.71955 | 4.04E-05 | 0.000338 |
| CLDN20 | 6.228054 | 4.05E-05 | 0.000338 |
| RNF11 | 0.704347 | 4.07E-05 | 0.00034 |
| RPN1 | -0.66844 | 4.11E-05 | 0.000343 |
| NOX4 | 0.763884 | 4.17E-05 | 0.000348 |
| BCAT1 | -0.78405 | 4.18E-05 | 0.000348 |
| NCOA7 | 0.813852 | 4.24E-05 | 0.000353 |
| ENSAPLG00000018956 | 4.641761 | 4.28E-05 | 0.000356 |
| ENSAPLG00000009553 | 1.233603 | 4.30E-05 | 0.000358 |
| GPR27 | 0.837495 | 4.31E-05 | 0.000358 |
| PLEKHB2 | 0.799132 | 4.32E-05 | 0.000359 |
| RPL22L1 | 1.025903 | 4.34E-05 | 0.000359 |
| ENSAPLG00000021234 | -1.57031 | 4.33E-05 | 0.000359 |
| HMGB2 | -1.21795 | 4.36E-05 | 0.000361 |
| RGP1 | -1.17062 | 4.36E-05 | 0.000361 |
| TCF12 | 0.681159 | 4.38E-05 | 0.000362 |
| DCN | -0.84925 | 4.41E-05 | 0.000364 |
| NKX2-5 | 6.203565 | 4.44E-05 | 0.000367 |
| ALG8 | -0.85638 | 4.50E-05 | 0.000371 |
| EEF2 | -0.64334 | 4.52E-05 | 0.000373 |
| ABCA4 | 2.497476 | 4.53E-05 | 0.000373 |
| ENSAPLG00000020680 | 0.73261 | 4.59E-05 | 0.000378 |
| LRTM2 | 5.788753 | 4.66E-05 | 0.000383 |
| ENSAPLG00000020417 | -0.83399 | 4.66E-05 | 0.000383 |
| ENSAPLG00000010727 | -0.97028 | 4.70E-05 | 0.000386 |
| ATIC | -0.71359 | 4.74E-05 | 0.000389 |
| GCNT4 | 1.148779 | 4.76E-05 | 0.00039 |
| NUP133 | -0.91122 | 4.77E-05 | 0.000391 |
| MVP | -0.87656 | 4.83E-05 | 0.000396 |
| AGO4 | 0.92842 | 4.90E-05 | 0.000401 |
| ZNF236 | 0.79996 | 5.05E-05 | 0.000413 |
| PRKAA1 | -0.8709 | 5.08E-05 | 0.000415 |
| ENSAPLG00000016876 | 4.490849 | 5.11E-05 | 0.000417 |
| BAK1 | 0.744213 | 5.12E-05 | 0.000417 |
| BLOC1S6 | 0.724247 | 5.14E-05 | 0.000419 |
| CLPX | -0.75143 | 5.23E-05 | 0.000426 |
| TSPAN13 | 2.894119 | 5.26E-05 | 0.000429 |
| CPSF7 | 1.473949 | 5.28E-05 | 0.00043 |
| ENSAPLG00000011141 | 1.277955 | 5.33E-05 | 0.000433 |
| LDLRAP1 | -0.81309 | 5.34E-05 | 0.000434 |
| ENSAPLG00000029258 | 2.21107 | 5.35E-05 | 0.000435 |
| FHL2 | -1.05813 | 5.38E-05 | 0.000436 |
| QTRT2 | -0.75097 | 5.41E-05 | 0.000438 |
| CBFA2T3 | 1.907867 | 5.41E-05 | 0.000438 |
| CUBN | 0.900593 | 5.43E-05 | 0.000439 |

| | | | |
|---|---|---|---|
| ENSAPLG00000016202 | -0.88863 | 5.43E-05 | 0.000439 |
| EMILIN2 | -0.60646 | 5.48E-05 | 0.000443 |
| NUDT9 | -1.01798 | 5.49E-05 | 0.000443 |
| ENSAPLG00000001882 | -0.91028 | 5.49E-05 | 0.000443 |
| LATS2 | 0.928917 | 5.57E-05 | 0.000449 |
| ACADS | -0.90548 | 5.58E-05 | 0.00045 |
| RFLNA | 6.188028 | 5.60E-05 | 0.000451 |
| SLC5A7 | 0.964501 | 5.62E-05 | 0.000452 |
| EIF2B1 | -0.77259 | 5.66E-05 | 0.000455 |
| BZW1 | -0.68175 | 5.72E-05 | 0.000459 |
| PSMD5 | -0.68717 | 5.73E-05 | 0.00046 |
| CD47 | -1.00696 | 5.74E-05 | 0.00046 |
| ENSAPLG00000008466 | -0.80504 | 5.74E-05 | 0.00046 |
| RMND5A | -0.69123 | 5.79E-05 | 0.000463 |
| NR0B2 | 2.010454 | 5.83E-05 | 0.000466 |
| UBE2O | 1.255954 | 5.86E-05 | 0.000468 |
| IMPA2 | 1.01323 | 5.86E-05 | 0.000468 |
| YOD1 | 0.946836 | 5.92E-05 | 0.000473 |
| LEKR1 | 2.17647 | 5.96E-05 | 0.000476 |
| ENSAPLG00000023154 | 1.034718 | 5.97E-05 | 0.000476 |
| SAT1 | 0.83529 | 5.98E-05 | 0.000476 |
| EFCAB7 | -1.7187 | 5.99E-05 | 0.000477 |
| SLC10A4 | -0.81807 | 5.99E-05 | 0.000477 |
| ENSAPLG00000020378 | 0.686342 | 6.01E-05 | 0.000478 |
| ENSAPLG00000003903 | -0.68156 | 6.03E-05 | 0.000479 |
| ANO10 | -0.9326 | 6.06E-05 | 0.000481 |
| NELL1 | -0.88721 | 6.06E-05 | 0.000481 |
| ENSAPLG00000005363 | -0.7374 | 6.09E-05 | 0.000483 |
| ENSAPLG00000013069 | -0.69587 | 6.17E-05 | 0.000489 |
| PHF20 | 0.871159 | 6.24E-05 | 0.000493 |
| BAG2 | 0.883913 | 6.23E-05 | 0.000493 |
| APOOL | -0.96832 | 6.23E-05 | 0.000493 |
| ATP1B3 | -0.65667 | 6.27E-05 | 0.000495 |
| IKZF5 | 0.993028 | 6.28E-05 | 0.000496 |
| DCUN1D3 | 0.775802 | 6.33E-05 | 0.0005 |
| LPAR3 | 5.020967 | 6.40E-05 | 0.000504 |
| URM1 | 0.949999 | 6.41E-05 | 0.000505 |
| ACO2 | -0.67737 | 6.44E-05 | 0.000507 |
| NUP35 | -0.81849 | 6.48E-05 | 0.00051 |
| COG7 | -0.74287 | 6.49E-05 | 0.00051 |
| ENSAPLG00000010185 | -0.80563 | 6.50E-05 | 0.000511 |
| NABP1 | -0.66921 | 6.56E-05 | 0.000515 |
| RXFP1 | 6.118434 | 6.57E-05 | 0.000515 |
| LAMP2 | -0.64966 | 6.58E-05 | 0.000515 |
| ENSAPLG00000017902 | 3.190421 | 6.58E-05 | 0.000515 |

| | | | |
|---|---|---|---|
| ENSAPLG00000014346 | -1.02274 | 6.60E-05 | 0.000517 |
| MYO3B | 6.208048 | 6.73E-05 | 0.000527 |
| THAP4 | 0.682481 | 6.79E-05 | 0.00053 |
| CALD1 | -0.88093 | 6.86E-05 | 0.000536 |
| PACS2 | 0.730057 | 6.90E-05 | 0.000538 |
| SLC7A11 | -0.87572 | 6.91E-05 | 0.000539 |
| STK39 | -0.99472 | 6.98E-05 | 0.000544 |
| TPM4 | -1.07867 | 6.99E-05 | 0.000544 |
| TLK2 | 0.85006 | 7.00E-05 | 0.000545 |
| ALAS1 | -0.73766 | 7.04E-05 | 0.000547 |
| INTS11 | -0.64557 | 7.06E-05 | 0.000549 |
| PTCH1 | 1.752605 | 7.11E-05 | 0.000552 |
| ANXA1 | -0.88554 | 7.14E-05 | 0.000554 |
| INTS13 | 0.689881 | 7.26E-05 | 0.000563 |
| SLC16A5 | 6.1034 | 7.27E-05 | 0.000563 |
| MRM1 | -1.08609 | 7.28E-05 | 0.000564 |
| STX18 | -1.47061 | 7.37E-05 | 0.00057 |
| ATP5MF | 0.889665 | 7.39E-05 | 0.000571 |
| PRRG3 | 2.040169 | 7.40E-05 | 0.000571 |
| FABP6 | 1.691897 | 7.42E-05 | 0.000573 |
| TM9SF2 | -0.64128 | 7.45E-05 | 0.000575 |
| ACTA1 | -0.96069 | 7.47E-05 | 0.000576 |
| ACTR6 | 0.669411 | 7.47E-05 | 0.000576 |
| ITFG1 | -0.64003 | 7.49E-05 | 0.000577 |
| RAPGEF2 | 0.757464 | 7.50E-05 | 0.000577 |
| STK11 | 1.123286 | 7.54E-05 | 0.00058 |
| TXNL4B | -1.11107 | 7.59E-05 | 0.000583 |
| CWF19L1 | -0.68617 | 7.62E-05 | 0.000585 |
| ENSAPLG00000004100 | 1.432924 | 7.67E-05 | 0.000587 |
| CCT8 | -0.68145 | 7.66E-05 | 0.000587 |
| ZC3H18 | 0.946356 | 7.66E-05 | 0.000587 |
| ENSAPLG00000028144 | -0.79269 | 7.72E-05 | 0.000591 |
| PXDN | -1.19121 | 7.75E-05 | 0.000593 |
| SMG1 | 0.924881 | 7.76E-05 | 0.000593 |
| SLC49A4 | -0.72268 | 7.79E-05 | 0.000595 |
| WLS | -0.66318 | 7.80E-05 | 0.000595 |
| NCKIPSD | 0.983503 | 7.83E-05 | 0.000597 |
| SOX10 | 6.117251 | 7.83E-05 | 0.000597 |
| ENSAPLG00000010396 | -0.95534 | 7.87E-05 | 0.0006 |
| ENSAPLG00000024194 | -1.48851 | 7.88E-05 | 0.0006 |
| ENSAPLG00000004542 | -1.03067 | 7.89E-05 | 0.0006 |
| ACTR3 | -0.65113 | 7.94E-05 | 0.000604 |
| COQ9 | -0.82196 | 7.97E-05 | 0.000605 |
| GLIS2 | 1.478616 | 8.02E-05 | 0.000608 |
| ST7L | -0.7361 | 8.03E-05 | 0.000609 |

| | | | |
|---|---|---|---|
| ENSAPLG00000021525 | -1.15938 | 8.06E-05 | 0.000611 |
| MOB2 | 0.725067 | 8.08E-05 | 0.000612 |
| VCP | -0.60964 | 8.13E-05 | 0.000615 |
| ENSAPLG00000003190 | 1.068437 | 8.18E-05 | 0.000619 |
| CSDC2 | 2.477717 | 8.34E-05 | 0.00063 |
| SIAH1 | -0.73392 | 8.40E-05 | 0.000635 |
| ENSAPLG00000011920 | 1.061739 | 8.43E-05 | 0.000636 |
| LRRC47 | -0.90699 | 8.45E-05 | 0.000637 |
| ENSAPLG00000027821 | 1.147596 | 8.46E-05 | 0.000638 |
| FRY | 1.56979 | 8.50E-05 | 0.00064 |
| LRFN5 | 1.262183 | 8.50E-05 | 0.00064 |
| MARCHF4 | 2.045574 | 8.54E-05 | 0.000642 |
| TRIM24 | 0.756422 | 8.55E-05 | 0.000643 |
| ADCK1 | -0.92941 | 8.57E-05 | 0.000644 |
| FAAH | -1.87026 | 8.60E-05 | 0.000646 |
| HADHB | -0.6972 | 8.64E-05 | 0.000648 |
| ENSAPLG00000009803 | -1.17432 | 8.68E-05 | 0.000651 |
| GRAMD1C | -0.83429 | 8.70E-05 | 0.000652 |
| CCNH | -0.92909 | 8.74E-05 | 0.000654 |
| ENSAPLG00000007453 | 3.00298 | 8.74E-05 | 0.000654 |
| RANGAP1 | -0.70022 | 8.82E-05 | 0.000659 |
| KIFBP | -0.79376 | 8.90E-05 | 0.000665 |
| ENSAPLG00000008356 | 1.053726 | 8.92E-05 | 0.000666 |
| SLC19A1 | -1.118 | 8.94E-05 | 0.000667 |
| RFLNB | 1.220291 | 9.05E-05 | 0.000675 |
| PCDH8 | 1.064756 | 9.05E-05 | 0.000675 |
| SGCE | -0.90375 | 9.09E-05 | 0.000677 |
| EXPH5 | 1.412939 | 9.10E-05 | 0.000677 |
| HMGXB4 | 0.859441 | 9.21E-05 | 0.000685 |
| HACD2 | -0.82053 | 9.25E-05 | 0.000687 |
| FGF12 | 1.601212 | 9.28E-05 | 0.000689 |
| ASB7 | 0.713273 | 9.42E-05 | 0.000699 |
| ENSAPLG00000020873 | 2.381192 | 9.43E-05 | 0.000699 |
| CYP20A1 | -0.67824 | 9.55E-05 | 0.000707 |
| SLC24A1 | 6.234836 | 9.59E-05 | 0.00071 |
| NUP37 | -0.70784 | 9.66E-05 | 0.000715 |
| EEPD1 | -0.77328 | 9.82E-05 | 0.000727 |
| FAM219A | 1.036643 | 9.87E-05 | 0.000729 |
| ENSAPLG00000008568 | -0.74572 | 0.000101 | 0.000743 |
| PCNA | -0.75076 | 0.000101 | 0.000748 |
| COPS4 | -0.67023 | 0.000102 | 0.000753 |
| KARS1 | -0.66651 | 0.000102 | 0.000753 |
| USP6NL | 0.753335 | 0.000103 | 0.00076 |
| ENSAPLG00000001119 | 2.86115 | 0.000104 | 0.000762 |
| OSGIN2 | -1.18608 | 0.000104 | 0.000765 |

| | | | |
|---|---|---|---|
| LACTB2 | -0.73223 | 0.000104 | 0.000765 |
| GOLPH3 | 0.67358 | 0.000104 | 0.000765 |
| ENSAPLG00000002946 | -0.76029 | 0.000105 | 0.000768 |
| PLBD2 | -0.79584 | 0.000105 | 0.000769 |
| TARDBP | -0.65092 | 0.000105 | 0.000771 |
| ENSAPLG00000009118 | 0.986755 | 0.000105 | 0.000771 |
| STYK1 | 3.088683 | 0.000106 | 0.000774 |
| TES | -0.61022 | 0.000107 | 0.000782 |
| PCCB | -0.93518 | 0.000107 | 0.000785 |
| ENSAPLG00000010383 | -0.98107 | 0.000108 | 0.000787 |
| SPTLC2 | -0.71866 | 0.000108 | 0.000787 |
| NGLY1 | -0.93294 | 0.000108 | 0.000788 |
| DHCR7 | -0.69335 | 0.000108 | 0.000791 |
| ENSAPLG00000002824 | 4.437089 | 0.000109 | 0.000792 |
| VTA1 | -0.72706 | 0.000109 | 0.000796 |
| ENSAPLG00000004058 | -0.81959 | 0.000109 | 0.000796 |
| ASPHD2 | 1.894829 | 0.000109 | 0.000796 |
| STYX | 0.768899 | 0.00011 | 0.000797 |
| TBC1D8B | -1.08437 | 0.00011 | 0.000799 |
| TIPIN | -0.9545 | 0.00011 | 0.0008 |
| GPR153 | 1.481743 | 0.00011 | 0.0008 |
| DYNLRB1 | 0.79256 | 0.00011 | 0.0008 |
| ENSAPLG00000002035 | 1.193933 | 0.000113 | 0.000818 |
| ENSAPLG00000019624 | 1.79198 | 0.000114 | 0.000826 |
| SMAP1 | -0.84082 | 0.000115 | 0.000832 |
| ENSAPLG00000007040 | 0.846316 | 0.000116 | 0.000837 |
| GLIS3 | -1.31841 | 0.000116 | 0.000841 |
| ZMIZ1 | 1.327911 | 0.000116 | 0.000841 |
| COL1A2 | -0.88444 | 0.000118 | 0.000852 |
| ENSAPLG00000017178 | 1.16856 | 0.00012 | 0.000863 |
| NAGA | -1.12783 | 0.00012 | 0.000865 |
| NR1D2 | 0.626113 | 0.00012 | 0.000865 |
| FAM13B | 0.815548 | 0.000121 | 0.00087 |
| ENSAPLG00000030799 | -1.05692 | 0.000121 | 0.000872 |
| ENSAPLG00000008335 | 0.657899 | 0.000122 | 0.000879 |
| BTBD10 | -0.64174 | 0.000123 | 0.000881 |
| MDH2 | -0.74667 | 0.000123 | 0.000886 |
| ENSAPLG00000021858 | 0.927705 | 0.000124 | 0.000893 |
| PKP4 | 0.695781 | 0.000126 | 0.000901 |
| CTNNB1 | -0.66999 | 0.000126 | 0.000901 |
| MSL2 | 1.245399 | 0.000126 | 0.000902 |
| MSH6 | -1.10523 | 0.000126 | 0.000903 |
| PNRC2 | 0.703957 | 0.000127 | 0.000907 |
| KIFAP3 | -0.66181 | 0.000127 | 0.000908 |
| SNX17 | 0.62892 | 0.000128 | 0.000911 |

| | | | |
|---|---|---|---|
| VEZF1 | 0.987571 | 0.000128 | 0.000914 |
| ENSAPLG00000023116 | 1.441064 | 0.000128 | 0.000914 |
| ENSAPLG00000011673 | 0.758339 | 0.000129 | 0.000919 |
| SAMD11 | 1.15535 | 0.000129 | 0.000921 |
| BEND5 | -0.82749 | 0.00013 | 0.000921 |
| SSX2IP | 1.834053 | 0.00013 | 0.000926 |
| KIF5B | -0.61425 | 0.00013 | 0.000926 |
| RECK | -0.78605 | 0.000132 | 0.000934 |
| NCDN | -0.92409 | 0.000132 | 0.000934 |
| LGALS3 | 0.937939 | 0.000132 | 0.000934 |
| SCLY | -1.27729 | 0.000133 | 0.00094 |
| ENSAPLG00000025225 | 0.714662 | 0.000133 | 0.00094 |
| CPE | -1.01444 | 0.000134 | 0.000948 |
| ENSAPLG00000008170 | -1.12215 | 0.000134 | 0.000948 |
| PER3 | 1.082265 | 0.000134 | 0.00095 |
| ERO1A | -0.87364 | 0.000135 | 0.000951 |
| MRAS | 0.993642 | 0.000135 | 0.000951 |
| OPHN1 | 1.297682 | 0.000135 | 0.000952 |
| NISCH | 0.824964 | 0.000136 | 0.000956 |
| CCT4 | -0.67386 | 0.000137 | 0.000967 |
| MBD5 | 0.766569 | 0.000138 | 0.000971 |
| TNFAIP3 | 0.94833 | 0.000139 | 0.000977 |
| TRIB1 | 0.756083 | 0.00014 | 0.000987 |
| ENSAPLG00000006355 | 0.892095 | 0.00014 | 0.000987 |
| ADTRP | 1.643242 | 0.000141 | 0.000991 |
| CREBRF | 0.714496 | 0.000141 | 0.000991 |
| CLTA | -0.92523 | 0.000141 | 0.000991 |
| PYGL | -0.7386 | 0.000141 | 0.000991 |
| KLF6 | 0.66559 | 0.000142 | 0.000996 |
| ENSAPLG00000004341 | 0.856309 | 0.000143 | 0.000997 |
| TAF1B | -1.10248 | 0.000142 | 0.000997 |
| IRAK2 | 0.654787 | 0.000143 | 0.000998 |
| ENSAPLG00000008465 | 4.326798 | 0.000143 | 0.000998 |
| APPBP2 | -0.66878 | 0.000144 | 0.001009 |
| ENSAPLG00000016924 | 1.639997 | 0.000146 | 0.001017 |
| ENSAPLG00000024554 | 1.248147 | 0.000147 | 0.001024 |
| TOB1 | 0.73257 | 0.000147 | 0.001025 |
| ZDHHC6 | 0.631904 | 0.000147 | 0.001025 |
| MTRF1L | -1.02496 | 0.000148 | 0.001027 |
| GTF2IRD1 | 0.711004 | 0.000148 | 0.001029 |
| ENSAPLG00000015887 | -0.84827 | 0.000148 | 0.001029 |
| PTP4A3 | 0.836452 | 0.000148 | 0.001029 |
| ENSAPLG00000026915 | -0.75608 | 0.000149 | 0.001035 |
| FYN | 0.588132 | 0.00015 | 0.001039 |
| TEX11 | 1.26952 | 0.00015 | 0.001041 |

| | | | |
|---|---|---|---|
| ENSAPLG00000018666 | 2.570651 | 0.00015 | 0.001041 |
| RAC2 | 1.148775 | 0.000151 | 0.001045 |
| DHDDS | -0.68559 | 0.000152 | 0.001052 |
| PLPP6 | -1.08994 | 0.000153 | 0.001059 |
| ZBTB20 | 0.95135 | 0.000154 | 0.001063 |
| LLGL1 | 0.732157 | 0.000154 | 0.001063 |
| NLRX1 | 0.867801 | 0.000155 | 0.001069 |
| NRARP | 1.289429 | 0.000155 | 0.00107 |
| AXIN1 | 0.784994 | 0.000158 | 0.001091 |
| SLC7A1 | -1.14423 | 0.000159 | 0.001093 |
| POGLUT2 | -0.73147 | 0.00016 | 0.001102 |
| RORB | 0.72721 | 0.00016 | 0.001102 |
| ENSAPLG00000002445 | -0.84624 | 0.00016 | 0.001102 |
| AJM1 | 2.439989 | 0.00016 | 0.001102 |
| RPL36AL | 0.790624 | 0.000161 | 0.001104 |
| EXOC6 | -0.76708 | 0.000161 | 0.001106 |
| BTD | -1.19093 | 0.000162 | 0.001109 |
| RAP2C | 0.792354 | 0.000162 | 0.001109 |
| SNAP29 | -0.77869 | 0.000162 | 0.001109 |
| VPS45 | -0.93372 | 0.000162 | 0.001109 |
| ID2 | 0.838346 | 0.000163 | 0.001112 |
| ENSAPLG00000028568 | 1.123328 | 0.000164 | 0.001124 |
| SMU1 | -0.62941 | 0.000165 | 0.001126 |
| SEMA3C | -0.66806 | 0.000165 | 0.001127 |
| DHTKD1 | -0.88648 | 0.000165 | 0.001127 |
| ALDH7A1 | -0.67944 | 0.000165 | 0.001127 |
| EIF5 | 0.69004 | 0.000165 | 0.001127 |
| GRK7 | 2.170874 | 0.000166 | 0.001127 |
| TSR1 | -0.70391 | 0.000166 | 0.001127 |
| LHFPL2 | 0.625059 | 0.000166 | 0.001127 |
| ENSAPLG00000004266 | -0.95896 | 0.000171 | 0.00116 |
| CCDC92 | 0.947768 | 0.000171 | 0.00116 |
| ENSAPLG00000023707 | 1.468292 | 0.000171 | 0.001164 |
| ATF1 | 0.68655 | 0.000172 | 0.001168 |
| ARF6 | 0.748995 | 0.000174 | 0.001183 |
| CUL2 | -0.58704 | 0.000175 | 0.00119 |
| BACH1 | 0.793736 | 0.000177 | 0.001197 |
| MTPN | 0.706923 | 0.000177 | 0.0012 |
| FAR1 | -0.75619 | 0.000178 | 0.001205 |
| GCDH | -0.8329 | 0.000178 | 0.001206 |
| MBNL3 | 0.759883 | 0.000178 | 0.001206 |
| MOSMO | 1.001786 | 0.000181 | 0.001219 |
| ANXA7 | -0.80197 | 0.000181 | 0.001219 |
| ENSAPLG00000004823 | 0.695067 | 0.000181 | 0.001219 |
| ENSAPLG00000008315 | 1.194506 | 0.000181 | 0.001219 |

| | | | |
|---|---|---|---|
| ACOX3 | -0.81696 | 0.000181 | 0.001219 |
| ENSAPLG00000014933 | 1.202922 | 0.000181 | 0.001219 |
| SELENOM | 0.789069 | 0.000181 | 0.001219 |
| WDR1 | -0.56931 | 0.000181 | 0.001219 |
| HMG20B | -0.69009 | 0.000182 | 0.001222 |
| FGF8 | 4.913751 | 0.000182 | 0.001222 |
| ANAPC13 | 1.083206 | 0.000182 | 0.001224 |
| NAA35 | -0.66309 | 0.000182 | 0.001224 |
| STARD13 | 0.870338 | 0.000183 | 0.001224 |
| RIPOR1 | 0.897378 | 0.000183 | 0.001226 |
| HPD | 1.102502 | 0.000184 | 0.001229 |
| ZNF704 | 0.776757 | 0.000184 | 0.00123 |
| TMEM39A | 0.649358 | 0.000187 | 0.001249 |
| ENSAPLG00000014620 | -1.0979 | 0.000187 | 0.001249 |
| TADA2A | -0.91163 | 0.000189 | 0.001261 |
| PNPLA8 | -0.73765 | 0.000189 | 0.001261 |
| LRRC40 | -0.87231 | 0.00019 | 0.001269 |
| ACSL1 | -0.71941 | 0.00019 | 0.001269 |
| ENSAPLG00000003176 | 0.886482 | 0.000191 | 0.001273 |
| MACROH2A1 | -0.72353 | 0.000191 | 0.001274 |
| ENSAPLG00000029768 | 1.177387 | 0.000191 | 0.001274 |
| GTF3C4 | -1.03363 | 0.000192 | 0.001275 |
| ADSL | -0.72573 | 0.000192 | 0.001275 |
| LPAR2 | 0.791947 | 0.000192 | 0.001276 |
| ALDH1A3 | -1.35903 | 0.000193 | 0.001284 |
| SNAP47 | 0.797741 | 0.000194 | 0.001285 |
| KALRN | -0.86826 | 0.000194 | 0.001286 |
| HMGCR | -0.64391 | 0.000194 | 0.001287 |
| ENSAPLG00000024805 | 0.926424 | 0.000196 | 0.001297 |
| HINT3 | 0.735654 | 0.000196 | 0.001297 |
| SCO1 | -0.6512 | 0.000196 | 0.001297 |
| ASAH1 | -0.67126 | 0.000197 | 0.0013 |
| SYNPO | 1.22558 | 0.000199 | 0.001312 |
| ENSAPLG00000021254 | 1.117394 | 0.000199 | 0.001316 |
| CLBA1 | 0.874423 | 0.0002 | 0.001319 |
| TMEM214 | -0.6894 | 0.0002 | 0.001319 |
| SLC2A9 | 4.880371 | 0.000201 | 0.001325 |
| RECQL5 | -0.99378 | 0.000202 | 0.001331 |
| ENSAPLG00000004586 | -0.95765 | 0.000202 | 0.001331 |
| ATP2B1 | -0.824 | 0.000203 | 0.001332 |
| ENSAPLG00000013405 | -0.65948 | 0.000204 | 0.00134 |
| HEYL | 1.953255 | 0.000205 | 0.001346 |
| GPT2 | -1.09762 | 0.00021 | 0.001376 |
| ARHGAP29 | -0.73734 | 0.00021 | 0.00138 |
| PPP1R3C | -0.85748 | 0.000211 | 0.001381 |

| | | | |
|---|---|---|---|
| EPRS1 | -0.7021 | 0.000211 | 0.001382 |
| HSPA9 | -0.59888 | 0.000211 | 0.001382 |
| IQUB | 2.5576 | 0.000211 | 0.001382 |
| TFPI2 | -0.8199 | 0.000211 | 0.001382 |
| ESRRG | 0.769441 | 0.000212 | 0.001389 |
| SEC14L1 | 0.649979 | 0.000213 | 0.001395 |
| ENSAPLG00000007797 | -1.05434 | 0.000214 | 0.001395 |
| TK1 | -1.22218 | 0.000215 | 0.001402 |
| TMEM177 | -1.2121 | 0.000215 | 0.001402 |
| DNAJB5 | 1.577215 | 0.000215 | 0.001403 |
| SRSF4 | 0.961916 | 0.000215 | 0.001403 |
| ENSAPLG00000030894 | 1.035306 | 0.000216 | 0.001409 |
| ENSAPLG00000016592 | 0.965793 | 0.000217 | 0.001415 |
| HIVEP2 | 1.00271 | 0.000218 | 0.001418 |
| NIPA2 | 0.611076 | 0.000218 | 0.001418 |
| RPL13 | 0.775831 | 0.000219 | 0.001422 |
| SENP6 | 0.664314 | 0.00022 | 0.001429 |
| ENSAPLG00000008423 | 1.061557 | 0.00022 | 0.001429 |
| CHAMP1 | 1.063741 | 0.000221 | 0.00143 |
| ENSAPLG00000019499 | 0.756851 | 0.000222 | 0.001436 |
| DOK7 | 0.740397 | 0.000223 | 0.001442 |
| TMEM245 | -1.00246 | 0.000223 | 0.001446 |
| ENSAPLG00000000577 | -0.89037 | 0.000224 | 0.001446 |
| DRAM2 | 0.753409 | 0.000225 | 0.001455 |
| ASAP1 | 0.740119 | 0.000225 | 0.001455 |
| TAF5L | 0.745472 | 0.000227 | 0.001466 |
| SLC4A3 | 0.928473 | 0.000228 | 0.001467 |
| SPATA5L1 | -1.04113 | 0.000227 | 0.001467 |
| SIX2 | 1.188198 | 0.000228 | 0.001467 |
| AP3M2 | -0.66045 | 0.000228 | 0.001471 |
| HNF1B | 3.396397 | 0.000229 | 0.001473 |
| NSD2 | 0.653941 | 0.000229 | 0.001476 |
| DPH5 | -0.91202 | 0.00023 | 0.001476 |
| MARK1 | 0.596256 | 0.000231 | 0.001482 |
| ENSAPLG00000028972 | 1.526787 | 0.000232 | 0.001489 |
| CDK5RAP3 | -1.32275 | 0.000233 | 0.001498 |
| RBMX2 | 0.786202 | 0.000234 | 0.0015 |
| P3H2 | -1.39839 | 0.000235 | 0.001507 |
| MRPL37 | -1.01673 | 0.000235 | 0.001509 |
| SUCLG1 | -0.6772 | 0.000236 | 0.00151 |
| GPC1 | -0.70879 | 0.000238 | 0.00152 |
| SLC25A6 | -0.66192 | 0.000238 | 0.001521 |
| SELENOP | 0.663537 | 0.000238 | 0.001522 |
| CMC4 | 2.441849 | 0.00024 | 0.001532 |
| CRYBB2 | 2.121198 | 0.000241 | 0.001538 |

| | | | |
|---|---|---|---|
| CACNA1I | 5.996643 | 0.000241 | 0.001538 |
| JOSD1 | 1.034807 | 0.000242 | 0.001545 |
| ERMARD | -1.10796 | 0.000243 | 0.001546 |
| VTI1B | -0.80869 | 0.000243 | 0.001548 |
| OBI1 | -0.81195 | 0.000244 | 0.001552 |
| ENSAPLG00000003199 | 0.798033 | 0.000245 | 0.001561 |
| ENSAPLG00000009757 | 0.785393 | 0.000247 | 0.001568 |
| BNC2 | 0.985369 | 0.000248 | 0.001579 |
| DNM1L | -0.60871 | 0.00025 | 0.00159 |
| DHCR24 | -1.25857 | 0.000251 | 0.001591 |
| TPT1 | 0.731413 | 0.000253 | 0.001605 |
| ENSAPLG00000018470 | 5.915732 | 0.000254 | 0.001608 |
| TUB | 0.612867 | 0.000254 | 0.00161 |
| SP3 | 0.618502 | 0.000255 | 0.001616 |
| ERLEC1 | -0.66254 | 0.000257 | 0.001628 |
| RNF166 | 1.490832 | 0.000257 | 0.001628 |
| ENSAPLG00000003368 | -1.07913 | 0.000258 | 0.001629 |
| ENSAPLG00000027896 | 1.015229 | 0.000258 | 0.001629 |
| ZBTB1 | 0.576269 | 0.000258 | 0.00163 |
| ENSAPLG00000013205 | 0.80581 | 0.000259 | 0.001633 |
| NECTIN3 | -0.69037 | 0.00026 | 0.001639 |
| BPGM | 0.748335 | 0.000261 | 0.001645 |
| ADK | -0.63875 | 0.000262 | 0.001652 |
| SLC41A2 | 0.792029 | 0.000267 | 0.001679 |
| ENSAPLG00000015531 | -0.78274 | 0.000267 | 0.001681 |
| ENSAPLG00000011770 | -1.05859 | 0.000268 | 0.001682 |
| LAMTOR5 | 0.874785 | 0.000267 | 0.001682 |
| ARHGEF12 | 0.878582 | 0.000268 | 0.001685 |
| SLC7A5 | -0.65766 | 0.00027 | 0.001698 |
| EIF2B2 | -0.79347 | 0.000271 | 0.001703 |
| PPARG | -0.79478 | 0.000274 | 0.001718 |
| TBX22 | 5.821919 | 0.000274 | 0.001718 |
| ELF1 | 0.686289 | 0.000275 | 0.001721 |
| SLC25A3 | -0.59523 | 0.000275 | 0.001721 |
| PRDM2 | 0.704972 | 0.000278 | 0.00174 |
| TULP4 | 1.64775 | 0.00028 | 0.001751 |
| PFKL | -0.72356 | 0.000282 | 0.001763 |
| CTSS | -1.36706 | 0.000284 | 0.001772 |
| HIVEP1 | 0.741824 | 0.000285 | 0.001779 |
| COL6A2 | -0.76959 | 0.000286 | 0.001785 |
| ADAM8 | -0.66573 | 0.000287 | 0.001789 |
| GBE1 | -0.69227 | 0.000288 | 0.001792 |
| ID4 | 0.699439 | 0.000291 | 0.001811 |
| GFOD2 | 1.240554 | 0.000291 | 0.001814 |
| MXI1 | 0.651134 | 0.000293 | 0.001823 |

| | | | |
|---|---|---|---|
| FDPS | -0.83393 | 0.000294 | 0.00183 |
| PRKD3 | 0.891841 | 0.000296 | 0.00184 |
| PBX1 | 0.957037 | 0.000296 | 0.001841 |
| TIMP3 | -1.02685 | 0.000299 | 0.001855 |
| TEKT2 | 3.151486 | 0.000301 | 0.001867 |
| PGRMC2 | -0.66267 | 0.000301 | 0.001867 |
| TMCC2 | 1.411446 | 0.000302 | 0.001869 |
| D2HGDH | -1.38377 | 0.000303 | 0.001874 |
| TRA2B | -0.6741 | 0.000305 | 0.001886 |
| PKHD1 | 1.438269 | 0.000305 | 0.001886 |
| TPBG | -1.08046 | 0.000305 | 0.001886 |
| ENSAPLG00000027187 | -1.20082 | 0.000305 | 0.001886 |
| GNAO1 | 0.918558 | 0.000306 | 0.001887 |
| ACAP2 | -0.7049 | 0.000307 | 0.001896 |
| BMP2 | -0.62787 | 0.000308 | 0.0019 |
| MARCHF2 | 0.741455 | 0.000309 | 0.001906 |
| EEFSEC | -0.9445 | 0.000311 | 0.001917 |
| ENSAPLG00000011112 | 1.621596 | 0.000313 | 0.001926 |
| UTP6 | -0.65848 | 0.000316 | 0.001944 |
| TRMT9B | -0.77588 | 0.000316 | 0.001944 |
| TYR | 5.86467 | 0.000316 | 0.001944 |
| FAM168A | 0.949907 | 0.000321 | 0.001974 |
| RPP40 | -0.82522 | 0.000322 | 0.001976 |
| ENSAPLG00000006069 | 0.611224 | 0.000324 | 0.001988 |
| PSEN1 | -0.76009 | 0.000326 | 0.002002 |
| MBTD1 | 0.71215 | 0.000328 | 0.002011 |
| FKBP9 | -0.66566 | 0.000329 | 0.002016 |
| SEC22B | -0.62453 | 0.000329 | 0.002016 |
| G3BP2 | -0.6244 | 0.00033 | 0.002022 |
| SCYL3 | -0.64577 | 0.000331 | 0.002025 |
| ENSAPLG00000005706 | 0.849375 | 0.000332 | 0.002029 |
| TWSG1 | -0.60459 | 0.000333 | 0.002035 |
| IPO11 | -0.79363 | 0.000333 | 0.002036 |
| ENSAPLG00000003459 | 1.656942 | 0.000334 | 0.002039 |
| HERC4 | -0.6439 | 0.000334 | 0.002039 |
| COL8A1 | -1.01834 | 0.000335 | 0.002044 |
| RERG | 0.679536 | 0.000337 | 0.002051 |
| NR3C2 | 0.872886 | 0.000337 | 0.002051 |
| GRAMD4 | 0.659286 | 0.000337 | 0.002051 |
| SLC6A20 | 5.847416 | 0.000337 | 0.002051 |
| ENSAPLG00000025635 | 1.378926 | 0.000339 | 0.002062 |
| ERO1B | -0.68875 | 0.000341 | 0.00207 |
| BRD8 | -0.77787 | 0.000341 | 0.002072 |
| SLC25A13 | -0.67997 | 0.000342 | 0.002074 |
| TRPV3 | 1.276781 | 0.000346 | 0.002099 |

| | | | |
|---|---|---|---|
| RPS15A | 0.662039 | 0.000346 | 0.002099 |
| FIGNL1 | -1.77539 | 0.000347 | 0.002105 |
| EIF2S1 | -0.63137 | 0.000348 | 0.002108 |
| EIF2A | -0.63757 | 0.000349 | 0.002109 |
| ENSAPLG00000007729 | -1.42734 | 0.000349 | 0.002113 |
| ADGRV1 | 0.950851 | 0.00035 | 0.002114 |
| FAM20C | 0.88112 | 0.000351 | 0.002119 |
| ENSAPLG00000025723 | -0.80386 | 0.000356 | 0.002152 |
| MRPS22 | -0.73948 | 0.000358 | 0.002159 |
| TAF3 | 0.732368 | 0.000358 | 0.002161 |
| ENSAPLG00000002923 | 0.806742 | 0.000365 | 0.002198 |
| METTL22 | 0.795873 | 0.000365 | 0.002198 |
| RARB | 1.423139 | 0.000366 | 0.002207 |
| MICU2 | -0.66822 | 0.000368 | 0.002215 |
| ENSAPLG00000013751 | -0.68686 | 0.000373 | 0.002241 |
| ZDHHC7 | 0.656643 | 0.000373 | 0.002244 |
| ENSAPLG00000026691 | -0.57058 | 0.000374 | 0.002244 |
| TMEM129 | -0.85703 | 0.000374 | 0.002248 |
| STK25 | 0.702387 | 0.000375 | 0.002252 |
| L3HYPDH | -1.13728 | 0.000376 | 0.002255 |
| STX1A | 0.990309 | 0.000376 | 0.002256 |
| MID1IP1 | 0.805593 | 0.000379 | 0.002271 |
| NR4A1 | 1.587461 | 0.000379 | 0.002272 |
| CASK | -0.70687 | 0.000383 | 0.002291 |
| ZYG11A | -0.76714 | 0.000383 | 0.002294 |
| SERHL2 | -0.76429 | 0.000384 | 0.002297 |
| ENSAPLG00000007708 | -1.23393 | 0.000385 | 0.002299 |
| RFNG | -0.81359 | 0.000385 | 0.002299 |
| ENSAPLG00000030527 | 0.688971 | 0.000386 | 0.002301 |
| ENSAPLG00000003869 | 0.59375 | 0.000389 | 0.002319 |
| DYRK3 | 0.831524 | 0.000389 | 0.002321 |
| DAB2 | 0.6691 | 0.00039 | 0.002327 |
| STON2 | 1.161204 | 0.000391 | 0.002329 |
| BRWD3 | 0.674014 | 0.000394 | 0.002344 |
| XYLB | -0.79708 | 0.000397 | 0.002364 |
| PCNP | 0.665288 | 0.000399 | 0.002373 |
| NFATC2 | 0.817418 | 0.000404 | 0.002401 |
| ITGA1 | -0.7319 | 0.000406 | 0.002408 |
| F2RL2 | 0.735732 | 0.000406 | 0.002409 |
| UBE2QL1 | -1.69889 | 0.000406 | 0.002409 |
| ENSAPLG00000009918 | -0.62224 | 0.000409 | 0.002423 |
| GALC | -0.83579 | 0.00041 | 0.002431 |
| RNF141 | 0.676933 | 0.00041 | 0.002431 |
| NAV2 | 1.085833 | 0.000412 | 0.002435 |
| ENSAPLG00000015238 | -0.71932 | 0.000412 | 0.002435 |

| | | | |
|---|---|---|---|
| ENSAPLG00000016521 | 1.611142 | 0.000412 | 0.002435 |
| AFG3L2 | -0.58404 | 0.000415 | 0.002455 |
| TCF3 | 0.586897 | 0.000416 | 0.002459 |
| UQCC1 | -0.61827 | 0.000421 | 0.002485 |
| ENSAPLG00000020305 | 1.029945 | 0.000422 | 0.002489 |
| MANEAL | -1.28918 | 0.000422 | 0.002489 |
| ENSAPLG00000024437 | -0.6871 | 0.000424 | 0.002496 |
| MCCC1 | -1.33092 | 0.000429 | 0.002525 |
| PPP4R2 | 0.667281 | 0.000429 | 0.002529 |
| MEGF11 | 1.990317 | 0.00043 | 0.002529 |
| ENSAPLG00000024581 | 0.636437 | 0.00043 | 0.002529 |
| SLC9A9 | -1.19721 | 0.000432 | 0.002541 |
| TDP1 | -0.87615 | 0.000436 | 0.002562 |
| ENSAPLG00000010480 | 1.042201 | 0.000437 | 0.002565 |
| MKKS | 0.722729 | 0.000437 | 0.002566 |
| TSC22D3 | 1.630527 | 0.000438 | 0.002568 |
| MCM9 | -1.7062 | 0.000439 | 0.00257 |
| TMEM184A | 1.54966 | 0.000439 | 0.00257 |
| TSC22D2 | 0.796116 | 0.000439 | 0.00257 |
| ENSAPLG00000023711 | 0.963852 | 0.000442 | 0.00259 |
| OSCP1 | -0.88091 | 0.000444 | 0.002595 |
| GGH | -0.70997 | 0.000444 | 0.002595 |
| ZBTB47 | 1.578171 | 0.000445 | 0.002602 |
| SPRY4 | 3.806124 | 0.000446 | 0.002604 |
| SMARCAD1 | -0.72555 | 0.000447 | 0.002609 |
| BCL9L | 1.890711 | 0.000447 | 0.002609 |
| SARAF | -0.69851 | 0.00045 | 0.002627 |
| TXLNB | 4.084845 | 0.000451 | 0.002628 |
| MRPL28 | -1.18992 | 0.000451 | 0.002628 |
| ADPGK | -0.74162 | 0.000455 | 0.002647 |
| SGCB | -0.62116 | 0.000456 | 0.002654 |
| KDM2A | 0.755864 | 0.000457 | 0.002659 |
| PRICKLE1 | -0.95285 | 0.000457 | 0.002659 |
| TMEM248 | 0.673746 | 0.00046 | 0.002673 |
| ENSAPLG00000029535 | 1.016275 | 0.000461 | 0.002677 |
| LRRC8C | -0.66388 | 0.000465 | 0.002697 |
| PKIA | 1.057923 | 0.000466 | 0.002704 |
| SEPTIN6 | -0.58727 | 0.000468 | 0.002711 |
| METTL6 | 0.633093 | 0.000469 | 0.002715 |
| ENSAPLG00000004588 | 0.638719 | 0.000469 | 0.002715 |
| KIF4A | 1.348401 | 0.000469 | 0.002715 |
| BTRC | -0.8504 | 0.000471 | 0.002723 |
| POMK | -0.7024 | 0.000471 | 0.002725 |
| CELF2 | -0.72494 | 0.000472 | 0.002725 |
| TUBB1 | 1.165123 | 0.000473 | 0.00273 |

| | | | |
|---|---|---|---|
| TTK | -0.9414 | 0.000475 | 0.002743 |
| ROR2 | -0.99328 | 0.000476 | 0.002744 |
| ZNF827 | 1.945179 | 0.000477 | 0.00275 |
| ENSAPLG00000005608 | 1.082994 | 0.000479 | 0.002761 |
| FAM185A | -0.72328 | 0.000479 | 0.002761 |
| SMPDL3A | -0.66164 | 0.00048 | 0.002762 |
| XDH | 3.841662 | 0.00048 | 0.002762 |
| ENSAPLG00000002886 | -1.02103 | 0.000481 | 0.002766 |
| NT5C1A | 3.656902 | 0.000484 | 0.00278 |
| PPP4R1 | 0.658607 | 0.000486 | 0.002794 |
| HOXD8 | 1.685029 | 0.000487 | 0.002797 |
| RNF220 | 0.723902 | 0.000489 | 0.002805 |
| MMP24 | 0.819106 | 0.00049 | 0.002806 |
| UNC93B1 | -0.93863 | 0.00049 | 0.002806 |
| DNAJC24 | -1.07245 | 0.00049 | 0.002806 |
| ENSAPLG00000031231 | 5.744408 | 0.000492 | 0.002816 |
| BDH1 | -0.70985 | 0.000496 | 0.002841 |
| ENSAPLG00000019504 | 0.879716 | 0.000498 | 0.002849 |
| KRAS | 0.765717 | 0.000502 | 0.002871 |
| ENSAPLG00000008830 | -1.1973 | 0.000503 | 0.002875 |
| ENSAPLG00000029764 | 1.828137 | 0.000503 | 0.002876 |
| ENSAPLG00000013916 | 4.620164 | 0.000505 | 0.002885 |
| TMEM178B | 0.944023 | 0.000507 | 0.002892 |
| RSPRY1 | 0.795858 | 0.000507 | 0.002892 |
| ST6GALNAC4 | 0.585538 | 0.000508 | 0.002895 |
| DLGAP4 | 0.629677 | 0.000511 | 0.002912 |
| ADCK2 | -1.83378 | 0.000513 | 0.002923 |
| ENSAPLG00000025150 | -0.75249 | 0.000514 | 0.002923 |
| ZDHHC16 | -0.78825 | 0.000514 | 0.002924 |
| SPECC1 | -0.63199 | 0.000517 | 0.002937 |
| LMOD3 | 1.816258 | 0.000517 | 0.002937 |
| PHEX | -1.14804 | 0.000524 | 0.002975 |
| GSDME | -0.78971 | 0.000525 | 0.002978 |
| ENSAPLG00000015426 | -0.78681 | 0.000526 | 0.002983 |
| ENSAPLG00000021773 | 0.737215 | 0.000526 | 0.002983 |
| CREM | 0.832859 | 0.000528 | 0.002991 |
| ENSAPLG00000017604 | 1.558485 | 0.000528 | 0.002991 |
| CLIC2 | 0.706926 | 0.000528 | 0.002992 |
| NEDD9 | 0.679019 | 0.000529 | 0.002992 |
| SCAMP1 | -0.60212 | 0.000529 | 0.002992 |
| ENSAPLG00000004765 | -1.48585 | 0.00053 | 0.002992 |
| MAB21L1 | 1.097584 | 0.000529 | 0.002992 |
| HAL | 3.041824 | 0.000533 | 0.003007 |
| ENSAPLG00000004064 | 1.476164 | 0.000533 | 0.003007 |
| LOXL1 | 0.731811 | 0.000533 | 0.003009 |

| | | | |
|---|---|---|---|
| MYEF2 | -0.75359 | 0.000534 | 0.003012 |
| CDK14 | 0.638478 | 0.000536 | 0.003021 |
| TUBA8 | 3.588729 | 0.000536 | 0.003021 |
| NUP93 | 0.58186 | 0.000537 | 0.003022 |
| CACTIN | 0.586615 | 0.00054 | 0.003037 |
| CILK1 | 1.019144 | 0.00054 | 0.003038 |
| SYNGR1 | 0.838078 | 0.000542 | 0.003047 |
| DELE1 | -1.82698 | 0.000543 | 0.003053 |
| DIAPH2 | -0.64355 | 0.000548 | 0.003076 |
| ARFIP2 | 0.649893 | 0.000548 | 0.003077 |
| HMGCS1 | -0.55702 | 0.000555 | 0.003112 |
| STXBP1 | 0.83913 | 0.000559 | 0.003134 |
| RAB9A | 0.637536 | 0.000561 | 0.00314 |
| CTSK | -0.87393 | 0.000561 | 0.00314 |
| LRIT2 | 5.721531 | 0.000561 | 0.003142 |
| GSK3B | 0.666148 | 0.000564 | 0.003155 |
| MPRIP | 0.900306 | 0.000566 | 0.003167 |
| NBR1 | -0.98344 | 0.000568 | 0.003175 |
| ARSK | -1.13596 | 0.000571 | 0.003191 |
| ZC3H13 | 0.600331 | 0.000575 | 0.00321 |
| ENSAPLG00000028225 | 0.834777 | 0.000581 | 0.00324 |
| CCT6B | -0.57808 | 0.000583 | 0.003252 |
| MCMBP | -0.54331 | 0.000585 | 0.003263 |
| ENSAPLG00000010086 | 0.730292 | 0.000586 | 0.003265 |
| LINS1 | -1.97052 | 0.000603 | 0.003357 |
| TENT4B | 0.641761 | 0.000605 | 0.003368 |
| GATA3 | 3.703621 | 0.000606 | 0.003373 |
| ENSAPLG00000025867 | -0.56544 | 0.000609 | 0.003384 |
| CRISPLD2 | 1.185234 | 0.000609 | 0.003386 |
| YWHAZ | -0.65808 | 0.00061 | 0.003387 |
| SLC25A5 | -0.82104 | 0.000617 | 0.003424 |
| NARS2 | -0.83628 | 0.000618 | 0.003431 |
| CAPN9 | 0.791051 | 0.000621 | 0.003445 |
| ENSAPLG00000009361 | 1.757698 | 0.000622 | 0.003448 |
| ENSAPLG00000004917 | 1.096828 | 0.000625 | 0.003464 |
| PTGER4 | 1.276819 | 0.000627 | 0.003475 |
| ENSAPLG00000025374 | -0.64636 | 0.000629 | 0.003479 |
| ENSAPLG00000021250 | 5.705613 | 0.00063 | 0.003488 |
| CANX | -0.54801 | 0.000633 | 0.003497 |
| TPRKB | -0.69741 | 0.000633 | 0.003499 |
| CEP57 | -0.75179 | 0.000636 | 0.003513 |
| ENSAPLG00000005367 | 1.850319 | 0.000638 | 0.00352 |
| FPGT | -0.86875 | 0.00064 | 0.00353 |
| IL1R2 | 0.678617 | 0.000643 | 0.003544 |
| DARS2 | -0.87661 | 0.000644 | 0.00355 |

| | | | |
|---|---|---|---|
| ENSAPLG00000015505 | 0.973451 | 0.000645 | 0.003551 |
| NR6A1 | 0.927275 | 0.000645 | 0.003551 |
| ENSAPLG00000004424 | 5.799511 | 0.000646 | 0.003557 |
| RUVBL1 | -0.60568 | 0.00065 | 0.003576 |
| EGLN3 | 1.791932 | 0.000651 | 0.003582 |
| ENSAPLG00000018316 | 1.541253 | 0.000652 | 0.003582 |
| CYP39A1 | -1.00977 | 0.000655 | 0.0036 |
| FBXO44 | -1.1771 | 0.000658 | 0.00361 |
| MRPL41 | -0.72991 | 0.000664 | 0.003646 |
| ENSAPLG00000030810 | -1.02558 | 0.000665 | 0.003646 |
| SMARCA2 | -0.60252 | 0.000666 | 0.003654 |
| GPRC5B | 1.476514 | 0.000669 | 0.003663 |
| ENSAPLG00000028706 | -0.64405 | 0.00067 | 0.003671 |
| ENSAPLG00000012588 | -0.8137 | 0.000676 | 0.0037 |
| NSG1 | -0.82158 | 0.000677 | 0.003701 |
| ENSAPLG00000014522 | -0.7545 | 0.00068 | 0.003721 |
| ARL9 | -0.71962 | 0.000682 | 0.003729 |
| HTATSF1 | -0.65984 | 0.000683 | 0.003729 |
| FRMD3 | 2.531124 | 0.000683 | 0.003729 |
| ENSAPLG00000030819 | -1.17375 | 0.000683 | 0.003729 |
| APOO | -0.6735 | 0.000686 | 0.003738 |
| EPN2 | 0.631014 | 0.000686 | 0.003738 |
| ENSAPLG00000014299 | 1.151136 | 0.000691 | 0.003767 |
| CYP2R1 | -0.88227 | 0.000693 | 0.003773 |
| UGGT2 | -0.80425 | 0.000697 | 0.003797 |
| DCK | -0.83054 | 0.000698 | 0.003798 |
| FH | -0.61817 | 0.000701 | 0.00381 |
| ENSAPLG00000004770 | 0.598109 | 0.000706 | 0.003835 |
| FZD1 | 0.79075 | 0.000706 | 0.003837 |
| MSL1 | 0.824 | 0.00071 | 0.003854 |
| NAT10 | -0.70635 | 0.000712 | 0.003865 |
| RERE | 0.758055 | 0.000714 | 0.003872 |
| TP53BP2 | 0.798215 | 0.000714 | 0.003872 |
| SAV1 | 0.575564 | 0.000715 | 0.003872 |
| C1S | -0.85727 | 0.000719 | 0.003892 |
| OGDHL | 3.70102 | 0.000727 | 0.003934 |
| ENSAPLG00000016289 | -0.67254 | 0.000727 | 0.003935 |
| UBE2I | 0.623089 | 0.00073 | 0.003946 |
| RAPGEF1 | 0.702333 | 0.000732 | 0.003958 |
| CYTIP | 1.659208 | 0.00074 | 0.003999 |
| ENSAPLG00000021626 | 0.780995 | 0.00074 | 0.003999 |
| ATG7 | -0.63862 | 0.000742 | 0.004003 |
| ARMC10 | -1.00921 | 0.000742 | 0.004003 |
| KATNA1 | -0.92438 | 0.000743 | 0.004005 |
| ARHGAP28 | -0.66069 | 0.000747 | 0.004026 |

| | | | |
|---|---|---|---|
| SEC61G | 0.763918 | 0.000748 | 0.00403 |
| EPS15L1 | -0.70392 | 0.000755 | 0.004063 |
| ENSAPLG00000016639 | -1.04363 | 0.000755 | 0.004063 |
| TUSC2 | 1.115686 | 0.000755 | 0.004063 |
| PSMC3 | -0.60205 | 0.000756 | 0.004065 |
| LASP1 | 0.758641 | 0.000757 | 0.004066 |
| SGPP1 | -0.99058 | 0.000759 | 0.004076 |
| CCNDBP1 | -0.59566 | 0.000765 | 0.004107 |
| GEN1 | 0.871511 | 0.000766 | 0.00411 |
| KLHL36 | 0.620232 | 0.000766 | 0.00411 |
| GPATCH2L | 0.875452 | 0.000767 | 0.00411 |
| RAPGEF5 | 1.164875 | 0.000768 | 0.00411 |
| LRP6 | 0.689525 | 0.000767 | 0.00411 |
| ENSAPLG00000023413 | 4.510283 | 0.000768 | 0.00411 |
| SESTD1 | -0.59605 | 0.000771 | 0.004125 |
| AKR1A1 | -0.79978 | 0.000771 | 0.004125 |
| USP14 | -0.64496 | 0.000773 | 0.00413 |
| ENSAPLG00000007074 | 0.734871 | 0.000777 | 0.004152 |
| HACL1 | -0.87155 | 0.00078 | 0.004165 |
| C16orf72 | 0.655426 | 0.00078 | 0.004165 |
| METAP1 | -0.62788 | 0.000782 | 0.004172 |
| CNPPD1 | -0.65517 | 0.000785 | 0.004186 |
| PHC2 | 0.745739 | 0.000786 | 0.004189 |
| SLC12A2 | -1.01635 | 0.000787 | 0.004191 |
| LCT | 4.553998 | 0.000789 | 0.004197 |
| MED21 | 0.680512 | 0.000789 | 0.004197 |
| ENSAPLG00000030678 | -0.90716 | 0.000792 | 0.004214 |
| C2orf69 | -0.72185 | 0.000802 | 0.004263 |
| HDLBP | 0.848042 | 0.000809 | 0.004297 |
| GOLGA5 | -0.57836 | 0.000816 | 0.004333 |
| HEATR3 | -0.61817 | 0.000816 | 0.004333 |
| INPP5D | 1.93139 | 0.000817 | 0.004335 |
| KIF5C | -0.90858 | 0.000818 | 0.004337 |
| LGR5 | 2.309514 | 0.00082 | 0.004348 |
| ENSAPLG00000005803 | -0.83336 | 0.000821 | 0.00435 |
| NEPRO | -1.02865 | 0.000823 | 0.004357 |
| ENSAPLG00000029667 | 0.739416 | 0.000823 | 0.004359 |
| ENSAPLG00000010306 | -0.99162 | 0.000825 | 0.004364 |
| ENSAPLG00000011857 | 3.539164 | 0.000826 | 0.004366 |
| INTS10 | -0.57951 | 0.000832 | 0.004399 |
| RPL34 | 0.683968 | 0.000833 | 0.004401 |
| GNG2 | 0.782348 | 0.000833 | 0.004401 |
| CDK10 | -0.72364 | 0.000839 | 0.004429 |
| ENSAPLG00000023922 | 1.495541 | 0.00084 | 0.004429 |
| FUBP1 | -0.56787 | 0.000845 | 0.004455 |

| | | | |
|---|---|---|---|
| XPO7 | 0.669482 | 0.000847 | 0.004464 |
| SMARCAL1 | -0.87342 | 0.000849 | 0.00447 |
| ZDHHC9 | 0.560203 | 0.000855 | 0.004501 |
| CYTH1 | 0.614282 | 0.000856 | 0.004505 |
| UBE2V2 | 0.606967 | 0.000862 | 0.004535 |
| ARHGEF3 | -0.63816 | 0.000865 | 0.004547 |
| CLUAP1 | -0.68993 | 0.000866 | 0.004551 |
| BACH2 | 1.307649 | 0.000867 | 0.004556 |
| GALNT11 | 0.552351 | 0.000868 | 0.004558 |
| ENSAPLG00000021804 | 0.740942 | 0.00087 | 0.004563 |
| RAI14 | -0.557 | 0.00087 | 0.004564 |
| WDR70 | -0.63367 | 0.000872 | 0.004569 |
| FNDC4 | 1.005234 | 0.000873 | 0.004574 |
| ASB5 | 0.649376 | 0.000876 | 0.004586 |
| COL12A1 | -0.60378 | 0.000877 | 0.004587 |
| TTC9 | 1.362286 | 0.000877 | 0.004587 |
| IL13RA1 | -1.302 | 0.000885 | 0.004626 |
| PUM2 | 0.577673 | 0.000885 | 0.004626 |
| NARF | 0.893909 | 0.000892 | 0.004659 |
| BCS1L | -0.69421 | 0.000896 | 0.00468 |
| C1orf112 | -1.01538 | 0.000898 | 0.004684 |
| EDN1 | -0.65313 | 0.000898 | 0.004684 |
| ENSAPLG00000016181 | -1.07296 | 0.000898 | 0.004685 |
| PGR | 2.392561 | 0.000899 | 0.004688 |
| DNALI1 | -1.12945 | 0.000904 | 0.004708 |
| RRM2 | -2.00914 | 0.000905 | 0.004714 |
| BEGAIN | 4.042158 | 0.00091 | 0.004734 |
| SEC16A | 0.920418 | 0.00091 | 0.004734 |
| NIT2 | -0.88996 | 0.00091 | 0.004734 |
| ENSAPLG00000022704 | 1.146295 | 0.000913 | 0.004744 |
| ENSAPLG00000030287 | -1.15624 | 0.000914 | 0.004748 |
| NELFA | 0.907873 | 0.000917 | 0.004756 |
| ENSAPLG00000009060 | 0.962458 | 0.000916 | 0.004756 |
| ATP6V1B2 | -0.51821 | 0.000917 | 0.004756 |
| CRNKL1 | -0.54857 | 0.00092 | 0.004768 |
| NAALADL2 | -0.79455 | 0.000922 | 0.004778 |
| PPM1E | 0.66602 | 0.000922 | 0.004778 |
| NR2C2 | 0.682829 | 0.000925 | 0.004791 |
| RFXANK | 0.606248 | 0.000926 | 0.004791 |
| RAPGEF4 | -1.69492 | 0.00093 | 0.00481 |
| ARHGAP10 | -0.61375 | 0.000934 | 0.004826 |
| HSPD1 | -0.60281 | 0.000934 | 0.004827 |
| SOS2 | 0.941427 | 0.00094 | 0.004857 |
| TMEM43 | -0.52827 | 0.000945 | 0.004881 |
| ENSAPLG00000007885 | -1.57724 | 0.000947 | 0.004886 |

| | | | |
|---|---|---|---|
| LGALS1 | 0.772654 | 0.000947 | 0.004886 |
| SERP1 | 0.826355 | 0.000954 | 0.004917 |
| MAP3K7CL | 0.8335 | 0.000957 | 0.00493 |
| GABRR1 | 4.457808 | 0.000959 | 0.004938 |
| ENSAPLG00000015060 | 0.755622 | 0.000961 | 0.004949 |
| ENSAPLG00000003643 | -0.52225 | 0.000964 | 0.004959 |
| ENSAPLG00000006994 | 3.588895 | 0.000972 | 0.005 |
| METTL4 | -0.75414 | 0.000974 | 0.005009 |
| TTC26 | -0.67852 | 0.000986 | 0.005068 |
| RTKN2 | 1.662137 | 0.000994 | 0.005105 |
| FILIP1L | 0.698685 | 0.000998 | 0.005122 |
| ENSAPLG00000017150 | 1.7989 | 0.000998 | 0.005122 |
| CCNK | 0.679949 | 0.001 | 0.005128 |
| FAHD1 | -0.82746 | 0.001009 | 0.005172 |
| BPNT1 | -0.77849 | 0.001015 | 0.005202 |
| ENSAPLG00000008904 | 1.285669 | 0.001016 | 0.005202 |
| LCLAT1 | 0.620376 | 0.001017 | 0.005204 |
| TJP1 | -0.72358 | 0.00102 | 0.005219 |
| C2orf76 | 0.828303 | 0.001029 | 0.005261 |
| MPST | -0.6439 | 0.001029 | 0.005261 |
| ENSAPLG00000010371 | -0.62716 | 0.001034 | 0.005279 |
| TBCD | -0.82555 | 0.001033 | 0.005279 |
| ENSAPLG00000003521 | -0.89168 | 0.001035 | 0.005281 |
| SGMS2 | 0.616189 | 0.001035 | 0.005281 |
| ENSAPLG00000007825 | -0.82882 | 0.001037 | 0.005288 |
| OAZ2 | 0.672935 | 0.001038 | 0.005292 |
| CREBL2 | 0.736055 | 0.001046 | 0.005327 |
| ALDOC | -0.58207 | 0.001046 | 0.005327 |
| ENSAPLG00000029063 | -1.14227 | 0.001049 | 0.005338 |
| KPNA4 | -0.51719 | 0.001054 | 0.005364 |
| PDGFRL | -0.70174 | 0.001056 | 0.005371 |
| GIT1 | 1.450605 | 0.001057 | 0.005373 |
| ENSAPLG00000006173 | -0.71018 | 0.00106 | 0.005383 |
| CCDC126 | 0.548823 | 0.00106 | 0.005383 |
| RFX6 | 4.549559 | 0.001063 | 0.005393 |
| ENSAPLG00000008051 | 0.584824 | 0.001068 | 0.005419 |
| PIK3C3 | -0.69368 | 0.001073 | 0.005442 |
| C6orf62 | 0.600983 | 0.001076 | 0.005453 |
| SDF2 | -0.86068 | 0.001077 | 0.005457 |
| ENSAPLG00000016133 | -0.77013 | 0.00108 | 0.005467 |
| TRUB1 | -0.85208 | 0.001081 | 0.005468 |
| PHB | -0.57506 | 0.001081 | 0.005468 |
| LSM5 | 0.643578 | 0.001081 | 0.005469 |
| MAT2B | -0.56226 | 0.001091 | 0.005513 |
| CIDEA | -0.95704 | 0.001093 | 0.005521 |

| | | | |
|---|---|---|---|
| ID3 | 0.796661 | 0.001098 | 0.005545 |
| TP53I3 | -0.89137 | 0.001101 | 0.005558 |
| USP31 | 1.067597 | 0.001103 | 0.005563 |
| AGAP3 | 0.603413 | 0.001103 | 0.005563 |
| ENSAPLG00000012564 | -0.62617 | 0.001112 | 0.005605 |
| PM20D1 | -0.83464 | 0.001114 | 0.005614 |
| IGF2BP2 | 0.60284 | 0.001119 | 0.005635 |
| MCEE | -0.94758 | 0.001122 | 0.005644 |
| IREB2 | -0.58456 | 0.001122 | 0.005644 |
| STK17B | -0.68093 | 0.001122 | 0.005645 |
| MAP3K5 | 0.70352 | 0.001124 | 0.00565 |
| HMGA2 | 0.837138 | 0.00113 | 0.005678 |
| CTSB | -0.56576 | 0.001143 | 0.005741 |
| PHETA1 | -0.91373 | 0.001144 | 0.005745 |
| LATS1 | 0.694309 | 0.001149 | 0.005764 |
| IGF2BP3 | -0.72222 | 0.00115 | 0.005765 |
| FRMD4B | 0.82792 | 0.001152 | 0.005772 |
| PACC1 | -0.72483 | 0.001152 | 0.005772 |
| NOC3L | -0.67042 | 0.001154 | 0.00578 |
| RPS6KC1 | -0.80421 | 0.001166 | 0.005836 |
| TMEM169 | 0.722163 | 0.001166 | 0.005836 |
| IFT57 | -0.67283 | 0.00117 | 0.00585 |
| BPNT2 | -0.56191 | 0.00118 | 0.005898 |
| SNCG | 3.119962 | 0.001181 | 0.005899 |
| CRACD | 1.117668 | 0.001186 | 0.005922 |
| ZEB2 | 0.676732 | 0.00119 | 0.005938 |
| ARPP19 | 0.508658 | 0.00119 | 0.005938 |
| URI1 | 0.537957 | 0.001196 | 0.005966 |
| PCOLCE2 | -0.6076 | 0.0012 | 0.005981 |
| FNDC7 | 2.31697 | 0.001202 | 0.00599 |
| EPHB1 | 0.866366 | 0.001206 | 0.006001 |
| PEX5 | -0.72722 | 0.001206 | 0.006001 |
| STRAP | -0.542 | 0.001206 | 0.006001 |
| MCOLN1 | 0.600189 | 0.001208 | 0.006009 |
| CDCA4 | 0.640737 | 0.001212 | 0.006024 |
| ENSAPLG00000023996 | -0.90619 | 0.001212 | 0.006024 |
| SLC4A7 | -0.58182 | 0.001214 | 0.006031 |
| RNF10 | 0.840711 | 0.001218 | 0.006045 |
| SPTLC3 | -0.9301 | 0.001221 | 0.006059 |
| FAM98A | -0.67987 | 0.001222 | 0.006061 |
| ENSAPLG00000019789 | -0.66659 | 0.001227 | 0.006082 |
| FJX1 | 1.558374 | 0.001228 | 0.006084 |
| UQCRB | 0.70457 | 0.001229 | 0.00609 |
| MAMLD1 | 0.712505 | 0.00123 | 0.006092 |
| ENSAPLG00000005846 | 0.786064 | 0.001232 | 0.006095 |

| | | | |
|---|---|---|---|
| ENSAPLG00000018473 | -0.73421 | 0.001234 | 0.006104 |
| ENSAPLG00000027541 | 0.849639 | 0.001238 | 0.006121 |
| PPFIBP1 | 0.649911 | 0.001239 | 0.006125 |
| FAM204A | 0.693609 | 0.001244 | 0.006143 |
| ARHGEF6 | 0.611373 | 0.001244 | 0.006143 |
| MAP3K7 | 0.565335 | 0.00125 | 0.006164 |
| GTF2B | -0.69053 | 0.00125 | 0.006164 |
| LMF2 | -1.13857 | 0.001249 | 0.006164 |
| PELO | -0.5955 | 0.001251 | 0.006166 |
| CDC42BPA | 0.681743 | 0.001253 | 0.006172 |
| BBS12 | -0.77279 | 0.001257 | 0.006186 |
| ENSAPLG00000006553 | 1.199023 | 0.001257 | 0.006186 |
| CCT5 | -0.54999 | 0.001258 | 0.006188 |
| SIRT5 | -1.03755 | 0.001263 | 0.006208 |
| CHRNB2 | 1.166889 | 0.001263 | 0.006208 |
| AIMP1 | -0.821 | 0.001269 | 0.006237 |
| SORBS1 | -1.65672 | 0.001283 | 0.006303 |
| RWDD3 | -0.80166 | 0.001285 | 0.006306 |
| GRIA3 | 2.452745 | 0.001293 | 0.006346 |
| SEPTIN11 | -0.54743 | 0.0013 | 0.006377 |
| ACSBG1 | 0.701235 | 0.00131 | 0.00642 |
| BOC | 0.74301 | 0.00131 | 0.00642 |
| DHX40 | -0.74919 | 0.001313 | 0.006429 |
| ME1 | -0.68491 | 0.001317 | 0.006447 |
| PABPC4 | 0.686397 | 0.001323 | 0.006473 |
| OTUD1 | 0.725047 | 0.001323 | 0.006473 |
| DUSP12 | -0.85204 | 0.00133 | 0.006503 |
| AIDA | 0.579119 | 0.001333 | 0.006513 |
| MYLK | -0.7286 | 0.001335 | 0.006517 |
| ENSAPLG00000026904 | 0.946473 | 0.001334 | 0.006517 |
| DUS1L | -0.57643 | 0.001338 | 0.006532 |
| MSN | -0.54186 | 0.001341 | 0.006539 |
| ENSAPLG00000013698 | 0.652038 | 0.001344 | 0.00655 |
| ZNF608 | 1.884045 | 0.001343 | 0.00655 |
| DCAF13 | -0.55914 | 0.00135 | 0.00657 |
| MYO1E | 0.660508 | 0.00135 | 0.00657 |
| NIF3L1 | -0.80102 | 0.00135 | 0.00657 |
| SNX1 | -0.53432 | 0.001351 | 0.006571 |
| ENSAPLG00000021285 | 2.31637 | 0.001351 | 0.006571 |
| ETV4 | 0.561851 | 0.001354 | 0.006583 |
| ATP6AP2 | -0.54578 | 0.001358 | 0.006597 |
| SPATA7 | -1.02087 | 0.001362 | 0.006613 |
| EDNRA | -0.67595 | 0.001364 | 0.006623 |
| BCKDHB | -0.79501 | 0.001373 | 0.006662 |
| ENSAPLG00000002685 | -2.10387 | 0.001376 | 0.006668 |

| | | | |
|---|---|---|---|
| SLC9A5 | 1.16351 | 0.001376 | 0.006668 |
| SKIL | 0.65264 | 0.001376 | 0.006668 |
| ABHD17B | -0.60436 | 0.001379 | 0.006678 |
| ENSAPLG00000002945 | -0.95535 | 0.001386 | 0.006709 |
| ZDHHC14 | 0.635774 | 0.001393 | 0.006743 |
| STT3A | -0.57005 | 0.0014 | 0.006772 |
| HTR2C | 0.878357 | 0.001407 | 0.006802 |
| PAPOLA | 0.513645 | 0.00141 | 0.006813 |
| IGF2R | -0.89677 | 0.001411 | 0.006817 |
| ADAM22 | 3.459978 | 0.001412 | 0.006818 |
| ATG13 | 0.73238 | 0.001416 | 0.006835 |
| RCC1 | 1.045122 | 0.001418 | 0.006839 |
| MVK | -0.6285 | 0.001423 | 0.006859 |
| RPAP3 | -0.6297 | 0.001423 | 0.006859 |
| EMC3 | -0.6327 | 0.001426 | 0.006866 |
| MYO9A | 0.941729 | 0.001426 | 0.006866 |
| GLA | -0.742 | 0.00144 | 0.00693 |
| EFNA5 | 1.064403 | 0.001441 | 0.006935 |
| IL1RL2 | 0.704469 | 0.001449 | 0.006969 |
| RGS2 | -0.82333 | 0.00145 | 0.006969 |
| CCDC47 | -0.54551 | 0.001453 | 0.006981 |
| KCNS2 | 2.162344 | 0.00147 | 0.007059 |
| VPS26A | -0.55184 | 0.001476 | 0.007085 |
| ENSAPLG00000018541 | 0.753018 | 0.00148 | 0.007102 |
| TGFB3 | -0.58026 | 0.001483 | 0.007112 |
| ENSAPLG00000020546 | 0.658909 | 0.001483 | 0.007112 |
| ENSAPLG00000018877 | 0.719975 | 0.001487 | 0.007124 |
| LIPA | -0.55261 | 0.001496 | 0.007168 |
| TAB2 | 0.711636 | 0.001497 | 0.007169 |
| GNA12 | 0.671716 | 0.0015 | 0.007177 |
| ENSAPLG00000003727 | 1.084286 | 0.001506 | 0.007197 |
| ENSAPLG00000011995 | -0.60418 | 0.001506 | 0.007197 |
| PIM3 | -0.54426 | 0.001505 | 0.007197 |
| MARK3 | 0.597262 | 0.001511 | 0.007217 |
| RPLP0 | 0.632765 | 0.001512 | 0.007221 |
| SLC37A4 | -0.83515 | 0.001515 | 0.007233 |
| KTN1 | -0.60391 | 0.00152 | 0.007253 |
| TNFAIP8L1 | -1.21879 | 0.001531 | 0.0073 |
| ENSAPLG00000001261 | 0.544459 | 0.001532 | 0.007303 |
| ADNP2 | 0.723895 | 0.001533 | 0.007303 |
| PSPC1 | -0.56356 | 0.001533 | 0.007303 |
| WNT4 | 1.59684 | 0.001535 | 0.007307 |
| HDHD2 | -0.66105 | 0.001539 | 0.007322 |
| COG2 | -0.59524 | 0.00154 | 0.007325 |
| IFFO2 | 0.636889 | 0.001544 | 0.00734 |

| | | | |
|---|---|---|---|
| ENSAPLG00000009809 | 0.552372 | 0.001557 | 0.007398 |
| EML5 | 2.752532 | 0.001559 | 0.007404 |
| ETV3 | 0.998193 | 0.001564 | 0.007428 |
| IARS1 | -0.64144 | 0.001569 | 0.007448 |
| COPS9 | 4.301755 | 0.001571 | 0.007451 |
| NADSYN1 | -0.60123 | 0.001574 | 0.007464 |
| YPEL5 | 0.55116 | 0.001575 | 0.007467 |
| UBA5 | -0.72673 | 0.001579 | 0.007484 |
| MED13 | 0.769463 | 0.00158 | 0.007484 |
| ENSAPLG00000008902 | -0.68538 | 0.001582 | 0.007487 |
| TRIO | 0.766701 | 0.001582 | 0.007488 |
| ENSAPLG00000013039 | 0.693853 | 0.001589 | 0.007518 |
| CBFA2T2 | 0.692117 | 0.001593 | 0.00753 |
| STAU2 | -0.66021 | 0.001598 | 0.007552 |
| ENSAPLG00000018685 | 1.369135 | 0.001599 | 0.007552 |
| ALKBH3 | 0.617948 | 0.0016 | 0.007553 |
| ENSAPLG00000003298 | 0.889583 | 0.001611 | 0.007605 |
| ENSAPLG00000011877 | 0.709218 | 0.001612 | 0.007607 |
| CHTOP | 0.658974 | 0.001616 | 0.007621 |
| ENSAPLG00000005376 | 3.88211 | 0.001621 | 0.007642 |
| APPL2 | -0.67845 | 0.001623 | 0.007649 |
| SHANK3 | 1.194045 | 0.001625 | 0.007652 |
| ENSAPLG00000009571 | -1.0292 | 0.001626 | 0.007656 |
| NTPCR | -0.90449 | 0.001638 | 0.007706 |
| ECHDC3 | -1.067 | 0.001639 | 0.007706 |
| ENSAPLG00000030814 | -0.89084 | 0.001639 | 0.007706 |
| MXRA5 | 2.050029 | 0.001648 | 0.007742 |
| PHF3 | 0.588013 | 0.001655 | 0.007773 |
| TWIST2 | 2.051408 | 0.001664 | 0.007811 |
| PTCHD3 | -1.05165 | 0.001669 | 0.007831 |
| ENSAPLG00000004124 | -0.5497 | 0.001681 | 0.007884 |
| ENSAPLG00000006340 | 2.600828 | 0.001687 | 0.00791 |
| CASP3 | -0.58988 | 0.001692 | 0.007931 |
| GAS6 | -0.92758 | 0.001702 | 0.007974 |
| ARV1 | -0.71248 | 0.001705 | 0.007985 |
| RBMS3 | 0.562132 | 0.001712 | 0.008014 |
| GPC3 | -0.81375 | 0.001723 | 0.00806 |
| MB21D2 | -0.88629 | 0.001736 | 0.008118 |
| TGFBR1 | -0.51385 | 0.001744 | 0.008151 |
| TRA2A | -0.56327 | 0.001744 | 0.008151 |
| ENSAPLG00000008001 | -0.53761 | 0.001755 | 0.008197 |
| RPSA | 0.634364 | 0.001761 | 0.008223 |
| ENSAPLG00000003498 | -0.82777 | 0.001766 | 0.00824 |
| SLC12A4 | -0.75162 | 0.001778 | 0.008294 |
| MAP1LC3A | 0.727016 | 0.001779 | 0.008294 |

| | | | |
|---|---|---|---|
| TCEANC2 | -0.87009 | 0.001786 | 0.008326 |
| BMP15 | 4.927037 | 0.001793 | 0.008354 |
| GPX7 | -0.91873 | 0.001804 | 0.008402 |
| ALOX5 | -0.9388 | 0.001809 | 0.008417 |
| RFC4 | -0.6569 | 0.001808 | 0.008417 |
| CHRNA3 | 3.846357 | 0.00181 | 0.008419 |
| SLC13A4 | 1.35175 | 0.001812 | 0.008424 |
| SGMS1 | 0.577777 | 0.001815 | 0.008425 |
| WARS1 | -0.54708 | 0.001813 | 0.008425 |
| PHACTR2 | 1.021347 | 0.001814 | 0.008425 |
| GRIP2 | 1.047969 | 0.001817 | 0.008431 |
| TLE3 | 1.573239 | 0.001822 | 0.008454 |
| PALD1 | 0.630265 | 0.001829 | 0.008472 |
| ENSAPLG00000018454 | 0.733289 | 0.001827 | 0.008472 |
| ENSAPLG00000019298 | -0.82611 | 0.001828 | 0.008472 |
| TOP1MT | -0.58915 | 0.001836 | 0.0085 |
| NPRL2 | -0.62334 | 0.001836 | 0.0085 |
| MIIP | -1.02719 | 0.001842 | 0.008523 |
| GFUS | -0.88339 | 0.001846 | 0.008537 |
| MBNL2 | 0.567617 | 0.001849 | 0.008548 |
| ENSAPLG00000006357 | 2.275405 | 0.00185 | 0.008552 |
| DTWD1 | -0.9135 | 0.001861 | 0.008598 |
| ENSAPLG00000018683 | 0.784391 | 0.001862 | 0.008598 |
| RSBN1 | 0.629091 | 0.001868 | 0.008624 |
| AMIGO2 | 1.523225 | 0.001869 | 0.008625 |
| ENSAPLG00000004582 | -1.41681 | 0.001873 | 0.008632 |
| ARFGAP2 | -0.68805 | 0.001872 | 0.008632 |
| PRRC2B | 0.812347 | 0.001876 | 0.008645 |
| ENSAPLG00000022799 | 1.538449 | 0.00188 | 0.008661 |
| MAX | 2.015022 | 0.001891 | 0.008707 |
| ITPRIPL2 | 0.717217 | 0.001895 | 0.008721 |
| POLR2K | 0.784286 | 0.001901 | 0.008745 |
| TUBE1 | 0.743237 | 0.001903 | 0.008752 |
| RALBP1 | 0.5905 | 0.001914 | 0.008794 |
| ENSAPLG00000015697 | 0.81248 | 0.001914 | 0.008794 |
| UROD | -0.80491 | 0.001917 | 0.008804 |
| ENSAPLG00000007167 | 0.678894 | 0.001923 | 0.008825 |
| PFDN4 | 0.613854 | 0.001923 | 0.008825 |
| INTS12 | -0.65898 | 0.001925 | 0.008827 |
| ENSAPLG00000012987 | 0.623753 | 0.001926 | 0.008827 |
| ENSAPLG00000019806 | 1.998742 | 0.001926 | 0.008827 |
| TFB1M | -0.78601 | 0.001944 | 0.008903 |
| AHRR | 0.844139 | 0.001961 | 0.008978 |
| IFT46 | 0.615164 | 0.001963 | 0.008984 |
| HDAC10 | -0.69366 | 0.001968 | 0.009001 |

| | | | |
|---|---|---|---|
| NLGN1 | -1.37939 | 0.001971 | 0.009015 |
| ENSAPLG00000024173 | -0.92273 | 0.001974 | 0.009024 |
| ALAD | -1.07628 | 0.001988 | 0.009084 |
| TLK1 | -0.64371 | 0.001991 | 0.009093 |
| DNAJA2 | -0.52605 | 0.001993 | 0.009098 |
| ENSAPLG00000012312 | -0.83413 | 0.001996 | 0.009105 |
| NEK7 | -0.57366 | 0.001996 | 0.009105 |
| AGO1 | 0.794074 | 0.002004 | 0.009138 |
| ENSAPLG00000003513 | 0.801022 | 0.002011 | 0.009165 |
| TBC1D15 | -0.5349 | 0.002013 | 0.009172 |
| LBH | 0.466389 | 0.002025 | 0.009221 |
| SOX5 | -0.89544 | 0.002039 | 0.009279 |
| MMP27 | -1.14586 | 0.002039 | 0.009279 |
| CEP350 | 0.793416 | 0.00204 | 0.00928 |
| ULK2 | 0.660769 | 0.002042 | 0.009282 |
| GTF2H1 | 0.587655 | 0.002046 | 0.009296 |
| IQSEC1 | 0.696347 | 0.002048 | 0.009301 |
| CCNF | 0.667739 | 0.002058 | 0.009342 |
| ENSAPLG00000013790 | -0.63945 | 0.002063 | 0.009361 |
| DLL1 | 0.864561 | 0.002064 | 0.009363 |
| PPP2CB | 0.539812 | 0.002066 | 0.009369 |
| DDX59 | 0.593891 | 0.002068 | 0.009375 |
| SLC25A21 | -1.12263 | 0.00207 | 0.009381 |
| PSMD1 | -0.48232 | 0.002078 | 0.009408 |
| OTUB1 | -0.58774 | 0.002078 | 0.009408 |
| CHCHD7 | 0.806723 | 0.002093 | 0.00947 |
| PKM | -0.53781 | 0.002101 | 0.009483 |
| MBTPS1 | 0.619041 | 0.002097 | 0.009483 |
| ENSAPLG00000006354 | 1.541889 | 0.0021 | 0.009483 |
| SLC4A5 | 3.751908 | 0.0021 | 0.009483 |
| FAM13C | -1.16803 | 0.0021 | 0.009483 |
| ENSAPLG00000027048 | 1.613964 | 0.0021 | 0.009483 |
| TRMT1L | -0.55772 | 0.002102 | 0.009485 |
| WSB2 | 0.524154 | 0.00211 | 0.009515 |
| SNX11 | -0.69656 | 0.002116 | 0.00954 |
| SERTM1 | 1.015 | 0.002123 | 0.009569 |
| ENSAPLG00000005844 | -0.64644 | 0.002127 | 0.009581 |
| FDX1 | -0.71407 | 0.002129 | 0.009587 |
| POLR2F | 0.740272 | 0.002137 | 0.009619 |
| NSUN2 | -0.54702 | 0.002141 | 0.009634 |
| MAPK12 | -1.19956 | 0.002142 | 0.009634 |
| FLAD1 | -0.89742 | 0.002144 | 0.009638 |
| ENSAPLG00000006799 | -0.56567 | 0.002146 | 0.009642 |
| UBAC2 | 0.576198 | 0.002151 | 0.00966 |
| PABIR2 | 0.531174 | 0.002151 | 0.00966 |

| | | | |
|---|---|---|---|
| ENSAPLG00000024940 | 0.815085 | 0.002152 | 0.00966 |
| GYPC | 0.940926 | 0.002157 | 0.009676 |
| EFEMP1 | -0.54746 | 0.002159 | 0.009678 |
| FKBP4 | -0.52168 | 0.002159 | 0.009678 |
| ENSAPLG00000004972 | 0.687635 | 0.002162 | 0.009689 |
| ENSAPLG00000005834 | -0.6097 | 0.002166 | 0.009701 |
| ANKRD13D | 0.645038 | 0.002167 | 0.009702 |
| HIKESHI | 0.670689 | 0.002168 | 0.009705 |
| LRRC3B | -1.0833 | 0.002178 | 0.009741 |
| CETP | 1.525825 | 0.002178 | 0.009741 |
| CHRM4 | 1.937811 | 0.002182 | 0.009751 |
| RNF14 | -0.68122 | 0.002187 | 0.00977 |
| CDS2 | -0.58637 | 0.002189 | 0.009776 |
| MFSD11 | 0.579124 | 0.002199 | 0.009816 |
| MRPL21 | 0.597247 | 0.0022 | 0.009817 |
| TADA1 | -0.65437 | 0.002206 | 0.00984 |
| FAM78B | 1.587781 | 0.002216 | 0.009882 |
| ISCA1 | 0.698253 | 0.002219 | 0.009891 |
| GJC2 | 0.767123 | 0.002226 | 0.009917 |
| MBP | 0.793312 | 0.002227 | 0.009917 |
| PPM1K | -0.64079 | 0.00223 | 0.009927 |
| AK2 | -0.57173 | 0.002231 | 0.009927 |
| GP1BB | 2.34595 | 0.002234 | 0.009934 |
| SPARC | -0.60331 | 0.002234 | 0.009934 |
| SLC9A7 | 0.63019 | 0.002239 | 0.009951 |
| STK38L | 0.58681 | 0.002241 | 0.009954 |
| KANSL3 | 0.56867 | 0.002246 | 0.009973 |
| WASF2 | 1.04457 | 0.002253 | 0.010001 |
| ANKLE2 | 0.652812 | 0.002254 | 0.010003 |
| SNRNP40 | -0.54742 | 0.00226 | 0.010019 |
| UFSP2 | -0.51337 | 0.002259 | 0.010019 |
| FERMT1 | -0.73148 | 0.002262 | 0.010026 |
| NRBF2 | -0.64338 | 0.002276 | 0.010084 |
| MRPL1 | -0.79521 | 0.002285 | 0.01012 |
| ENSAPLG00000001572 | 1.478495 | 0.002299 | 0.010177 |
| ARNT | 0.968046 | 0.002302 | 0.010184 |
| SWAP70 | 0.548549 | 0.002306 | 0.0102 |
| BEND7 | 0.982724 | 0.002311 | 0.010215 |
| RELN | -1.11121 | 0.002315 | 0.010231 |
| ENPP4 | -0.7478 | 0.002318 | 0.01024 |
| NKD1 | 1.369929 | 0.002333 | 0.0103 |
| CEPT1 | -0.53038 | 0.002338 | 0.010314 |
| PIGU | -0.59009 | 0.002337 | 0.010314 |
| ACOT11 | 0.65025 | 0.002344 | 0.010339 |
| RSAD2 | 2.54187 | 0.002356 | 0.010387 |

| | | | |
|---|---|---|---|
| CYB5R2 | -0.71082 | 0.002364 | 0.010415 |
| ENSAPLG00000014992 | -0.88664 | 0.00237 | 0.010437 |
| AURKA | -1.04871 | 0.002371 | 0.010439 |
| CLTC | -0.71251 | 0.002372 | 0.010439 |
| HMBS | -0.61379 | 0.002376 | 0.010451 |
| ARHGEF7 | 0.543744 | 0.002377 | 0.010454 |
| ENSAPLG00000020051 | -0.6756 | 0.002385 | 0.010484 |
| PNISR | 0.633344 | 0.002388 | 0.010489 |
| ETNK2 | 1.753511 | 0.002387 | 0.010489 |
| SMAD3 | 0.738641 | 0.002389 | 0.01049 |
| UHRF1BP1L | 0.602504 | 0.002395 | 0.010513 |
| LIMCH1 | -0.85422 | 0.002404 | 0.010545 |
| EOMES | -2.03523 | 0.00241 | 0.010567 |
| SMOX | 0.533475 | 0.00242 | 0.010606 |
| ACTR5 | -0.71335 | 0.00242 | 0.010606 |
| AP3D1 | 0.544164 | 0.002422 | 0.01061 |
| CNP | 0.831852 | 0.002429 | 0.010631 |
| MTHFD1L | -0.82324 | 0.002428 | 0.010631 |
| GLTP | -0.88183 | 0.002433 | 0.010643 |
| LRP2 | 1.940055 | 0.002456 | 0.010741 |
| PACSIN2 | -0.54088 | 0.002458 | 0.010745 |
| ATP5PB | -0.62091 | 0.002465 | 0.01077 |
| ENSAPLG00000020705 | -0.86297 | 0.002467 | 0.010776 |
| CRIPT | 0.597105 | 0.00247 | 0.010786 |
| COL8A2 | -0.71973 | 0.002483 | 0.010838 |
| TNFSF13B | 2.933492 | 0.002487 | 0.010848 |
| GALM | -0.66528 | 0.002488 | 0.01085 |
| HOXA3 | 1.308142 | 0.002503 | 0.01091 |
| ENSAPLG00000009349 | -0.48837 | 0.002504 | 0.010912 |
| ENSAPLG00000002088 | 4.191438 | 0.002513 | 0.010946 |
| ENSAPLG00000011319 | 0.829572 | 0.002528 | 0.011004 |
| HNRNPM | -0.52968 | 0.002535 | 0.011034 |
| ADGRL2 | -0.98614 | 0.002537 | 0.011037 |
| ENSAPLG00000029470 | 3.262023 | 0.002542 | 0.011052 |
| SIAH3 | 4.272015 | 0.002549 | 0.01108 |
| YPEL1 | 0.612396 | 0.002559 | 0.011117 |
| WDR26 | -0.69045 | 0.002563 | 0.011131 |
| PHB2 | -0.50157 | 0.002569 | 0.011155 |
| ITGA11 | -0.65904 | 0.002572 | 0.011159 |
| PPA1 | -0.55229 | 0.002572 | 0.011159 |
| SMARCE1 | 0.518252 | 0.002575 | 0.011166 |
| ZSWIM5 | 0.668467 | 0.00259 | 0.011226 |
| SCARF1 | 3.044322 | 0.0026 | 0.011266 |
| ENSAPLG00000011821 | 0.670614 | 0.002604 | 0.011277 |
| TMEM33 | -0.49179 | 0.002626 | 0.011371 |

| | | | |
|---|---|---|---|
| LZTS2 | 0.767605 | 0.002634 | 0.0114 |
| EPAS1 | 0.796453 | 0.002643 | 0.011435 |
| SLC25A38 | 0.948739 | 0.00266 | 0.011502 |
| SCRN3 | -0.63914 | 0.002666 | 0.011522 |
| ENSAPLG00000012742 | -0.56563 | 0.002671 | 0.011541 |
| ENSAPLG00000027775 | -0.65508 | 0.002676 | 0.011557 |
| DLEC1 | 0.711764 | 0.002677 | 0.01156 |
| ENSAPLG00000020856 | -0.52623 | 0.002681 | 0.011571 |
| THRB | 0.580234 | 0.002686 | 0.01159 |
| COX18 | -0.62652 | 0.002699 | 0.011637 |
| ENSAPLG00000008548 | 0.641233 | 0.002715 | 0.011704 |
| UVRAG | 0.559776 | 0.002717 | 0.011706 |
| PLAGL2 | 0.748738 | 0.002719 | 0.011711 |
| INO80D | 0.797157 | 0.002738 | 0.011788 |
| PRPF40A | -0.53583 | 0.002746 | 0.011813 |
| PLEKHM2 | 0.827921 | 0.002746 | 0.011813 |
| TSPAN14 | -0.55229 | 0.002751 | 0.01183 |
| NFYB | -0.60532 | 0.002763 | 0.011878 |
| TECTA | 3.43907 | 0.00277 | 0.011905 |
| MAP4K4 | 0.654327 | 0.002779 | 0.011936 |
| TMEM59 | -0.68282 | 0.002782 | 0.011946 |
| USP13 | 2.711901 | 0.002788 | 0.011966 |
| MARF1 | 0.718818 | 0.002791 | 0.011975 |
| RIOX1 | -0.67775 | 0.002794 | 0.011982 |
| TPRA1 | -0.75725 | 0.002796 | 0.011985 |
| TRABD | -0.49285 | 0.002797 | 0.011986 |
| ENSAPLG00000011046 | 0.541848 | 0.002815 | 0.012058 |
| MBNL1 | 0.793332 | 0.002819 | 0.012071 |
| NOS2 | 0.977988 | 0.002822 | 0.012079 |
| CDH20 | 1.555176 | 0.002834 | 0.012107 |
| NDUFAF1 | 0.504887 | 0.002832 | 0.012107 |
| LRIG1 | 1.016491 | 0.002834 | 0.012107 |
| ENSAPLG00000028515 | -1.47128 | 0.002832 | 0.012107 |
| ENSAPLG00000029553 | -0.63234 | 0.002833 | 0.012107 |
| GATD1 | -0.80201 | 0.00284 | 0.012127 |
| ADCY3 | 2.964317 | 0.002842 | 0.012128 |
| RHOJ | 0.973192 | 0.002843 | 0.012128 |
| AP5Z1 | -0.62574 | 0.002852 | 0.012164 |
| BSG | -0.53847 | 0.002858 | 0.012186 |
| PUM3 | -0.54143 | 0.002879 | 0.01227 |
| ENSAPLG00000030615 | 1.189076 | 0.00288 | 0.01227 |
| SPATS2L | -0.66875 | 0.002902 | 0.012355 |
| ZNF518A | 0.620663 | 0.002903 | 0.012355 |
| NSUN4 | -0.68137 | 0.002906 | 0.012366 |
| SARDH | -0.94257 | 0.002908 | 0.01237 |

| | | | |
|---|---|---|---|
| RNF185 | 0.506178 | 0.002916 | 0.0124 |
| PARD6A | 0.740682 | 0.002918 | 0.012403 |
| TMEM87A | -0.47915 | 0.002931 | 0.012452 |
| KDM4C | -0.79901 | 0.002944 | 0.012503 |
| SLC35F5 | -0.57542 | 0.002945 | 0.012503 |
| NEFL | 1.065946 | 0.002966 | 0.012585 |
| RDH14 | -0.70108 | 0.002976 | 0.012625 |
| ENSAPLG00000014617 | 4.745055 | 0.00298 | 0.012634 |
| SLC40A1 | 0.750133 | 0.002987 | 0.012661 |
| CLIC6 | 4.297648 | 0.002989 | 0.012662 |
| ENSAPLG00000010495 | -0.59334 | 0.002993 | 0.012675 |
| NAA25 | -0.51867 | 0.003007 | 0.012729 |
| METTL15 | -0.79233 | 0.003012 | 0.012748 |
| HDC | 3.881392 | 0.003014 | 0.012752 |
| MFSD8 | -0.82927 | 0.003018 | 0.012762 |
| TLE4 | 0.627648 | 0.00303 | 0.012806 |
| FRMPD3 | 3.234187 | 0.003049 | 0.012882 |
| GPX1 | 0.73366 | 0.003057 | 0.012913 |
| SLC7A14 | 3.719809 | 0.003073 | 0.012973 |
| SMARCA1 | -0.61601 | 0.003086 | 0.013024 |
| EIPR1 | -0.57252 | 0.003087 | 0.013024 |
| CANT1 | -0.63592 | 0.003098 | 0.013065 |
| ENSAPLG00000019867 | 0.655661 | 0.003112 | 0.013119 |
| CCSER1 | -0.77265 | 0.003118 | 0.013139 |
| KIF20A | -1.406 | 0.003123 | 0.013156 |
| HMOX2 | -0.58238 | 0.003141 | 0.013226 |
| CCDC12 | 0.635841 | 0.003157 | 0.01329 |
| ENSAPLG00000004098 | 0.685236 | 0.003168 | 0.013329 |
| CAPG | -0.70156 | 0.003173 | 0.013344 |
| FAM3C | -0.56015 | 0.00318 | 0.013371 |
| UBE2D3 | 0.530515 | 0.003193 | 0.013417 |
| SLC25A14 | 0.509091 | 0.003195 | 0.013422 |
| TRIP13 | -0.81809 | 0.003198 | 0.013428 |
| ENSAPLG00000001730 | -0.87374 | 0.003215 | 0.01349 |
| PHACTR3 | 1.104012 | 0.003214 | 0.01349 |
| PLS3 | -0.45519 | 0.00322 | 0.013506 |
| PTCD3 | -0.68376 | 0.003226 | 0.013522 |
| TMEM266 | 1.364 | 0.003226 | 0.013522 |
| ENSAPLG00000004298 | 1.096123 | 0.003231 | 0.013538 |
| PWP1 | -0.53858 | 0.003239 | 0.013565 |
| FSTL5 | 1.454365 | 0.00326 | 0.013649 |
| ENSAPLG00000014237 | 0.790436 | 0.003262 | 0.013651 |
| ENSAPLG00000019315 | 0.795587 | 0.003282 | 0.013729 |
| AUP1 | -0.53924 | 0.003291 | 0.01376 |
| ENSAPLG00000003490 | 1.065803 | 0.003297 | 0.013782 |

| | | | |
|---|---|---|---|
| ENSAPLG00000007401 | 0.807344 | 0.003327 | 0.0139 |
| LUM | -0.79951 | 0.003328 | 0.0139 |
| ENSAPLG00000016512 | 0.86471 | 0.003338 | 0.013938 |
| TRMT44 | -0.65201 | 0.003342 | 0.013947 |
| PDZRN4 | -1.14379 | 0.00335 | 0.013975 |
| RNF130 | 0.475703 | 0.003352 | 0.01398 |
| FAM222A | 4.784149 | 0.003358 | 0.013999 |
| RAB33B | 0.602481 | 0.003373 | 0.014057 |
| ENSAPLG00000009817 | 1.717289 | 0.003393 | 0.014135 |
| FNDC5 | 1.004532 | 0.003402 | 0.014165 |
| ENSAPLG00000009985 | -0.67015 | 0.003404 | 0.014167 |
| TOPBP1 | -0.87928 | 0.003405 | 0.014168 |
| GHR | 0.845798 | 0.003411 | 0.014187 |
| CXXC5 | 0.927797 | 0.003414 | 0.014196 |
| AKAP6 | 0.866892 | 0.00342 | 0.014212 |
| ENSAPLG00000017524 | -0.5658 | 0.003422 | 0.014219 |
| ACOT7 | 0.5267 | 0.003434 | 0.014259 |
| ENSAPLG00000005925 | 0.647739 | 0.003439 | 0.014276 |
| DNAJB9 | -0.6519 | 0.003441 | 0.01428 |
| KIAA1328 | 0.843496 | 0.00345 | 0.014312 |
| RHEB | 0.508741 | 0.003464 | 0.014364 |
| WRAP73 | -0.77484 | 0.003473 | 0.014388 |
| ERAP1 | -0.907 | 0.003474 | 0.014388 |
| ZNF207 | -0.50144 | 0.003471 | 0.014388 |
| ENSAPLG00000012884 | -0.84384 | 0.003493 | 0.014461 |
| SNX21 | 0.54121 | 0.003507 | 0.014516 |
| CAPN2 | -0.52856 | 0.003514 | 0.014536 |
| TRAPPC13 | -0.56222 | 0.003517 | 0.014545 |
| ZBTB14 | 0.829859 | 0.003522 | 0.014558 |
| ENSAPLG00000017467 | 0.573255 | 0.003536 | 0.014611 |
| TMED4 | -0.51551 | 0.003537 | 0.014611 |
| ENSAPLG00000013000 | -0.48174 | 0.00354 | 0.014618 |
| POU2F3 | 2.537396 | 0.003544 | 0.014627 |
| ENSAPLG00000003738 | 0.86498 | 0.00358 | 0.01477 |
| NSMCE2 | 0.609629 | 0.003586 | 0.01479 |
| ATN1 | 1.167689 | 0.003604 | 0.014857 |
| SLC16A8 | 4.125483 | 0.003617 | 0.014906 |
| DAGLA | 0.925646 | 0.003627 | 0.014941 |
| ENSAPLG00000003917 | -0.58251 | 0.003631 | 0.014953 |
| RAD51 | -0.89791 | 0.00364 | 0.014987 |
| CALML3 | 4.093136 | 0.003646 | 0.015005 |
| HNRNPDL | -0.56161 | 0.003651 | 0.015018 |
| SLC7A6 | -0.52443 | 0.003658 | 0.015037 |
| ITGA6 | -0.73398 | 0.003657 | 0.015037 |
| GRHL2 | 0.995117 | 0.003673 | 0.015091 |

| | | | |
|---|---|---|---|
| ENSAPLG00000011298 | -0.5957 | 0.003695 | 0.015176 |
| TMEM230 | 0.550244 | 0.003702 | 0.0152 |
| ENSAPLG00000003871 | -1.23215 | 0.00371 | 0.01522 |
| CPD | -0.82136 | 0.003709 | 0.01522 |
| DPY19L3 | -0.61488 | 0.00372 | 0.015256 |
| CPSF3 | -0.48516 | 0.003726 | 0.015276 |
| ENSAPLG00000029307 | 1.078498 | 0.003768 | 0.015442 |
| CADM1 | -1.29603 | 0.003786 | 0.015511 |
| MEX3B | 0.534767 | 0.003793 | 0.015532 |
| HYAL2 | -0.76103 | 0.003796 | 0.01554 |
| ZCCHC17 | 0.626277 | 0.003806 | 0.015575 |
| PDE9A | -0.74471 | 0.003826 | 0.015651 |
| ZNF622 | -0.50015 | 0.003832 | 0.015667 |
| ENSAPLG00000027094 | -0.6508 | 0.003842 | 0.015703 |
| ENSAPLG00000009678 | 1.498971 | 0.003848 | 0.015723 |
| SNX14 | -0.50926 | 0.00385 | 0.015724 |
| XXYLT1 | 0.504387 | 0.003868 | 0.015791 |
| ELOVL6 | -0.72884 | 0.003874 | 0.015813 |
| GRHPR | -0.91548 | 0.003877 | 0.015816 |
| TGFBR2 | 0.605368 | 0.003882 | 0.015831 |
| CDC42SE2 | 0.592136 | 0.00389 | 0.015858 |
| CDCA7L | -0.48982 | 0.0039 | 0.015895 |
| SURF4 | -0.48992 | 0.003905 | 0.01591 |
| RBM24 | 0.670229 | 0.003909 | 0.015919 |
| C17orf75 | -0.75404 | 0.003913 | 0.015926 |
| RIOK3 | -0.47332 | 0.00392 | 0.015949 |
| VRK2 | -0.68178 | 0.003935 | 0.016004 |
| RTKN | 1.025237 | 0.003941 | 0.016026 |
| ITPK1 | -0.68606 | 0.003944 | 0.016031 |
| CARNMT1 | 0.496286 | 0.003946 | 0.016031 |
| UNC119 | 0.71606 | 0.003953 | 0.016057 |
| ARPC1B | -0.57916 | 0.003965 | 0.0161 |
| ZBTB33 | 0.838685 | 0.003969 | 0.016109 |
| ENSAPLG00000006877 | -0.75707 | 0.003972 | 0.016115 |
| STOM | -0.74063 | 0.003977 | 0.016123 |
| ATP9A | 1.02924 | 0.003977 | 0.016123 |
| WNK2 | 0.868012 | 0.003997 | 0.016198 |
| SOS1 | 0.768776 | 0.004004 | 0.01622 |
| FAM241A | 0.963443 | 0.004006 | 0.016222 |
| CDC37L1 | -0.63657 | 0.004014 | 0.016249 |
| BMPR2 | 0.758379 | 0.004022 | 0.016264 |
| PPP1CA | -0.56541 | 0.004022 | 0.016264 |
| LMOD1 | -0.7551 | 0.004023 | 0.016264 |
| SZRD1 | 1.245902 | 0.004024 | 0.016264 |
| ENSAPLG00000019645 | -1.01799 | 0.004036 | 0.016307 |

| | | | |
|---|---|---|---|
| GALNT2 | -1.87244 | 0.004077 | 0.016465 |
| ITM2C | 3.001709 | 0.00408 | 0.016471 |
| ENSAPLG00000000604 | 0.769444 | 0.004096 | 0.016533 |
| METAP2 | -0.47826 | 0.004125 | 0.016643 |
| IMPA1 | -0.54221 | 0.004133 | 0.01666 |
| SRM | -0.4651 | 0.004131 | 0.01666 |
| FGFR3 | 0.694702 | 0.004134 | 0.01666 |
| SLC4A2 | 0.714005 | 0.004144 | 0.016695 |
| SNX2 | -0.54526 | 0.004147 | 0.0167 |
| ENSAPLG00000008759 | -0.62697 | 0.004156 | 0.016718 |
| PARK7 | -0.56107 | 0.004156 | 0.016718 |
| BNIP1 | -0.64251 | 0.004154 | 0.016718 |
| FAM114A2 | -0.54561 | 0.004167 | 0.016754 |
| ENSAPLG00000010147 | -0.54755 | 0.004172 | 0.016768 |
| RAB40C | 0.837953 | 0.004174 | 0.016773 |
| ENSAPLG00000020237 | 4.69467 | 0.004181 | 0.016793 |
| ICE2 | 0.887117 | 0.004187 | 0.016809 |
| WASHC4 | -0.66648 | 0.004188 | 0.016809 |
| TENM4 | 4.732299 | 0.004195 | 0.01683 |
| RAB12 | 0.473228 | 0.004196 | 0.01683 |
| OXSM | -0.76079 | 0.0042 | 0.016839 |
| EFHD2 | -0.73955 | 0.004207 | 0.016859 |
| ASB3 | -0.70572 | 0.004208 | 0.01686 |
| TRAPPC12 | -0.54941 | 0.004212 | 0.016867 |
| GHSR | 4.114787 | 0.004217 | 0.01688 |
| SHQ1 | -0.53923 | 0.004226 | 0.016911 |
| ENSAPLG00000002508 | 0.939661 | 0.00423 | 0.016922 |
| PDXK | 0.646784 | 0.004243 | 0.016966 |
| GLCCI1 | 0.815557 | 0.004259 | 0.017025 |
| PLIN2 | -0.46746 | 0.004273 | 0.017076 |
| MBTPS2 | -0.81783 | 0.004276 | 0.01708 |
| ENSAPLG00000023537 | 0.566626 | 0.004286 | 0.017116 |
| PKP1 | 1.993923 | 0.00429 | 0.017123 |
| LDHA | -0.56473 | 0.004293 | 0.017128 |
| WDR11 | -0.75708 | 0.004297 | 0.017137 |
| GIN1 | -0.91747 | 0.004324 | 0.017241 |
| GNS | -0.75522 | 0.004332 | 0.017266 |
| LPCAT1 | -0.57659 | 0.004335 | 0.017271 |
| RGS4 | 2.755801 | 0.004338 | 0.017279 |
| RPL8 | -0.53627 | 0.004343 | 0.017288 |
| DCUN1D2 | -0.73305 | 0.004344 | 0.017288 |
| USP34 | 0.574723 | 0.004359 | 0.017341 |
| RCAN2 | 0.852462 | 0.004361 | 0.017344 |
| RTF1 | -0.57796 | 0.004365 | 0.017355 |
| SOD2 | -0.52126 | 0.004392 | 0.017456 |

| | | | |
|---|---|---|---|
| NONO | -0.43247 | 0.004398 | 0.017473 |
| SOSTDC1 | 4.789843 | 0.004402 | 0.017482 |
| PDLIM5 | -0.47359 | 0.004412 | 0.017511 |
| ZNF503 | 0.818881 | 0.004413 | 0.017511 |
| SDE2 | -0.58922 | 0.004415 | 0.017513 |
| HIC2 | 0.674909 | 0.004422 | 0.017536 |
| PLEK2 | 0.587854 | 0.004432 | 0.017569 |
| MREG | 0.828499 | 0.004441 | 0.017599 |
| USH2A | 2.773111 | 0.004447 | 0.017616 |
| ENSAPLG00000011659 | 0.632327 | 0.00445 | 0.017622 |
| NEK3 | -1.05819 | 0.004454 | 0.017626 |
| ENSAPLG00000022853 | 0.661739 | 0.004455 | 0.017626 |
| ENSAPLG00000008521 | -0.49083 | 0.004463 | 0.017653 |
| FAM118B | -0.50208 | 0.004468 | 0.017658 |
| YIPF4 | 0.453172 | 0.004466 | 0.017658 |
| ENSAPLG00000002443 | -0.94217 | 0.004471 | 0.017665 |
| SLC27A4 | -0.54252 | 0.004474 | 0.017672 |
| PKNOX1 | 0.499291 | 0.00449 | 0.017726 |
| APPL1 | -0.64966 | 0.004495 | 0.017735 |
| MMP16 | -0.61875 | 0.004494 | 0.017735 |
| FOXK1 | 0.592648 | 0.004519 | 0.017822 |
| CAPN10 | -0.65885 | 0.004525 | 0.01784 |
| WDR41 | -0.75859 | 0.004527 | 0.017841 |
| ADAT1 | 0.873392 | 0.004534 | 0.01786 |
| ENSAPLG00000003614 | 0.625451 | 0.004544 | 0.017883 |
| DUSP7 | 0.523664 | 0.004542 | 0.017883 |
| C3orf14 | 0.666123 | 0.004544 | 0.017883 |
| ENSAPLG00000005509 | 1.479408 | 0.004547 | 0.017887 |
| ENSAPLG00000002748 | 1.41789 | 0.004549 | 0.017889 |
| TLN1 | -0.58153 | 0.004556 | 0.017909 |
| GTF3C5 | -0.66211 | 0.004572 | 0.017967 |
| CEP19 | 0.68807 | 0.004576 | 0.017977 |
| DDX24 | -0.66063 | 0.00461 | 0.018105 |
| RPL27A | 0.551554 | 0.004618 | 0.01812 |
| YES1 | 0.57676 | 0.004618 | 0.01812 |
| NEURL1 | 0.924055 | 0.004624 | 0.018138 |
| ENSAPLG00000013325 | 0.79392 | 0.004626 | 0.018139 |
| ENSAPLG00000027780 | 0.698447 | 0.004635 | 0.018166 |
| FARSB | -0.47224 | 0.004642 | 0.01817 |
| SALL3 | 1.340981 | 0.004643 | 0.01817 |
| ENSAPLG00000013932 | -0.81439 | 0.004644 | 0.01817 |
| ENSAPLG00000021679 | 0.678451 | 0.004637 | 0.01817 |
| EIF4EBP2 | 0.951809 | 0.004641 | 0.01817 |
| DKK3 | -0.62055 | 0.004654 | 0.018203 |
| ENSAPLG00000017564 | -1.13659 | 0.004656 | 0.018203 |

| | | | |
|---|---|---|---|
| TANC1 | 0.673635 | 0.004666 | 0.018235 |
| CDR2 | 0.514114 | 0.004706 | 0.018381 |
| TXK | 4.626811 | 0.004706 | 0.018381 |
| HPSE | -3.97675 | 0.004718 | 0.018416 |
| CLDN5 | 0.601873 | 0.004718 | 0.018416 |
| GPR107 | -0.47997 | 0.004725 | 0.018433 |
| GALNT5 | 0.871832 | 0.004726 | 0.018433 |
| RABL3 | -0.60127 | 0.004729 | 0.018436 |
| ENSAPLG00000030207 | 0.693256 | 0.00474 | 0.018475 |
| FAM53B | 1.567656 | 0.004759 | 0.018542 |
| UST | 0.573864 | 0.004763 | 0.01855 |
| STRN | -0.55926 | 0.004776 | 0.018595 |
| ENSAPLG00000021321 | -0.51812 | 0.004782 | 0.01861 |
| ENSAPLG00000007746 | -0.44096 | 0.004784 | 0.018612 |
| PIGC | -0.63859 | 0.00479 | 0.018628 |
| HSF2 | -0.45291 | 0.004792 | 0.018628 |
| MTREX | -0.50944 | 0.004794 | 0.018628 |
| NPR3 | -1.35525 | 0.004795 | 0.018628 |
| IPMK | 0.626636 | 0.004809 | 0.018664 |
| DUSP4 | 0.696557 | 0.004809 | 0.018664 |
| RBM22 | -0.63538 | 0.004808 | 0.018664 |
| HDGFL3 | 0.728686 | 0.004816 | 0.018681 |
| HSPBAP1 | -0.71521 | 0.004826 | 0.018713 |
| SEC14L5 | 1.039225 | 0.004829 | 0.01872 |
| ENSAPLG00000022070 | 0.655581 | 0.004849 | 0.018792 |
| CDK5RAP2 | 0.615168 | 0.004854 | 0.018805 |
| TNRC6B | 0.756059 | 0.004865 | 0.018837 |
| ENSAPLG00000019878 | -0.66611 | 0.004879 | 0.018886 |
| CLTCL1 | -0.71313 | 0.0049 | 0.01896 |
| MT4 | 0.735518 | 0.004915 | 0.019013 |
| SRP54 | -0.45717 | 0.004923 | 0.019036 |
| ENSAPLG00000011790 | -0.49221 | 0.004928 | 0.019049 |
| ENSAPLG00000001741 | -0.5569 | 0.004935 | 0.01907 |
| ENSAPLG00000008180 | 0.618867 | 0.004941 | 0.019085 |
| KIF21A | 1.036219 | 0.004944 | 0.019092 |
| LIPT1 | -0.5749 | 0.004949 | 0.019096 |
| MATK | 1.314489 | 0.004947 | 0.019096 |
| ENSAPLG00000020795 | -0.73941 | 0.004959 | 0.019129 |
| BCL2L13 | 0.596381 | 0.00498 | 0.0192 |
| KLB | 4.602653 | 0.004983 | 0.0192 |
| ENSAPLG00000022197 | 4.603744 | 0.004982 | 0.0192 |
| ENSAPLG00000013772 | -0.63371 | 0.005012 | 0.019305 |
| CEP41 | -0.97243 | 0.005016 | 0.019314 |
| TPI1 | -0.5222 | 0.005032 | 0.019364 |
| DDX1 | -0.43892 | 0.005033 | 0.019364 |

| | | | |
|---|---|---|---|
| ITGA8 | -0.76029 | 0.005041 | 0.019388 |
| CNGA3 | 4.59686 | 0.005049 | 0.019415 |
| CCSER2 | 0.525383 | 0.005079 | 0.019521 |
| ENSAPLG00000023658 | 2.777828 | 0.005085 | 0.019539 |
| GJD4 | 4.588333 | 0.005116 | 0.01965 |
| CFAP58 | 4.594027 | 0.005119 | 0.019655 |
| MTX2 | -0.5103 | 0.005127 | 0.019671 |
| UBASH3B | 0.881306 | 0.005125 | 0.019671 |
| SLC26A2 | -1.27778 | 0.005146 | 0.019737 |
| ADAL | -0.63023 | 0.005172 | 0.019831 |
| PCARE | 4.591638 | 0.005174 | 0.019832 |
| TM9SF4 | -0.48604 | 0.00518 | 0.019846 |
| ATPAF1 | -0.7327 | 0.005196 | 0.0199 |
| MDN1 | 0.59067 | 0.005218 | 0.019978 |
| CREG1 | -0.78427 | 0.005225 | 0.01999 |
| SH3BGRL | 0.616094 | 0.005225 | 0.01999 |
| THAP1 | 1.169779 | 0.005227 | 0.019993 |
| ENSAPLG00000030815 | 1.026519 | 0.005242 | 0.020044 |
| FIGNL2 | 1.752545 | 0.005247 | 0.020053 |
| ENSAPLG00000004813 | 0.632842 | 0.00525 | 0.02006 |
| SLC9B2 | -0.90759 | 0.005252 | 0.02006 |
| PUS7 | -0.48226 | 0.005271 | 0.020124 |
| ENSAPLG00000031181 | -1.57003 | 0.005277 | 0.020141 |
| LAPTM4A | -0.52254 | 0.005282 | 0.020155 |
| HDAC1 | -0.45315 | 0.005298 | 0.020208 |
| VPS39 | -0.79133 | 0.005307 | 0.020234 |
| MCOLN3 | 0.836736 | 0.00532 | 0.020269 |
| LSS | -0.50561 | 0.005318 | 0.020269 |
| CERS1 | 1.325944 | 0.00533 | 0.020302 |
| ENSAPLG00000008971 | -0.57025 | 0.005334 | 0.02031 |
| ESRRB | 2.606526 | 0.00535 | 0.020362 |
| ACP6 | -0.64476 | 0.005364 | 0.020407 |
| CRYL1 | -1.03903 | 0.005381 | 0.020464 |
| ENSAPLG00000017182 | 0.512997 | 0.005383 | 0.020467 |
| GATA6 | 0.509716 | 0.00539 | 0.020485 |
| ENSAPLG00000007524 | 0.481117 | 0.005392 | 0.020485 |
| TMC6 | -0.9651 | 0.005395 | 0.02049 |
| PPP1R21 | -0.59474 | 0.00541 | 0.020539 |
| SPIRE1 | 0.588222 | 0.005424 | 0.020586 |
| PARG | 0.642396 | 0.005449 | 0.020676 |
| CPEB2 | 0.923522 | 0.005452 | 0.02068 |
| FRRS1 | -0.71929 | 0.005465 | 0.020719 |
| CCNY | 1.318655 | 0.005476 | 0.020755 |
| SETD6 | -0.48235 | 0.005495 | 0.020819 |
| TRPS1 | 0.547528 | 0.005509 | 0.020858 |

| | | | |
|---|---|---|---|
| ENSAPLG00000028687 | 0.758493 | 0.005508 | 0.020858 |
| ENSAPLG00000013301 | -0.55936 | 0.005519 | 0.020889 |
| XPO6 | 0.670008 | 0.00553 | 0.020924 |
| AMPD3 | -0.63033 | 0.005534 | 0.02093 |
| TBC1D4 | -0.72682 | 0.005542 | 0.020934 |
| INPP4A | 0.843816 | 0.005539 | 0.020934 |
| RIPK1 | -0.63236 | 0.005542 | 0.020934 |
| ENSAPLG00000017972 | 2.273937 | 0.005542 | 0.020934 |
| ENSAPLG00000018261 | 2.755281 | 0.005549 | 0.020951 |
| ELOVL5 | -0.4473 | 0.005553 | 0.020958 |
| RNASEH2B | -0.56218 | 0.005564 | 0.020993 |
| LZIC | 0.48969 | 0.005579 | 0.021042 |
| ENSAPLG00000015141 | 0.826148 | 0.005585 | 0.021057 |
| ENSAPLG00000030677 | -0.75365 | 0.005598 | 0.021101 |
| CASP8AP2 | 0.695666 | 0.005613 | 0.021148 |
| NYX | 2.223672 | 0.005626 | 0.021184 |
| ENSAPLG00000027307 | 0.511974 | 0.005626 | 0.021184 |
| S100Z | 3.001441 | 0.005641 | 0.021234 |
| TBC1D12 | 0.636732 | 0.005643 | 0.021234 |
| ENSAPLG00000006338 | 0.454123 | 0.005658 | 0.021275 |
| GET1 | -0.50235 | 0.005657 | 0.021275 |
| PHLDA1 | 0.8466 | 0.00566 | 0.021276 |
| CEP43 | -0.62536 | 0.005679 | 0.021339 |
| ENSAPLG00000012071 | 0.846591 | 0.005685 | 0.021346 |
| ENSAPLG00000016444 | 3.227792 | 0.005685 | 0.021346 |
| ENSAPLG00000013847 | 0.576897 | 0.005692 | 0.021367 |
| PTK7 | -0.67183 | 0.005705 | 0.021408 |
| OAZ1 | 0.618747 | 0.00571 | 0.021417 |
| ARMC9 | -0.83077 | 0.00572 | 0.021449 |
| NUP62 | -0.58304 | 0.00573 | 0.02148 |
| MLST8 | 0.467459 | 0.005739 | 0.021504 |
| GNAS | 0.55921 | 0.005751 | 0.021542 |
| TECPR2 | -0.55508 | 0.005776 | 0.021629 |
| GNG11 | 1.337778 | 0.005781 | 0.02164 |
| MAPKAPK3 | 1.073294 | 0.005792 | 0.021673 |
| JMJD1C | 0.52482 | 0.005824 | 0.021787 |
| ZBTB41 | 0.558626 | 0.005826 | 0.021787 |
| KCNC1 | 4.517428 | 0.005839 | 0.021822 |
| MLYCD | -1.29141 | 0.00584 | 0.021822 |
| EFNB2 | 0.605585 | 0.005848 | 0.021847 |
| FAM110B | -0.65952 | 0.005889 | 0.021993 |
| ARIH2 | -0.46045 | 0.005903 | 0.022035 |
| C1QTNF5 | -1.3514 | 0.005924 | 0.022108 |
| GNA13 | 0.535312 | 0.005934 | 0.022135 |
| RNF149 | -0.47341 | 0.005936 | 0.022138 |

| | | | |
|---|---|---|---|
| PIGS | -1.0047 | 0.005948 | 0.022174 |
| ITM2A | -0.47071 | 0.005956 | 0.022194 |
| NID2 | -0.95938 | 0.005957 | 0.022194 |
| SLC8A3 | 1.957357 | 0.00597 | 0.022235 |
| SESN1 | -0.56955 | 0.005981 | 0.022266 |
| CTPS2 | -0.81013 | 0.005989 | 0.022288 |
| LRRN2 | 4.740387 | 0.005995 | 0.022303 |
| ENDOV | -0.82217 | 0.006014 | 0.022367 |
| ATF2 | 0.459332 | 0.006035 | 0.022435 |
| BCL11B | 0.806708 | 0.006047 | 0.022473 |
| ENSAPLG00000014048 | -0.52707 | 0.006052 | 0.022486 |
| KLF5 | 0.477344 | 0.006063 | 0.022518 |
| ENSAPLG00000000603 | 0.592456 | 0.006071 | 0.022539 |
| FECH | -0.5717 | 0.006089 | 0.022598 |
| RBM5 | 0.462456 | 0.00612 | 0.022708 |
| BTBD7 | 0.705567 | 0.006136 | 0.022756 |
| DEGS1 | -0.48754 | 0.006143 | 0.022776 |
| ITPA | -0.67111 | 0.006151 | 0.022798 |
| C6orf120 | 0.511641 | 0.006173 | 0.02287 |
| KIFC3 | 0.683053 | 0.006178 | 0.022876 |
| TIPRL | -0.45364 | 0.006177 | 0.022876 |
| PUM1 | 0.566511 | 0.006195 | 0.022931 |
| ENSAPLG00000011608 | 0.865532 | 0.006211 | 0.022983 |
| ENSAPLG00000026440 | 0.555616 | 0.006216 | 0.02299 |
| GMNN | -0.80926 | 0.006228 | 0.023027 |
| SMPD3 | 3.469842 | 0.006236 | 0.023048 |
| HIBADH | -0.62295 | 0.006244 | 0.02307 |
| OSR2 | 1.023223 | 0.006266 | 0.023144 |
| GOLM1 | -0.78534 | 0.00629 | 0.023225 |
| PRRC1 | -0.49852 | 0.006316 | 0.023316 |
| IDE | -0.64136 | 0.006342 | 0.023404 |
| CDK19 | 0.640697 | 0.006361 | 0.023465 |
| INTS8 | -0.50053 | 0.006367 | 0.023478 |
| SGSH | -0.57474 | 0.006388 | 0.02354 |
| SACM1L | -0.50763 | 0.006386 | 0.02354 |
| RNF123 | -0.76692 | 0.006393 | 0.023551 |
| BMERB1 | 0.635398 | 0.006401 | 0.023573 |
| NPC1 | -0.66842 | 0.006404 | 0.023574 |
| YME1L1 | -0.60364 | 0.006409 | 0.023585 |
| ENSAPLG00000004797 | 0.473065 | 0.006425 | 0.023637 |
| LTBP1 | -0.6583 | 0.006431 | 0.023652 |
| DUSP19 | 0.53294 | 0.006434 | 0.023653 |
| TMCO1 | -0.54576 | 0.006456 | 0.023725 |
| EGFR | -0.52038 | 0.006495 | 0.023862 |
| ENSAPLG00000021637 | 1.050208 | 0.006499 | 0.023868 |

| | | | |
|---|---|---|---|
| RSPH1 | 2.221724 | 0.006505 | 0.023881 |
| AOPEP | 0.600316 | 0.006513 | 0.023886 |
| RIMKLB | -0.70119 | 0.00651 | 0.023886 |
| UHRF1 | -0.45693 | 0.006512 | 0.023886 |
| AAK1 | 0.698888 | 0.006529 | 0.023932 |
| MANF | -0.65587 | 0.006529 | 0.023932 |
| ENSAPLG00000010941 | 2.358655 | 0.006565 | 0.024056 |
| SYNGR3 | 2.16861 | 0.006582 | 0.024108 |
| TBL3 | -0.50117 | 0.006601 | 0.024169 |
| CCN4 | -0.66974 | 0.006622 | 0.024239 |
| MOCS2 | -0.68841 | 0.00663 | 0.024258 |
| ZFAND4 | 0.534403 | 0.006657 | 0.024343 |
| TMEM144 | -0.54947 | 0.006656 | 0.024343 |
| ZNF800 | 1.140592 | 0.006687 | 0.024445 |
| MRPL13 | 0.528352 | 0.006694 | 0.024452 |
| RUSC2 | 0.910057 | 0.006692 | 0.024452 |
| PEX6 | -1.293 | 0.006746 | 0.024633 |
| MORN2 | 1.539515 | 0.006749 | 0.024637 |
| AP4B1 | -0.60961 | 0.006764 | 0.024685 |
| RFC3 | -0.62451 | 0.006771 | 0.024701 |
| ENSAPLG00000003863 | -2.04368 | 0.006781 | 0.024729 |
| ENSAPLG00000026309 | -1.02773 | 0.006789 | 0.024752 |
| ASXL2 | 0.616856 | 0.006797 | 0.024772 |
| BTG1 | -0.53407 | 0.006817 | 0.024836 |
| CORIN | 3.620217 | 0.00682 | 0.024837 |
| PHKA2 | -0.77107 | 0.006827 | 0.024857 |
| UBXN2A | 0.550658 | 0.00684 | 0.024895 |
| NET1 | -0.50404 | 0.006853 | 0.024932 |
| ENSAPLG00000006872 | -0.5346 | 0.006861 | 0.024954 |
| SSTR3 | 1.822718 | 0.006868 | 0.024972 |
| SLC35E2B | 0.537049 | 0.006893 | 0.025046 |
| OSBPL3 | 0.583691 | 0.006893 | 0.025046 |
| ENSAPLG00000012085 | -0.46088 | 0.006918 | 0.025123 |
| EIF3L | -0.45151 | 0.006919 | 0.025123 |
| TAF5 | 0.539336 | 0.00695 | 0.025226 |
| CLCC1 | -0.74129 | 0.006955 | 0.025237 |
| ENSAPLG00000011792 | 0.46173 | 0.006988 | 0.02535 |
| SRP72 | -0.45541 | 0.006992 | 0.025356 |
| ATG4B | -0.49274 | 0.006997 | 0.025363 |
| ENSAPLG00000027201 | 3.961264 | 0.007012 | 0.025411 |
| ADPRM | -0.86137 | 0.007068 | 0.025605 |
| PANK2 | 0.544697 | 0.007073 | 0.025613 |
| MGST1 | -0.70983 | 0.007075 | 0.025613 |
| ENSAPLG00000012655 | -0.52167 | 0.007082 | 0.025629 |
| TCEA1 | -0.54398 | 0.007131 | 0.025789 |

| | | | |
|---|---|---|---|
| DGKZ | 0.916378 | 0.007129 | 0.025789 |
| TRAPPC6B | -0.64196 | 0.007134 | 0.025791 |
| IL18RAP | 2.007391 | 0.007145 | 0.025824 |
| CHN1 | 0.539943 | 0.007155 | 0.025843 |
| ASH1L | 0.542228 | 0.007153 | 0.025843 |
| LONP1 | -0.53748 | 0.00716 | 0.02585 |
| EIF1B | 0.516846 | 0.007172 | 0.025872 |
| ENSAPLG00000005550 | 0.596498 | 0.007173 | 0.025872 |
| CLSTN3 | -0.61814 | 0.00717 | 0.025872 |
| ENSAPLG00000002657 | 0.454571 | 0.00718 | 0.025891 |
| ENSAPLG00000019377 | 0.96565 | 0.007189 | 0.025913 |
| NAGLU | -0.73083 | 0.007224 | 0.026031 |
| PCSK6 | -0.90169 | 0.007248 | 0.02611 |
| ENSAPLG00000025986 | 0.843396 | 0.00726 | 0.026145 |
| ENSAPLG00000009253 | 0.72587 | 0.007265 | 0.026154 |
| ENSAPLG00000028746 | 1.369989 | 0.007274 | 0.026176 |
| ATP9B | -0.49244 | 0.007282 | 0.026196 |
| TYW3 | -0.82206 | 0.007284 | 0.026196 |
| ENSAPLG00000004283 | -0.56277 | 0.007311 | 0.026283 |
| HPRT1 | 0.492816 | 0.007349 | 0.026411 |
| SRGAP3 | 0.679042 | 0.007358 | 0.026437 |
| NUP85 | -0.48297 | 0.0074 | 0.026576 |
| ETV5 | -0.61429 | 0.007405 | 0.026585 |
| EVC2 | -0.76403 | 0.007412 | 0.026585 |
| ENSAPLG00000006235 | -0.48613 | 0.007411 | 0.026585 |
| STARD3 | 0.775756 | 0.007409 | 0.026585 |
| SAP30 | 0.587011 | 0.007421 | 0.0266 |
| SLC25A20 | 0.459388 | 0.007421 | 0.0266 |
| NFKB1 | -0.62364 | 0.007453 | 0.026705 |
| SLC44A3 | -0.74652 | 0.007466 | 0.026744 |
| S100A4 | 0.556789 | 0.007473 | 0.026754 |
| IL1RAP | -0.52155 | 0.007474 | 0.026754 |
| NOLC1 | -0.53241 | 0.007478 | 0.026761 |
| NOL11 | -0.45315 | 0.007486 | 0.02678 |
| SHMT1 | -0.96602 | 0.007493 | 0.026786 |
| MYO9B | 0.627373 | 0.00749 | 0.026786 |
| ENSAPLG00000006408 | 0.497456 | 0.00752 | 0.026865 |
| ENSAPLG00000019239 | -1.27003 | 0.007519 | 0.026865 |
| SEMA7A | 1.139929 | 0.007524 | 0.026868 |
| ENSAPLG00000017897 | 0.730286 | 0.007525 | 0.026868 |
| EYA2 | -1.21251 | 0.007543 | 0.026908 |
| SMG7 | 0.694134 | 0.007544 | 0.026908 |
| ENSAPLG00000026321 | -1.06651 | 0.00754 | 0.026908 |
| CDC5L | -0.50748 | 0.00757 | 0.026993 |
| TMEM132C | -0.77405 | 0.007591 | 0.02705 |

| | | | |
|---|---|---|---|
| ENSAPLG00000012475 | -0.64555 | 0.007589 | 0.02705 |
| TST | -0.63779 | 0.007657 | 0.027275 |
| MPPE1 | -0.61998 | 0.007662 | 0.027283 |
| RAB9B | 0.68371 | 0.007709 | 0.027444 |
| CUX1 | 0.527093 | 0.007723 | 0.027475 |
| NOL10 | -0.47587 | 0.007723 | 0.027475 |
| DLAT | -0.5118 | 0.007742 | 0.027534 |
| CYBC1 | -0.86499 | 0.007746 | 0.027538 |
| PIBF1 | -0.68308 | 0.007755 | 0.027563 |
| TRIM23 | 0.835118 | 0.00777 | 0.027604 |
| ABCD3 | -0.54486 | 0.007782 | 0.027639 |
| FAM207A | -0.59176 | 0.007787 | 0.027647 |
| SORCS2 | 4.556562 | 0.007805 | 0.027703 |
| C11orf54 | -0.64725 | 0.007817 | 0.027736 |
| REXO4 | -0.59671 | 0.00783 | 0.027775 |
| TMEM167A | 0.733286 | 0.007849 | 0.027831 |
| TAF8 | -0.54765 | 0.007863 | 0.027872 |
| MN1 | 2.276217 | 0.007906 | 0.028015 |
| KIF23 | -1.27094 | 0.007924 | 0.028072 |
| TBK1 | -0.63897 | 0.007969 | 0.028222 |
| LARP4B | 0.637661 | 0.00802 | 0.028384 |
| ENSAPLG00000017636 | -0.59389 | 0.00802 | 0.028384 |
| ENSAPLG00000022082 | -0.51528 | 0.008054 | 0.028495 |
| TTC7A | -0.65492 | 0.0081 | 0.028648 |
| ZUP1 | -0.59288 | 0.00811 | 0.028674 |
| GRIK1 | 0.814168 | 0.00814 | 0.028759 |
| ENSAPLG00000026608 | 0.807741 | 0.008138 | 0.028759 |
| ACSL4 | -0.46993 | 0.008149 | 0.028784 |
| RASL10B | 3.621084 | 0.008154 | 0.028792 |
| NRF1 | 0.713194 | 0.008183 | 0.028885 |
| NELFB | -0.43703 | 0.008198 | 0.028927 |
| ACSF3 | -1.96183 | 0.008215 | 0.02898 |
| MTFP1 | 0.83366 | 0.008264 | 0.029142 |
| JDP2 | 0.640379 | 0.008314 | 0.029309 |
| COBL | -0.7163 | 0.008321 | 0.029324 |
| BRI3 | 0.631158 | 0.008328 | 0.02934 |
| SLK | 0.503373 | 0.008335 | 0.029355 |
| SMARCD1 | -0.54326 | 0.008341 | 0.029366 |
| DCP1B | -0.5338 | 0.008347 | 0.029376 |
| GINM1 | -0.55817 | 0.008388 | 0.029513 |
| ENSAPLG00000019197 | 0.656333 | 0.008397 | 0.029535 |
| FLT4 | 1.062901 | 0.008427 | 0.029621 |
| ANKRD46 | -0.51595 | 0.008425 | 0.029621 |
| PES1 | -0.41889 | 0.008486 | 0.029819 |
| EFCAB14 | 0.583905 | 0.008502 | 0.029865 |

| | | | |
|---|---|---|---|
| CTR9 | -0.4877 | 0.00851 | 0.029885 |
| ENSAPLG00000014082 | -0.48504 | 0.008515 | 0.02989 |
| DNAJB4 | 0.410269 | 0.008529 | 0.029912 |
| ENSAPLG00000014428 | 1.908827 | 0.008524 | 0.029912 |
| ENSAPLG00000020023 | 0.810718 | 0.008528 | 0.029912 |
| SNAPC3 | -0.81008 | 0.008534 | 0.029919 |
| LYAR | -0.57615 | 0.00854 | 0.029932 |
| PIGK | -0.46551 | 0.008554 | 0.029972 |
| NCOA6 | 0.487049 | 0.008563 | 0.029993 |
| QSOX1 | -0.67991 | 0.008572 | 0.030015 |
| SFRP2 | -0.76274 | 0.008598 | 0.030095 |
| WWC1 | 0.536092 | 0.008621 | 0.030167 |
| CTHRC1 | -0.69826 | 0.008633 | 0.030198 |
| ECHDC1 | -1.6273 | 0.008637 | 0.030204 |
| GEMIN8 | -0.7255 | 0.008641 | 0.030208 |
| PDIK1L | 0.601602 | 0.008652 | 0.030235 |
| SLC39A8 | -0.89056 | 0.008667 | 0.030279 |
| GCH1 | 0.804167 | 0.008672 | 0.030286 |
| IBTK | 0.466725 | 0.00868 | 0.030305 |
| SEPTIN8 | -0.48405 | 0.008692 | 0.030336 |
| DUT | -0.81907 | 0.008702 | 0.030358 |
| PSMG4 | 0.738842 | 0.008704 | 0.030358 |
| SEPSECS | -0.64573 | 0.008716 | 0.030391 |
| MACROH2A2 | -0.58562 | 0.00872 | 0.030396 |
| FBXL2 | 0.751853 | 0.008731 | 0.030426 |
| ENSAPLG00000006105 | -0.63073 | 0.00875 | 0.03048 |
| NEDD4L | 0.614017 | 0.008752 | 0.03048 |
| ENSAPLG00000008574 | -0.74124 | 0.008782 | 0.030571 |
| ECI2 | -0.67929 | 0.008793 | 0.0306 |
| QRSL1 | -0.52497 | 0.008835 | 0.03072 |
| CBLB | 0.534622 | 0.008836 | 0.03072 |
| PTPDC1 | -0.85003 | 0.008838 | 0.03072 |
| MINDY1 | 0.600767 | 0.008838 | 0.03072 |
| NAF1 | -0.70154 | 0.008844 | 0.030731 |
| IPO13 | -0.78773 | 0.008877 | 0.030835 |
| ENSAPLG00000003922 | 0.70792 | 0.008899 | 0.03089 |
| CEP89 | -0.65822 | 0.008897 | 0.03089 |
| SNAPC1 | -0.54751 | 0.008929 | 0.030987 |
| IFT27 | -0.6127 | 0.008936 | 0.031001 |
| ENSAPLG00000004830 | 3.854353 | 0.008964 | 0.031086 |
| CACNA1C | 0.477296 | 0.008977 | 0.031123 |
| PSMB7 | -0.52529 | 0.009 | 0.031191 |
| PSTPIP2 | 2.75202 | 0.009009 | 0.031212 |
| ENSAPLG00000013508 | 0.586487 | 0.009016 | 0.031227 |
| CACNA1D | 0.71488 | 0.009025 | 0.031248 |

| | | | |
|---|---|---|---|
| GSR | 0.462972 | 0.009028 | 0.03125 |
| TRAPPC11 | -0.65196 | 0.009056 | 0.031335 |
| CCDC125 | 3.126889 | 0.009069 | 0.031342 |
| EPC1 | 0.554717 | 0.009062 | 0.031342 |
| ENSAPLG00000025146 | 0.537168 | 0.009066 | 0.031342 |
| ENSAPLG00000029386 | -0.76958 | 0.009067 | 0.031342 |
| PRKG2 | 1.557392 | 0.009092 | 0.031412 |
| SLX4 | 0.915861 | 0.009107 | 0.031452 |
| RBM45 | -0.43392 | 0.009133 | 0.031534 |
| DVL1 | 0.994122 | 0.009144 | 0.03156 |
| RSRC1 | -0.69902 | 0.009168 | 0.031632 |
| ZNF598 | 0.480384 | 0.009173 | 0.031639 |
| ENSAPLG00000025801 | -1.49687 | 0.009202 | 0.031731 |
| SCNN1G | 0.620893 | 0.009227 | 0.031806 |
| CNOT1 | 0.66036 | 0.009242 | 0.03185 |
| TRIM71 | 2.316297 | 0.009278 | 0.031964 |
| GOLT1B | 0.476286 | 0.009289 | 0.031991 |
| DNAJC6 | 1.411675 | 0.009292 | 0.031992 |
| TAF2 | 0.584591 | 0.009297 | 0.031997 |
| RPA2 | -0.49195 | 0.009301 | 0.032001 |
| ANGEL2 | -0.45594 | 0.009315 | 0.032039 |
| VDR | 0.851421 | 0.009323 | 0.032056 |
| SLC66A3 | -1.03578 | 0.00933 | 0.032072 |
| ENSAPLG00000008728 | 0.540547 | 0.009334 | 0.032075 |
| ENSAPLG00000015024 | 1.058888 | 0.009351 | 0.032122 |
| CASP8 | -0.44006 | 0.009375 | 0.032196 |
| CSE1L | -0.47607 | 0.009386 | 0.032224 |
| EIF3A | -0.55968 | 0.009434 | 0.032374 |
| ZFAND6 | 0.48977 | 0.009436 | 0.032374 |
| ENSAPLG00000020340 | -0.52759 | 0.009451 | 0.032415 |
| PEAK1 | 0.653899 | 0.009458 | 0.03243 |
| SERPINB8 | -0.54929 | 0.009469 | 0.032455 |
| JUP | -0.45267 | 0.009501 | 0.032547 |
| ENSAPLG00000023763 | 0.690926 | 0.009501 | 0.032547 |
| ENSAPLG00000027469 | 0.54067 | 0.009512 | 0.032573 |
| P2RY1 | 2.426466 | 0.00952 | 0.032589 |
| PSMC2 | -0.46482 | 0.009523 | 0.03259 |
| ENSAPLG00000018959 | 0.520557 | 0.009542 | 0.032644 |
| NSMCE4A | -0.59792 | 0.00958 | 0.032763 |
| MIEF1 | 1.029843 | 0.009594 | 0.032801 |
| DYNLT2 | 0.863919 | 0.009634 | 0.03293 |
| LIF | 0.708072 | 0.00964 | 0.03294 |
| ENSAPLG00000005093 | 2.497729 | 0.009698 | 0.033126 |
| MAD1L1 | -0.57132 | 0.009734 | 0.033229 |
| CTBS | -0.58213 | 0.009733 | 0.033229 |

| | | | |
|---|---|---|---|
| ENSAPLG00000016934 | -0.83393 | 0.009744 | 0.033253 |
| MTR | -0.89046 | 0.009784 | 0.03338 |
| WNK1 | 0.562582 | 0.009809 | 0.033454 |
| HAT1 | -0.46579 | 0.009824 | 0.033492 |
| HOXA7 | 0.784714 | 0.009839 | 0.033534 |
| ENSAPLG00000015499 | 0.527578 | 0.009857 | 0.033584 |
| C11orf49 | -0.97456 | 0.009867 | 0.033608 |
| CDC123 | -0.64618 | 0.0099 | 0.033711 |
| ENSAPLG00000009638 | 2.158497 | 0.009919 | 0.033761 |
| AP3B1 | -0.48856 | 0.009921 | 0.033761 |
| ENSAPLG00000005997 | -0.45878 | 0.009948 | 0.03384 |
| CHST3 | -0.51957 | 0.00995 | 0.03384 |
| PRKD1 | 0.472302 | 0.010055 | 0.034186 |
| PMP22 | 0.651276 | 0.01006 | 0.03419 |
| JCAD | 1.491512 | 0.010069 | 0.034212 |
| LAMA5 | 0.898054 | 0.010084 | 0.034252 |
| LAMB1 | -0.67524 | 0.010096 | 0.034281 |
| ZFYVE26 | 0.899732 | 0.01011 | 0.03432 |
| MED7 | -0.54732 | 0.01015 | 0.034444 |
| PCBD1 | 0.877804 | 0.010159 | 0.034465 |
| RBKS | 0.603896 | 0.010186 | 0.034545 |
| OTX1 | 2.779316 | 0.010226 | 0.034669 |
| KYAT3 | -0.53705 | 0.010257 | 0.034764 |
| PGAP2 | 0.568212 | 0.010291 | 0.034867 |
| TTC27 | -0.42645 | 0.010316 | 0.034941 |
| TMTC2 | -0.47904 | 0.010327 | 0.034968 |
| NDRG1 | -0.4637 | 0.010338 | 0.034994 |
| DYNC1LI1 | -0.43206 | 0.010349 | 0.03501 |
| ENSAPLG00000019153 | 1.316909 | 0.010348 | 0.03501 |
| NVL | -0.50402 | 0.01038 | 0.035103 |
| ENSAPLG00000014622 | -0.67272 | 0.010421 | 0.035231 |
| ENSAPLG00000006149 | 3.436281 | 0.01043 | 0.03525 |
| ENSAPLG00000002871 | 2.694071 | 0.010468 | 0.035362 |
| ENSAPLG00000018624 | 1.337138 | 0.010469 | 0.035362 |
| ENSAPLG00000001151 | 2.396403 | 0.010498 | 0.035448 |
| JADE3 | 0.6826 | 0.010518 | 0.035505 |
| ENSAPLG00000004420 | -0.57974 | 0.010523 | 0.035511 |
| CD81 | -0.66182 | 0.010553 | 0.035597 |
| ENSAPLG00000025387 | -0.80375 | 0.010555 | 0.035597 |
| TNFAIP8L3 | 0.645942 | 0.010581 | 0.035673 |
| NEMP1 | -0.90059 | 0.010591 | 0.035696 |
| BTBD2 | -0.94173 | 0.010605 | 0.03573 |
| GLB1L | -0.52436 | 0.010632 | 0.035811 |
| PDRG1 | 0.580546 | 0.010637 | 0.035819 |
| PTCHD4 | 3.855395 | 0.010647 | 0.035841 |

| | | | |
|---|---|---|---|
| SNX9 | -0.48161 | 0.010673 | 0.035917 |
| DOP1A | -0.52477 | 0.010681 | 0.035932 |
| RAB22A | 0.542744 | 0.010688 | 0.035946 |
| ENSAPLG00000008204 | 3.043188 | 0.010696 | 0.035961 |
| ENSAPLG00000028703 | 0.468714 | 0.01074 | 0.036098 |
| TIAL1 | -0.45844 | 0.010749 | 0.036117 |
| ENSAPLG00000020030 | 0.573422 | 0.01076 | 0.036144 |
| ENSAPLG00000006701 | -0.4851 | 0.010781 | 0.036198 |
| NUDT12 | -0.95309 | 0.010785 | 0.036198 |
| ENSAPLG00000019038 | -1.82095 | 0.01079 | 0.036198 |
| C7orf50 | -0.9428 | 0.010788 | 0.036198 |
| MTDH | -0.45339 | 0.010804 | 0.036235 |
| MGP | -0.49123 | 0.010835 | 0.036325 |
| NDUFV2 | -0.51651 | 0.010901 | 0.036536 |
| ZNF488 | 0.566624 | 0.010943 | 0.036668 |
| FGF14 | 0.739894 | 0.01097 | 0.036745 |
| ADCY9 | 0.628615 | 0.011046 | 0.036989 |
| PXDC1 | 1.242956 | 0.011072 | 0.037063 |
| ENSAPLG00000005277 | -0.90592 | 0.011075 | 0.037063 |
| DNAH17 | 0.590739 | 0.011099 | 0.037131 |
| SYT4 | 0.745063 | 0.011123 | 0.037202 |
| MAPKAPK2 | 0.720493 | 0.011127 | 0.037203 |
| TLNRD1 | 0.51224 | 0.011144 | 0.037248 |
| MAP3K14 | 0.807922 | 0.011155 | 0.037262 |
| TMEM183A | 0.471111 | 0.011155 | 0.037262 |
| THOC3 | -0.46838 | 0.01117 | 0.037301 |
| CUL5 | 0.50196 | 0.011189 | 0.037351 |
| SPPL3 | 0.467312 | 0.011228 | 0.03747 |
| PSMG3 | 0.437826 | 0.011274 | 0.037601 |
| NIPSNAP1 | -0.67454 | 0.011272 | 0.037601 |
| CNGB1 | 1.5211 | 0.011281 | 0.037615 |
| UBN1 | 0.574905 | 0.011309 | 0.037694 |
| ENSAPLG00000023268 | 0.847827 | 0.011351 | 0.037823 |
| ORC5 | -0.48433 | 0.011415 | 0.038026 |
| TAOK3 | 0.484251 | 0.011429 | 0.038059 |
| CRHBP | 2.571197 | 0.011466 | 0.038171 |
| ARRDC1 | 0.492685 | 0.011519 | 0.038337 |
| CD276 | -0.64707 | 0.011542 | 0.038403 |
| RAB39A | 0.85455 | 0.011563 | 0.038461 |
| MAPK8IP3 | 0.699172 | 0.011572 | 0.038478 |
| ARL6IP5 | 0.435315 | 0.011661 | 0.038762 |
| AGPAT5 | -0.52343 | 0.011681 | 0.038816 |
| MFAP1 | -0.54185 | 0.011693 | 0.038843 |
| ENSAPLG00000025790 | 0.571504 | 0.011712 | 0.038895 |
| ENSAPLG00000004837 | 0.747737 | 0.011744 | 0.038992 |

| | | | |
|---|---|---|---|
| ENSAPLG00000010203 | -0.60172 | 0.011827 | 0.039254 |
| EIF3D | -0.43176 | 0.011833 | 0.039262 |
| ENSAPLG00000009201 | -0.39284 | 0.011839 | 0.039271 |
| GPD1L | -0.57079 | 0.011863 | 0.039337 |
| CHAF1B | -0.65458 | 0.01187 | 0.039348 |
| MPP5 | -0.49398 | 0.011881 | 0.039373 |
| RFC2 | -0.5909 | 0.011906 | 0.039444 |
| MYO6 | -0.42242 | 0.011914 | 0.039458 |
| XRCC2 | -0.64395 | 0.011918 | 0.039459 |
| NUF2 | -0.6981 | 0.011955 | 0.03957 |
| CDCP2 | 3.243425 | 0.011965 | 0.039591 |
| ADAMTS18 | 0.957838 | 0.012056 | 0.039871 |
| ENSAPLG00000027932 | -1.93048 | 0.012057 | 0.039871 |
| ENSAPLG00000008994 | -0.64808 | 0.012081 | 0.039938 |
| ENSAPLG00000018906 | 0.473086 | 0.012105 | 0.040005 |
| CPN1 | -0.65868 | 0.01211 | 0.04001 |
| NTNG2 | 1.23247 | 0.01213 | 0.040064 |
| AP3S1 | 0.478541 | 0.01217 | 0.040185 |
| ENSAPLG00000008559 | 0.428301 | 0.01219 | 0.040231 |
| ENSAPLG00000008916 | 0.672866 | 0.012192 | 0.040231 |
| SPOPL | 0.440951 | 0.012203 | 0.040255 |
| SRPK1 | 0.430661 | 0.012221 | 0.040304 |
| ENSAPLG00000008369 | -0.43514 | 0.012254 | 0.0404 |
| TTBK2 | 0.79404 | 0.012302 | 0.040546 |
| WWTR1 | 0.886181 | 0.01231 | 0.040559 |
| PGGHG | -0.96504 | 0.012319 | 0.040579 |
| SESN2 | 1.294647 | 0.01233 | 0.040602 |
| TFG | -0.4778 | 0.012356 | 0.040675 |
| PSD | 1.181264 | 0.012362 | 0.040684 |
| CASR | 3.757896 | 0.012371 | 0.0407 |
| SRSF5 | -0.71352 | 0.012408 | 0.040808 |
| ENSAPLG00000018082 | -0.72429 | 0.012462 | 0.040976 |
| COPE | -0.43955 | 0.012474 | 0.041003 |
| ISOC1 | -0.52463 | 0.012497 | 0.041064 |
| MTCL1 | 0.529725 | 0.012516 | 0.041114 |
| CHID1 | -0.6806 | 0.012541 | 0.041184 |
| ACTN1 | -0.42933 | 0.012558 | 0.041229 |
| HSF1 | 1.022811 | 0.012587 | 0.04131 |
| RNF207 | 0.894666 | 0.012645 | 0.041489 |
| RGCC | -0.71419 | 0.012662 | 0.041531 |
| ENSAPLG00000028437 | -0.74974 | 0.012688 | 0.041604 |
| FAM160A1 | 0.578842 | 0.012696 | 0.041619 |
| ENSAPLG00000006348 | -0.70568 | 0.01271 | 0.041652 |
| ST13 | -0.40161 | 0.012732 | 0.04171 |
| MXD4 | 0.570161 | 0.012764 | 0.041804 |

| | | | |
|---|---|---|---|
| FBXO42 | 0.600444 | 0.012786 | 0.041861 |
| TPR | -0.46777 | 0.012844 | 0.042041 |
| SUPV3L1 | -0.41416 | 0.012858 | 0.042075 |
| MALRD1 | 1.058779 | 0.012898 | 0.042192 |
| MAPRE1 | -0.52807 | 0.012925 | 0.042266 |
| ENSAPLG00000026602 | 0.5259 | 0.012944 | 0.042316 |
| FAM217B | 0.530117 | 0.012955 | 0.042339 |
| CLUH | -0.44048 | 0.012984 | 0.042422 |
| RGS3 | 0.55803 | 0.012997 | 0.042452 |
| MYOD1 | 1.730273 | 0.013007 | 0.042472 |
| IRF7 | 0.770466 | 0.01304 | 0.042567 |
| ENSAPLG00000020571 | 1.266892 | 0.013048 | 0.042569 |
| ENSAPLG00000022933 | 1.802787 | 0.013049 | 0.042569 |
| ENSAPLG00000021761 | 0.934684 | 0.013065 | 0.04261 |
| CENPN | -0.94928 | 0.013072 | 0.042619 |
| ENSAPLG00000007461 | -0.59849 | 0.013089 | 0.042664 |
| IRAK4 | -0.59251 | 0.013102 | 0.042693 |
| SPATS2 | 0.459117 | 0.013128 | 0.042764 |
| ENSAPLG00000006575 | -0.61209 | 0.013148 | 0.042809 |
| AGA | -0.51882 | 0.01315 | 0.042809 |
| TMEM259 | 0.531746 | 0.013162 | 0.042837 |
| OXTR | 3.367915 | 0.013202 | 0.042953 |
| SBNO1 | 0.413566 | 0.013208 | 0.042961 |
| DCTN4 | -0.48734 | 0.013214 | 0.042968 |
| ENSAPLG00000003325 | -0.47119 | 0.013267 | 0.043123 |
| FSBP | -0.72856 | 0.01327 | 0.043123 |
| ERCC8 | 0.549864 | 0.013282 | 0.04315 |
| DNM3 | 0.99979 | 0.01329 | 0.043164 |
| KDF1 | 2.193238 | 0.013302 | 0.043177 |
| FNIP2 | -0.44648 | 0.013302 | 0.043177 |
| APMAP | -0.4441 | 0.013309 | 0.043187 |
| PDHX | -0.49278 | 0.013325 | 0.043225 |
| ENSAPLG00000018609 | -0.59774 | 0.013335 | 0.043244 |
| PLEKHA2 | 2.827645 | 0.01335 | 0.043281 |
| TMEM128 | -0.56774 | 0.013357 | 0.04329 |
| MYB | 0.475349 | 0.013363 | 0.043295 |
| DERL3 | -0.66813 | 0.013385 | 0.043356 |
| ENSAPLG00000009244 | 3.488203 | 0.013435 | 0.043503 |
| ENSAPLG00000015538 | 1.983243 | 0.013439 | 0.043503 |
| HPS5 | -0.7044 | 0.013484 | 0.043637 |
| CCNL1 | 0.519358 | 0.013547 | 0.043829 |
| SLC52A3 | -0.54395 | 0.013552 | 0.043831 |
| KLHL30 | 3.719106 | 0.013572 | 0.043881 |
| ATP6V1C1 | -0.43524 | 0.01359 | 0.043927 |
| MED9 | 0.623211 | 0.013607 | 0.043969 |

| | | | |
|---|---|---|---|
| ENSAPLG00000013548 | 0.544248 | 0.013629 | 0.044028 |
| FEN1 | -0.50838 | 0.013666 | 0.044135 |
| KAT2B | 0.478196 | 0.013679 | 0.044164 |
| LSM3 | 0.538043 | 0.013688 | 0.044177 |
| ENSAPLG00000017976 | 0.895395 | 0.013713 | 0.044247 |
| TMED3 | -0.50447 | 0.01376 | 0.044386 |
| CRADD | 0.836471 | 0.013792 | 0.044474 |
| VPS51 | -0.67676 | 0.013797 | 0.044478 |
| SNRPC | 0.687938 | 0.013865 | 0.044685 |
| ENSAPLG00000011088 | -0.72968 | 0.013989 | 0.04507 |
| MSANTD3 | -0.78526 | 0.013995 | 0.045076 |
| TMEM120A | -0.46658 | 0.014031 | 0.045178 |
| NMU | 0.48337 | 0.014058 | 0.045253 |
| ENSAPLG00000030115 | -0.50466 | 0.014129 | 0.045468 |
| SMAD7 | 0.718917 | 0.014153 | 0.045533 |
| MYH9 | -0.5905 | 0.014159 | 0.045538 |
| ENSAPLG00000010464 | 2.182404 | 0.014244 | 0.045797 |
| NR4A2 | 0.774138 | 0.014286 | 0.045918 |
| ZBTB10 | -0.46872 | 0.014302 | 0.045957 |
| PSMD14 | -0.42533 | 0.014335 | 0.046047 |
| ZW10 | -0.53077 | 0.014344 | 0.046065 |
| AVEN | -0.60916 | 0.01438 | 0.046152 |
| ZNF148 | 0.498434 | 0.014378 | 0.046152 |
| GRIN3B | 1.219564 | 0.014416 | 0.04624 |
| ABL1 | 1.257016 | 0.014413 | 0.04624 |
| IL6ST | -0.60058 | 0.014424 | 0.046253 |
| STAMBPL1 | -0.73116 | 0.014481 | 0.046422 |
| CDKAL1 | -0.47186 | 0.014516 | 0.04652 |
| STXBP3 | -0.45619 | 0.014534 | 0.046537 |
| MAN1A1 | -0.73984 | 0.014527 | 0.046537 |
| ENSAPLG00000027237 | 0.521063 | 0.014534 | 0.046537 |
| NDUFS6 | 0.626454 | 0.014585 | 0.046686 |
| VPS50 | -0.52128 | 0.014593 | 0.046699 |
| ENSAPLG00000022299 | -0.55338 | 0.014616 | 0.046757 |
| MRPS2 | 0.561065 | 0.014636 | 0.046809 |
| GPATCH3 | 0.619771 | 0.014662 | 0.046866 |
| ENSAPLG00000027519 | 0.598722 | 0.014662 | 0.046866 |
| SLC25A15 | -0.69613 | 0.0147 | 0.046968 |
| WAPL | 0.40739 | 0.014703 | 0.046968 |
| ENSAPLG00000004192 | -0.59469 | 0.014709 | 0.04697 |
| OSTM1 | -0.62675 | 0.014712 | 0.04697 |
| SIK2 | 0.753213 | 0.014719 | 0.046978 |
| ZNF711 | 0.411559 | 0.014765 | 0.047111 |
| TOR4A | 1.514839 | 0.014777 | 0.047132 |
| CSMD2 | 2.816788 | 0.014782 | 0.047132 |

| | | | |
|---|---|---|---|
| ENSAPLG00000026011 | -0.85025 | 0.014785 | 0.047132 |
| SOX6 | 1.10889 | 0.014794 | 0.047147 |
| ZBED4 | 0.562908 | 0.01481 | 0.047184 |
| STN1 | -1.36819 | 0.014834 | 0.047244 |
| BMPR1A | -0.55863 | 0.014837 | 0.047244 |
| SUN1 | -0.54388 | 0.014858 | 0.047296 |
| CYTH4 | 0.651635 | 0.014894 | 0.047398 |
| MSH3 | -0.5826 | 0.014912 | 0.04744 |
| SEC61A1 | -0.42224 | 0.015007 | 0.047729 |
| RASSF8 | 0.493366 | 0.015044 | 0.047834 |
| NFYA | -0.49838 | 0.015054 | 0.047838 |
| WDR91 | -0.44305 | 0.015054 | 0.047838 |
| DRAXIN | 2.074086 | 0.015071 | 0.047876 |
| ENSAPLG00000020498 | 0.655926 | 0.015133 | 0.048058 |
| DPY30 | 0.502086 | 0.015146 | 0.048086 |
| ENSAPLG00000004132 | -0.55667 | 0.015157 | 0.048107 |
| NOB1 | -0.44354 | 0.015185 | 0.048183 |
| CDC14A | -0.60243 | 0.0152 | 0.048216 |
| STAG2 | 0.444892 | 0.015234 | 0.048308 |
| UBN2 | 0.636536 | 0.015272 | 0.048414 |
| TRAP1 | -0.42628 | 0.015276 | 0.048414 |
| APIP | -0.48436 | 0.015296 | 0.048465 |
| ENSAPLG00000006590 | -0.44072 | 0.015312 | 0.048502 |
| ENSAPLG00000001050 | -0.69251 | 0.015347 | 0.048596 |
| DTNBP1 | -0.55416 | 0.01542 | 0.048814 |
| ENSAPLG00000002870 | -0.43973 | 0.015446 | 0.048865 |
| RAB41 | 0.683983 | 0.015449 | 0.048865 |
| MPP4 | 3.274114 | 0.015447 | 0.048865 |
| MCAT | -0.93246 | 0.015503 | 0.04902 |
| PHKA1 | 0.784205 | 0.015518 | 0.049054 |
| MLLT1 | 0.694336 | 0.01553 | 0.049078 |
| ELP2 | -0.44166 | 0.015549 | 0.049124 |
| HOMER1 | 0.578858 | 0.015567 | 0.049167 |
| DCAF7 | -0.47687 | 0.015585 | 0.049208 |
| INKA2 | -0.56667 | 0.015601 | 0.049244 |
| AKT3 | 0.579858 | 0.015634 | 0.049335 |
| SUGCT | -1.44941 | 0.015665 | 0.049403 |
| ENSAPLG00000011360 | 0.553071 | 0.015665 | 0.049403 |
| RFX3 | -0.65637 | 0.01568 | 0.049435 |
| NFATC1 | 0.621084 | 0.0157 | 0.049486 |
| DYRK1A | 0.658405 | 0.0158 | 0.049787 |
| ENSAPLG00000002087 | -0.58036 | 0.015808 | 0.049797 |
| TMEM38B | -0.8883 | 0.015827 | 0.049842 |
| ENSAPLG00000014904 | 0.591742 | 0.015845 | 0.049885 |

**Supplementary Table 9: Differentially expressed cytokine or cytokine related genes in infected avian endothelial cells**

| Cytokines | Black swan | | Chicken | | Duck | |
|---|---|---|---|---|---|---|
| **Chemokine** | **Up regulated** | **Down regulated** | **Up regulated** | **Down regulated** | **Up regulated** | **Down regulated** |
| CC subfamily | *CCL20, CXCL1,* | | *CCL4, CCL4L1, CCL4L2, CCL17, CCR5, CCL20* | | *CCR5* | |
| CXC subfamily | *CXCL8, CXCR5,* | *CXCR4* | *CXCL8, CXCL12, CXCL14, CX3CL1* | *CXCR2, CXCR4, CXCR7* | *CXCL8, CXCL12,* | |
| **The class I helical cytokines** | | | | | | |
| ɤ-chain-utilizing | *IL21R* | | *IL7, IL7R, IL9R, IL15* | | | |
| IL4-like | *CSF2RA* | | *CSF2RB, CSF2RB, CSF2,* | *IL2RB* | | *IL13RA1* |
| Prolactin family | | | *GHR* | | *GHR* | |
| IL6/12-like | *IL6, IL11RA, IL12RB2, LIF* | | *IL6, IL11RA, IL12, IL35, CLCF1, OSMR* | *IL6ST, IL12RB1* | *LIF* | *IL6ST* |

| | | | | | | |
|---|---|---|---|---|---|---|
| **The class II helical cytokines** | | | | | | |
| IL10/28-like | | | *IL10RA, IL10RB, IL20, IL20RA, IL28A, IL28B* | | | |
| Interferon family | *IFNB1* | | *IFNAR2, IFNGR2* | *IFNAR1* | | |
| **IL1-like cytokines** | *IL18R2, ST2* | | *IL1R1, IL1RAP, IL1R2, IL1RL2, IL18, IL18R1, IL18RAP* | | *IL1R1, IL1R2, IL18RAP* | |
| **IL17-like** | | | *IL17RA* | | | |
| **Non-classified** | *IL16* | | *IL34* | *IL16, CSFR1* | | |
| **TNF family** | *DR6, EDAR, CD30L, GITRL, BAFF, (TNSF8, TNFAIP2, TNSF13B, TNFSF21)* | *CD40* | *VEGI, DR4, DR5, DR6, EDA, CD30L, BAFF* | *TNFR1, DCR3, FAS, XEDAR, RANK, RANKL, OPG, CD40, BAFFR, TROY* | *DR6, CD30L, BAFF* | *TNFR1, TRAIL, NGF* |

| | | | | | |
|---|---|---|---|---|---|
| **TGF-β family** | *ACVR2A, BMPR2, BPMR1B* | *TGFB1, TGFB2, BMP10, ACVR2A, INHBB, AMH AMHR2, BMPR1B, ACVR1, BMP8,* | *TGFBR1, TGFBR2, BMPR1A, GDF9, BMP15* | *TGFBR2, GDF2, BMPR2, INHBB, NODAL, BMP15* | *TGFBR1, TGFB3, BMPR1A, BMPR1B* |

**Supplementary Table 10: 113 GO terms were significantly enriched in infected black swan endothelial cells**

| Term | Weight |
|---|---|
| cleavage involved in rRNA processing | 0.000057 |
| defense response to virus | 0.000085 |
| positive regulation of transforming | 0.00012 |
| ribonucleoprotein complex assembly | 0.00019 |
| snRNA 3'-end processing | 0.00019 |
| maturation of 5.8S rRNA from | 0.00047 |
| somitogenesis | 0.00077 |
| positive regulation of histone | 0.00088 |
| lens morphogenesis in camera-type eye | 0.00184 |
| centrosome localization | 0.00194 |
| negative regulation of MAP kinase | 0.00221 |
| ribosome assembly | 0.00245 |
| peptide hormone secretion | 0.00281 |
| maternal process involved in female | 0.00301 |
| positive regulation of cell cycle | 0.00346 |
| positive regulation of phosphorylation | 0.00353 |
| tRNA metabolic process | 0.0036 |
| positive regulation of cytoskeleton | 0.00373 |
| regulation of G2/M transition of | 0.0042 |
| social behavior | 0.00567 |
| nuclear-transcribed mRNA catabolic | 0.00605 |
| intraciliary transport | 0.00607 |
| negative regulation of intrinsic | 0.00607 |
| regulation of cytosolic calcium ion | 0.00618 |
| ribosomal large subunit biogenesis | 0.00747 |
| negative regulation of transcription by | 0.00748 |
| cytoplasmic microtubule organization | 0.00924 |
| RNA phosphodiester bond hydrolysis | 0.00978 |
| spleen development | 0.00978 |
| positive regulation of telomere | 0.00978 |
| regulation of cell population | 0.00991 |
| regulation of cell cycle | 0.01028 |
| ciliary basal body-plasma membrane | 0.01068 |
| snRNA transcription by RNA polymerase II | 0.01105 |
| negative regulation of JNK cascade | 0.01105 |
| positive regulation of cellular protein | 0.01125 |
| regulation of cellular response to heat | 0.0114 |
| positive regulation of cell population | 0.01183 |
| metaphase plate congression | 0.01233 |
| ganglioside metabolic process | 0.01239 |
| RNA 5'-end processing | 0.01239 |

| | |
|---|---|
| modulation by host of viral process | 0.01239 |
| positive regulation of protein | 0.01239 |
| protein localization to chromosome | 0.01239 |
| establishment of protein localization | 0.01239 |
| nuclear-transcribed mRNA catabolic | 0.01239 |
| ceramide catabolic process | 0.01239 |
| regulation of necroptotic process | 0.01239 |
| peptidyl-threonine dephosphorylation | 0.01239 |
| anion transmembrane transport | 0.01257 |
| membrane docking | 0.01382 |
| megakaryocyte differentiation | 0.01456 |
| regulation of JUN kinase activity | 0.01456 |
| positive regulation of establishment of | 0.01485 |
| negative regulation of microtubule | 0.01487 |
| regulation of cardiac muscle cell | 0.01487 |
| negative regulation of cellular | 0.01565 |
| protein folding | 0.01568 |
| nitrogen compound transport | 0.0157 |
| positive regulation of mitotic cell | 0.01571 |
| negative regulation of small molecule | 0.01756 |
| positive regulation of cell growth | 0.0197 |
| RNA localization | 0.01973 |
| cellular response to gamma radiation | 0.01979 |
| developmental induction | 0.01979 |
| extrinsic apoptotic signaling pathway | 0.01987 |
| focal adhesion assembly | 0.01998 |
| maintenance of location in cell | 0.0201 |
| regulation of translational initiation | 0.02123 |
| neuron maturation | 0.02147 |
| positive regulation of stress fiber | 0.02147 |
| positive regulation of interleukin-2 | 0.02152 |
| positive regulation of microtubule | 0.02152 |
| negative regulation of ERK1 and ERK2 | 0.02203 |
| RNA processing | 0.02409 |
| histone deacetylation | 0.02434 |
| regulation of viral genome replication | 0.02485 |
| positive regulation of cell cycle G2/M | 0.025 |
| positive regulation of lipase activity | 0.0251 |
| positive regulation of receptor | 0.02513 |
| DNA damage response signal | 0.02545 |
| modification-dependent macromolecule | 0.02637 |
| DNA damage checkpoint | 0.02928 |
| regulation of multicellular organismal | 0.02931 |
| regulation of transcription from RNA | 0.02935 |
| zymogen activation | 0.02954 |

| | |
|---|---|
| embryonic camera-type eye morphogenesis | 0.02959 |
| positive regulation of cell cycle | 0.02963 |
| actin filament-based movement | 0.02968 |
| cellular response to interferon-gamma | 0.02983 |
| maturation of SSU-rRNA from | 0.02993 |
| regulation of organ growth | 0.03064 |
| CD4-positive alpha-beta T cell | 0.03091 |
| rRNA processing | 0.03466 |
| locomotory behavior | 0.03578 |
| transcription, DNA-templated | 0.03752 |
| spermatid development | 0.03783 |
| negative regulation of catalytic | 0.03904 |
| multi-multicellular organism process | 0.03998 |
| protein exit from endoplasmic reticulum | 0.04023 |
| transcription-coupled nucleotide-excisio | 0.04023 |
| oocyte development | 0.04042 |
| response to interleukin-1 | 0.04042 |
| protein localization to microtubule | 0.04046 |
| regulation of intracellular steroid | 0.04049 |
| neurogenesis | 0.04146 |
| thymus development | 0.04196 |
| protein stabilization | 0.04276 |
| developmental growth | 0.0465 |
| adult behavior | 0.04788 |
| carbohydrate metabolic process | 0.04809 |
| cell population proliferation | 0.04815 |
| positive regulation of translation | 0.04881 |

**Supplementary Table 11: 123 GO terms were significantly enriched in infected chicken endothelial cells**

| Term | Weight |
|---|---|
| mitochondrial RNA metabolic process | 0.0001 |
| tRNA aminoacylation for protein | 0.00026 |
| regulation of protein tyrosine kinase | 0.00033 |
| translation | 0.00062 |
| ribosomal large subunit biogenesis | 0.00076 |
| negative regulation of intrinsic | 0.00114 |
| regulation of cyclin-dependent protein | 0.00118 |
| RNA methylation | 0.00308 |
| protein targeting | 0.00322 |
| cytosolic calcium ion transport | 0.00329 |
| negative regulation of apoptotic | 0.00347 |
| mitochondrial gene expression | 0.00386 |
| cytoplasmic translation | 0.00444 |
| early endosome to late endosome | 0.00454 |
| intracellular protein transport | 0.00515 |
| establishment of protein localization | 0.00525 |
| cytoplasmic pattern recognition | 0.00528 |
| intrinsic apoptotic signaling pathway | 0.00528 |
| Golgi to plasma membrane transport | 0.00528 |
| establishment of cell polarity | 0.00536 |
| cellular response to UV | 0.0058 |
| regulation of stem cell differentiation | 0.00581 |
| protein ubiquitination | 0.00614 |
| negative regulation of leukocyte | 0.00635 |
| regulation of epidermal growth factor | 0.00635 |
| positive regulation of viral process | 0.00644 |
| regulation of phosphatidylinositol 3-kin | 0.00645 |
| mRNA polyadenylation | 0.00698 |
| axonal transport | 0.00698 |
| RNA phosphodiester bond hydrolysis | 0.00737 |
| ER-nucleus signaling pathway | 0.00791 |
| translational elongation | 0.00791 |
| myeloid leukocyte migration | 0.00841 |
| alpha-amino acid biosynthetic process | 0.00852 |
| aerobic respiration | 0.00852 |
| rRNA processing | 0.00869 |
| lysosomal transport | 0.00881 |
| dicarboxylic acid metabolic process | 0.00901 |
| regulation of leukocyte chemotaxis | 0.01066 |
| regulation of osteoclast differentiation | 0.01241 |
| positive regulation of innate immune | 0.01255 |
| endoplasmic reticulum unfolded protein | 0.01255 |
| regulation of synaptic plasticity | 0.01257 |
| negative regulation of response to DNA | 0.01271 |

| | |
|---|---|
| cellular response to hypoxia | 0.01383 |
| positive regulation of protein | 0.01508 |
| sulfur compound metabolic process | 0.0155 |
| substrate adhesion-dependent cell | 0.01626 |
| mitochondrial membrane organization | 0.01677 |
| ncRNA processing | 0.01692 |
| mitochondrial transport | 0.01718 |
| localization within membrane | 0.01735 |
| mononuclear cell migration | 0.01737 |
| chaperone-mediated protein folding | 0.01737 |
| spliceosomal snRNP assembly | 0.01737 |
| positive regulation of cellular protein | 0.01742 |
| cellular response to virus | 0.01779 |
| interaction with symbiont | 0.01813 |
| mRNA catabolic process | 0.01815 |
| chromatin organization | 0.01826 |
| autophagosome assembly | 0.01826 |
| microtubule organizing center | 0.01829 |
| apoptotic process | 0.01959 |
| mRNA splicing | 0.02116 |
| regulation of protein dephosphorylation | 0.02172 |
| microtubule cytoskeleton organization | 0.02176 |
| positive regulation of translation | 0.02329 |
| telomere maintenance | 0.02394 |
| regulation of GTPase activity | 0.02509 |
| positive regulation of binding | 0.02523 |
| osteoclast differentiation | 0.02523 |
| regulation of mRNA stability | 0.0254 |
| natural killer cell activation | 0.0254 |
| regulation of extracellular matrix | 0.0254 |
| cholesterol biosynthetic process | 0.0254 |
| protein import | 0.02546 |
| spindle assembly | 0.02565 |
| positive regulation of protein | 0.02566 |
| long-chain fatty acid metabolic process | 0.02566 |
| positive regulation of proteasomal | 0.02724 |
| protein localization to cell periphery | 0.028 |
| ubiquitin-dependent ERAD pathway | 0.02896 |
| cell population proliferation | 0.02948 |
| peptidyl-amino acid modification | 0.02987 |
| protein N-linked glycosylation | 0.03069 |
| protein modification by small protein | 0.03085 |
| positive regulation of signal | 0.03171 |
| regulation of cell cycle | 0.03192 |
| cell migration involved in sprouting | 0.03224 |
| organelle transport along microtubule | 0.03239 |
| RNA 3'-end processing | 0.03348 |
| protein methylation | 0.03397 |

| | |
|---|---|
| DNA packaging | 0.03409 |
| camera-type eye morphogenesis | 0.0342 |
| cellular response to abiotic stimulus | 0.03452 |
| leukocyte mediated cytotoxicity | 0.03538 |
| endothelial cell development | 0.03538 |
| purine ribonucleoside metabolic process | 0.03538 |
| regulation of blood pressure | 0.03607 |
| positive regulation of lymphocyte | 0.0363 |
| regulation of cell adhesion | 0.03636 |
| regulation of lymphocyte migration | 0.0364 |
| L-alpha-amino acid transmembrane | 0.0364 |
| regulation of double-strand break | 0.0364 |
| cellular iron ion homeostasis | 0.0364 |
| regulation of DNA binding | 0.03646 |
| BMP signaling pathway | 0.03649 |
| cellular response to external stimulus | 0.0365 |
| regulation of organelle assembly | 0.03762 |
| cation transmembrane transport | 0.03809 |
| positive regulation of intracellular traunsduction | 0.04161 |
| carboxylic acid catabolic process | 0.04558 |
| regulation of oxidative stress-induced | 0.04589 |
| ribosome assembly | 0.04634 |
| cellular response to interleukin-1 | 0.04634 |
| mitochondrial translation | 0.04634 |
| regulation of vesicle-mediated transport | 0.04672 |
| peptide metabolic process | 0.04757 |
| regulation of G1/S transition of | 0.04857 |
| positive regulation of Notch signaling | 0.04888 |
| establishment of spindle localization | 0.04888 |
| cellular calcium ion homeostasis | 0.04906 |
| negative regulation of translation | 0.04975 |

**Supplementary Table 12: 150 GO terms were significantly enriched in infected duck endothelial cells**

| Term | Weight |
|---|---|
| endodermal cell differentiation | 0.00046 |
| angiogenesis | 0.00048 |
| peptidyl-proline hydroxylation | 0.00087 |
| trabecula formation | 0.00087 |
| inactivation of MAPK activity | 0.00087 |
| negative regulation of cell population p... | 0.00093 |
| cell-substrate adhesion | 0.00179 |
| response to hypoxia | 0.00199 |
| IMP metabolic process | 0.00229 |
| serine family amino acid biosynthetic pr... | 0.00229 |
| sphingomyelin metabolic process | 0.00229 |
| dicarboxylic acid metabolic process | 0.0023 |
| tricarboxylic acid cycle | 0.00246 |
| lymphocyte proliferation | 0.00312 |
| negative regulation of transcription by ... | 0.00331 |
| in utero embryonic development | 0.00352 |
| nitrogen compound metabolic process | 0.00377 |
| negative regulation of canonical Wnt sig... | 0.00427 |
| thymus development | 0.00445 |
| protein homotetramerization | 0.00445 |
| negative regulation of apoptotic process | 0.00456 |
| regulation of osteoblast proliferation | 0.005 |
| positive regulation of phospholipid meta... | 0.00504 |
| tRNA aminoacylation for protein translat... | 0.00511 |
| purine ribonucleoside monophosphate bios... | 0.00548 |
| endoplasmic reticulum calcium ion homeos... | 0.00548 |
| regulation of ERK1 and ERK2 cascade | 0.00579 |
| cerebral cortex radial glia guided migra... | 0.00587 |
| regulation of stem cell proliferation | 0.00588 |
| mitochondrial transmembrane transport | 0.00679 |
| skeletal system morphogenesis | 0.00698 |
| regulation of blood pressure | 0.0071 |
| skin development | 0.00791 |
| proteolysis involved in cellular protein... | 0.00801 |
| retrograde protein transport | 0.00802 |
| response to toxic substance | 0.00818 |
| long-chain fatty acid metabolic process | 0.00914 |
| neuron apoptotic process | 0.00919 |
| myelination | 0.00944 |
| positive regulation of DNA-binding trans... | 0.00975 |
| cardiac chamber morphogenesis | 0.00982 |
| regulation of T cell proliferation | 0.00991 |
| meiotic cell cycle process | 0.00994 |
| outflow tract septum morphogenesis | 0.01051 |

| | |
|---|---|
| heart trabecula morphogenesis | 0.01051 |
| post-embryonic development | 0.01091 |
| glycolytic process | 0.01136 |
| cell fate commitment involved in formati... | 0.01186 |
| negative regulation of striated muscle c... | 0.01186 |
| rhythmic process | 0.01193 |
| cartilage development | 0.01246 |
| endosomal transport | 0.01285 |
| collagen metabolic process | 0.01313 |
| nuclear envelope organization | 0.01318 |
| regulation of cardiac muscle cell differ... | 0.01318 |
| regulation of keratinocyte differentiati... | 0.01341 |
| intracellular receptor signaling pathway | 0.01346 |
| response to unfolded protein | 0.01371 |
| dendritic spine organization | 0.0138 |
| mesenchyme development | 0.01403 |
| Golgi localization | 0.01427 |
| histone H3-K36 methylation | 0.01427 |
| methionine metabolic process | 0.01427 |
| positive regulation of carbohydrate meta... | 0.01432 |
| mRNA catabolic process | 0.01473 |
| negative regulation of gene expression | 0.01476 |
| aromatic compound catabolic process | 0.01517 |
| myoblast differentiation | 0.01565 |
| negative regulation of ERK1 and ERK2 cas... | 0.0158 |
| cellular response to lipid | 0.01587 |
| negative regulation of gene expression | 0.01706 |
| negative regulation of cell differentiat... | 0.01789 |
| positive regulation of transforming grow... | 0.01837 |
| positive regulation of developmental gro... | 0.01871 |
| carboxylic acid metabolic process | 0.01876 |
| oxidation-reduction process | 0.02009 |
| cellular response to hypoxia | 0.02025 |
| epithelial cell proliferation | 0.02088 |
| regulation of protein stability | 0.02136 |
| negative regulation of cyclin-dependent ... | 0.02142 |
| sphingolipid catabolic process | 0.02142 |
| negative regulation of small molecule me... | 0.02268 |
| aorta morphogenesis | 0.02486 |
| one-carbon metabolic process | 0.02486 |
| regulation of calcium ion import | 0.02486 |
| protein N-linked glycosylation via aspar... | 0.02486 |
| extracellular matrix organization | 0.02508 |
| negative regulation of neurogenesis | 0.0252 |
| negative regulation of cellular metaboli... | 0.02534 |
| fatty acid beta-oxidation | 0.0268 |
| regulation of growth | 0.02776 |
| somite development | 0.02839 |

| | |
|---|---|
| peptide catabolic process | 0.02862 |
| sulfur amino acid biosynthetic process | 0.02862 |
| regulation of phospholipid biosynthetic ... | 0.02862 |
| interleukin-1-mediated signaling pathway | 0.02862 |
| folic acid-containing compound metabolic... | 0.02862 |
| positive regulation of glial cell differ... | 0.02862 |
| apoptotic process involved in morphogene... | 0.02862 |
| carboxylic acid biosynthetic process | 0.02902 |
| carbohydrate derivative catabolic proces... | 0.02941 |
| branching involved in blood vessel morph... | 0.02982 |
| embryonic hindlimb morphogenesis | 0.02982 |
| regulation of SMAD protein signal transd... | 0.02982 |
| regulation of hematopoietic progenitor c... | 0.02982 |
| positive regulation of transcription by ... | 0.03033 |
| epithelium development | 0.03049 |
| positive regulation of cell population p... | 0.03067 |
| response to endoplasmic reticulum stress | 0.03099 |
| protein localization to organelle | 0.03101 |
| negative regulation of cell growth | 0.03135 |
| embryonic eye morphogenesis | 0.0314 |
| regulation of transcription by RNA polym... | 0.03143 |
| establishment of protein localization to... | 0.03146 |
| regulation of interleukin-1 production | 0.03146 |
| organic anion transport | 0.03149 |
| regulation of immune response | 0.03215 |
| cellular response to fibroblast growth f... | 0.03231 |
| ER-nucleus signaling pathway | 0.03262 |
| actin nucleation | 0.03277 |
| tissue homeostasis | 0.03492 |
| antibiotic metabolic process | 0.03534 |
| branching involved in ureteric bud morph... | 0.03534 |
| positive regulation of T cell differenti... | 0.03536 |
| chondrocyte development | 0.03546 |
| sulfur amino acid metabolic process | 0.03551 |
| membrane protein proteolysis | 0.03569 |
| cardiac septum development | 0.03573 |
| extrinsic apoptotic signaling pathway in... | 0.03579 |
| regulation of phosphatidylinositol 3-kin... | 0.03583 |
| regulation of protein deacetylation | 0.03588 |
| prostate gland development | 0.03592 |
| regulation of histone acetylation | 0.03602 |
| myeloid cell differentiation | 0.03665 |
| sensory perception of mechanical stimulu... | 0.03685 |
| apoptotic mitochondrial changes | 0.03691 |
| cellular response to unfolded protein | 0.03692 |
| negative regulation of secretion | 0.03699 |
| postsynapse organization | 0.03701 |
| female pregnancy | 0.03707 |

| | |
|---|---|
| positive regulation of vasculature devel... | 0.03707 |
| cellular response to steroid hormone sti... | 0.03708 |
| synaptic vesicle recycling | 0.03709 |
| ceramide biosynthetic process | 0.03711 |
| regulation of interleukin-10 production | 0.03713 |
| regulation of G2/M transition of mitotic... | 0.03716 |
| peptidyl-lysine modification | 0.03745 |
| face morphogenesis | 0.03797 |
| protein stabilization | 0.03818 |
| cellular response to insulin stimulus | 0.03867 |

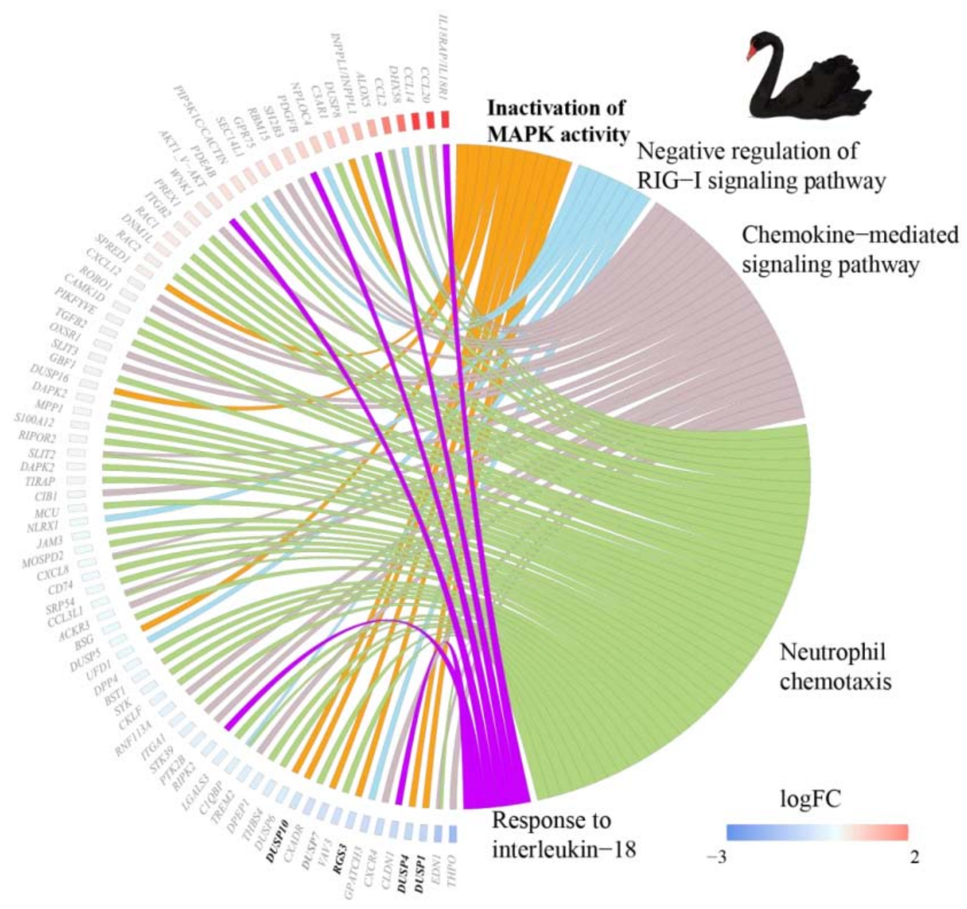


**Supplementary Figure 11: Differential regulation of DEGs linked to the “inactivation of MAPK’’ pathway in black swan endothelial cells in response to VN04 infection along with randomly selected GO terms.** Significant genes are in bold. The figure includes additional GO biological process terms associated with black swan endothelial cells’ DEGs to create a complete picture

**Supplementary Table 13: Dual-specificity phosphatases in black swan, chicken and duck endothelial cells infected with VN04***

| Gene | Black Swan | Chicken | Duck |
|---|---|---|---|
| *DUSP1* | | | |
| *DUSP4* | | NE | |
| DUSP5 | | | |
| DUSP6 | NE | | NE |
| DUSP7 | | | |
| DUSP8 | NE | NE | |
| DUSP10 | | | |
| DUSP12 | NE | | |
| DUSP15 | NE | | NE |
| DUSP16 | NE | NE | |
| DUSP19 | NE | | NE |
| DUSP23 | NE | | NE |
| RGS3 | | NE | |
| SPRED1 | NE | NE | |

* Green indicates significantly upregulated genes; red indicates significantly down regulated genes; NE = not differentially expression (adjusted p value >0.05) or not expressed

**Supplementary Table 14: Parameters used for the FALCON run.** Length cut-off was detected automatically for seed-read length

| Parameter | Value |
|---|---|
| Length cut-off | -1 |
| Length cut-off pre-reads | 5000 |
| Target | Assembly |
| Genome size | 1,4181 x 10^9 |
| Seed-coverage | 30 |
